# Hurricane Harvey Impacts on Water Quality and Microbial Communities in Houston, TX Waterbodies

**DOI:** 10.1101/2022.01.20.476661

**Authors:** Michael G. LaMontagne, Yan Zhang, George J. Guillen, Terry J. Gentry, Michael S. Allen

## Abstract

Hurricanes, and other extreme weather events, can temporarily alter the structure of coastal systems and generate floodwaters that are contaminated with fecal-associated microbes; however, every coastal system and extreme weather event is unique, so identification of trends and commonalities in these episodic events is challenging. To improve our understanding of the resilience of coastal systems to the disturbance of extreme weather events and the risk exposure to floodwaters poses to the public, we monitored water quality, fecal indicator bacteria (FIB) at three stations within Clear Lake, an estuary between Houston and Galveston, and three stations in bayous that feed into the estuary. Water samples were collected immediately before Hurricane Harvey (pre-HH, August 25^th^, 2017), immediately after (HH, August 30^th^) and then throughout the fall of 2017 (post-HH). FIB levels were monitored by culturing *E. coli* and *Enterococci*. Microbial community structure was profiled by high throughput sequencing of PCR-amplified 16S rRNA gene fragments. Water quality and FIB data was also compared to historical data for these water body segments. Before Harvey, salinity within Clear Lake ranged from 9 to 11 practical salinity units (PSU). Immediately after the storm, salinity dropped to < 1 PSU and then gradually increased to pre-storm and historical levels over two months. Dissolved inorganic nutrient levels were also relatively low immediately after Harvey and returned, within a couple of months, to pre-HH and historical levels. FIB levels were elevated immediately after the storm; however, after one week, *E. coli* levels had decreased to acceptable levels for freshwater. *Enterococci* levels collected several weeks after the storm were within the range of historical levels for these water bodies. Microbial community structure shifted from a system dominated by *Cyanobacteria* sp. before HH to a system dominated by *Proteobacteria* and *Bacteroidetes* immediately after. Further, several sequences observed only in floodwater showed similarity to sequences previously reported for samples collected following Hurricane Irene. These changes in beta diversity corresponded to salinity and nitrate/nitrite concentrations. Differential abundance analysis of metabolic pathways, predicted from 16S sequences, suggested that pathways associated with virulence and antibiotic resistance were elevated in floodwater. Overall, these results suggest that recovery of the Clear Lake system following Hurricane Harvey took at least a month and floodwater generated from these extreme events may have high levels of fecal contamination, antibiotic resistant bacteria and bacteria rarely observed in other systems.

## Introduction

Hurricane Harvey deluged the Houston metropolitan area in August of 2017 with over a meter of rain in less than 48 hours. This rainfall set a record for the continental United States [9], and exposed thousands, perhaps millions, of citizens and first responders to potentially contaminated floodwaters. In rural regions typical of areas north of Houston, flooding of agricultural land could release animal waste associated with areas used for animal grazing [18]. In suburban watersheds typical of the greater Houston-Galveston area, rainfall could accelerate the resuspension and transport of waste from onsite sewage facilities, such as residential septic tanks [35]. Indeed, waterways in the Houston-Galveston area frequently exceed fecal indicator bacteria (FIB) criteria during high flow and flood events [42, 55]. Little is known about the health risks associated with exposure to sewage and other human waste in floodwaters in urban, industrialized watersheds [1]. Human waste presents a particular health threat [52] and the perception that floodwater is contaminated with sewage could further alarm and mentally traumatize the public and hamper recovery efforts [16, 17]. These threats to public health are expected to worsen, as several models predict that the intensity, if not the frequency, of tropical cyclones and hurricanes will increase over the next few decades [29, 61].

Extreme weather events could also alter the quality of receiving waters. Flooding can result in release of petroleum products and other hazardous materials that could stress aquatic systems [20, 36, 47]. This environmental risk is high in the Galveston Bay systems; the Houston Ship Channel is the largest petrochemical complex in the United States [4]. Floodwaters can also temporarily alter nutrient cycles. For example, Hurricane Bob, a category 3 storm when it landed on Cape Cod, increased nutrient loading to estuaries in Cape Cod, Massachusetts, but the system appeared to recover rapidly [59]. Hurricane Ivan exacerbated eutrophication in Pensacola Bay, Florida temporarily, but the system recovered in a few days [21]. The extent to which these few studies can be extrapolated to other areas with unique geographies and conditions remains unknown. The current paucity of data reflects the inherent challenges of quantifying multiple stressors during extreme yet ephemeral events [12]. Metagenomic methods have the potential to provide additional insight into overall ecosystem health; however very few studies have applied these methods to assess the health of aquatic systems, particularly with respect to extreme weather events [19].

Here we apply metagenomics to determine the impact of Hurricane Harvey (HH) on the health of Clear Lake, an estuary between Houston and Galveston that connects with upper Galveston Bay. This estuary is popular with anglers and boaters and is routinely monitored by a consortium of state agencies, non-profits and academic institutions. We collected water samples immediately before and after landfall of Hurricane Harvey, and then weekly into the fall. These samples were analyzed for fecal indicator bacteria (FIB), dissolved inorganic nutrients (DIN) and microbial community structure, as assessed by metagenomic analysis. FIB counts, DIN concentrations and other environmental parameters were compared to data mined from public archives. Relative to pre-storm levels, and values typical for waterbodies sampled herein, HH elevated FIB counts and lowered DIN and salinity concentrations. The structure of the community shifted from a community dominated by *Cyanobacteria* and A*ctinobacteria* before the storm to a community dominated by the phyla *Proteobacteria* and *Bacteroidetes* immediately after the event. Shifts in the microbiological community structure corresponded to changes in salinity and NO_x_ concentrations.

## Methods

### Sampling Locations and Collection

We selected sites around Clear Lake (Fig. 1), an estuary between Houston and Galveston (Fig. S1), based on the availability of long-term water quality data for these locations and ease of access. This map was generated with scripts presented in file S1. Water samples were collected on the afternoon of August 24^th^, 2017, one day before Hurricane Harvey landed in the Corpus Christi area. These samples are designated “pre” in this work. A second set of samples was collected on August 30^th^. These samples are designated “HH” throughout, as they correspond to Hurricane Harvey samples. We added a second sampling site (N) when collecting the HH set to collect floodwater received by segment 1101C (Fig. 1). Starting on September 8^th^ we sampled six times to generate a “post” sample set. Samples collected in August and September were designated as summer season. Samples collected in October were designated as fall season. We also generated a mock sample by mixing raw sewage, collected as described previously [2], and surface water collected from station H (Fig. 1) in March of 2018. The sewage and water were mixed at a ratio of 1 part sewage with 9 parts surface water.

**Figure 1.**
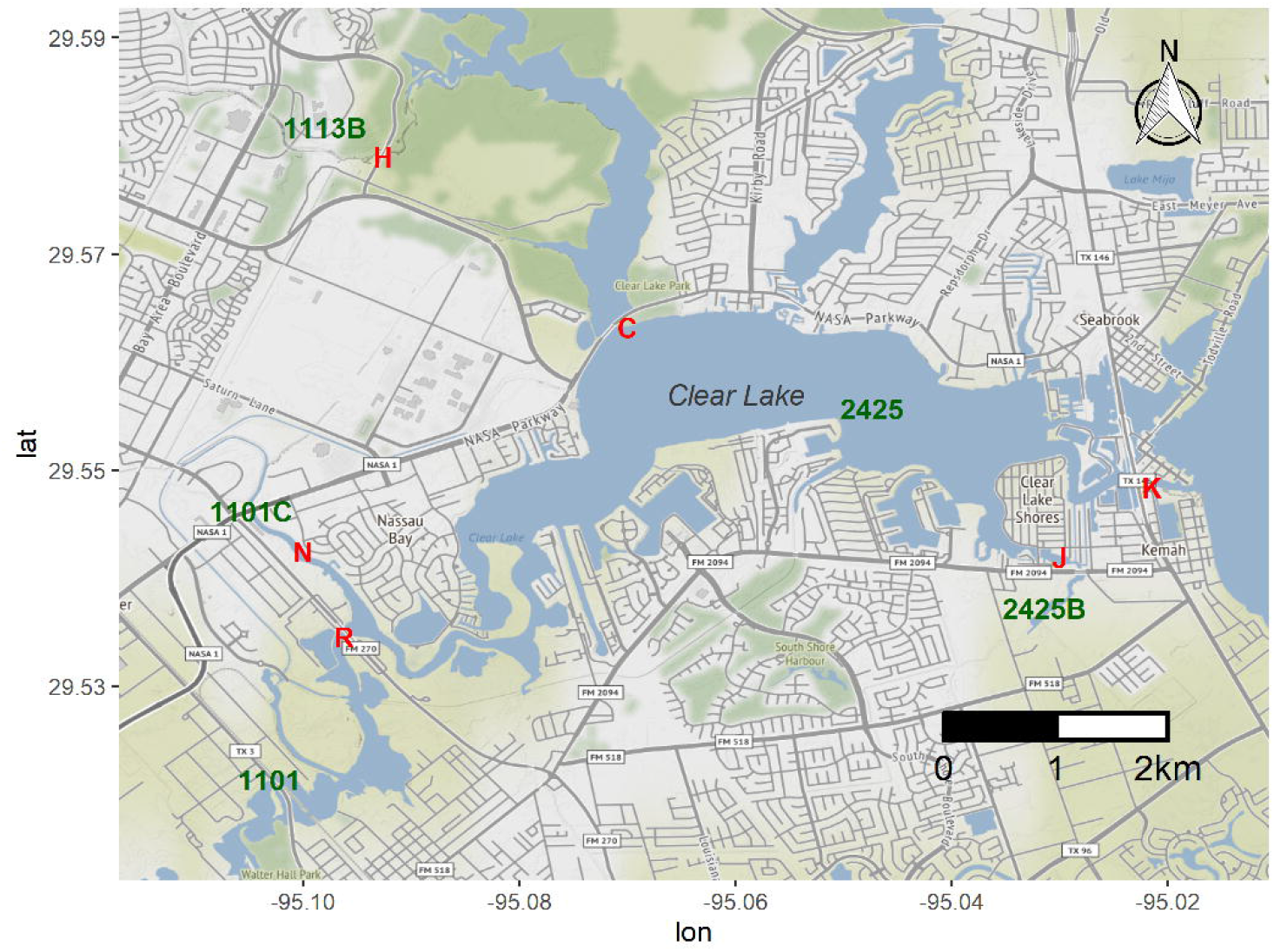
Sampling stations around Clear Lake. Stations sampled in this study are indicate by red letters (H, C, K, J, R, and N). Water body segments as defined by the TCEQ (1113B, 2424, etc.) are indicated in green. Figure was generated with scripts in File S01.

At each sampling station, we collected surface water samples with a bucket lowered from an overpass or dock. Temperature and dissolved oxygen were measured *in situ* at 3 – 5 cm beneath the surface with a YSI model 55 dissolved oxygen (DO) probe (YSI Inc., Young Spring, OH). Water samples were split in the field for FIB (*E. coli* and *Enterococci*), metagenomic and nutrient analysis. For FIB analysis, unfiltered samples were transported on wet ice and stored at 4 °C. Incubations for quantification of FIB were initiated within 24 hours of sampling. FIB samples from the pre-HH samples were discarded because we were locked out of our laboratory for several days and sampling holding times were exceeded.

Water samples collected before HH were stored on wet ice, returned to the laboratory and filtered to collect microbial samples for metagenomic analysis and to archive nutrient samples within 24 hours of collection. Following HH, all samples were filtered in the field immediately upon collection. For metagenomic analysis, water samples were pulled through a Sterivex SVGPL10RC 0.2 µm cartridge (EMD Millipore, Billerica MA) until refusal (no flow at 15 psi), with a hand vacuum pump, as described previously [30]. The volume filtered, which ranged from 75 and 300 ml, was measured with a graduated cylinder. Sample eluates and cartridges were transported on wet ice, temporarily stored at -20 °C, and archived at -80 °C. Two technical replicates were generated by collecting duplicate samples from station J on the eve of the storm and from the sewage-spiked samples described above.

### Laboratory Methods

Dissolved inorganic nutrient analysis for ammonium, orthophosphate, and nitrate were done by colorimetric analysis in microplates, as described previously [46]. *E. coli* and *Enterococci* were enumerated using Colilert and Enterolert in the Quanti-Tray/2000 format following manufacturer recommendations (IDEXX, Westbrook, ME). Microbial community DNA for metagenomic analysis was recovered from the Sterivex cartridges as described previously [2] and assessed for molecular weight by agarose gel electrophoresis. These crude extracts were subsequently purified by passage through a OneStep PCR Inhibitor Removal column (Zymo, D6030) and purity was assessed by UV-spectra. Metagenomic analysis followed protocols outlined in the Earth Microbiome Project [8]. Briefly, the V4 region of the 16S rRNA gene was amplified to generate an amplicon library. This library was multiplexed using Illumina designed indices, pooled with equal amounts, and sequenced on an Illumina MiSeq instrument as described by Caporaso et al. [7].

### Data Analysis

Water quality data were analyzed and figures were generated with custom scripts presented in supplemental files S2 and S3. These scripts included functions from packages from ggplot2 [62]. This data set included water quality data collected as described above and public data previously collected by the Texas Commission for Environmental Quality (TCEQ) and cooperating organizations. Data from the TCEQ archive was limited to samples collected between January 1^st^, 2011 and May 6^th^, 2021. For this time period, the mean for each of these five segments was calculated by grouping by segment, month and year. This data range was then merged with water quality data generated from samples collected in this study to generate supplemental file S4.

MiSeq data were processed to determine alpha diversity of the microbial community using functions from DADA2 v. 1.20.0, [5], with custom scripts presented in supplemental file S5. Briefly, reads were filtered, trimmed, denoised, and merged to yield sequences from 251 to 253 nucleotides long. Chimeras were then removed with the function removeBimeraDenovo in DADA2 and putative nonchimeric sequences were assigned taxonomy and aligned with the functions IdTaxa and AlignSeqs in Decipher v 2.20.0 [64]. Amplicon sequence variants (ASVs) and taxonomic identifications were merged to create a phylogseq-class object - available as file S6 - with functions in phyloseq v 1.36.0 [33]. ASVs with uncertain taxonomic identification at the phylum level were then removed before fitting the alignments into a phylogentic tree with functions in phangom v 2.7.1 [50]. Technical replicates (two samples collected at the same time and place but processed independently) were then merged and meta-data (volume filtered, environmental conditions, FIB counts, DIN, etc.) and reference sequences were combined to create a phylogseq-class object - available as file S7 - with functions in phyloseq and Biostrings (v 2.60.2 [38]). Reference sequences were also exported in fasta format and compared to public sequences with the Seqmatch application (RDP Taxonomy 18) available through the Ribosomal Database Project [11]. Default settings were used in Seqmatch. MiSeq data is available in the NCBI SRA under accession number/Bioproject ID: PRJNA795782.

Alpha diversity (richness and Shannon indices) was estimated with the plot_richness function after sewage-spiked samples were removed. Analysis of variance, calculated with a core function in R version 4.1.1, was used to test for significance of differences between samples collected before HH made land fall (sampled August 25^th^), versus samples collected immediately after the storm (August 30^th^) and in September and October. Significance of differences was assessed with a Tukey test using the function HSD.test in the R package agricoloe v 1.3.5[13]. Coverage of the library of reads used for diversity analysis was visualized with the function rarecure in vegan v 2.5.7 [37].

The relationship between microbial community structure and nutrient levels was determined with correspondence analysis with custom scripts presented in R markdown file in S8. This workflow started with a phyloseq object (S7). A prevalence threshold of 10% was set to remove rare taxa. Counts of the remaining 1,616 ASVs were transformed with Hellinger option prior to nonmetric multidimensional scaling analysis (NMDS) with functions in vegan v 2.5.7 [37]. NMDS was first run with all 44 samples, prior to removal of sewage-spiked samples. Goodness of fit of NMDS ordination was visualized with a Shepard plot generated prior to fitting meta-data to the ordination with functions in vegan. The resulting ordination plots were visualized with functions available in R package ggordiplots v 0.4.0 [45].

Functional composition was predicted from the 1,616 ASVs used in NMDS analysis (above) with PICRUSt2 v 2.3.0-b [14]. To prepare the data, an ASV abundance table and fasta files were exported from a phyloseq object (S09) to a biom file (S10) and a sequence file (S11) using R package biomformat v 1.20.0 [34]. This pipeline, including bash scripts used in PICRUCSt2 analysis, are presented in File S12.

Differential abundance of pathways (Files S13 and S14) predicted from ASVs and ASVs themselves was assessed using functions in R package ANCOM-BC v 1.2.2 [32], following scripts presented in file S15. Pathways that were differentially abundant were plotted with functions available in R package Heatplus v 3.0.0 [43], following scripts presented in file S16. Pathway functions and expected taxonomic range associated with them were identified with the web application MetaCyc v 25.5 [10].

## Results

### Environmental Conditions

On the eve of HH (August 25^th^, 2017), surface salinity at stations C and K, which correspond to water body segment 2425 (i.e. Clear Lake) (Fig. 1), were 9 and 12 practical salinity units (PSU), respectively. These pre-HH salinity levels are within the 95 % confidence interval of the ten-year average for salinity for records for this water body segment (2425, Fig. S2). Immediately after the storm, salinity dropped to < 1 PSU at all stations sampled herein and then gradually increased to pre-storm levels over the next two months (Fig. 2).

**Figure 2.**
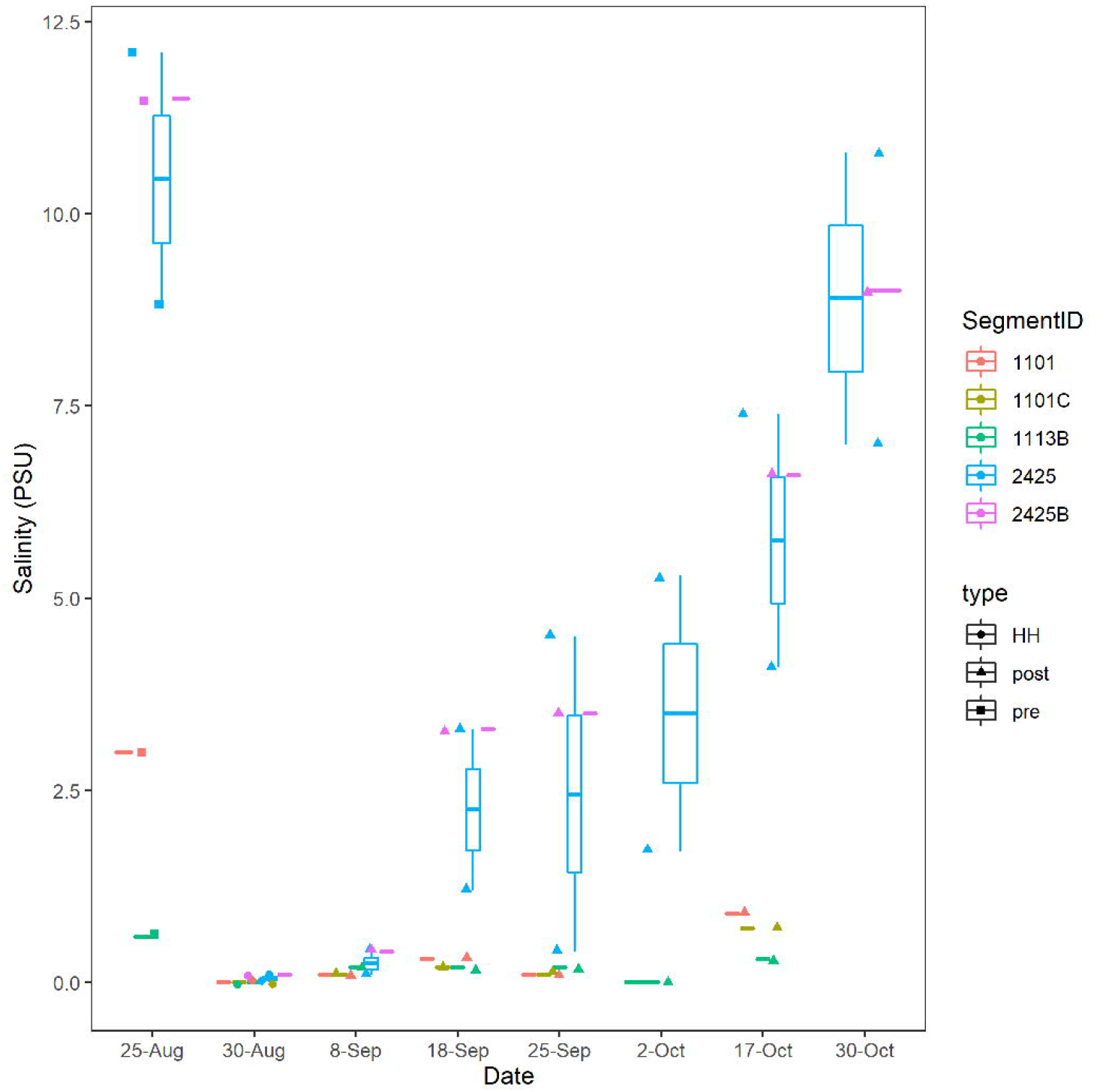
Salinity concentrations in practical salinity units (PSU). All dates are 2017. Boxes indicate 25 and 75% quantiles. Whiskers indicate range. Horizontal lines indicate median or, for segments where only one sample was taken, the value for that individual sample. Water body segments are as Figure 1. Figure was generated with scripts in File S03.

Hurricane Harvey lowered the concentration of dissolved inorganic nutrients (DIN). On the eve of the storm, nitrate/nitrite (NO_x_) ranged from 1 – 59 µM (Fig. S3). These pre-HH NO_x_ levels are in the range for records for the last ten years for these segments (Fig. S3). Immediately after the storm, NO_x_ ranged from 1 – 9 µM. NO_x_ varied significantly between segments (p < 0.001) and type (pre-storm, HH, and post-storm). Highest levels of DIN were observed for samples collected from segment 1113B, which is approximately 50 meters downstream of the outfall pipe of a wastewater treatment plant.

NO_x_ showed a non-conservative mixing relationship with salinity (Fig. 3). That is the system is a sink NO_x_. High concentrations (> 10 µM) were associated with samples that showed salinities of 3 PSU or less. Samples that showed salinities greater than 3 PSU, showed NO_x_ levels of 3 µM or less, which suggests a freshwater source. Ammonium and phosphate levels showed a similar pattern with salinity as NO_x_. High concentrations of ammonium (Fig. S4) and phosphate (Fig. S5) were associated with low salinities. DIN/P ratios were generally below 16 (Fig. S6). These ratios were on average highest (7 – 8) for segments 1101C and 1113B, respectively, and < 2 for segments 2425 and 2425B.

**Figure 3.**
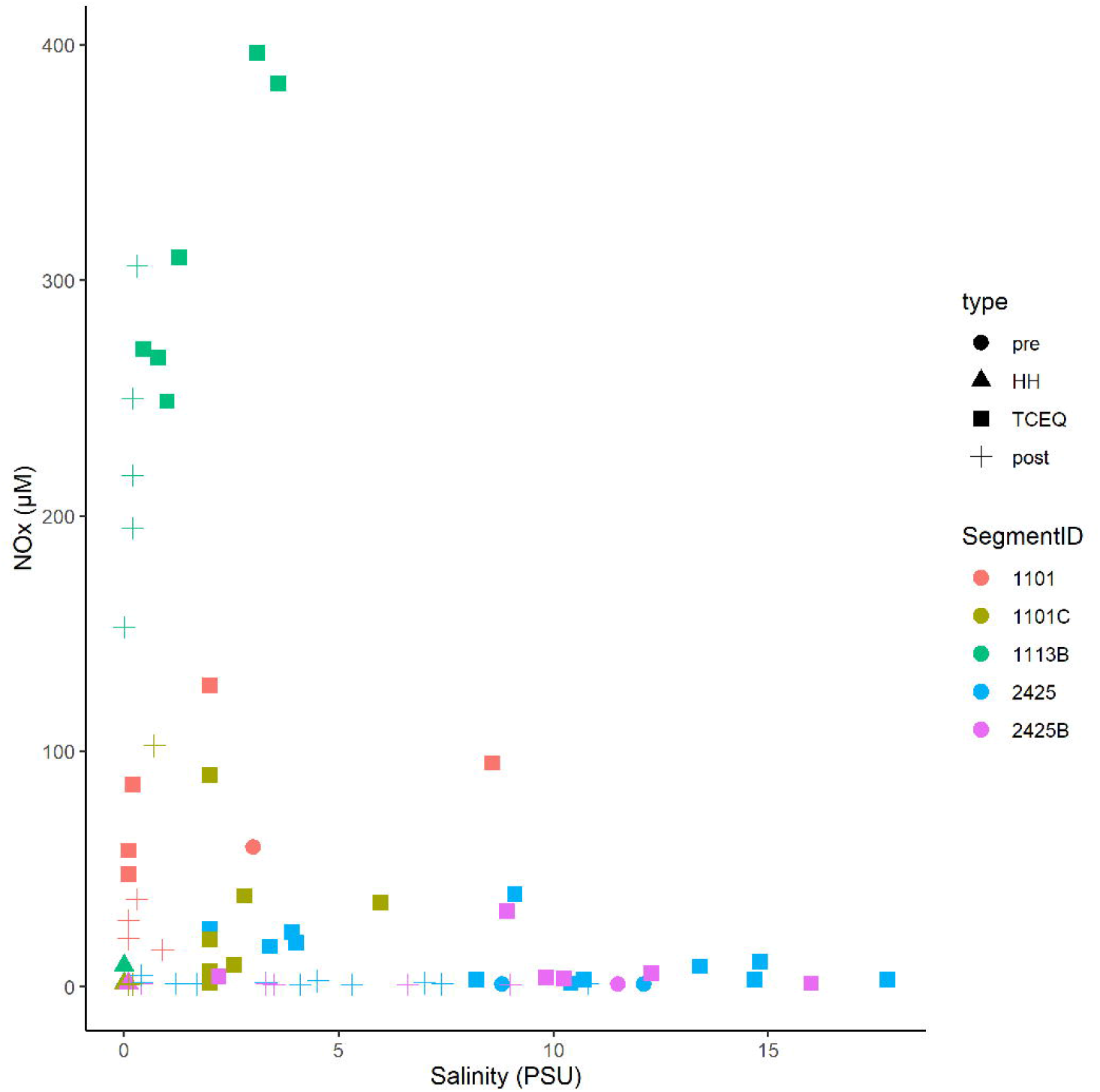
Mixing diagram of salinity versus nitrate/nitrite for the Clear Lake system. Sample type “pre” indicates samples collected on August 25^th^ (before HH). Type “HH” indicates samples collected on August 30^th^ (immediately after HH). Type “post” indicates samples collected from September 8^th^ to October 30^th^. Type “TCEQ” indicates historical data collected by TCEQ and partner agencies during 2011 – 2021. Note DIN data is not available for the sample collected before HH from waterbody segment 1113B. Figure was generated with scripts in File S03.

Oxygen levels in Clear Lake and tributaries to that system did not show a strong response to HH. Across all segments, oxygen levels averaged 6.8 mg/L before HH and 5.2 mg/L after. This temporal difference was not significant (p = 0.131); however, spatial differences between segments were significant (p = 0.0003). Oxygen levels averaged 6.9 – 7.1 mg/L for segments 2425 and 2425B, respectively, and less than 6 mg/L for stations 1101C, 1101, and 1113B (Fig. 4). Lowest oxygen concentrations were observed at station H in waterbody segment 1113B, where DIN concentrations are relatively high.

**Figure 4.**
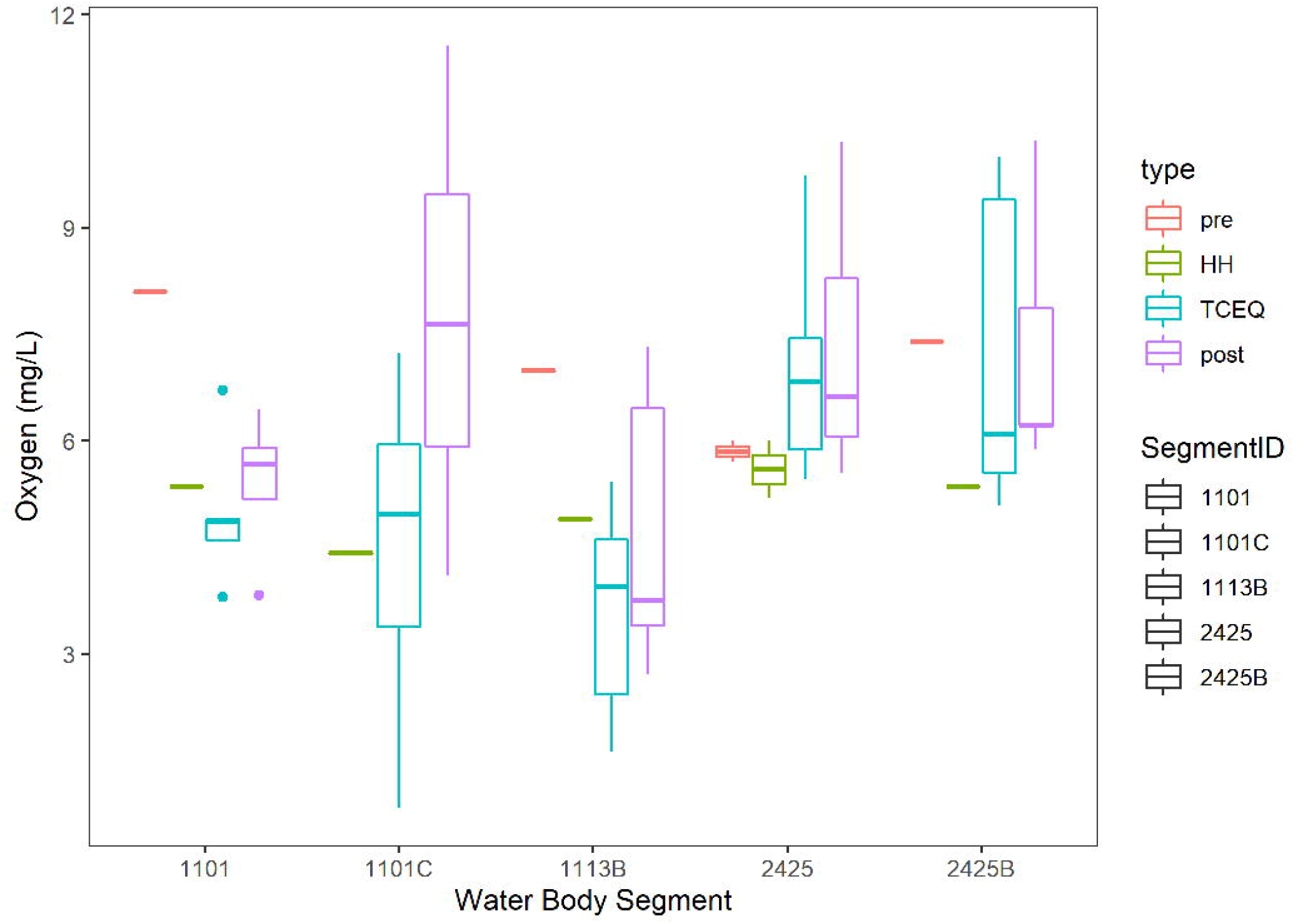
Boxplots of oxygen levels in the Clear Lake system. Segments are depicted in Figure 1. Sample types are as Figure 3. Boxes, whiskers and lines indicate are as Figure 2. Figure was generated with scripts in File S03.

### Fecal indicator bacteria

*E. coli* levels ranged from 488 to 1,733 MPN/100 ml for the six stations sampled on September 1^st^, 2017 (Fig. 5), which was 72 hours after HH pasted over the study area. The geometric mean (GM) for this set of samples was 1,018 MPN/100 ml. These values exceeded the statistical threshold value (STV) for single samples and GM recommended for water contact by the EPA [58] and by the TCEQ for these particular water body segments [55]. After one week, the *E. coli* levels had decreased to <100 MPN/100 mL and remained relatively low until the end of October, when levels spiked again.

**Figure 5.**
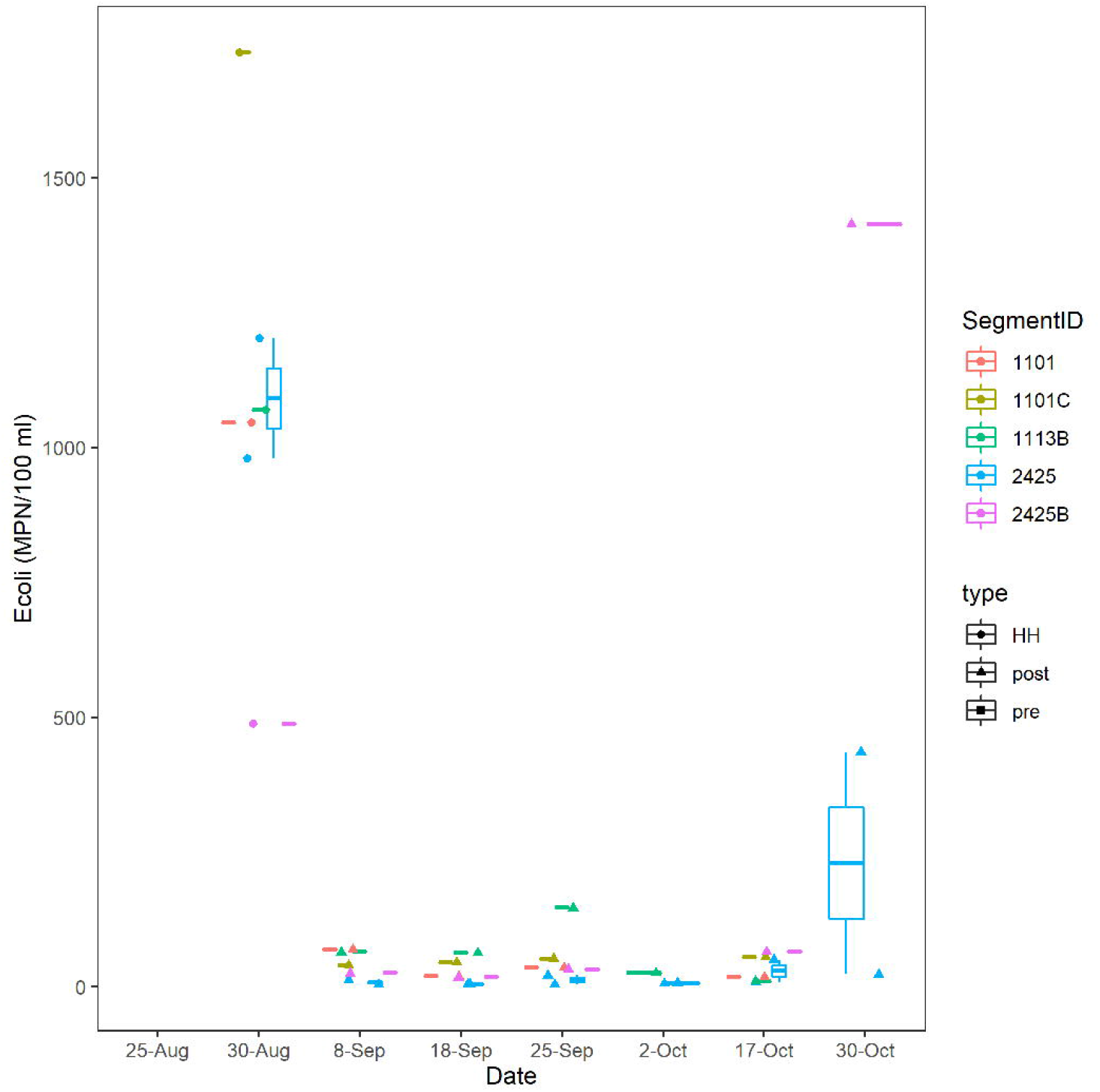
*E. coli* concentrations over time. MPN/100 ml were assessed by culturing as described in Methods. Dates are as Figure 2. Figure was generated with scripts in File S03.

*Enterococci* levels ranged from 63 to 3,050 MPN/100 ml for samples collected in the fall of 2017; however, because of logistical issues, *Enterococci* levels were not measured until September 18^th^. For this period (post-HH), *Enterococci* levels did not differ between segments sampled (Fig. S07), and 22 of 24 samples exceeded the STV for single samples recommended for recreational water contact by the EPA [58]; GM (495 MPN/100 ml) exceeded, by an order of magnitude, the GM recommended for recreational water contact by the EPA [58]. FIB counts of samples taken in the post-HH period, were significantly higher (p < 0.001) than counts for the same segments collected over the last decade, where the GM of *Enterococci* was 74 MPN/100 ml.

### Microbial Diversity

Alpha diversity of the bacterial and archaeal community did not differ significantly between samples collected before and after HH (Fig. 6). Average Shannon diversity indices ranged from 4.95 to 4.89 for samples after the event and averaged 4.44 for samples collected immediately before the storm. Diversity was relatively lower for samples collected before HH at stations 1101 and 1113B but only one sample was collected at that time point (Fig. S8). Average richness ranged from 761 to 633 for samples collected after the event and 488 for samples collected before (Fig. S9). Rarefaction analysis suggested the sequence library appeared to have the depth to describe alpha diversity (Fig. S10). After quality control, which included removing sequences that did not classify at the phylum level, average depth of the library was 112,178 reads. In other words, all 44 samples reached an asymptote.

**Figure 6.**
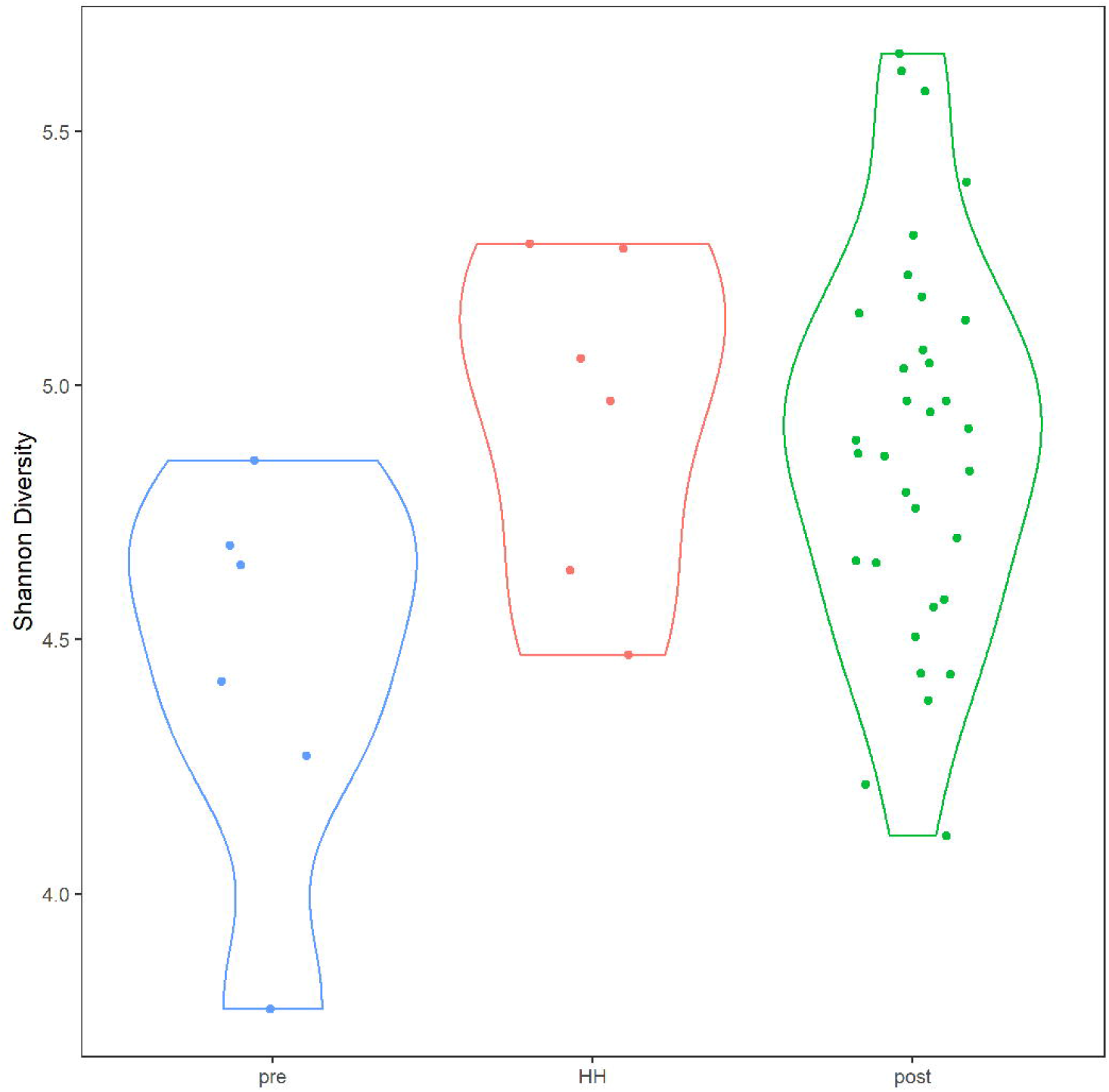
Alpha diversity in Clear Lake system before and after Hurricane Harvey. Y-axis indicates Shannon diversity indices. Categories correspond to before (pre), immediately after (HH) and more than a week after (post). Figure was generated with scripts in File S05.

Beta diversity of the bacterial and archaeal community structure, as assessed by NMDS, differed significantly between samples collected before and immediately after HH (Fig. 7). This ordination analysis was supported by a good fit between the dissimilarity of samples to their ordination distance (Fig. S11) and appeared driven by NO_x_ and salinity/conductivity (Fig. 8). Salinity appeared strongly (p = 0.001) associated with pre-HH samples and NO_x_ appeared strongly (p = 0.019) associated with post-HH samples. Phosphate also appeared associated with post-HH samples but the significance was weak (P = 0.099).

**Figure 7.**
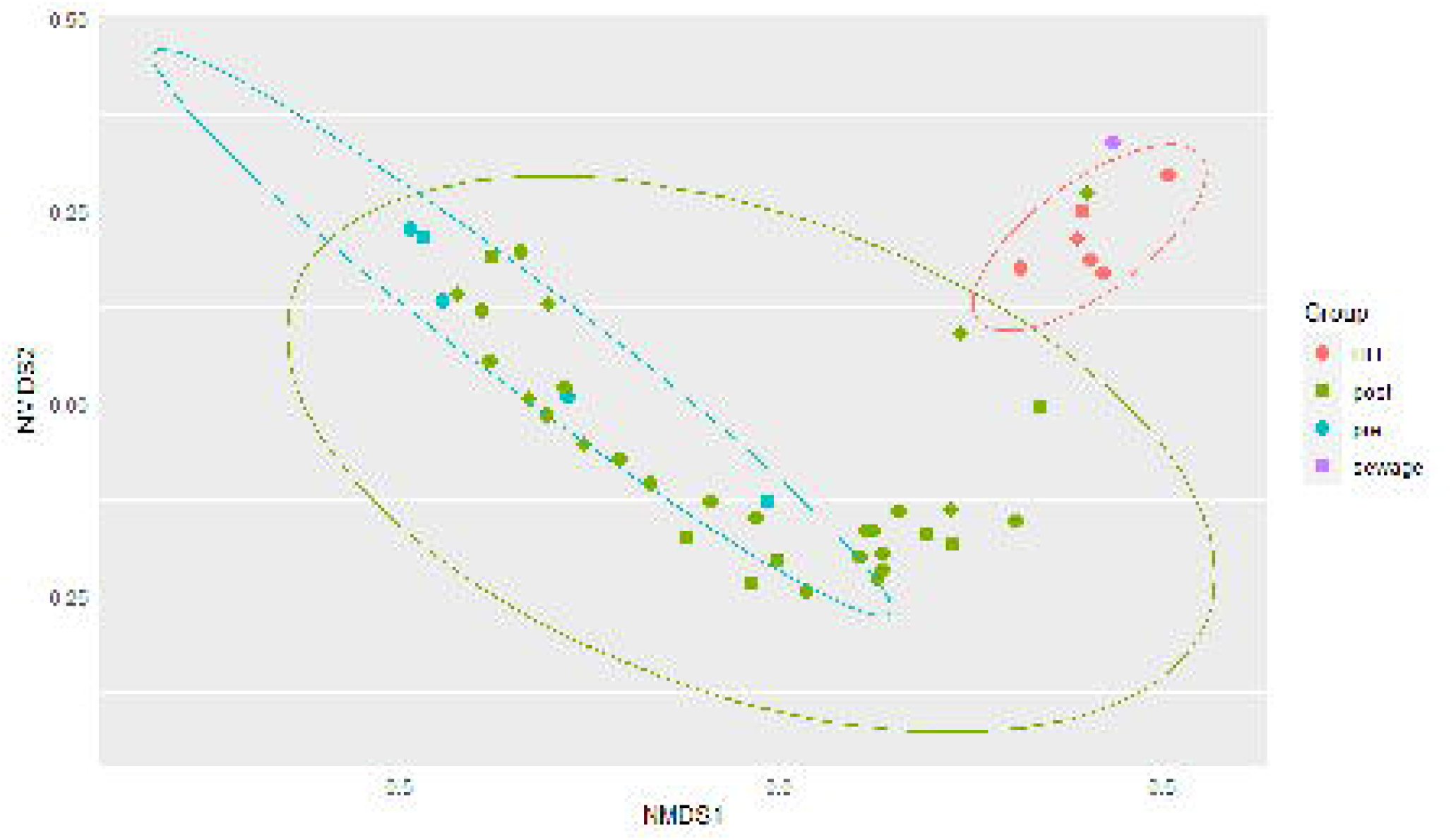
Nonmetric multidimensional scaling analysis (NMDS) of microbial community structure for samples collected before and after Hurricane Harvey. NMDS was run on the abundance of amplicon sequence variants as described in Methods. Group indicates type of samples (see Figure 2) or a sewage spike sample. Figure was generated with scripts in File S08.

**Figure 8.**
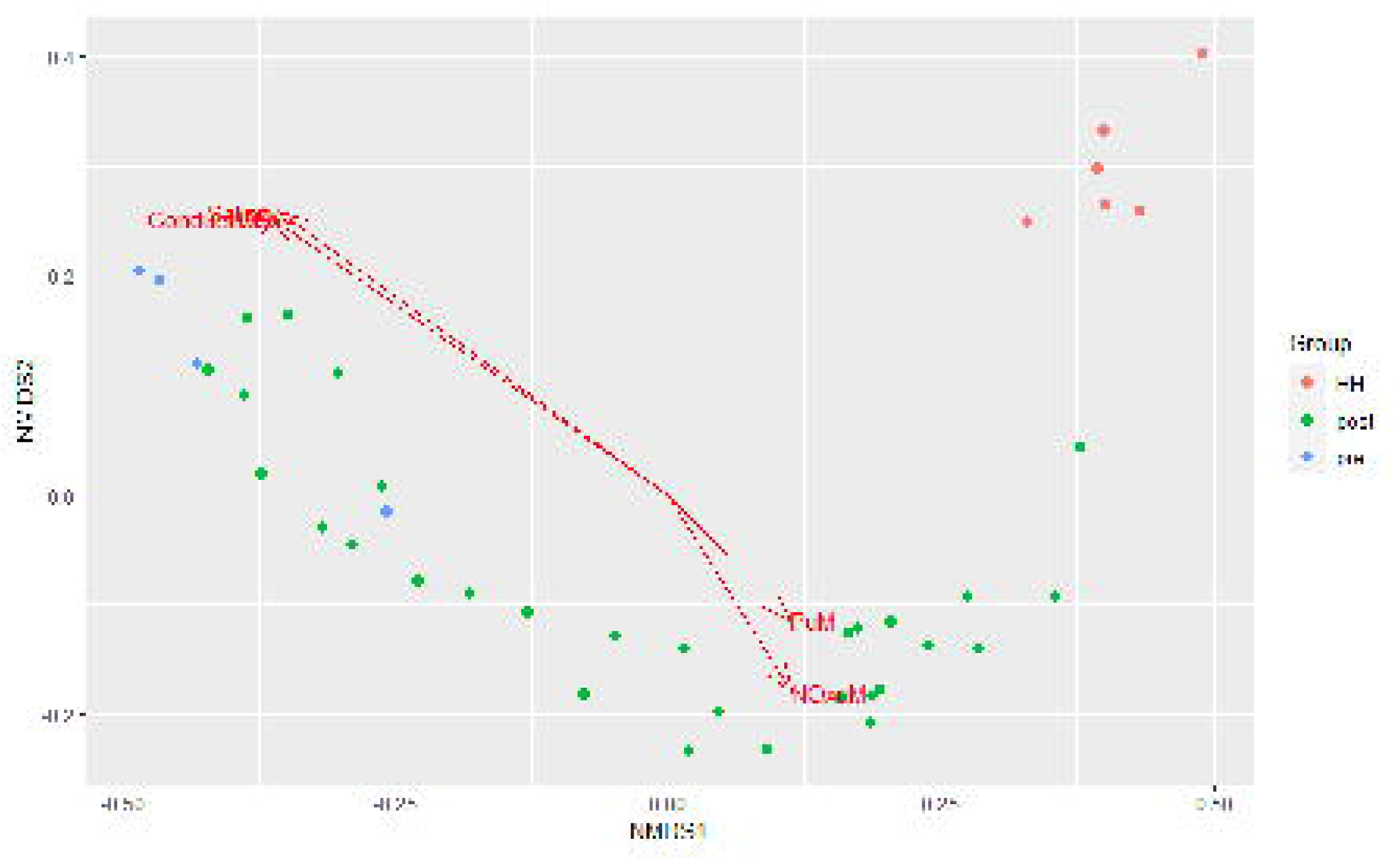
Fit of environmental data to NMDS model of microbial community structure. Environmental variables that showed a significant (P < 0.10) relationship with community structure are shown. Only samples with DIN data available are shown. Note conductivity and salinity vectors were practically identical to each other and are necessarily plotted on together. Figure was generated with scripts in File S08.

Bacterial community structure of the Clear Lake system shifted from a system dominated by Cyanobacteria before HH to a system dominated by *Proteobacteria* and *Bacteroidetes* immediately after (Fig 9). A total of 59 phyla were detected in 7,491 ASVs generated from 44 samples. Almost all of these ASVs (7,410/7,491) classified as bacteria. After removing taxa with relatively low (<10 %) prevalence, almost all of the ASVs (1534/1617) were found to be differentially abundant in a model that tested the factors: salinity, NOx and sample type (pre, HH & post). In other words, these factors predicted the abundance of 95% ASVs.

**Figure 9.**
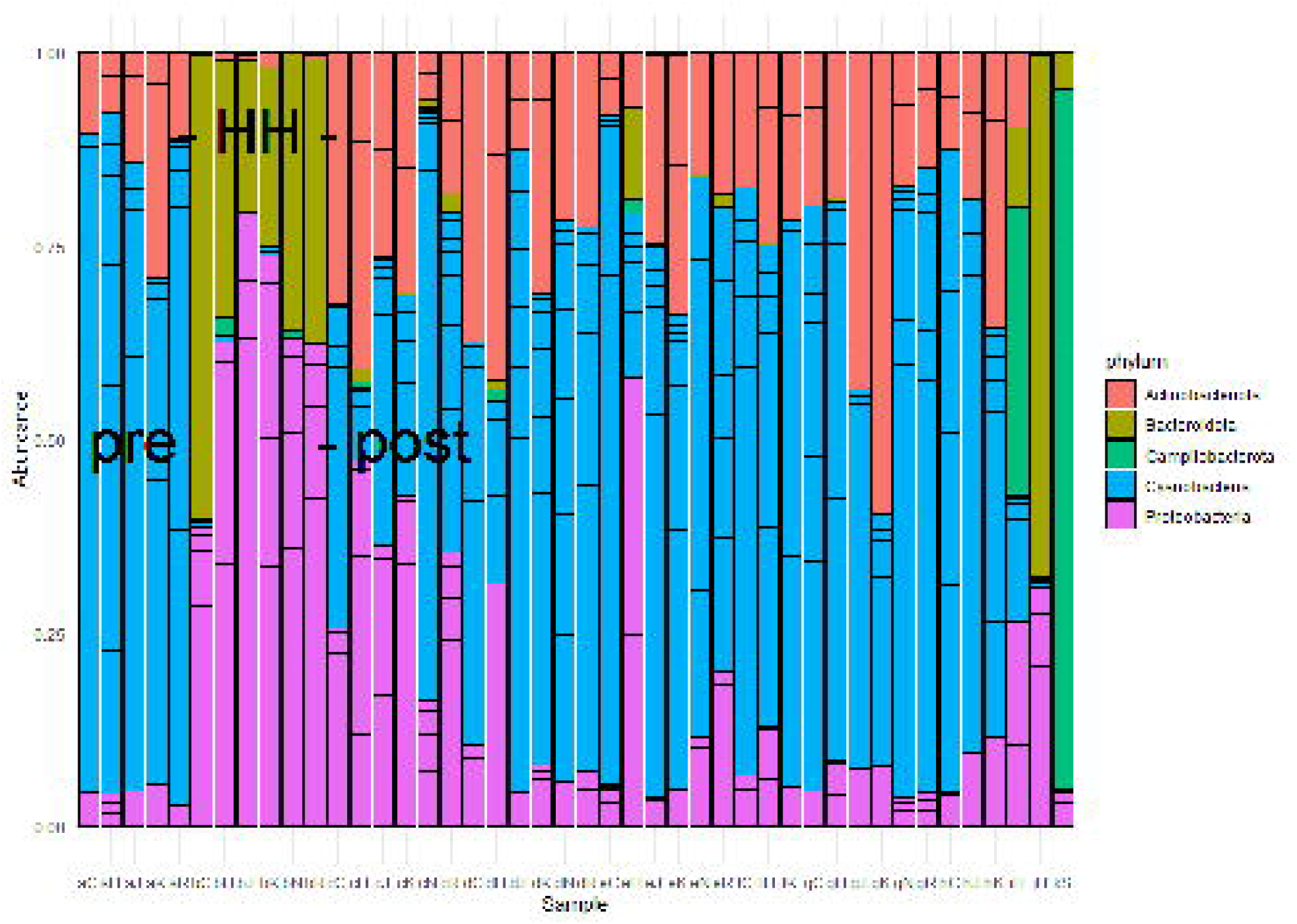
Relative abundance of numerically dominant microbes in Clear Lake for samples collected from August 2017 to March 2018. Samples are plotted chronologically from before Hurricane Harvey to 2018. Sample codes indicate date and location sampled respectively. The first letter (lower case) indicates dates in Figure 2, where a = August 25^th^, b = August 30^th^ etc. The second letter (upper case) indicates stations in Figure 1. Sample jH indicates a sample collected in March 2018 at station H. Sample kS corresponds to sample jH spiked with sewage. Only the twenty most abundant ASV are shown. Figure was generated with scripts in File S05.

Sample type predicted the majority of abundances. For example, the abundance of 819 (51%) ASVs differed between pre-HH and HH samples and the abundance of 1007 (62%) ASVs differed between pre-HH and post-HH samples. Of the ten most abundant ASVs observed in samples collected immediately after HH, nine classified as γ-*Proteobacteria*. Most of these (7/9) classified within the family *Comamonadaceae* and showed similarity to bacteria typically observed in freshwater systems. For example, ASV18 showed similarity to *Limnohabitans curvus* MWH-C1a, which was isolated from a lake [24]. The other two highly abundant ASVs (ASV64 and ASV23) showed similarity to *γ-Proteobacteria* isolated from rhizosphere soil [25] and freshwater systems [22], respectively. ASV6, the most abundant ASV in libraries generated from floodwater samples, accounted for 5 – 17 % of the reads generated in those six libraries. This ASV showed similarity to *Aquirufa* strains isolated from lakes [23, 31]. The ASV showed the greatest differential abundance between pre-HH and HH samples (ASV103) showed high similarity to two uncultured bacteria (KP686762 and KP686755) generated from floodwater collected in North Carolina immediately after Hurricane Irene [3].

PICRUSt2 analysis predicted the abundance of 418 metabolic pathways from 1,616 ASVs generated from 44 samples. Differential abundance analysis suggested that 76 of these pathways were significantly different between samples. Cluster analysis, based on the relative abundance of these 76 pathways, suggested that floodwater samples formed a coherent group (Fig. 10). That is, with one exception (sample eH), floodwater samples were relatively similar to each other in terms of predicted pathways. The outlying sample was also similar to samples collected immediately after HH in terms of numerically abundant phyla (Fig. 9). In particular, eH and floodwater samples showed relatively high proportions of *Proteobacteria* and *Bacteroidetes*.

**Figure 10.**
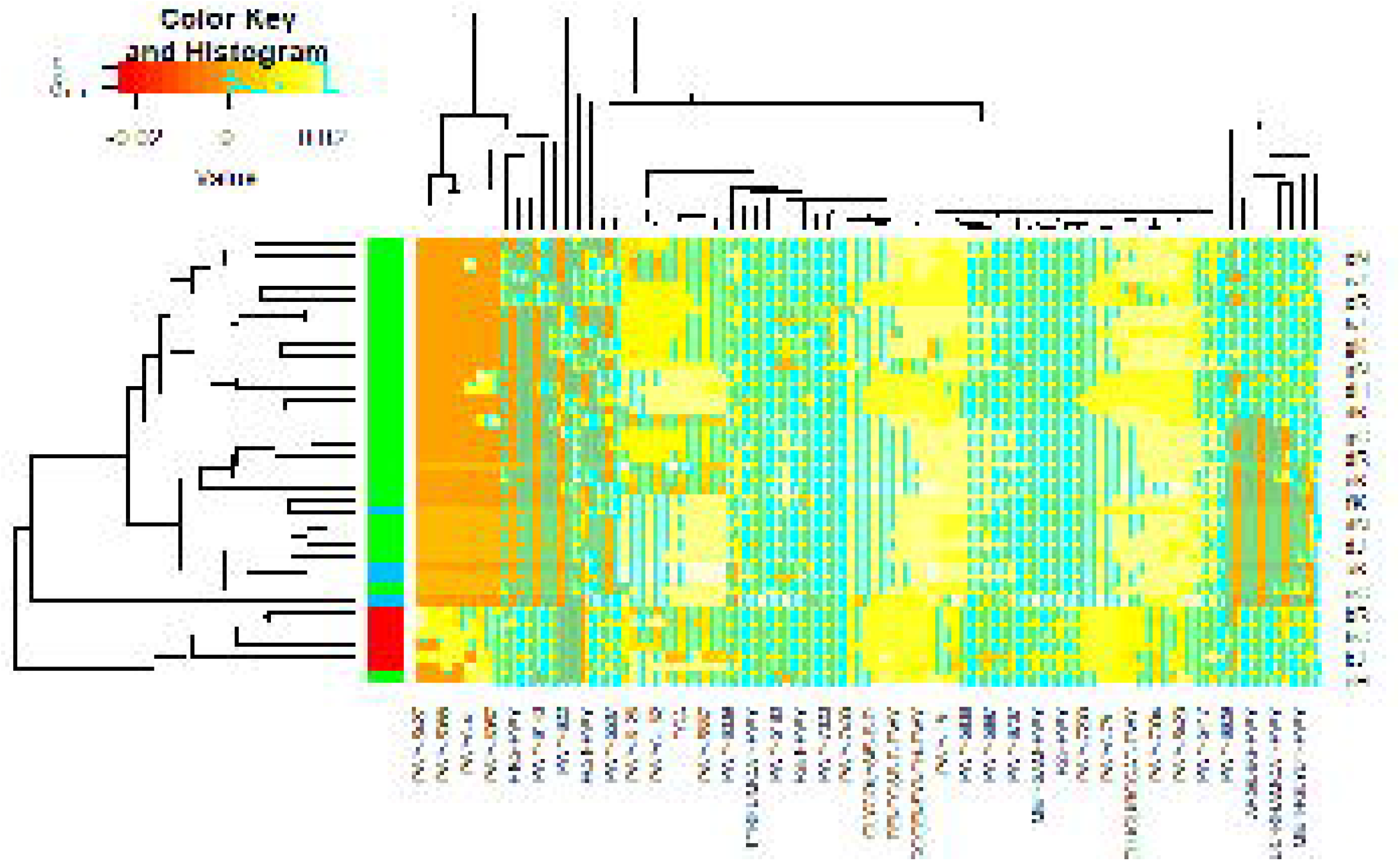
Heatmap of Metabolic Pathway Abundances. Pathways on lower horizontal axis were predicted from ASVs with PICRUSt2 as described in Methods. Representative sample codes (see Figure 9) are indicated on right vertical axis. Sample types are indicated by colors in block on left vertical axis, where blue = pre, red = HH and green = post. Figure was generated with scripts in File S16.

Comparison of pathways predicted from samples collected immediately before and after HH, identified 29 differentially abundant pathways (Fig. S14); 14 of these were significantly higher in samples collected before the storm and 15 were significantly higher after the storm. Half (7/14) of the pathways associated with samples collected before HH were biosynthesis pathways. These include PWY-5347, which produces methionine and PWY-5840, which produces menaquinol-7. In contrast, only 3 of 14 pathways that were differentially abundant in samples collected immediately after HH were biosynthesis pathways, and two of these biosynthetic pathways are associated with virulence. PWY0-1338 confers resistance to the antibiotic polymyxin and PWY-6143 produces pseudaminic acid, which is associated with pathogenic Gram negative bacteria [49]. The vast majority (11/15) of pathways that were more abundant in samples collected immediately after HH were degradation pathways. These include pathways ORNDEG-PWY, ARGDEG-PWY and ORNARGDEG-PWY, which are associated with degradation of L-arginine, putrescine, 4-aminobutanoate and L-ornithine [10].

Salinity and NO_x_ levels appeared associated with the abundance of 30 pathways. Of these 12 of were associated positively with salinity and 13 were associated negatively (Fig. S13). All but one of the pathways positively associated with salinity were biosynthesis pathways. These included five pathways (PWY-6165, 6349, -6350, -6654, -6167) associated with archaea and PWY-622, which is associated with starch biosynthesis by photoautotrophs [10]. In contrast, 6 of 13 pathways negatively associated with salinity were degradation pathways. These included two pathways (PWY-5427, - 6956) associated with naphthalene degradation by bacteria and PWY-5088, which is associated with glutamate degradation by members of the Firmicutes phylum [10]. NO_x_ concentrations appeared associated with the abundance of five pathways (Fig. S14). The two positively associated pathways were degradation pathways; both are associated with mandelate degradation by *Proteobacteria*. Pathways negatively associated with NO_x_ concentrations include PWY-6174, which is associated with the mevalonate pathway in archaea, and PWY-5183, which is associated with toluene degradation by *Proteobacteria* [10].

## Discussion

Rising sea levels and warming waters, associated with global warming, are predicted to increase the frequency of coastal flooding [60]. Global warming is also expected to increase the severity of hurricanes [28]. These climate driven changes could alter the structure of coastal systems and offshore systems [51] more frequently bring many people into contact with floodwater, which creates a public health risk [6, 16]. The response of the system and risks to the populace will vary depending on the system and storm. Here we studied the water quality and microbial communities of samples collected from the Clear Lake system, a rapidly developing area between Houston and Galveston. Hurricane Harvey temporarily shifted the structure of the Clear Lake system from a mesosaline (∼ 10 PSU), estuary, fed by eutrophic, fresh tributaries, to a freshwater system, with little difference between the lake and tributaries in terms of salinity, nutrients and other chemical parameters.

The temporary shift to a freshwater system was accompanied with a dramatic decrease in cyanobacteria and in increase in *γ-Proteobacteria* typically observed in soils and freshwater systems. This pattern of dilution and recovery is consistent with a model of the recovery time for salinity in that system [15], but is a few weeks slower for the time reported for salinity recovery for Galveston Bay [53]. Overall, the recovery of the Galveston Bay system appears slower than estuaries impacted by Hurricane Bob [59] and estuaries impacted by multiple hurricanes in North Carolina [40] and the shift in bacterial community structure is consistent with changes reported following HH for Galveston Bay [65].

Bacteria dominated this system, as assessed by metagenomic analysis of PCR-amplified 16S rRNA gene fragments, before and after HH and the structure of this community corresponded to salinity and nutrient concentrations. The relationship between salinity and bacterial community structure parallels a report that salinity corresponded to changes in viral community structure in Galveston Bay following HH [63]. These results agree with previous studies of systems in Louisiana impacted by Hurricanes Katrina and Rita [2]. These results also agree with previous reports for estuaries in general [56], and the dogma that nitrogen limits productivity in coastal systems. That is, if nitrogen limits primary production, we would expect that a change in nitrogen availability would change the entire system. Indeed, low N/P ratios suggests that nitrogen limits productivity in Clear Lake, which is consistent with Ryther and Dunstan’s dogma [48]; however, only inorganic nutrients were measured herein. Organic matter also contains significant pools of nitrogen and phosphate. For example, in Galveston Bay total nitrogen concentrations were about 5X higher than DIN concentrations for samples collected following HH [53].

Oxygen was not depleted significantly in water segments sampled herein following HH, relative to pre-storm levels and historical records, and hypoxic conditions (< 3 mg/L) were only observed once in this study. This agrees with a previous report for Bayous in the Houston-Galveston area, where relatively rural watersheds receiving waters, like Peach Creek, did not go hypoxic, with the exception of the headwaters of Clear Creek [27]. The general lack of hypoxia in this system contrasts with previous reports for other systems in the Gulf of Mexico. For example, hypoxia persisted in Pensacola Bay for months following Hurricane Ivan [21] and floodwaters overlying New Orleans were hypoxic following Hurricane Katrina [39].

High *E. coli* MPNs for samples collected immediately after HH, suggests that floodwaters were contaminated with fecal matter. These elevated MPNs agree with previous reports for Bayous within the Galveston Bay system [27, 66, 67], for the Guadalupe River [26], which was also in the path of HH, and the report of Enterobacteriaceae in marine sponges offshore of Galveston Bay [51]. The EPA and TCEQ [55] recommend *Enterococci* for estuaries and coastal waters; however, because of logistical issues, *Enterococci* MPNs were not available for several weeks after HH. Levels of these FIB remained elevated relative to typical levels for this system for weeks (Fig. S07). These high MPNs agree with the observation that bacteria typically observed in human waste, such as *Bacteroides* spp., abounded in libraries generated from all samples collected immediately following HH (Fig. 9).

PICRUSt2 analysis suggested that flooding also enriched for antibiotic resistant genes (ARG), virulence factors and carbon cycling pathways. These predictions of functional genes from rRNA sequences, and the inference of microbial community structure from targeted metagenomic analysis in general, should be treated with caution. Every step in targeted metagenomic analysis, from sampling to data analysis is fraught with bias [44]. In particular, PICRUSt2 depends on reference genomes, which are largely derived from the human gut microbiome. This creates a bias depending on the sample type [54]. For example, PICRUSt2 underestimates certain pathways in soil systems [57].

Nevertheless, prediction of ARG and virulence factors with PICRUSt2 did agree with previously published qPCR measurements of ARG in samples collected from soils flooded during HH [41], in samples collected within Galveston Bay two weeks after HH [66], and ARG and pathogens in floodwaters and bayous following HH [67]. Prediction of increase in carbon cycling bacteria similarly agrees with reports that loading of dissolved organic carbon (DOC) during extreme weather events can enhance carbon cycling by bacterial communities in receiving waters [3] and high DOC levels in Galveston Bay following HH [53, 65].

## Conclusions

The massive influx of freshwater from Hurricane Harvey into the Clear Lake system temporarily changed the system from an estuary, with relatively low levels of FIB and a microbial community dominated by primary producers, to a freshwater system, with high levels of FIB. The microbial community observed immediately following the hurricane included bacteria that have also been reported in estuaries following hurricanes, but rarely elsewhere, and enrichment of antibiotic resistant bacteria. It took the system a couple of months to recover, in terms of nutrients and salinity, to pre-storm conditions.

## Acknowledgments

We thank Diep Le and Theodore Richardson for assistance with laboratory and data analysis support respectively. This work was supported by NSF awards 1759542 and 1759540.

## Supplemental Figure Legends

**Figure S01.**
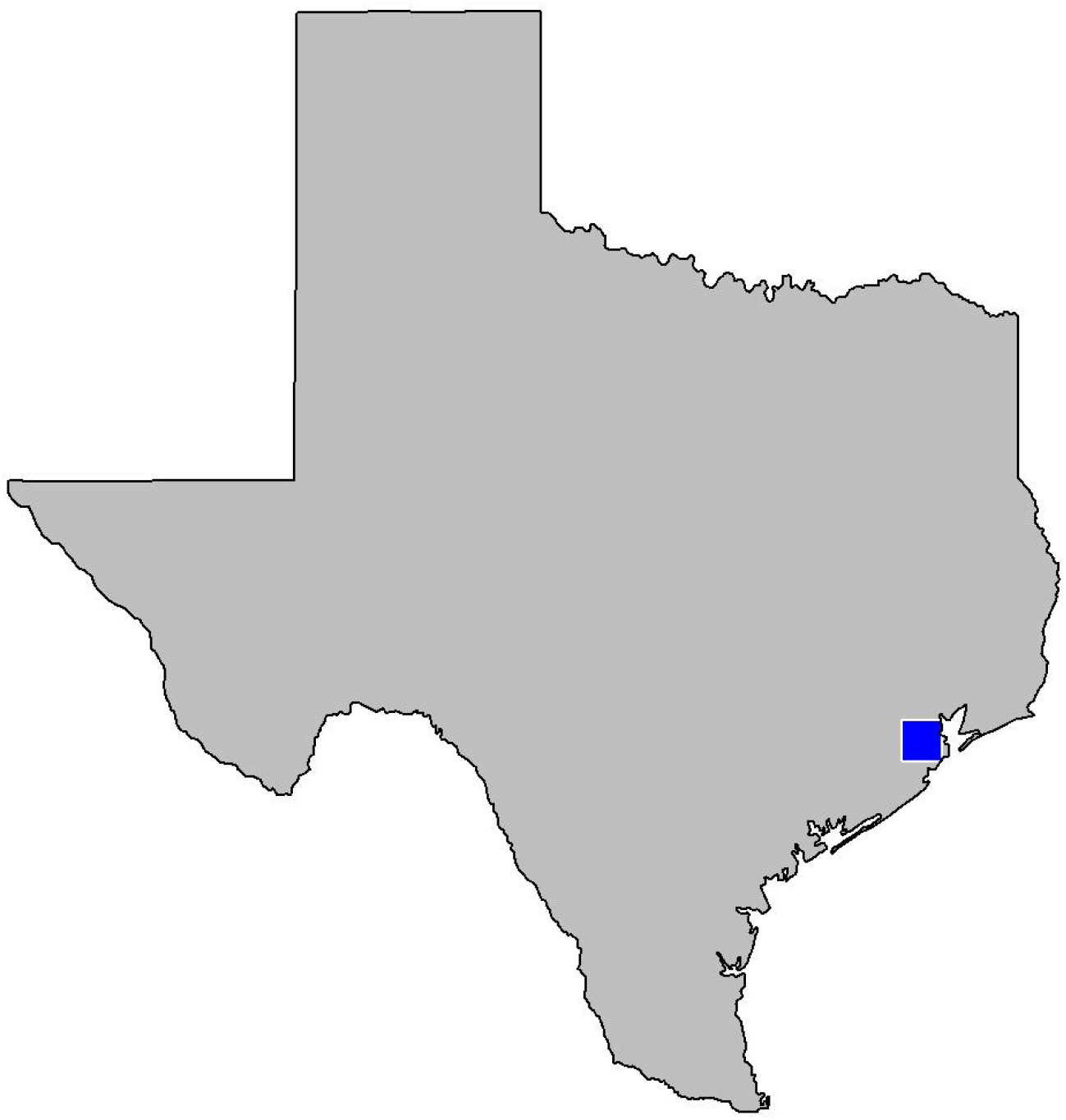
Study location. Figure was generated with scripts in File S01.

**Figure S02.**
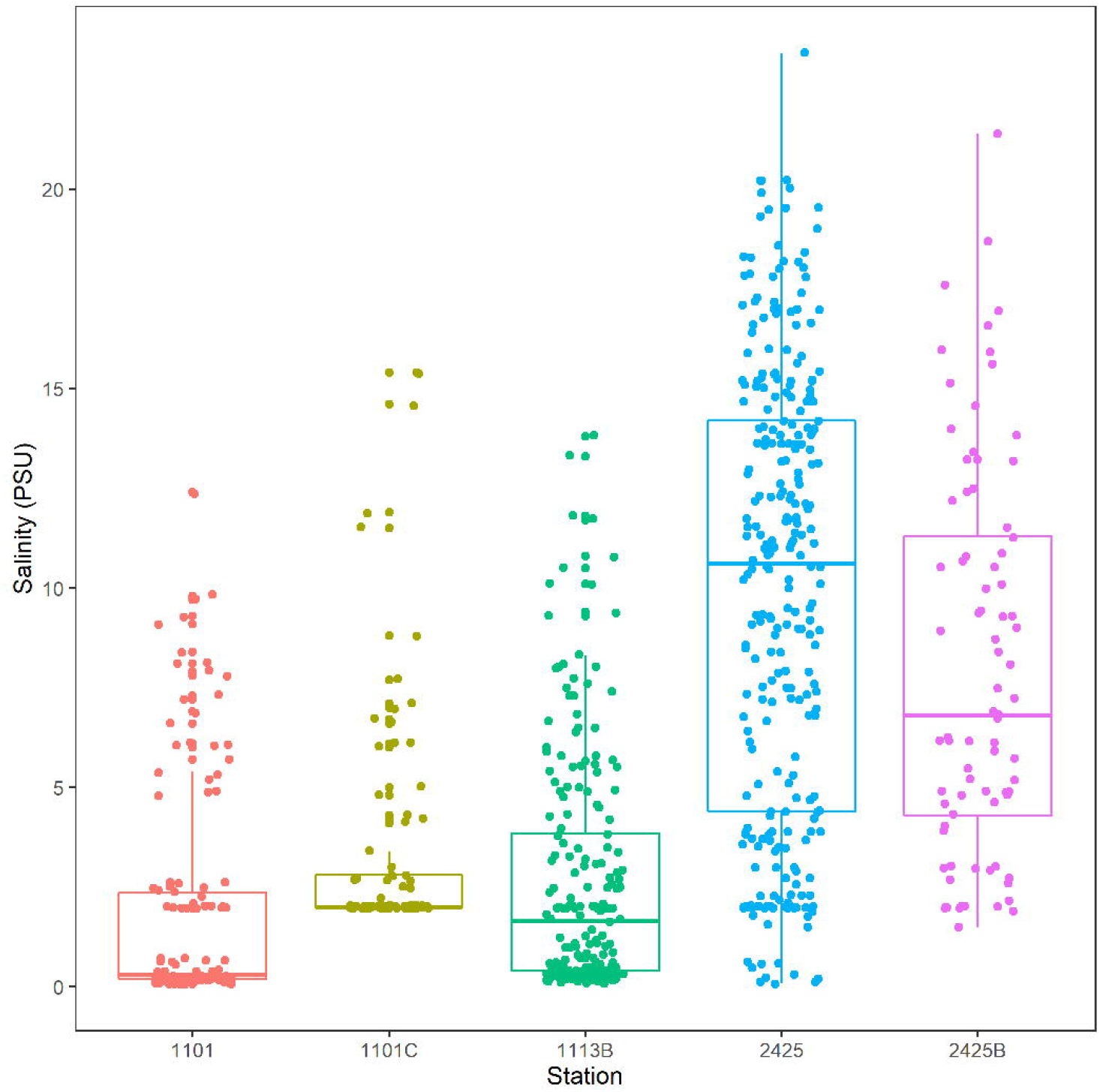
Historical salinity data for Clear Lake system. Stations correspond to segments in Figure 1. Data is from 2011 to 2021. Figure was generated with scripts in File S02.

**Figure S03.**
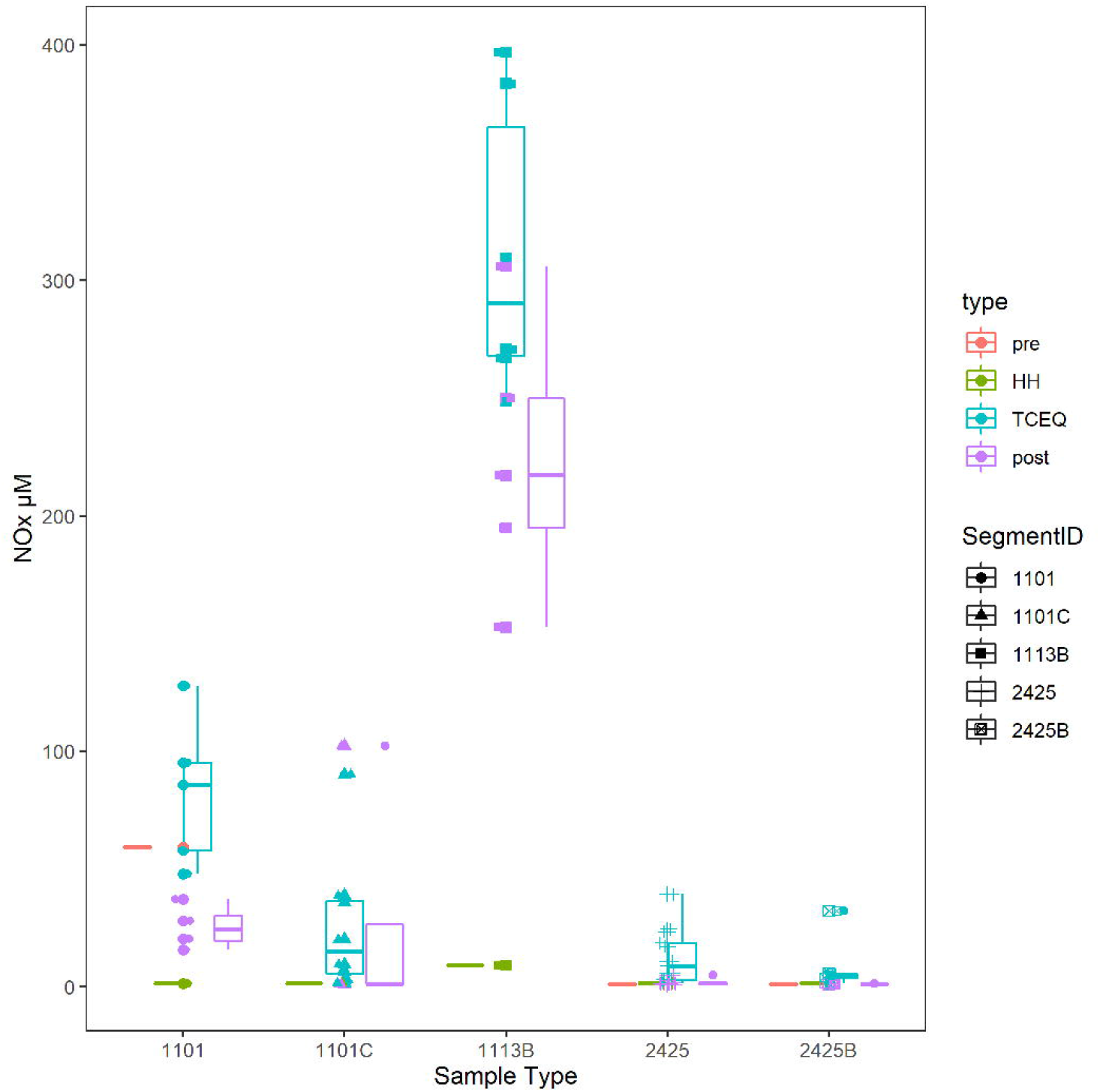
Nitrate/nitrite concentration by water body segments and sample type. Type and segment are as Figure 4. Figure was generated with scripts in File S03

**Figure S04.**
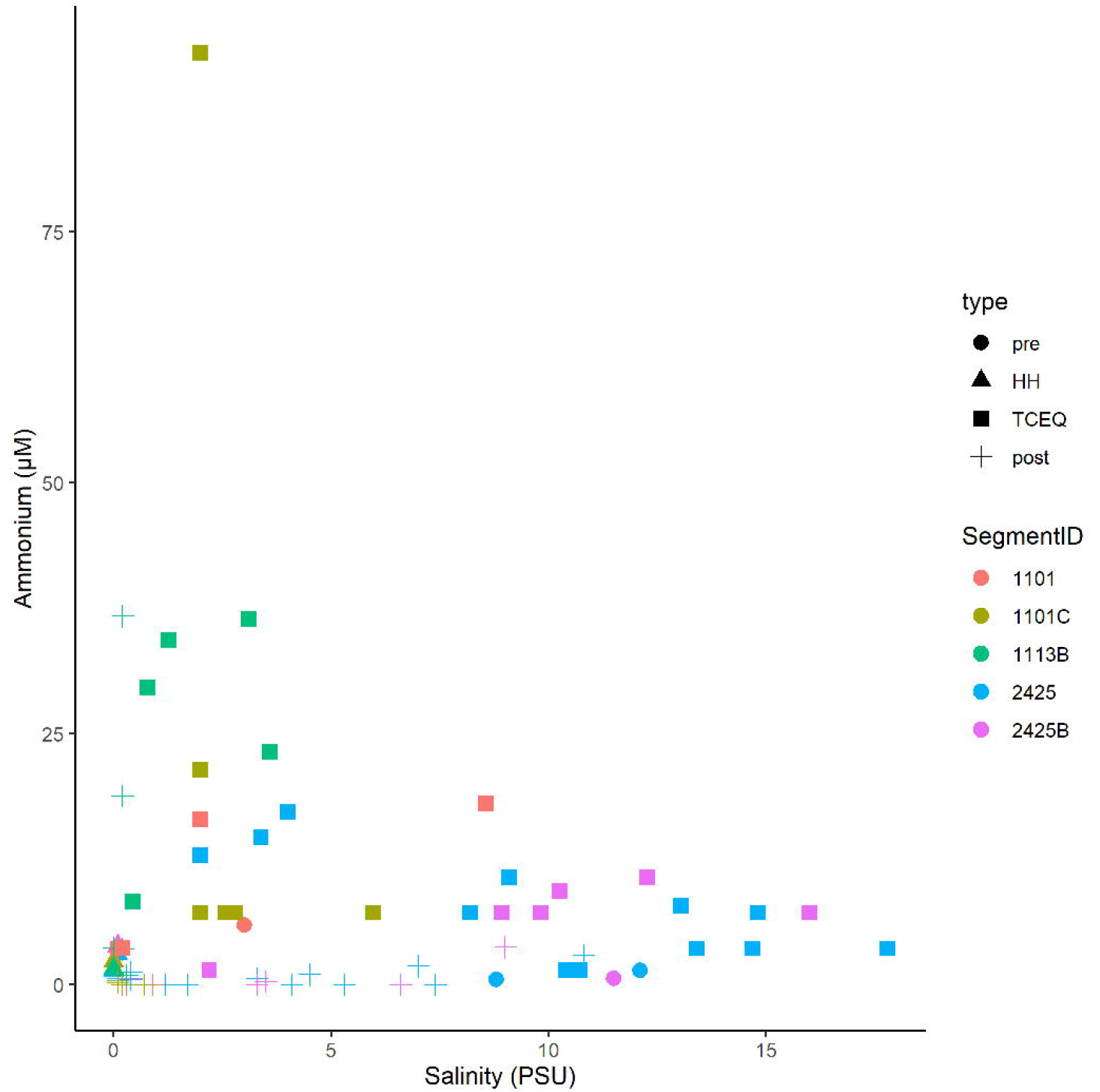
Mixing diagram of salinity versus ammonium for the Clear Lake system. Symbols are as Figure 3. Figure was generated with scripts in File S03.

**Figure S05.**
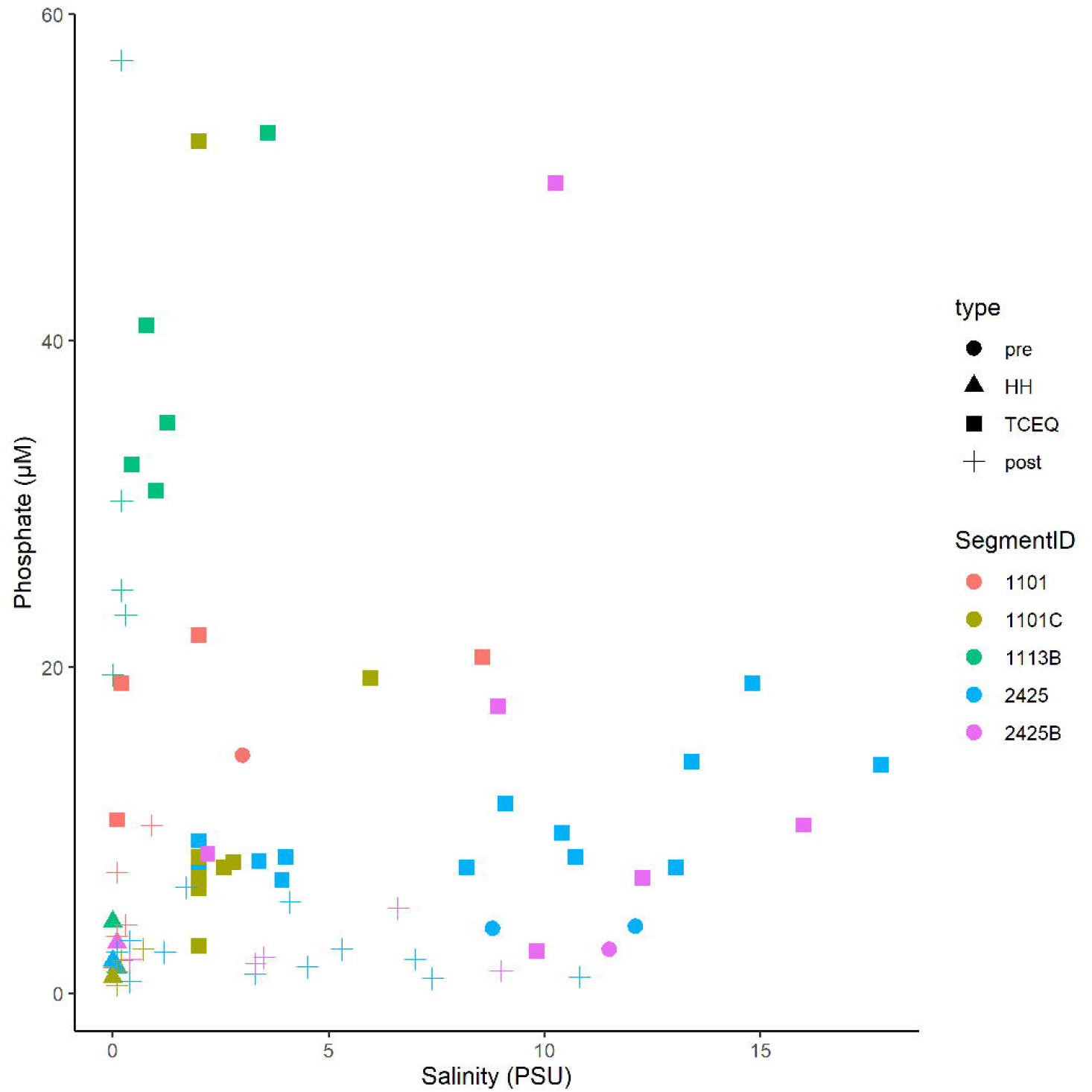
Mixing diagram of salinity versus phosphate for the Clear Lake system. Symbols are as Figure 3. Figure was generated with scripts in File S03.

**Figure S06.**
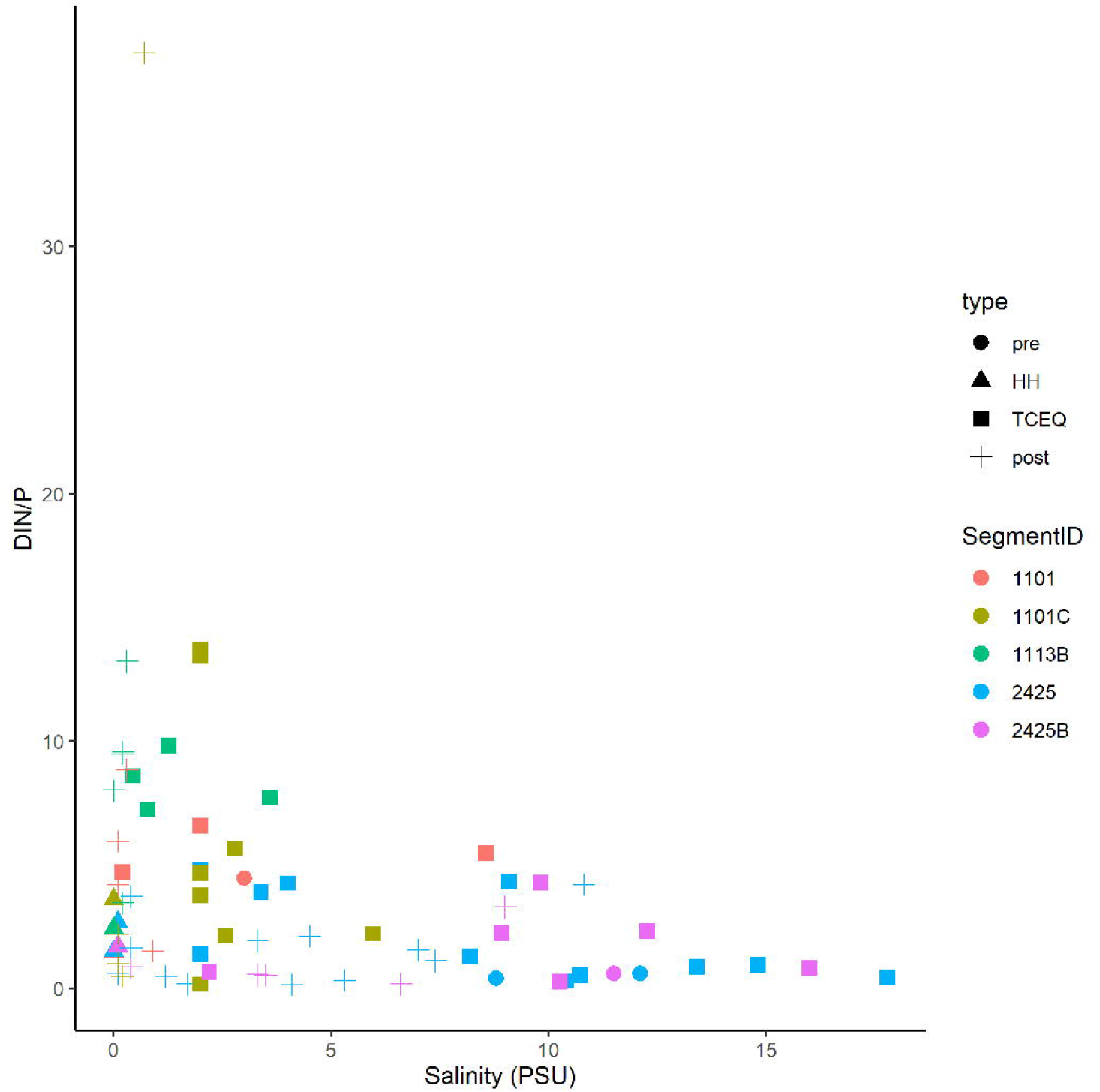
Mixing diagram of N/P ratio versus phosphate for the Clear Lake system. Symbols are as Figure 3. Figure was generated with scripts in File S03.

**Figure S07.**
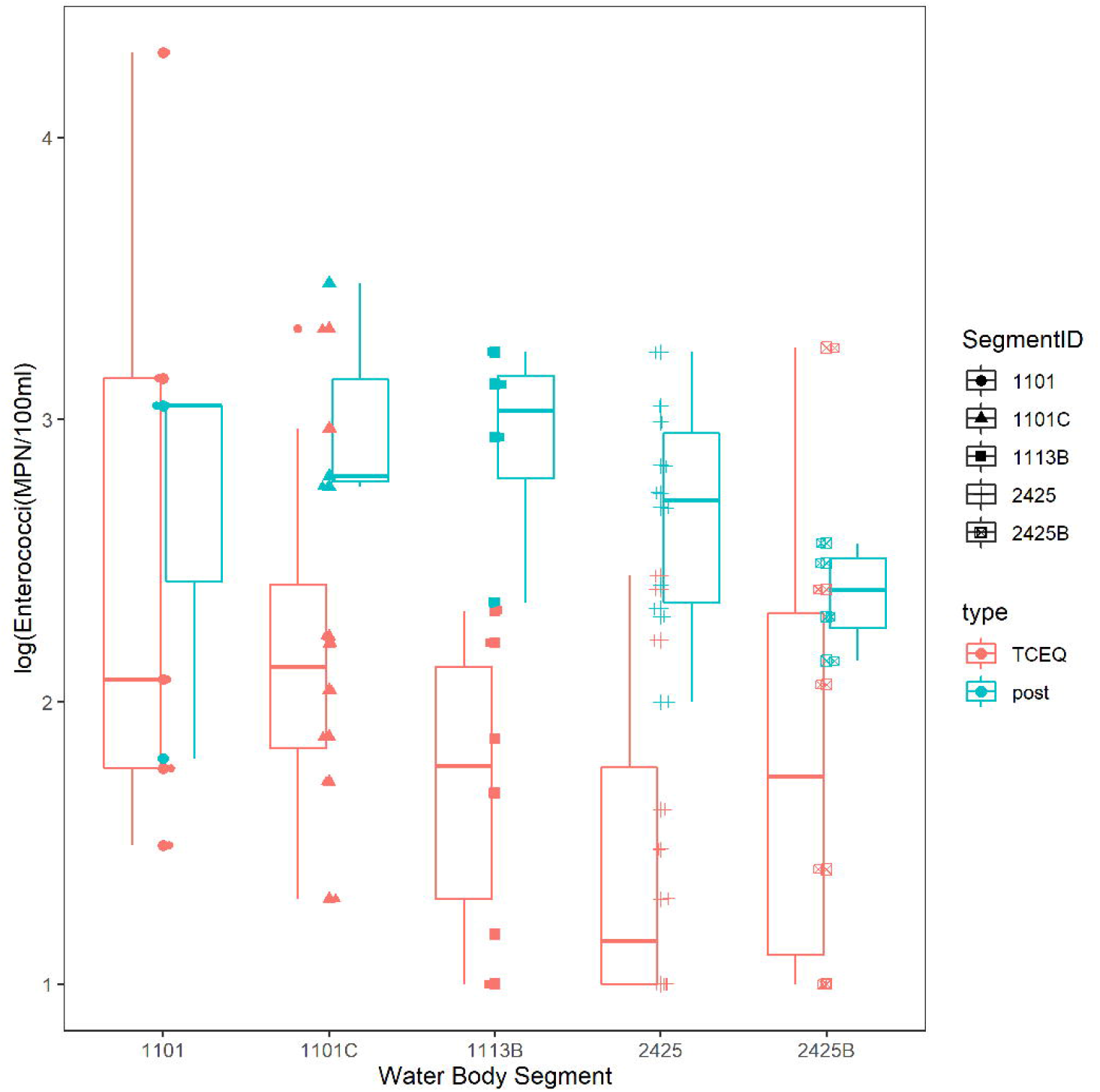
Comparison of historical *Enterococci* concentrations versus concentrations following Hurricane Harvey. Symbols are as Figure 3. Figure was generated with scripts in File S03.

**Figure S08.**
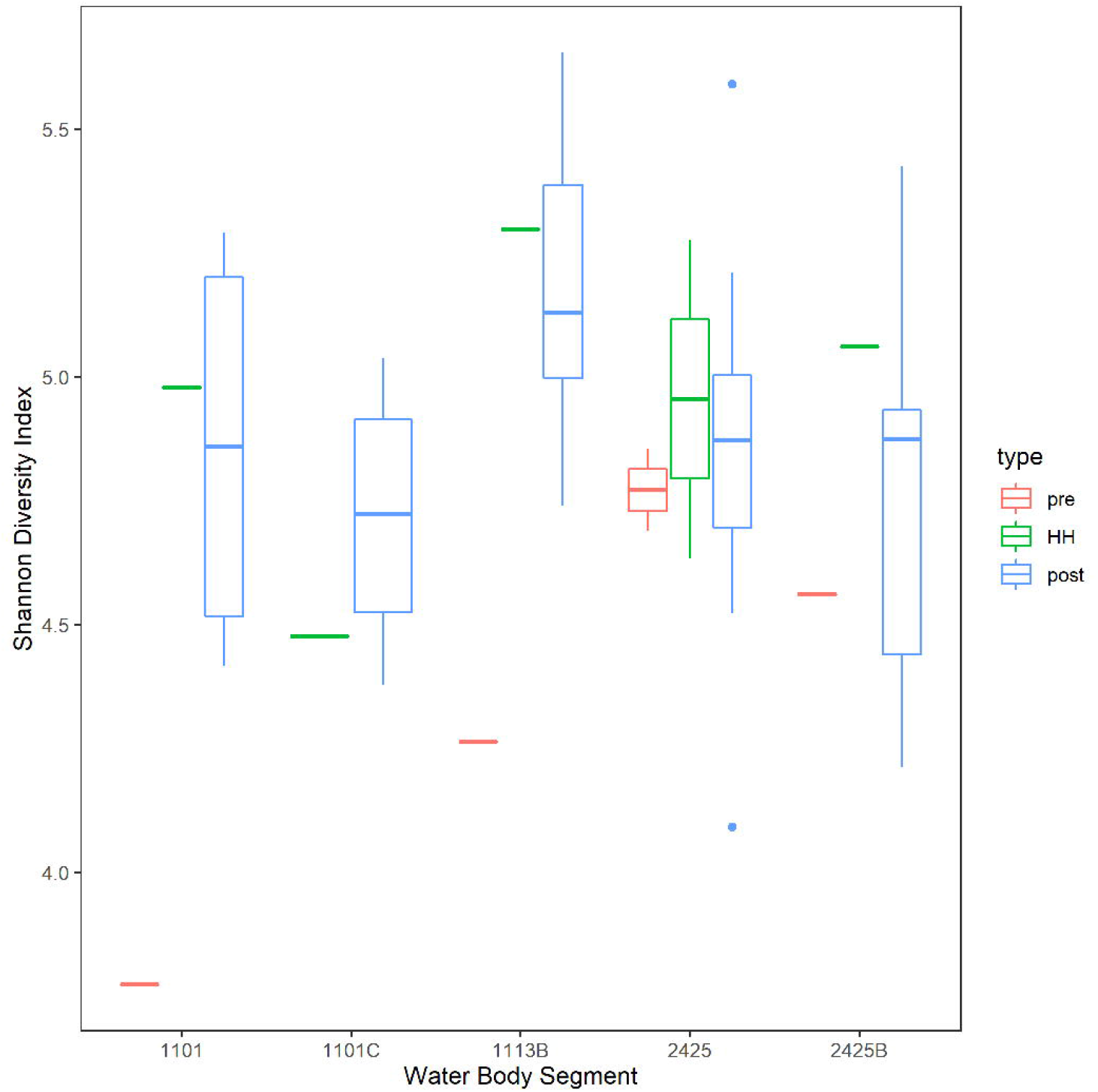
Alpha diversity of bacterial communities in the Clear Lake system before and after Hurricane Harvey. Shannon diversity indices were calculated by targeted metagenomic analysis, as described in Methods. Boxes, whiskers and horizontal lines are described in Figure 2. Sample types are described in Figure 3. Figure was generated with scripts in File S05.

**Figure S09.**
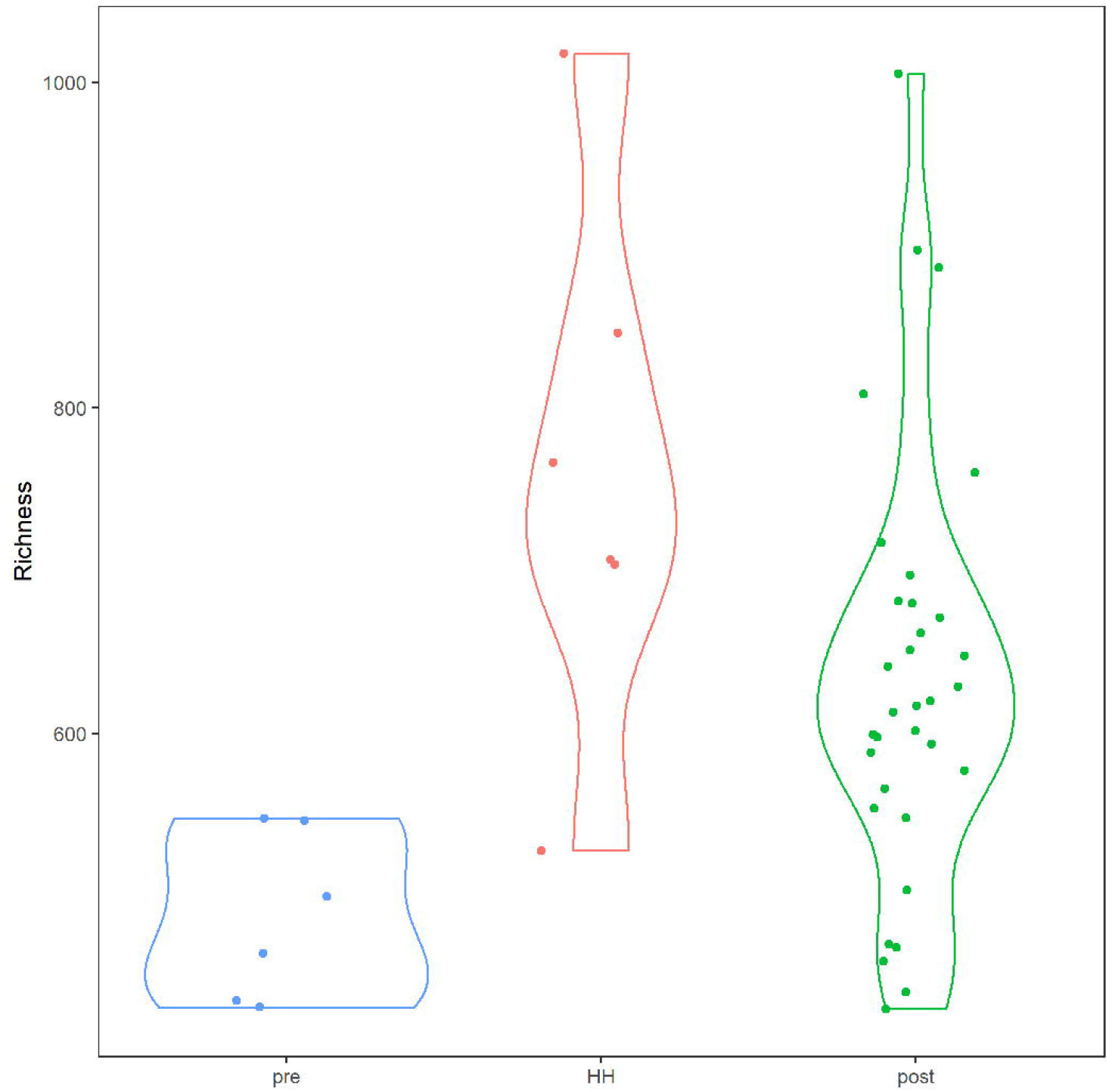
Microbial richness in Clear Lake system before and after Hurricane Harvey. X-axis indicates number of amplicon sequence variants. Categories correspond to before (pre), immediately after (HH) and more than a week after (post). Figure was generated with scripts in File S05.

**Figure S10.**
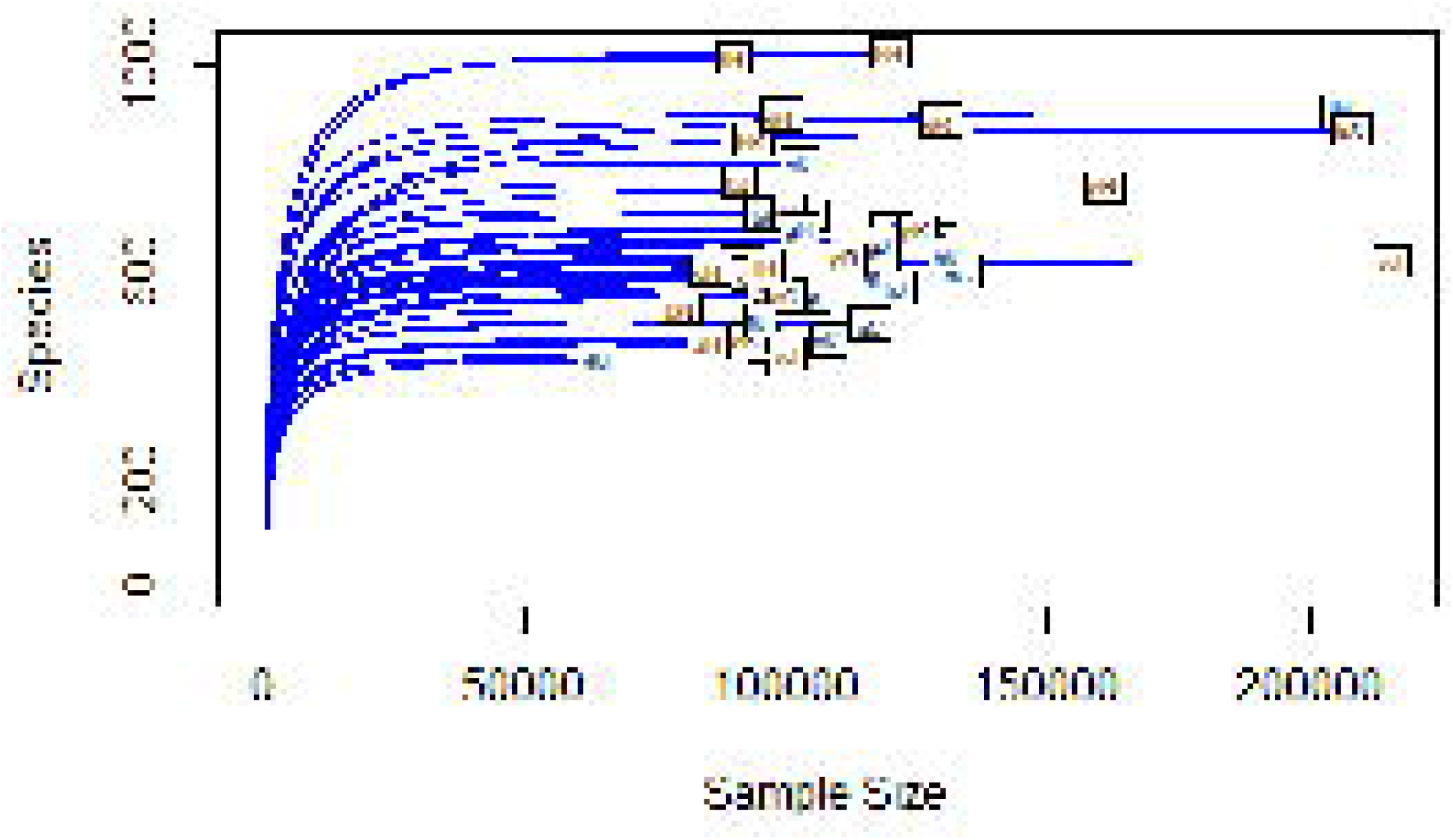
Rarefaction analysis of richness by number of reads. Symbols are as Figure 9. Figure was generated with scripts in File S05.

**Figure S11.**
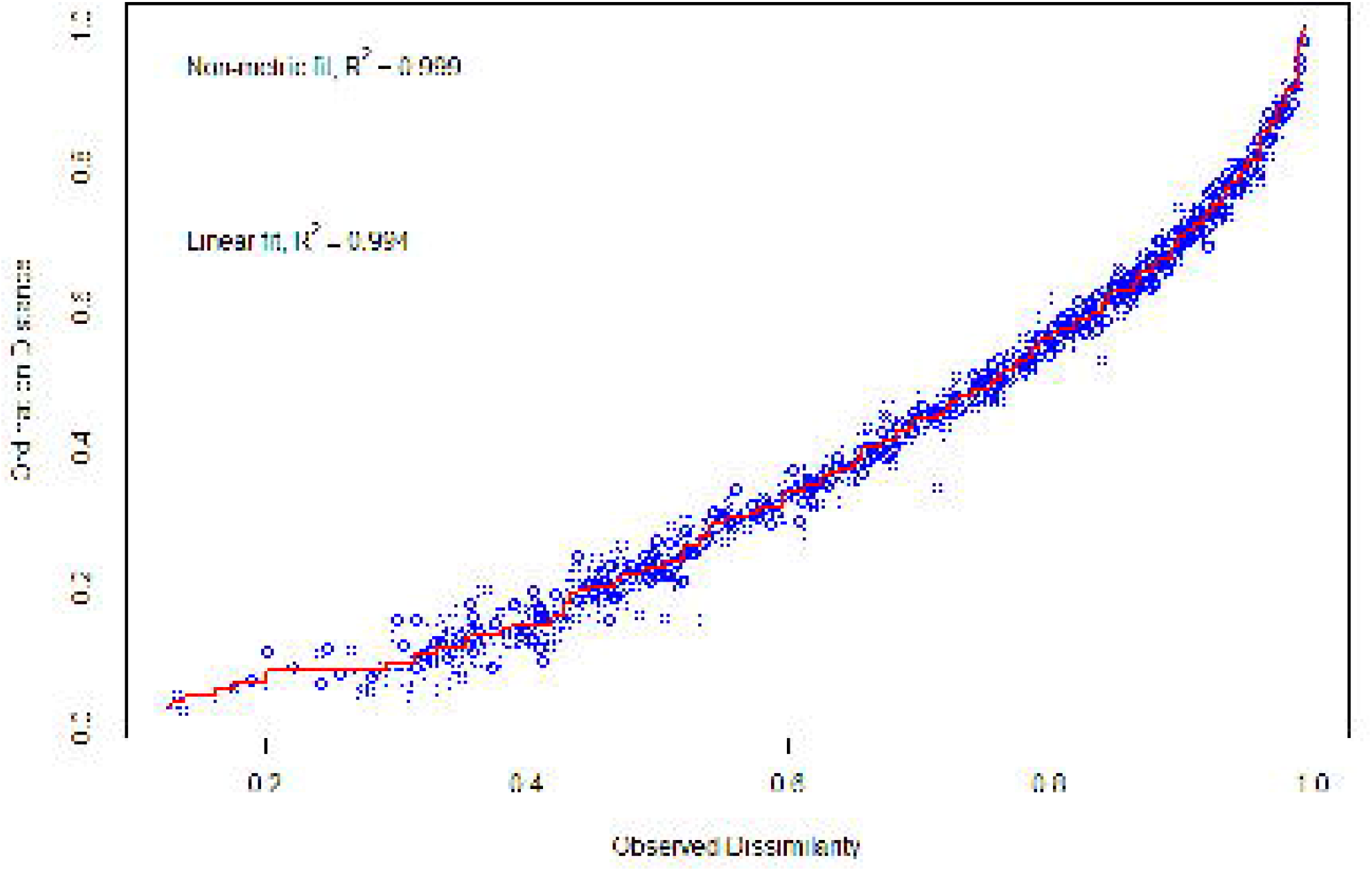
Shepards diagram showing fit of NMDS to dissimilarity of any two pairs of samples. Figure was generated with scripts in File S08.

**Figure S12.**
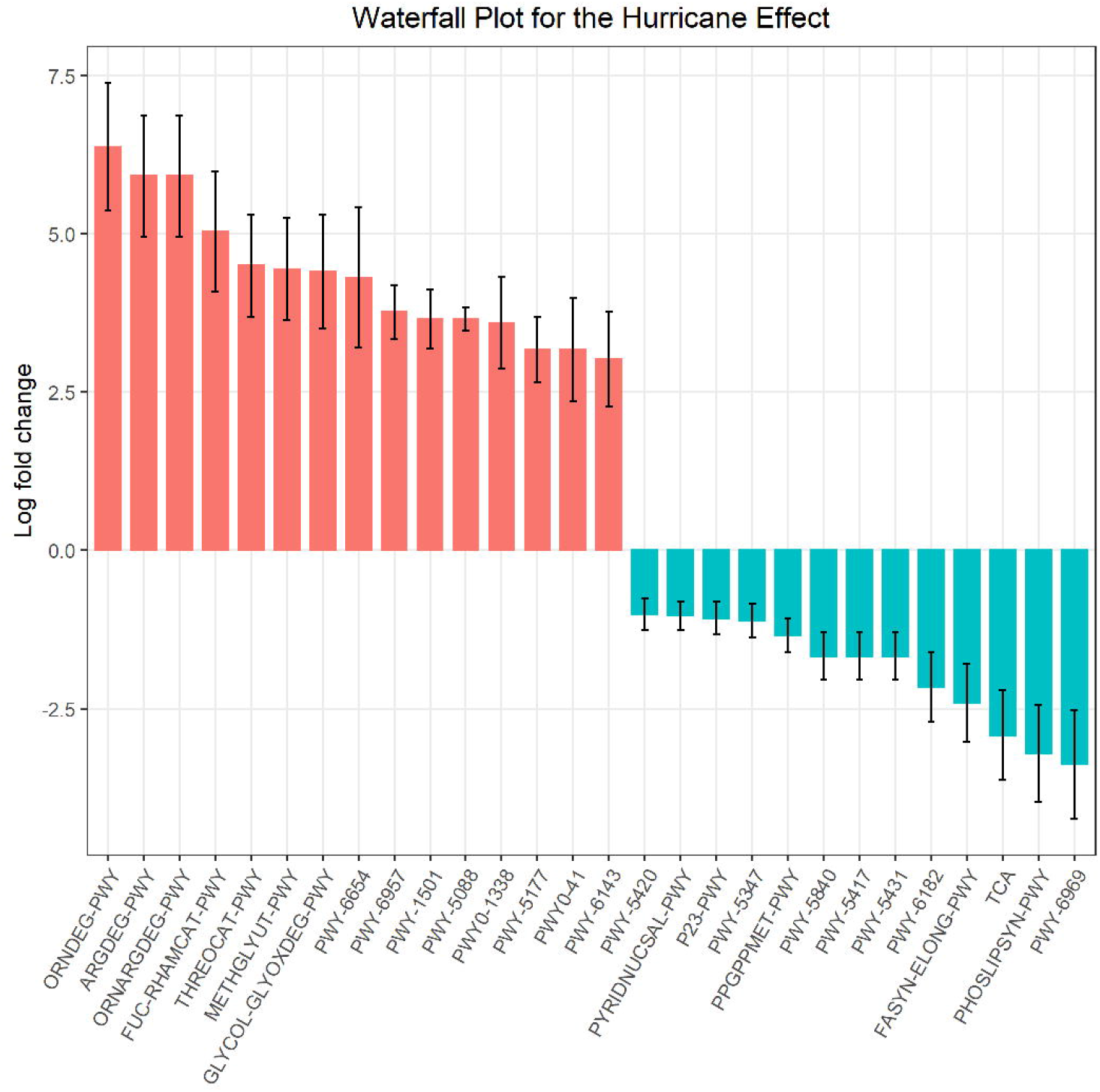
Waterfall plot showing pathways enriched (red) or depleted (teal) in comparsion of samples collected before or immediately after Hurricane Harvey. Figure was generated with scripts in File S15.

**Figure S13.**
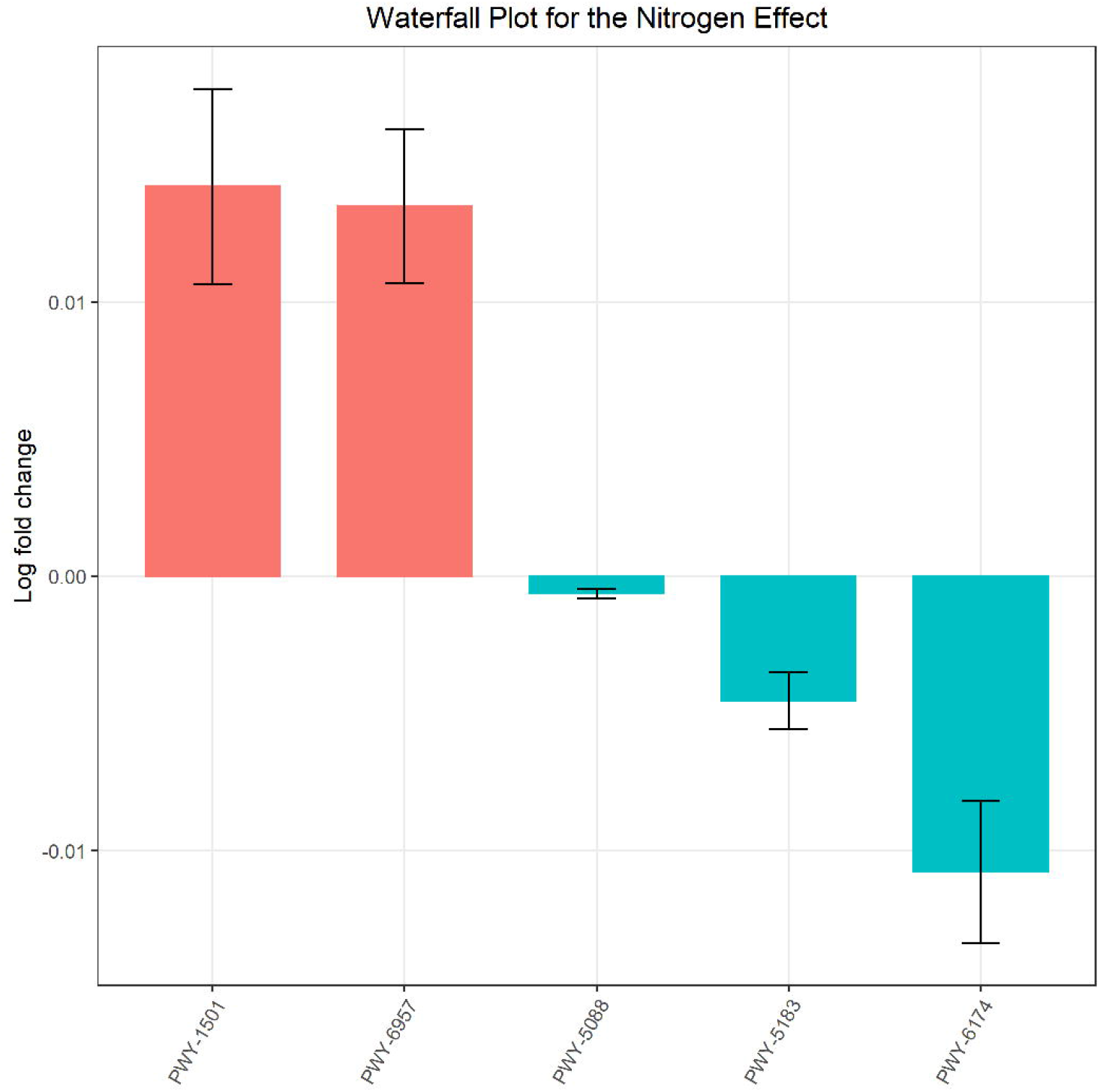
Waterfall plot showing pathways enriched (red) or depleted (teal) as function of nitrate concentration. Figure was generated with scripts in File S15.

