## Supplemental File S01 for "Hurricane Harvey Impacts on Water Quality and Microbial Communities in Houston, TX Waterbodies": S01_map.html

HHmap


### HHmap

###### Michael G. LaMontagne

#### 11/21/2021

#### R Markdown

This R Markdown document presents generation of maps showing region of study and sampling stations.

```
library(devtools)
```

```
## Loading required package: usethis
```

```
library(ggplot2)
library(ggmap)
```

```
## Google's Terms of Service: https://cloud.google.com/maps-platform/terms/.
```

```
## Please cite ggmap if you use it! See citation("ggmap") for details.
```

```
library(maps)
library(mapdata)
library(dplyr)
```

```
## 
## Attaching package: 'dplyr'
```

```
## The following objects are masked from 'package:stats':
## 
##     filter, lag
```

```
## The following objects are masked from 'package:base':
## 
##     intersect, setdiff, setequal, union
```

```
sessionInfo()
```

```
## R version 4.1.1 (2021-08-10)
## Platform: x86_64-w64-mingw32/x64 (64-bit)
## Running under: Windows 10 x64 (build 22000)
## 
## Matrix products: default
## 
## locale:
## [1] LC_COLLATE=English_United States.1252 
## [2] LC_CTYPE=English_United States.1252   
## [3] LC_MONETARY=English_United States.1252
## [4] LC_NUMERIC=C                          
## [5] LC_TIME=English_United States.1252    
## 
## attached base packages:
## [1] stats     graphics  grDevices utils     datasets  methods   base     
## 
## other attached packages:
## [1] dplyr_1.0.7     mapdata_2.3.0   maps_3.4.0      ggmap_3.0.0.903
## [5] ggplot2_3.3.5   devtools_2.4.2  usethis_2.1.3  
## 
## loaded via a namespace (and not attached):
##  [1] Rcpp_1.0.7          lattice_0.20-44     tidyr_1.1.4        
##  [4] prettyunits_1.1.1   png_0.1-7           ps_1.6.0           
##  [7] assertthat_0.2.1    rprojroot_2.0.2     digest_0.6.27      
## [10] utf8_1.2.2          R6_2.5.1            plyr_1.8.6         
## [13] evaluate_0.14       httr_1.4.2          pillar_1.6.4       
## [16] RgoogleMaps_1.4.5.3 rlang_0.4.11        rstudioapi_0.13    
## [19] callr_3.7.0         jquerylib_0.1.4     rmarkdown_2.11     
## [22] desc_1.4.0          stringr_1.4.0       munsell_0.5.0      
## [25] compiler_4.1.1      xfun_0.25           pkgconfig_2.0.3    
## [28] pkgbuild_1.2.0      htmltools_0.5.2     tidyselect_1.1.1   
## [31] tibble_3.1.4        fansi_0.5.0         crayon_1.4.1       
## [34] withr_2.4.2         bitops_1.0-7        grid_4.1.1         
## [37] jsonlite_1.7.2      gtable_0.3.0        lifecycle_1.0.1    
## [40] DBI_1.1.1           magrittr_2.0.1      scales_1.1.1       
## [43] cli_3.0.1           stringi_1.7.5       cachem_1.0.6       
## [46] fs_1.5.0            remotes_2.4.1       sp_1.4-5           
## [49] testthat_3.1.0      bslib_0.3.1         ellipsis_0.3.2     
## [52] generics_0.1.1      vctrs_0.3.8         tools_4.1.1        
## [55] glue_1.4.2          purrr_0.3.4         jpeg_0.1-9         
## [58] processx_3.5.2      pkgload_1.2.3       fastmap_1.1.0      
## [61] yaml_2.2.1          colorspace_2.0-2    sessioninfo_1.1.1  
## [64] memoise_2.0.0       knitr_1.36          sass_0.4.0
```

#### Get state data

```
# get state data
states <- map_data("state")
# just TX
tx_df <- subset(states, region == "texas")
head(tx_df)
```

```
##            long      lat group order region subregion
## 12203 -94.49792 33.66700    50 12203  texas      <NA>
## 12204 -94.48074 33.65554    50 12204  texas      <NA>
## 12205 -94.47501 33.63262    50 12205  texas      <NA>
## 12206 -94.46355 33.61544    50 12206  texas      <NA>
## 12207 -94.45209 33.59824    50 12207  texas      <NA>
## 12208 -94.42345 33.59824    50 12208  texas      <NA>
```

#### Plot state with no frills

```
tx_base <- ggplot(data = tx_df, mapping = aes(x = long, y = lat, group = group)) + 
  coord_fixed(1.3) + 
  geom_polygon(color = "black", fill = "gray")
tx_base + theme_nothing()
```

#### Include study region

```
mapHGA <- as.data.frame(read.table("mapRegion.csv", sep=",", header = TRUE, check.names = TRUE))
tx_base + theme_nothing() + 
  geom_polygon(data = mapHGA, fill = "blue", color = "white") +
  geom_polygon(color = "black", fill = NA)  # get the state border back on top
```

#### Save Plot

```
ggsave("Fig_S1_txRegion.png")
```

```
## Saving 7 x 5 in image
```

#### Import stations and make box

```
# UHCL stations
mapHH <- as.data.frame(read.table("hhStations.csv", sep=",", header = TRUE, check.names = TRUE))
head(mapHH)
```

```
##   Station Latitude Longitude
## 1       C  29.5632  -95.0700
## 2       H  29.5790  -95.0926
## 3       J  29.5420  -95.0300
## 4       K  29.5484  -95.0215
## 5       N  29.5425  -95.1000
## 6       R  29.5346  -95.0962
```

```
saveRDS(mapHH, "mapHHstations.rds")
```

#### Show coordinates for seqments

```
tceqMeta <- readRDS("tceqMeta.rds")
n <- c(1:2,12:13)
tceqStation <- tceqMeta[,n]
tceqStation <- unique(tceqStation)
#row.names(tceqStation) <- NULL
#n <- c(1:2,7:11)tceqStation <- na.omit(tceqStation)
tail(tceqStation)
```

```
##       SegmentID StationID Latitude Longitude
## 6058       2425     13335 29.55555 -95.04806
## 8192      1101C     17928 29.54622 -95.10481
## 9118      1113B     11409 29.58350 -95.10356
## 10029     1113B     17317 29.57992 -95.09236
## 11091     2425B     16476 29.54056 -95.02805
## 11819     2425B     16485 29.53389 -95.03481
```

```
saveRDS(tceqStation, "tceqStation.rds")
write.csv(tceqStation, "tceqStation.csv")
```

#### Merge by SegmentID to create new spots

```
mapTCEQsegment <- tceqStation %>%
  group_by(SegmentID) %>%
  summarise(across(everything(), mean))
# mapTCEQsegment <- mapTCEQsegment[ -c(3) ]
mapTCEQsegment
```

```
## # A tibble: 5 x 4
##   SegmentID StationID Latitude Longitude
##   <chr>         <dbl>    <dbl>     <dbl>
## 1 1101         11446      29.5     -95.1
## 2 1101C        17928      29.5     -95.1
## 3 1113B        14363      29.6     -95.1
## 4 2425         13334.     29.6     -95.0
## 5 2425B        16480.     29.5     -95.0
```

```
write.csv(mapTCEQsegment, "mapTCEQsegment.csv")
```

#### Make box

```
# 
sbbox <- make_bbox(lon = tceqStation$Longitude, lat = tceqStation$Latitude, f = .15)
sbbox
```

```
##      left    bottom     right       top 
## -95.11709  29.51206 -95.01066  29.59282
```

#### Plot stations and segments

```
sq_map <- get_stamenmap(sbbox, maptype = "terrain", zoom = 14)
```

```
## Map tiles by Stamen Design, under CC BY 3.0. Data by OpenStreetMap, under ODbL.
```

```
ggmap(sq_map)
```

```
hhMapStations <- ggmap(sq_map) + 
  geom_text(data = mapHH, mapping = aes(x = Longitude, y = Latitude, label=Station), color = "red", fontface = "bold", check_overlap = FALSE) +
  geom_text(data= tceqStation, aes(x=Longitude, y=Latitude, label=SegmentID),
            color = "darkblue", fontface = "bold", check_overlap = FALSE) +
  geom_text(data= mapTCEQsegment, aes(x=Longitude, y=Latitude, label=SegmentID),
            color = "darkGreen", fontface = "bold", check_overlap = FALSE) +
  annotate(geom = "text", x = -95.06, y = 29.557, label = "Clear Lake", 
         fontface = "italic", color = "grey22", size = 4)
hhMapStations
```

#### Including Plots

```
sq_map <- get_stamenmap(sbbox, maptype = "terrain", zoom = 14)
```

```
## Map tiles by Stamen Design, under CC BY 3.0. Data by OpenStreetMap, under ODbL.
```

```
ggmap(sq_map)
```

```
hhMapSegments <- ggmap(sq_map) + 
  geom_text(data = mapHH, mapping = aes(x = Longitude, y = Latitude, label=Station), color = "red", fontface = "bold", check_overlap = FALSE) +
  geom_text(data= mapTCEQsegment, aes(x=Longitude, y=Latitude, label=SegmentID),
            color = "darkGreen", fontface = "bold", check_overlap = FALSE) +
  annotate(geom = "text", x = -95.06, y = 29.557, label = "Clear Lake", 
         fontface = "italic", color = "grey22", size = 4)
hhMapSegments
```

#### End R markdown

```
hhMapSegments
```

```
ggsave("Fig_1_hhMapSegments.png")
```

```
## Saving 7 x 5 in image
```
