## Supplemental File S02 for "Hurricane Harvey Impacts on Water Quality and Microbial Communities in Houston, TX Waterbodies": S02_TCEQ.html

TCEQ meta data


### TCEQ meta data

###### Michael G. LaMontagne

#### 12/19/2021

#### R Markdown

This is R Markdown document presents workflow for processing data downloaded from https://www80.tceq.texas.gov/SwqmisPublic/index.htm. Data from 2011 - 2021 for four segments (1101C, 1113B, 2425 & 2425B) were appended and imported.

```
library(stringr)
library(dplyr)
```

```
## 
## Attaching package: 'dplyr'
```

```
## The following objects are masked from 'package:stats':
## 
##     filter, lag
```

```
## The following objects are masked from 'package:base':
## 
##     intersect, setdiff, setequal, union
```

```
library(ggplot2)
library(ggbreak)
```

```
## ggbreak v0.0.7
## 
## If you use ggbreak in published research, please cite the following paper:
## 
## S Xu, M Chen, T Feng, L Zhan, L Zhou, G Yu. Use ggbreak to effectively utilize plotting space to deal with large datasets and outliers. Frontiers in Genetics. 2021, 12:774846. doi: 10.3389/fgene.2021.774846
```

```
library(agricolae)
sessionInfo()
```

```
## R version 4.1.1 (2021-08-10)
## Platform: x86_64-w64-mingw32/x64 (64-bit)
## Running under: Windows 10 x64 (build 22000)
## 
## Matrix products: default
## 
## locale:
## [1] LC_COLLATE=English_United States.1252 
## [2] LC_CTYPE=English_United States.1252   
## [3] LC_MONETARY=English_United States.1252
## [4] LC_NUMERIC=C                          
## [5] LC_TIME=English_United States.1252    
## 
## attached base packages:
## [1] stats     graphics  grDevices utils     datasets  methods   base     
## 
## other attached packages:
## [1] agricolae_1.3-5 ggbreak_0.0.7   ggplot2_3.3.5   dplyr_1.0.7    
## [5] stringr_1.4.0  
## 
## loaded via a namespace (and not attached):
##  [1] tidyselect_1.1.1   xfun_0.25          bslib_0.3.1        purrr_0.3.4       
##  [5] lattice_0.20-44    haven_2.4.3        labelled_2.8.0     ggfun_0.0.4       
##  [9] colorspace_2.0-2   vctrs_0.3.8        generics_0.1.1     miniUI_0.1.1.1    
## [13] htmltools_0.5.2    yaml_2.2.1         AlgDesign_1.2.0    utf8_1.2.2        
## [17] gridGraphics_0.5-1 rlang_0.4.11       later_1.3.0        jquerylib_0.1.4   
## [21] pillar_1.6.4       glue_1.4.2         withr_2.4.2        DBI_1.1.1         
## [25] lifecycle_1.0.1    questionr_0.7.5    munsell_0.5.0      combinat_0.0-8    
## [29] gtable_0.3.0       evaluate_0.14      forcats_0.5.1      knitr_1.36        
## [33] fastmap_1.1.0      httpuv_1.6.3       fansi_0.5.0        highr_0.9         
## [37] Rcpp_1.0.7         xtable_1.8-4       promises_1.2.0.1   scales_1.1.1      
## [41] jsonlite_1.7.2     mime_0.12          hms_1.1.1          klaR_0.6-15       
## [45] digest_0.6.27      aplot_0.1.1        stringi_1.7.5      shiny_1.7.1       
## [49] grid_4.1.1         tools_4.1.1        yulab.utils_0.0.4  magrittr_2.0.1    
## [53] sass_0.4.0         patchwork_1.1.1    tibble_3.1.4       cluster_2.1.2     
## [57] crayon_1.4.1       pkgconfig_2.0.3    ellipsis_0.3.2     MASS_7.3-54       
## [61] ggplotify_0.1.0    rstudioapi_0.13    assertthat_0.2.1   rmarkdown_2.11    
## [65] R6_2.5.1           nlme_3.1-152       compiler_4.1.1
```

```
# import data
tceqMeta <- as.data.frame(read.table("tceqMeta.csv", sep=",", header = TRUE, check.names = TRUE))
head(tceqMeta)
```

```
##   SegmentID StationID        StationDescription  EndDate EndTime EndDepth
## 1      1101     11446 CLEAR CREEK TIDAL AT SH 3 4/2/2013   10:53      0.3
## 2      1101     11446 CLEAR CREEK TIDAL AT SH 3 4/2/2013   10:53      0.3
## 3      1101     11446 CLEAR CREEK TIDAL AT SH 3 4/2/2013   10:53      0.3
## 4      1101     11446 CLEAR CREEK TIDAL AT SH 3 4/2/2013   10:53      0.3
## 5      1101     11446 CLEAR CREEK TIDAL AT SH 3 4/2/2013   10:53      0.3
## 6      1101     11446 CLEAR CREEK TIDAL AT SH 3 4/2/2013   10:53      0.3
##                            SubmittingEntity     CollectingEntity
## 1 TEXAS COMMISSION ON ENVIRONMENTAL QUALITY TCEQ REGIONAL OFFICE
## 2 TEXAS COMMISSION ON ENVIRONMENTAL QUALITY TCEQ REGIONAL OFFICE
## 3 TEXAS COMMISSION ON ENVIRONMENTAL QUALITY TCEQ REGIONAL OFFICE
## 4 TEXAS COMMISSION ON ENVIRONMENTAL QUALITY TCEQ REGIONAL OFFICE
## 5 TEXAS COMMISSION ON ENVIRONMENTAL QUALITY TCEQ REGIONAL OFFICE
## 6 TEXAS COMMISSION ON ENVIRONMENTAL QUALITY TCEQ REGIONAL OFFICE
##                                                             ParameterName
## 1                                                     PH (STANDARD UNITS)
## 2 EVIDENCE OF PRIMARY CONTACT RECREATION (1 = OBSERVED, 0 = NOT OBSERVED)
## 3                            DEPTH OF BOTTOM OF WATER BODY AT SAMPLE SITE
## 4                                   DAYS SINCE PRECIPITATION EVENT (DAYS)
## 5                                           SALINITY - PARTS PER THOUSAND
## 6                                                OXYGEN, DISSOLVED (MG/L)
##   ParameterCode Value Latitude Longitude month day year
## 1           400   7.7 29.52138 -95.10315     4   2 2013
## 2         89979   1.0 29.52138 -95.10315     4   2 2013
## 3         82903   3.3 29.52138 -95.10315     4   2 2013
## 4         72053   0.0 29.52138 -95.10315     4   2 2013
## 5           480   6.0 29.52138 -95.10315     4   2 2013
## 6           300   5.8 29.52138 -95.10315     4   2 2013
```

```
# remove station 20014, marina in Seabrook
tceqMeta <- tceqMeta[which(tceqMeta$StationID != 20014),]
# remove station 16576, which is west of I45
tceqMeta <- tceqMeta[which(tceqMeta$StationID != 16576),]
# remove station 16573, which is mouth of tributatry
tceqMeta <- tceqMeta[which(tceqMeta$StationID != 16573),]
# remove station 16475, which is different tributatry
#tceqMeta <- tceqMeta[which(tceqMeta$StationID != 16475),]
# remove station 16486, which is different tributatry
tceqMeta <- tceqMeta[which(tceqMeta$StationID != 16486),]
# remove station 16486, which is different tributatry
tceqMeta <- tceqMeta[which(tceqMeta$StationID != 16486),]
# remove station stations 21400 - 21409, which are different segments
tceqMeta <- tceqMeta[-which(tceqMeta$Latitude < 29.521),]
saveRDS(tceqMeta, "tceqMeta.rds")
```

#### Select temperature and take mean by Segment & month

```
# tceqMeta <- readRDS("tceqMeta.rds")
tceqTemp <- tceqMeta %>% filter(ParameterCode == 10)
monthTCEQ <- tceqTemp %>%
  group_by(SegmentID, month, year, ParameterName, ParameterCode) %>%
  summarize(meanValue = mean(Value, na.rm = TRUE), n = n())
```

```
## `summarise()` has grouped output by 'SegmentID', 'month', 'year', 'ParameterName'. You can override using the `.groups` argument.
```

```
monthTCEQ$SegMonth <- paste(monthTCEQ$SegmentID,"_",   monthTCEQ$month,"_",monthTCEQ$year)
head(monthTCEQ)
```

```
## # A tibble: 6 x 8
## # Groups:   SegmentID, month, year, ParameterName [6]
##   SegmentID month  year ParameterName     ParameterCode meanValue     n SegMonth
##   <chr>     <int> <int> <chr>                     <int>     <dbl> <int> <chr>   
## 1 1101          1  2014 TEMPERATURE, WAT~            10      7.74     5 1101 _ ~
## 2 1101          1  2020 TEMPERATURE, WAT~            10     15        1 1101 _ ~
## 3 1101          3  2015 TEMPERATURE, WAT~            10     20.4      5 1101 _ ~
## 4 1101          3  2016 TEMPERATURE, WAT~            10     22.5      5 1101 _ ~
## 5 1101          3  2017 TEMPERATURE, WAT~            10     20.8      5 1101 _ ~
## 6 1101          3  2018 TEMPERATURE, WAT~            10     21.0      6 1101 _ ~
```

#### Plot temperature as sanity check

```
names(monthTCEQ)[6] <- "Temperature"
# select site description
monthTCEQsum <- monthTCEQ[,c(1:3,6,8)]
head(monthTCEQsum)
```

```
## # A tibble: 6 x 5
## # Groups:   SegmentID, month, year [6]
##   SegmentID month  year Temperature SegMonth       
##   <chr>     <int> <int>       <dbl> <chr>          
## 1 1101          1  2014        7.74 1101 _ 1 _ 2014
## 2 1101          1  2020       15    1101 _ 1 _ 2020
## 3 1101          3  2015       20.4  1101 _ 3 _ 2015
## 4 1101          3  2016       22.5  1101 _ 3 _ 2016
## 5 1101          3  2017       20.8  1101 _ 3 _ 2017
## 6 1101          3  2018       21.0  1101 _ 3 _ 2018
```

```
plot(monthTCEQsum$month,monthTCEQsum$Temperature)
```

#### Mean by month and segment for salinity

```
tceqTemp <- tceqMeta %>% filter(ParameterCode == 480)
monthTCEQ <- tceqTemp %>%
  group_by(SegmentID, month, year, ParameterName, ParameterCode) %>%
  summarize(meanValue = mean(Value, na.rm = TRUE), n = n())
```

```
## `summarise()` has grouped output by 'SegmentID', 'month', 'year', 'ParameterName'. You can override using the `.groups` argument.
```

```
monthTCEQ$SegMonth <- paste(monthTCEQ$SegmentID,"_",   monthTCEQ$month,"_",monthTCEQ$year)
head(monthTCEQ)
```

```
## # A tibble: 6 x 8
## # Groups:   SegmentID, month, year, ParameterName [6]
##   SegmentID month  year ParameterName    ParameterCode meanValue     n SegMonth 
##   <chr>     <int> <int> <chr>                    <int>     <dbl> <int> <chr>    
## 1 1101          1  2014 SALINITY - PART~           480     5.42      5 1101 _ 1~
## 2 1101          1  2020 SALINITY - PART~           480     2.1       1 1101 _ 1~
## 3 1101          3  2015 SALINITY - PART~           480     0.1       5 1101 _ 3~
## 4 1101          3  2016 SALINITY - PART~           480     0.2       5 1101 _ 3~
## 5 1101          3  2017 SALINITY - PART~           480     0.2       5 1101 _ 3~
## 6 1101          3  2018 SALINITY - PART~           480     0.667     6 1101 _ 3~
```

#### Add salinity and test for significance

```
names(monthTCEQ)[6] <- "Salinity (PSU)"
monthTCEQsum2 <- monthTCEQ[,c(1:3,6,8)]
head(monthTCEQsum2)
```

```
## # A tibble: 6 x 5
## # Groups:   SegmentID, month, year [6]
##   SegmentID month  year `Salinity (PSU)` SegMonth       
##   <chr>     <int> <int>            <dbl> <chr>          
## 1 1101          1  2014            5.42  1101 _ 1 _ 2014
## 2 1101          1  2020            2.1   1101 _ 1 _ 2020
## 3 1101          3  2015            0.1   1101 _ 3 _ 2015
## 4 1101          3  2016            0.2   1101 _ 3 _ 2016
## 5 1101          3  2017            0.2   1101 _ 3 _ 2017
## 6 1101          3  2018            0.667 1101 _ 3 _ 2018
```

#### join temp & salinity

```
monthTCEQsum3 <- full_join(monthTCEQsum, monthTCEQsum2, by = "SegMonth")
names(monthTCEQsum3)[1] <- "SegmentID"
names(monthTCEQsum3)[2] <- "month"
names(monthTCEQsum3)[3] <- "year"
dim(monthTCEQsum3)
```

```
## [1] 174   9
```

```
head(monthTCEQsum3)
```

```
## # A tibble: 6 x 9
##   SegmentID month  year Temperature SegMonth        SegmentID.y month.y year.y
##   <chr>     <int> <int>       <dbl> <chr>           <chr>         <int>  <int>
## 1 1101          1  2014        7.74 1101 _ 1 _ 2014 1101              1   2014
## 2 1101          1  2020       15    1101 _ 1 _ 2020 1101              1   2020
## 3 1101          3  2015       20.4  1101 _ 3 _ 2015 1101              3   2015
## 4 1101          3  2016       22.5  1101 _ 3 _ 2016 1101              3   2016
## 5 1101          3  2017       20.8  1101 _ 3 _ 2017 1101              3   2017
## 6 1101          3  2018       21.0  1101 _ 3 _ 2018 1101              3   2018
## # ... with 1 more variable: Salinity (PSU) <dbl>
```

#### join salinity

```
monthTCEQsum4 <- monthTCEQsum3[,c(1:3,5,4,9)]
head(monthTCEQsum4)
```

```
## # A tibble: 6 x 6
##   SegmentID month  year SegMonth        Temperature `Salinity (PSU)`
##   <chr>     <int> <int> <chr>                 <dbl>            <dbl>
## 1 1101          1  2014 1101 _ 1 _ 2014        7.74            5.42 
## 2 1101          1  2020 1101 _ 1 _ 2020       15               2.1  
## 3 1101          3  2015 1101 _ 3 _ 2015       20.4             0.1  
## 4 1101          3  2016 1101 _ 3 _ 2016       22.5             0.2  
## 5 1101          3  2017 1101 _ 3 _ 2017       20.8             0.2  
## 6 1101          3  2018 1101 _ 3 _ 2018       21.0             0.667
```

#### calculate mean for Enterococci

```
tceqTemp <- tceqMeta %>% filter(ParameterCode == 31701)
monthTCEQ <- tceqTemp %>%
  group_by(SegmentID, month, year, ParameterName, ParameterCode) %>%
  summarize(meanValue = mean(Value, na.rm = TRUE), n = n())
```

```
## `summarise()` has grouped output by 'SegmentID', 'month', 'year', 'ParameterName'. You can override using the `.groups` argument.
```

```
monthTCEQ$SegMonth <- paste(monthTCEQ$SegmentID,"_",   monthTCEQ$month,"_",monthTCEQ$year)
head(monthTCEQ)
```

```
## # A tibble: 6 x 8
## # Groups:   SegmentID, month, year, ParameterName [6]
##   SegmentID month  year ParameterName     ParameterCode meanValue     n SegMonth
##   <chr>     <int> <int> <chr>                     <int>     <dbl> <int> <chr>   
## 1 1101          1  2014 ENTEROCOCCI, ENT~         31701        30     1 1101 _ ~
## 2 1101          1  2020 ENTEROCOCCI, ENT~         31701        31     1 1101 _ ~
## 3 1101          3  2015 ENTEROCOCCI, ENT~         31701       340     1 1101 _ ~
## 4 1101          3  2016 ENTEROCOCCI, ENT~         31701       110     1 1101 _ ~
## 5 1101          3  2017 ENTEROCOCCI, ENT~         31701       420     1 1101 _ ~
## 6 1101          3  2018 ENTEROCOCCI, ENT~         31701        10     1 1101 _ ~
```

#### join Enterococci

```
names(monthTCEQ)[6] <- "Enterococci (MPN/100ml)"
monthTCEQsum2 <- monthTCEQ[,c(1:3,6,8)]
head(monthTCEQsum2)
```

```
## # A tibble: 6 x 5
## # Groups:   SegmentID, month, year [6]
##   SegmentID month  year `Enterococci (MPN/100ml)` SegMonth       
##   <chr>     <int> <int>                     <dbl> <chr>          
## 1 1101          1  2014                        30 1101 _ 1 _ 2014
## 2 1101          1  2020                        31 1101 _ 1 _ 2020
## 3 1101          3  2015                       340 1101 _ 3 _ 2015
## 4 1101          3  2016                       110 1101 _ 3 _ 2016
## 5 1101          3  2017                       420 1101 _ 3 _ 2017
## 6 1101          3  2018                        10 1101 _ 3 _ 2018
```

#### join Enterococci

```
monthTCEQsum3 <- full_join(monthTCEQsum4, monthTCEQsum2, by = "SegMonth")
names(monthTCEQsum3)[1] <- "SegmentID"
names(monthTCEQsum3)[2] <- "month"
names(monthTCEQsum3)[3] <- "year"
dim(monthTCEQsum3)
```

```
## [1] 175  10
```

```
head(monthTCEQsum3)
```

```
## # A tibble: 6 x 10
##   SegmentID month  year SegMonth        Temperature `Salinity (PSU)` SegmentID.y
##   <chr>     <int> <int> <chr>                 <dbl>            <dbl> <chr>      
## 1 1101          1  2014 1101 _ 1 _ 2014        7.74            5.42  1101       
## 2 1101          1  2020 1101 _ 1 _ 2020       15               2.1   1101       
## 3 1101          3  2015 1101 _ 3 _ 2015       20.4             0.1   1101       
## 4 1101          3  2016 1101 _ 3 _ 2016       22.5             0.2   1101       
## 5 1101          3  2017 1101 _ 3 _ 2017       20.8             0.2   1101       
## 6 1101          3  2018 1101 _ 3 _ 2018       21.0             0.667 1101       
## # ... with 3 more variables: month.y <int>, year.y <int>,
## #   Enterococci (MPN/100ml) <dbl>
```

#### summary with Enterococci

```
monthTCEQsum5 <- monthTCEQsum3[,c(1:6,10)]
head(monthTCEQsum5)
```

```
## # A tibble: 6 x 7
##   SegmentID month  year SegMonth  Temperature `Salinity (PSU)` `Enterococci (MP~
##   <chr>     <int> <int> <chr>           <dbl>            <dbl>             <dbl>
## 1 1101          1  2014 1101 _ 1~        7.74            5.42                 30
## 2 1101          1  2020 1101 _ 1~       15               2.1                  31
## 3 1101          3  2015 1101 _ 3~       20.4             0.1                 340
## 4 1101          3  2016 1101 _ 3~       22.5             0.2                 110
## 5 1101          3  2017 1101 _ 3~       20.8             0.2                 420
## 6 1101          3  2018 1101 _ 3~       21.0             0.667                10
```

#### mean of nitrate/nitrite

```
tceqTemp <- tceqMeta %>% filter(ParameterCode == 630)
tceqTemp$Value <- 1000*(tceqTemp$Value/14)
tceqTemp$ParameterName <- "NOx (uM)"
monthTCEQ <- tceqTemp %>%
  group_by(SegmentID, month, year, ParameterName, ParameterCode) %>%
  summarize(meanValue = mean(Value, na.rm = TRUE), n = n())
```

```
## `summarise()` has grouped output by 'SegmentID', 'month', 'year', 'ParameterName'. You can override using the `.groups` argument.
```

```
monthTCEQ$SegMonth <- paste(monthTCEQ$SegmentID,"_",   monthTCEQ$month,"_",monthTCEQ$year)
head(monthTCEQ)
```

```
## # A tibble: 6 x 8
## # Groups:   SegmentID, month, year, ParameterName [6]
##   SegmentID month  year ParameterName ParameterCode meanValue     n SegMonth    
##   <chr>     <int> <int> <chr>                 <int>     <dbl> <int> <chr>       
## 1 1101          1  2014 NOx (uM)                630     370       1 1101 _ 1 _ ~
## 2 1101          1  2020 NOx (uM)                630     228.      1 1101 _ 1 _ ~
## 3 1101          3  2015 NOx (uM)                630      21.4     1 1101 _ 3 _ ~
## 4 1101          3  2016 NOx (uM)                630      73.6     1 1101 _ 3 _ ~
## 5 1101          3  2017 NOx (uM)                630      26.4     1 1101 _ 3 _ ~
## 6 1101          3  2018 NOx (uM)                630     218.      1 1101 _ 3 _ ~
```

#### join nitrate/nitrite

```
names(monthTCEQ)[6] <- "NOx (uM)"
monthTCEQsum2 <- monthTCEQ[,c(1:3,6,8)]
head(monthTCEQsum2)
```

```
## # A tibble: 6 x 5
## # Groups:   SegmentID, month, year [6]
##   SegmentID month  year `NOx (uM)` SegMonth       
##   <chr>     <int> <int>      <dbl> <chr>          
## 1 1101          1  2014      370   1101 _ 1 _ 2014
## 2 1101          1  2020      228.  1101 _ 1 _ 2020
## 3 1101          3  2015       21.4 1101 _ 3 _ 2015
## 4 1101          3  2016       73.6 1101 _ 3 _ 2016
## 5 1101          3  2017       26.4 1101 _ 3 _ 2017
## 6 1101          3  2018      218.  1101 _ 3 _ 2018
```

#### merge nitrate/nitrite

```
monthTCEQsum3 <- full_join(monthTCEQsum5, monthTCEQsum2, by = "SegMonth")
names(monthTCEQsum3)[1] <- "SegmentID"
names(monthTCEQsum3)[2] <- "month"
names(monthTCEQsum3)[3] <- "year"
dim(monthTCEQsum3)
```

```
## [1] 175  11
```

```
head(monthTCEQsum3)
```

```
## # A tibble: 6 x 11
##   SegmentID month  year SegMonth  Temperature `Salinity (PSU)` `Enterococci (MP~
##   <chr>     <int> <int> <chr>           <dbl>            <dbl>             <dbl>
## 1 1101          1  2014 1101 _ 1~        7.74            5.42                 30
## 2 1101          1  2020 1101 _ 1~       15               2.1                  31
## 3 1101          3  2015 1101 _ 3~       20.4             0.1                 340
## 4 1101          3  2016 1101 _ 3~       22.5             0.2                 110
## 5 1101          3  2017 1101 _ 3~       20.8             0.2                 420
## 6 1101          3  2018 1101 _ 3~       21.0             0.667                10
## # ... with 4 more variables: SegmentID.y <chr>, month.y <int>, year.y <int>,
## #   NOx (uM) <dbl>
```

#### Summary with nitrate

```
monthTCEQsum6 <- monthTCEQsum3[,c(1:7,11)]
head(monthTCEQsum6)
```

```
## # A tibble: 6 x 8
##   SegmentID month  year SegMonth  Temperature `Salinity (PSU)` `Enterococci (MP~
##   <chr>     <int> <int> <chr>           <dbl>            <dbl>             <dbl>
## 1 1101          1  2014 1101 _ 1~        7.74            5.42                 30
## 2 1101          1  2020 1101 _ 1~       15               2.1                  31
## 3 1101          3  2015 1101 _ 3~       20.4             0.1                 340
## 4 1101          3  2016 1101 _ 3~       22.5             0.2                 110
## 5 1101          3  2017 1101 _ 3~       20.8             0.2                 420
## 6 1101          3  2018 1101 _ 3~       21.0             0.667                10
## # ... with 1 more variable: NOx (uM) <dbl>
```

#### add ammonium

```
tceqTemp <- tceqMeta %>% filter(ParameterCode == 610)
tceqTemp$Value <- 1000*(tceqTemp$Value/14)
tceqTemp$ParameterName <- "NH4 (uM)"
monthTCEQ <- tceqTemp %>%
  group_by(SegmentID, month, year, ParameterName, ParameterCode) %>%
  summarize(meanValue = mean(Value, na.rm = TRUE), n = n())
```

```
## `summarise()` has grouped output by 'SegmentID', 'month', 'year', 'ParameterName'. You can override using the `.groups` argument.
```

```
monthTCEQ$SegMonth <- paste(monthTCEQ$SegmentID,"_",   monthTCEQ$month,"_",monthTCEQ$year)
names(monthTCEQ)[6] <- "NH4 (uM)"
monthTCEQsum2 <- monthTCEQ[,c(1:3,6,8)]
#head(monthTCEQsum2)
monthTCEQsum3 <- full_join(monthTCEQsum6, monthTCEQsum2, by = "SegMonth")
names(monthTCEQsum3)[1] <- "SegmentID"
names(monthTCEQsum3)[2] <- "month"
names(monthTCEQsum3)[3] <- "year"
#dim(monthTCEQsum3)
#head(monthTCEQsum3)
monthTCEQsum7 <- monthTCEQsum3[,c(1:8,12)]
head(monthTCEQsum7)
```

```
## # A tibble: 6 x 9
##   SegmentID month  year SegMonth  Temperature `Salinity (PSU)` `Enterococci (MP~
##   <chr>     <int> <int> <chr>           <dbl>            <dbl>             <dbl>
## 1 1101          1  2014 1101 _ 1~        7.74            5.42                 30
## 2 1101          1  2020 1101 _ 1~       15               2.1                  31
## 3 1101          3  2015 1101 _ 3~       20.4             0.1                 340
## 4 1101          3  2016 1101 _ 3~       22.5             0.2                 110
## 5 1101          3  2017 1101 _ 3~       20.8             0.2                 420
## 6 1101          3  2018 1101 _ 3~       21.0             0.667                10
## # ... with 2 more variables: NOx (uM) <dbl>, NH4 (uM) <dbl>
```

#### add phosphate

```
tceqTemp <- tceqMeta %>% filter(ParameterCode == 665)
tceqTemp$Value <- 1000*(tceqTemp$Value/31)
tceqTemp$ParameterName <- "PO4 (uM)"
monthTCEQ <- tceqTemp %>%
  group_by(SegmentID, month, year, ParameterName, ParameterCode) %>%
  summarize(meanValue = mean(Value, na.rm = TRUE), n = n())
```

```
## `summarise()` has grouped output by 'SegmentID', 'month', 'year', 'ParameterName'. You can override using the `.groups` argument.
```

```
monthTCEQ$SegMonth <- paste(monthTCEQ$SegmentID,"_",   monthTCEQ$month,"_",monthTCEQ$year)
names(monthTCEQ)[6] <- "PO4 (uM)"
monthTCEQsum2 <- monthTCEQ[,c(1:3,6,8)]
#add phosphate 
monthTCEQsum3 <- full_join(monthTCEQsum7, monthTCEQsum2, by = "SegMonth")
names(monthTCEQsum3)[1] <- "SegmentID"
names(monthTCEQsum3)[2] <- "month"
names(monthTCEQsum3)[3] <- "year"
#head(monthTCEQsum3)
monthTCEQsum8 <- monthTCEQsum3[,c(1:9,13)]
head(monthTCEQsum8)
```

```
## # A tibble: 6 x 10
##   SegmentID month  year SegMonth  Temperature `Salinity (PSU)` `Enterococci (MP~
##   <chr>     <int> <int> <chr>           <dbl>            <dbl>             <dbl>
## 1 1101          1  2014 1101 _ 1~        7.74            5.42                 30
## 2 1101          1  2020 1101 _ 1~       15               2.1                  31
## 3 1101          3  2015 1101 _ 3~       20.4             0.1                 340
## 4 1101          3  2016 1101 _ 3~       22.5             0.2                 110
## 5 1101          3  2017 1101 _ 3~       20.8             0.2                 420
## 6 1101          3  2018 1101 _ 3~       21.0             0.667                10
## # ... with 3 more variables: NOx (uM) <dbl>, NH4 (uM) <dbl>, PO4 (uM) <dbl>
```

#### add pH

```
tceqTemp <- tceqMeta %>% filter(ParameterCode == 400)
tceqTemp$ParameterName <- "pH"
monthTCEQ <- tceqTemp %>%
  group_by(SegmentID, month, year, ParameterName, ParameterCode) %>%
  summarize(meanValue = mean(Value, na.rm = TRUE), n = n())
```

```
## `summarise()` has grouped output by 'SegmentID', 'month', 'year', 'ParameterName'. You can override using the `.groups` argument.
```

```
monthTCEQ$SegMonth <- paste(monthTCEQ$SegmentID,"_",   monthTCEQ$month,"_",monthTCEQ$year)
names(monthTCEQ)[6] <- "pH"
monthTCEQsum2 <- monthTCEQ[,c(1:3,6,8)]
#join to previous sum 
monthTCEQsum3 <- full_join(monthTCEQsum8, monthTCEQsum2, by = "SegMonth")
names(monthTCEQsum3)[1] <- "SegmentID"
names(monthTCEQsum3)[2] <- "month"
names(monthTCEQsum3)[3] <- "year"
#head(monthTCEQsum3)
monthTCEQsum9 <- monthTCEQsum3[,c(1:10,14)]
head(monthTCEQsum9)
```

```
## # A tibble: 6 x 11
##   SegmentID month  year SegMonth  Temperature `Salinity (PSU)` `Enterococci (MP~
##   <chr>     <int> <int> <chr>           <dbl>            <dbl>             <dbl>
## 1 1101          1  2014 1101 _ 1~        7.74            5.42                 30
## 2 1101          1  2020 1101 _ 1~       15               2.1                  31
## 3 1101          3  2015 1101 _ 3~       20.4             0.1                 340
## 4 1101          3  2016 1101 _ 3~       22.5             0.2                 110
## 5 1101          3  2017 1101 _ 3~       20.8             0.2                 420
## 6 1101          3  2018 1101 _ 3~       21.0             0.667                10
## # ... with 4 more variables: NOx (uM) <dbl>, NH4 (uM) <dbl>, PO4 (uM) <dbl>,
## #   pH <dbl>
```

#### add conductivity

```
tceqTemp <- tceqMeta %>% filter(ParameterCode == 94)
tceqTemp$ParameterName <- "cond"
monthTCEQ <- tceqTemp %>%
  group_by(SegmentID, month, year, ParameterName, ParameterCode) %>%
  summarize(meanValue = mean(Value, na.rm = TRUE), n = n())
```

```
## `summarise()` has grouped output by 'SegmentID', 'month', 'year', 'ParameterName'. You can override using the `.groups` argument.
```

```
monthTCEQ$SegMonth <- paste(monthTCEQ$SegmentID,"_",   monthTCEQ$month,"_",monthTCEQ$year)
names(monthTCEQ)[6] <- "Cond"
monthTCEQsum2 <- monthTCEQ[,c(1:3,6,8)]
#join to previous sum 
monthTCEQsum3 <- full_join(monthTCEQsum9, monthTCEQsum2, by = "SegMonth")
names(monthTCEQsum3)[1] <- "SegmentID"
names(monthTCEQsum3)[2] <- "month"
names(monthTCEQsum3)[3] <- "year"
#head(monthTCEQsum3)
monthTCEQsum10 <- monthTCEQsum3[,c(1:11,15)]
head(monthTCEQsum10)
```

```
## # A tibble: 6 x 12
##   SegmentID month  year SegMonth  Temperature `Salinity (PSU)` `Enterococci (MP~
##   <chr>     <int> <int> <chr>           <dbl>            <dbl>             <dbl>
## 1 1101          1  2014 1101 _ 1~        7.74            5.42                 30
## 2 1101          1  2020 1101 _ 1~       15               2.1                  31
## 3 1101          3  2015 1101 _ 3~       20.4             0.1                 340
## 4 1101          3  2016 1101 _ 3~       22.5             0.2                 110
## 5 1101          3  2017 1101 _ 3~       20.8             0.2                 420
## 6 1101          3  2018 1101 _ 3~       21.0             0.667                10
## # ... with 5 more variables: NOx (uM) <dbl>, NH4 (uM) <dbl>, PO4 (uM) <dbl>,
## #   pH <dbl>, Cond <dbl>
```

#### add oxygen

```
tceqTemp <- tceqMeta %>% filter(ParameterCode == 300)
tceqTemp$ParameterName <- "oxygen"
monthTCEQ <- tceqTemp %>%
  group_by(SegmentID, month, year, ParameterName, ParameterCode) %>%
  summarize(meanValue = mean(Value, na.rm = TRUE), n = n())
```

```
## `summarise()` has grouped output by 'SegmentID', 'month', 'year', 'ParameterName'. You can override using the `.groups` argument.
```

```
monthTCEQ$SegMonth <- paste(monthTCEQ$SegmentID,"_",   monthTCEQ$month,"_",monthTCEQ$year)
names(monthTCEQ)[6] <- "Oxygen"
monthTCEQsum2 <- monthTCEQ[,c(1:3,6,8)]
#join to previous sum 
monthTCEQsum3 <- full_join(monthTCEQsum10, monthTCEQsum2, by = "SegMonth")
names(monthTCEQsum3)[1] <- "SegmentID"
names(monthTCEQsum3)[2] <- "month"
names(monthTCEQsum3)[3] <- "year"
#head(monthTCEQsum3)
monthTCEQsum11 <- monthTCEQsum3[,c(1:12,16)]
head(monthTCEQsum11)
```

```
## # A tibble: 6 x 13
##   SegmentID month  year SegMonth  Temperature `Salinity (PSU)` `Enterococci (MP~
##   <chr>     <int> <int> <chr>           <dbl>            <dbl>             <dbl>
## 1 1101          1  2014 1101 _ 1~        7.74            5.42                 30
## 2 1101          1  2020 1101 _ 1~       15               2.1                  31
## 3 1101          3  2015 1101 _ 3~       20.4             0.1                 340
## 4 1101          3  2016 1101 _ 3~       22.5             0.2                 110
## 5 1101          3  2017 1101 _ 3~       20.8             0.2                 420
## 6 1101          3  2018 1101 _ 3~       21.0             0.667                10
## # ... with 6 more variables: NOx (uM) <dbl>, NH4 (uM) <dbl>, PO4 (uM) <dbl>,
## #   pH <dbl>, Cond <dbl>, Oxygen <dbl>
```

#### plot nutrients versus salinity

```
monthTCEQsum <- monthTCEQsum11
n <- c(6,8:12)
pairs(monthTCEQsum[,n], pch = 19)
```

#### export data

```
monthTCEQsum$type <- "TCEQ"
monthTCEQsum$season <- "NA"
monthTCEQsum$factorDate <- "15-Month"
n <- c(1,14,2,15,8:12,6,13,5,7,16)
monthTCEQsum <- monthTCEQsum[,n]
# saveRDS(monthTCEQsum, "monthTCEQsum.rds")
# based on variable values
monthTCEQsum2 <- monthTCEQsum[ which(monthTCEQsum$month > 7
& monthTCEQsum$month < 11), ]
write.csv(monthTCEQsum2, "monthTCEQ.csv")
```

#### test salinity for significance

```
tceqTemp <- tceqMeta %>% filter(ParameterCode == 480)
# ANOVA
fit <- aov(Value ~ SegmentID*month, data=tceqTemp)
summary(fit)
```

```
##                  Df Sum Sq Mean Sq F value Pr(>F)    
## SegmentID         4   9602  2400.4 133.389 <2e-16 ***
## month             1     62    61.8   3.432 0.0643 .  
## SegmentID:month   4    158    39.6   2.200 0.0674 .  
## Residuals       772  13893    18.0                   
## ---
## Signif. codes:  0 '***' 0.001 '**' 0.01 '*' 0.05 '.' 0.1 ' ' 1
```

#### Plot salinity by station

```
# compare means
tceqTemp$SegmentID <- as.factor(tceqTemp$SegmentID)
ggplot(tceqTemp, aes(x = SegmentID, y = Value, color = SegmentID)) + 
  geom_violin(fill = NA) +
  geom_jitter(position = position_jitter(0.2)) +
  xlab("Station") + ylab(tceqTemp[1,9]) + theme_bw() +
 theme(panel.grid.major = element_blank(), panel.grid.minor = element_blank()) +
  theme(legend.position = "none")
```

#### Name change

```
ggsave("Fig_S2_SalinityTCEQ.png")
```

```
## Saving 7 x 5 in image
```

```
# select site description
```

#### calculate geometric mean for Enterococci

```
tceqTemp <- tceqMeta %>% filter(ParameterCode == 31701)
# hist
tceqTemp$logV <- log10(tceqTemp$Value)
hist(tceqTemp$logV)
```

#### Plot by Segment

```
tceqTemp$Segment <- as.factor(tceqTemp$SegmentID)
EC <- ggplot(tceqTemp, aes(x = Segment, y = logV, color = SegmentID)) +
  geom_violin(fill = NA) +
  geom_jitter(position = position_jitter(0.2)) +
  xlab("Segment") + ylab("log(Enterococci) MPN/100") + theme_bw() +
 theme(panel.grid.major = element_blank(), panel.grid.minor = element_blank()) +
  theme(legend.position = "none")
EC
```

#### Plot seasonal pattern

```
ggsave("Fig_5_Enterococci_HH.png")
```

```
## Saving 7 x 5 in image
```

```
fit <- aov(logV ~ SegmentID, data=tceqTemp)
summary(fit)
```

```
##              Df Sum Sq Mean Sq F value   Pr(>F)    
## SegmentID     4  33.71   8.427   16.59 5.12e-12 ***
## Residuals   240 121.91   0.508                     
## ---
## Signif. codes:  0 '***' 0.001 '**' 0.01 '*' 0.05 '.' 0.1 ' ' 1
```

#### ANOVA of Enterococci

```
tukeyTest <- HSD.test(fit, trt = 'SegmentID')
tukeyTest
```

```
## $statistics
##     MSerror  Df     Mean       CV
##   0.5079751 240 1.702146 41.87206
## 
## $parameters
##    test    name.t ntr StudentizedRange alpha
##   Tukey SegmentID   5         3.887148  0.05
## 
## $means
##           logV       std  r Min      Max      Q25      Q50      Q75
## 1101  1.924662 0.8057510 25   1 4.301030 1.477121 1.763428 2.531479
## 1101C 2.307196 0.7806265 30   1 4.380211 1.799341 2.204120 2.705420
## 1113B 1.782413 0.7310181 53   1 4.204120 1.301030 1.716003 2.113943
## 2425  1.239513 0.4274708 86   1 2.869232 1.000000 1.000000 1.301030
## 2425B 1.933871 0.9522618 51   1 4.913814 1.301030 1.612784 2.332321
## 
## $comparison
## NULL
## 
## $groups
##           logV groups
## 1101C 2.307196      a
## 2425B 1.933871     ab
## 1101  1.924662     ab
## 1113B 1.782413      b
## 2425  1.239513      c
## 
## attr(,"class")
## [1] "group"
```

#### Plot seasonal pattern

```
plot(tceqTemp$month,tceqTemp$logV)
```

#### ANOVA nitrate

```
tceqTemp <- tceqMeta %>% filter(ParameterCode == 630)
# change units to umol
tceqNOx <- tceqTemp
tceqNOx$Value <- 1000*(tceqTemp$Value/14)
tceqNOx$ParameterName <- "NOx (uM)"
# ANOVA
fit <- aov(Value ~ SegmentID*month, data=tceqNOx)
summary(fit)
```

```
##                  Df  Sum Sq Mean Sq F value Pr(>F)    
## SegmentID         4 8125133 2031283  62.973 <2e-16 ***
## month             1    1121    1121   0.035  0.852    
## SegmentID:month   4   91059   22765   0.706  0.589    
## Residuals       243 7838351   32257                   
## ---
## Signif. codes:  0 '***' 0.001 '**' 0.01 '*' 0.05 '.' 0.1 ' ' 1
```

```
##
```

#### run HSD

```
tukeyTest <- HSD.test(fit, trt = 'SegmentID')
tukeyTest
```

```
## $statistics
##    MSerror  Df    Mean      CV
##   32256.59 243 129.203 139.007
## 
## $parameters
##    test    name.t ntr StudentizedRange alpha
##   Tukey SegmentID   5         3.886782  0.05
## 
## $means
##           Value       std  r       Min        Max        Q25        Q50
## 1101  126.17143  82.12833 25 21.428571  370.00000  77.142857 101.428571
## 1101C  21.52381  28.73092 30  1.428571  132.85714   6.607143   9.642857
## 1113B 452.70936 367.28657 58 51.428571 1764.28571 234.642857 333.928571
## 2425   19.96697  20.69729 93  1.428571   97.14286   2.857143  10.714286
## 2425B  16.47416  18.66876 47  1.428571   70.00000   2.857143   7.857143
##             Q75
## 1101  192.85714
## 1101C  27.50000
## 1113B 600.35714
## 2425   34.28571
## 2425B  23.57143
## 
## $comparison
## NULL
## 
## $groups
##           Value groups
## 1113B 452.70936      a
## 1101  126.17143      b
## 1101C  21.52381      b
## 2425   19.96697      b
## 2425B  16.47416      b
## 
## attr(,"class")
## [1] "group"
```

#### Plot nitrate by segment to generate Fig S3

```
ggplot(tceqNOx, aes(x = SegmentID, y = Value, color = SegmentID)) + 
  geom_violin(fill = NA) +
  geom_jitter(position = position_jitter(0.2)) +
  xlab("Segment") + ylab("NOx \u00B5M") + theme_bw() +
 theme(panel.grid.major = element_blank(), panel.grid.minor = element_blank()) +
  theme(legend.position = "none")
```

#### Save nitrate

```
ggsave("Fig_S3_NOx.png")
```

```
## Saving 7 x 5 in image
```

#### ANOVA Ammonium

```
tceqTemp <- tceqMeta %>% filter(ParameterCode == 610)
tceqNH4 <- tceqTemp
tceqNH4$Value <- 1000*(tceqTemp$Value/14)
tceqNH4$ParameterName <- "NH4 (uM)"
# ANOVA
fit <- aov(Value ~ SegmentID*month, data=tceqNH4)
summary(fit)
```

```
##                  Df Sum Sq Mean Sq F value   Pr(>F)    
## SegmentID         4  21079    5270   7.597 8.84e-06 ***
## month             1     66      66   0.095    0.759    
## SegmentID:month   4    221      55   0.080    0.989    
## Residuals       241 167167     694                     
## ---
## Signif. codes:  0 '***' 0.001 '**' 0.01 '*' 0.05 '.' 0.1 ' ' 1
```

```
##
```

#### Tukey test ammonium by seqment

```
tukeyTest <- HSD.test(fit, trt = 'SegmentID')
tukeyTest
```

```
## $statistics
##    MSerror  Df    Mean       CV
##   693.6374 241 14.3669 183.3171
## 
## $parameters
##    test    name.t ntr StudentizedRange alpha
##   Tukey SegmentID   5         3.887025  0.05
## 
## $means
##           Value       std  r      Min       Max      Q25       Q50       Q75
## 1101  13.238000  7.150559 25 2.664286  30.00000 8.571429 12.142857 16.428571
## 1101C 15.238095 17.469520 30 7.142857  92.85714 7.142857  7.142857 14.285714
## 1113B 30.602041 52.864482 56 1.428571 279.28571 3.571429 11.071429 23.392857
## 2425   6.629213  3.525317 89 1.428571  19.28571 3.571429  7.142857  7.142857
## 2425B 10.084034  7.177981 51 1.428571  50.00000 7.142857  7.142857 14.285714
## 
## $comparison
## NULL
## 
## $groups
##           Value groups
## 1113B 30.602041      a
## 1101C 15.238095     ab
## 1101  13.238000     ab
## 2425B 10.084034      b
## 2425   6.629213      b
## 
## attr(,"class")
## [1] "group"
```

#### Plot Ammonium

```
ggplot(tceqNH4, aes(x = SegmentID, y = Value, color = SegmentID)) + 
  geom_violin(fill = NA) +
  geom_jitter(position = position_jitter(0.2)) +
 # geom_boxplot(fill = NA) +
  xlab("Segment") + ylab("Ammonium (\u00B5M)") + theme_bw() +
 theme(panel.grid.major = element_blank(), panel.grid.minor = element_blank()) +
  theme(legend.position = "none")
```

#### save ammonium figure

```
ggsave("Fig_S4_NH4.png")
```

```
## Saving 7 x 5 in image
```

#### ANOVA phosphorous

```
tceqTemp <- tceqMeta %>% filter(ParameterCode == 665)
tceqP <- tceqTemp
tceqP$Value <- 1000*(tceqTemp$Value/31)
tceqP$ParameterName <- "Phosphate (uM)"
# ANOVA
fit <- aov(Value ~ SegmentID*month, data=tceqP)
summary(fit)
```

```
##                  Df Sum Sq Mean Sq F value Pr(>F)  
## SegmentID         4  30821    7705   2.698 0.0314 *
## month             1    329     329   0.115 0.7346  
## SegmentID:month   4   2700     675   0.236 0.9176  
## Residuals       239 682474    2856                 
## ---
## Signif. codes:  0 '***' 0.001 '**' 0.01 '*' 0.05 '.' 0.1 ' ' 1
```

```
##
```

#### Tukey test phosphate by seqment

```
tukeyTest <- HSD.test(fit, trt = 'SegmentID')
tukeyTest
```

```
## $statistics
##    MSerror  Df     Mean       CV
##   2855.539 239 24.17101 221.0799
## 
## $parameters
##    test    name.t ntr StudentizedRange alpha
##   Tukey SegmentID   5         3.887272  0.05
## 
## $means
##          Value       std  r       Min       Max       Q25       Q50      Q75
## 1101  21.83731  9.637172 23 8.3870968  43.87097 14.258065 20.612903 27.12903
## 1101C 18.88172 36.512125 30 2.5806452 204.83871  6.209677  9.193548 18.70968
## 1113B 45.18817 31.663130 54 3.3548387 109.35484 17.177419 36.774194 71.04839
## 2425  17.48352 78.919763 91 0.6451613 761.29032  6.774194  8.709677 11.45161
## 2425B 18.01392 24.945257 51 2.2580645 150.00000  7.580645 10.322581 16.45161
## 
## $comparison
## NULL
## 
## $groups
##          Value groups
## 1113B 45.18817      a
## 1101  21.83731     ab
## 1101C 18.88172     ab
## 2425B 18.01392     ab
## 2425  17.48352      b
## 
## attr(,"class")
## [1] "group"
```

#### Plot phosphorous

```
ggplot(tceqTemp, aes(x = SegmentID, y = Value, color = SegmentID)) + 
  geom_violin(fill = NA) +
  geom_jitter(position = position_jitter(0.2)) +
#  geom_boxplot(fill = NA) +
  xlab("Segment") + ylab("Phosphate (\u00B5M)") + theme_bw() +
 theme(panel.grid.major = element_blank(), panel.grid.minor = element_blank()) +
  theme(legend.position = "none")
```

### save phosphate figure

```
ggsave("Fig_S5_PO4.png")
```

```
## Saving 7 x 5 in image
```

#### ANOVA pH

```
tceqTemp <- tceqMeta %>% filter(ParameterCode == 400)
# ANOVA
fit <- aov(Value ~ SegmentID*month, data=tceqTemp)
summary(fit)
```

```
##                  Df Sum Sq Mean Sq F value  Pr(>F)    
## SegmentID         4  39.19   9.797  61.370 < 2e-16 ***
## month             1   0.53   0.526   3.295 0.06989 .  
## SegmentID:month   4   2.78   0.696   4.361 0.00171 ** 
## Residuals       761 121.48   0.160                    
## ---
## Signif. codes:  0 '***' 0.001 '**' 0.01 '*' 0.05 '.' 0.1 ' ' 1
```

```
##
```

#### Tukey test pH by seqment

```
tukeyTest <- HSD.test(fit, trt = 'SegmentID')
tukeyTest
```

```
## $statistics
##     MSerror  Df     Mean       CV
##   0.1596345 761 7.988586 5.001422
## 
## $parameters
##    test    name.t ntr StudentizedRange alpha
##   Tukey SegmentID   5         3.866925  0.05
## 
## $means
##          Value       std   r Min Max Q25 Q50 Q75
## 1101  7.690769 0.3667258 130 7.1 8.6 7.5 7.6 7.7
## 1101C 7.944944 0.4535244  89 7.2 8.8 7.6 7.9 8.3
## 1113B 7.804569 0.3965790 197 7.0 9.0 7.5 7.7 8.1
## 2425  8.263139 0.4231302 274 7.2 9.3 8.0 8.3 8.5
## 2425B 8.033333 0.3471311  81 7.3 8.7 7.8 8.1 8.3
## 
## $comparison
## NULL
## 
## $groups
##          Value groups
## 2425  8.263139      a
## 2425B 8.033333      b
## 1101C 7.944944      b
## 1113B 7.804569      c
## 1101  7.690769      c
## 
## attr(,"class")
## [1] "group"
```

#### Plot pH

```
ggplot(tceqTemp, aes(x = SegmentID, y = Value, color = SegmentID)) + 
  geom_violin(fill = NA) +
  geom_jitter(position = position_jitter(0.2)) +
#  geom_boxplot(fill = NA) +
  xlab("Segment") + ylab(tceqTemp[1,9]) + theme_bw() +
 theme(panel.grid.major = element_blank(), panel.grid.minor = element_blank()) +
  theme(legend.position = "none")
```

### save pH figure

```
ggsave("Fig_S6_pH.png")
```

```
## Saving 7 x 5 in image
```

#### ANOVA conductivity

```
tceqTemp <- tceqMeta %>% filter(ParameterCode == 94)
## ANOVA
fit <- aov(Value ~ SegmentID*month, data=tceqTemp)
summary(fit)
```

```
##                  Df    Sum Sq   Mean Sq F value Pr(>F)    
## SegmentID         4 2.757e+10 6.893e+09 137.227 <2e-16 ***
## month             1 1.634e+08 1.634e+08   3.253 0.0717 .  
## SegmentID:month   4 4.800e+08 1.200e+08   2.389 0.0496 *  
## Residuals       772 3.878e+10 5.023e+07                   
## ---
## Signif. codes:  0 '***' 0.001 '**' 0.01 '*' 0.05 '.' 0.1 ' ' 1
```

#### Tukey test conductivity by seqment

```
tukeyTest <- HSD.test(fit, trt = 'SegmentID')
tukeyTest
```

```
## $statistics
##    MSerror  Df     Mean     CV
##   50234236 772 9424.514 75.204
## 
## $parameters
##    test    name.t ntr StudentizedRange alpha
##   Tukey SegmentID   5         3.866793  0.05
## 
## $means
##           Value      std   r Min   Max    Q25   Q50     Q75
## 1101   3206.600 4880.507 130 225 21000  398.5   445  4325.0
## 1101C  4119.034 5852.273  89 141 25200  442.0  1130  5190.0
## 1113B  4499.711 5160.478 204 258 22900  669.0  1950  7342.5
## 2425  16457.317 9102.347 278 214 37000 7857.5 17900 23800.0
## 2425B 13499.296 7890.998  81 421 34000 7760.0 12000 19100.0
## 
## $comparison
## NULL
## 
## $groups
##           Value groups
## 2425  16457.317      a
## 2425B 13499.296      b
## 1113B  4499.711      c
## 1101C  4119.034      c
## 1101   3206.600      c
## 
## attr(,"class")
## [1] "group"
```

#### Plot conductivity

```
ggplot(tceqTemp, aes(x = SegmentID, y = Value, color = SegmentID)) + 
  geom_violin(fill = NA) +
  geom_jitter(position = position_jitter(0.2)) +
#  geom_boxplot(fill = NA) +
  xlab("Segment") + ylab(tceqTemp[1,9]) + theme_bw() +
 theme(panel.grid.major = element_blank(), panel.grid.minor = element_blank()) +
  theme(legend.position = "none")
```

### save conductivity figure

```
ggsave("Fig_S6_Cond.png")
```

```
## Saving 7 x 5 in image
```

#### add Oxygen

```
tceqTemp <- tceqMeta %>% filter(ParameterCode == 300)
# ANOVA
fit <- aov(Value ~ SegmentID*month, data=tceqTemp)
summary(fit)
```

```
##                  Df Sum Sq Mean Sq F value   Pr(>F)    
## SegmentID         4   1285   321.3   45.75  < 2e-16 ***
## month             1    347   346.7   49.37 4.72e-12 ***
## SegmentID:month   4    344    85.9   12.23 1.21e-09 ***
## Residuals       761   5345     7.0                     
## ---
## Signif. codes:  0 '***' 0.001 '**' 0.01 '*' 0.05 '.' 0.1 ' ' 1
```

#### Tukey test oxygen by seqment

```
tukeyTest <- HSD.test(fit, trt = 'SegmentID')
tukeyTest
```

```
## $statistics
##    MSerror  Df     Mean       CV
##   7.023898 761 6.621271 40.02651
## 
## $parameters
##    test    name.t ntr StudentizedRange alpha
##   Tukey SegmentID   5         3.866925  0.05
## 
## $means
##          Value      std   r Min  Max  Q25  Q50    Q75
## 1101  5.624615 1.751857 130 2.9 11.8 4.60 5.15  6.775
## 1101C 6.031325 3.298445  83 0.2 15.4 3.60 5.70  7.700
## 1113B 5.225980 2.809390 204 0.0 16.2 3.30 5.35  6.825
## 2425  8.239209 3.126920 278 0.8 18.5 6.10 7.35 10.000
## 2425B 6.797368 2.413682  76 2.1 11.8 4.95 6.80  8.325
## 
## $comparison
## NULL
## 
## $groups
##          Value groups
## 2425  8.239209      a
## 2425B 6.797368      b
## 1101C 6.031325     bc
## 1101  5.624615      c
## 1113B 5.225980      c
## 
## attr(,"class")
## [1] "group"
```

#### Tukey test oxygen by month

```
tukeyTest <- HSD.test(fit, trt = 'month')
tukeyTest
```

```
## $statistics
##    MSerror  Df     Mean       CV
##   7.023898 761 6.621271 40.02651
## 
## $parameters
##    test name.t ntr StudentizedRange alpha
##   Tukey  month  12         4.636192  0.05
## 
## $means
##        Value      std   r Min  Max   Q25  Q50    Q75
## 1  10.142718 3.065066 103 5.7 18.5 7.700 9.80 11.800
## 2   7.347500 3.123032  40 2.3 16.0 5.850 6.65  8.000
## 3   5.853571 2.355880  56 0.3 14.1 4.800 5.60  6.950
## 4   7.421951 3.481918  41 0.6 15.4 4.400 6.90 10.500
## 5   6.097000 2.794998 100 0.1 16.2 4.775 6.25  7.200
## 6   4.893750 1.709225  48 1.7 10.4 3.600 4.45  5.775
## 7   5.813084 2.720574 107 0.4 15.7 3.950 5.20  6.600
## 8   4.468966 3.075324  58 0.0 10.6 1.500 4.65  6.775
## 9   5.573171 1.144776  41 3.8  9.2 4.900 5.60  6.000
## 10  6.385075 2.507278  67 0.4 13.6 4.600 6.20  7.650
## 11  6.373529 2.507875  68 0.5 11.5 4.975 6.55  7.800
## 12  7.590476 1.634955  42 4.0 12.4 6.800 7.40  7.975
## 
## $comparison
## NULL
## 
## $groups
##        Value groups
## 1  10.142718      a
## 12  7.590476      b
## 4   7.421951      b
## 2   7.347500     bc
## 10  6.385075    bcd
## 11  6.373529    bcd
## 5   6.097000    bcd
## 3   5.853571   bcde
## 7   5.813084    cde
## 9   5.573171    cde
## 6   4.893750     de
## 8   4.468966      e
## 
## attr(,"class")
## [1] "group"
```

#### Plot Oxygen

```
ggplot(tceqTemp, aes(x = SegmentID, y = Value, color = SegmentID)) + 
  geom_violin(fill = NA) +
  geom_jitter(position = position_jitter(0.2)) +
#  geom_boxplot(fill = NA) +
  xlab("Segment") + ylab("Oxygen (mg/L)") + theme_bw() +
 theme(panel.grid.major = element_blank(), panel.grid.minor = element_blank()) +
  theme(legend.position = "none")
```

### save oxygen figure

```
ggsave("Fig_S7_oxygen.png")
```

```
## Saving 7 x 5 in image
```

#### Plot oxygen by month

```
tceqTemp$month <- as.factor(tceqTemp$month)
ggplot(tceqTemp, aes(x = month, y = Value, color = SegmentID)) + 
  geom_boxplot(fill = NA) +
  geom_jitter(position = position_jitter(0.2)) +
  xlab("Month") + ylab("Oxygen (mg/L)") + theme_bw() +
 theme(panel.grid.major = element_blank(), panel.grid.minor = element_blank()) +
  theme(legend.position = "right")
```

### save oxygen by month figure

```
ggsave("Fig_S8_oxygen_month.png")
```

```
## Saving 7 x 5 in image
```

#### plot N/P

```
monthTCEQsum11$DIN <- monthTCEQsum11$`NOx (uM)` + monthTCEQsum11$`NH4 (uM)`
monthTCEQsum11$NP <- monthTCEQsum11$DIN / monthTCEQsum11$`PO4 (uM)`
head(monthTCEQsum11)
```

```
## # A tibble: 6 x 15
##   SegmentID month  year SegMonth  Temperature `Salinity (PSU)` `Enterococci (MP~
##   <chr>     <int> <int> <chr>           <dbl>            <dbl>             <dbl>
## 1 1101          1  2014 1101 _ 1~        7.74            5.42                 30
## 2 1101          1  2020 1101 _ 1~       15               2.1                  31
## 3 1101          3  2015 1101 _ 3~       20.4             0.1                 340
## 4 1101          3  2016 1101 _ 3~       22.5             0.2                 110
## 5 1101          3  2017 1101 _ 3~       20.8             0.2                 420
## 6 1101          3  2018 1101 _ 3~       21.0             0.667                10
## # ... with 8 more variables: NOx (uM) <dbl>, NH4 (uM) <dbl>, PO4 (uM) <dbl>,
## #   pH <dbl>, Cond <dbl>, Oxygen <dbl>, DIN <dbl>, NP <dbl>
```

```
saveRDS(monthTCEQsum11, "monthTCEQsum11.rds")
```

#### ANOVA N/P

```
fit <- aov(NP ~ SegmentID*month, data=monthTCEQsum11)
summary(fit)
```

```
##                  Df Sum Sq Mean Sq F value Pr(>F)    
## SegmentID         4 1277.5   319.4  45.729 <2e-16 ***
## month             1    0.0     0.0   0.003  0.955    
## SegmentID:month   4   33.8     8.4   1.210  0.309    
## Residuals       145 1012.7     7.0                   
## ---
## Signif. codes:  0 '***' 0.001 '**' 0.01 '*' 0.05 '.' 0.1 ' ' 1
## 20 observations deleted due to missingness
```

#### Tukey test oxygen by seqment

```
tukeyTest <- HSD.test(fit, trt = 'SegmentID')
tukeyTest
```

```
## $statistics
##    MSerror  Df     Mean       CV
##   6.984341 145 4.822503 54.80122
## 
## $parameters
##    test    name.t ntr StudentizedRange alpha
##   Tukey SegmentID   5         3.906632  0.05
## 
## $means
##              NP      std  r        Min       Max       Q25       Q50       Q75
## 1101   6.291095 2.194853 21 2.34453782 10.333333 4.5761905  6.089286  8.378378
## 1101C  3.849975 3.509755 30 0.16402116 13.688312 1.0222485  3.141298  5.005473
## 1113B 10.690946 3.145573 26 6.37799043 19.383776 8.5161400 10.252246 11.889923
## 2425   2.993875 2.169584 52 0.06511511 10.333333 1.1598639  2.583333  4.428571
## 2425B  2.547290 2.054606 26 0.25881262  8.965157 0.8877143  2.274768  3.182102
## 
## $comparison
## NULL
## 
## $groups
##              NP groups
## 1113B 10.690946      a
## 1101   6.291095      b
## 1101C  3.849975      c
## 2425   2.993875      c
## 2425B  2.547290      c
## 
## attr(,"class")
## [1] "group"
```

#### Plot N/P

```
ggplot(monthTCEQsum11, aes(x = SegmentID, y = NP, color = SegmentID)) + 
  geom_violin(fill = NA) +
  geom_jitter(position = position_jitter(0.2)) +
#  geom_boxplot(fill = NA) +
  xlab("Segment") + ylab("N/P") + theme_bw() +
 theme(panel.grid.major = element_blank(), panel.grid.minor = element_blank()) +
  theme(legend.position = "none")
```

```
## Warning: Removed 19 rows containing non-finite values (stat_ydensity).
```

```
## Warning: Groups with fewer than two data points have been dropped.
```

```
## Warning: Removed 19 rows containing missing values (geom_point).
```

#### save N/P

```
ggsave("Fig_S9_NP.png")
```

```
## Saving 7 x 5 in image
```

```
## Warning: Removed 19 rows containing non-finite values (stat_ydensity).
```

```
## Warning: Groups with fewer than two data points have been dropped.
```

```
## Warning: Removed 19 rows containing missing values (geom_point).
```
