## Supplemental File S03 for "Hurricane Harvey Impacts on Water Quality and Microbial Communities in Houston, TX Waterbodies": S03_metaData.html

HHmetaData


### HHmetaData

###### Michael G. LaMontagne

#### 11/25/2021

#### R Markdown

This is a R Markdown document presents environmental data (salinity, nutrients..) and FIB counts for samples collected from August 25 through October 30th.

#### Import HH metadata and plot nitrate

```
library(stringr)
library(dplyr)
```

```
## 
## Attaching package: 'dplyr'
```

```
## The following objects are masked from 'package:stats':
## 
##     filter, lag
```

```
## The following objects are masked from 'package:base':
## 
##     intersect, setdiff, setequal, union
```

```
library(ggplot2)
library(ggbreak)
```

```
## ggbreak v0.0.7
## 
## If you use ggbreak in published research, please cite the following paper:
## 
## S Xu, M Chen, T Feng, L Zhan, L Zhou, G Yu. Use ggbreak to effectively utilize plotting space to deal with large datasets and outliers. Frontiers in Genetics. 2021, 12:774846. doi: 10.3389/fgene.2021.774846
```

```
library(forcats)
library(agricolae)
sessionInfo()
```

```
## R version 4.1.1 (2021-08-10)
## Platform: x86_64-w64-mingw32/x64 (64-bit)
## Running under: Windows 10 x64 (build 22000)
## 
## Matrix products: default
## 
## locale:
## [1] LC_COLLATE=English_United States.1252 
## [2] LC_CTYPE=English_United States.1252   
## [3] LC_MONETARY=English_United States.1252
## [4] LC_NUMERIC=C                          
## [5] LC_TIME=English_United States.1252    
## 
## attached base packages:
## [1] stats     graphics  grDevices utils     datasets  methods   base     
## 
## other attached packages:
## [1] agricolae_1.3-5 forcats_0.5.1   ggbreak_0.0.7   ggplot2_3.3.5  
## [5] dplyr_1.0.7     stringr_1.4.0  
## 
## loaded via a namespace (and not attached):
##  [1] tidyselect_1.1.1   xfun_0.25          bslib_0.3.1        purrr_0.3.4       
##  [5] lattice_0.20-44    haven_2.4.3        labelled_2.8.0     ggfun_0.0.4       
##  [9] colorspace_2.0-2   vctrs_0.3.8        generics_0.1.1     miniUI_0.1.1.1    
## [13] htmltools_0.5.2    yaml_2.2.1         AlgDesign_1.2.0    utf8_1.2.2        
## [17] gridGraphics_0.5-1 rlang_0.4.11       later_1.3.0        jquerylib_0.1.4   
## [21] pillar_1.6.4       glue_1.4.2         withr_2.4.2        DBI_1.1.1         
## [25] lifecycle_1.0.1    questionr_0.7.5    munsell_0.5.0      combinat_0.0-8    
## [29] gtable_0.3.0       evaluate_0.14      knitr_1.36         fastmap_1.1.0     
## [33] httpuv_1.6.3       fansi_0.5.0        highr_0.9          Rcpp_1.0.7        
## [37] xtable_1.8-4       promises_1.2.0.1   scales_1.1.1       jsonlite_1.7.2    
## [41] mime_0.12          hms_1.1.1          klaR_0.6-15        digest_0.6.27     
## [45] aplot_0.1.1        stringi_1.7.5      shiny_1.7.1        grid_4.1.1        
## [49] tools_4.1.1        yulab.utils_0.0.4  magrittr_2.0.1     sass_0.4.0        
## [53] patchwork_1.1.1    tibble_3.1.4       cluster_2.1.2      crayon_1.4.1      
## [57] pkgconfig_2.0.3    ellipsis_0.3.2     MASS_7.3-54        ggplotify_0.1.0   
## [61] rstudioapi_0.13    assertthat_0.2.1   rmarkdown_2.11     R6_2.5.1          
## [65] nlme_3.1-152       compiler_4.1.1
```

#### Including Plots

```
HHmetaData <- readRDS("metaHHfactor.rds")
tail(HHmetaData)
```

```
##    Station SegmentID   type month season       Date NOxuM NH4uM PuM   pH
## hC     C34      2425   post    10   fall 2017-11-01   1.5   1.8 2.1 7.83
## hJ     J76     2425B   post    10   fall 2017-11-01   0.9   3.7 1.4 7.48
## hK     K45      2425   post    10   fall 2017-11-01   1.3   2.9 1.0 7.31
## iH     H08     1113B   post     3 spring 2018-03-17    NA    NA  NA   NA
## jH     H08     1113B   post     4 spring 2018-04-25    NA    NA  NA   NA
## kS    SH13     1113B sewage     4 spring 2018-04-25    NA    NA  NA   NA
##    Conductivity   TDS Salinity SterivexML Oxygen   Temp Total Ecoli Enterococii
## hC        12.16 10935      7.0        112 10.205 17.375  4640    23        1733
## hJ        15.30 13772      9.0        125 10.235 17.205 22820  1414         310
## hK        18.07 16263     10.8        146  8.245 17.420 27550   435         100
## iH           NA    NA       NA         NA     NA     NA    NA    NA          NA
## jH           NA    NA       NA         NA     NA     NA    NA    NA          NA
## kS           NA    NA       NA         NA     NA     NA    NA    NA          NA
##    factorDate
## hC     30-Oct
## hJ     30-Oct
## hK     30-Oct
## iH     15-Mar
## jH     23-Apr
## kS     23-Apr
```

```
# write.csv(HHmetaData, "HHmetaData.csv")
```

#### HH and TCEQ

```
HHmetaData3 <- read.csv("monthTCEQ_HH.csv")
head(HHmetaData3)
```

```
##   SegmentID type month season      NOxuM     NH4uM     PO4uM       pH
## 1      2425 TCEQ     8   <NA>   2.857143  3.571429        NA 8.566667
## 2      2425 TCEQ     8   <NA>   1.428571  1.428571  9.854839 8.575000
## 3      2425 TCEQ     8   <NA>   2.857143  3.571429 14.032258 8.583333
## 4     1101C TCEQ     8   <NA>   2.857143 21.428571  6.451613 7.566667
## 5     1113B TCEQ     8   <NA> 396.785714 36.428571        NA 7.716667
## 6     1113B TCEQ     8   <NA> 267.142857 29.607143 40.967742 7.933333
##   Conductivity   Salinity    Oxygen Temperature Ecoli EnterococciMPN factorDate
## 1    24150.000 14.6833333 6.8000000    30.36667    NA           10.0     15-Aug
## 2    17550.000 10.4000000 7.0750000    32.22500    NA             NA     15-Aug
## 3    29000.000 17.7833333 7.1833333    31.80000    NA           10.0     15-Aug
## 4     1773.333  2.0000000 0.8333333    30.30000    NA          160.0     15-Aug
## 5     5461.667  3.1000000 3.9166667    30.16667    NA           74.0     15-Aug
## 6     1455.333  0.7833333 1.6333333    31.08333    NA          161.5     15-Aug
##     Day
## 1 44423
## 2 44423
## 3 44423
## 4 44423
## 5 44423
## 6 44423
```

#### Define factors

```
#HHmetaData2 <- HHmetaData[-which(HHmetaData$season == "spring"),]
#names(HHmetaData2)[16] <- "Temperature"
#names(HHmetaData2)[9] <- "PO4uM"
HHmetaData3$DINuM <- HHmetaData3$NOxuM + HHmetaData3$NH4uM
HHmetaData3$NP <- HHmetaData3$DINuM / HHmetaData3$PO4uM
HHmetaData3$logEC <- log10(HHmetaData3$EnterococciMPN)
head(HHmetaData3)
```

```
##   SegmentID type month season      NOxuM     NH4uM     PO4uM       pH
## 1      2425 TCEQ     8   <NA>   2.857143  3.571429        NA 8.566667
## 2      2425 TCEQ     8   <NA>   1.428571  1.428571  9.854839 8.575000
## 3      2425 TCEQ     8   <NA>   2.857143  3.571429 14.032258 8.583333
## 4     1101C TCEQ     8   <NA>   2.857143 21.428571  6.451613 7.566667
## 5     1113B TCEQ     8   <NA> 396.785714 36.428571        NA 7.716667
## 6     1113B TCEQ     8   <NA> 267.142857 29.607143 40.967742 7.933333
##   Conductivity   Salinity    Oxygen Temperature Ecoli EnterococciMPN factorDate
## 1    24150.000 14.6833333 6.8000000    30.36667    NA           10.0     15-Aug
## 2    17550.000 10.4000000 7.0750000    32.22500    NA             NA     15-Aug
## 3    29000.000 17.7833333 7.1833333    31.80000    NA           10.0     15-Aug
## 4     1773.333  2.0000000 0.8333333    30.30000    NA          160.0     15-Aug
## 5     5461.667  3.1000000 3.9166667    30.16667    NA           74.0     15-Aug
## 6     1455.333  0.7833333 1.6333333    31.08333    NA          161.5     15-Aug
##     Day      DINuM        NP    logEC
## 1 44423   6.428571        NA 1.000000
## 2 44423   2.857143 0.2899228       NA
## 3 44423   6.428571 0.4581281 1.000000
## 4 44423  24.285714 3.7642857 2.204120
## 5 44423 433.214286        NA 1.869232
## 6 44423 296.750000 7.2435039 2.208173
```

#### Salinity

```
n <- c(5:8, 10)
pairs(HHmetaData3[,n], pch = 19)
```

#### Set factors

```
HHmetaData3$SegmentID <- as.factor(HHmetaData3$SegmentID)
HHmetaData3$type <- as.factor(HHmetaData3$type)
HHmetaData3$factorDate <- as.factor(HHmetaData3$factorDate)
head(HHmetaData3)
```

```
##   SegmentID type month season      NOxuM     NH4uM     PO4uM       pH
## 1      2425 TCEQ     8   <NA>   2.857143  3.571429        NA 8.566667
## 2      2425 TCEQ     8   <NA>   1.428571  1.428571  9.854839 8.575000
## 3      2425 TCEQ     8   <NA>   2.857143  3.571429 14.032258 8.583333
## 4     1101C TCEQ     8   <NA>   2.857143 21.428571  6.451613 7.566667
## 5     1113B TCEQ     8   <NA> 396.785714 36.428571        NA 7.716667
## 6     1113B TCEQ     8   <NA> 267.142857 29.607143 40.967742 7.933333
##   Conductivity   Salinity    Oxygen Temperature Ecoli EnterococciMPN factorDate
## 1    24150.000 14.6833333 6.8000000    30.36667    NA           10.0     15-Aug
## 2    17550.000 10.4000000 7.0750000    32.22500    NA             NA     15-Aug
## 3    29000.000 17.7833333 7.1833333    31.80000    NA           10.0     15-Aug
## 4     1773.333  2.0000000 0.8333333    30.30000    NA          160.0     15-Aug
## 5     5461.667  3.1000000 3.9166667    30.16667    NA           74.0     15-Aug
## 6     1455.333  0.7833333 1.6333333    31.08333    NA          161.5     15-Aug
##     Day      DINuM        NP    logEC
## 1 44423   6.428571        NA 1.000000
## 2 44423   2.857143 0.2899228       NA
## 3 44423   6.428571 0.4581281 1.000000
## 4 44423  24.285714 3.7642857 2.204120
## 5 44423 433.214286        NA 1.869232
## 6 44423 296.750000 7.2435039 2.208173
```

#### Remove TCEQ data

```
HHmetaData2 <- HHmetaData3[which(HHmetaData3$type != "TCEQ"),]
dim(HHmetaData2)
```

```
## [1] 41 19
```

#### Plot salinity by segement and date to generate Figure 2

```
HHmetaData2 %>%
  dplyr::mutate(factorDate = fct_reorder(factorDate, -dplyr::desc(Day))) %>%
ggplot(aes(x = factorDate, y = Salinity, shape = type, color = SegmentID)) + 
  geom_boxplot(fill = NA) +
  geom_jitter(position = position_jitter(0.2)) +
  xlab("Date") + ylab("Salinity (PSU)") + theme_bw() +
  theme(panel.grid.major = element_blank(), panel.grid.minor = element_blank()) +
  theme(legend.position = "right")
```

#### Save Figure 2

```
ggsave("Fig_2_SalinityHH.png")
```

```
## Saving 7 x 5 in image
```

#### Plot salinity and DIN

```
n <- c(5:7,10,17,18)
pairs(HHmetaData3[,n], pch = 19)
```

#### Plot nitrate by segment to compare TCEQ and HH data

```
HHmetaData3 %>%
  dplyr::mutate(type = fct_reorder(type, -dplyr::desc(Day))) %>%
ggplot(aes(x = SegmentID, y = NOxuM, color = type, shape = SegmentID)) + geom_boxplot(fill = NA) + geom_point(size = 2) +
  geom_jitter(position = position_jitter(0.05)) +
  xlab("Sample Type") + ylab("NOx \u00B5M") + theme_bw() +
  theme(panel.grid.major = element_blank(), panel.grid.minor = element_blank()) +
  theme(legend.position = "right")
```

```
## Warning: Removed 2 rows containing non-finite values (stat_boxplot).
```

```
## Warning: Removed 2 rows containing missing values (geom_point).

## Warning: Removed 2 rows containing missing values (geom_point).
```

#### ANOVA nitrate

```
ggsave("Fig_S3_NOxHH.png")
```

```
## Saving 7 x 5 in image
```

```
## Warning: Removed 2 rows containing non-finite values (stat_boxplot).
```

```
## Warning: Removed 2 rows containing missing values (geom_point).

## Warning: Removed 2 rows containing missing values (geom_point).
```

```
head(HHmetaData3)
```

```
##   SegmentID type month season      NOxuM     NH4uM     PO4uM       pH
## 1      2425 TCEQ     8   <NA>   2.857143  3.571429        NA 8.566667
## 2      2425 TCEQ     8   <NA>   1.428571  1.428571  9.854839 8.575000
## 3      2425 TCEQ     8   <NA>   2.857143  3.571429 14.032258 8.583333
## 4     1101C TCEQ     8   <NA>   2.857143 21.428571  6.451613 7.566667
## 5     1113B TCEQ     8   <NA> 396.785714 36.428571        NA 7.716667
## 6     1113B TCEQ     8   <NA> 267.142857 29.607143 40.967742 7.933333
##   Conductivity   Salinity    Oxygen Temperature Ecoli EnterococciMPN factorDate
## 1    24150.000 14.6833333 6.8000000    30.36667    NA           10.0     15-Aug
## 2    17550.000 10.4000000 7.0750000    32.22500    NA             NA     15-Aug
## 3    29000.000 17.7833333 7.1833333    31.80000    NA           10.0     15-Aug
## 4     1773.333  2.0000000 0.8333333    30.30000    NA          160.0     15-Aug
## 5     5461.667  3.1000000 3.9166667    30.16667    NA           74.0     15-Aug
## 6     1455.333  0.7833333 1.6333333    31.08333    NA          161.5     15-Aug
##     Day      DINuM        NP    logEC
## 1 44423   6.428571        NA 1.000000
## 2 44423   2.857143 0.2899228       NA
## 3 44423   6.428571 0.4581281 1.000000
## 4 44423  24.285714 3.7642857 2.204120
## 5 44423 433.214286        NA 1.869232
## 6 44423 296.750000 7.2435039 2.208173
```

#### ANOVA nitrate

```
fit <- aov(NOxuM ~ SegmentID*type, data=HHmetaData3)
summary(fit)
```

```
##                Df Sum Sq Mean Sq F value   Pr(>F)    
## SegmentID       4 572741  143185 159.398  < 2e-16 ***
## type            3  32016   10672  11.880 3.30e-06 ***
## SegmentID:type 10  64814    6481   7.215 2.37e-07 ***
## Residuals      60  53897     898                     
## ---
## Signif. codes:  0 '***' 0.001 '**' 0.01 '*' 0.05 '.' 0.1 ' ' 1
## 2 observations deleted due to missingness
```

#### ANOVA of nitrate/nitrite

```
tukeyTest <- HSD.test(fit, trt = 'type')
tukeyTest
```

```
## $statistics
##    MSerror Df     Mean       CV
##   898.2852 60 53.01438 56.53449
## 
## $parameters
##    test name.t ntr StudentizedRange alpha
##   Tukey   type   4         3.737089  0.05
## 
## $means
##          NOxuM        std  r      Min      Max      Q25      Q50    Q75
## HH    2.666667   3.153834  6 1.300000   9.1000 1.300000  1.35000  1.550
## post 45.010000  86.499067 30 0.900000 306.0000 0.925000  1.35000 25.975
## pre  15.625000  29.116705  4 1.000000  59.3000 1.075000  1.10000 15.650
## TCEQ 71.218985 112.535620 38 1.428571 396.7857 4.017857 19.28571 78.750
## 
## $comparison
## NULL
## 
## $groups
##          NOxuM groups
## TCEQ 71.218985      a
## post 45.010000      b
## pre  15.625000     bc
## HH    2.666667      c
## 
## attr(,"class")
## [1] "group"
```

#### Tukey test

```
tukeyTest <- HSD.test(fit, trt = 'SegmentID')
tukeyTest
```

```
## $statistics
##    MSerror Df     Mean       CV
##   898.2852 60 53.01438 56.53449
## 
## $parameters
##    test    name.t ntr StudentizedRange alpha
##   Tukey SegmentID   5         3.977418  0.05
## 
## $means
##            NOxuM        std  r Min       Max     Q25        Q50        Q75
## 1101   52.344156  38.138767 11 1.3 127.85714  24.050  47.857143  72.507143
## 1101C  23.891209  34.676964 13 0.9 102.20000   1.300   6.428571  35.714286
## 1113B 250.494048 103.770614 12 9.1 396.78571 211.525 258.521429 306.910714
## 2425    6.360345   9.334970 29 0.9  39.28571   1.200   1.700000   5.714286
## 2425B   4.490110   8.466351 13 0.9  32.14286   1.000   1.400000   3.928571
## 
## $comparison
## NULL
## 
## $groups
##            NOxuM groups
## 1113B 250.494048      a
## 1101   52.344156      b
## 1101C  23.891209     bc
## 2425    6.360345      c
## 2425B   4.490110      c
## 
## attr(,"class")
## [1] "group"
```

#### Including Plot nitrate/nitrite versus salinity to generate Figure 3.

```
HHmetaData3 %>%
  dplyr::mutate(type = fct_reorder(type, -dplyr::desc(Day))) %>%
ggplot(aes(x=Salinity, y = NOxuM, color = SegmentID, shape = type)) + geom_point(size = 3) + xlab("Salinity (PSU)") + ylab("NOx (\u00B5M)") + theme_bw() + theme_classic()
```

```
## Warning: Removed 2 rows containing missing values (geom_point).
```

#### Save plots for nitrate vs salinity

```
ggsave("Fig_3_Salinity_NOxHH.png")
```

```
## Saving 7 x 5 in image
```

```
## Warning: Removed 2 rows containing missing values (geom_point).
```

#### Plots for ammonium vs salinity

```
HHmetaData3 %>%
  dplyr::mutate(type = fct_reorder(type, -dplyr::desc(Day))) %>%
ggplot(aes(x=Salinity, y = NH4uM, color = SegmentID, shape = type)) + geom_point(size = 3) + xlab("Salinity (PSU)") + ylab("Ammonium (\u00B5M)") + theme_bw() + theme_classic()
```

```
## Warning: Removed 4 rows containing missing values (geom_point).
```

#### Save plot for ammonium vs salinity

```
ggsave("Fig_S4_Salinity_NH4.png")
```

```
## Saving 7 x 5 in image
```

```
## Warning: Removed 4 rows containing missing values (geom_point).
```

```
write.csv(HHmetaData3, "HHmetaData3.csv")
```

#### Plots for phosphate vs salinity

```
HHmetaData3 %>%
  dplyr::mutate(type = fct_reorder(type, -dplyr::desc(Day))) %>%
ggplot(aes(x=Salinity, y = PO4uM, color = SegmentID, shape = type)) + geom_point(size = 3) + xlab("Salinity (PSU)") + ylab("Phosphate (\u00B5M)") + theme_bw() + theme_classic()
```

```
## Warning: Removed 4 rows containing missing values (geom_point).
```

#### Save plot for phosphate vs salinity

```
ggsave("Fig_S5_Salinity_PO4.png")
```

```
## Saving 7 x 5 in image
```

```
## Warning: Removed 4 rows containing missing values (geom_point).
```

#### Plots for N/P vs salinity

```
HHmetaData3 %>%
  dplyr::mutate(type = fct_reorder(type, -dplyr::desc(Day))) %>%
ggplot(aes(x=Salinity, y = NP, color = SegmentID, shape = type)) + geom_point(size = 3) + xlab("Salinity (PSU)") + ylab("DIN/P") + theme_bw() + theme_classic()
```

```
## Warning: Removed 8 rows containing missing values (geom_point).
```

#### Save plot for DIN/P vs salinity

```
ggsave("Fig_S6_DIN_Salinity.png")
```

```
## Saving 7 x 5 in image
```

```
## Warning: Removed 8 rows containing missing values (geom_point).
```

#### ANOVA DIN/P

```
fit <- aov(NP ~ SegmentID*type, data=HHmetaData3)
summary(fit)
```

```
##                Df Sum Sq Mean Sq F value  Pr(>F)   
## SegmentID       4  506.0  126.50   5.076 0.00152 **
## type            3   31.3   10.45   0.419 0.73994   
## SegmentID:type 10   94.3    9.43   0.379 0.95075   
## Residuals      54 1345.8   24.92                   
## ---
## Signif. codes:  0 '***' 0.001 '**' 0.01 '*' 0.05 '.' 0.1 ' ' 1
## 8 observations deleted due to missingness
```

#### Tukey DIN/P

```
tukeyTest <- HSD.test(fit, trt = 'SegmentID')
tukeyTest
```

```
## $statistics
##    MSerror Df     Mean       CV
##   24.92267 54 3.864045 129.1978
## 
## $parameters
##    test    name.t ntr StudentizedRange alpha
##   Tukey SegmentID   5         3.991024  0.05
## 
## $means
##             NP       std  r       Min       Max       Q25       Q50      Q75
## 1101  4.799792  2.331134  9 1.4736842  8.833333 4.2027027 4.6957869 5.942857
## 1101C 6.992615 10.255805 13 0.1640212 37.851852 2.1220238 3.6000000 5.668571
## 1113B 7.956454  3.124326 10 2.4318182 13.228448 7.3608171 8.3070116 9.535472
## 2425  1.712150  1.498758 27 0.1607143  4.797619 0.4954945 1.2916667 2.406250
## 2425B 1.408966  1.277849 13 0.1730769  4.290179 0.5555556 0.8303571 2.234600
## 
## $comparison
## NULL
## 
## $groups
##             NP groups
## 1113B 7.956454      a
## 1101C 6.992615      a
## 1101  4.799792     ab
## 2425  1.712150      b
## 2425B 1.408966      b
## 
## attr(,"class")
## [1] "group"
```

#### Plot for E.coli

```
HHmetaData2 %>%
  dplyr::mutate(factorDate = fct_reorder(factorDate, -dplyr::desc(Day))) %>%
ggplot(aes(x = factorDate, y = Ecoli, shape = type, color = SegmentID)) + 
  geom_boxplot(fill = NA) +
  geom_jitter(position = position_jitter(0.2)) +
  xlab("Date") + ylab("Ecoli (MPN/100 ml)") + theme_bw() +
  theme(panel.grid.major = element_blank(), panel.grid.minor = element_blank()) +
  theme(legend.position = "right")
```

```
## Warning: Removed 5 rows containing non-finite values (stat_boxplot).
```

```
## Warning: Removed 5 rows containing missing values (geom_point).
```

#### Save plot for Ecoli

```
ggsave("Fig_4_Ecoli_HH.png")
```

```
## Saving 7 x 5 in image
```

```
## Warning: Removed 5 rows containing non-finite values (stat_boxplot).
```

```
## Warning: Removed 5 rows containing missing values (geom_point).
```

#### Plot for Enterococci

```
HHmetaData3 %>%
  dplyr::mutate(type = fct_reorder(type, -dplyr::desc(Day))) %>%
ggplot(aes(x = SegmentID, y = logEC, color = type, shape = SegmentID)) + geom_boxplot(fill = NA) + geom_point(size = 2) +
  geom_jitter(position = position_jitter(0.05)) +
  xlab("Water Body Segment") + ylab("log(Enterococci(MPN/100ml)") + theme_bw() +
  theme(panel.grid.major = element_blank(), panel.grid.minor = element_blank()) +
  theme(legend.position = "right")
```

```
## Warning: Removed 19 rows containing non-finite values (stat_boxplot).
```

```
## Warning: Removed 19 rows containing missing values (geom_point).

## Warning: Removed 19 rows containing missing values (geom_point).
```

#### ANOVA of Enterococci

```
ggsave("Fig_S7_Enterococci_segment.png")
```

```
## Saving 7 x 5 in image
```

```
## Warning: Removed 19 rows containing non-finite values (stat_boxplot).
```

```
## Warning: Removed 19 rows containing missing values (geom_point).

## Warning: Removed 19 rows containing missing values (geom_point).
```

```
fit <- aov(logEC ~ SegmentID + type, data=HHmetaData3)
summary(fit)
```

```
##             Df Sum Sq Mean Sq F value   Pr(>F)    
## SegmentID    4  2.807   0.702   1.662    0.172    
## type         1 11.125  11.125  26.347 3.85e-06 ***
## Residuals   55 23.223   0.422                     
## ---
## Signif. codes:  0 '***' 0.001 '**' 0.01 '*' 0.05 '.' 0.1 ' ' 1
## 19 observations deleted due to missingness
```

#### ANOVA of Enterococci

```
tukeyTest <- HSD.test(fit, trt = 'type')
tukeyTest
```

```
## $statistics
##     MSerror Df     Mean       CV
##   0.4222452 55 2.196069 29.58942
## 
## $parameters
##    test name.t ntr StudentizedRange alpha
##   Tukey   type   2         2.834147  0.05
## 
## $means
##         logEC       std  r      Min     Max      Q25      Q50      Q75
## post 2.695361 0.4297366 24 1.799341 3.48430 2.346740 2.750730 3.049218
## TCEQ 1.872203 0.8000618 37 1.000000 4.30103 1.176091 1.763428 2.230449
## 
## $comparison
## NULL
## 
## $groups
##         logEC groups
## post 2.695361      a
## TCEQ 1.872203      b
## 
## attr(,"class")
## [1] "group"
```

#### Plot for Oxygen

```
HHmetaData2 %>%
  dplyr::mutate(factorDate = fct_reorder(factorDate, -dplyr::desc(Day))) %>%
ggplot(aes(x = factorDate, y = Oxygen, shape = type, color = SegmentID)) + 
  geom_boxplot(fill = NA) +
  geom_jitter(position = position_jitter(0.2)) +
  xlab("Date") + ylab("Oxygen (mg/L)") + theme_bw() +
  theme(panel.grid.major = element_blank(), panel.grid.minor = element_blank()) +
  theme(legend.position = "right")
```

#### Oxygen by segment

```
HHmetaData3 %>%
  dplyr::mutate(type = fct_reorder(type, -dplyr::desc(Day))) %>%
ggplot(aes(x = SegmentID, y = Oxygen, color = type, shape = SegmentID)) + geom_boxplot(fill = NA) + # geom_point(size = 2) +
  # geom_jitter(position = position_jitter(0.05)) +
  xlab("Water Body Segment") + ylab("Oxygen (mg/L)") + theme_bw() +
  theme(panel.grid.major = element_blank(), panel.grid.minor = element_blank()) +
  theme(legend.position = "right")
```

```
## Warning: Removed 2 rows containing non-finite values (stat_boxplot).
```

#### Oxygen by segment

```
ggsave("Fig_4_Oxygen_segment.png")
```

```
## Saving 7 x 5 in image
```

```
## Warning: Removed 2 rows containing non-finite values (stat_boxplot).
```

```
fit <- aov(Oxygen ~ SegmentID + type, data=HHmetaData3)
summary(fit)
```

```
##             Df Sum Sq Mean Sq F value   Pr(>F)    
## SegmentID    4  74.68  18.669   6.091 0.000289 ***
## type         3  17.83   5.945   1.940 0.131122    
## Residuals   70 214.54   3.065                     
## ---
## Signif. codes:  0 '***' 0.001 '**' 0.01 '*' 0.05 '.' 0.1 ' ' 1
## 2 observations deleted due to missingness
```

#### Oxygen by segment

```
tukeyTest <- HSD.test(fit, trt = 'SegmentID')
tukeyTest
```

```
## $statistics
##    MSerror Df     Mean       CV
##   3.064865 70 6.095151 28.72243
## 
## $parameters
##    test    name.t ntr StudentizedRange alpha
##   Tukey SegmentID   5         3.960014  0.05
## 
## $means
##         Oxygen      std  r       Min    Max      Q25      Q50      Q75
## 1101  5.451667 1.282853 11 3.8000000  8.100 4.733333 5.350000 6.087500
## 1101C 5.590000 2.813262 12 0.8333333 11.570 4.015833 5.133333 6.758333
## 1113B 4.404377 1.811488 13 1.6333333  7.320 3.410000 3.971429 5.428571
## 2425  6.860385 1.335836 30 5.2000000 10.205 5.890625 6.397500 7.457500
## 2425B 7.108750 1.852057 12 5.1000000 10.235 5.797500 6.212500 8.248750
## 
## $comparison
## NULL
## 
## $groups
##         Oxygen groups
## 2425B 7.108750      a
## 2425  6.860385      a
## 1101C 5.590000     ab
## 1101  5.451667     ab
## 1113B 4.404377      b
## 
## attr(,"class")
## [1] "group"
```

#### Oxygen by type

```
tukeyTest <- HSD.test(fit, trt = 'type')
tukeyTest
```

```
## $statistics
##    MSerror Df     Mean       CV
##   3.064865 70 6.095151 28.72243
## 
## $parameters
##    test name.t ntr StudentizedRange alpha
##   Tukey   type   4         3.721978  0.05
## 
## $means
##        Oxygen       std  r       Min   Max      Q25    Q50      Q75
## HH   5.203333 0.5259911  6 4.4200000  6.00 4.975000 5.2750 5.350000
## post 6.606167 2.0473404 30 2.7100000 11.57 5.728750 6.2125 7.728750
## pre  6.838000 0.9910701  5 5.7000000  8.10 6.000000 6.9900 7.400000
## TCEQ 5.725048 2.1065287 37 0.8333333 10.00 4.828571 5.6000 6.885714
## 
## $comparison
## NULL
## 
## $groups
##        Oxygen groups
## pre  6.838000      a
## post 6.606167      a
## TCEQ 5.725048      a
## HH   5.203333      a
## 
## attr(,"class")
## [1] "group"
```
