## Supplemental File S05 for "Hurricane Harvey Impacts on Water Quality and Microbial Communities in Houston, TX Waterbodies": S05_HHdada.html

Alpha Diversity Hurricane Harvey UHCL


### Alpha Diversity Hurricane Harvey UHCL

###### Michael G. LaMontagne

#### 12/31/2021

#### R Markdown

This is R Markdown document generates ASVs from 16S amplicon reads in DADA2 and generates a phyloseq object that is passed off to phyloseq. That phyloseq object will be used in correspondence analysis to determine environmental variable that correspond to bacteria communities associated with floodwaters.

```
library(philentropy)
packageVersion("philentropy")
```

```
## [1] '0.5.0'
```

```
library(dada2)
```

```
## Loading required package: Rcpp
```

```
packageVersion("dada2")
```

```
## [1] '1.20.0'
```

```
library(gridExtra)
packageVersion("gridExtra")
```

```
## [1] '2.3'
```

```
library(ggplot2)
packageVersion("ggplot2")
```

```
## [1] '3.3.5'
```

```
library(phyloseq)
```

```
## 
## Attaching package: 'phyloseq'
```

```
## The following object is masked from 'package:philentropy':
## 
##     distance
```

```
packageVersion("phyloseq")
```

```
## [1] '1.36.0'
```

```
library(DECIPHER); packageVersion("DECIPHER")
```

```
## Loading required package: Biostrings
```

```
## Loading required package: BiocGenerics
```

```
## Loading required package: parallel
```

```
## 
## Attaching package: 'BiocGenerics'
```

```
## The following objects are masked from 'package:parallel':
## 
##     clusterApply, clusterApplyLB, clusterCall, clusterEvalQ,
##     clusterExport, clusterMap, parApply, parCapply, parLapply,
##     parLapplyLB, parRapply, parSapply, parSapplyLB
```

```
## The following object is masked from 'package:gridExtra':
## 
##     combine
```

```
## The following objects are masked from 'package:stats':
## 
##     IQR, mad, sd, var, xtabs
```

```
## The following objects are masked from 'package:base':
## 
##     anyDuplicated, append, as.data.frame, basename, cbind, colnames,
##     dirname, do.call, duplicated, eval, evalq, Filter, Find, get, grep,
##     grepl, intersect, is.unsorted, lapply, Map, mapply, match, mget,
##     order, paste, pmax, pmax.int, pmin, pmin.int, Position, rank,
##     rbind, Reduce, rownames, sapply, setdiff, sort, table, tapply,
##     union, unique, unsplit, which.max, which.min
```

```
## Loading required package: S4Vectors
```

```
## Loading required package: stats4
```

```
## 
## Attaching package: 'S4Vectors'
```

```
## The following objects are masked from 'package:base':
## 
##     expand.grid, I, unname
```

```
## Loading required package: IRanges
```

```
## 
## Attaching package: 'IRanges'
```

```
## The following object is masked from 'package:phyloseq':
## 
##     distance
```

```
## The following object is masked from 'package:philentropy':
## 
##     distance
```

```
## The following object is masked from 'package:grDevices':
## 
##     windows
```

```
## Loading required package: XVector
```

```
## Loading required package: GenomeInfoDb
```

```
## 
## Attaching package: 'Biostrings'
```

```
## The following object is masked from 'package:base':
## 
##     strsplit
```

```
## Loading required package: RSQLite
```

```
## [1] '2.20.0'
```

```
library(phangorn)
```

```
## Loading required package: ape
```

```
## 
## Attaching package: 'ape'
```

```
## The following object is masked from 'package:Biostrings':
## 
##     complement
```

```
packageVersion("phangorn")
```

```
## [1] '2.7.1'
```

```
library(vegan)
```

```
## Loading required package: permute
```

```
## Loading required package: lattice
```

```
## This is vegan 2.5-7
```

```
## 
## Attaching package: 'vegan'
```

```
## The following objects are masked from 'package:phangorn':
## 
##     diversity, treedist
```

```
packageVersion("vegan")
```

```
## [1] '2.5.7'
```

```
library(agricolae)
```

```
## Registered S3 methods overwritten by 'klaR':
##   method      from 
##   predict.rda vegan
##   print.rda   vegan
##   plot.rda    vegan
```

```
## 
## Attaching package: 'agricolae'
```

```
## The following object is masked from 'package:ape':
## 
##     consensus
```

```
packageVersion("agricolae")
```

```
## [1] '1.3.5'
```

```
library(dplyr)
```

```
## 
## Attaching package: 'dplyr'
```

```
## The following objects are masked from 'package:Biostrings':
## 
##     collapse, intersect, setdiff, setequal, union
```

```
## The following object is masked from 'package:GenomeInfoDb':
## 
##     intersect
```

```
## The following object is masked from 'package:XVector':
## 
##     slice
```

```
## The following objects are masked from 'package:IRanges':
## 
##     collapse, desc, intersect, setdiff, slice, union
```

```
## The following objects are masked from 'package:S4Vectors':
## 
##     first, intersect, rename, setdiff, setequal, union
```

```
## The following objects are masked from 'package:BiocGenerics':
## 
##     combine, intersect, setdiff, union
```

```
## The following object is masked from 'package:gridExtra':
## 
##     combine
```

```
## The following objects are masked from 'package:stats':
## 
##     filter, lag
```

```
## The following objects are masked from 'package:base':
## 
##     intersect, setdiff, setequal, union
```

```
packageVersion("dplyr")
```

```
## [1] '1.0.7'
```

```
sessionInfo()
```

```
## R version 4.1.1 (2021-08-10)
## Platform: x86_64-w64-mingw32/x64 (64-bit)
## Running under: Windows 10 x64 (build 22000)
## 
## Matrix products: default
## 
## locale:
## [1] LC_COLLATE=English_United States.1252 
## [2] LC_CTYPE=English_United States.1252   
## [3] LC_MONETARY=English_United States.1252
## [4] LC_NUMERIC=C                          
## [5] LC_TIME=English_United States.1252    
## 
## attached base packages:
## [1] stats4    parallel  stats     graphics  grDevices utils     datasets 
## [8] methods   base     
## 
## other attached packages:
##  [1] dplyr_1.0.7         agricolae_1.3-5     vegan_2.5-7        
##  [4] lattice_0.20-44     permute_0.9-5       phangorn_2.7.1     
##  [7] ape_5.5             DECIPHER_2.20.0     RSQLite_2.2.8      
## [10] Biostrings_2.60.2   GenomeInfoDb_1.28.2 XVector_0.32.0     
## [13] IRanges_2.26.0      S4Vectors_0.30.0    BiocGenerics_0.38.0
## [16] phyloseq_1.36.0     ggplot2_3.3.5       gridExtra_2.3      
## [19] dada2_1.20.0        Rcpp_1.0.7          philentropy_0.5.0  
## 
## loaded via a namespace (and not attached):
##   [1] colorspace_2.0-2            hwriter_1.3.2              
##   [3] ellipsis_0.3.2              GenomicRanges_1.44.0       
##   [5] rstudioapi_0.13             bit64_4.0.5                
##   [7] fansi_0.5.0                 codetools_0.2-18           
##   [9] splines_4.1.1               cachem_1.0.6               
##  [11] knitr_1.36                  ade4_1.7-18                
##  [13] jsonlite_1.7.2              Rsamtools_2.8.0            
##  [15] cluster_2.1.2               png_0.1-7                  
##  [17] shiny_1.7.1                 compiler_4.1.1             
##  [19] assertthat_0.2.1            Matrix_1.3-4               
##  [21] fastmap_1.1.0               later_1.3.0                
##  [23] htmltools_0.5.2             tools_4.1.1                
##  [25] igraph_1.2.7                gtable_0.3.0               
##  [27] glue_1.4.2                  GenomeInfoDbData_1.2.6     
##  [29] reshape2_1.4.4              ShortRead_1.50.0           
##  [31] fastmatch_1.1-3             Biobase_2.52.0             
##  [33] jquerylib_0.1.4             vctrs_0.3.8                
##  [35] rhdf5filters_1.4.0          multtest_2.48.0            
##  [37] nlme_3.1-152                iterators_1.0.13           
##  [39] xfun_0.25                   stringr_1.4.0              
##  [41] mime_0.12                   miniUI_0.1.1.1             
##  [43] lifecycle_1.0.1             zlibbioc_1.38.0            
##  [45] MASS_7.3-54                 scales_1.1.1               
##  [47] hms_1.1.1                   promises_1.2.0.1           
##  [49] MatrixGenerics_1.4.3        SummarizedExperiment_1.22.0
##  [51] biomformat_1.20.0           rhdf5_2.36.0               
##  [53] RColorBrewer_1.1-2          yaml_2.2.1                 
##  [55] memoise_2.0.0               sass_0.4.0                 
##  [57] labelled_2.8.0              latticeExtra_0.6-29        
##  [59] stringi_1.7.5               AlgDesign_1.2.0            
##  [61] highr_0.9                   klaR_0.6-15                
##  [63] foreach_1.5.1               BiocParallel_1.26.2        
##  [65] rlang_0.4.11                pkgconfig_2.0.3            
##  [67] matrixStats_0.61.0          bitops_1.0-7               
##  [69] evaluate_0.14               purrr_0.3.4                
##  [71] Rhdf5lib_1.14.2             GenomicAlignments_1.28.0   
##  [73] bit_4.0.4                   tidyselect_1.1.1           
##  [75] plyr_1.8.6                  magrittr_2.0.1             
##  [77] R6_2.5.1                    generics_0.1.1             
##  [79] combinat_0.0-8              DelayedArray_0.18.0        
##  [81] DBI_1.1.1                   haven_2.4.3                
##  [83] pillar_1.6.4                withr_2.4.2                
##  [85] mgcv_1.8-36                 survival_3.2-11            
##  [87] RCurl_1.98-1.5              tibble_3.1.4               
##  [89] crayon_1.4.1                questionr_0.7.5            
##  [91] utf8_1.2.2                  rmarkdown_2.11             
##  [93] jpeg_0.1-9                  grid_4.1.1                 
##  [95] data.table_1.14.2           blob_1.2.2                 
##  [97] forcats_0.5.1               digest_0.6.27              
##  [99] xtable_1.8-4                httpuv_1.6.3               
## [101] RcppParallel_5.1.4          munsell_0.5.0              
## [103] bslib_0.3.1                 quadprog_1.5-8
```

#### list fastq

```
path <- "C:/Users/mglam/Dropbox/BIOINFO/Rscripts/HHdada/UNTreads" 
head(list.files(path))
```

```
## [1] "A6A_S33_L001_R1_001.fastq.gz"      "A6A_S33_L001_R2_001.fastq.gz"     
## [3] "C34SEP25_S40_L001_R1_001.fastq.gz" "C34SEP25_S40_L001_R2_001.fastq.gz"
## [5] "C35OCT2_S46_L001_R1_001.fastq.gz"  "C35OCT2_S46_L001_R2_001.fastq.gz"
```

#### Including Plots

```
# Forward and reverse fastq filenames have format: SAMPLENAME_R1_001.fastq and SAMPLENAME_R2_001.fastq
fnFs <- sort(list.files(path, pattern="_R1_001.fastq.gz", full.names = TRUE))
fnRs <- sort(list.files(path, pattern="_R2_001.fastq.gz", full.names = TRUE))
# Extract sample names, assuming filenames have format: SAMPLENAME_XXX.fastq
sample.names <- sapply(strsplit(basename(fnFs), "_"), `[`, 1)
## plot some quality profiles from forward
plotQualityProfile(fnFs[1:2])
```

```
#saveRDS(fnFs, "HHfnFs.rds")
## and reverse
plotQualityProfile(fnRs[1:2])
```

```
#saveRDS(fnRs, "HHfnRs.rds")
```

#### Filter reads

```
# Place filtered files in filtered/ subdirectory
filtFs <- file.path(path, "filtered", paste0(sample.names, "_F_filt.fastq.gz"))
filtRs <- file.path(path, "filtered", paste0(sample.names, "_R_filt.fastq.gz"))
head(filtFs)
```

```
## [1] "C:/Users/mglam/Dropbox/BIOINFO/Rscripts/HHdada/UNTreads/filtered/A6A_F_filt.fastq.gz"     
## [2] "C:/Users/mglam/Dropbox/BIOINFO/Rscripts/HHdada/UNTreads/filtered/C34SEP25_F_filt.fastq.gz"
## [3] "C:/Users/mglam/Dropbox/BIOINFO/Rscripts/HHdada/UNTreads/filtered/C35OCT2_F_filt.fastq.gz" 
## [4] "C:/Users/mglam/Dropbox/BIOINFO/Rscripts/HHdada/UNTreads/filtered/C6C_F_filt.fastq.gz"     
## [5] "C:/Users/mglam/Dropbox/BIOINFO/Rscripts/HHdada/UNTreads/filtered/CA25_F_filt.fastq.gz"    
## [6] "C:/Users/mglam/Dropbox/BIOINFO/Rscripts/HHdada/UNTreads/filtered/CA30_F_filt.fastq.gz"
```

#### Filter and trim (skip in markdown)

```
t1 <- 219
t2 <- 160
t3 <- 20
#outHH <- filterAndTrim(fnFs, filtFs, fnRs, filtRs, trimLeft=t3, truncLen=c(t1,t2),
#                     maxN=0, maxEE=c(2,2), truncQ=2, rm.phix=TRUE,
#                     compress=TRUE, multithread=FALSE)
# saveRDS(outHH, "outHH.rds")
outHH <- readRDS("outHH.rds")
head(outHH)
```

```
##                                   reads.in reads.out
## A6A_S33_L001_R1_001.fastq.gz        151699    139654
## C34SEP25_S40_L001_R1_001.fastq.gz   142049    132852
## C35OCT2_S46_L001_R1_001.fastq.gz    164534    152672
## C6C_S34_L001_R1_001.fastq.gz        188069    176819
## CA25_S2_L001_R1_001.fastq.gz        191428    178167
## CA30_S8_L001_R1_001.fastq.gz        153976    140764
```

#### Learn error rates (skip in markdown)

```
#errR <- learnErrors(filtRs, randomize = TRUE, multithread=TRUE, nbases = 1e8)
#errF <- learnErrors(filtFs, randomize = TRUE, multithread=TRUE, nbases = 1e8)
#saveRDS(errR, "errRhh.rds")
#saveRDS(errF, "errFhh.rds")
errR <- readRDS("errRhh.rds")
errF <- readRDS("errFhh.rds")
plotErrors(errF, nominalQ=TRUE)
```

#### Dereplication

```
#derepFs <- derepFastq(filtFs, verbose=TRUE)
#derepRs <- derepFastq(filtRs, verbose=TRUE)
## Name the derep-class objects by the sample names
#names(derepFs) <- sample.names
#names(derepRs) <- sample.names
## run the core sample inference program
#dadaFs <- dada(derepFs, err=errF, multithread=TRUE)
#dadaRs <- dada(derepRs, err=errR, multithread=TRUE)
## insepct one
#dadaFs[[1]]
## merge paired reads
#mergers <- mergePairs(dadaFs, derepFs, dadaRs, derepRs, verbose=TRUE)
# Inspect the merger data.frame from the first sample
#saveRDS(mergers, "mergersHH.rds")
mergers <- readRDS("mergersHH.rds")
head(mergers[1])
```

```
## $A6A
##                                                                                                                                                                                                                                                                                                                  sequence
## 1                                                            ACGTAGGGTGCAAGCGTTAATCGGAATTACTGGGCGTAAAGCGTGCGCAGGCGGACTTTTAAGCCAGATGTGAAAGCCCCGAGCTTAACTTGGGAATTGCGTTTGGAACTGGGAGTCTAGAGTCTGTCAGAGGGGGATGGAATTCCACGTGTAGCAGTGAAATGCGTAGAGATGTGGAGGAACACCGATGGCGAAGGCAGTCCCCTGGGATAAGACTGACGCTCATGCACGAAAGCGTGGGGAGCAAACAGG
## 2                                                            ACAGAGGATGCAAGCGTTATCCGGAATCACTGGGCATAAAGCGTCTGTAGGTGGTTTGGTAAGTCTGCTGTTAAAGACTGGGGCTCAACCCCAGAAAAGCAGTGGAAACTGCTAGACTTGAGTGTGGTAGAGGTAGAGGGAATTCCTAGTGTAGCGGTGAAATGCGTAGATATTAGGAAGAACACCAATGGCGAAGGCACTCTACTGGACCATAACTGACACTGAGAGACGACAGCTAGGGGAGCAAATGGG
## 3                                                            ACGAAGGGTGCAAGCGTTATTCGGAATAATTGGGCGTAAAGGGTTCGTAGGTTGGATTGTAAGTCAAGTGTGAAATCCCCAGGCTTAACCTGGGACGTGCATTTGAAACTGTGATTCTTGAGTATTGGAGAGGGTAGTGGAATTGTAGGTGTAGGAGTGACATCCGTAGAGATCTGCAGGAACATCAGAGGCGAAGGCGACTACCTGGCCGATTACTGACACTGAGGAACGAAAGCGTGGGGAGCAAACAGG
## 4                                                            ACGGGGGGTGCAAGCGTTACTCGGAATCACTGGGCGTAAAGAGCATGTAGGCGGATTAATAAGTTTGAAGTGAAATCCTATAGCTTAACTATAGAACTGCTTTGAAAACTGTTAATCTAGAATGTGGGAGAGGTAGATGGAATTTCTGGTGTAGGGGTAAAATCCGTAGAGATCAGAAGGAATACCGATTGCGAAGGCGATCTACTGGAACATTATTGACGCTGAGATGCGAAAGCGTGGGGAGCAAACAGG
## 5                                                            ACGTAGGGTGCAAGCGTTAATCGGAATTACTGGGCGTAAAGCGTGCGCAGGCGGTTATATAAGTCAGATGTGAAATCCCCGGGCTCAACCTGGGAACTGCATTTGAGACTGTATAGCTAGAGTACGGTAGAGGGGGATGGAATTCCGCGTGTAGCAGTGAAATGCGTAGATATGCGGAGGAACACCGATGGCGAAGGCAATCCCCTGGACCTGTACTGACGCTCATGCACGAAAGCGTGGGGAGCAAACAGG
## 6                                                            ACATAGGGTGCAAGCGTTGTCCGGAATTATTGGGCGTAAAGAGCTCGTAGGTGGTTCGTCACGTCGGATGTGAAACTCTGGGGCTTAACCCCAGACCTGCATTCGATACGGGCGAGCTTGAGTATGGTAGGGGAGTCTGGAATTCCTGGTGTAGCGGTGGAATGCGCAGATATCAGGAGGAACACCAATGGCGAAGGCAGGACTCTGGGCCATTACTGACACTGAGGAGCGAAAGCGTGGGGAGCGAACAGG
## 7                                                            ACGTAGGGTGCGAGCGTTAATCGGAATTACTGGGCGTAAAGCGTGCGCAGGCGGTTTCGTAAGACAGAGGTGAAATCCCCGGGCTCAACCTGGGAACTGCCTTTGTGACTGCGAGGCTAGAGTATGGCAGAGGGGGGTGGAATTCCACGTGTAGCAGTGAAATGCGTAGAGATGTGGAGGAACACCGATGGCGAAGGCAGCCCCCTGGGCCAATACTGACGCTCATGCACGAAAGCGTGGGGAGCAAACAGG
## 8                                                            ACGTAGGGTGCAAGCGTTAATCGGAATTACTGGGCGTAAAGCGTGCGCAGGCGGTTATGTAAGACAGAGGTGAAATCCCCGGGCTCAACCTGGGAACTGCCTTTGTGACTGCATAGCTGGAGTGCGGCAGAGGGGGATGGAATTCCGCGTGTAGCAGTGAAATGCGTAGATATGCGGAGGAACACCGATGGCGAAGGCAATCCCCTGGGCCTGCACTGACGCTCATGCACGAAAGCGTGGGGAGCAAACAGG
## 9                                                            ACGAGTGCCTCAAGCGTTACCCGGAATCACTGGGCGTAAAGGTTGTGTAGGTGGTAACATTAGTCTTTTGTGAAAGCCCGCGGCTCAACCGCGGATCCGCAAAGGAAACGGTGTAACTAGAGGATGCAAGAGGTGTGCGGAACTCATAGTGTAGCGGTGAAATGCGTTGATATTATGGGGAACACCAAAAGCGAAGGCAGCACACTGGTGCACTCCTGACACTGAAACAAGAAAGCGTAGGTAGCGAATGGG
## 10                                                           ACGTAGGGTGCAAGCGTTAATCGGAATTACTGGGCGTAAAGCGTGCGCAGGCGGTTATGCAAGACAGAGGTGAAATCCCCGGGCTCAACCTGGGAACTGCCTTTGTGACTGCATAGCTAGAGTACGGCAGAGGGGGATGGAATTCCGCGTGTAGCAGTGAAATGCGTAGATATGCGGAGGAACACCGATGGCGAAGGCAATCCCCTGGGCCTGTACTGACGCTCATGCACGAAAGCGTGGGGAGCAAACAGG
## 11                                                           ACGTAGGGTGCAAGCGTTAATCGGAATTACTGGGCGTAAAGCGTGCGCAGGCGGCTTTGCAAGACAGATGTGAAATCCCCGGGCTCAACCTGGGAACTGCATTTGTGACTGCAAGGCTAGAGTACGGTAGAGGGGGATGGAATTCCGCGTGTAGCAGTGAAATGCGTAGATATGCGGAGGAACACCAATGGCGAAGGCAATCCCCTGGACCTGTACTGACGCTCATGCACGAAAGCGTGGGGAGCAAACAGG
## 12                                                           ACGTAGGGTGCAAGCGTTAATCGGAATTACTGGGCGTAAAGCGTGCGCAGGCGGTTATGTAAGACAGAGGTGAAATCCCCGGGCTCAACCTGGGAACTGCCTTTGTGACTGCATAGCTAGAGTACGGCAGAGGGGGATGGAATTCCGCGTGTAGCAGTGAAATGCGTAGATATGCGGAGGAACACCGATGGCGAAGGCAATCCCCTGGGCCTGTACTGACGCTCATGCACGAAAGCGTGGGGAGCAAACAGG
## 13                                                           ACGTAGGGTGCAAGCGTTAATCGGAATTACTGGGCGTAAAGCGTGCGCAGGCGGTTATGTAAGACAGAGGTGAAATCCCCGGGCTCAACCTGGGAACTGCCTTTGTGACTGCATAGCTAGAGTACGGTAGAGGGGGATGGAATTCCGCGTGTAGCAGTGAAATGCGTAGATATGCGGAGGAACACCGATGGCGAAGGCAATCCCCTGGACCTGTACTGACGCTCATGCACGAAAGCGTGGGGAGCAAACAGG
## 14                                                           ACATAGGGTGCAAGCGTTGTCCGGAATTATTGGGCGTAAAGAGCTCGTAGGTCGTTTGTTACGTCGGATGTGAAAACCTGAGGCTCAACCTCAGGCCTGCATTCGATACGGGCAAACTAGAGTTTGGTAGGGGAGACTGGAATTCCTGGTGTAGCGGTGGAATGCGCAGATATCAGGAGGAACACCAATGGCGAAGGCAGGTCTCTGGGCCAATACTGACACTGAGGAGCGAAAGTCTGGGGAGCGAACAGG
## 15                                                           ACGTAGGGTGCGAGCGTTAATCGGAATTACTGGGCGTAAAGCGTGCGCAGGCGGTTATACAAGACAGGCGTGAAATCCCCGGGCTTAACCTGGGAATGGCGTCTGTGACTGTATAGCTAGAGTGTGTCAGAGGGGGGTGGAATTCCACGTGTAGCAGTGAAATGCGTAGATATGTGGAGGAACACCAATGGCGAAGGCAGCCCCCTGGGATAACACTGACGCTCATGCACGAAAGCGTGGGGAGCAAACAGG
## 16                                                           ACGGAGGGTGCAAGCGTTGTCCGGATTTATTGGGTTTAAAGGGTGCGCAGGTGGTTTATTAAGTCAGTGGTGAAAGACGGTCGCTCAACGATTGCAGTGCCATTGAAACTGGTAGACTTGAGTAAAGTAGAGGTGGGCGGAATTGATAGTGTAGCGGTGAAATGCATAGATATTATCAAGAACTCCAATTGCGTAGGCAGCTCACTTGGCTTTTACTGACACTCATGCACGAAAGTGTGGGTATCAAACAGG
## 17                                                           ACGGAGGGTGCAAGCGTTATCCGGATTCACTGGGTTTAAAGGGTGCGTAGGCGGGTAGGTAAGTCAGTGGTGAAATCCTGGAGCTCAACTCCAGAACTGCCATTGATACTATCTATCTTGAATATTGTGGAGGTAAGCGGAATATGTCATGTAGCGGTGAAATGCTTAGATATGACATAGAACACCCATTGCGAAGGCAGCTTACTACGCATATATTGACGCTGAGGCACGAAAGCGTGGGGATCAAACAGG
## 18                                                           ACGTAGGGTGCAAGCGTTAATCGGAATTACTGGGCGTAAAGCGTGCGCAGGCGGTTATATAAGACAGATGTGAAATCCCCGGGCTCAACCTGGGAACTGCATTTGTGACTGTATAGCTAGAGTACGGTAGAGGGGGATGGAATTCCGCGTGTAGCAGTGAAATGCGTAGATATGCGGAGGAACACCGATGGCGAAGGCAATCCCCTGGACCTGTACTGACGCTCATGCACGAAAGCGTGGGGAGCAAACAGG
## 19                                                           ACGTAGGGTGCAAGCGTTAATCGGAATTACTGGGCGTAAAGCGTGCGCAGGCGGTGATGTAAGACAGATGTGAAATCCCCGGGCTCAACCTGGGAACTGCATTTGTGACTGCATCGCTGGAGTGCGGCAGAGGGGGATGGAATTCCGCGTGTAGCAGTGAAATGCGTAGATATGCGGAGGAACACCGATGGCGAAGGCAATCCCCTGGGCCTGCACTGACGCTCATGCACGAAAGCGTGGGGAGCAAACAGG
## 20                                                           ACGTAGGGTGCGAGCGTTAATCGGAATTACTGGGCGTAAAGCGTGCGCAGGCGGTTTCGTAAGACAGACGTGAAATCCCCGGGCTCAACCTGGGAACTGCGTTTGTGACTGCGAGGCTAGAGTATGGCAGAGGGGGGTGGAATTCCACGTGTAGCAGTGAAATGCGTAGAGATGTGGAGGAACACCGATGGCGAAGGCAGCCCCCTGGGCCAATACTGACGCTCATGCACGAAAGCGTGGGGAGCAAACAGG
## 21                                                           ACGAAGGGTGCTAGCGTTATTCGGAATCACTGGGCGTAAAGAGCGTGTAGGCGGTTTGTTAAGTCAACTGTTAAAGCACCCAGCTTAACTGGGTAAAGGCAGGTGAAACTGGCAGGCTAGAGTGCAAGAGAGAGAAGTGGAATTGTCGGAGTAGCGGTAAAATGCGTAGATCTCGACAGGAACACCGGTTGCGAAGGCGGCTTCTTGGCTTGTAACTGACGCTAAAGCGCGAAAGCGTGGGGAGCAAACAGG
## 22                                                           ACATAGGGTGCAAGCGTTGTCCGGAATTATTGGGCGTAAAGAGCTCGTAGGTCGTTTGTCGCGTCGATTGTGAAAATCTGAGGCTCAACCTCAGACCTGCAGTCGATACGGGCAAACTAGAGTGTGGTAGGGGAGACTGGAATTCCTGGTGTAGCGGTGGAATGCGCAGATATCAGGAGGAACACCAATGGCGAAGGCAGGTCTCTGGGCCATAACTGACACTGAGGAGCGAAAGTGCGGGGAGCGAACAGG
## 23                                                           ACGTAGGGTGCGAGCGTTAATCGGAATTACTGGGCGTAAAGCGTGCGCAGGCGGTTTTGTAAGTCGGATGTGAAATCCCCGGGCTCAACCTGGGAACTGCGTTCGAAACTGCAAGGCTAGAGTGTGTCAGAGGGGGGTAGAATTCCACGTGTAGCAGTGAAATGCGTAGAGATGTGGAGGAATACCAATGGCGAAGGCAGCCCCCTGGGATAACACTGACGCTCATGCACGAAAGCGTGGGGAGCAAACAGG
## 24                                                           ACGTAGGGTGCAAGCGTTAATCGGAATTACTGGGCGTAAAGCGTGCGCAGGCGGTTATATAAGTCAGATGTGAAATCCCCGGGCTCAACCTGGGAACTGCATTTGAGACTGTATAGCTAGAGTACGGCAGAGGGGGATGGAATTCCGCGTGTAGCAGTGAAATGCGTAGATATGCGGAGGAACACCGATGGCGAAGGCAATCCCCTGGGCCTGTACTGACGCTCATGCACGAAAGCGTGGGGAGCAAACAGG
## 25                                                           ACGTAGGGTGCAAGCGTTAATCGGAATTACTGGGCGTAAAGCGTGCGCAGGCGGATGTTTAAGCCAGATGTGAAAGCCCCGAGCTTAACTTGGGAATTGCGTTTGGAACTGAGCATCTAGAGTCTGTCAGAGGGGGATGGAATTCCACGTGTAGCAGTGAAATGCGTAGAGATGTGGAGGAACACCGATGGCGAAGGCAGTCCCCTGGGATAAGACTGACGCTCATGCACGAAAGCGTGGGGAGCAAACAGG
## 26                                                           ACGTAGGGTGCAAGCGTTAATCGGAATTACTGGGCGTAAAGCGTGCGCAGGCGGTTCGGAAAGAAAGGTGTGAAATCCCAGGGCTTAACCTTGGAACTGCACTTTTAACTACCGAGCTAGAGTATGTCAGAGGGGGGTGGAATTCCGCGTGTAGCAGTGAAATGCGTAGATATGCGGAGGAACACCGATGGCGAAGGCAGCCCCCTGGGATAATACTGACGCTCATGCACGAAAGCGTGGGGAGCAAACAGG
## 27                                                           ACAGAGGGTGCAAGCGTTAATCGGAATTACTGGGCGTAAAGCGTGCGTAGACGGTTATCTAAGTCGGATGTGAAATCCCCGGGCTCAACCTGGGAATTGCATTCGAGACTGAATAGCTAGGGTGCGGAAGAGGGAAGCGGAATTTCCGGTGTAGCGGTGAAATGCGTAGATATCGGAAGGAACATCAGTGGCGAAAGCGGCTTCCTGGTCCAGCACCGACGTTCAGGCACGAAAGCGTGGGGAGCAAACAGG
## 28                                                           ACATAGGGTGCAAGCGTTGTCCGGAATTATTGGGCGTAAAGAGCTCGTAGGTGGTTCGATACGTCGGATGTGAAAATCAGGGGCTCAACCCCTGACCTGCATCCGATACGGTCGAGCTAGAGTTTGGTAGGGGAGACTGGAATTCCTGGTGTAGCGGTGGAATGCGCAGATATCAGGAGGAACACCGATGGCGAAGGCAGGTCTCTGGGCCAATACTGACACTGAGGAGCGAAAGCGTGGGGAGCGAACAGG
## 29                                                           ACGGAGGATGCAAGTGTTATCCGGAATCACTGGGCGTAAAGCGTCTGTAGGTGGTTTAATAAGTCAACTGTTAAATCTTGAGGCTCAACTTCAAAATCGCAGCTGAAACTATTAGACTAGAGTATAGTAGAGGTAAAGGGAATTTCCAGTGGAGCGGTGAAATGCGTAGATATTGGAAAGAACACCGATGGCGAAAGCACTTTACTGGGCTATTACTAACACTCAGAGACGAAAGCTAGGGTAGCAAATGGG
## 30                                                           ACGTAGGGTGCGAGCGTTAATCGGAATTACTGGGCGTAAAGCGTGCGCAGGCGGTTATACAAGACAGGCGTGAAATCCCCGGGCTTAACCTGGGAATGGCGCCTGTGACTGTATAGCTAGAGTGTGTCAGAGGGGGGTAGAATTCCACGTGTAGCAGTGAAATGCGTAGATATGTGGAGGAATACCAATGGCGAAGGCAGCCCCCTGGGATAACACTGACGCTCATGCACGAAAGCGTGGGGAGCAAACAGG
## 31                                                           ACGGAGGATCCAAGCGTTATCCGGATTTATTGGGTTTAAAGGGTCCGTAGGCGGGTCTTTAAGTCAGTGGTGAAAGCCCACAGCTCAACTGTGGAACTGCCATTGATACTGGAGATCTTGAGTGTAGTAGAAGTAGGCGGAATAGGGCATGTAGCGGTGAAATGCATAGATATGCCCTAGAACACCGATTGCGAAGGCAGCTTACTATGTTACAACTGACGCTGAGGGACGAAAGCGTGGGGAGCAAACAGG
## 32                                                           ACGGAGGATCCAAGCGTTATCCGGATTTATTGGGTTTAAAGGGAGCGTAGGTTGACATATAAGTCAGCTGTGAAAGTTTACGGCTCAACCGTGAAATTGCAGTTGATACTGTATGTCTTGAGTGTACAAGAGGTGGGCGGAATTCGTGGTGTAGCGGTGAAATGCTTAGATATCACGAAGAACTCCAATTGCGAAGGCAGCTCACTGGGGTACAACTGACACTGAGGCTCGAAAGTGTGGGTATCAAACAGG
## 33                                                           ACGGAGGGTGCAAGCGTTATCCGGATTCACTGGGTTTAAAGGGTGCGTAGGCGGGCAGGTAAGTCAGTGGTGAAATCCCCGAGCTTAACTCGGGAACTGCCGTTGATACTATCTGTCTTGAATACTGTGGAGGTAAGCGGAATATGTCATGTAGCGGTGAAATGCTTAGATATGACATAGAACACCAATTGCGAAGGCAGCTTGCTACACGGTCATTGACGCTGAGGCACGAAAGCGTGGGGATCAAACAGG
## 34                                                           ACGGAGGGTGCAAGCGTTAATCGGAATTACTGGGCGTAAAGCGCACGCAGGCGGTTGGATAAGTTAGATGTGAAAGCCCCGGGCTCAACCTGGGAATTGCATTTAAAACTGTCCAGCTAGAGTCTTGTAGAGGGGGGTAGAATTCCAGGTGTAGCGGTGAAATGCGTAGAGATCTGGAGGAATACCGGTGGCGAAGGCGGCCCCCTGGACAAAGACTGACGCTCAGGTGCGAAAGCGTGGGGAGCAAACAGG
## 35                                                           ACAGAGGATGCAAGCGTTATCCGGAATCACTGGGCATAAAGCGTCTGTAGGTGGTTTGGTAAGTCTGCTGTTAAAGACTGGGGCTCAACCCCAGAAAAGCAGTGGAAACTGCCAGACTTGAGTGTGGTAGAGGTAAAGGGAATTCCTAGTGTAGCGGTGAAATGCGTAGATATTAGGAAGAACACCAATGGCGAAGGCACTTTACTGGGCCATAACTGACACTGAGAGACGACAGCTAGGGGAGCAAATGGG
## 36                                                           ACGGAGGGTGCAAGCGTTAATCGGAATAACTGGGCGTAAAGCGCACGCAGGCGGTTAGATAAGTCAGATGTGAAATCCCCGGGCTCAACCTGGGAACTGCATTTGAAACTGTCTGACTAGAGTCTTGTAGAGGGGGGTAGAATTCCAGGTGTAGCGGTGAAATGCGTAGAGATCTGGAGGAATACCGGTGGCGAAGGCGGCCCCCTGGACAAAGACTGACGCTCAGGTGCGAAAGCGTGGGGAGCAAACAGG
## 37                                                           ACGTAGGGTGCAAGCGTTAATCGGAATTACTGGGCGTAAAGCGTGCGCAGGCGGTTATATAAGACAGATGTGAAATCCCCGGGCTCAACCTGGGAACTGCATTTGTGACTGTATAGCTAGAGTACGGCAGAGGGGGATGGAATTCCGCGTGTAGCAGTGAAATGCGTAGATATGCGGAGGAACACCGATGGCGAAGGCAATCCCCTGGGCCTGTACTGACGCTCATGCACGAAAGCGTGGGGAGCAAACAGG
## 38                                                           ACGGAGGGTGCAAGCGTTAATCGGAATAACTGGGCGTAAAGCGCACGCAGGCGGTTAGATAAGTCAGATGTGAAAGCCCCGGGCTCAACCTGGGAACTGCATTTGAAACTGTCTGACTAGAGTCTTGTAGAGGGGGGTAGAATTCCAGGTGTAGCGGTGAAATGCGTAGAGATCTGGAGGAATACCGGTGGCGAAGGCGGCCCCCTGGACAAAGACTGACGCTCAGGTGCGAAAGCGTGGGGAGCAAACAGG
## 39                                                           ACATAGGGTGCAAGCGTTGTCCGGAATTATTGGGCGTAAAGAGCTCGTAGGTGGTTCGATACGTCGGATGTGAAAATCAGGGGCTCAACCCCTGACCTGCATTCGATACGGTCGAGCTAGAGTTTGGTAGGGGAGACTGGAATTCCTGGTGTAGCGGTGGAATGCGCAGATATCAGGAGGAACACCGATGGCGAAGGCAGGTCTCTGGGCCAATACTGACACTGAGGAGCGAAAGCGTGGGGAGCGAACAGG
## 40                                                           ACGGAGGGTCCAAGCGTTAATCGGAATTACTGGGCGTAAAGCGCGCGTAGGCGGTTAATTAAGCGAGATGTGAAATCCCTGGGCTCAACCTAGGAATTGCGTTTCGAACTGATTAGCTAGAGTGCAGTAGAGGGTGGTGGAATTTCCGGTGTAGCGGTGAAATGCGTAGAGATCGGAAGGAACATCAGTGGCGAAGGCGACTGCCTGGACTGACACTGACGCTGAGGTGCGAAAGCGTGGGGAGCAAACAGG
## 41                                                           ACGTAGGGAGCAAGCGTTGTCCGGAATTACTGGGTGTAAAGGGAGCGTAGGCGGGACGGCAAGTCAGATGTGAAATTTATGGGCTCAACTCATAACCTGCATTTGAAACTGCTGTTCTTGAGTGAAGTAGAGGTAAGCGGAATTCCTGGTGTAGCGGTGAAATGCGTAGATATCAGGAGGAACATCGGTGGCGAAGGCGGCTTACTGGGCTTTTACTGACGCTGAGGCTCGAAAGCGTGGGGAGCAAACAGG
## 42                                                           ACGGAGGGTGCAAGCGTTATCCGGATTCACTGGGTTTAAAGGGTGCGTAGGCGGGCAGGTAAGTCAGTGGTGAAATCCTGGAGCTCAACTCCAGAACTGCCATTGATACTATCTGTCTTGAATATTGTGGAGGTAAGCGGAATATGTCATGTAGCGGTGAAATGCTTAGATATGACATAGAACACCCATTGCGAAGGCAGCTTACTACGCATATATTGACGCTGAGGCACGAAAGCGTGGGGATCAAACAGG
## 43                                                           ACGTAGGATCCGAGCGTTGTCCGGAGTTACTGGGTGTAAAGGGTGCGTAGGCTGTTCTGTGCGTCAGAGGTGAAATCCACGGGCTTAACTCGTGGGGTGCCTTTGATACGGCAGGGCTTGAGTACGAGAGAGGTGCGTGGAATTCCTGGTGTAGCAGTGAAATGCGTAGATATCAGGAGGAACACCGGTGGCGAAGGCGGCGCACTGGCTCGTAACTGACGCTGAGGCACGAAAGCGTGGGGATCAAACAGG
## 44                                                           ACGAAAGGTGCAAGCGTTGTTCGGATTGACTGGGCGTAAAGAGCACGTAGGTGGGTCTTTAAGTCGATTGTGAAATCCCGGAGCTCAACTTCGGACTGGCAATCGAAACTGTAGATCTTGAATACCCAAGGGGTTGGTGGAATTGCACATGTAGCAGTGAAATGCGTAGATATGTGCAGGAACACCGGAGGCGAAGGCGGCCAACTGGTGGTGTATTGACGCTGAGGTGCGAAAGCGCGGGGAGCAAACAGG
## 45                                                           ACGGAGGATCCGAGCGTTATCCGGATTTATTGGGTTTAAAGGGTGCGTAGGTGGTTTGATAAGTCAGCGGTGAAAGTTTGCAGCTTAACTGTAAAAATGCCGTTGAAACTGTCGGACTTGAGTGTAAATGAGGTAGGCGGAATGCGTGGTGTAGCGGTGAAATGCATAGATATCACGCAGAACTCCGATTGCGAAGGCAGCTTACTAAGCTACAACTGACACTGAAGCACGAAAGCGTGGGGATCAAACAGG
## 46                                                           ACGTAGGGTGCAAGCGTTAATCGGAATTACTGGGCGTAAAGCGTGCGCAGGCGGTTATGTAAGACAGGCGTGAAATCCCCGGGCTTAACCTGGGAATTGCGCTTGTGACTGCATAGCTAGAGTATGTCAGAGGGGGGTAGAATTCCACGTGTAGCAGTGAAATGCGTAGAGATGTGGAGGAATACCAATGGCGAAGGCAGCCCCCTGGGATAATACTGACGCTCATGCACGAAAGCGTGGGGAGCAAACAGG
## 47                                                           ACGGAGGGAGCTAGCGTTGTTCGGAATCACTGGGCGTAAAGCGTGCGTAGGCGGTGACTCAAGTCAGAGGTGAAAGCCTGGAGCTCAACTCCAGAACTGCCTTTGAAACTAGGTCGCTAGAACATCGGAGAGGTGAGTGGAATTCCGAGTGTAGAGGTGAAATTCGTAGATATTCGGAAGAACACCAGTGGCGAAGGCGGCTCACTGGACGATTGTTGACGCTGAGGCACGAAAGCGTGGGGAGCAAACAGG
## 48                                                           ACGTAGGGTGCAAGCGTTAATCGGAATTACTGGGCGTAAAGCGTGCGCAGGCGGTTGTGCAAGACAGGTGTGAAATCCCCGGGCTTAACCTGGGAACTGCACTTGTGACTGCACGGCTTGAGTGCGGCAGAGGGGGATGGAATTCCGCGTGTAGCAGTGAAATGCGTAGATATGCGGAGGAACACCGATGGCGAAGGCAATCCCCTGGGCCTGCACTGACGCTCATGCACGAAAGCGTGGGGAGCAAACAGG
## 49                                                           ACGAGTGCCCCGAGCGTTATCCGGAATCATTGGGCGTAAAGGGTGTGTAGGTGGTCTTGTTAGTCAATTGTTAAAACCCGGGGCTTAACCCCGGAAGTGCGATTGAAACGGCAAAACTAGAGGATGGCAGGGATGTATGGAACTCATGGAGTAGGGGTGAAATCCGTTGATATCATGGGGAACACCAAAAGCGAAGGCAGTACATTGGGCCACTCCTGACACTGAAACACGAAAGCGTGGGTAGCGAATGGG
## 50                                                           ACGTAGGGTGCAAGCGTTAATCGGAATTACTGGGCGTAAAGCGTGCGCAGGCGGTTATGCAAGACAGGTGTGAAATCCCCGAGCTCAACTTGGGAACTGCACTTGTGACTGCATAGCTAGAGTACGGTAGAGGGGAGTGGAATTCCGCGTGTAGCAGTGAAATGCGTAGATATGCGGAGGAACACCGATGGCGAAGGCAGCTCCCTGGACCTGTACTGACGCTCATGCACGAAAGCGTGGGGAGCAAACAGG
## 51                                                           ACGTAGGGTGCGAGCGTTAATCGGAATTACTGGGCGTAAAGCGTGCGCAGGCGGTTATGTAAGACAGATGTGAAATCCCCGGGCTCAACCTGGGAACTGCGTTTGTGACTGCATAACTAGAGTACGGCAGAGGGAGGTGGAATTCCGCGTGTAGCAGTGAAATGCGTAGATATGCGGAGGAACACCGATGGCGAAGGCAGCCTCCTGGGCCAGTACTGACGCTCATGCACGAAAGCGTGGGGAGCAAACAGG
## 52                                                           ACGGAGGGTGCAAGCGTTAATCGGAATTACTGGGCGTAAAGCGCGCGTAGGCGGCTAGTTAAGCTAGATGTGAAATCCCTGGGCTCAACCTGGGAACTGCATTTAGAACTGACTGGCTAGAGTGCAGAAGAGGAGTGTGGAATTTCCGGTGTAGCGGTGAAATGCGTAGAGATCGGAAGGAACATCAGTGGCGAAGGCGACACTCTGGTCTGACACTGACGCTGAGGTGCGAAAGCGTGGGTAGCAAACAGG
## 53                                                           ACGTAGGGTGCAAGCGTTAATCGGAATTACTGGGCGTAAAGCGTGCGCAGGCGGTTTTATAAGACAGAGGTGAAATCCCCGGGCTCAACCTGGGAACTGCCTTTGTGACTGTAAGGCTAGAGTGTGTCAGAGGGGGGTAGAATTCCACGTGTAGCAGTGAAATGCGTAGAGATGTGGAGGAATACCGATGGCGAAGGCAGCCCCCTGGGATAACACTGACGCTCATGCACGAAAGCGTGGGGAGCAAACAGG
## 54                                                           ACGGAGGGTGCAAGCGTTGTCCGGATTTATTGGGTTTAAAGGGTGCGTAGGCGGCTGAATAAGTCAGCGGTGAAAGACTTCGGCTTAACCGGAGCAGTGCCGTTGATACTGTTTGGCTTGAGTGTTGGAGGGGTACATGGAATTGATGGTGTAGCGGTGAAATGCATAGATACCATCAGGAACACCGATAGCGAAGGCATTGTACTGGCCAACAACTGACGCTGAGGCACGAAAGTGTGGGGATCGAACAGG
## 55                                                           ACGGAGGATGCAAGTGTTATCCGGAATCACTGGGCGTAAAGCGTCTGTAGGTGGTTTGATAAGTCAACTGTTAAATCTTGAGGCTCAACTTCAAAATCGCAGTCGAAACTATTAGACTAGAGTATAGTAGAGGTAAAGGGAATTTCCAGTGGAGCGGTGAAATGCGTAGATATTGGAAAGAACACCGATGGCGAAAGCACTTTACTGGGCTATTACTAACACTCAGAGACGAAAGCTAGGGTAGCAAATGGG
## 56                                                           ACGTAGGGTGCAAGCGTTAATCGGAATTACTGGGCGTAAAGCGTGCGCAGGCGGTTGATTAAGTTTGATGTGAAATCCCCGGGCTTAACCTGGGAACTGCATTGAATACTGGTCAGCTAGAGTGTGTCAGAGGGGGGTAGAATTCCACGTGTAGCAGTGAAATGCGTAGAGATGTGGAGGAATACCGATGGCGAAGGCAGCCCCCTGGGATAACACTGACGCTCATGCACGAAAGCGTGGGGAGCAAACAGG
## 57                                                           ACGGAGGGTGCAAGCGTTATCCGGATTCACTGGGTTTAAAGGGTGCGTAGGCGGGCAGGTAAGTCAGTGGTGAAATCCTGGAGCTCAACTCCAGAACTGCCATTGATACTATCTGTCTTGAATATTGTGGAGGTAAGCGGAATATGTCATGTAGCGGTGAAATGCTTAGATATGACATAGAACACCTATTGCGAAGGCAGCTTACTACGCATATATTGACGCTGAGGCACGAAAGCGTGGGGATCAAACAGG
## 58                                                           ACGTAGGGTGCGAGCGTTAATCGGAATTACTGGGCGTAAAGCGTGCGCAGGCGGTTGTGCAAGACAGATGTGAAATCCCCGGGCTTAACCTGGGAATTGCATTTGTGACTGCACGGCTAGAGTGTGTCAGAGGGGGGTAGAATTCCACGTGTAGCAGTGAAATGCGTAGATATGTGGAGGAATACCGATGGCGAAGGCAGCCCCCTGGGATAACACTGACGCTCATGCACGAAAGCGTGGGGAGCAAACAGG
## 59                                                           ACATAGGGTGCAAGCGTTGTCCGGAATTATTGGGCGTAAAGAGCTCGTAGGTTGTTTGTTACGTCGGATGTGAAAACCTGAGGCTCAACCTCAGGCCTGCATTCGATACGGGCAAACTAGAGTTTGGTAGGGGAGACTGGAATTCCTGGTGTAGCGGTGGAATGCGCAGATATCAGGAGGAACACCAATGGCGAAGGCAGGTCTCTGGGCCAATACTGACACTGAGGAGCGAAAGTCTGGGGAGCGAACAGG
## 60                                                           ACGAAGGTGGCAAGCGTTGTTCGGATTCACTGGGCGTACAGGGTGTGTAGGCGGTTTGGTAAGCCTTCTGTTAAAGCTTCGGGCCCAACCCGGAAAGCGCAGAGGGTACTGCCAGGCTAGAGGGTGGGAGAGGAGCGCGGAATTCCCGGTGTAGCGGTGAAATGCGTAGAGATCGGGAGGAAGGCCGGTGGCGAAGGCGGCGCTCTGGAACATACCTGACGCTGAGACACGAAAGCGTGGGGAGCAAACAGG
## 61                                                           ACGTAGGGTGCGAGCGTTAATCGGAATTACTGGGCGTAAAGCGTGCGCAGGCGGTTTTGTAAGACAGATGTGAAATCCCCGGGCTTAACCTGGGAACTGCATTTGTGACTGCAAGACTAGAGTGTGTCAGAGGGAGGTGGAATTCCGCGTGTAGCAGTGAAATGCGTAGAGATGCGGAAGAACACCGATGGCGAAGGCAGCCTCCTGGGATAACACTGACGCTCAGGCACGAAAGCGTGGGGAGCAAACAGG
## 62                                                           ACAGAGGATGCAAGCGTTATCCGGAATCACTGGGCATAAAGCGTCTGTAGGTTGCTTGCCAAGTCTGCTGTTAAAGACTAGGGCCTAACCCTGGGAAAGCAGTGGAAACTAGCAGGCTTGAGTGTGGTAGAGGTAGAGGGAATTCCTGGTGTAGCGGTGAAATGCGTAGATATTAGGAAGAACACCAATGGCGAAAGCACTCTGCTGGGCCACAACTGACACTGAGAGACGACAGCTAGGGGAGCAAATGGG
## 63                                                           ACATAGGGTGCAAGCGTTGTCCGGAATTATTGGGCGTAAAGAGCTCGTAGGTGGTTCGATACGTCGGATGTGAAAATCAGGGGCTCAACCCCTGACCTGCATTCGATACGGTCGAGCTAGAGTGTGGTAGGGGAGACTGGAATTCCTGGTGTAGCGGTGGAATGCGCAGATATCAGGAGGAACACCGATGGCGAAGGCAGGTCTCTGGGCCATTACTGACACTGAGGAGCGAAAGCGTGGGGAGCGAACAGG
## 64                                                           ACGTAGGGTGCGAGCGTTAATCGGAATTACTGGGCGTAAAGCGTGCGCAGGCGGTTTTGTAAGACAGGCGTGAAATCCCCGGGCTCAACCTGGGAACTGCGCTTGTGACTGCAAGGCTAGAGTACGGCAGAGGGGGGTGGAATTCCACGTGTAGCAGTGAAATGCGTAGAGATGTGGAGGAACACCAATGGCGAAGGCAGCCCCCTGGGTCGATACTGACGCTCATGCACGAAAGCGTGGGTAGCAAACAGG
## 65                                                          ACGGAGGATCCAAGCGTTATCCGGAATTACTGGGCGTAAAGAGTTGCGTAGGTGGCATTGTAAGTCTGTAGTGAAAGCGTGGGGCTCAACCCCATACACATTATGGAAACTGCAAAGCTAGAGGACAAGAGAGGTTATTGGAATTCCTAGTGTAGGAGTGAAATCCGTAGATATTAGGAGGAACACCGATGGCGTAGGCAGATAACTGGCTTGTTCCTGACACTAAGGCACGAAAGCATGGGTAGCAAACGGG
## 66                                                           ACGGAGGGTGCAAGCGTTATCCGGAATCATTGGGTTTAAAGGGTCCGCAGGCGGTCTGATAAGTCAGTGGTGAAAGCCTATTGCTCAACAATAGAACTGCCATTGATACTGTTAGACTTGAATTAGGTCGGAGTGGGCGGAATATGACATGTAGCGGTGAAATGCTTAGATATGTCATAGAACACCGATAGCGAAGGCAGCTCACTAGCCCTAAATTGACGCTCATGGACGAAAGCGTGGGGAGCAAACAGG
## 67                                                          ACGAGTGCTTCAAGCGTTATCCGGAATCATTGGGCGTAAAGGGTGTGTAGGCGGTTATGTTAGTCTCTCGTCAAATCTTTCGGCTCAACCGAGAGTCCGCGAGGGAAACGGCATGACTAAGAGGTTGGAAGAGGTCTCTGGAACTCATGGTGTAGCGGTGAAATGCGTTGATATCATGGGGAACACCAAAAGCGAAGGCAAGAGACTGGTCCATACCTGACGCTGAAACACGAAAGCGTGGGTCGCGAATGGG
## 68                                                           ACGGAGGATGCAAGTGTTATCCGGAATCACTGGGCGTAAAGCGTCTGTAGGTGGTTTAATAAGTCAACTGTTAAATCTTGAGGCTTAACTTCAAAATCGCAGTCGAAACTATTAGACTAGAGTATAGTAGGGGTAAAGGGAATTTCCAGTGGAGCGGTGAAATGCGTAGATATTGGAAAGAACACCGATGGCGAAAGCACTTTACTGGGCTATTACTAACACTCAGAGACGAAAGCTAGGGTAGCAAATGGG
## 69                                                           ACGTAGGGTGCAAGCGTTAATCGGAATTACTGGGCGTAAAGCGTGCGCAGGCGGTTATGTAAGACAGATGTGAAATCCCCGGGCTCAACCTGGGAACTGCATTTGTGACTGCATAGCTAGAGTACGGTAGAGGGGGATGGAATTCCGCGTGTAGCAGTGAAATGCGTAGATATGCGGAGGAACACCGATGGCGAAGGCAATCCCCTGGACCTGTACTGACGCTCATGCACGAAAGCGTGGGGAGCAAACAGG
## 70                                                           ACGGAGGGTGCAAGCGTTATCCGGATTCACTGGGTTTAAAGGGTGCGTAGGCGGGCTAGTAAGTCAGTGGTGAAATCCCCGAGCTTAACTTGGGAACTGCCGTTGATACTATTAGTCTTGAATATCGTGGAGGTAAGCGGAATATGTCATGTAGCGGTGAAATGCTTAGATATGACATAGAACACCAATTGCGAAGGCAGCTTGCTACACGATTATTGACGCTGAGGCACGAAAGCGTGGGGATCAAACAGG
## 71                                                           ACGTAGGGTGCGAGCGTTAATCGGAATTACTGGGCGTAAAGCGTGCGCAGGCGGTTACATAAGACAGATGTGAAATCCCCGGGCTTAACCTGGGAACTGCGTTTGTGACTGTGTGACTAGAGTACGGTAGAGGGGGGTGGAATTCCTGGTGTAGCAGTGAAATGCGTAGATATCAGGAGGAACATCGATGGCGAAGGCAGCCCCCTGGACCTGTACTGACGCTCATGCACGAAAGCGTGGGGAGCAAACAGG
## 72                                                           ACGTAGGGTGCAAGCGTTAATCGGAATTACTGGGCGTAAAGCGTGCGCAGGCGGTTTCGTAAGACAGAGGTGAAATCCCCGGGCTCAACCTGGGAACTGCCTTTGTGACTGCGAGGCTAGAGTATGGCAGAGGGGGGTGGAATTCCACGTGTAGCAGTGAAATGCGTAGAGATGTGGAGGAACACCGATGGCGAAGGCAGCCCCCTGGGCCAATACTGACGCTCATGCACGAAAGCGTGGGGAGCAAACAGG
## 73                                                           ACGTAGGGTGCGAGCGTTAATCGGAATTACTGGGCGTAAAGCGTGCGCAGGCGGTTTTGTAAGTCAGATGTGAAATCCCCGGGCTCAACCTGGGAACTGCGTTTGAAACTACAAGGCTAGAGTATGGTAGAGGGGGGTAGAATTCCACGTGTAGCAGTGAAATGCGTAGAGATGTGGAGGAATACCAATGGCGAAGGCAGCCCCCTGGACTAATACTGACGCTCATGCACGAAAGCGTGGGGAGCAAACAGG
## 74                                                           ACGGAGGGTGCGAGCGTTATCCGGATTCACTGGGTTTAAAGGGTGCGTAGGCGGGCTAGTAAGTCAGTGGTGAAATCCCCGAGCTTAACTTGGGAACTGCCGTTGATACTATTAGTCTTGAATATCGTGGAGGTAAGCGGAATATGTCATGTAGCGGTGAAATGCTTAGATATGACATAGAACACCAATTGCGAAGGCAGCTTGCTACACGATTATTGACGCTGAGGCACGAAAGCGTGGGGATCAAACAGG
## 75                                                           ACGTAGGGTGCAAGCGTTAATCGGAATTACTGGGCGTAAAGCGTGCGCAGGCGGTTATGCAAGACAGAGGTGAAATCCCCGGGCTCAACCTGGGAACTGCCTTTGTGACTGCATGACTAGAGTACGGCAGAGGGGGATGGAATTCCGCGTGTAGCAGTGAAATGCGTAGATATGCGGAGGAACACCGATGGCGAAGGCAATCCCCTGGGCCTGTACTGACGCTCATGCACGAAAGCGTGGGGAGCAAACAGG
## 76                                                           ACAGAGAGGGCAAGCGTTGTCCGGAATGACTGGGCGTAAAGCGCGCGCAGGCGGTTTTTTAAGTTGGATGTGAAATCCCCGGGCTTAACCTGGGAACTGCATACGATACTGGGAAACTGGAGGCCGAGAGAGGAAAGCGGAATGACGAGTGTAGAGGTGAAATTCGTAGATATTCGTCGGAACACCGGAGGCGAAGGCGGCTTTCTGGCTCGGACCTGACGCTGAGGCGCGAAAGCGTGGGGAGCAAACAGG
## 77                                                           ACGTAGGGTGCAAGCGTTAATCGGAATTACTGGGCGTAAAGCGTGCGCAGGCGGTTATGTAAGACAGAGGTGAAATCCCCGGGCTCAACCTGGGAACGGCCTTTGTGACTGCATAGCTAGAGTACGGTAGAGGGGGATGGAATTCCGCGTGTAGCAGTGAAATGCGTAGATATGCGGAGGAACACCGATGGCGAAGGCAATCCCCTGGACCTGTACTGACGCTCATGCACGAAAGCGTGGGGAGCAAACAGG
## 78                                                           ACGTAGGGTGCGAGCGTTAATCGGAATTACTGGGCGTAAAGCGTGCGCAGGCGGTCTTGTAAGACAGATGTGAAATCCCCGGGCTCAACCTGGGACCTGCATTTGTGACTGCAAGGCTGGAGTACGGCAGAGGGGGATGGAATTCCGCGTGTAGCAGTGAAATGCGTAGATATGCGGAGGAACACCGATGGCGAAGGCAATCCCCTGGGCCTGTACTGACGCTCATGCACGAAAGCGTGGGGAGCAAACAGG
## 79                                                           ACGGAGGATCCAAGCGTTATCCGGAATCATTGGGTTTAAAGGGTCCGTAGGCGGTTTATTAAGTCAGTGGTGAAATCTGGTCGCTCAACGATCAAACGGCCATTGATACTGATAGACTTGAATTACTGGGAAGTAACTAGAATATGTAGTGTAGCGGTGAAATGCTTAGATATTACATGGAATACCAATTGCGAAGGCAGGTTACTACCAGTGGATTGACGCTGATGGACGAAAGCGTGGGGAGCGAACAGG
## 80                                                          ACGAACCAAGCGAACGTTATTCGGAATTACTGGGCTTAAAGCGCGTGTAGGCGGACTGCCACGTCGGTTGTTGAAATCCCCCGGCTTAACCGGGGAACAGGCACCGATACGAGTAGTCTTGAGGTGGGTAGGGGAGACTGGAACTTCCGGTGGAGCGGTGAAATGCGTTGAGATCGGAAGGAACGCCCGCGGCGAAAGCGAGTCTCTGGCCCCATACTGACGCTGAGACGCGAAAGCCAGGGGAGCAAACGGG
## 81                                                           ACGGAGGGTGCAAGCGTTATCCGGAATCATTGGGTTTAAAGGGTCCGCAGGCGGATTTATAAGTCAGTGGTGAAATCCTATCGCTTAACGATAGAACTGCCATTGATACTGTAAGTCTTGAATTCGGTCGGAGTGGGCGGAATGTGTAGTGTAGCGGTGAAATGCATAGATATTACACAGAACACCGATAGCGAAGGCAGCTCACTAGGCCTGAATTGACGCTCAGGGACGAAAGCGTGGGGATCAAACAGG
## 82                                                           ACGGAGGATCCGAGCGTTATCCGGATTTATTGGGTTTAAAGGGTGCGTAGGCGGCCTATTAAGTCAGGGGTGAAAACTTGCAGCTTAACTGTAAGCGTGCCTTTGATACTGATGGGCTTGAATTAAGATGAAGTAGGCGGAATGTGACAAGTAGCGGTGAAATGCATAGATATGTCACAGAACACCAATTGCGAAGGCAGCTTACTAAGGTTAAATTGACGCTGAGGCACGAAAGCGTGGGGATCAAACAGG
## 83                                                          ACGTAGGGGGCGAGCGTTGTCCGAAGTTACTGGGCGTAAAGCGCGCGTAGGCGGTTATTCAAGTCTGGGGTGAAAGGTTCAGCGCTCAACGTGAACAGTGCCTTGGATACTGGATGACTTGAGTCTGGAAGGGGGAAGGGGAATTCCTGGTGTAGCGGTGAAATGCGTAGATATCAGGAGGAACACCGGTGGCGAAGGCGCCTTCCTGGTCTAAGACTGACGCTGAGGTGCGAAAGCGTGGGGAGCAAACGGG
## 84                                                           ACGAGTGCCTCAAGCGTTATCCGGAATCATTGGGCGTAAAGGTTGTGTAGGTGGTTATATTAGTCTCTTGTTAAATTCTTCGGCTTAACCGGGGACATGCAAGAGAAACGGTATGACTAGAGGATGCGAGGGGTCTGCGGAATTCATAGTGTAGCGGTAAAATGCGTTGATATTATGGAGAACACCAAAAGCGAAGGCAGCAGACTGGAGCACTCCTGACACTGAAACAAGAAAGCGTGGGTCGCGAATGGG
## 85                                                           ACAGAGACCTCAAGCGTTATCCGGATTTATTGGGCGTAAAGCGTCCGCAGGTGGTTCGTTAAGTCAATGGTCAAATCACGGGGCTCAACCCCGCAACTGCCGTTGATACTGGCGAACTTGAGGCCGGTAGAGGTAAGCGGAATTACAGGTGTAGCGGTAAAATGCGTTTATATCTGTAAGAACACCAAAGGCGAAAGCAGCTTACTGGAACGGTCCTGACACTCATGGACGAAAGCGTGGGGAGCGAAAAGG
## 86                                                           ACATAGGGTGCAAGCGTTGTCCGGAATTATTGGGCGTAAAGAGCTCGTAGGTGGTTCGATACGTCGGATGTGAAAATCAGGGGCTCAACCCCTGACCTGCATTCGATACGGTCGAGCTGGAGTTTGGTAGGGGAGACTGGAATTCCTGGTGTAGCGGTGGAATGCGCAGATATCAGGAGGAACACCGATGGCGAAGGCAGGTCTCTGGGCCAATACTGACACTGAGGAGCGAAAGCGTGGGGAGCGAACAGG
## 87                                                           ACGAGGGGTGCAAACGTTATTCGGAATTACTGGGTGTAAAGGGTACGTAGACGGCATCTTAAGTCAATTGTTAAATTCCCCGGCCTAACTGGGGGTAGGCGATTGAAACTAAGAAGCTAGAGAGTGGAAGAGAGAAGTGGAATTCTCGGAGTAGCGGTAAAATGCGTAGATCTCGAGAGGAACACCGATGGCGAAGGCAGCTTCTTGGTCCATATCTGACGTTGAGGTACGAAAGCGTGGGGAGCAAACAGG
## 88                                                           ACGGAGGGTGCAAGCGTTGTCCGGATTTATTGGGTTTAAAGGGTGCGTAGGCGGCTGAATAAGTCAGCGGTGAAAGACTTCGGCTTAACCGGAGCAGTGCCGTTGATACTGTTTAGCTTGAGTGTTGGAGGGGTACATGGAATTGATGGTGTAGCGGTGAAATGCATAGATACCATCAGGAACACCGATAGCGAAGGCATTGTACTGGCCAACAACTGACGCTGAGGCACGAAAGTGTGGGGATCGAACAGG
## 89                                                           ACGGAGGGTGCGAGCGTTAATCGGAATTACTGGGCGTAAAGCGCGCGTAGGCGGTTGGCTAAGTTTGCTGTGAAAGCCCCGGGCTTAACCTGGGAACTGCAGTGAATACTGGTCAGCTAGAGTATGGTAGAGGGTAGTGGAATTTCCGGTGTAGCAGTGAAATGCGTAGAGATCGGAAGGAACACCAGTGGCGAAGGCGGCTATCTGGACCAATACTGACGCTGAGGTGCGAAAGCGTGGGGAGCAAACAGG
## 90                                                           ACGTAGGGTGCAAGCGTTAATCGGAATTACTGGGCGTAAAGCGTGCGCAGGCGGTTTTGTAAGACAGAGGTGAAATCCCCGGGCTCAACCTGGGAACTGCCTTTGTGACTGCAAGGCTGGAGTACGGCAGAGGGGGATGGAATTCCGCGTGTAGCAGTGAAATGCGTAGATATGCGGAGGAACACCGATGGCGAAGGCAATCCCCTGGGCCTGTACTGACGCTCATGCACGAAAGCGTGGGGAGCAAACAGG
## 91                                                           ACGTAGGGTGCAAGCGTTAATCGGAATTACTGGGCGTAAAGGGTGCGCAGGCGGTATTGTAAGCCAGATGTGAAATCCCCGGGCTCAACCTGGGAACTGCGTTTGGAACTGCAATGCTAGAGTGTGTCAGAGGGGGGTGGAATTCCACGTGTAGCAGTGAAATGCGTAGAGATGTGGAGGAACACCGATGGCGAAGGCAGCCCCCTGGGATAACACTGACGCTCAGGCACGAAAGCGTGGGGAGCAAACAGG
## 92                                                           ACGTAGGGTGCAAGCGTTAATCGGAATTACTGGGCGTAAAGCGTGCGCAGGCGGTTATATAAGACAGATGTGAAATCCCCGGGCTCAACCTGGGAACTGCATTTGTGACTGTATAGCTAGAGTGCGGCAGAGGGGGATGGAATTCCGCGTGTAGCAGTGAAATGCGTAGATATGCGGAGGAACACCGATGGCGAAGGCAATCCCCTGGGCCTGCACTGACGCTCATGCACGAAAGCGTGGGGAGCAAACAGG
## 93                                                           ACGGAGGGTGCAAGCGTTAATCGGAATTACTGGGCGTAAAGCGCACGTAGGCTGTAGTGTAAGTCAGGGGTGAAATCCCACGGCTCAACCGTGGAACTGCCTTTGATACTGCACAACTTGAATCCGGGAGAGGGTGGCGGAATTCCAGGTGTAGGAGTGAAATCCGTAGATATCTGGAGGAACATCAGTGGCGAAGGCGGCCACCTGGACCGGTATTGACGCTGAGGTGCGAAAGCGTGGGGAGCAAACAGG
## 94                                                           ACGTAGGGTGCGAGCGTTAATCGGAATTACTGGGCGTAAAGCGTGCGCAGGCGGTTGTGCAAGACAGATGTGAAATCCCCGGGCTCAACCTGGGAATTGCATTTGTGACTGCACGGCTAGAGTGTGTCAGAGGGGGGTAGAATTCCACGTGTAGCAGTGAAATGCGTAGATATGTGGAGGAATACCGATGGCGAAGGCAGCCCCCTGGGATAACACTGACGCTCATGCACGAAAGCGTGGGGAGCAAACAGG
## 95                                                           ACGAAGGATCCAAGCGTTGTCCGGATTTACTGGGTTTAAAGGGTGCGTAGGCGGAAAATTAAGTCAGTGGTGAAAGCCCGCAGCTCAACTGTGGAACTGCCATTGAAACTGATTTTCTTGAATATAGCTGAGGCAGATGGAATATAACATGTAGCGGTGAAATGCTTAGATATGTTATAGAACACCGATTGCGAAGGCAGTCTGCTAAACTATTATTGACGCTGAGGCACGAAAGCGTGGGGAGCGAACAGG
## 96                                                           ACGTAGGGTGCAAGCGTTAATCGGAATTACTGGGCGTAAAGCGTGCGCAGGTGGTTCGTTAAGACAGTTGTGAAATCCCCGGGCTTAACCTGGGAACTGCAATTGTGACTGGCGGACTAGAGTTTGGCAGAGGGGGGTGGAATTCCTGGTGTAGCAGTGAAATGCGTAGATATCAGGAGGAACACCGATGGCGAAGGCAGCCCCCTGGGCCATGACTGACACTCATGCACGAAAGCGTGGGGAGCAAACAGG
## 97                                                           ACGTAGGGTGCGAGCGTTAATCGGAATTACTGGGCGTAAAGCGTGCGCAGGCGGACTTTTAAGCCAGATGTGAAAGCCCCGAGCTTAACTTGGGAATTGCGTTTGGAACTGGGAGTCTAGAGTCTGTCAGAGGGGGATGGAATTCCACGTGTAGCAGTGAAATGCGTAGAGATGTGGAGGAACACCGATGGCGAAGGCAGTCCCCTGGGATAAGACTGACGCTCATGCACGAAAGCGTGGGGAGCAAACAGG
## 98                                                           ACATAGGGTGCAAGCGTTGTCCGGAATTATTGGGCGTAAAGAGCTCGTAGGTTGTTTGTCGCGTCGATTGTGAAAATCTGAGGCTCAACCTCAGACCTGCAGTCGATACGGGCAAACTAGAGTGTGGTAGGGGAGACTGGAATTCCTGGTGTAGCGGTGGAATGCGCAGATATCAGGAGGAACACCAATGGCGAAGGCAGGTCTCTGGGCCATAACTGACACTGAGGAGCGAAAGTGCGGGGAGCGAACAGG
## 99                                                           ACGGAGGGTGCAAGCGTTATCCGGATTTATTGGGTTTAAAGGGTCCGTAGGCGGATCTGTAAGTCAGTGGTGAAATCTCACAGCTTAACTGTGAAACTGCCATTGATACTGCAGGTCTTGAGTGTTGTTGAAGTAGCTGGAATAAGTAGTGTAGCGGTGAAATGCATAGATATTACTTAGAACACCAATTGCGAAGGCAGGTTACTAAGCAACAACTGACGCTGATGGACGAAAGCGTGGGGAGCGAACAGG
## 100                                                          ACGAAGGGGGCTAGCGTTGCTCGGAATTACTGGGCGTAAAGGGAGCGTAGGCGGATAGTTTAGTCAGAGGTGAAAGCCCAGGGCTCAACCTTGGAAATGCCTTTGATACTGGCTATCTTGAGTATGGGAGAGGTAAGCGGAATTCCGAGTGTAGAGGTGAAATTCGTAGATATTCGGAAGAACACCAGTGGCGAAGGCGGCTTACTGGCCCATTACTGACGCTGAGGCTCGAAAGCGTGGGGAGCAAACAGG
## 101                                                          ACGTAGGGTGCAAGCGTTAATCGGAATTACTGGGCGTAAAGCGTGCGCAGGCGGTTTGTTAAGACAGTCGTGAAATCCCCGGGCTTAACCTGGGAACTGCGATTGTGACTGGCAAGCTAGAGTACGGCAGAGGGGGGTGGAATTCCTGGTGTAGCAGTGAAATGCGTAGAGATCAGGAGGAACACCGATGGCGAAGGCAGCCCCCTGGGCTAGTACTGACGCTCATGCACGAAAGCGTGGGGAGCAAACAGG
## 102                                                          ACGTAGGGTGCAAGCGTTAATCGGAATTACTGGGCGTAAAGCGTGCGCAGGCGGTTATATAAGACAGATGTGAAATCCCCGGGCTCAACCTGGGAACTGCATTTGTGACTGTATAGCTGGAGTGCGGCAGAGGGGGATGGAATTCCGCGTGTAGCAGTGAAATGCGTAGATATGCGGAGGAACACCGATGGCGAAGGCAATCCCCTGGGCCTGCACTGACGCTCATGCACGAAAGCGTGGGGAGCAAACAGG
## 103                                                          ACGGAGGATCCGAGCGTTATCCGGATTTATTGGGTTTAAAGGGAGCGTAGGTTGACGTATAAGTCAGCTGTGAAAGTTTACGGCTCAACCGTGAAATTGCAGTTGATACTGTATGTCTTGAGTGTACAAGAGGTGGGCGGAATTCGTGGTGTAGCGGTGAAATGCTTAGATATCACGAAGAACTCCAATTGCGAAGGCAGCTCACTGGGGTACAACTGACACTGAGGCTCGAAAGTGTGGGTATCAAACAGG
## 104                                                          ACGGAGGATCCAAGCGTTATCCGGAATCATTGGGTTTAAAGGGTCCGTAGGCGGTCTTATAAGTCAGTGGTGAAATCCCATCGCTCAACGATGGAACTGCCATTGATACTGTAAGACTTGAATGCTTAGGAAGTAACTAGAATATGTAGTGTAGCGGTGAAATGCTTAGATATTACATGGAATACCAATTGCGAAGGCAGGTTACTACTAAGTGATTGACGCTGATGGACGAAAGCGTGGGGAGCGAACAGG
## 105                                                        ACGTAGGGTACAAGCATTAATCGGAATGATTGGGCGTAAAGGGCGCGTAGGCGGCCTTGTAAGTCAGGTGTGAAATTCCAAAGCTCAACTTTGGAGCTGCACTTGAAACTACAAGGCTAAAGAGGTAAGACGGAGAAAGCGGAATTCCACGTGTAGCGGTGAAATGCGTAGATATGTGGAGGAACACCTGTGGCGAAAGCGGCTTTCTAGTTTTAACCTGACGCTGAGGCGCGAGAGCAAGGGGAGCAAACAGG
## 106                                                          ACGTAGGTGACAAGCGTTATCCGGATTTACTGGGTGTAAAGGGCGTGTAGGCGGTTTTGCAAGTCAGATGTGAAATTCCCGGGCTCAACTCGGGCGCTGCATCTGAAACTGTAAAACTTGAGTACTGGAGAGGATAGCGGAATTCCTAGTGTAGCGGTAAAATGCGCAGATATTAGGAGGAACACCGGTGGCGAAGGCGGCTATCTGGACAGTAACTGACGCTGAGGCGCGAAAGCGTGGGGAGCAAACAGG
## 107                                                          ACGAAGGGTGCGAGCGTTGTTCGGAATTACTGGGCGTAAAGGGTTCGTAGGCGGGAATGCAAGTCAAGTGTGAAATCCCCAGGCTTAACCTGGGACGTGCATTTGAGACTGTGTTTCTTGAGTTTCGGAGAGGGTGGTGGAATTGCTGGTGTAGGAGTGACATCCGTAGAGATCAGCAGGAACACCGGAGGCGAAGGCGACCACCTGGCCGGATACTGACGCTGAGGAACGAAAGCGTGGGGAGCAAACAGG
## 108                                                          ACGGAGGATCCGAGCGTTATCCGGATTTATTGGGTTTAAAGGGAGCGTAGATGGATGTTTAAGTCAGTTGTGAAAGTTTGCGGCTCAACCGTAAAATTGCAGTTGATACTGGATATCTTGAGTGCAGTTGAGGCAGGCGGAATTCGTGGTGTAGCGGTGAAATGCTTAGATATCACGAAGAACTCCGATTGCGAAGGCAGCCTGCTAAGCTGCAACTGACATTGAGGCTCGAAAGTGTGGGTATCAAACAGG
## 109                                                          ACGTAGGGTGCGAGCGTTAATCGGAATTACTGGGCGTAAAGCGTGCGCAGGTGGTTGTTTAAGACAGATGTGAAATCCCCGGGCTTAACCTGGGAACTGCGTTTGTGACTGGACAGCTAGAGTGCGGCAGAGGGGGGTGGAATTCCACGTGTAGCAGTGAAATGCGTAGAGATGTGGAGGAACACCGATGGCGAAGGCAGCCCCCTGGGCCAGCACTGACGCTCATGCACGAAAGCGTGGGGAGCAAACAGG
## 110                                                         ACGTAGGCTCCAAGCGTTGTCCGGATTTATTGGGCGTAAAGAGCTCGTAGGCGGTTGAGTAAGTCGGGTGTGAAAACTCTGGGCTTAACCCGGAGACGCCATCCGATACTGCTCTGACTAGAGTTCAGGAGGGGAGTGGGGAATTCCTAGTGTAGCGGTGAAATGCGCAGATATTAGGAGGAACACCGGTGGCGAAGGCGCCACTCTGGACTGAAACTGACGCTGAGGAGCGAAAGCATGGGTAGCAAACAGG
## 111                                                          ACGGAGGATGCAAGTGTTATCCGGAATCACTGGGCGTAAAGCGTCTGTAGGTGGTTTAATAAGTCAACTGTTAAATCTTGAGGCTCAACTTCAAAATCGCAGTCGAAACTATTAGACTAGAGTATAGTAGAGGTAAAGGGAATTTCCAGTGGAGCGGTGAAATGCGTAGATATTGGAAAGAACACCGATGGCGAAAGCACTTTACTGGGCTATTACTAACACTCAGAGACGAAAGCTAGGGTAGCAAATGGG
## 112                                                          ACGTAGGGTGCGAGCGTTAATCGGAATTACTGGGCGTAAAGCGTGCGCAGGCGGTTATATAAGACAGTTGTGAAATCCCCGGGCTCAACCTGGGAATTGCATCTGTGACTGTATAGCTAGAGTACGGTAGAGGGGGATGGAATTCCGCGTGTAGCAGTGAAATGCGTAGATATGCGGAGGAACACCGATGGCGAAGGCAATCCCCTGGACCTGTACTGACGCTCATGCACGAAAGCGTGGGGAGCAAACAGG
## 113                                                          ACGTAGGGTGCAAGCGTTAATCGGAATTACTGGGCGTAAAGCGTGCGCAGGCGGTTATACAAGACAGGCGTGAAATCCCCGGGCTTAACCTGGGAATGGCGTCTGTGACTGTATGACTAGAGTGTGTCAGAGGGGGGTAGAATTCCACGTGTAGCAGTGAAATGCGTAGATATGTGGAGGAATACCAATGGCGAAGGCAGCCCCCTGGGATAACACTGACGCTCATGCACGAAAGCGTGGGGAGCAAACAGG
## 114                                                          ACGAAGGGTGCAAGCGTTAATCGGAATTACTGGGCGTAAAGCGCGCGTAGGCGGTTTGGTTAGTTGGATGTGAAAGCCCCGGGCTCAACCTGGGAACTGCATCCAATACTGCCAAGCTAGAGTACGGTAGAGGGGGGTAGAATTCCACGTGTAGCGGTGAAATGCGTAGAGATGTGGAGGAATACCAGTGGCGAAGGCGGCCCCCTGGATCGATACTGACGCTGAGGTGCGAAAGCGTGGGGAGCAAACAGG
## 115                                                          ACGTAGGGTGCGAGCGTTAATCGGAATTACTGGGCGTAAAGCGTGCGCAGGCGGCTTTGCAAGACAGAGGTGAAATCCCCGGGCTCAACCTGGGAACTGCCTTTGTGACTGCAAGGCTAGAGTACGGCAGAGGGAGATGGAATTCCGCGTGTAGCAGTGAAATGCGTAGATATGCGGAGGAACACCGATGGCGAAGGCAATCTCCTGGGCCTGTACTGACGCTCATGCACGAAAGCGTGGGGAGCAAACAGG
## 116                                                          ACGTAGGGTGCAAGCGTTAATCGGAATTACTGGGCGTAAAGCGTGCGCAGGCGGTTTGAAAAGTCAGCTGTGAAAGCCCCGGGCTCAACCTGGGAACTGCGGTTGAAACTCTCAAGCTAGAGTGCGTCAGAGGGGGGTGGAATTCCACGTGTAGCAGTGAAATGCGTAGAGATGTGGAGGAACACCAATGGCGAAGGCAGCCCCCTGGGATGACACTGACGCTCATGCACGAAAGCGTGGGGAGCAAACAGG
## 117                                                          ACGTAGGGTGCAAGCGTTAATCGGAATTACTGGGCGTAAAGCGTGCGCAGGCGGTTATGTAAGACAGAGGTGAAATCCCCGGGCTCAACCTGGGAACTGCCTTTGTGACTGCGAGGCTAGAGTATGGCAGAGGGGGGTGGAATTCCACGTGTAGCAGTGAAATGCGTAGAGATGTGGAGGAACACCGATGGCGAAGGCAGCCCCCTGGGCCAATACTGACGCTCATGCACGAAAGCGTGGGGAGCAAACAGG
## 118                                                          ACAGAGGTCTCAAGCGTTGTTCGGAATCACTGGGCGTAAAGGGTGCGTAGGTTGCGTGGTAAGTCAGATGTGAAAGCCTGGGGCTCAACCTCAGAATTGCATCCGATACTGCCGTGCTAGAGTACTGAAGAGGTGACTAGAATTCTAGGTGTAGCAGTGAAATGCGTAGATATCTAGAGGAATACCAAAGGCGTAGGCAGGTCACTGGGCAGTTACTGACACTGAGGCACGAAGGCCAGGGGAGCAAACGGG
## 119                                                          ACGGAGGGTGCAAGCGTTATCCGGATTTACTGGGTTTAAAGGGTGCGTAGGCGGACAGGTAAGTCAGTGGTGAAATCCCCGAGCTTAACTCGGGAACTGCCGTTGATACTATCTGTCTTGAATATCGTGGAGGTAAGCGGAATATGTCATGTAGCGGTGAAATGCTTAGATATGACATAGAACACCAATTGCGAAGGCAGCTTGCTACACGGTCATTGACGCTGAGGCACGAAAGCGTGGGGATCAAACAGG
## 120                                                          ACGTAGGGTGCAAGCGTTAATCGGAATTACTGGGCGTAAAGCGTGCGCAGGCGGTTATGCAAGACAGATGTGAAATCCCCGGGCTCAACCTGGGAACTGCATTTGTGACTGCATAGCTAGAGTACGGTAGAGGGGGATGGAATTCCGCGTGTAGCAGTGAAATGCGTAGATATGCGGAGGAACACCGATGGCGAAGGCAATCCCCTGGACCTGTACTGACGCTCATGCACGAAAGCGTGGGGAGCAAACAGG
## 121                                                          ACGTAGGGTGCGAGCGTTAATCGGAATTACTGGGCGTAAAGCGTGCGCAGGCGGTTATGTAAGACAGATGTGAAATCCCCGGGCTCAACCTGGGAACTGCATTTGTGACTGCATAGCTTGAGTGCGGCAGAGGGGGATGGAATTCCGCGTGTAGCAGTGAAATGCGTAGATATGCGGAGGAACACCGATGGCGAAGGCAATCCCCTGGGCCTGCACTGACGCTCATGCACGAAAGCGTGGGGAGCAAACAGG
## 122                                                          ACGTAGGGTGCAGGCGTTAATCGGAATTACTGGGCGTAAAGCGTGCGCAGGCGGTTTGCTAAGACAGGTGTGAAATCCCCGGGCTTAACCTGGGAATTGCGCTTGTAACTGGCAAGCTAGAGTGTGGCAGAGGGGGGTGGAATTCCACGTGTAGCAGTGAAATGCGTAGATATGTGGAGGAACACCGATGGCGAAGGCAGCCCCCTGGGTTAACACTGACGCTCAGGCACGAAAGCGTGGGGAGCAAACAGG
## 123                                                          ACGTAGGGTGCGAGCGTTAATCGGAATTACTGGGCGTAAAGCGTGCGCAGGCGGTTTTGTAAGACAGAGGTGAAATCCCCGGGCTCAACCTGGGAACTGCCTTTGTGACTGCAAGGCTAGAGTGCGGCAGAGGGGGATGGAATTCCGCGTGTAGCAGTGAAATGCGTAGATATGCGGAGGAACACCGATGGCGAAGGCAATCCCCTGGGCCTGCACTGACGCTCATGCACGAAAGCGTGGGGAGCAAACAGG
## 124                                                          ACAGAGGGTGCAAGCGTTAATCGGAATTACTGGGCGTAAAGCGCGCGTAGGCGGTTGTGTAAGTTGGATGTGAAATCCCCGGGCTTAACCTGGGCACTGCATTCAAAACTGCACGGCTAGAGTATGGGAGAGGAAGGTAGAATTCCAGGTGTAGCGGTGAAATGCGTAGAGATCTGGAGGAATACCGATGGCGAAGGCAGCCTTCTGGCCTAATACTGACGCTGAGGTGCGAAAGCATGGGGAGCAAACAGG
## 125                                                          ACGTAGGGTGCAAGCGTTAATCGGAATTACTGGGCGTAAAGCGTGCGCAGGCGGTTATGTAAGACAGAGGTGAAATCCCCGGGCTCAACCTGGGAACTGCCTTTGTGACTGCATGACTAGAGTACGGCAGAGGGGGATGGAATTCCGCGTGTAGCAGTGAAATGCGTAGATATGCGGAGGAACACCGATGGCGAAGGCAATCCCCTGGGCCTGTACTGACGCTCATGCACGAAAGCGTGGGGAGCAAACAGG
## 126                                                          ACAGAGGGTGCGAGCGTTAATCGGAATTACTGGGCGTAAAGCGCATGCAGATGGTTTGACAAGTCGGATGTGAAAGCCCCGGGCTTAACCTGGGAACGGCACTCGAAACTGTCAGACTAGAATGTGGAAGAGGGAAGTGGAATTCCGGGTGTAGCAGTGAAATGCGTAGATATCCGGAGGAACATCAGTGGCGAAGGCGACTTCCTGGTCCAACATTGACATTCAGATGCGAAAGCGTGGGGAGCAAACAGG
## 127                                                          ACGTAGGGTGCAAGCGTTAATCGGAATTACTGGGCGTAAAGCGTGCGCAGGCGGTTTCGTAAGACAGACGTGAAATCCCCGGGCTCAACCTGGGAACTGCGTTTGTGACTGCGAGGCTAGAGTATGGCAGAGGGGGGTGGAATTCCACGTGTAGCAGTGAAATGCGTAGAGATGTGGAGGAACACCGATGGCGAAGGCAGCCCCCTGGGCCAATACTGACGCTCATGCACGAAAGCGTGGGGAGCAAACAGG
## 128                                                          ACGTAGGGTGCAAGCGTTAATCGGAATTACTGGGCGTAAAGCGTGCGCAGGCGGTGATGTAAGACAGGTGTGAAATCCCCGGGCTTAACCTGGGAACTGCACTTGTGACTGCATCGCTGGAGTACGGCAGAGGGGGATGGAATTCCGCGTGTAGCAGTGAAATGCGTAGATATGCGGAGGAACACCGATGGCGAAGGCAATCCCCTGGGCCTGTACTGACGCTCATGCACGAAAGCGTGGGGAGCAAACAGG
## 129                                                          ACGAAGGGTGCAAGCGTTACTCGGAATTACTGGGCGTAAAGCGTGCGTAGGTGGTTTGTTAAGTCAGATGTGAAAGCCCCGGGCTCAACCTGGGAATTGCATTTGATACTGGCAGACTAGAGTGCGGTAGAGGAGAGTGGAATTCCCGGTGTAGCAGTGAAATGCGTAGAGATCGGGAGGAACATCAGTTGCGAAGGCGGCTCTCTGGACCAGCACTGACACTGAGGCACGAAAGCGTGGGGAGCAAACAGG
## 130                                                          ACGTAGGGTGCGAGCGTTAATCGGAATTACTGGGCGTAAAGCGTGCGCAGGCGGTTTTGAAAGTCAGATGTGAAATCCCCGGGCTCAACCTGGGAACTGCGTTTGAAACTCCAAAGCTAGAGTATGGGAGAGGGAGGTAGAATTCCACGTGTAGCAGTGAAATGCGTAGAGATGTGGAGGAATACCAATGGCGAAGGCAGCCTCCTGGCCTAATACTGACGCTCATGCACGAAAGCGTGGGGAGCAAACAGG
## 131                                                          ACGGAGGATCCAAGCGTTATCCGGAATCATTGGGTTTAAAGGGTCCGTAGGCGGTTTAATAAGTCAGTGGTGAAATCTGGTCGCTCAACGATCAAACGGCCATTGATACTGTTAGACTTGAATTACTGGGAAGTAACTAGAATATGTAGTGTAGCGGTGAAATGCTTAGATATTACATGGAATACCAATTGCGAAGGCAGGTTACTACCAGTGGATTGACGCTGATGGACGAAAGCGTGGGGAGCGAACAGG
## 132                                                          ACGTAGGGTGCAAGCGTTAATCGGAATTACTGGGCGTAAAGCGTGCGCAGGCGGTTCGGAAAGAGAGGTGTGAAATCCCAGGGCTCAACCTTGGAACTGCACTTTTAACTACCGAGCTAGAGTATGTCAGAGGGGGGTGGAATTCCGCGTGTAGCAGTGAAATGCGTAGATATGCGGAGGAACACCGATGGCGAAGGCAGCCCCCTGGGATAATACTGACGCTCATGCACGAAAGCGTGGGGAGCAAACAGG
## 133                                                          ACGAAGGTGGCAAGCGTTACCCGGATTGATTGGGCGTAAAGCGTCCGCAGACTGTATGATAAGTTCTTGGTCAAATCCTGAAGCTCAACTTCAGGGCCGCTGAGAATACTGTCATGCTAGAGACTGGGAGAGGTAAGCGGAATTACCGGTGTAGCAGTAAAATGCGTTAAGATCGGTAGGAACACCAAATGCGAAGGCAGCTTACTGGAACAGTTCTGACGTTATAGGACGAAAGCGTGGGGAGCGAATGGG
## 134                                                          ACAGAGGTCTCAAGCGTTGTTCGGATTCATTGGGCGTAAAGGGTGCGTAGGTGGCAAGGTAAGTCAGATGTGAAATCCCCGGGCTCAACCTGGGAACTGCATTTGATACTGCCGTGCTCGAGTGCTGGAGGGGAGACTGGAATTCACGGTGTAGCAGTGAAATGCGTAGATATCGTGAGGAAGACCGGTGGCGAAGGCGAGTCTCTGGACAGCAACTGACACTGAGGCACGAAGGCTAGGGGAGCAAACGGG
## 135                                                          ACGGAGGGTGCAAGCGTTATCCGGAATCATTGGGTTTAAAGGGTCCGCAGGCGGTCTTATAAGTCAGTGGTGAAATCTCGTAGCTTAACTATGAAACTGCCATTGATACTGTAGGACTTGAATTCGGTCGAAGTGGGCGGAATATGACATGTAGCGGTGAAATGCGTAGATATGTCATAGAACACCGATAGCGAAGGCAGCTCACTAGGCCTGGATTGACGCTCAGGGACGAAAGCGTGGGGAGCAAACAGG
## 136                                                          ACGTAGGGTGCAAGCGTTAATCGGAATTACTGGGCGTAAAGCGTGCGCAGGCGGTTTTGTAAGACAGGCGTGAAATCCCCGGGCTCAACCTGGGAATGGCGCTTGTGACTGCAAAGCTGGAGTGCGGCAGAGGGGGATGGAATTCCGCGTGTAGCAGTGAAATGCGTAGATATGCGGAGGAACACCGATGGCGAAGGCAATCCCCTGGGCCTGCACTGACGCTCATGCACGAAAGCGTGGGGAGCAAACAGG
## 137                                                          ACGGAGGGTGCAAGCGTTAATCGGAATTACTGGGCGTAAAGCGCACGCAGGCGGCTTGTTAAGTCAGATGTGAAAGCCCCGGGCTCAACCTGGGAACTGCATTTGAAACTGACAAGCTAGAGTCTTGTAGAGGGGGGTAGAATTCCAGGTGTAGCGGTGAAATGCGTAGAGATCTGGAGGAATACCGGTGGCGAAGGCGGCCCCCTGGACAAAGACTGACGCTCAGGTGCGAAAGCGTGGGGAGCAAACAGG
## 138                                                          ACAGAGGATGCAAGCGTTATCCGGAATCACTGGGCATAAAGCGTCTGTAGGTTGCTTGTCAAGTCTGCTGTTAAAGATCAGGGCCTAACCCTGGAAAAGCAGTGGAAACTAATAGGCTTGAGTGTGGTAGAGGTAGAGGGAATTCCTGGTGTAGCGGTGAAATGCGTAGATATTAGGAAGAACACCAATGGCGAAAGCACTCTACTGGGCCATAACTGACACTGAGAGACGACAGCTAGGGGAGCAAATGGG
## 139                                                          ACAGAGACTGCAAGCGTTATTCGGATTCACTGGGCGTAAAGGGTGCGCAGGCGGCCGTGTGTGTTAGGCGTGAAAGCCCGAGGCTCAACCTCGGAATTGCACCTAAAACTACACGGCTAGAGTACTGGAGAGGGTAGCGGAATTCACGGTGTAGCAGTGAAATGCGTAGATATCGTGAGGAACACCAATGGCGAAGGCAGCTACCTGGACAGTAACTGACGCTCAGGCACGAAAGCGTGGGGAGCAAAAGGG
## 140                                                          ACGAAGGATCCAAGCGTTGTCCGGATTTACTGGGTTTAAAGGGTGCGTAGGCGGAAAATTAAGTCAGTGGTGAAAGCCCGCAGCTCAACTGCGGAACTGCCATTGAAACTGATTTTCTTGAATACGGTTGAGGTAGATGGAATATAACATGTAGCGGTGAAATGCTTAGATATGTTATAGAACACCAATTGCGAAGGCAGTCTACTAAGCCGTGATTGACGCTGAGGCACGAAAGCGTGGGGAGCGAACAGG
## 141                                                          ACGTAGGGTGCGAGCGTTAATCGGAATTACTGGGCGTAAAGCGTGCGCAGGTGGTTTTGTAAGCTTGATGTGAAATCCCCGGGCTCAACCTGGGAACTGCATTGAGGACTGCAAGACTAGAGTGTGGCAGAGGGGGGTGGAATTCCACGTGTAGCAGTGAAATGCGTAGAGATGTGGAGGAACACCGATGGCGAAGGCAGCCCCCTGGGCTAACACTGACACTCATGCACGAAAGCGTGGGGAGCAAACAGG
## 142                                                          ACGTAGGGTGCGAGCGTTAATCGGAATTACTGGGCGTAAAGGGTGCGCAGGCGGCGCCATAAGACAGCTGTGAAATCCCCGGGCTTAACCTGGGAACTGCGGTTGTGACTGTGGTGCTTGAGTGCGGCAGAGGGGGGTGGAATTCCACGTGTAGCAGTGAAATGCGTAGAGATGTGGAGGAACACCGATGGCGAAGGCAGCCCCCTGGGTCGACACTGACGCTCATGCACGAAAGCGTGGGGAGCAAACAGG
## 143                                                          ACGGAGGATCCAAGCGTTATCCGGAATCATTGGGTTTAAAGGGTCCGTAGGCGGTCTAATAAGTCAGTGGTGAAAGCCCATCGCTCAACGATGGAACGGCCATTGATACTGTTAGACTTGAATTACTGGGAAGTAACTAGAATATGTAGTGTAGCGGTGAAATGCTTAGATATTACATGGAATACCAATTGCGAAGGCAGGTTACTACCAGTGGATTGACGCTGATGGACGAAAGCGTGGGGAGCGAACAGG
## 144                                                          ACGGAGGGGGCTAGCGTTATTCGGAATTACTGGGCGTAAAGCGCACGTAGGCGGACCAGAAAGTCAGAGGTGAAAGCCCGGGGCTCAACCCCGGAACTGCCTTTGAAACTCCTGGTCTTGAGGTCGAGAGAGGTGAGTGGAATTCCGAGTGTAGAGGTGAAATTCGTAGATATTCGGAGGAACACCAGTGGCGAAGGCGGCTCACTGGCTCGATACTGACGCTGAGGTGCGAAAGCGTGGGGAGCAAACAGG
## 145                                                          ACAGAGGGTGCAAGCGTTATCCGGAATTATTGGGCGTAAAGCGTCCGCAGGTGGTTTGTTAAGTCGAGTATTAAAGACGGCAGCTCAACTGTCGGAATGTATTCGATACTGGCAAGCTAGAGTCTTGGAGGGGTGAGCAGAATTCTGCATGTAGGGGTAAAATCCGTAGATATGCAGAGGAATACCAAAAGCGAAGGCAGCTCACTGGCCAAGTACTGACACTCATGGACGAAAGCGTGGGGAGCAAACAGG
## 146                                                          ACGGAGGGTGCAAGCGTTAATCGGAATTACTGGGCGTAAAGCGCACGTAGGTGGTTTGTTAAGCCAGCTGTGAAATCCCCGGGCTCAACCTGGGCACTGCAGTTGGAACTGGCAAGCTAGAGTAGGGTAGAGGGGTGTGGAATTCCAGGTGTAGCGGTGAAATGCGTAGATATCTGGAGGAACATCAGTGGCGAAGGCGACACCCTGGACTCATACTGACACTGAGGTGCGAAAGCGTGGGGAGCAAACAGG
## 147                                                          ACGGAGGGAGCTAGCGTTGTTCGGAATTACTGGGCGTAAAGCGTGCGTAGGCGGCTATTCAAGTCAGAGGTGAAAGCCTGGAGCTCAACTCCAGAACTGCCTTTGAAACTAGATAGCTAGAATCTTGGAGAGGTGAGTGGAATTCCGAGTGTAGAGGTGAAATTCGTAGATATTCGGAAGAACACCAGTGGCGAAGGCGACTCACTGGACAAGTATTGACGCTGAGGTACGAAAGCGTGGGGAGCAAACAGG
## 148                                                          ACGGAGGGTGCAAGCGTTACTCGGAATCACTGGGCGTAAAGAGCGTGTAGGCGGGTATATAAGTCAGAAGTGAAATCCTATAGCTTAACTATAGAACTGCTTTTGAAACTGTATACCTAGAATGTGGGAGAGGTAGATGGAATTTCTGGTGTAGGGGTAAAATCCGTAGAGATCAGAAGGAATACCGATTGCGAAGGCGATCTACTGGAACATTATTGACGCTGAGACGCGAAAGCGTGGGGAGCAAACAGG
## 149                                                          ACGTAGGGTGCAAGCGTTAATCGGAATTACTGGGCGTAAAGCGTGCGCAGGTGGTTTGTTAAGACAGTTGTGAAATCCCCGGGCTTAACCTGGGAACTGCAATTGTGACTGGCAGACTAGAGTGAGGCAGAGGGGGGCGGAATTCCTGGTGTAGCAGTGAAATGCGTAGATATCAGGAGGAACACCGATGGCGAAGGCAGCCCCCTGGGCCATCACTGACACTCATGCACGAAAGCGTGGGGAGCAAACAGG
## 150                                                          ACGTAGGGTGCGAGCGTTAATCGGAATTACTGGGCGTAAAGCGTGCGCAGGCGGTTATATAAGACAGATGTGAAATCCCCGGGCTCAACCTGGGAACTGCATTTGTGACTGTATAGCTAGAGTGCGGCAGAGGGGGATGGAATTCCGCGTGTAGCAGTGAAATGCGTAGATATGCGGAGGAACACCGATGGCGAAGGCAATCCCCTGGGCCTGCACTGACGCTCATGCACGAAAGCGTGGGGAGCAAACAGG
## 151                                                          ACAGAGGTCCCGAGCGTTGTTCGGATTCACTGGGCGTAAAGGGTGCGTAGGTGGTTGGTTAAGTTTGAGGTGAAAGCTCCGAGCTCAACTCGGAAAATGCCTTGAAGACTATCTGACTAGAGGATTGGAGGGGAGATTGGAATTCTCGGTGTAGCAGTGAAATGCGTAGATATCGAGAGGAACACCAGTGGCGAAGGCGAATCTCTGGACAATTCCTGACACTGAGGCACGAAAGCCAAGGGAGCAAACGGG
## 152                                                          ACGTAGGGTGCGAGCGTTAATCGGAATTACTGGGCGTAAAGCGTGCGCAGGCGGTTTTGTAAGACAGATGTGAAATCCCCGGGCTCAACCTGGGACCTGCATTTGTGACTGCAAGGCTGGAGTACGGCAGAGGGGGATGGAATTCCGCGTGTAGCAGTGAAATGCGTAGATATGCGGAGGAACACCGATGGCGAAGGCAATCCCCTGGGCCTGTACTGACGCTCATGCACGAAAGCGTGGGGAGCAAACAGG
## 153                                                          CCCAGCGGTCCAAGTCGCAACCACAATTATTGGGTCTAAAACATCCGTAGCTTGCTTGATAAGTCTCTTGTGAAATCCGGCCTCTTAAGGGTCGAAGAGCGAGAGATACTGTCGGGCTAGAGACCGGGAGAGGTAAGAAGTACGGTCGGGGTACCGGTAAAATGGGTTGATCCCGTCTGGACTAACAACAGCGAAGGCATCTTACCAGCACGGATCTGACAGTAAGGGATGAAGGCTAGGGGCGCAAAAGGG
## 154                                                          ACATAGGGTGCAAGCGTTGTCCGGAATTATTGGGCGTAAAGAGCTCGTAGGTGGTTCGATACGTCGGATGTGAAAATCAGGGGCTCAACCCCTGACCTGCATCCGATACGGTCGAGCTGGAGTTTGGTAGGGGAGACTGGAATTCCTGGTGTAGCGGTGGAATGCGCAGATATCAGGAGGAACACCGATGGCGAAGGCAGGTCTCTGGGCCAATACTGACACTGAGGAGCGAAAGCGTGGGGAGCGAACAGG
## 155                                                          ACATAGGTCCCAAGCGTTATCCGGAATTACTGGGCGTAAAGCGTCTGCAGGTGGAAAAGTGTGTGAGATGTGAAAGACCGGGGCTCAACCCCGTGTTTGTGTCTCAAACTGCTTTTCTAGAGTGAGTAAGAGGTATGCGGAATTTATGGAGTAGGAGTGCAATCCGTTGACACCATAAAGAACACCAAAAGCGAAGGCAGCATACTGGGGCTCTACTGACACTCAGAGACGAAAGCGTGGGGAGCGAAAGGG
## 156                                                          ACGAAGGATCCAAGCGTTGTCCGGATTTACTGGGTTTAAAGGGTGCGTAGGCGGAAAATTAAGTCAGTGGTGAAAGCCCGCAGCTCAACTGTGGAACTGCCATTGAAACTGGTTTTCTTGAATATAGCTGAGGCAGATGGAATATAACATGTAGCGGTGAAATGCTTAGATATGTTATAGAACACCGATTGCGAAGGCAGTCTGCTAAACTATTATTGACGCTGAGGCACGAAAGCGTGGGGAGCGAACAGG
## 157                                                          ACGTAGGGTGCGAGCGTTAATCGGAATTACTGGGCGTAAAGCGTGCGCAGGCGGTGATGTAAGACAGATGTGAAATCCCCGGGCTCAACCTGGGAACTGCGTTTGTGACTGCATCACTCGAGTACGGCAGAGGGAGGTGGAATTCCACGTGTAGCAGTGAAATGCGTAGAGATGTGGAGGAACACCGATGGCGAAGGCAGCCTCCTGGGCCAGTACTGACGCTCATGCACGAAAGCGTGGGGAGCAAACAGG
## 158                                                          ACGGAGGGTGCAAGCGTTAATCGGAATTACTGGGCGTAAAGCGCACGCAGGCGGTTTGTTAAGTCAGATGTGAAATCCCCGGGCTCAACCTGGGAACTGCATCTGATACTGGCAAGCTTGAGTCTCGTAGAGGGGGGTAGAATTCCAGGTGTAGCGGTGAAATGCGTAGAGATCTGGAGGAATACCGGTGGCGAAGGCGGCCCCCTGGACGAAGACTGACGCTCAGGTGCGAAAGCGTGGGGAGCAAACAGG
## 159                                                          ACGTAGGGTGCAAGCGTTAATCGGAATTACTGGGCGTAAAGCGTGCGCAGGCGGTCTTGTAAGACAGATGTGAAATCCCCGGGCTCAACCTGGGACCTGCATTTGTGACTGCAAGGCTGGAGTACGGCAGAGGGGGATGGAATTCCGCGTGTAGCAGTGAAATGCGTAGATATGCGGAGGAACACCGATGGCGAAGGCAATCCCCTGGGCCTGTACTGACGCTCATGCACGAAAGCGTGGGGAGCAAACAGG
## 160                                                         ACGTGAGTGGCAAGCGTTATCCGGAATCATTGGGCGTAAAGCGTCCGTAGGGGGGTGTTTAAGTGAAATGTTAAATTCTTCGACCCAATCGAAGGCCTGCGTTTCATACTGGACACCTAGAAGATGGGAGAGGTAAATAGAATTTCTGGAGTAGGGGTAATATCCGTAGAGACCAGAAGGAATACCAATAGCGAAGGCAATTTACTGGCCCATTCTTGACCCTCAAGTGACGAAAGCGTGGGGAGCAAACAGG
## 161                                                          ACGAGTGCCCCAAGCGTTATCCGGAATTACTGGGCGTAAAGCGTGCGTAGGTGGCATGGTTAGTCTCCTGTTAAAGCTCTCGGCTTAACCGAGAAAGTGCGGGAGATACGGCCAAGCTCGAGGTGGTGAGAGGTCTATGGAACTCATGGTGTAGGGGTGAAATCCGTTGATATCATGGGGAACACCAAATGCGAAGGCAATAGACTGGCACCATACTGACACTGGAGCACGAAAGCGTGGGGATCAAACGGG
## 162                                                          ACGGAGGGTGCAAGCGTTATCCGGATTTATTGGGTTTAAAGGGTCCGTAGGCGGACTTATAAGTCAGTGGTGAAAGCCTGTCGCTTAACGATAGAACTGCCATTGATACTGTAAGTCTTGAGTATATTTGAGGTAGCTGGAATAAGTAGTGTAGCGGTGAAATGCATAGATATTACTTAGAACACCAATTGCGAAGGCAGGTTACCAAGATATAACTGACGCTGAGGGACGAAAGCGTGGGGAGCGAACAGG
## 163                                                          ACGGAGGGTGCAAGCGTTATCCGGATTCACTGGGTTTAAAGGGTGCGTAGGCGGGCAGGTAAGTCAGTGGTGAAATCCTAGAGCTTAACTCTAGAACTGCCATTGATACTATCTGTCTTGAATATTGTGGAGGTAAGCGGAATATGTCATGTAGCGGTGAAATGCTTAGATATGACATAGAACACCTATTGCGAAGGCAGCTTACTACGCATATATTGACGCTGAGGCACGAAAGCGTGGGGATCAAACAGG
## 164                                                          ACGTAGGGTGCAAGCGTTAATCGGAATTACTGGGCGTAAAGCGTGCGCAGGCGGACTTTTAAGCCAGATGTGAAAGCCCCGAGCTTAACTTGGGAATTGCGTTTGGAACTGGGAGTCTAGAGTCTGTCAGAGGGGGATGGAATTCCACGTGTAGCAGTGAAATGCGTAGATATGCGGAGGAACACCGATGGCGAAGGCAATCCCCTGGGCCTGTACTGACGCTCATGCACGAAAGCGTGGGGAGCAAACAGG
## 165                                                          ACGAAGGGTGCAAGCGTTAATCGGAATTACTGGGCGTAAAGCGTGCGTAGGTGGTTTGTTAAGTCAGATGTGAAAGCCCTGGGCTCAACCTGGGAACTGCATTTGATACTGGCAGGCTAGAGTGCGGTAGAGGGTGGTGGAATTCCCGGTGTAGCGGTGAAATGCGTAGAGATCGGGAGGAACATTCGTGGCGAAGGCGGCCACCTGGACCAGCACTGACACTGAGGCACGAAAGCGTGGGGAGCAAACAGG
## 166                                                          ACGGAGGATCCAAGCGTTGTCCGGATTTATTGGGTTTAAAGGGTGCGTAGGCGGATCTTTAAGTCAGTGGTGAAAGCCTGCAGCTTAACTGTAGAACAGCCATTGAAACTGAAGATCTTGAATTTGGTTAAAGTAGGCGGAATGTATCATGTAGCGGTGAAATGCTTAGATATGATACAGAACACCGATAGCGAAGGCAGCTTGCTGAACCAATATTGACGCTGAGGCACGAAAGCGTGGGGAGCGAACAGG
## 167                                                          ACGTAGGGTGCAAGCGTTAATCGGAATTACTGGGCGTAAAGCGTGCGCAGGCGGTTTTGTAAGTCGGATGTGAAATCCCCGGGCTCAACCTGGGAACTGCGTTCGAAACTGCAAGGCTAGAGTGTGTCAGAGGGGGGTAGAATTCCACGTGTAGCAGTGAAATGCGTAGAGATGTGGAGGAATACCAATGGCGAAGGCAGCCCCCTGGGATAACACTGACGCTCATGCACGAAAGCGTGGGGAGCAAACAGG
## 168                                                          ACGTAGGGTGCGAGCGTTAATCGGAATTACTGGGCGTAAAGCGTGCGCAGGCGGTTATATAAGACAGATGTGAAATCCCCGGGCTTAACCTGGGACCTGCATTTGTGACTGTATAGCTAGAGTACGGTAGAGGGGGATGGAATTCCGCGTGTAGCAGTGAAATGCGTAGATATGCGGAGGAACACCGATGGCGAAGGCAATCCCCTGGACCTGTACTGACGCTCATGCACGAAAGCGTGGGGAGCAAACAGG
## 169                                                          ACGGAGGGTGCAAGCGTTGTCCGGATTTATTGGGTTTAAAGGGTGCGCAGGTGGTTTATTAAGTCAGTGGTGAAAGACGGTCGCTCAACGATTGCAGTGCCATTGATACTGGTAGACTTGAGTAAAGTAGAGGTGGGCGGAATTGATAGTGTAGCGGTGAAATGCATAGATATTATCAAGAACTCCAATTGCGTAGGCAGCTCACTTGGCTTTTACTGACACTCATGCACGAAAGTGTGGGTATCAAACAGG
## 170                                                          ACAGAGGGTGCAAGCGTTAATCGGAATTACTGGGCGTAAAGCGCACGTAGGCGGTTTTTTAAGTCAGATGTGAAAGCCCCGGGCTCAACCTGGGAATTGCATTTGAAACTGGAAAACTAGAGTGTGTGAGAGGGGGGTAGAATTCCAAGTGTAGCGGTGAAATGCGTAGAGATTTGGAGGAATACCAGTGGCGAAGGCGGCCCCCTGGCACAACACTGACGCTCAGGTGCGAAAGCGTGGGGAGCAAACAGG
## 171                                                          ACGTAGGGTGCGAGCGTTAATCGGAATTACTGGGCGTAAAGCGTGCGCAGGCGGCTTTGCAAGACAGATGTGAAATCCCCGGGCTCAACCTGGGAACTGCATTTGTGACTGCAAGGCTAGAGTACGGTAGAGGGGGATGGAATTCCGCGTGTAGCAGTGAAATGCGTAGATATGCGGAGGAACACCAATGGCGAAGGCAATCCCCTGGACCTGTACTGACGCTCATGCACGAAAGCGTGGGGAGCAAACAGG
## 172                                                          ACGTAGGGTGCAAGCGTTAATCGGAATTACTGGGCGTAAAGCGTGCGCAGGCGGTTTTGTAAGCCAGATGTGAAATCCCCGAGCTTAACTTGGGAACTGCGTTTGGAACTACAAGACTAGAGTGTGTCAGAGGGGGGTAGAATTCCACGTGTAGCAGTGAAATGCGTAGAGATGTGGAGGAATACCGATGGCGAAGGCAGCCCCCTGGGATAACACTGACGCTCATGCACGAAAGCGTGGGGAGCAAACAGG
## 173                                                         ACGTAGGATCCAAGCGTTATCCGGAATTACTGGGCGTAAAGCGTGCGCAGACGGTTTGGTAGGTCTGGTGCGAAATCCGGTGGCTCAACCACCTGGACTGTACCAGAAACCCCCTGACTCGAGGTCGGTAGAGGCAAGTGGAATTGCTGGTGTAGGGGTGACATCCGTAGATATCAGCAGGAACACCAATGGCGAAGGCAGCTTGCTGGGCCGTACCTGACGTTCAGGCACGAAAGCGTGGGGAGCGAACAGG
## 174                                                          ACAGAGGATGCAAGCGTTATCCGGAATCACTGGGCATAAAGCGTCTGTAGGTTGCTTGCCAAGTCTGCTGTTAAAGACTAGGGCCTAACCCTGGGAAAGCAGTGGAAACTAGCAAGCTTGAGTGTGGTAGAGGTAGAGGGAATTCCTGGTGTAGCGGTGAAATGCGTAGATATTAGGAAGAACACCAATGGCGAAAGCACTCTGCTGGGCCACAACTGACACTGAGAGACGACAGCTAGGGGAGCAAATGGG
## 175                                                          ACGTAGGGTGCGAGCGTTAATCGGAATTACTGGGCGTAAAGCGTGCGCAGGTGGTTACGTAAGTCTGATGTGAAATCCCCGAGCTTAACTTGGGAACTGCATTGGAGACTGCGTGGCTAGAGTGTGTCAGAGGGGGGTGGAATTCCACGTGTAGCAGTGAAATGCGTAGAGATGTGGAGGAACACCGATGGCGAAGGCAGCCCCCTGGGATAACACTGACACTCATGCACGAAAGCGTGGGGAGCAAACAGG
## 176                                                          ACGTAGGGTGCGAGCGTTAATCGGAATTACTGGGCGTAAAGCGTGCGCAGGCGGCTTTTTAAGCCAGATGTGAAATCCCCGGGCTTAACCTGGGAACTGCATTTGGAACTGGAAGGCTAGAGTGCGGCAGAGGGGGGTAGAATTCCACGTGTAGCAGTGAAATGCGTAGAGATGTGGAGGAATACCGATGGCGAAGGCAGCCCCCTGGGTCGACACTGACGCTCATGCACGAAAGCGTGGGGAGCAAACAGG
## 177                                                          ACGGAGGGTGCAAGCGTTATCCGGAATCACTGGGTTTAAAGGGTGCGTAGGCGGCCAGATAAGTCAGAGGTGAAAGTTTCCGGCTCAACCGGGAGATTGCCTTTGATACTGTTTGGCTTGAATCAGATTGAGGTTGGCGGAATGTGACATGTAGCGGTGAAATGCATAGATATGTCATAGAACATCGATTGCGAAGGCAGCTGGCTGGATCTGTATTGACGCTGAGGCACGAAAGCGTGGGGAGCAAACAGG
## 178                                                          ACGGAGGGAGCTAGCGTTGTTCGGAATTACTGGGCGTAAAGAGTACGTAGGCGGTTATTCAAGTCAGAGGTGAAAGCCTGGAGCTCAACTCCAGAACTGCCTTTGAAACTAGATAGCTAGAATTATGGAGAGGTTAGTGGAATTCCGAGTGTAGAGGTGAAATTCGTAGATATTCGGAAGAACACCAGTGGCGAAGGCGACTAACTGGACATATATTGACGCTGAGGTACGAAAGCGTGGGGAGCAAACAGG
## 179                                                          ACGGAGGGTGCAAGCGTTAATCGGAATTACTGGGCGTAAAGCGCGCGTAGGCGGCTGATTAAGTCGGATGTGAAAGCCCCGGGCTCAACCTGGGAACTGCATACGATACTGATCGGCTAGAGTATGAGAGAGGGAGGTAGAATTCCACGTGTAGCGGTGAAATGCGTAGATATGTGGAGGAATACCGGTGGCGAAGGCGGCCTCCTGGCTTAATACTGACGCTGAGGTGCGAAAGCGTGGGGAGCAAACAGG
## 180                                                          ACGGAGGGTGCAAGCGTTACTCGGAATCACTGGGCGTAAAGGATGCGTAGGCTGGAAATCAAGTCGAGAGTGAAATCCAACGGCTCAACCGTTGAACTGCTCTCGAAACTGGTTACCTAGAATATGGGAGAGGTAGATGGAATTGGTGGTGTAGGGGTAAAATCCGTAGATATCACCAGGAATACCGATTGCGAAGGCGATCTACTGGAACATTATTGACGCTGAGGCATGAAAGCGTGGGGAGCAAACAGG
## 181                                                          ACGTAGGGTGCAAGCGTTAATCGGAATTACTGGGCGTAAAGCGTGCGCAGGCGGTTATATAAGTCAGATGTGAAATCCCCGGGCTCAACCTGGGAACTGCCTTTGTGACTGCATAGCTAGAGTACGGCAGAGGGGGATGGAATTCCGCGTGTAGCAGTGAAATGCGTAGATATGCGGAGGAACACCGATGGCGAAGGCAATCCCCTGGGCCTGTACTGACGCTCATGCACGAAAGCGTGGGGAGCAAACAGG
## 182                                                          ACGAAGGGTGCAAGCGTTAATCGGAATTACTGGGCGTAAAGCGCGCGTAGGTGGTTCAGCAAGTTGGAGGTGAAATCCCCGGGCTCAACCTGGGAACTGCCTCCAAAACTACTGAGCTAGAGTACGGTAGAGGGTAGTGGAATTTCCTGTGTAGCGGTGAAATGCGTAGATATAGGAAGGAACACCAGTGGCGAAGGCGACTACCTGGACTGATACTGACACTGAGGTGCGAAAGCGTGGGGAGCAAACAGG
## 183                                                          ACGTAGGGTGCAAGCGTTAATCGGAATTACTGGGCGTAAAGCGTGCGCAGGCGGTTATGTAAGACAGATGTGAAATCCCCGGGCTCAACCTGGGAACTGCATTTGTGACTGCATAGCTGGAGTGCGGCAGAGGGGGATGGAATTCCGCGTGTAGCAGTGAAATGCGTAGATATGCGGAGGAACACCGATGGCGAAGGCAATCCCCTGGGCCTGCACTGACGCTCATGCACGAAAGCGTGGGGAGCAAACAGG
## 184                                                          ACATAGGTCCCAAGCGTTATCCGGAATTACTGGGCGTAAAGCGTCTGCAGGTTGAAAAGTGTGTGAGATGTGAAAGACCGGGGCTCAACCCCGCGTTTGTGTCTCAAACTGCTTTTCTCGAGTGAGCAAGAGGTATGCGGAATTTATGGAGTAGGAGTGCAATCCGTTGACACCATAAAGAACACCAAAAGCGAAGGCAGCATACTGGGGCTCTACTGACACTCAGAGACGAAAGCGTGGGGAGCGAAAGGG
## 185                                                          ACGGAGGATCCAAGCGTTATCCGGAATCATTGGGTTTAAAGGGTCCGTAGGCGGTTTAATAAGTCAGTGGTGAAATCTGGTCGCTCAACGATCAAACGGCCATTGATACTGTTAGACTTGAATTACTAGGAAGTAACTAGAATATGTAGTGTAGCGGTGAAATGCTTAGATATTACATGGAATACCAATTGCGAAGGCAGGTTACTACTAGTGGATTGACGCTGATGGACGAAAGCGTGGGGAGCGAACAGG
## 186                                                          ACGTAGGGTGCGAGCGTTAATCGGAATTACTGGGCGTAAAGCGTGCGCAGGCGGTTGTGTAAGTCAGATGTGAAATCCCCGGGCTCAACCTGGGAACTGCATTTGAGACTGCACGGCTAGAGTGTGACAGAGGGGGGTAGAATTCCACGTGTAGCAGTGAAATGCGTAGAGATGTGGAGGAATACCGATGGCGAAGGCAGCCCCCTGGGTTACTACTGACGCTCATGCACGAAAGCGTGGGGAGCAAACAGG
## 187                                                          ACGGAGGGTGCAAGCGTTGTCCGGATTTATTGGGTTTAAAGGGTGCGCAGGTGGTTTATTAAGTCAGTGGTGAAAGACGGTCGCTCAACGATTGCAGTGCCATTGAAACTAGTAGACTTGAGTAAAGTAGAGGTGGGCGGAATTGATAGTGTAGCGGTGAAATGCATAGATATTATCAAGAACTCCAATTGCGTAGGCAGCTCACTTGGCTTTTACTGACACTCATGCACGAAAGTGTGGGTATCAAACAGG
## 188                                                          ACGGAGGGGGCTAGCGTTGTTCGGAATTACTGGGCGTAAAGGGCGCGTAGGCGGATTAGTAAGTTGGGAGTGAAAGCCCGGGGCTTAACCTCGGAACTGCTTTCAAAACTGCTAGTCTTGAGTGAAGTAGGGGATGATGGAATTCCTAGTGTAGAGGTGAAATTCTTAGATATTAGGAGGAACACCGGTGGCGAAGGCGGTCATCTGGACTTCAACTGACGCTGAGGCGCGAAAGCGTGGGGAGCAAACAGG
## 189                                                          ACGGAGGATCCGAGCGTTATCCGGATTTATTGGGTTTAAAGGGAGCGTAGGTTGACATATAAGTCAGCTGTGAAAGTTTACGGCTCAACCGTGAAATTGCAGTTGATACTGTATGTCTTGAGTGTACAAGAGGTGGGCGGAATTCGTGGTGTAGCGGTGAAATGCTTAGATATCACGAAGAACTCCAATTGCGAAGGCAGCTCACTGGGGTACAACTGACACTGAGGCTCGAAAGTGTGGGTATCAAACAGG
## 190                                                          ACATAGGGTGCAAGCGTTGTCCGGAATTATTGGGCGTAAAGAGCTCGTAGGTGGTTCGTTACGTCGGATGTGAAAATCTGGGGCTCAACCCCAGACCTGCATCCGATACGGTCGAGCTAGAGTTTGGTAGGGGAGACTGGAATTCCTGGTGTAGCGGTGGAATGCGCAGATATCAGGAGGAACACCAATGGCGAAGGCAGGTCTCTGGGCCAATACTGACACTGAGGAGCGAAAGCGTGGGGAGCGAACAGG
## 191                                                          ACAGAGACTGCAAGCGTTACTCGGATTCACTGGGCGTAAAGGGAGCGCAGGCGGGCGGGTGTGTCATATGTGAAATCCCGGGGCTCAACCCCGGGGCTGCATGTGAAACTACCTGTCTAGAGATTCGGAGGGGTAAGCGGAATTCCTGGTGGAGCAGTGAAATGCGTAGATATCAGGAGGAACACCAACGGCGAAGGCAGCTTACTGGACGAAATCTGACGCTCAGGCTCGAAAGCATGGGGAGCAAAAGGG
## 192                                                          ACGGAGGATCCAAGCGTTATCCGGATTTATTGGGTTTAAAGGGTCCGTAGGCGGGTCTTTAAGTCAGTGGTGAAAGCCTGCAGCTTAACTGTGGAAATGCCATTGATACTGGAGGCCTTGAGTGTAGTGGAAGTAGGCGGAATAGGGCATGTAGCGGTGAAATGCATAGATATGCCCTAGAACACCGATTGCGAAGGCAGCTTACTACGTTACAACTGACGCTGAGGGACGAAAGCGTGGGGAGCAAACAGG
## 193                                                          ACGGAGGGTGCAAGCGTTAATCGGAATTACTGGGCGTAAAGCGTGCGTAGGCGGCTCTTTAAGTCAGATGTGAAATCCCCGGGCTCAACCTGGGAACTGCATCTGATACTGGGGAGCTAGAATGTGGGAGAGGGAAGTGGAATTTCCGGTGTAGCGGTGAAATGCATAGAGATCGGAAGGAACATCAGTGGCGAAGGCGACTTCCTGGACCAACATTGACGCTGAGGCACGAAAGCGTGGGTAGCAAACAGG
## 194                                                          ACATAGGGTGCAAGCGTTGTCCGGAATTATTGGGCGTAAAGAGCTCGTAGGTGGTTTGTTACGTCGGTTGTGAAATTCAGGGGCTCAACCCCTGACTTGCAGCCGATACGGGCAAGCTAGAGTTTGGTAGGGGAGACTGGAATTCCTGGTGTAGCGGTGGAATGCGCAGATATCAGGAGGAACACCGATGGCGAAGGCAGGTCTCTGGGCCAATACTGACACTGAGGAGCGAAAGCGTGGGGAGCGAACAGG
## 195                                                          ACAGAGGGTGCAAGCGTTAATCGGAATTACTGGGCGTAAAGCGTGCGTAGACGGTTATTTAAGTCGGATGTGAAATCCCCGGGCTCAACCTGGGAATTGCATTCGAGACTGAATAGCTAGGGTGCGGAAGAGGGAAGCGGAATTTCCGGTGTAGCGGTGAAATGCGTAGATATCGGAAGGAACATCAGTGGCGAAAGCGGCTTCCTGGTCCAGCACCGACGTTCAGGCACGAAAGCGTGGGGAGCAAACAGG
## 196                                                          ACGGAGGGGGCTAGCGTTGTTCGGAATTACTGGGCGTAAAGCGCACGTAGGCGGACTGGAAAGTCAGAGGTGAAATCCCAGGGCTCAACCTTGGAACTGCCTTTGAAACTCCCAGTCTTGAGGTCGAGAGAGGTGAGTGGAATTCCGAGTGTAGAGGTGAAATTCGTAGATATTCGGAGGAACACCAGTGGCGAAGGCGGCTCACTGGCTCGATACTGACGCTGAGGTGCGAAAGCGTGGGGAGCAAACAGG
## 197                                                          ACGGAGGATCCAAGCGTTATCCGGAATCATTGGGTTTAAAGGGTCCGTAGGCGGCCTTATAAGTCAGTGGTGAAATCTCCCGGCTCAACCGGGAAATGGCCATTGATACTGTAGGGCTTGAATTATTAGGAAGTAACTAGAATATGTAGTGTAGCGGTGAAATGCTTAGAGATTACATGGAATACCAATTGCGAAGGCAGGTTACTACTAATGGATTGACGCTGATGGACGAAAGCGTGGGTAGCGAACAGG
## 198                                                          ACAGAGACTGCAAGCGTTACTCGGATTCACTGGGCGTAAAGGGAGCGCAGGCGGGCGGGTGTGTCATATGTGAAATCCCGGGGCTCAACCCCGGGGCTGCATCTGAAACTACCTGTCTAGAGATTCGGAGGGGTAAGCGGAATTCCTGGTGGAGCAGTGAAATGCGTAGATATCAGGAGGAACACCAACGGCGAAGGCAGCTTACTGGACGAAATCTGACGCTCAGGCTCGAAAGCATGGGGAGCAAAAGGG
## 199                                                          ACGAGTGGCCCAAGCGTTATCCGGAATTATTGGGCGTAAAGGATGTGTAGGTGGTTGTATTAGTCTTTGGTCTAACCTCTGTGCTTAACATGGAGACAGCCGAGGAAACGGTACGACTAGAGTAGAGTAGGGGTAAGCGGAACTCATAGTGTAGGGGTGAAATCCGTTGATATTATGGGGAACACCAAAAGCGAAGGCAGCTTACTGGACTCTTACTGACACTGAAACATGAAAGCGTGGGTAGCGAATGGG
## 200                                                          ACGAGTGCCTCAAGCGTTATCCGGAATCATTGGGCGTAAAGGATGTGTAGGTGGTCGTGTTAGTCTCTCGTTAAATTCTTCGGCTTAACCGGGGGCATGCGGGGGAAACGGCACGACTAGAGGGTGCAAGGGGTCTATGGAACTCATAGTGTAGCGGTGAAATGCGTTGATATTATGGGGAACACCAAAAGCGAAGGCAGTAGACTAGGGCACTCCTGACACTGAAACATGAAAGCGTGGGTAGCGAATGGG
## 201                                                          ACGGAGGGGGCTAGCGTTATTCGGAATTACTGGGCGTAAAGCGCACGTAGGCGGATTAGAAAGTCAGAGGTGAAATCCCAGGGCTCAACCTTGGAACTGCCTTTGAAACTCCTAGTCTTGAGGTCGAGAGAGGTGAGTGGAATTCCGAGTGTAGAGGTGAAATTCGTAGATATTCGGAGGAACACCAGTGGCGAAGGCGGCTCACTGGCTCGATACTGACGCTGAGGTGCGAAAGCGTGGGGAGCAAACAGG
## 202                                                          ACGGAGGATGCAAGCGTTATCCGGATTTATTGGGTTTAAAGGGTGCGTAGGCGGTTGGATAAGTCAGCGGTGAAAGTTTTCTGCTTAACAGGAAAAATGCCGTTGATACTGTCTGGCTAGAATATGGTTGCTGTGAGTGGAATGTGTGGTGTAGCGGTGAAATGCTTAGATATCACACAGAATATCGATTGCGAAGGCAGCTCACAAAGCCATTATTGACGCTGAGGCACGAAAGTGTGGGGATCAAACAGG
## 203                                                          ACGGAGGGTGCGAGCGTTGTCCGGAATCACTGGGCGTAAAGGGCGCGTAGGCGGTCTGCTAAGCGTGTGGTGAAAGCTCGGGGCTCAACCCTGAGTCGGCCATGCGAACTGGTGGACTTGAGCACTGTAGAGGCAGGTGGAATTCCGGGTGTAGCGGTGGAATGCGTAGAGATCCGGAAGAACACCGGTGGCGAAGGCGGCCTGCTGGGCAGTAGCTGACGCTGAGGCGCGACAGCGTGGGGAGCAAACAGG
## 204                                                          ACGAAGGGTGCAAGCGTTGTTCGGAATTACTGGGCGTAAAGCGCGCGTAGGCGGTCTACTAAGTCAGACGTGAAAGCCCTGGGCTTAACCCGGGAATGGCGTTTGATACTGGTAGACTAGAGATTCAGAGAGGTGAGTGGAATTCCTAGTGTAGAGGTGAAATTCGTAGATATTAGGAGGAACATCAGTGGCGAAGGCGGCTCACTGGCTGAATACTGACGCTAAGGCGCGAAAGCGTGGGGAGCAAACAGG
## 205                                                          ACGGAGGGTGCAAGCGTTAATCGGAATTACTGGGCGTAAAGCGCACGCAGGCGGTCTGTCAAGTCGGATGTGAAATCCCCGGGCTCAACCTGGGAACTGCATTCGAAACTGGCAGGCTAGAGTCTTGTAGAGGGGGGTAGAATTCCAGGTGTAGCGGTGAAATGCGTAGAGATCTGGAGGAATACCGGTGGCGAAGGCGGCCCCCTGGACAAAGACTGACGCTCAGGTGCGAAAGCGTGGGGAGCAAACAGG
## 206                                                          ACGTAGGGTGCGAGCGTTAATCGGAATTACTGGGCGTAAAGCGTGCGCAGGCGGTTATGCAAGACAGAGGTGAAATCCCCGGGCTCAACCTGGGAACTGCCTTTGTGACTGCATAGCTAGAGTACGGCAGAGGGGGATGGAATTCCGCGTGTAGCAGTGAAATGCGTAGATATGCGGAGGAACACCGATGGCGAAGGCAATCCCCTGGGCCTGTACTGACGCTCATGCACGAAAGCGTGGGGAGCAAACAGG
## 207                                                          ACGGAGGGTGCAAGCGTTATCCGGATTCACTGGGTTTAAAGGGAGCGTAGGTGGGCTTTTAAGTCAGTGGTGAAATCTTTGGGCTTAACCCGAAAACTGCCATTGATACTATTAGTCTTGAATTCTCTGGAGGTAAGCGGAATATGTCATGTAGCGGTGAAATGCTTAGATATGACATAGAACACCCATTGCGAAGGCAGCTTACTACGGAGACATTGACACTGAGGCTCGAAAGCGTGGGGATCAAACAGG
## 208                                                          ACGAAGGGTGCAAGCGTTAATCGGAATTACTGGGCGTAAAGCGCGCGTAGGTGGTTTTGTAAGTTGGAGGTGAAATCCCCGGGCTCAACCTGGGAACTGCCTCCAAAACTGCATGACTAGAGTACGGTAGAGGGTGGTGGAATTTCCTGTGTAGCGGTGAAATGCGTAGATATAGGAAGGAACACCAGTGGCGAAGGCGACCACCTGGACTGATACTGACACTGAGGTGCGAAAGCGTGGGGAGCAAACAGG
## 209                                                          ACGGAGGGTGCAAGCGTTAATCGGAATAACTGGGCGTAAAGCGCACGCAGGCGGTTGGATAAGTCAGATGTGAAAGCCCCGGGCTCAACCTGGGAACTGCATTTGAAACTGTCTGACTAGAGTCTTGTAGAGGGGGGTAGAATTCCAGGTGTAGCGGTGAAATGCGTAGAGATCTGGAGGAATACCGGTGGCGAAGGCGGCCCCCTGGACAAAGACTGACGCTCAGGTGCGAAAGCGTGGGGAGCAAACAGG
## 210                                                          ACAGAGGGTGCAAGCGTTAATCGGAATTACTGGGCGTAAAGCGCACGTAGGCGGTTGTTTAAGTTGGATGTGAAAGCCCCGGGCTCAACCTGGGAATTGCATTCAAAACTGGACAACTAGAGTGTGTGAGAGGGGGGTAGAATTCCAAGTGTAGCGGTGAAATGCGTAGAGATTTGGAGGAATACCAGTGGCGAAGGCGGCCCCCTGGCACAACACTGACGCTCAGGTGCGAAAGCGTGGGGAGCAAACAGG
## 211                                                          ACGTAGGGTGCGAGCGTTAATCGGAATTACTGGGCGTAAAGCGTGCGCAGGCGGTTATATAAGACAGTTGTGAAATCCCCGGGCTCAACCTGGGAATTGCATCTGTGACTGTGTAGCTAGAGTACGGTAGAGGGGGATGGAATTCCGCGTGTAGCAGTGAAATGCGTAGATATGCGGAGGAACACCGATGGCGAAGGCAATCCCCTGGACCTGTACTGACGCTCATGCACGAAAGCGTGGGGAGCAAACAGG
## 212                                                          ACATAGGGTGCAAGCGTTGTCCGGAATTATTGGGCGTAAAGAGCTCGTAGGTGGTTAGATACGTCGGATGTGAAAATCAGGGGCTCAACCCCTGACCTGCATTCGATACGGTCTAGCTAGAGTGTGGTAGGGGAGACTGGAATTCCTGGTGTAGCGGTGGAATGCGCAGATATCAGGAGGAACACCGATGGCGAAGGCAGGTCTCTGGGCCATTACTGACACTGAGGAGCGAAAGCGTGGGGAGCGAACAGG
## 213                                                          ACGAAGGATGCAAGCGTTATCCGGATTCATTGGGTTTAAAGGGTGCGTAGGCGGACTTGTAAGTCAGTGGTGAAATCTCTTTGCTTAACAAAGAAACTGCCATTGATACTGCAGGTCTAGAGTATAGATGACGTTGGCGGAATATGACATGTAGTGGTGAAATACTTAGATATGTCATAGAACACCGATTGCGAAGGCAGCTAACGAAACTATAACTGACGCTGAGGCACGAAAGTGCGGGGATCAAACAGG
## 214                                                          ACGTAGGGTGCAAGCGTTAATCGGAATTACTGGGCGTAAAGCGTGCGCAGGCGGTTATGTAAGACAGAGGTGAAATCCCCGGGCTCAACCTGGGAACTGCCTTTGTGACTGCATAGCTTGAGTGCGGCAGAGGGGGATGGAATTCCGCGTGTAGCAGTGAAATGCGTAGATATGCGGAGGAACACCGATGGCGAAGGCAATCCCCTGGGCCTGCACTGACGCTCATGCACGAAAGCGTGGGGAGCAAACAGG
## 215                                                          ACGGAGGGTGCAAGCGTTATCCGGATTTATTGGGTTTAAAGGGTCCGTAGGCGGGCCGATAAGTCAGTGGTGAAAGCCCATAGCTCAACTATGGAACTGCCATTGATACTGTCGGTCTTGAGTACAAGTGAGGTTGGCGGAATGTGTGGTGTAGCGGTGAAATGCTTAGATATCACACAGAACACCGATTGCGAAGGCAGCTGGCCAACTTGTAACTGACGCTGAGGGACGAAAGTGTGGGGATCAAACAGG
## 216                                                          ACGGAAGGTCCAGGCGTTATCCGGATTTATTGGGTTTAAAGGGAGCGTAGGCCGTCTGTTAAGCGTGTTGTGAAATGTAGATGCTCAACATCTGAATTGCAGCGCGAACTGGCAGACTTGAGTGTGCGCAACGTAGGCGGAATTCGTGGTGTAGCGGTGAAATGCTTAGATATCACGAAGAACTCCGATTGCGAAGGCAGCTTACGGGAGCACAACTGACGCTGAAGCTCGAAAGTGCGGGTATCGAACAGG
## 217                                                          ACGAAGGGTGCAAGCGTTGCTCGGAATTATTGGGCGTAAAGGGTAGGTAGGTGGTTGCACATGTCTGGGGTGAAATCCCTGAGCTTAACTCAGGAAGTGCCTTGGAAACGGTGTAACTAGAGTACTAGAGAGGTTCGTGGAATTCTTGGTGTAGCGGTGAAATGCGTAGAGATCAAGAGGAACATCAGCGGCGAAGGCGGCGAACTGGATAGTAACTGACACTCAACTACGAAAGCGTGGGGAGCAAACAGG
## 218                                                          ACGAGTGCCTCGAGCGTTATCCGGAATTATTGGGCGTAAAGGGTGTGTAGGTGGTTATGTTAGTCTCCTGTCAAATTCTTCGGCTTAACCGGGGGTCCGCAGGGGAAACGGCATGACTTGAGGATGCGAGAGGTAAATGGAACTCATAGTGTAGGGGTAAAATCCGTTGATATTATGGGGAACACCAAAAGCGAAGGCAATTTACTGGCGCACTCCTGACACTGAAACACGAAAGCGTGGGTAGCGAATGGG
## 219                                                          ACGAAGGGGGCTAGCGTTGCTCGGAATTACTGGGCGTAAAGGGCGCGTAGGCGGCTTAGTCAGTCAGGCGTGAAATTCCTGGGCTTAACCTGGGGTCTGCGTTTGAGACGGCTAGGCTAGAGGATAGGAGAGGGTCGTGGAATTCCCAGTGTAGAGGTGAAATTCGTAGATATTGGGAAGAACACCGGTGGCGAAGGCGGCGACCTGGCCTATTACTGACGCTGAGGCGCGACAGCGTGGGGAGCAAACAGG
## 220                                                          ACGTAGGGGGCAAGCGTTATCCGGATTTACTGGGTGTAAAGGGAGCGTAGACGGCGATGCAAGTCAGATGTGAAAACCCATGGCTCAACCATGGGATTGCATTTGAAACTGTGTTGCTAGAGTGCAGGAGAGGTAAGCGGAATTCCTGGTGTAGCGGTGAAATGCGTAGATATCAGGAGGAACACCGGTGGCGAAGGCGGCTTACTAGACTGTAACTGACGTTGAGGCTCGAAGGCGTGGGTAGCAAACAGG
## 221                                                          ACGTAGGGTGCGAGCGTTAATCGGAATTACTGGGCGTAAAGCGTGCGCAGGCGGCGTGACAAGTCAGATGTGAAATCCCCGAGCTCAACTTGGGAACTGCGTTTGAAACTGTCAGGCTAGAATATGTCAGAGGGGGGTAGAATTCCACGTGTAGCAGTGAAATGCGTAGAGATGTGGAGGAATACCAATGGCGAAGGCAGCCCCCTGGGATAATATTGACGCTCATGCACGAAAGCGTGGGGAGCAAACAGG
## 222                                                          ACGAAGGATCCAAGCGTTATCCGGATTCATTGGGTTTAAAGGGTGCGTAGGCGGATAGATAAGTCAGTGGTGAAAGCCGGTCGCTCAACGATCGAATTGCCATTGATACTGTTTATCTAGAATATAGATGATGTTGGCGGAATATGACATGTAGCGGTGAAATGCATAGATATGTCATAGAACACCTATTGCGAAGGCAGCTGACAAAACTATTATTGACGCTGATGCACGAAAGTGCGGGGATCAAACAGG
## 223                                                          ACGTAGGGTGCGAGCGTTAATCGGAATTACTGGGCGTAAAGCGTGCGCAGGCGGTTATGTAAGACAGAGGTGAAATCCCCGGGCTCAACCTGGGAACTGCCTTTGTGACTGCATAGCTGGAGTGCGGCAGAGGGGGATGGAATTCCGCGTGTAGCAGTGAAATGCGTAGATATGCGGAGGAACACCGATGGCGAAGGCAATCCCCTGGGCCTGCACTGACGCTCATGCACGAAAGCGTGGGGAGCAAACAGG
## 224                                                          ACGTAGGCTTCAAGCGTTATCCGGAATTACTGGGTGTAAAGCGTCTGTAGGCGGCAGAGTAAGTTTGGCATGAAAGACCGGGGCTTAACCCCGTGTTTGTGTCGAAAACTGCTCAGCTAGAATCTGGGAGAGGGAAGCAGAATGATGTGAGTAGGGGTGCAATCCGTTGATACACATCAGAATACCAAAAGCGAAGGCAGCTTCCTGGAACAGTATTGACGCTCAGAGACGAAAGCGTGGGGAGCGAAAAGG
## 225                                                          ACGGAGGGTGCAAGCGTTACTCGGAATCACTGGGCGTAAAGAGCGTGTAGGCGGATTGATAAGTTTGAAGTGAAATCCTATAGCTTAACTATAGAACTGCTTTGAAAACTGTTAATCTAGAATGTGGGAGAGGTAGATGGAATTTCTGGTGTAGGGGTAAAATCCGTAGAGATCAGAAGGAATACCGATTGCGAAGGCGATCTACTGGAACAATATTGACGCTGAGACGCGAAAGCGTGGGGAGCAAACAGG
## 226                                                          ACGGAGGGTGCAAGCGTTGTCCGGATTTATTGGGTTTAAAGGGTGCGTAGGCGGCTATTTAAGTCAGCGGTGAAAGACTTCAGCTTAACTGGAGCAGTGCCATTGATACTGGATAGCTTGAGTGTTGGAGGGGTACATGGAATTGATGGTGTAGCGGTGAAATGCATAGATACCATCAGGAACACCGATAGCGAAGGCATTGTACTGGCCAACAACTGACGCTGAGGCACGAAAGTGTGGGGATCGAACAGG
## 227                                                          ACGTAGGGTGCAAGCGTTAATCGGAATTACTGGGCGTAAAGCGTGCGTAGACGGTTATCTAAGTCGGATGTGAAATCCCCGGGCTCAACCTGGGAATTGCATTCGAGACTGAATAGCTAGGGTGCGGAAGAGGGAAGCGGAATTTCCGGTGTAGCGGTGAAATGCGTAGATATCGGAAGGAACATCAGTGGCGAAAGCGGCTTCCTGGTCCAGCACCGACGTTCAGGCACGAAAGCGTGGGGAGCAAACAGG
## 228                                                          ACGTAGGGTGCGAGCGTTAATCGGAATTACTGGGCGTAAAGCGTGCGCAGGCGGTTATGTAAGACAGATGTGAAATCCCCGGGCTCAACCTGGGACCTGCATTTGTGACTGCATAGCTAGAGTACGGTAGAGGGGGATGGAATTCCGCGTGTAGCAGTGAAATGCGTAGATATGCGGAGGAACACCGATGGCGAAGGCAATCCCCTGGACCTGTACTGACGCTCATGCACGAAAGCGTGGGGAGCAAACAGG
## 229                                                          ACGTAGGGTGCAAGCGTTAATCGGAATTACTGGGCGTAAAGCGTGCGCAGGTGGTTTGTTAAGACAGTTGTGAAATCCCCGGGCTCAACCTGGGAACTGCAATTGTGACTGACAGACTAGAGTTTGGCAGAGGGGGGTGGAATTCCTGGTGTAGCAGTGAAATGCGTAGATATCAGGAGGAACACCGATGGCGAAGGCAGCCCCCTGGGCCATGACTGACACTCATGCACGAAAGCGTGGGGAGCAAACAGG
## 230                                                          ACGGAAGGTCCGGGCGTTATCCGGATTTATTGGGTTTAAAGGGAGCGTAGGCGGAGCGTCAAGCCAGCAGTGAAATGTAGCGGCCCAACCGCTGCACTGCTGTTGGAACTGGCGCCCTTGAGTGCACACGAGGCATGCGGAATTTGTGGTGTAGCGGTGAAATGCTTAGATATCACGAAGAACTCCGATTGCGAAGGCAGCGTGCCGGAGTGCAACTGACGCTGAAGCTCGAAAGTGCGGGTATCGAACAGG
## 231                                                          ACGGAGGGTGCAAGCGTTATCCGGAATCATTGGGTTTAAAGGGTCCGCAGGCGGGCTTATAAGTCAGTGGTGAAAGCCTACAGCTTAACTGTAGAACTGCCATTGATACTGTAAGTCTTGAATTCGGTCGAAGTGGGCGGAATGTGTAGTGTAGCGGTGAAATGCTTAGATATTACACAGAACACCGATAGCGAAGGCAGCTCACTAGGCCTGAATTGACGCTCATGGACGAAAGCGTGGGGAGCAAACAGG
## 232                                                          ACGGGAGTGGCAAGCGTTATCCGGAATTATTGGGCGTAAAGCGTCCGCAGGCGGTCTTGTAAGTCTGTTGTTAAAGCGTGGAGCTTAACTCCATTTCAGCAATGGAAACTGTAAGACTAGAGTGTGGTAGGGGCAGAGGGAATTCCCGGTGTAGCGGTGAAATGCGTAGATATCGGGAAGAACACCAGTGGCGAAGGCGCTCTGCTGGGCCATAACTGACGCTCATGGACGAAAGCCAGGGGAGCGAAAGGG
## 233                                                          ACAGAGGGTGCGAGCGTTAATCGGAATTACTGGGCGTAAAGCGTGCGTAGGCGGTTTGATAAGCTGGATGTGAAATCCCCGGGCTTAACCTGGGAACTGCATTCAGGACTGTTTGACTAGAGTCTGGGAGAGGGTGGTGGAATTTCCTGTGTAGCGGTGAAATGCGTAGATATAGGAAGGAACATCAGTGGCGAAGGCGACTGCCTGGCCCAAGACTGACGCTGAGGCACGAAAGCGTGGGTAGCAAACAGG
## 234                                                          ACAGAGGGTGCAAGCGTTGTTCGGACTTACTGGGCGTAAAGCGCGTGTAGGCGGTCTTTTAAGTCTGGTGTGAAAGCCTGAGGCTTAACCTCAGAAGTGCACTGGATACTGGAAGACTAGAGGGTGAAAGAGGAGAGTGGAATTCATGGTGTAGGGGTGAAATCCGTAGATATCATGAGGAACATCAGTGGCGAAGGCGGCTCTCTGGTTCACTACTGACGCTCAGACGCGAAAGCATGGGGAGCAAACAGG
## 235                                                          ACGAGTGCTCCGAGCGTTATCCGGAATCATTGGGCGTAAAGGGTGTGTAGGTGGTAATACTAGTCTTCAGTTAAATTCTTCGGCTCAACCGGGGACATGCTGGGGAAACGGTATTACTAGAGGATGCGAGAGGTGTGTGGAACTCTATGTGTAGGGGTGAAATCCGTTGATATATAGGGGAACACCAAAAGCGAAGGCAGCACACTGGCGCACTCCTGACACTGAAACACTAAAGCGTGGGTAGCGAATGGG
## 236                                                          ACGGAGGGTGCAAGCGTTATCCGGATTTACTGGGTTTAAAGGGTGCGTAGGCGGGCAGGTAAGTCAGTGGTGAAATCCCCGAGCTTAACTCGGGAACTGCCGTTGATACTATCTGTCTTGAATATAGTGGAGGTAAGCGGAATATGTCATGTAGCGGTGAAATGCTTAGATATGACATAGAACACCAATTGCGAAGGCAGCTTGCTACACTATCATTGACGCTGAGGCACGAAAGCGTGGGGATCAAACAGG
## 237                                                          ACGTAGGGTGCGAGCGTTAATCGGAATTACTGGGCGTAAAGCGTGCGCAGGCGGTTTCGTAAGACAGACGTGAAATCCCCGGGCTCAACCTGGGAACTGCGTTTGTGACTGCGAGGCTAGAGTACGGCAGAGGGGGGTAGAATTCCACGTGTAGCAGTGAAATGCGTAGAGATGTGGAGGAATACCGATGGCGAAGGCAGCCCCCTGGGTTAGTACTGACGCTCATGCACGAAAGCGTGGGGAGCAAACAGG
## 238                                                          ACAGAGGGTGCAAGCGTTAATCGGAATGACTGGGCGTAAAGCGTGCGTAGGTGGCAAAATAAGTTAGTTGTGAAATCCCTGGGCTTAACCTAGGAACTGCAATTGATACTGTTTAGCTAGAGTATAGTAGAGGTAAGTGGAATTTCCGGTGTAGCGGTGAAATGCGTAGATATCGGAAGGAACACCAGTGGCGAAGGCGGCTTACTGGGCTAATACTGACACTGAGGCACGAAAGCGTGGGGAGCAAACAGG
## 239                                                          ACGTAGGGTGCGAGCGTTAATCGGAATTACTGGGCGTAAAGCGTGCGCAGGCGGTTATGTAAGACAGATGTGAAATCCCCGGGCTCAACCTGGGAACTGCGTTTGTGACTGCATAACTAGAGTACGGCAGAGGGAGGTGGAATTCCGCGTGTAGCAGTGAAATGCGTAGAGATGCGGAGGAACACCGATGGCGAAGGCAGCCTCCTGGGCCAGTACTGACGCTCATGCACGAAAGCGTGGGGAGCAAACAGG
## 240                                                          ACGTAGGGTCCAAGCGTTAATCGGAATTACTGGGCGTAAAGCGTGCGCAGGCGGCTGATTAAGCCAGATGTGAAATCCCCGGGCTCAACCTGGGAACTGCGTTTGGAACTGGTCAGCTAGAGTACGTCAGAGGGGGGTAGAATTCCACGTGTAGCAGTGAAATGCGTAGAGATGTGGAGGAATACCGATGGCGAAGGCAGCCCCCTGGGATGATACTGACGCTCATGCACGAAAGCGTGGGGAGCAAACAGG
## 241                                                          ACGTAGGGTGCGAGCGTTAATCGGAATTACTGGGCGTAAAGCGTGCGCAGGCGGTTTCTTAAGCCAGATGTGAAATCCCCGGGCTTAACCTGGGAACTGCATTTGGAACTGGGAGACTAGAGTGTGTCAGAGGGAGGTGGAATTCCACGTGTAGCAGTGAAATGCGTAGAGATGTGGAGGAACACCGATGGCGAAGGCAGCCTCCTGGGATGACACTGACGCTCATGCACGAAAGCGTGGGGAGCAAACAGG
## 242                                                          ACGTAGGGTGCAAGCGTTAATCGGAATTACTGGGCGTAAAGCGTGCGCAGGCGGTTTTGTAAGACAGGCGTGAAATCCCCGGGCTTAACCTGGGAATTGCGCTTGTGACTGCAAGGCTAGAGTACGGTAGAGGGGGGTAGAATTCCGCGTGTAGCAGTGAAATGCGTAGATATGCGGAGGAACACCGATGGCGAAGGCAGCCCCCTGGACCTGTACTGACGCTCATGCACGAAAGCGTGGGGAGCAAACAGG
## 243                                                      ACAGAGACCTCAAACGTTATCCGGAATCATTGGGCGTAAAGCGTACCGATAGGTGGTTTTATAAGTCAGAAGTGAAATCCTAAGGCTTAACCTTGGGACTGTCTTTTGAAACTATAAAACTAGAGGGGCAAAGAGGAAGCTGGAACAAACGGTGTAGTAGTGAAATGCGTTGATATCGTTTGGAACACCAATAGCGTAGGCAGGCTTCTGGGTGCCACCTGACACTGCTAGGACGAAAGCGTGGGGAGCGATAAGG
## 244                                                        ACGTAGGGTGCAAGCATTATCCGGAGTGACTGGGCGTAAAGAGTTGCGTAGGTGGTTATGTAAGTGAATAGTGAAACCTGGTGGCTCAACCATACAGACTATTATTCAAACTGCATAACTCGAGAGTGGTAGAGGTAACTGGAATTTCTTGTGTAGGAGTGAAATCCGTAGATATAAGAAGGAACACCAATGGCGTAGGCAGGTTACTGGACCATTTCTGACACTAAGGCACGAAAGCGTGGGGAGCGAACCGG
## 245                                                          ACGGAGGGTGCAAGCGTTATCCGGATTCACTGGGTTTAAAGGGTGCGTAGGCGGGCAGGTAAGTCAGTGGTGAAATCCCCGAGCTTAACTCGGGAACTGCCGTTGATACTATCTGTCTTGAATACCGTGGAGGTGAGCGGAATATGTCATGTAGCGGTGAAATGCTTAGATATGACATAGAACACCAATTGCGAAGGCAGCTCGCTACACGGTCATTGACGCTGAGGCACGAAAGCGTGGGGATCAAACAGG
## 246                                                          ACGTAGGGTGCGAGCGTTAATCGGAATTACTGGGCGTAAAGCGTGCGCAGGCGGTTTTGTAAGACAGGCGTGAAATCCCCGGGCTCAACCTGGGAACTGCGCTTGTGACTGCATCACTCGAGTACGGCAGAGGGGGGTGGAATTCCACGTGTAGCAGTGAAATGCGTAGAGATGTGGAGGAACACCGATGGCGAAGGCAGCCCCCTGGGCCGATACTGACGCTCATGCACGAAAGCGTGGGTAGCAAACAGG
## 247                                                          ACGGAGGATGCAAGTGTTATCCGGAATCACTGGGCGTAAAGCGTCTGTAGGTGGTTTGATAAGTCAACTGTTAAATCTTGAAGCTCAACTTCAAAATCGCAGTCGAAACTATTAGACTAGAGTATAGTAGGGGTAAAGGGAATTTCCAGTGGAGCGGTGAAATGCGTAGATATTGGAAAGAACACCGATGGCGAAGGCACTTTACTGGGCTATTACTGACACTCAGAGACGAAAGCTAGGGTAGCAAATGGG
## 248                                                          ACGGAGGGTGCGAGCGTTAATCGGAATTACTGGGCGTAAAGCGCACGTAGGCGGTTCTGTAAGTGGGATGTGAAACCCCCGGGCTTAACCTGGGAATTGCATACCAGACTGCAAAGCTAGAGTATGAGAGAGGGTGGTGGAATTTCCGGTGTAGCGGTGAAATGCGTAGAGATCGGAAGGAACATCAGTGGCGAAGGCGGCCACCTGGCTTAATACTGACGCTGAGGTGCGAAAGCGTGGGGATCAAACAGG
## 249                                                          ACAGAGACTGCAAGCGTTACTCGGATTCACTGGGCGTAAAGGGAGCGCAGGCGGACTCGTGTGTCGGACGTGAAATACCGGGGCTTAACCCCGGTGCTGCGTTCGAAACTACGAGTCTAGAGACTTGGAGGGGTAAGCGGAATTCTTGGTGGAGCAGTGAAATGCGTAGATATCAAGAGGAACACCAACGGCGAAGGCAGCTTACTGGACAAGATCTGACGCTCAGGCTCGAAAGCGTGGGGAGCAAAAGGG
## 250                                                          ACGTAGGGTGCGAGCGTTAATCGGAATTACTGGGCGTAAAGCGTGCGCAGGCGGTTTCGTAAGACAGAGGTGAAATCCCCGGGCTCAACCTGGGAACTGCCTTTGTGACTGCGAGGCTAGAGTATGGCAGAGGGGGGTGGAATTCCACGTGTAGCAGTGAAATGCGTAGATATGCGGAGGAACACCGATGGCGAAGGCAATCCCCTGGGCCTGTACTGACGCTCATGCACGAAAGCGTGGGGAGCAAACAGG
## 251                                                          ACGGAAGGTGCAAGCGTTAATCGGAATTACTGGGCGTAAAGCGCGCGTAGGCGGCTTGGTAAGTCGGATGTGAAAGCCCCGGGCTCAACCTGGGAACTGCATTCGATACTGCCGGGCTGGAGTACGAGAGAGGGGGGTGGAATTCCAGGTGTAGCGGTGAAATGCGTAGAGATCTGGAGGAACATCGGTGGCGAAGGCGGCCCCCTGGTTCGATACTGACGCTGAGGTGCGAAAGCGTGGGGAGCAAACAGG
## 252                                                          ACGTAGGGTGCGAGCGTTAATCGGAATTACTGGGCGTAAAGCGTGCGCAGGCGGTTGTGCAAGACAGATGTGAAATCCCCGGGCTCAACCTGGGAATGGCATTTGTGACTGCACGGCTAGAGTGTGTCAGAGGGGGGTAGAATTCCACGTGTAGCAGTGAAATGCGTAGATATGTGGAGGAATACCGATGGCGAAGGCAGCCCCCTGGGATAACACTGACGCTCATGCACGAAAGCGTGGGGAGCAAACAGG
## 253                                                      ACAGAGACCTCAAGCGTTATCCGGAATCATTGGGCGTAAAGCGTACCGACAGGTGGTCATGCAAGTCAGAGGTGAAATCTCCGAGCTTAACTCGGAATCTGTCCTTTGAAACTGCACGACTAGAGGGGCAAAGAGGAAGCTGGAACAAACGGTGTAGTAGTGAAATGCGTTGATATCGTTTGGAACACCAATAGCGAAGGCGGGCTTCTGGGTGCCACCTGACACTGCTAGGACGAAAGCGTGGGGAGCGAATGGG
## 254                                                          ACGGAGGGTGCAAGCGTTATCCGGATTCACTGGGTTTAAAGGGTGCGTAGGTGGATCTGTAAGTCAGAGGTGAAATCCCCGAGCTTAACTTGGGAACTGCCTTTGATACTATAGATCTTGAATATCGTGGAGGTAAGCGGAATATGTCATGTAGCGGTGAAATGCATAGAGATGACATAGAACACCAATTGCGAAGGCAGCTTGCTACACGATGATTGACACTGAGGCACGAAAGCGTGGGGAGCAAACAGG
## 255                                                          ACGTAGGGTGCGAGCGTTAATCGGAATTACTGGGCGTAAAGCGTGCGCAGGCGGTTCTATAAGACAGATGTGAAATCCCCGGGCTTAACCTGGGAACTGCGTTTGTGACTGTAGGACTCGAGTGTGGCAGAGGGGGGTGGAATTCCACGTGTAGCAGTGAAATGCGTAGAGATGTGGAGGAACACCGATGGCGAAGGCAGCCCCCTGGGTCAACACTGACGCTCATGCACGAAAGCGTGGGGAGCAAACAGG
## 256                                                          ACGGAGGGGGCTAGCGTTGTTCGGAATTACTGGGCGTAAAGCGCACGTAGGCGGATCAGAAAGTCAGAGGTGAAATCCCAGGGCTCAACCTTGGAACTGCCTTTGAAACTCCTGATCTTGAGGTCGAGAGAGGTGAGTGGAATTCCGAGTGTAGAGGTGAAATTCGTAGATATTCGGAGGAACACCAGTGGCGAAGGCGGCTCACTGGCTCGATACTGACGCTGAGGTGCGAAAGCGTGGGGAGCAAACAGG
## 257                                                          ACGGAGGGGGCTAGCGTTGTTCGGAATTACTGGGCGTAAAGCGCACGTAGGCGGACTGGAAAGTCAGAGGTGAAATCCCGGGGCTCAACCCCGGAACTGCCTTTGAAACTCCCAGTCTCGAGTCCGGGAGAGGTGAGTGGAATTCCGAGTGTAGAGGTGAAATTCGTAGATATTCGGAGGAACACCAGTGGCGAAGGCGGCTCACTGGCCCGGTACTGACGCTGAGGTGCGAAAGCGTGGGGAGCAAACAGG
## 258                                                          ACGAGGGGTGCAAGCGTTGTTCGGAATTACTGGGCGTAAAGCGCGTGTAGGTGGTTGAGTATGTCGAATGTGAAAGCCCCAGGCTTAACCTGGGAAGTGCGTTCGAAACTGCTTAACTCGAGTACGGGAGAGGAGGGCGGAATTCCCAGTGTAGAGGTGAAATTCGTAGATATTGGGAGGAACACCGGAGGCGAAAGCGGCTCTCTAGACCGTAACTGACACTGAGACGCGAAAGCGTGGGGATCAAACAGG
## 259                                                          ACGTAGGGTGCGAGCGTTAATCGGAATTACTGGGCGTAAAGCGTGCGCAGGCGGTTTTGAAAGTCAGATGTGAAATCCCCGAGCTCAACTTGGGAACTGCGTTTGAAACTCCAAAACTCGAATATGTCAGAGGGGGGTAGAATTCCACGTGTAGCAGTGAAATGCGTAGAGATGTGGAGGAATACCAATGGCGAAGGCAGCCCCCTGGGATAATATTGACGCTCATGCACGAAAGCGTGGGGAGCAAACAGG
## 260                                                          ACGTAGGGTGCAAGCGTTAATCGGAATTACTGGGCGTAAAGCGTGCGCAGGCGGACTTTTAAGCCAGATGTGAAAGCCCCGAGCTTAACTTGGGAATTGCGTTTGGAACTGGGAGTCTAGAGTCTGTCAGAGGGGGATGGAATTCCACGTGTAGCAGTGAAATGCGTAGAGATGTGGAGGAACACCGATGGCGAAGGCAGCCCCCTGGGCCAATACTGACGCTCATGCACGAAAGCGTGGGGAGCAAACAGG
## 261                                                           ACGGGAGGGGCAAGCGTTATCCAAAATTACTGGGCGTAAAGTGTCCGTAGACTGTAAATTAAGTTTTATGTTAAAATTTAAAGCTTAACTTTAAAAAAATATACAATACTGTTTTACTTGAGTTTTATACGGAAGAGTAGAATTTCATGTCAAAGAGTAAATTCTAAAAATACATGAAGGAATATCGAAAGCGAAGGCGACTCTTTAGTACAAACTGACGTTGAGGGACGAAAGTGTAGGGAGCGAACAGG
## 262                                                          ACGGAGGGTGCAAGCGTTATCCGGAATCATTGGGTTTAAAGGGTCCGCAGGCGGTCTTATAAGTCAGTGGTGAAAGCCTATTGCTCAACAATAGAACTGCCATTGATACTGTTAGACTTGAATTAGGTCGGAGTGGGCGGAATATGACATGTAGCGGTGAAATGCTTAGATATGTCATAGAACACCGATAGCGAAGGCAGCTCACTAGCCCTAAATTGACGCTCATGGACGAAAGCGTGGGGAGCAAACAGG
## 263                                                          ACAGAGGATGCAAGCGTTATCCGGAATCACTGGGCATAAAGCGTCTGTAGGTCGCCTACCAAGTCTGCTGTTAAAGATCAGGGCCTAACCCTGGGAAAGCAGTGGAAACTAGTAGGCTTGAGTATGGTAGAGGTAGAGGGAATTCCTGGTGTAGCGGTGAAATGCGTAGATATTAGGAAGAACACCAATGGCGAAAGCACTCTACTGGGCCATAACTGACACTGAGAGACGACAGCTAGGGGAGCAAATGGG
## 264                                                          ACGTGGAGTGCGAGCGTTATCCGGATTTACTGGGCGTAAAGAGTTCGTAGGCGTTTGATAAAGTTTCGTTTGAAATACCGAGGCTCAACTTCGGGAACGGACGAAATACTTATCAGATTGAGATATCTAGGGGGTACTGGAACTGACAGTGTAGCAGTGAAATGCGTTGATATTGTCAAGAACACCAAGGGCGAAGGCAGGTACCTGGGGATATTCTGACGCTGAGGAACGAAAGCTAGGGGAGCGAAAGGG
## 265                                                          ACGAAGGGGGCTAGCGTTGTTCGGAATTACTGGGCGTAAAGGGCGCGTAGGCGGCGCTGTAAGTGAGGTGTGAAAGACCTGGGCTCAACCTGGGAAGTGCATTTCAGACTGCAGTGCTAGAGTGCGAGAGAGGAAAGTGGAATGACGCGTGTAGAGGTGAAATTCGTAGATATGCGTTGGAACACCAGTGGCGAAGGCGACTTTCTGGCTCGTAACTGACGCTAAGGCGCGAAAGCGTGGGGAGCAAACAGG
## 266                                                          ACGTAGGATGCAAGCGTTATCCGGAATTATTGGGCGTAAAGCGTTCGTAGGTGGTTTGTTAAGTAGGGCGTTAAATCTTTGGGCTTAACCCAAAGCTGGCGTCCTAAACTGGCTCACTAGAGACAGGTAGAGGTAAGTGGAATTCTGTATGTAGGGGTAATATCCGTAGATATGCAGAGGAACACCAAAAGCGAAAGCAACTTACTGGACCTGTTCTGACACTCAAGAACGAAAGCGTGGGGAGCAAACGGG
## 267                                                          ACGTAGGGTGCAAGCATTAATCGGAATTATTGGGCGTAAAGGGCGCGTAGGCGGATTGATAAGTCAGATGTGAAATACCAAAGCTCAACTTTGGTGCTGCATTTGAAACTGTCTCTCTAGAGGATTGACGGAGAAAGGGGAATTCCACGTGTAGCGGTGAAATGCGTAGATACGTGGAGGAACACCGGTGGCGAAAGCGCCTTTCTAGTTTAGACCTGACGCTGAGGCGCGAGAGCAAGGGGAGCAAACAGG
## 268                                                          ACGGAGGATCCAAGCGTTATCCGGATTTATTGGGTTTAAAGGGAGCGTAGATGGATGTTTAAGTCAGTTGTGAAAGTTCGGGGCTCAACCCCGGAATTGCAGTTGATACTGGATATCTTGAGTACAGTTGAGGTAGGCGGAATTCGTGGTGTAGCGGTGAAATGCTTAGATATCACGAAGAACTCCGATTGCGAAGGCAGCTTACTAAGCTGTAACTGACATTGAGGCTCGAAAGTGTGGGTATCAAACAGG
## 269                                                          ACGAGTGCCTCAAGCGTTACCCGGAATCACTGGGCGTAAAGGTTGTGTAGGTGGTTGTATTAGTCTTTTGTTAAAGCCTGCGGCTTAACCGCAGAATCGCAAAGGAAACGGTACAACTAGAGGATGCAAGGGGTGTGTGGAACTCATAGTGTAGCGGTGAAATGCGTTGATATTATGGGGAACACCAAAAGCGAAGGCAGCACACTGGTGCACTCCTGACACTGAAACAAGAAAGCGTAGGTAGCGAATGGG
## 270                                                          ACAGAGGGTGCGAGCGTTAATCGGATTTACTGGGCGTAAAGCGTGCGTAGGCGGCTTTTTAAGTCGGATGTGAAATCCCTGAGCTTAACTTAGGAATTGCATTCGATACTGGGAAGCTAGAGTATGGGAGAGGATGGTAGAATTCCAGGTGTAGCGGTGAAATGCGTAGAGATCTGGAGGAATACCGATGGCGAAGGCAGCCATCTGGCCTAATACTGACGCTGAGGTACGAAAGCATGGGGAGCAAACAGG
## 271                                                          ACGGAGGGTGCAAGCGTTATCCGGATTTATTGGGTTTAAAGGGTCCGTAGGCGGATGTGTAAGTCAGTGGTGAAATCTCACAGCTTAACTGTGAAACTGCCATTGATACTGCATGTCTTGAGTGTTGTTGAAGTAGCTGGAATAAGTAGTGTAGCGGTGAAATGCATAGATATTACTTAGAACACCAATTGCGAAGGCAGGTTACTAAGCAACAACTGACGCTGATGGACGAAAGCGTGGGGAGCGAACAGG
## 272                                                         ACGAGGGCCCCGAGCGTTATCCGGAATTATTGGGCGTAAAGGGTGTGTAGGTGGTCATATTAGTCTTGTGTAAAAGCCTGTCGCCTAACGGCAGATCCGCATGGGAAACGGTATGACTGAGAGGGTGTAAGAGGTATATGGAACTCACGGTGTAGGGGTGAAATCCGTTGATATCGTGGGGAACGCCAAAAGCGAAAGCAATATACTGGTACATACCTGACACTGAAACACGAAAGCTAGGGTAGCGAATGGG
## 273                                                          ACGTAGGGTGCGAGCGTTAATCGGAATTACTGGGCGTAAAGCGTGCGCAGGCGGTTTTGTAAGACAGGCGTGAAATCCCCGGGCTCAACCTGGGAACTGCGTTTGTGACTGCAAGGCTAGAGTATGGCAGAGGGGGGTGGAATTCCACGTGTAGCAGTGAAATGCGTAGAGATGTGGAGGAACACCGATGGCGAAGGCAGCCCCCTGGGCCAATACTGACGCTCATGCACGAAAGCGTGGGGAGCAAACAGG
## 274                                                          ACGAAGGGGGCTAGCGTTGCTCGGAATCACTGGGCGTAAAGCGCACGTAGGCGGATCCTTAAGTCAGAGGTGAAATCCCAAGGCTCAACCTTGGAACGGCCTTTGATACTGGGGATCTCGAGTCCGGAAGAGGTTGGTGGAACTGCGAGTGTAGAGGTGAAATTCGTAGATATTCGCAAGAACACCAGTGGCGAAGGCGGCCAACTGGTCCGGCACTGACGCTGAGGTGCGAAAGCGTGGGGAGCAAACAGG
## 275                                                          ACGGAGGGTGCAAGCGTTGTTCGGAATTACTGGGCGTAAAGCGTGTCTAGGTGGATTTGTAAGTCAGGCGTGAAATCCCCCGGCTCAACCGGGGAGGGTCGCTTGATACTGCATATCTTGAGTTTCGGAGAGGTGGGCAGAATTCCCAGTGTAGAGGTGAAATTCGTAGAGATTGGGAGGAATACCGGAGACGAAGGTGGCTCACTAGCCGAATACTGACACTCAAGCACGAAAGCGTGGGTAGCAAACAGG
## 276                                                          ACAGAGGGTGCGAGCGTTAATCGGAATTACTGGGCGTAAAGCGCATGCAGATGGCTTTGTAAGTCGGATGTGAAATCCCCGGGCTTAACCTGGGAACTGCACTCGAAACTGCAAGGCTAGAATGTGGAAGAGGGAAGTGGAATTCCGGGTGTAGCAGTGAAATGCGTAGATATCCGGAGGAACATCAGTGGCGAAGGCGACTTCCTGGTCCAACATTGACATTCAGATGCGAAAGCGTGGGGAGCAAACAGG
## 277                                                          ACGTAGGGTGCAAGCGTTAATCGGAATTACTGGGCGTAAAGCGTGCGCAGGCGGTTATATAAGACAAATGTGAAATCCCCGGGCTCAACCTGGGAACTGCATATGTGACTGTATAGCTAGAGTGCGGCAGAGGGGGATGGAATTCCGCGTGTAGCAGTGAAATGCGTAGATATGCGGAGGAACACCGATGGCGAAGGCAATCCCCTGGGCCTGCACTGACGCTCATGCACGAAAGCGTGGGGAGCAAACAGG
## 278                                                          ACGTAGGGTGCAAGCGTTAATCGGAATTACTGGGCGTAAAGCGTGCGCAGGCGGTTATGCAAGACAGATGTGAAATCCCCGGGCTTAACCTGGGAACTGCATTTGTGACTGCATGGCTAGAGTGCGGCAGAGGGGGATGGAATTCCGCGTGTAGCAGTGAAATGCGTAGATATGCGGAGGAACACCGATGGCGAAGGCAATCCCCTGGGCCTGCACTGACGCTCATGCACGAAAGCGTGGGGAGCAAACAGG
## 279                                                         ACGAAGGGAGCAAGCGTTATCCTAAATGATTGGGCGTAAAGGGTTCGTAGGTTGTAATTGTTGTCGTATGTGAAATCTTGAAATTTTCTTTCAAATATGCATATGATACTCTATTACTTGAGTATATCAGGGGATAACAGAATGTTATATGGAGGAGTAAAATCCATTGAGCTATAATGGAATGCCAGTGGCGAAAGCGGTTATCTGGGATATATACTGACGCTGAGGAACGAAAGCGTGGGGAGCAAACAGG
## 280                                                          ACGTAGGGTGCGAGCGTTAATCGGAATTACTGGGCGTAAAGCGTGCGCAGGCGGTTTTGTAAGTCAGATGTGAAATCCCCGAGCTTAACTTGGGAACTGCGTTTGAAACTACAAGACTAGAGTGTGTCAGAGGGGGGTAGAATTCCACGTGTAGCAGTGAAATGCGTAGAGATGTGGAGGAATATCAATGGCGAAGGCAGCCCCCTGGGATAACACTGACGCTCATGCACGAAAGCGTGGGGAGCGAACAGG
## 281                                                          ACGTAGGGTGCGAGCGTTAATCGGAATTACTGGGCGTAAAGCGTGCGCAGGCGGCTTTGCAAGACAGATGTGAAATCCCCGGGCTCAACCTGGGAACTGCATTTGTGACTGCAAGGCTAGAGTACGGCAGAGGGGGATGGAATTCCGCGTGTAGCAGTGAAATGCGTAGATATGCGGAGGAACACCAATGGCGAAGGCAATCCCCTGGGCCTGTACTGACGCTCATGCACGAAAGCGTGGGGAGCAAACAGG
## 282                                                          ACGAGTGCTTCGAGCGTTATCCGGAATCATTGGGCGTAAAGGGTGTGTAGGTTGTTGTGTTAGTCTCTCGTCAAATCTTTCGGCTCAACCGAGAGTTCGCGAGGGAAACGGCACGACTAGAGGTTGGGAGAGGTTTCTGGAACTCATGGTGTAGCGGTGAAATGCGTTGATATCATGGGGAACACCAAAAGCGAAGGCAAGAAACTGGCCCATACCTGACACTGAAACACGAAAGCGTGGGTCGCGAATGGG
## 283                                                          ACGAAGGGTGCAAGCGTTGTTCGGATTTATTGGGCGTAAAGCGCGCGTAGGCGGACCTGTAAGTCAGATGTGAAATCTCGGTGCTCAACACCGAAACTGCGTCTGAAACTGCGGGTCTAGAATCTTGGAGGGGGAGTGGGAATTTCGCATGTAGGGGTAAAATCCGTAGAGATGCGAAGGAACACCAGAGGCGAAGGCGCACTCCTGGACAAGTATTGACGCTGAGGCGCGAAAGCGTGGGGATCAAACAGG
## 284                                                          ACGAAGGGGGCTAGCGTTGCTCGGAATCACTGGGCGTAAAGCGCACGTAGGCGGGCTTTTAAGTCGGAGGTGAAATCCCAGGGCTCAACCCTGGAACTGCCTTCGATACTGAAAGTCTTGAGTCCGGAAGAGGTAAGTGGAACTCCGAGTGTAGAGGTGAAATTCGTAGATATTCGGAAGAACACCAGTGGCGAAGGCGGCTTACTGGTCCGGTACTGACGCTGAGGTGCGAAAGCGTGGGGAGCAAACAGG
## 285                                                          ACGTAGGGAGCAAGCGTTATCCGGAATTACTGGGTGTAAAGGGAGTGTAGGCGGGATTACAAGTCAGATGTGAAAACTGTGGGCTCAACTCACAGATTGCATTTGAAACTGTAGTTCTTGAGTGAAGTAGAGGTAAGTGGAATTCCTAGTGTAGCGGTGAAATGCGTAGATATTAGGAGGAACATCAGTGGCGAAGGCGACTTACTGGGCTTTAACTGACGCTGAGGCTCGAAAGCGTGGGGAGCAAACAGG
## 286                                                          ACGGAGGATCCAAGCGTTATCCGGAATCATTGGGTTTAAAGGGTCCGTAGGCGGTTTAATAAGTCAGTGGTGAAAGCCCATCGCTCAACGGTGGAACGGCCATTGATACTGTTAAACTTGAATTATTAGGAAGTAACTAGAATATGTAGTGTAGCGGTGAAATGCTTAGAGATTACATGGAATACCAATTGCGAAGGCAGGTTACTACTAATGGATTGACGCTGATGGACGAAAGCGTGGGTAGCGAACAGG
## 287                                                         ACGAAGGGAGCAAGCGTTATCCTAAATGATTGGGCGTAAAGGGTTCGTAGGTTGTAATTGTTGTCGTATGTGAAATCTTGAAACTTTATTTCAAATATGCATATGATACTCTATTACTTGAGTATATCAGGGGATAACAGAATGTTATATGGAGGAGTAAAATCCATTGAGCTATAATGGAATGCCAGTGGCGAAAGCGGTTATCTGGGATATATACTGACGCTGAGGAACGAAAGCGTGGGGAGCAAACAGG
## 288                                                      ACAGAGACCTCAAGCGTTATCCGGAATCATTGGGCGTAAAGCGTACCGATAGGTGGCTTTACAAGTCAGAAGTGAAATCCTAAGGCTTAACTTTGGGACTGTCTTTTGAAACTGTAAAGCTAGAGGGGCAAAGAGGAAGCTGGAACGAACGGTGTAGTAGTGAAATGCGTTGATATCGTTCGGAACACCAATAGCGTAGGCAGGCTTCTGGGTGCCACCTGACACTGCTAGGACGAAAGCGTGGGTAGCGATAAGG
## 289                                                          ACGGAGGGTGCGAGCGTTGTCCGGAATCACTGGGCGTAAAGGGCGCGTAGGCGGTCTGCTAAGCGTGCGGTGAAAGCCCGGGGCTCAACCCCGGGTCGGCCGTGCGAACTGGTGGACTAGAGCACTGTAGAGGCAGGTGGAATTCCGGGTGTAGCGGTGGAATGCGTAGAGATCCGGAAGAACACCGGTGGCGAAGGCGGCCTGCTGGGCAGTAGCTGACGCTGAGGCGCGACAGCGTGGGGAGCAAACAGG
## 290                                                          ACGTAGGGTGCGAGCGTTAATCGGAATTACTGGGCGTAAAGCGTGCGCAGGCGGTTTCGTAAGACAGAGGTGAAATCCCCGGGCTCAACCTGGGAACTGCCTTTGTGACTGCATAGCTGGAGTGCGGCAGAGGGGGATGGAATTCCGCGTGTAGCAGTGAAATGCGTAGATATGCGGAGGAACACCGATGGCGAAGGCAATCCCCTGGGCCTGCACTGACGCTCATGCACGAAAGCGTGGGGAGCAAACAGG
## 291                                                          ACGAAGGATGCAAGCGTTATCCGGATTCATTGGGTTTAAAGGGTGCGTAGGCGGATTAGTAAGTCAGTGGTGAAATCCCATCGCTTAACGATGGTACTGCCATTGATACTGCTAGTCTTGAGTACATATGACGTTGGCGGAATATGACATGTAGCGGTGAAATGCTTAGATATGTCATAGAACACCGATTGCGAAGGCAGCTAACGAAAATGTAACTGACGCTGATGCACGAAAGTGCGGGGATCAAACAGG
## 293                                                          ACGGAGGGTGCAAGCGTTATCCGGAATCATTGGGTTTAAAGGGTCCGCAGGCGGATTTATAAGTCAGTGGTGAAAGCCTACAGCTCAACTGTAGAACTGCCATTGATACTGTAAGTCTTGAATTCGGTCGAAGTGGGCGGAATATGACATGTAGCGGTGAAATGCTTAGATATGTCATAGAACACCGATAGCGAAGGCAGCTCACTAGGCCTGGATTGACGCTCAGGGACGAAAGCGTGGGGAGCAAACAGG
## 294                                                          ACGGAGGGTGCGAGCGTTGTCCGGAATCATTGGGTGTAAAGGGTGCGTAGGCGGGTTCTTAAGTCTGGGGTGAAAGTCTGCGGCTCAACCGTAGGATTGCCTTGGATACTGGGAGTCTTGAGTATGACAGAGGCCGGTAGAATTCGTGGTGTAGCGGTGAAATGCATAGATATCACGAAGAATACCAGTGGCGTAGGCGGCCGGCTGGGTCAACACTGACGCTGAGGCACGAGAGTGTGGGGAGCAAACAGG
## 295                                                          ACATAGGGTGCAAGCGTTGTCCGGAATTATTGGGCGTAAAGAGCTCGTAGGTGGTTTGATACGTCGGATGTGAAATTCAGGGGCTCAACCCCTGACCTGCATTCGATACGGTCAAGCTAGAGTTTGGTAGGGGAGACTGGAATTCCTGGTGTAGCGGTGGAATGCGCAGATATCAGGAGGAACACCGATGGCGAAGGCAGGTCTCTGGGCCAATACTGACACTGAGGAGCGAAAGCGTGGGGAGCGAACAGG
## 296                                                          ACGTAGGGTGCGAGCGTTAATCGGAATTACTGGGCGTAAAGCGTGCGCAGGCGGCTTTGCAAGACAGATGTGAAATCCCCGGGCTCAACCTGGGAACTGCATTTGTGACTGCAAGGCTAGAGTACGGCAGAGGGGGATGGAATTCCGCGTGTAGCAGTGAAATGCGTAGATATGCGGAGGAACACCGATGGCGAAGGCAATCCCCTGGGCCTGTACTGACGCTCATGCACGAAAGCGTGGGGAGCAAACAGG
## 297                                                          ACAGAGGGTGCAAGCGTTAATCGGAATTACTGGGCGTAAAGCGCGCGTAGGCGGTTTGTTAAGTCGGATGTGAAATCCCTGGGCTCAACCTAGGCACTGCATCCGATACTGGCTAACTAGAGTGTGGGAGAGGAAGGTAGAATTCCAGGTGTAGCGGTGAAATGCGTAGAGATCTGGAGGAATACCGATGGCGAAGGCAGCCTTCTGGCCTAACACTGACGCTGAGGTGCGAAAGCATGGGGAGCAAACAGG
## 298                                                          ACGTAGGGTGCGAGCGTTAATCGGAATTACTGGGCGTAAAGCGTGCGCAGGCGGTCTTGCAAGACAGGTGTGAAATCCCCGGGCTTAACCTGGGAACTGCACTTGTGACTGCAAGGCTGGAGTACGGCAGAGGGGGATGGAATTCCGCGTGTAGCAGTGAAATGCGTAGATATGCGGAGGAACACCGATGGCGAAGGCAATCCCCTGGGCCTGTACTGACGCTCATGCACGAAAGCGTGGGGAGCAAACAGG
## 299                                                          ACGTAGGTGACAAGCGTTATCCGGATTTACTGGGCGTAAAGGGCGTGTAGGCGGCTTAGCAAGTCAGATGTGAAATTCTTGGGCTCAACCCGAGAGCTGCATCTGAAACTGCTGAGCTTGAGTGCTGGATGGGATAGCGGAATTCCTGGTGGAGCGGTAAAATGCGCAGATATCAGGAAGAACACCGGTGGCGAAGGCGGCTATCTGGACAGTAACTGACGCTGAGGCGCGAAAGCGTGGGGAGCAAACAGG
## 300                                                          ACGTAGGGTGCGAGCGTTAATCGGAATTACTGGGCGTAAAGCGTGCGCAGGCGGTTATATAAGACAGATGTGAAATCCCCGGGCTTAACCTGGGAACTGCGTTTGTGACTGTATAACTCGAGTGTGGCAGAGGGGGGTGGAATTCCACGTGTAGCAGTGAAATGCGTAGAGATGTGGAGGAACACCGATGGCGAAGGCAGCCCCCTGGGTCAACACTGACGCTCATGCACGAAAGCGTGGGGAGCAAACAGG
## 301                                                          ACGTAGGGTGCAAGCGTTAATCGGAATTACTGGGCGTAAAGCGTGCGCAGGCGGTTTTATAAGACAGAGGTGAAATCCCCGGGCTCAACCTGGGAACTGCCTTTGTGACTGTAAGGCTAGAGTGTGTCAGAGGGGGGTAGAATTCCACGTGTAGCAGTGAAATGCGTAGATATGTGGAGGAATACCGATGGCGAAGGCAGCCCCCTGGGATAACACTGACGCTCATGCACGAAAGCGTGGGGAGCAAACAGG
## 302                                                          ACAGAGGTGGCAAGCGTTGTTCGGATTTATTGGGTGTAAAGGGCAGGTAGGCGGTCAAGTAAGTCTGTTGTGAAATCCCGGGGCTCAACCCCGGAACTGCAACGGAAACTACTTGGCTGGAGCAATGTAGGGGTGAGGGGAATTCTCGGTGTAAGGGTGAAATCTGTAGATATCGAGAGGAACACCAATGGCGAAGGCACCTCACTGGACATTTGCTGACGCTGAGCTGCGAAAGTAGGGGGAGCAAACAGG
## 303                                                          ACGTAGGGTGCAAGCGTTAATCGGAATTACTGGGCGTAAAGCGTGCGCAGGCGGCTTTGCAAGACAGATGTGAAATCCCCGGGCTCAACCTGGGAACTGCATTTGTGACTGCATGGCTAGAGTGCGGCAGAGGGGGATGGAATTCCGCGTGTAGCAGTGAAATGCGTAGATATGCGGAGGAACACCGATGGCGAAGGCAATCCCCTGGGCCTGCACTGACGCTCATGCACGAAAGCGTGGGGAGCAAACAGG
## 304                                                          ACAGAGACCTCAAGCGTTATCCGGATTCATTGGGCGTAAAGCGTCCGCAGGTGGTTTTTCAAGTTGGGAGTCAAATCTTTGGGCTTAACCTAGAGACTGCTTCCAATACTGGGAAACTTGAGACTGGGAGAGGTATACGGAACTGTTGGTGTAGTAGTAAAATGCGTTGATATCAACAGGAACACCAAAGGCGAAAGCAGTATACTGGAACAGTTCTGACACTCATGGACGAAAGCGTGGGGAGCGAATGGG
## 305                                                          ACGTAGGGTGCGAGCGTTAATCGGAATTACTGGGCGTAAAGCGTGCGCAGGCGGCTTTGCAAGACAGAGGTGAAATCCCCGGGCTCAACCTGGGAACTGCCTTTGTGACTGCAAGGCTAGAGTACGGCAGAGGGGGATGGAATTCCGCGTGTAGCAGTGAAATGCGTAGATATGCGGAGGAACACCAATGGCGAAGGCAATCCCCTGGGCCTGTACTGACGCTCATGCACGAAAGCGTGGGGAGCAAACAGG
## 306                                                          ACGTAGGGTGCAAGCGTTGTCCGGAATTATTGGGCGTAAAGAGCTCGTAGGCGGTTTGTCGCGTCTGCTGTGAAAATCCGAGGCTCAACCTCGGACCTGCAGTGGGTACGGGCAAGCTAGAGTGCGGTAGGGGAGATGGGAATTCCTGGTGTAGCGGTGGAATGCGCAGATATCAGGAGGAACACCAATGGCGAAGGCACATCTCTGGGCCGTAACTGACGCTGAGGAGCGAAAGCGTGGGGAGCGAACAGG
## 307                                                          ACGTAGGGTGCAAGCGTTAATCGGAATTACTGGGCGTAAAGCGTGCGCAGGCGGTTATATAAGTCAGATGTGAAATCCCCGGGCTCAACCTGGGAACTGCATTTGAGACTGTATAGCTAGAGTACGGTAGAGGGGGATGGAATTCCGCGTGTAGCAGTGAAATGCGTAGATATGCGGAGGAACACCGATGGCGAAGGCAATCCCCTGGGCCTGTACTGACGCTCATGCACGAAAGCGTGGGGAGCAAACAGG
## 308                                                          ACGAAGGGGGCTAGCGTTGTTCGGAATTACTGGGCGTAAAGCGCACGTAGGCGGGTCGTTAAGTCAGGGGTGAAATCCCGGAGCTCAACTCCGGAACTGCCTTTGATACTGGCGATCTAGAGATCGGAAGAGGTGAGTGGAATTCCCAGTGTAGAGGTGAAATTCGTAGATATTGGGAAGAACACCAGTGGCGAAGGCGGCTCACTGGTCCGATACTGACGCTGAGGTGCGAAAGCGTGGGGAGCAAACAGG
## 309                                                          ACGAAGGGTGCAAGCGTTAATCGGAATTACTGGGCGTAAAGGGTGCGTAGGTGGTTTATTAAGTCGGATGTGAAATCCCTGGGCTTAACCTAGGAACTGCATACGATACTGGTAGACTAGAGTGCGGTAGAGGATGGTGGAATTCCCGGTGTAGCGGTGAAATGCGTAGAGATCGGGAGGAACATCAGTGGCGAAGGCGGCCATCTGGACCAGCACTGACACTAAAGCACGAAAGCGTGGGGAGCAAACAGG
## 310                                                          ACGAGGGATCCAAGCGTTGTTCGGAATCATTGGGCGTAAAGCGGGTGTAGGTGGCTATATAAGTCAGATGTGAAAGCCCAGGGCTTAACCCTGGAAGTGCATTTGATACTGTTTAGCTTGAGTGTGGGAGAGGTTACTGGAATTCCTGGTGTAGTGGTGAAATACGTAGATATCAGGAGGAATACCGGTGGCGAAGGCGGGTAACTGGCCCAACACTGACACTGAGACCCGAAAGCGTGGGGATCAAACAGG
## 311                                                          ACGGAGGGTGCAAGCGTTATTCGGAATTACTGGGCGTAAAGCGCACGTAGGCCGCTTTGTAAGTCAGGGGTGAAATCCCACGGCTCAACCGTGGAACTGCCCTTGAAACTGCAGAGCTCGAATCCTGGAGAGGGTGGCGGAATTCCTGGTGTAGGAGTGAAATCCGTAGATATCAGGAGGAACACCGGTGGCGAAGGCGGCCACCTGGACAGGTATTGACGCTGAGGTGCGAAAGTGTGGGGAGCAAACAGG
## 312                                                          ACGAAGGTGGCAAGCGTTGTTCGGATTTACTGGGCGTAAAGAGTGCGTAGGCGGTTTAATAAGTCAGGAGTGAAAGCCCAAGGCTCAACCTTGGAATTGCTTTTGAAACTATTTTACTTGAATCCGGTAGAGGTTGGCGGAATTCCCAGTGTAGAGGTGAAATTCGCAGATATTGGGAGGAACACCAGTGGCGTAAGCGGCCAACTGGACCGGTATTGACGCTGAGGCACGAAAGCGTGGGTAGCAAACAGG
## 313                                                          ACGAAGGGTGCTAGCGTTGTTCGGAATGACTGGGCGTAAAGGGTGCGTAGGCGGCCAGAAGCGTCTGATGTGAAATCCCTGGGCTTAACCTAGGAACTGCATTGGATACGGTCTGGCTAGAGTCCGCGAGAGGAAGATGGAATTGTGCGTGTAGAGGTGAAATTCGTAGATATGCACAAGAACACCGGTGGCGAAGGCGATCTTCTGGAGCGGTACTGACGCTAAGGCACGAAAGTGTGGGGAGCAAACAGG
## 314                                                          ACGAAGGGTGCTAGCGTTGTTCGGAATGACTGGGCGTAAAGGGCGCGTAGGCGGTCTTTTAAGTTAGGCGTGAAAGCCCTGGGCTTAACCCAGGACGTGCGCTTAAGACTGGAAGGCTTGAGTTTGGAAGAGGAAAGTGGAATTCCTAGTGTAGAGGTGAAATTCGTAGATATTAGGAAGAACACCAGTGGCGAAGGCGACTTTCTGGTCCAATACTGACGCTGAGGCGCGAAAGTGTGGGGAGCAAACAGG
## 315                                                          ACGTAGGGTGCGAGCGTTAATCGGAATTACTGGGCGTAAAGCGTGCGCAGGCGGTTTTGAAAGTCAGATGTGAAATCCCCGGGCTCAACCTGGGAACTGCGTTTGAAACTCCAAAGCTAGAGTATGGGAGAGGGAGGTAGAATTCCACGTGTAGCAGTGAAATGCGTAGAGATGTGGAGGAATACCAATGGCGAAGGCAGCCTCCTGGCTTAATACTGACGCTCATGCACGAAAGCGTGGGGAGCAAACAGG
## 316                                                          ACATAGGGTGCAAGCGTTGTCCGGAATTATTGGGCGTAAAGAGCTCGTAGGTCGTTTGACACGTCGGATGTGAAAATCTGAGGCTTAACCTCAGACCTGCATTCGATACGGTCAAACTAGAGTGTGGTAGGGGAGACTGGAATTCCTGGTGTAGCGGTGGAATGCGCAGATATCAGGAGGAACACCAATGGCGAAGGCAGGTCTCTGGGCCATTACTGACACTGAGGAGCGAAAGCGCGGGGAGCGAACAGG
## 317                                                          ACAGAGGGTGCAAGCGTTAATCGGAATTACTGGGCGTAAAGCGTGCGTAGGCGGTTGTTTAAGTCGGATGTGAAATCCCCGGGCTCAACCTGGGAATTGCATTCGATACTGGACAGCTAGAGTTCGGCAGAGGGAAGTGGAATTTCCGGTGTAGCGGTGAAATGCGTAGATATCGGAAGGAACATCAGTGGCGAAAGCGACTTCCTGGACCAGAACTGACGCTCAGGCACGAAAGCGTGGGGAGCAAACAGG
## 318                                                          ACGTAGGATGCAAGCGTTATCCGGAATCATTGGGCGTAAAGCGTCCGCAGGTGGAACAGTAAGTTGAGCGATAAATACCAGTGCCTAACATTGGTGCCTCGTTCAAAACTGCTGATCTAGAGGATGGGAGGGGCAGACGGAATTCCCGGTGTAGGGGTAAAATCCGTAGATATCGGGAAGAACACCAAAAGCGAAGGCAGTCTGCTGGAACATTCCTGACACTCATGGACGAAAGCGTGGGGAGCGAAAAGG
## 319                                                          ACGTAGGGTGCGAGCGTTAATCGGAATTACTGGGCGTAAAGCGTGCGCAGGCGGCCTTAAAAGTCAGATGTGAAATCCCCGAGCTCAACTTGGGAACTGCGTTTGAAACTCTAAGGCTAGAATATGTCAGAGGGGGGTAGAATTCCACGTGTAGCAGTGAAATGCGTAGAGATGTGGAGGAATACCAATGGCGAAGGCAGCCCCCTGGGATAATATTGACGCTCATGCACGAAAGCGTGGGGAGCAAACAGG
## 320                                                          ACGGAGGATCCAAGCGTTATCCGGAATCATTGGGTTTAAAGGGTCCGTAGGCGGTTTGGTAAGTCAGTGGTGAAAGCCCGCAGCTCAACTGTGGAACGGCCATTGATACTGCCAGACTTGAATTACAAGGAAGTAACTAGAATATGTAGTGTAGCGGTGAAATGCTTAGATATTACATGGAATACCAATTGCGAAGGCAGGTTACTACTTGTGGATTGACGCTGATGGACGAAAGCGTGGGGAGCGAACAGG
## 321                                                          ACGTAGGGTGCAAGCGTTAATCGGAATTACTGGGCGTAAAGCGTGCGCAGGCGGTCATGCAAGACAGATGTGAAATCCCCGGGCTCAACCTGGGAACTGCATTTGTGACTGCATGGCTTGAGTACGGCAGAGGGGGATGGAATTCCGCGTGTAGCAGTGAAATGCGTAGATATGCGGAGGAACACCGATGGCGAAGGCAATCCCCTGGGCCTGTACTGACGCTCATGCACGAAAGCGTGGGGAGCAAACAGG
## 322                                                          ACGTAGGGTGCGAGCGTTAATCGGAATTACTGGGCGTAAAGCGTGCGCAGGCGGTTTTGCAAGACAGTGGTGAAATCCCCGGGCTCAACCTGGGAACGGCCATTGTGACTGCAAGGCTGGAGTGCGGCAGAGGGGGATGGAATTTCGCGTGTAGCAGTGAAATGCGTAGATATGCGAAGGAACACCGATGGCGAAGGCAATCCCCTGGGCCTGCACTGACGCTCATGCACGAAAGCGTGGGGAGCAAACAGG
## 323                                                          ACGTAGGGTGCAAGCGTTAATCGGAATTACTGGGCGTAAAGCGTGCGCAGGCGGTTATATAAGTCAGATGTGAAATCCCCGGGCTCAACCTGGGAACTGCATTTGAGACTGTATAGCTAGAGTACGGTAGAGGGGGATGGAATTCCGCGTGTAGCAGTGAAATGCGTAGATATGCGGAGGAACACCGATGGCGAAGGCAATCCCCTGGGCCTGCACTGACGCTCATGCACGAAAGCGTGGGGAGCAAACAGG
## 324                                                          ACGTAGGGTGCAAGCGTTAATCGGAATTACTGGGCGTAAAGCGTGCGCAGGCGGTTTTGTAAGACAGAGGTGAAATCCCCGGGCTCAACCTGGGAACTGCCTTTGTGACTGCAAAGCTGGAGTACGGCAGAGGGGGATGGAATTCCGCGTGTAGCAGTGAAATGCGTAGATATGCGGAGGAACACCGATGGCGAAGGCAATCCCCTGGGCCTGTACTGACGCTCATGCACGAAAGCGTGGGGAGCAAACAGG
## 325                                                          ACGTAGGGTGCAAGCGTTAATCGGAATTACTGGGCGTAAAGCGTGCGCAGGCGGTTTTGTAAGACAGAGGTGAAATCCCCGGGCTCAACCTGGGAACTGCCTTTGTGACTGCAAAGCTGGAGTGCGGCAGAGGGGGATGGAATTCCGCGTGTAGCAGTGAAATGCGTAGATATGCGGAGGAACACCGATGGCGAAGGCAATCCCCTGGGCCTGCACTGACGCTCATGCACGAAAGCGTGGGGAGCAAACAGG
## 326                                                          ACGAAGGGTGCAAGCGTTGTTCGGAATCATTGGGCGTAAAGCGCGCGCAGGCGGATCAGCAAGTCAGATGTGAAATCTCGAAGCTCAACTTCGAAACTGCGTCTGAAACTGCTAGTCTAGAATGTCGGAGGGGGCAGGGGAATTTCACGTGTAGGGGTAAAATCCGTAGAGATGTGAAGGAACACCGGAGGCGAAGGCGCCTGCCTGGACGACTATTGACGCTGAGGCGCGAAAGCGTGGGGAGCAAACAGG
## 327                                                          ACGGAGGATCCAAGCGTTATCCGGAATCATTGGGTTTAAAGGGTCCGTAGGCGGTCTTATAAGTCAGTGGTGAAAGCCCATCGCTCAACGATGGAACTGCCATTGATACTGTAAGACTTGAATGCTTAGGAAGTAACTAGAATATGTAGTGTAGCGGTGAAATGCTTAGATATTACATGGAATACCAATTGCGAAGGCAGGTTACTACTAAGTGATTGACGCTGATGGACGAAAGCGTGGGGAGCGAACAGG
## 328                                                          ACGAAGGGTGTAAGCGTTGTTCGGATTTATTGGGCGTAAAGCGCGCGCAGGTGGATTTTTAAGTCAGATGTGAAATCTCGGGGCTCAACCCCGAACGTGCGTCTGAAACTGGAGATCTAGAATATTGGAGGGGGTAGAGGAATTTCACATGTAGGGGTAAAATCCGTAGAGATGTGAAGGAACACCAGAGGCGAAGGCGTCTACCTGGCCAACCATTGACACTGAGGCGCGAAAGCGTGGGGAGCAAACAGG
## 329                                                          ACGAAGGGTGCGAGCGTTGTTCGGAATGACTGGGCGTAAAGGGTTCGTAGGCGGGAGTGCAAGTCAAGTGTGAAATCCCCAGGCTTAACCTGGGACGTGCACTTGAAACTGCATTTCTTGAGTCGAGGAGAGGGTAATGGAATTGCTGGTGTAGGAGTGACATCCGTAGAGATCAGCAGGAACACCGGAGGCGAAGGCGATTACCTGGCCGAAGACTGACGCTGAGGAACGAAAGCGTGGGGAGCAAACAGG
## 330                                                          ACGGAGGATCCAAGCGTTATCCGGATTTATTGGGTTTAAAGGGTGCGTAGGCGGCTTATTAAGTCAGGGGTGAAAGACGGTGGCTCAACCATCGCAGTGCCCTTGATACTGATGAGCTTGAATACACTAGAGGTAGGCGGAATGAGACAAGTAGCGGTGAAATGCATAGATATGTCTCAGAACACCAATTGCGAAGGCAGCTTACTATGGTGTTATTGACGCTGAGGCACGAAAGCGTGGGGATCAAACAGG
## 331                                                          ACGGAGGGTGCAAGCGTTACTCGGAATCACTGGGCGTAAAGAGCGTGTAGGCGGGTGAATAAGTCAGAAGTGAAATCCAATAGCTCAACTATTGAACTGCTTTTGAAACTGTATACCTAGAATGTGGGAGAGGCAGATGGAATTTCTGGTGTAGGGGTAAAATCCGTAGAGATCAGAAGGAATACCGATTGCGAAGGCGATCTGCTGGAACATTATTGACGCTGAGACGCGAAAGCGTGGGGAGCAAACAGG
## 332                                                          ACGAAGGTGGCGAGCGTTACTCGGATTTACTAGGCGTAAAGCGTGGGCAGGCGGCTCGATTAGTCTCTTGTGAAAGCCTTGGGCTTAACCCAGGGAGGCCAAGAGATACTATCGGGCTTGGATGTGGGAGAGGAGACTGGAATTCCCGGTGTAGCGGTGAAATGCGTTGATATCGGGAGGAACACCAATGGCGAAAGCAAGTCTCTGGACCACCATCGACGCTCATCCACGAAAGCTGGGGGATCAAACAGG
## 333                                                          ACAGAGACCTCAAGCGTTATCCGGATTTATTGGGCGTAAAGCGCAGGTAGGCGGGTTTATTAGTCGGATGTCAAAGACCGGGGCTCAACCCCGGGAAGGCATTCGAAACGGTAAACCTAGAGAATGTGAGAGATGAGTGGAATTCATGGTGTAGTAGTGAAATGCGTTGATATCATGAGGAACACCGAAGGCGAAGGCAACTCATTGGCACATTTCTGACGCTGAGCTGCGAAAGCGTGGGGAGCGAATGGG
## 334                                                          ACGGAGGGTGCAAGCGTTATCCGGATTTATTGGGTTTAAAGGGTCCGTAGGCGGATCGTTAAGTCAGTGGTGAAAGCCTACCGCTCAACGGTAGAACTGCCATTGATACTGGCGATCTTGAATATGTATGAAGTTGGCGGAATGTGTGGTGTAGCGGTGAAATGCTTAGATATCACACAGAACACCGATTGCGAAGGCAGCTGACTAATACATGATTGACGCTGAGGGACGAAAGTGTGGGGATCAAACAGG
## 335                                                          ACGTAGGATGCAAGCGTTGTCCGGATTTATTGGGCGTAAAGAGTTCGTAGGTGGTTTGTTAAGTTTGGTGTTAAAGATTGGGGCTCAACCCTGAAACTGCACTGAATACTGGCAGACTCGAGTGTGGTAGAGGCTAGTGGAATTCCCAGTGTAGCGGTGAAATGCGTAGATATTGGGAAGAACACCGGTGGCGTAGGCGACTAGCTGGGCCATAACTGACACTGAGGAACGAAAGCCAGGGGAGCGAATGGG
## 336                                                          ACGTAGGATGCAAGCGTTGTCCGGATTTATTGGGCGTAAAGAGTTCGTAGGTGGTTTGTCAAGTCTGATGTTAAAGATCGGGGCTCAACCCTGGGACTGCATTGGATACTGGCAGACTTGAGTATGGTAGAGGCTAGTGGAATTCCCAGTGTAGCGGTGAAATGCGTAGATATTGGGAAGAACACCGGTGGCGTAGGCGACTAGCTGGGCCATAACTGACGCTGAGGAACGAAAGCCAGGGGAGCGAATGGG
## 337                                                          ACGGAGGATCCGAGCGTTATCCGGATTTATTGGGTTTAAAGGGTGCGCAGGCGGCTTTTTAAGTTGCCGGTGAAATGTCGGGGCTCAACTCCGTCACTGCCGGCGATACTGGAAGGCTTGAGAATGGAGGAGGATTCCGGAATGTGTGGTGTAGCGGTGAAATGCATAGATATCACACAGAACACCGATTGCGAAGGCAGGAGTCCACACCATATCTGACGCTGAGGCACGAAAGCGTGGGGATCGAACAGG
## 338                                                         ACGTAGGCACCAAGCGTTGTCCGGATTTATTGGGCGTAAAGAGCTCGTAGGCGGTTCAGCAAGTCGGGTGTGAAAACTCTGGGCTTAACCCAGAGCCTGCACCCGAAACTGCTGTGACTAGAGTTCGGTAGGGGAGCGGGGAATTCCTGGTGTAGCGGTGAAATGCGCAGATATCAGGAGGAACACCGGTGGCGAAGGCGCCGCTCTGGGCCGAAACTGACGCTGAGGAGCGAAAGCGTGGGTAGCAAACAGG
## 339                                                          ACGTAGGGTGCAAGCGTTAATCGGAATTACTGGGCGTAAAGCGTGCGCAGGCGGCTTTGCAAGACAGATGTGAAATCCCCGGGCTCAACCTGGGAACTGCATTTGAGACTGTATAGCTAGAGTACGGTAGAGGGGGATGGAATTCCGCGTGTAGCAGTGAAATGCGTAGATATGCGGAGGAACACCGATGGCGAAGGCAATCCCCTGGACCTGTACTGACGCTCATGCACGAAAGCGTGGGGAGCAAACAGG
## 340                                                          ACGTAGGGTGCGAGCGTTAATCGGAATTACTGGGCGTAAAGCGTGCGCAGGCGGTTTTGTAAGTCAGACGTGAAAGCCCCGGGCTTAACCTGGGAATTGCGTTTGAAACTGCAAGGCTAGAGTACGGCAGAGGGGGGTAGAATTCCACGTGTAGCAGTGAAATGCGTAGATATGTGGAGGAATACCGATGGCGAAGGCAGCCCCCTGGGCTAGTACTGACGCTCATGCACGAAAGCGTGGGGAGCAAACAGG
## 341                                                          ACGGGAGTGGCAAGCGTTATCCGGAATTATTGGGCGTAAAGCGTCCGCAGGCGGTCTTGTAAGTCTGTCGTTAAAGCGTGGAGCTTAACTCCATTTCAGCGATGGAAACTGCAAGACTAGAGTGTGGTAGGGGCAGAGGGAATTCCCGGTGTAGCGGTGAAATGCGTAGATATCGGGAAGAACACCAGTGGCGAAGGCGCTCTGCTGGGCCATAACTGACGCTCATGGACGAAAGCCAGGGGAGCGAAAGGG
## 342                                                          ACGAGGGATCCAAGCGTTGTTCGGAATCATTGGGCGTAAAGCGGGTGTAGGCGGCTCTATAAGTCAGATGTGAAAGCCCTGGGCTCAACCCGGGAAGTGCATTTGATACTGCAGAGCTTGAGTGTGGGAGAGGCTAGTGGAATTCCTGGTGTAGTGGTGAAATACGTAGATATCAGGAGGAACATCGGTGGCGAAGGCGGCTAGCTGGCCCAACACTGACGCTGAGATCCGAAAGCGTGGGGATCAAACAGG
## 343                                                          ACGTAGGTGACAAGCGTTATCCGGATTTACTGGGTGTAAAGGGCGTGTAGGCGGTTTTGCAAGTCAGATGTGAAATTCCCAGGCTTAACCTGGGCGCTGCATCTGAAACTGCAAGACTTGAGTACTGGAGAGGATAGTGGAATTCCTAGTGTAGCGGTAAAATGCGCAGATATTAGGAGGAACACCGGTGGCGAAGGCGACTATCTGGACAGTAACTGACGCTGAGGCGCGAAAGCGTGGGGAGCAAACAGG
## 344                                                          ACAGAGACTGCAAGCGTTATTCGGATTCACTGGGCGTAAAGGGTGCGCAGGCGGCCAAGTGTGTGAGGCGTGAAAGCCCGAGGCTCAACCTCGGAATTGCACCTCAAACTACATGGCTAGAGCATTGGAGAGGGTAGCAGAATTCATGGTGTAGCAGTGAAATGCGTAGATATCATGAGGAATACCAGAGGCGAAGGCGGCTACCTGGACAATTGCTGACGCTCAGGCACGAAAGCGTGGGGAGCAAAAGGG
## 345                                                          ACGAAGGGTGCTAGCGTTACTCGGAATTATTGGGCGTAAAGCGTGCGCAGGCGGTATTGTAAGTTAAAAGTGAAAGCCTCAGGCTCAACCTGAGAATGGCTTTTAATACTGCAGTACTAGAGTATCAGAAAAGATGGCGGAATTCCTAGTGTAGAGGTGAAATTCGTAGATATTAGGAGGAACACCAGTGGCGAAAGCGGCTATCTAGCTGAATACTGACGCTGATGCACGAAAGCGTGGGGAGCAAACAGG
## 346                                                          ACAGAGGGTGCAAGCGTTAATCGGAATTACTGGGCGTAAAGCGCGCGTAGGTGGTTTTGTAAGTTGGAGGTGAAATCCCCGGGCTCAACCTGGGAACTGCCTCCAAAACTGCATAGCTAGAGTACGGTAGAGGGTGGTGGAATTTCCTGTGTAGCGGTGAAATGCGTAGATATAGGAAGGAACACCAGTGGCGAAGGCGACCACCTGGACTGATACTGACACTGAGGTGCGAAAGCGTGGGGAGCAAACAGG
## 347                                                          ACGAGTGCCTCAAGCGTTATCCGGAATCATTGGGCGTAAAGGTTGTGTAGGTGGTTATATTAGTCTTTTGTTAAATTCTTCGGCTTAACCGGGGGCATGCAAAGGAAACGGTATGACTAGAGGATGCGAGGGGTCTGCGGAACTCATAGTGTAGCGGTAAAATGCGTTGATATTATGGGGAACACCAAAAGCGAAGGCAGCAGACTGGAGCACTCCTGACACTGAAACAAGAAAGCGTGGGTCGCGAATGGG
## 348                                                          ACGGAGGGAGCTAGCGTTGTTCGGAATTACTGGGCGTAAAGAGTACGTAGGCGGTTATTCAAGTCAGAGGTGAAAGCCTGGAGCTCAACTCCAGAACTGCCTTTGAAACTAGATAGCTAGAATCTTGGAGAGGTGAGTGGAATTCCGAGTGTAGAGGTGAAATTCGTAGATATTCGGAAGAACACCAGTGGCGAAGGCGACTCACTGGACAAGTATTGACGCTGAGGTACGAAAGCGTGGGGAGCAAACAGG
## 349                                                          ACGAAGGGTGCTAGCGTTGTTCGGAATCATTGGGCGTAAAGCGTACGTAGGCGGTTTAGTAAGTCGATTGTGAAATACCCGAGCTCAACTCAGGAATTGCAATCGAAACTGCTTTGCTAGAGTTTAGTGGGGGATAGTGGAATTCCTAGTGTAGGGGTGAAATCCGTAGAGATTAGGAGGAACATCAGTGGCGAAGGCGACTATCTACGCTAATACTGACGCTAAGGTACGAAAGCGTGGGGAGCAAACAGG
## 350                                                          ACGAGTGCCTCAAGCGTTATCCGGAATCATTGGGCGTAAAGGTTGTGTAGGTGGTCATATTAGTCTTTTGTTAAATTCTTCGGCTTAACCGGGGGCATGCAAGAGAAACGGTATGACTAGAGGATGCGAGGGGTCTGCGGAATTCATAGTGTAGCGGTAAAATGCGTTGATATTATGGAGAACACCAAAAGCGAAGGCAGCAGACTGGAGCACTCCTGACACTGAAACAAGAAAGCGTGGGTCGCGAATGGG
## 351                                                          ACGTAGGGTGCAAGCGTTAATCGGAATTACTGGGCGTAAAGCGTGCGCAGGCGGTGATGTAAGACAGGCGTGAAATCCCCGGGCTTAACCTGGGAACTGCGCTTGTGACTGCATCGCTGGAGTGCGGCAGAGGGGGATGGAATTCCGCGTGTAGCAGTGAAATGCGTAGATATGCGGAGGAACACCGATGGCGAAGGCAATCCCCTGGGCCTGCACTGACGCTCATGCACGAAAGCGTGGGGAGCAAACAGG
## 352                                                          ACGTAGGGTGCGAGCGTTAATCGGAATTACTGGGCGTAAAGCGTGCGCAGGCGGTTATGTAAGACAGATGTGAAATCCCCGGGCTCAACCTGGGAACTGCGTTTGTGACTGCATAACTAGAGTACGGCAGAGGGAGGTGGAATTCCACGTGTAGCAGTGAAATGCGTAGAGATGTGGAGGAACACCGATGGCGAAGGCAGCCTCCTGGGCCAGTACTGACGCTCATGCACGAAAGCGTGGGGAGCAAACAGG
## 353                                                          ACGTGAGCCCCAAGCGTTATCCGGAATTACTGGGCGTAAAGAGTGCGTAGGCGGTTATGTTAGTCTTCTGTTTAATCCATCGGCTCAACCGGTGAACCGCAGAGGAAACGGCATGACTGGAGTGTGTGAGAGGTTTGCGGAACTCATGGTGTAGGGGTGAAATCCGTTGATATCATGGGGAACACCAAATGCGAAGGCAGCAAACTGGCACATTACTGACGCTGAGGCACGAAAGCGTGGGTAGCGAATGGG
## 354                                                          ACGAAGGATTCAAGCGTTGTCCGGATTTACTGGGTTTAAAGGGTGCGTAGGCGGAAAGCTAAGTCAGTGGTGAAAGCCCGCAGCTCAACTGCGGAACTGCCATTGAAACTGGCCTTCTTGAATACAGTGGAGGTAGATGGAATATAACATGTAGCGGTGAAATGCTTAGATATGTTATAGAACACCGATTGCGAAGGCAGTCTGCTACACTGTCATTGACGCTGAGGCACGAAAGCGTGGGGAGCGAACAGG
## 355                                                          ACGTAGGGTGCGAGCGTTAATCGGAATTACTGGGCGTAAAGGGTGCGCAGGCGGATATATAAGTTAGATGTGAAATACCTGGGCTTAACCTAGGAACGGCATTTAATACTGTATATCTAGAGTTTGTCAGAGGGTGGTAGAATTCCAAGTGTAGCAGTGAAATGCGTAGAGATTTGGAGGAATACCGGTGGCGAAGGCGGCCACCTGGGATAAAACTGACGCTCATGCACGAAAGCGTGGGGAGCAAACAGG
## 356                                                          ACGTAGGGTGCAAGCGTTAATCGGAATTACTGGGCGTAAAGCGTGCGCAGGCGGTTGTGCAAGACAGGTGTGAAATCCCCGGGCTTAACCTGGGAACTGCACTTGTGACTGCACAGCTGGAGTACGGCAGAGGGGGATGGAATTCCGCGTGTAGCAGTGAAATGCGTAGATATGCGGAGGAACACCGATGGCGAAGGCAATCCCCTGGGCCTGTACTGACGCTCATGCACGAAAGCGTGGGGAGCAAACAGG
## 357                                                          ACGGAGGGTCCGAGCGTTATCCGGAATTATTGGGTTTAAAGGGTCCGTAGGCGGGCTATTAAGTCAGGGGTGAAAGTTTGCAGCTCAACTGTAAAATTGCCTTTGATACTGGTAGTCTTGAGTTGTATTGAAGTGGCTGGAATATGTAGTGTAGCGGTGAAATGCATAGATATTACATAGAACACCGATTGCGAAGGCAGGTCACTAAGTACACACTGACGCTGATGGACGAAAGCGTGGGGAGCGAACAGG
## 358                                                          ACGTAGGGTGCAAGCGTTAATCGGAATTATTGGGCGTAAAGGGTGCGCAGGCTGTGCTGTAAGTCAGATGTGAAATCCCCGGGCTTAACCTGGGAATTGCGTTTGAAACTGCAGTGCTAGAGTGTGATAGAGGAGAGTGGAATTCCATGTGTAGCAGTGAAATGCGTAGAGATGTGGAAGAACACCAATGGCGAAGGCAGCTCTCTGGATTAACACTGACGCTCATGCACGAAAGCGTGGGGAGCAAACAGG
## 359                                                          ACGAAGGTGGCGAGCGTTACTCGGAATTACTAGGCGTAAAGCGTGGGCAGGCGGCTTGATAAGTCTTTTGTGAAAGCCTTGGGCTTAACCCAGGGAGGCCAGAAGATACTGTCAGGCTTGGATGTGGGAGAGGAGACTGGAATTCCCGGTGTAGCGGTGAAATGCGTTGATATCGGGAGGAACACCAATGGCGAAAGCAAGTCTCTGGACCACCATCGACGCTCATCCACGAAAGCTGGGGGATCAAACAGG
## 360                                                          ACGTAGGGTGCGAGCGTTAATCGGAATTACTGGGCGTAAAGCGTGCGCAGGCGGTTGGGTAAGTCAGATGTGAAATCCCCGGGCTCAACCTGGGAACTGCATTTGAGACTGCCTAGCTGGAGTTTGGCAGAGGGGGGTGGAATTCCACGTGTAGCAGTGAAATGCGTAGAGATGTGGAGGAACACCGATGGCGAAGGCAGCCCCCTGGGCCAATACTGACGCTCATGCACGAAAGCGTGGGGAGCAAACAGG
## 361                                                          ACGGAGGGTGCGAGCGTTAATCGGAATTACTGGGCGTAAAGCGCGCGTAGGCGGTCGGCTAAGTCTGCTGTGAAAGCCCTGGGCTTAACCTGGGAACTGCAGTGGATACTGGTCGGCTAGAGTATGGTAGAGGAGAGTGGAATTTCCGGTGTAGCAGTGAAATGCGTAGAGATCGGAAGGAACACCAGTGGCGAAGGCGGCTCTCTGGACCAATACTGACGCTGAGGTGCGAAAGCGTGGGGAGCAAACAGG
## 362                                                          ACGGAGGGTGCAAGCGTTAATCGGAATAACTGGGCGTAAAGCGCACGCAGGCGGTTATATAAGTCAGATGTGAAAGCCCCGGGCTCAACCTGGGAACTGCATTTGAAACTGTGTAACTAGAGTCTTGTAGAGGGGGGTAGAATTCCAGGTGTAGCGGTGAAATGCGTAGAGATCTGGAGGAATACCGGTGGCGAAGGCGGCCCCCTGGACAAAGACTGACGCTCAGGTGCGAAAGCGTGGGGAGCAAACAGG
## 363                                                          ACGGAAGGTGCAAGCGTTAATCGGAATTACTGGGCGTAAAGCGCGCGTAGGCGGCTTGGTAAGTCGGATGTGAAAGCCCCGGGCTTAACCTGGGAACTGCATTCGATACTGCTGGGCTGGAGTACGAGAGAGGGGGGTGGAATTCCAGGTGTAGCGGTGAAATGCGTAGAGATCTGGAGGAACATCGGTGGCGAAGGCGGCCCCCTGGCTCGATACTGACGCTGAGGTGCGAAAGCGTGGGGAGCAAACAGG
## 364                                                          ACGGAGGGTGCAAGCGTTACTCGGAATCACTGGGCGTAAAGGACGCGTAGGCTGTTTAGCAAGTCAGATGTGAAATCCAATGGCTCAACCATTGAACTGCATTTGAAACTGCTAGACTAGAGTATGGGAGGGGGAGATGGAATTAGTGGTGTAGGGGTAAAATCCGTAGATATCACTAGGAATATCTAAAGCGAAGGCGATCTCCTGGAACATTACTGACGCTAAGGCGTGAAAGCGTGGGGAGCAAACAGG
## 365                                                          ACGGAGGGTGCAAGCGTTATCCGGATTTACTGGGTTTAAAGGGTGCGTAGGCGGGCAGGTAAGTCAGTGGTGAAATCCCCGAGCTTAACTCGGGAACTGCCGTTGATACTATCTGTCTTGAATATCGTGGAGGTAAGCGGAATATGTCATGTAGCGGTGAAATGCTTAGATATGACATAGAACACCAATTGCGAAGGCAGCTTGCTACACGATTATTGACGCTGAGGCACGAAAGCGTGGGGATCAAACAGG
## 366                                                          ACAGAGGATGCAAGCGTTATCCGGAATCACTGGGCATAAAGCGTCTGTAGGTGGTTTGATAAGTCTGCTGTTAAAGACTAGGGCTTAACCCTAGGACAGCAGTGGAAACTATTAGACTTGAGTATGGTAGAGGTAGAGGGAATTCCTAGTGTAGCGGTGAAATGCGTAGATATTAGGAAGAACACCAATGGCGAAAGCACTCTACTGGGCCACAACTGACACTGAGAGACGACAGCTAGGGGAGCAAATGGG
## 367                                                          ACGTAGGGTGCGAGCGTTAATCGGAATTACTGGGCGTAAAGCGTGCGCAGGCGGTTTGGTAAGACAGGCGTGAAATCCCCGGGCTCAACCTGGGAACTGCGCTTGTGACTGCCTCACTAGAGTACGGCAGAGGGGGGTGGAATTCCACGTGTAGCAGTGAAATGCGTAGAGATGTGGAGGAACACCGATGGCGAAGGCAGCCCCCTGGGCCGATACTGACGCTCATGCACGAAAGCGTGGGTAGCAAACAGG
## 368                                                          ACAGAGGGTGCAAGCGTTAATCGGAATTACTGGGCGTAAAGCGCGCGTAGGCGGTTTGTTAAGTCGGATGTGAAATCCCCGGGCTCAACCTGGGCACTGCATCCGATACTGGCTGACTAGAGTGTGGGAGAGGAAGGTAGAATTCCAGGTGTAGCGGTGAAATGCGTAGAGATCTGGAGGAATACCGATGGCGAAGGCAGCCTTCTGGCCTAACACTGACGCTGAGGTGCGAAAGCATGGGGAGCAAACAGG
## 369                                                          ACGTAGGGTGCGAGCGTTAATCGGAATTACTGGGCGTAAAGCGTGCGCAGGCGGTTTAGTAAGACAGGCGTGAAATCCCCGGGCTCAACCTGGGAACTGCGCTTGTGACTGCTAAGCTAGAGTACGGCAGAGGGGGGTGGAATTCCACGTGTAGCAGTGAAATGCGTAGAGATGTGGAGGAACACCGATGGCGAAGGCAGCCCCCTGGGCCGATACTGACGCTCATGCACGAAAGCGTGGGTAGCAAACAGG
## 370                                                          ACGTAGGGAGCGAGCGTTGTCCGGAATTACTGGGTGTAAAGGGAGTGTAGGCGGGACGGTAAGTCAGGTGTGAAATTTATGGGCTCAACTCATAAACTGCACTTGAAACTGCTGTTCTTGAGTGAAGTAGAGGTTGGCGGAATTCCTAGTGTAGCGGTGAAATGCGTAGATATTAGGAGGAACATCAGTGGCGAAGGCGGCCAACTGGGCTTTTACTGACGCTGAGGCTCGAAAGCGTGGGGAGCAAACAGG
## 371                                                           ACGTAGGGGGCAAGCGTTATCCGGAATTACTGGGCGTAAAGGGTGCGTAGGCGGTTTATCAAGTCAGGGGTGAAAGGCTACGGCTCAACCGTAGTGAGCCTCTGAAACTGGTGAACTTGAGTGTAGGAGAGGAAAGTGGAATTCCCAGTGTAGCGGTGAAATGCGTAGATATTGGGAGGAACACCAGTGGCGAAGGCGACTTTCTGGACTACAACTGACGCTGAGGCACGAAAGCGTGGGGAGCAAACAGG
## 373                                                          ACGAAGGTGGCAAGCGTTGTTCGGATTTACTGGGCGTAAAGAGTGCGTAGGCGGTTTAACAAGTCAGGAGTGAAATCCCGAGGCTCAACCTTGGAATTGCTTTTGAAACTGTTTTACTTGAGTCCGGGAGAGGTTGGCGGAATTCCGAGTGTAGAGGTGAAATTCGCAGATATTCGGAGGAACACCAGTGGCGTAAGCGGCCAACTGGACCGGTACTGACGCTGAGGCACGAAAGCGTGGGGAGCAAACAGG
## 374                                                          ACGTAGGGTGCAAGCGTTAATCGGAATTACTGGGCGTAAAGCGTGCGCAGGCGGTTTTGTAAGACTGTCGTGAAATCCCCGGGCTCAACCTGGGAATGGCGATGGTGACTGCAAGGCTAGAGTTTGGCAGAGGGGGGTAGAATTCCACGTGTAGCAGTGAAATGCGTAGATATGTGGAGGAACACCGATGGCGAAGGCAGCCCCCTGGGTCAAAACTGACGCTCATGCACGAAAGCGTGGGGAGCAAACAGG
## 375                                                          ACAGAGGGTGCGAGCGTTAATCGGAATTACTGGGCGTAAAGGGTGCGTAGGTGGTTTTCTAAGTTGTTTGTGAAATATCCGGGCTTAACCTGGGTGCTGCGAACAATACTGGGAGACTAGAGTAGGGTAGAGGGAAGCGGAATTTCCGGTGTAGCAGTGAAATGCGTAGATATCGGAAGGAACACCAGTGGCGAAGGCGGCTTCCTGGACTCATACTGACACTGAGGCACGAAAGCGTGGGGATCAAACAGG
## 376                                                          ACATAGGGTGCAAGCGTTGTCCGGAATTATTGGGCGTAAAGAGCTCGTAGGTGGTTTGTTACGTCGGATGTGAAATTCAGGGGCTCAACCCCTGACTTGCATCCGATACGGGCAAGCTAGAGTTTGGTAGGGGAGACTGGAATTCCTGGTGTAGCGGTGGAATGCGCAGATATCAGGAGGAACACCGATGGCGAAGGCAGGTCTCTGGGCCAATACTGACACTGAGGAGCGAAAGCGTGGGGAGCGAACAGG
## 377                                                          TCGAGGAAGGCAAGCGTTATCCGGAATCATTGGGCGTAAAGCGTGCGCAAGCGGTTATATAAGTGAGGTGTCAAATTCCAACGCTCAACGTTGGACTCGCGCTTCATACTGTATGACTTGAGAATAGTAGAGGCAAACGGAATTCCCGGTGTAGTAGTGAAATGCGTTAATATCGGGAGGAACGCCAAAAGCGAAAGCAGTTTGCTGGGCTATTTCTGACGCCGTTGCACGACAGCGTGGGGAGCAAACAGG
## 378                                                           ACGTAGGGGGCTAGCGTTATCCGGAATTACTGGGCGTAAAGGGTGCGTAGGTGGTTTCTTAAGTCAGAGGTGAAAGGCTACGGCTCAACCGTAGTAAGCCTTTGAAACTGGGAAACTTGAGTGCAGGAGAGGAGAGTGGAATTCCTAGTGTAGCGGTGAAATGCGTAGATATTAGGAGGAACACCAGTTGCGAAGGCGGCTCTCTGGACTGTAACTGACACTGAGGCACGAAAGCGTGGGGAGCAAACAGG
## 379                                                          ACGGGGGATGCAAGCGTTATCCGGAATGATTGGGCGTAAAGCGCCTGTAGGTTGTTTATTAAGTCCGTTGTTAAAGCCTAGGGCTCAACCCTAGAAAAGCAATGGAAACTAGTAGACTAGAGTATGGCAGGGGTAGAGGGAATTTCTAGTGTAGCGGTGAAATGCGTAGATATTAGAAAGAACACCGGTGGCGAAAGCGCTCTACTGGACCATTACTGACACTCAGAGGCGAAAGCTAGGGTAGCAAAAGGG
## 380                                                          ACGTAGGGTGCAAGCGTTAATCGGAATTACTGGGCGTAAAGCGTGCGCAGGCGGTCATGCAAGACAGATGTGAAATCCCCGGGCTCAACCTGGGAACTGCATTTGTGACTGCATGGCTTGAGTGCGGCAGAGGGGGATGGAATTCCGCGTGTAGCAGTGAAATGCGTAGATATGCGGAGGAACACCGATGGCGAAGGCAATCCCCTGGGCCTGCACTGACGCTCATGCACGAAAGCGTGGGGAGCAAACAGG
## 381                                                          ACGAAGGATGCAAGCGTTATCCGGATTCATTGGGTTTAAAGGGAGCGTAGGTGGACTGATAAGTCAGTGGTGAAATCTTCGAGCTTAACTCGGAAATTGCCATTGATACTGTCGGTCTTGAGTACAGTTGCTGTGGGCGGAATATGACATGTAGTGGTGAAATACATAGAGATGTCATAGAACACCGATTGCGAAGGCAGCTCACAAAGCTGTAACTGACACTGAGGCTCGAAAGTGTGGGGATCAAACAGG
## 382                                                          ACAGAGGGTGCGAGCGTTAATCGGATTTACTGGGCGTAAAGCGCGCGTAGGTGGTTAATTAAGTCAAATGTGAAATCCCCGAGCTTAACTTGGGAATTGCATTCGATACTGGTTAGCTAGAGTATGGGAGAGGATGGTAGAATTCCAGGTGTAGCGGTGAAATGCGTAGAGATCTGGAGGAATACCGATGGCGAAGGCAGCCATCTGGCCTAATACTGACACTGAGGTGCGAAAGCATGGGGAGCAAACAGG
## 383                                                          ACGGAGGATCCAAGCGTTATCCGGATTTATTGGGTTTAAAGGGTGCGTAGGTGGATTAATAAGTCAGTGGTGAAAGCCTGCAGCTTAACTGTAGAACTGCCATTGATACTGTTAGTCTTGAATGTAGTTGAGGTGGGCGGAATATGACATGTAGCGGTGAAATGCTTAGATATGTCATAGAACACCGATTGCGAAGGCAGCTCACTAAGCTATAATTGACACTGAGGCACGAAAGCGTGGGGAGCAAACAGG
## 384                                                          ACGGGAGTGGCAAGCGTTATCCGGAATTATTGGGCGTAAAGCGTCCGCAGGCGGTCTTGTAAGTCTGTCGTTAAAGCGTGGAGCTTAACTCCATTTGAGCGATGGAAACTGCAAGACTAGAGTGTGGTAGGGGCAGAGGGAATTCCCGGTGTAGCGGTGAAATGCGTAGATATCGGGAAGAACACCAGTGGCGAAGGCGCTCTGCTGGGCCATAACTGACGCTCATGGACGAAAGCCAGGGGAGCGAAAGGG
## 385                                                          ACGTAGGGTGCAAGCGTTAATCGGAATTACTGGGCGTAAAGCGTGCGCAGGCGGTTCTATAAGACAGAGGTGAAATCCCCGGGCTCAACCTGGGAACTGCCTTTGTGACTGTAGGGCTAGAGTGCGGCAGAGGGGGATGGAATTCCGCGTGTAGCAGTGAAATGCGTAGATATGCGGAGGAACACCGATGGCGAAGGCAATCCCCTGGGCCTGCACTGACGCTCATGCACGAAAGCGTGGGGAGCAAACAGG
## 386                                                          ACGGAGGGTGCAAGCGTTATCCGGAATCATTGGGTTTAAAGGGTCCGCAGGCGGATTTATAAGTCAGTGGTGAAATCTCACAGCTCAACTGTGAAACTGCCATTGATACTGTAAGTCTTGAATTCGGTCGAAGTGGGCGGAATATGACATGTAGCGGTGAAATGCTTAGATATGTCATAGAACACCGATAGCGAAGGCAGCTCACTAGGCCTGGATTGACGCTCAGGGACGAAAGCGTGGGGAGCAAACAGG
## 387                                                          ACGTAGGGTGCAAGCGTTAATCGGAATTACTGGGCGTAAAGCGTGCGCAGGCGGTTTTGTAAGACAGGCGTGAAATCCCCGGGCTCAACCTGGGAATGGCGCTTGTGACTGCAAGGCTAGAGTGCGGCAGAGGGGGATGGAATTCCGCGTGTAGCAGTGAAATGCGTAGATATGCGGAGGAACACCGATGGCGAAGGCAATCCCCTGGGCCTGCACTGACGCTCATGCACGAAAGCGTGGGGAGCAAACAGG
## 388                                                          ACAGAGGATGCAAGCGTTATCCGGAATCACTGGGCATAAAGCGTCTGTAGGTGGTTTGGTAAGTCTGCTGTTAAAGACTGGGGCTCAACCCCAGAAAAGCAGTGGAAACTGCTAGACTTGAGTGTGGTAGAGGTAGAGGGAATTCCTAGTGTAGCGGTGAAATGCGTAGATATTAGGAAGAACACCAATGGCGAAGGCACTCTACTGGACCATAACTGACACTGAGAGACGACAGCTAGGGGAGCAAACAGG
## 389                                                          ACGAAGGGTGCAAGCGTTGTTCGGAATCATTGGGCGTAAAGCGCGCGCAGGCGGATCAGCAAGTCAGATGTGAAATCTCGAAGCTCAACTTCGAAACTGCGTCTGAAACTGCTAGTCTAGAATGTCGGAGGGGGTAGGGGAATTTCACGTGTAGGGGTAAAATCCGTAGAGATGTGAAGGAACACCGGAGGCGAAGGCGCCTGCCTGGACGACTATTGACGCTGAGGCGCGAAAGCGTGGGGAGCAAACAGG
## 390                                                          ACGGAGGATCCGAGCGTTATCCGGATTTATTGGGTTTAAAGGGTGCGTAGGCGGAATAATAAGTCAGTTGTGAAAGTTTGCGGCTCAACCGTAAAATTGCAGTTGATACTGTTATTCTTGAGTGTACATAAGGTAGGCGGAATTCGTGGTGTAGCGGTGAAATGCTTAGATATCACGAAGAACTCCAATTGCGAAGGCAGCTTACCGGGGTACAACTGACGCTGAGGCACGAAAGTGTGGGTATCAAACAGG
## 392                                                          ACATAGGGTGCAAGCGTTGTCCGGATTTATTGGGCGTAAAGAGTTCGTAGGCGGTTTTTCGCGTCGGATGTGAAAACTCAGGGCTCAACCCTGAGCCTGCATTCGATACGGGAAGACTCGAGGACGGCAGGGGAGACTGGAACTTCTGGTGTAGCGGTGGAATGCGCAGATATCAGAAAGAACACCAATGGCGAAGGCAGGTCTCTGGGCCGATCCTGACGCTGAGGAACGAAAGCGTGGGGAGCAAACAGG
## 393                                                          ACGGAGGGTGCAAGCGTTGTCCGGAATCATTGGGCGTAAAGAGTTCGTAGGTGGCATGTTAAGTCTGGTGTTAAAGCCCGAAGCTCAACTTCGGTTCGGCACTGGATACTGGCAAGCTTGAATGCGGTAGAGGTAAAGGGAATTCCTGGTGTAGCGGTGAAATGCGTAGATATCAGGAGGAACATCGGTGGCGTAAGCGCTTTACTGGGCCGTAATTGACACTGAGGAACGAAAGCCGGGGTAGCAAATGGG
## 394                                                          ACGAAGGATGCAAGCGTTATCCGGATTCATTGGGTTTAAAGGGAGCGTAGGCGGTCTGATAAGTCAGTGGTGAAATCTCTCAGCTTAACTGAGAAACTGCCATTGATACTGTTAGACTTGAGTACGGTTGCTGTGGGCGGAATATGACATGTAGTGGTGAAATACATAGAGATGTCATAGAACACCGATTGCGAAGGCAGCTCACAAAGCCGTAACTGACGCTGAGGCTCGAAAGTGCGGGGATCAAACAGG
## 395                                                          ACAGAGACCTCAAGCGTTATCCGGATTCATTGGGCGTAAAGCGTCCGCAGGTGGTTTTTCAAGTTGGGAGTCAAATCTTTGGGCTTAACCTAGAGACTGCTTTCAATACTGGGAAACTTGAGACTGGGAGAGGTATACGGAACTGTTGGTGTAGTAGTAAAATGCGTTGATATCAACAGGAACACCAAAGGCGAAAGCAGTATACTGGAACAGTTCTGACACTCATGGACGAAAGCGTGGGGAGCGAATGGG
## 396                                                          ACGGAGGGTGCAAGCGTTATCCGGATTTATTGGGTTTAAAGGGTCCGTAGGCGGGCCGATAAGTCAGTGGTGAAAGCCCATAGCTCAACTATGGAACTGCCATTGATACTGTCGGTCTTGAGTATATGTGATGTTGGCGGAATGTGTAGTGTAGCGGTGAAATGCTTAGATATTACACAGAACACCGATTGCGAAGGCAGCTGACAAATATATAACTGACGCTGAGGGACGAAAGTGTGGGGATCAAACAGG
## 397                                                          ACAGAGGGTGCGAGCGTTAATCGGATTTACTGGGCGTAAAGCGTGCGTAGGCGGCTGATTAAGTCGGATGTGAAATCCCTGAGCTTAACTTAGGAATTGCATTCGATACTGGTCAGCTAGAGTATGGGAGAGGATGGTAGAATTCCAGGTGTAGCGGTGAAATGCGTAGAGATCTGGAGGAATACCGATGGCGAAGGCAGCCATCTGGCCTAATACTGACGCTGAGGTACGAAAGCATGGGGAGCAAACAGG
## 398                                                          ACGGAGGGTGCAAGCGTTATCCGGATTTACTGGGTTTAAAGGGTGCGTAGGCGGGCAGGTAAGTCAGTGGTGAAATCCCCGAGCTTAACTCGGGAACTGCCGTTGATACTATCTGTCTTGAATATAGTGGAGGTAAGCGGAATATGTCATGTAGCGGTGAAATGCTTAGATATGACATAGAACACCAATTGCGAAGGCAGCTTGCTACACTATTATTGACGCTGAGGCACGAAAGCGTGGGGATCAAACAGG
## 399                                                          ACGGAGGATGCAAGTGTTATCCGGAATCACTGGGCGTAAAGCGTCTGTAGGTGGTTTAATAAGTCAACTGTTAAATCTTGAGGCTCAACTTCAAAATCGCAGTCGAAACTATTAGACTAGAGTATAGTAGAGGTAAAGGGAATTTCCAGTGGAGCGGTGAAATGCGTAGAGATTGGAAAGAACACCGATGGCGAAAGCACTTTACTGGGCTATTACTAACACTCAGAGACGAAAGCTCGGGTAGCAAATGGG
## 400                                                          ACGTAGGGTGCAAGCGTTAATCGGAATTACTGGGCGTAAAGCGTGCGCAGGCGGTTTTGTAAGACAGATGTGAAAGCCCCGGGCTCAACCTGGGAAGTGCATTTGTGACTGCAAGACTAGAGTGTGTCAGAGGGGGGTGGAATTCCACGTGTAGCAGTGAAATGCGTAGAGATGTGGAGGAACACCGATGGCGAAGGCAGCCCCCTGGGATAACACTGACGCTCATGCACGAAAGCGTGGGGAGCAAACAGG
## 401                                                           ACGGAGGGTGCAAGCGTTATTCGGATTTATTGGGTTTAAAGGGTGTGTAGGCGGTCTTGTAAGTCAGTGGTGAAATCGGGAGGCTCAACTTCGCGATTGCCATTGATACTGTTAGGCTAGAGTATAGCCGGCTTGAGAGGAATGCGACAAGTAGCGGTGAAATGCTTAGATATGTCGCAGAACACCGATAGCGAAGGCATCTCAAGAGGTTATACTGACGCTGAGGCACGAAAGCGTGGGGATCAAACAGG
## 402                                                          ACGAAGGGTGCAAGCGTTGTTCGGAATCATTGGGCGTAAAGCGCGCGCAGGCGGATCAGCAAGTCAGATGTGAAATCTCGAAGCTCAACTTCGAAACTGCGTCTGAAACTGCTAGTCTAGAATGTCGGAGGGGGCAGGGGAATTTCACGTGTAGGGGTAAAATCCGTAGAGATGTGAAGGAACACCGGGGGCGAAGGCGCCTGCCTGGACGACTATTGACGCTGAGGCGCGAAAGCGTGGGGAGCAAACAGG
## 403                                                          ACGTAGGGTGCAAGCGTTAATCGGAATTACTGGGCGTAAAGCGTGCGCAGGCGGTTATGTAAGACAGAGGTGAAATCCCCGGGCTCAACCTGGGAACTGCCTTTGTGACTGCATAGCTGGAGTGCGGCAGAGGGGGATGGAATTCCGCGTGTAGCAGTGAAATGCGTAGATATGCGGAGGAACACCGATGGCGAAGGCAATCCCCTGGACCTGTACTGACGCTCATGCACGAAAGCGTGGGGAGCAAACAGG
## 404                                                          ACGAAGGGTGCAAGCGTTACTCGGAATTACTGGGCGTAAAGCGTGCGTAGGTGGTTCGTTAAGTCTGATGTGAAAGCCCTGGGCTCAACCTGGGAATGGCATTGGATACTGGCGGGCTAGAGTGCGGTAGAGGGTAGTGGAATTCCCGGTGTAGCAGTGAAATGCGTAGAGATCGGGAGGAACATCCGTGGCGAAGGCGACTACCTGGACCAGCACTGACACTGAGGCACGAAAGCGTGGGGAGCAAACAGG
## 405                                                          ACAGAGGATGCAAGCGTTATCCGGAATGATTGGGCGTAAAGCGTCTGTAGGTGGCTCTCCAAGTCTCCTGTCAAATCCCAGAGCTTAACTTTGGATCGGCACGAGAAACTCGAGAGCTTGAGTACGGTAGGGGCAGAGGGAATTCCCGGTGTAGCGGTGAAATGCGTAGAGATCGGGAAGAACACCGATGGCGAAAGCACTCTGCTGGGCCGTTACTGACACTGAGAGACGAAAGCTAAGGGAGCGAATGGG
## 407                                                          ACGAGTGCCCCAAGCGTTATCCGGAATTATTGGGTGTAAAGGGTGCGTAGGTGGTTTTATTAGTCTTGTGTTAAAGCCTGGAGCTTAACTCCAGAAATGCATAGGAAACGGTAAAACTAGAGGATGTGAGAGGTCTGTGGAACTCATAGTGTAGGGGTGAAATCCGTTGATATTATGGGGAACACCAAAAGCGAAGGCAGCAGACTGGCACATTCCTGACATTGAGGCACGAAAGCGTAGGTAGCGAATGGG
## 408                                                          ACAGAGGGGGCAAGCGTTGTCCGGAGTTACTGGGCGTAAAGCGCACGTAGGCGGTTCTGCAAGTGGTGTGTGAAACCAATCAGCTTAACTGATTGACGCCATGCCAAACTGCAGGACTCGAGTCATGGAGAGGTAGTTCGAATTGCTGGTGTAGTGGTGAAATGCGTAGATATCAGCAGGAAGACCAAGGGAGAAATCAGACTACTGGCCATGAACTGACGCTCAGGTGCGAAAGCGTGGGGAGCGAACTGG
## 409                                                          ACGGAGGGTGCAAGCGTTATCCGGATTCACTGGGTTTAAAGGGTGCGTAGGTGGGCAGTTAAGTCAGTGGTGAAATCTCCGGGCTTAACCCGGAAACTGCCATTGATACTATCTGTCTTGAATATCCTGGAGGTGAGCGGAATATGTCATGTAGCGGTGAAATGCTTAGATATGACATAGAACACCGATTGCGAAGGCAGCTCGCTACGGGATTATTGACACTGAGGCACGAAAGCGTGGGGATCAAACAGG
## 410                                                          ACGGAGGGTGCAAGCGTTATCCGGAATCATTGGGTTTAAAGGGTCCGCAGGCGGATTTATAAGTCAGTGGTGAAAGCCTATCGCTTAACGATAGAACTGCCATTGATACTGTAAGTCTTGAATTCGGTCGAAGTGGGCGGAATGTGTAGTGTAGCGGTGAAATGCATAGATATTACACAGAACACCGATAGCGAAGGCAGCTCACTAGGCCTGGATTGACGCTCAGGGACGAAAGCGTGGGGATCAAACAGG
## 411                                                          ACGGAGGGTGCAAGCGTTAATCGGAATTACTGGGCGTAAAGCGCACGCAGGCGGTCTGTCAAGTCGGATGTGAAATCCCCGGGCTCAACCTGGGAACTGCATCCGAAACTGGCAGGCTAGAGTCTTGTAGAGGGGGGTAGAATTCCAGGTGTAGCGGTGAAATGCGTAGAGATCTGGAGGAATACCGGTGGCGAAGGCGGCCCCCTGGACAAAGACTGACGCTCAGGTGCGAAAGCGTGGGGAGCAAACAGG
## 412                                                          ACATAGGGTGCAAGCGTTGTCCGGAATTATTGGGCGTAAAGAGCTCGTAGGTGGTCGATCACGTCGGATGTGAAATTCTGAGGCTTAACCTCGGACCTGCATTCGATACGGGTTGACTAGAGTGTGGTAGGGGAGACTGGAATTCCTGGTGTAGCGGTGGAATGCGCAGATATCAGGAGGAACACCGATGGCGAAGGCAGGTCTCTGGGCCATTACTGACACTGAGGAGCGAAAGCGTGGGGAGCGAACAGG
## 413                                                          ACGGAGGGTGCAAGCGTTAATCGGAATTACTGGGCGTAAAGCGCACGCAGGCGGTCAGATAAGTCAGATGTGAAAGCCCCGGGCTCAACCTGGGAACTGCATTTGAAACTGTCTGACTAGAGTCTTGTAGAGGGGGGTAGAATTCCAGGTGTAGCGGTGAAATGCGTAGAGATCTGGAGGAATACCGGTGGCGAAGGCGGCCCCCTGGACAAAGACTGACGCTCAGGTGCGAAAGCGTGGGGAGCAAACAGG
## 414                                                          ACAGAGACCCCGAGCGTTGACCGGATTTATTGGGCGTAAAGCGTACGCAGGCTGTCTTGTAAGTCATTTGTTAAACCTTCAGGCTCAACCTGGAGACTGCGGGTGATACTGCAAGACTAGAGTATGGGAGAGGTCGGCGGAATGCCTAGTGTAGGGGTAAAATCCGTTAATATTAGGTAGAACACCAAATGTGAAAACAGCCGACTGGAACATTACTGACGCTGATGTACGAAAGCATGGGTAGCGAATGGG
## 415                                                          ACAGAGGGTGCAAGCGTTAATCGGAATTACTGGGCGTAAAGCGCGCGTAGGCGGTTTGTTAAGTCGGATGTGAAATCCCTGGGCTCAACCTAGGCACTGCATCCGATACTGGCTGACTAGAGTGTGGGAGAGGAAGGTAGAATTCCAGGTGTAGCGGTGAAATGCGTAGAGATCTGGAGGAATACCGATGGCGAAGGCAGCCTTCTGGCCTAACACTGACGCTGAGGTGCGAAAGCATGGGGAGCAAACAGG
## 416                                                          ACGTAGGGTGCAAGCGTTGTCCGGAATTATTGGGCGTAAAGAGCTCGTAGGCGGTTTGTCGCGTCTGCTGTGAAATCCCGAGGCTCAACCTCGGGCCTGCAGTGGGTACGGGCAAACTAGAGTGTGGTAGGGGAGATTGGAACTCCTGGTGTAGCGGTGGAATGCGCAGATATCAGGAAGAACACCGATGGCGAAGGCAGATCTCTGGGCCATTACTGACGCTGAGGAGCGAAAGCGTGGGGAGCGAACAGG
## 417                                                           ACGGGAGGGGCAAGCGTTATCCGAAATTACTGGGCGTAAAGTGTCCGTAGATTGTAATATAAGTTATATGTTAAAATTTAAAGCTTAACTTTAAAAAAATATACAATACTGTTTTACTTGAGTTTTATACGGAAGAGTAGAATTTTACGTTAAGGAGTAAATTCCAAGAATACGTAGAGGAATATCATTAGCGAAGGCGACTCTTTAGTACAAACTGACATTGAGGGACGAAAGTGTAGGGAGCAAACAGG
## 418                                                          ACGTAGGGTGCAAGCGTTAATCGGAATTACTGGGCGTAAAGCGTGCGCAGGCGGTTATGCAAGACAGAGGTGAAATCCCCGGGCTCAACCTGGGAACTGCCTTTGTGACTGCATAGCTAGAGTACGGCAGAGGGGGATGGAATTCCGCGTGTAGCAGTGAAATGCGTAGATATGCGGAGGAACACCGATGGCGAAGGCAATCCCCTGGACCTGTACTGACGCTCATGCACGAAAGCGTGGGGAGCAAACAGG
## 419                                                          ACGTAGGGTGCAAGCGTTGTCCGGAATTATTGGGCGTAAAGAGCTCGTAGGCGGTTTGTCACGTCTGCTGTGAAATTTCGAGGCTCAACCTCGAACTTGCAGTGGGTACGGGCAGGCTAGAGTGCGGTAGGGGAGATGGGAATTCCTGGTGTAGCGGTGGAATGCGCAGATATCAGGAGGAACACCAATGGCGAAGGCACATCTCTGGGCCGTAACTGACGCTGAGGAGCGAAAGCGTGGGGAGCGAACAGG
## 420                                                          ACATAGGGTGCAAGCGTTGTCCGGAATTATTGGGCGTAAAGAGCTCGTAGGTGGTTTGATACGTCGGGTGTGAAATTCAGGGGCTCAACCCCTGACCTGCATTCGATACGGTCAAGCTAGAGTTTGGTAGGGGAGACTGGAATTCCTGGTGTAGCGGTGGAATGCGCAGATATCAGGAGGAACACCGATGGCGAAGGCAGGTCTCTGGGCCAATACTGACACTGAGGAGCGAAAGCGTGGGGAGCGAACAGG
## 421                                                          ACGAGTGCCCCAAGCGTTATCCGGAATCATTGGGCGTAAAGGGTGTGTAGGTGGTTTTGTTAGTCTTATGTTAAAACCCGAGGCTTAACTTCGGACATGCATTTGATACGGCAAAACTAGAGGATGACAGAGGTGTGCGGAACTCATGGAGTAGGGGTGAAATCCGTTGATATCATGGGGAACACCAAAAGCGAAGGCAGCACACTGGGTCATTCCTGACACTGAAACACGAAAGCGTGGGGAGCAAAAAGG
## 422                                                          ACGTAGGGTGCGAGCGTTAATCGGAATTACTGGGCGTAAAGCGTGCGCAGGCGGTTTCGTAAGACAGACGTGAAATCCCCGGGCTCAACCTGGGAACTGCCTTTGTGACTGCATAGCTAGAGTACGGCAGAGGGGGATGGAATTCCGCGTGTAGCAGTGAAATGCGTAGATATGCGGAGGAACACCGATGGCGAAGGCAATCCCCTGGGCCTGTACTGACGCTCATGCACGAAAGCGTGGGGAGCAAACAGG
## 423                                                          ACGTAGGGTGCAAGCGTTAATCGGAATTACTGGGCGTAAAGCGTGCGCAGGTGGTTGTATAAGACAGATGTGAAATCCCCGGGCTCAACCTGGGAACTGCATTTGTGACTGTACGGCTAGAGTGTGTCAGAGGGAGGTGGAATTCCACGTGTAGCAGTGAAATGCGTAGATATGTGGAGGAACACCAATGGCGAAGGCAGCCTCCTGGGATAACACTGACGCTCATGCACGAAAGCGTGGGGAGCAAACAGG
## 424                                                          ACGTAGGGTGCAAGCGTTAATCGGAATTACTGGGCGTAAAGCGTGCGCAGGCGGACTTTTAAGCCAGATGTGAAAGCCCCGAGCTTAACTTGGGAATTGCGTTTGGAACTGGGAGTCTAGAGTCTGTCAGAGGGGGATGGAATTCCACGTGTAGCAGTGAAATGCGTAGAGATGTGGAGGAACACCGATGGCGAAGGCAATCCCCTGGACCTGTACTGACGCTCATGCACGAAAGCGTGGGGAGCAAACAGG
## 425                                                          ACATAGGGTGCAAGCGTTGTCCGGAATTATTGGGCGTAAAGAGCTCGTAGGTGGTTTGTTACGTCGGATGTGAAATGCAGGGGCTCAACCCCTGACTTGCATCCGATACGGGCAAGCTAGAGTTTGGTAGGGGAGACTGGAATTCCTGGTGTAGCGGTGGAATGCGCAGATATCAGGAGGAACACCGATGGCGAAGGCAGGTCTCTGGGCCAATACTGACACTGAGGAGCGAAAGCGTGGGGAGCGAACAGG
## 426                                                          ACGTGAGGTGCTAACATTACCCGGATTTATTGGGCGTAAAGAGTTCGTAGGCGGCAAGGTAAGTCAGATATTAAATCCTGAGGCTCAACTTCAGGTCTGTATCTGAAACTGTCTAGCTAGAGTCTTGGAGAGGTGAGTGGAATTCCTCATGTAGGGGTAAAATCCGTAGATATGAGGAGGAACACCAAAAGCGAAGGCAGCTCACTGGCCAAGTACTGACGCTAAAGAACGAAAGCATGGGTAGCGAACGGG
## 427                                                          ACGTAGGGTGCAAGCGTTAATCGGAATTACTGGGCGTAAAGCGCGCGTAGGCGGTTAATTAAGCGAGATGTGAAATCCCTGGGCTCAACCTAGGAATTGCGTTTCGAACTGATTAGCTAGAGTGCAGTAGAGGGTGGTGGAATTTCCGGTGTAGCGGTGAAATGCGTAGAGATCGGAAGGAACATCAGTGGCGAAGGCGACTGCCTGGACTGACACTGACGCTGAGGTGCGAAAGCGTGGGGAGCAAACAGG
## 428                                                          ACGTAGGGTGCGAGCGTTAATCGGAATTACTGGGCGTAAAGCGTGCGCAGGCGGTTGTGTAAGACAGATGTGAAATCCCCGGGCTCAACCTGGGAACTGCGTTTGTGACTGCACAACTAGAGTACGGCAGAGGGAGGTGGAATTCCGCGTGTAGCAGTGAAATGCGTAGAGATGCGGAGGAACACCGATGGCGAAGGCAGCCTCCTGGGCCAGTACTGACGCTCATGCACGAAAGCGTGGGGAGCAAACAGG
## 429                                                          ACGTAGGGTGCAAGCGTTAATCGGAATTACTGGGCGTAAAGCGTGCGCAGGCGGTTCGGAAAGAAAGGTGTGAAATCCCAGGGCTTAACCTTGGAACTGCACTTTTAACTACCGAGCTAGAGTATGTCAGAGGGGGGTGGAATTCCGCGTGTAGCAGTGAAATGCGTAGATATGCGGAGGAACACCGATGGCGAAGGCAATCCCCTGGACCTGTACTGACGCTCATGCACGAAAGCGTGGGGAGCAAACAGG
## 430                                                          ACGAAGGATGCAAGCGTTATCCGGATTCATTGGGTTTAAAGGGAGCGTAGGCGGTCTTATAAGTCAGTGGTGAAAGCTCTTCGCTTAACGAAGAAATTGCCATTGATACTGTAGGACTTGAGTACAGTTGCTGTGGGCGGAATATGACATGTAGTGGTGAAATACATAGAGATGTCATAGAACACCGATTGCGAAGGCAGCTCACAAAACTGTAACTGACGCTGAGGCTCGAAAGTGCGGGGATCAAACAGG
## 431                                                          ACGGAGGGTGCAAGCATTAATCGGAATTATTGGGCGTAAAGGGCGCGTAGGCGGATAGGATAGTCAGATGTGAAATTTCGGAGCTCAACTTCGAAGCTGCATTTGAAACTACTTATCTGGAGGGTAGGCGGAGAAAACGGAATTCCACATGTAGCGGTGAAATGCGTAGAGATGTGGAAGAACACCTGTGGCGAAAGCGGTTTTCTAGCTTATTCCTGACGCTGAGGCGCGAAAGCAAGGGGAGCAAACAGG
## 432                                                          ACGGAGGATCCAAGCGTTATCCGGATTTATTGGGTTTAAAGGGTGCGTAGGCGGGCCCTTAAGTCAGTGGTGAAAGTTTGCAGCTTAACTGTAAAATTGCCATTGAAACTGAGGGTCTTGAGTGTAAATAAGGTAGGCGGAATGTGTTGTGTAGCGGTGAAATGCTTAGATATAACACAGAACACCAATTGCGAAGGCAGCTTACTGGGATACAACTGACGCTGAGGCACGAAAGCGTGGGGATCAAACAGG
## 433                                                          ACGGAGGGTGCAAGCGTTAATCGGAATAACTGGGCGTAAAGCGCACGCAGGCGGTCGGATAAGTCAGATGTGAAAGCCCCGGGCTCAACCTGGGAACTGCATTTGAAACTGTTCGACTAGAGTCTTGTAGAGGGGGGTAGAATTCCAGGTGTAGCGGTGAAATGCGTAGAGATCTGGAGGAATACCGGTGGCGAAGGCGGCCCCCTGGACAAAGACTGACGCTCAGGTGCGAAAGCGTGGGGAGCAAACAGG
## 434                                                          ACGAAGGGTGTAAGCGTTGTTCGGATTTATTGGGCGTAAAGCGCGCGCAGGTGGATTTTTAAGTCAGATGTGAAATCTCGGGGCTCAACCCCGAACGTGCGTCTGAAACTGGAGATCTAGAATATTGGAGGGGGTAGAGGAATTTCACATGTAGGGGTAAAATCCGTAGAGATGTGAAGGAACACCAGAGGCGAAGGCGTCTACCTGGCCAAACATTGACACTGAGGCGCGAAAGCGTGGGGAGCAAACAGG
## 435                                                          ACGTAGGGTGCGAGCGTTAATCGGAATTACTGGGCGTAAAGCGTGCGCAGGCGGCTTTGCAAGACAGAGGTGAAATCCCCGGGCTCAACCTGGGAACTGCCTTTGTGACTGCAAGGCTAGAGTACGGCAGAGGGGGATGGAATTCCGCGTGTAGCAGTGAAATGCGTAGATATGCGGAGGAACACCGATGGCGAAGGCAATCCCCTGGGCCTGTACTGACGCTCATGCACGAAAGCGTGGGGAGCAAACAGG
## 436                                                          ACGGAGGATCCAAGCGTTATCCGGATTTATTGGGTTTAAAGGGTCCGTAGGCGGGTCTTTAAGTCAGTGGTGAAAGCCTGCAGCTTAACTGTGGAAATGCCATTGATACTGGAGGCCTTGAGTGTAGTAGAAGTAGGCGGAATAGGGCATGTAGCGGTGAAATGCATAGATATGCCCTAGAACACCGATTGCGAAGGCAGCTTACTATGTTACAACTGACGCTGAGGGACGAAAGCGTGGGGAGCAAACAGG
## 437                                                          ACGAAGGGTGCAAGCGTTATTCGGAATGACTGGGCGTAAAGAGCGTGAAGGCGGCTTCTTAAGTCAACTGTTAAAGGGCTCGGCTTAACTGAGCAACTGCGGGTGAAACTGAGAAGCTAGAGTGCAGAAGAGAAAGGTAGAATTCTCGGAGTAGCGGTAAAATGCGTAGATCTCGAGAGGAATACCGGTTGCGAAGGCGGCCTTTTGGTCTGTAACTGACGCTCAAGCGCGAAAGCGTGGGGAGCAAACAGG
## 439                                                          ACGGAGGGTGCAAGCGTTACTCGGAATCACTGGGCGTAAAGGATGCGTAGGCTGGAAATCAAGTCGAGAGTGAAATCCAACGGCTCAACCGTTGAACTGCTCTCGAAACTGATTACCTAGAATATGGGAGAGGTAGATGGAATTGGTGGTGTAGGGGTAAAATCCGTAGATATCACCAGGAATACCGATTGCGAAGGCGATCTACTGGAACATTATTGACGCTGAGGCATGAAAGCGTGGGGAGCAAACAGG
## 440                                                          ACGGAGGATGCAAGTGTTATCCGGAATCACTGGGCGTAAAGCGTCTGTAGGTGGTCAAATAAGTCAACTGTTAAATCTTGAGGCTCAACCTCAAAATCGCAGTCGAAACTATTAGACTAGAGTATAGTAGGGGTAAAGGGAATTTCCAGTGGAGCGGTGAAATGCGTAGAGATTGGAAAGAACACCGATGGCGAAGGCACTTTACTGGGCTATTACTAACACTGAGAGACGAAAGCTAGGGTAGCAAATGGG
## 441                                                          ACGTAGGGTGCAAGCGTTAATCGGAATTACTGGGCGTAAAGCGTGCGCAGGCGGTTGTGCAAGACAGATGTGAAATCCCCGGGCTTAACCTGGGAATGGCATTTGTGACTGCACGGCTAGAGTGTGTCAGAGGGGGGTAGAATTCCACGTGTAGCAGTGAAATGCGTAGATATGTGGAGGAATACCGATGGCGAAGGCAGCCCCCTGGGATAACACTGACGCTCATGCACGAAAGCGTGGGGAGCAAACAGG
## 442                                                          ACAGAGACTGCAAGCGTTATTCGGATTCACTGGGCGTAAAGGGTGCGCAGGCGGTTGCGTGTGTTGGATGTGAAAGCCCGGGGCTCAACCCCGGAATTGCGTCCAAAACTACGCGACTAGAGTACCGGAGAGGATAGCGGAATTCACGGTGTAGCAGTGAAATGCGTAGATATCGTGAAGAACACCAGCGGCGAAGGCGGCTATCTGGAAGGTAACTGACGCTCAGGCACGAAAGCGTGGGGAGCAAAAGGG
## 443                                                          ACAGAGGGTGCAAACGTTGCTCGGAATTACTGGGCGTAAAGCGCGTGTAGGTGGACCAGCAAGTCAGACGTGAAAGCCCTGGGCTCAACCCAGGAACTGCATTTGAAACTGCTGGTCTGGAGTGGCGGAGAGGATCGTGGAATTCCTGGTGTAGAGGTGAAATTCGTAGATATCAGGAGGAACACCGGTGGCGAAGGCGGCGATCTGGACGCTTACTGACACTGAGACGCGAAAGCGTGGGGAGCAAACAGG
## 444                                                          ACAGAGGTGGCAAGCGTTGTTCGGATTTATTGGGTGTAAAGGGCAGGTAGGCGGCTAAGTAAGTCTGTTGTGAAATCCCGGGGCTCAACCCCGGAACTGCAACGGAAACTACTTAGCTGGAACATAGTAGGGGTGAGGGGAATTCTCGGTGTAAGGGTGAAATCTGTAGATATCGAGAGGAACACCAATGGCGAAGGCACCTCACTGGACTATTGTTGACGCTGAGCTGCGAAAGTAGGGGGAGCAAACAGG
## 445                                                         ACGAGGGCCCCGAGCGTTATCCGGAATTATTGGGCGTAAAGGGTGTGTAGGTGGTCATATTAGTCTTGTGTAAAAGCCTGTCGCCTTACGACAGATCCGCACGGGAAACGGTATGACTGAGAGGGTGCAAGAGGTATATGGAACTCACGGTGTAGGGGTGAAATCCGTTGATATCGTGGGGAACGCCAAAAGCGAAAGCAATATACTGGTGCATACCTGACACTGAAACACGAAAGCCAGGGTAGCGAATGGG
## 447                                                          ACGTAGGTGGCGAGCGTTGTCCGGATTTATTGGGTTTAAAGGGTGCGTAGGCGGCTTATTAAGTCAGTGGTGAAAGACGGCAGCTTAACTGTCGAGGTGCCATTGATACTGATGAGCTAGAGTACTGATGAGGTAGGCGGAATGGACGGTGTAGCGGTGAAATGCATAGATATCGTCCAGAACACCGATAGCGAAGGCAGCTTACTAAGGAGTAACTGACGCTGAGGCACGAAAGTGTGGGGATCAAACAGG
## 448                                                          ACGAGTGCCCCGAGCGTTATCCGGATTTATTGGGCGTAAAGCGTCCGCAGGCGGTCTGTAACATCTTTTGTTAAATCACAAGGCTTAACTTTGTGACCGCGAGAGATATGGACAGACTAGAGACTGGGAGAGGTAAGCGGAATTGCTGGTGTAGGGGTTAAATCCGTTAATATCAGCAGGAACACCAAATGCGAAGGCAGCTTACTGGAACAGTTCTGACGCTCATGGACGAAAGCGTGGGTAGCGAATGGG
## 449                                                          ACGGAGGGGGCTAGCGTTGTTCGGAATTACTGGGCGTAAAGCGCACGTAGGCGGACTGGAAAGTCAGGGGTGAAATCCCGGGGCTCAACCCCGGAACTGCCTTTGAAACTCCCAGTCTTGAGGTCGAGAGAGGTGAGTGGAATTCCGAGTGTAGAGGTGAAATTCGTAGATATTCGGAGGAACACCAGTGGCGAAGGCGGCTCACTGGCTCGATACTGACGCTGAGGTGCGAAAGCGTGGGGAGCAAACAGG
## 450                                                          ACAGAGGGTGCGAGCGTTGTCCGGAATCACTGGGCGTAAAGGGCGCGTAGGTGGCTTGGTAAGCGTGTGGTGAAAGTCCGGGGCTCAACCCCGGATCTGCCGTGCGAACTGCTGAGCTCGAGCGATGTAGAGGCAGGTGGAATTTCGGGTGTAGCGGTGGAATGCGTAGATATCCGAAAGAACACCAGTGGCGAAGGCGGCCTGCTGGGCATCAGCTGACACTGAGGCGCGACAGCGTGGGGAGCAAACAGG
## 451                                                          ACAGAGACCTCAAACATTGTCCGGATTCATTGGGCGTAAAGCGTCCGCAGGTGGCTTAATAAGTTAGGGGTTAAAGCTTCCGACCTAATCGGAAAACTGCTTCTAATACTATTAAGCTCGAGGTTGGGAGAGGCATATGGAACTGCCGGTGTAGTAGTAAAATGCGTTGATATCGGCAGGAACACCAAAGGCGAAGGCAATATGCTGGAACAACTCTGACACTCAGGGACGAAAGCGTGGGGAGCGAATGGG
## 452                                                          ACGTAGGGTGCGAGCGTTAATCGGAATTACTGGGCGTAAAGCGTGCGCAGGCGGTTTGTTAAGACAGTTGTGAAATCCCCGGGCTTAACCTGGGAACTGCAATTGTGACTGGCAGGCTAGAGTTTGGCAGAGGGGGGTGGAATTCCTGGTGTAGCAGTGAAATGCGTAGATATCAGGAGGAACACCGATGGCGAAGGCAGCCCCCTGGGCCATGACTGACGCTCATGCACGAAAGCGTGGGGAGCAAACAGG
## 453                                                           ACGTAGGTGGCGAGCGTTGTCCGGAATTATTGGGCGTAAAGGGAGCGCAGGTGGGCTGTCAAGTCTGTCTTAAAAGTGCGGGGCTTAACCCCGTGATGGGATAGAAACTGGCAGTCTTGAGTGCAGGAGAGGAAAGCGGAATTCCTAGTGTAGCGGTGAAATGCGTAGATATTAGGAGGAACACCAGTGGCGAAGGCGGCTTTCTGGACTGTAACTGACACTGAGGCTCGAAAGCCAGGGGAGCGAACGGG
## 454                                                          ACAGAGGTCCCAAGCGTTGTTCGGATTCACTGGGCGTAAAGGGTGCGTAGGCGGTGGGGTAAGTTTGAGGTGAAATCTCCGGGCTCAACCCGGAAAATGCCTTGAAGACTATCCTGCTAGAGGATCGGAGGGGAGACTGGAATTCTCGGTGTAGCAGTGAAATGCGTAGATATCGAGAGGAACACCAGTGGCGAAGGCGAGTCTCTGGACGATTCCTGACGCTGAGGCACGAAAGCCAGGGGAGCAAACGGG
## 455                                                          ACGGAGGGTGCAAGCGTTGTCCGGATTTATTGGGTTTAAAGGGTGCGTAGGCGGCTGAATAAGTCAGCGGTGAAAGACTCCGGCTTAACCGGAGCAGTGCCGTTGATACTGTTTAGCTTGAGTTCTGCAGGGGTACATGGAATTGATGATGTAGCGGTGAAATGCATAGATATCATCAGGAACACCGATAGCGAAGGCATTGTACTGGGCAGCGACTGACGCTGAGGCACGAAAGTGTGGGGATCGAACAGG
## 456                                                          ACATAGGGTGCAAGCGTTGTCCGGAATTATTGGGCGTAAAGAGCTCGTAGGTGGTTCGTTACGTCGGATGTGAAAACCTGAGGCTCAACCTCAGGCCTGCATTCGATACGGGCGAACTGGAGTGTGGTAGGGGAGACTGGAATTCCTGGTGTAGCGGTGGAATGCGCAGATATCAGGAGGAACACCAATGGCGAAGGCAGGTCTCTGGGCCACTACTGACACTGAGGAGCGAAAGCGTGGGGAGCAAACAGG
## 457                                                          ACGTAGGGGGCGAGAGTTGTCCGGATTTACTGGGTGTAAAGGGTGCTCAGGTGGATTTGTAAGTCAGAGGTGAAATCCCAAAGCTCAACTTTGGTGCTGCCTTTGATACTGCAAGTCTTGAGTTTGGAAGAGGGTAATGGAATATCTGGTGTAGCAGTGAAATGCGTAGATATCAGATAGAACACCAATGGCGAAGGCAGTTACCTGGTCCAAAACTGACACTAAAGCACGAAAGTGTGGGGAGCAAACAGG
## 458                                                          ACAGAGGGTGCAAGCGTTAATCGGAATTACTGGGCGTAAAGCGCACGTAGGCGGCTACGTAAGTCAGGTGTGAAATCCCCGGGCTCAACCTGGGAACTGCACTTGAGACTGCGTGGCTAGAGTATGGGAGAGGAAGGCGGAATTCCCGGTGTAGCGGTGAAATGCGTAGATATCGGGAGGAACATCAGTGGCGAAGGCGGCCTTCTGGACCAATACTGACGCTCAGGTGCGAAAGCGTGGGGAGCAAACAGG
## 459                                                          ACGAAGGTGGCAAGCGTTGTTCGGATTTACTGGGCGTAAAGAGTGCGTAGGCGGTTTAATAAGTCAGAAGTGAAAGCCCAGGGCTCAACCTTGGAATTGCTTTTGAAACTGTTTTACTTGAATCCGGTAGAGGTTGGCGGAATTCTCAGTGTAGAGGTGAAATTCGCAGATATTGAGAGGAACACCAGTGGCGTAAGCGGCCAACTGGACCGGTATTGACGCTGAGGCACGAAAGCATGGGGAGCAAACAGG
## 460                                                          ACGTAGGGTGCGAGCGTTAATCGGAATTACTGGGCGTAAAGGGTGCGCAGGCGGTTTTGTAAGTCAGATGTGAAATCCCCGGGCTTAACCTGGGAATTGCGTTTGAAACTACAAATCTAGAGTGTGGCAGAGGGAGGTGGAATTCCATGTGTAGCAGTGAAATGCGTAGAGATATGGAAGAACATCGATGGCGAAGGCAGCCTCCTGGGTTAACACTGACGCTCATGCACGAAAGCGTGGGGAGCAAACAGG
## 461                                                          ACGTAGGGTGCAAGCGTTAATCGGAATTACTGGGCGTAAAGCGTGCGCAGGCGGTTATGTAAGACAGAGGTGAAATCCCCGGGCTCAACCTGGGAACTGCCTTTGTGACTGCATAGCTAGAGTACGGTAGAGGGGGATGGAATTCCGCGTGTAGCAGTGAAATGCGTAGATATGCGGAGGAACACCGATGGCGAAGGCAATCCCCTGGGCCTGTACTGACGCTCATGCACGAAAGCGTGGGGAGCAAACAGG
## 462                                                          ACGGAGGGTGCAAGCGTTATCCGGAATCATTGGGTTTAAAGGGTCCGCAGGCGGACTTATAAGTCAGTGGTGAAAGCCTACAGCTCAACTGTAGAACTGCCATTGATACTGTAAGTCTTGAATATGTTCGGAGTGGGCGGAATGTGACATGTAGCGGTGAAATGCTTAGATATGTCACAGAACACCGATAGCGAAGGCAGCTCACTAGACCATTATTGACGCTCATGGACGAAAGCGTGGGGAGCAAACAGG
## 463                                                          ACGTAGGGTGCAAGCGTTAATCGGAATTACTGGGCGTAAAGCGTGCGCAGGCGGTGATGTAAGACAGAGGTGAAATCCCCGGGCTCAACCTGGGAACTGCCTTTGTGACTGCATCGCTGGAGTACGGCAGAGGGGGATGGAATTCCGCGTGTAGCAGTGAAATGCGTAGATATGCGGAGGAACACCGATGGCGAAGGCAATCCCCTGGGCCTGTACTGACGCTCATGCACGAAAGCGTGGGGAGCAAACAGG
## 464                                                          ACGGAGGGTGCAAGCGTTATCCGGATTCACTGGGTTTAAAGGGTGTGTAGGCGGGCTTGTAAGTCCGAGGTGAAATCTCCGAGCTCAACTTGGAAACTGCCTTGGATACTATAGGTCTTGAATATCGTTGAGGTAGGCGGAATATGTCATGTAGCGGTGAAATGCTTAGATATGACATAGAACACCGATTGCGAAGGCAGCTTGCTAAACGATTATTGACGCTGAGGCACGAAAGCGTGGGGATCAAACAGG
## 466                                                          ACGAAGGATCCAAGCGTTGTCCGGATTTACTGGGTTTAAAGGGTGCGTAGGCGGGAAATTAAGTCAGTGGTGAAAGCTGGTAGCTCAACTATCAAATTGCCATTGAAACTGATTTTCTCGAGTACGGTTGAGGTAGCTGGAATGTAACATGTAGCGGTGAAATGCTTAGATATGTTACAGAACACCAATTGCGAAGGCAGGCTACTAAACCGTAACTGACGCTGAGGCACGAAAGCGTGGGGATCAAACAGG
## 467                                                          ACGGAGGGTGCAAGCGTTATCCGGATTCACTGGGTTTAAAGGGTGCGTAGGCGGGCAGGTAAGTCAGTGGTGAAATCTTCGAGCTTAACTCGGAAACTGCCGTTGATACTATCTGTCTTGAATTTAGTGGAGGTAAGCGGAATATGTCATGTAGCGGTGAAATGCTTAGATATGACATAGAACACCAATTGCGAAGGCAGCTTGCTACACTAATATTGACGCTGAGGCACGAAAGCGTGGGGATCAAACAGG
## 468                                                          ACGGAGGGTGCAAGCGTTACTCGGAATCACTGGGCGTAAAGAGCGTGTAGGCGGATATATAAGTCAGAAGTGAAATCCAATAGCTTAACTATTGAACTGCTTTTGAAACTGTATATCTAGAATGTGGGAGAGGTAGATGGAATTTCTGGTGTAGGGGTAAAATCCGTAGAGATCAGAAGGAATACCGATTGCGAAGGCGATCTACTGGAACATTATTGACGCTGAGACGCGAAAGCGTGGGGAGCAAACAGG
## 469                                                          ACGGAGGGTGCTAGCGTTGTTCGGAATTACTGGGCGTAAAGCGCGCGTAGGCGGTCTAGTAAGTTGGAAGTGAAATCCCAGAGCTTAACTCTGGAATTGCTTTCAAAACTGTTAGACTTGAGAATGGTAGGGGAAAGTGGAATTCCTAGTGTAGAGGTGAAATTCGTAGATATTAGGAGGAACACCAGAGGCGAAAGCGGCTTTCTGGACCATTTCTGACGCTAAGGCGCGAAAGCGTGGGGAGCAAACAGG
## 470                                                          ACGGAGGATCCGAGCGTTATCCGGATTTATTGGGTTTAAAGGGAGCGTAGGCGGACGCTTAAGTCAGTTGTGAAAGTTTGCGGCTCAACCGTAAAATTGCAGTTGATACTGGGTGTCTTGAGTACAGTAGAGGCAGGCGGAATTCGTGGTGTAGCGGTGAAATGCTTAGATATCACGAAGAACTCCGATTGCGAAGGCAGCTTGCTGGACTGTAACTGACGCTGATGCTCGAAAGTGTGGGTATCAAACAGG
## 471                                                          ACAGAGGATGCAAGCGTTATCCGGAATCACTGGGCATAAAGCGTCTGTAGGTTGCTTGCCAAGTCTGCTGTTAAAGATCAGGGCTTAACCCTGGGAAAGCAGTGGAAACTAGTAGGCTTGAGTGTGGTAGAGGTAGAGGGAATTCCTGGTGTAGCGGTGAAATGCGTAGATATTAGGAAGAACACCAATGGCGAAAGCACTCTACTGGGCCACAACTGACACTGAGAGACGACAGCTAGGGGAGCAAATGGG
## 472                                                          ACGAAGGATCCAAGCGTTGTCCGGATTTACTGGGTTTAAAGGGTGCGTAGGCGGACTTTTAAGTCAGTGGTGAAAGCCGGTAGCTTAACTATCGAATTGCCATTGAAACTGAAAGTCTCGAGTATGGTTGAGGTGGCCGGAATGTATCATGTAGCGGTGAAATGCTTAGATATGATACAGAACACCAATTGCGAAGGCAGGTCACTAAGCCATAACTGACGCTGAGGCACGAAAGCGTGGGGAGCAAACAGG
## 473                                                          ACGTAGGGTGCGAGCGTTAATCGGAATTACTGGGCGTAAAGCGTGCGCAGGCGGTTATGCAAGACAGAGGTGAAATCCCCGGGCTCAACCTGGGAACTGCCTTTGTGACTGCATGACTAGAGTACGGCAGAGGGAGATGGAATTCCGCGTGTAGCAGTGAAATGCGTAGATATGCGGAGGAACACCGATGGCGAAGGCAATCTCCTGGGCCTGTACTGACGCTCATGCACGAAAGCGTGGGGAGCAAACAGG
## 474                                                           ACGTATGTCGCAAGCGTTATCCGGAATTATTGGGCATAAAGGGCGTCTAGGCGGTTTAGAAAGTCTGATGTGAAAATGCGGGGCTCAACCCCGTATTGCGTTGGAAACTACTAGACTAGAGTATTGGAGAGGTAAGCGGAACTACAAGTGTAGAGGTGAAATTCGTAGATATTTGTAGGAATGCCAATTGGGAAGCCAGCTTACTGGACAAATACTGACGCTGAAGCGCGAAAGCTAGGGGAGCAAACAGG
## 475                                                          ACGGAGAGTGCAAACGTTGTTCGGAATTATTGGGCGTAAAGGGCGCGTAGGCGGCTCGGTAAGTCAGATGTGAAAGCCCGGGGCTCAACCCTGGAAGTGCGTTTGAAACTGCTGAGCTTGAGTGCAGGAGGGAGAGGTGGAATTCCAGGTGTAGGAGTGAAATCCGTAGATATCTGGAGGAACACCGGTGGCGAAGGCGACCTCTTGGCCTGAAACTGACGCTGAGGCGCGAAGGCGTGGGGAGCAAACAGG
## 476                                                          ACAGAGGATGCAAGCGTTATCCGGAATCACTGGGCATAAAGCGTCTGTAGGTTGCTTAGCAAGTCTGCTGTTAAAGATCAGGGCCTAACCCTGGGAAAGCAGTGGAAACTAGTAGGCTTGAGTGTGGTAGAGGTAGAGGGAATTCCTGGTGTAGCGGTGAAATGCGTAGATATTAGGAAGAACACCAATGGCGAAAGCACTCTACTGGGCCACAACTGACACTAAGAGACGACAGCTAGGGGAGCAAATGGG
## 478                                                          ACGTAGGGTGCAAGCGTTAATCGGAATTACTGGGCGTAAAGCGTGCGCAGGCGGTTGTGCAAGACAGGTGTGAAATCCCCGGGCTTAACCTGGGAACTGCACTTGTGACTGCACAGCTTGAGTGCGGCAGAGGGGGATGGAATTCCGCGTGTAGCAGTGAAATGCGTAGATATGCGGAGGAACACCGATGGCGAAGGCAATCCCCTGGGCCTGCACTGACGCTCATGCACGAAAGCGTGGGGAGCAAACAGG
## 480                                                          ACGTGAGTGGCAAGCGTTATCCGGAATCATTGGGCGTAAAGCGTCAGCAGGGGGATGTTCAAGTGAAATGTTAAAGCCTTCGACCCAATCGAAGATCTGTGTTTCATACTGTTCATCTAGAAGATAGGAGAGGTAAATAGAATTTCCGGAGTAGGGGTAATATCCGTAGATACCGGAAGGAACACCAATAGCGAAGGCAATTTACTGGCCTATTCTTGACCCTCAGTGACGAAAGCGTGGGTAGCAAACAGG
## 481                                                          ACAGAGGGTGCAAGCGTTAATCGGAATTACTGGGCGTAAAGCGCGCGTAGGCGGCTGCCTAAGTCGGATGTGAAATCCCCGGGCTTAACCTGGGAACTGCATTCGATACTGGGCGGCTAGAGTATGAGAGAGGGAGGTAGAATTCCACGTGTAGCGGTGAAATGCGTAGATATGTGGAGGAATACCGGTGGCGAAGGCGGCCTCCTGGCTTAATACTGACGCTGAGGTGCGAAAGCGTGGGGAGCAAACAGG
## 482                                                          ACGAAGGGACCTAGCGTAGTTCGGAATTACTGGGCGTAAAGCGCGCGTAGGCCGTTGAGTTAGTTAATTGTGAAATCCCAAAGCTTAACTTTGGAACTGCAATTAAAACTGCTCGACTAGAGTTTGATAGAGGAAAGCGGAATACATAGTGTAGAGGTGAAATTCGTAGATATTATGTAGAACACCAGTTGCGAAGGCGGCTTTCTGGATCAACACTGACGCTGAGGCGCGAAAGTATGGGTAGCAAAGAGG
## 483                                                          ACGAGTGCCCCAAGCGTTATCCGGAATTATTGGGCGTAAAGGGTGCGTAGGTGGTTTGATTAGTCGTTTGTTAAAAGTCCCGGCTTAACCGGGAAGATGCAGGCGAAACGGTCAGACTAGAGAGAGTGAGAGGCCGGTGGAACTCATGGTGTAGGGGTGAAATCCGTTGATATCATGGGGAACGCCAAAAGCGAAGGCAGCCGGCTGGCGCTTATCTGACACTGAGGCACGAAAGCGTGGGTAGCGAACGGG
## 484                                                          ACGGGAGTGGCAAGCGTTATCCGGAATTATTGGGCGTAAAGCGTCCGCAGGCGGTTTTACAAGTCTGTCGTTAAAGCGTGGAGCTTAACTCCATTTCAGCGATGGAAACTGTAAGACTAGAGTGTGGTAGGGGCAGAGGGAATTCCCGGTGTAGCGGTGAAATGCGTAGATATCGGGAAGAACACCAGTGGCGAAGGCGCTCTGCTGGGCCATAACTGACGCTCATGGACGAAAGCCAGGGGAGCGAAAGGG
## 485                                                          ACGGAGGGTGCAAGCGTTATCCGGATTCACTGGGTTTAAAGGGTGCGTAGGTGGGTTGATAAGTCAGTGGTGAAAGCTCCGAGCTTAACTTGGAAACTGCCATTGATACTGTCAGTCTTGAATGTCGTAGAGGTAAGCGGAATGTTGCATGTAGCGGTGAAATGCTTAGATATGTTACAGAACACCAATTGCGAAGGCAGCTTACTATGCGATGATTGACGCTGAGGCACGAAAGCGTGGGGAGCAAACAGG
## 486                                                          ACGTAGGGTGCAAGCGTTAATCGGAATTACTGGGCGTAAAGCGTGCGCAGGCGGTCTTGTAAGACAGAGGTGAAATCCCCGGGCTCAACCTGGGAACTGCCTTTGTGACTGCAAGGCTTGAGTGCGGCAGAGGGGGATGGAATTCCGCGTGTAGCAGTGAAATGCGTAGATATGCGGAGGAACACCGATGGCGAAGGCAATCCCCTGGGCCTGCACTGACGCTCATGCACGAAAGCGTGGGGAGCAAACAGG
## 487                                                          ACGTAGGGTGCGAGCGTTAATCGGAATTACTGGGCGTAAAGCGTGCGCAGGCGGTTTCGTAAGACAGAGGTGAAATCCCCGGGCTCAACCTGGGAACTGCCTTTGTGACTGCATAGCTAGAGTACGGCAGAGGGGGATGGAATTCCGCGTGTAGCAGTGAAATGCGTAGATATGCGGAGGAACACCGATGGCGAAGGCAATCCCCTGGGCCTGTACTGACGCTCATGCACGAAAGCGTGGGGAGCAAACAGG
## 488                                                          ACGTAGGGTGCGAGCGTTAATCGGAATTACTGGGCGTAAAGGGTGCGCAGGTGGTTTTGCAAGACTGACGTGAAAGCCCCGGGCTCAACCTGGGAACGGCGTTAGTGACTGCAAGACTAGAGTGTGGCAGAGGGGGGTGGAATTCCACGTGTAGCAGTGAAATGCGTAGAGATGTGGAGGAACACCGATGGCGAAGGCAGCCCCCTGGGCTAACACTGACACTCATGCACGAAAGCGTGGGGAGCAAACAGG
## 490                                                          ACGAAGGGTGCAAGCGTTGTTCGGAATCATTGGGCGTAAAGCGCGCGCAGGCGGATTACTAAGTCAGATGTGAAATCTCGGGGCTCAACCCCGAAACTGCGTCTGAAACTGGTAGTCTAGAATGTCGGAGGGGGCAGGGGAATTTCACGTGTAGGGGTAAAATCCGTAGAGATGTGAAGGAACACCGGGGGCGAAGGCGCCTGCCTGGACGACTATTGACGCTGAGGCGCGAAAGCGTGGGGAGCAAACAGG
## 491                                                          ACGAAGGGTGCAAGCGTTAATCGGAATTACTGGGCGTAAAGCGTGCGTAGATGGTTCGCTAAGTCGGATGTGAAAGCCCCGGGCTCAACCTGGGAACTGCATCCGATACTGGCGGACTAGAGTGTGATAGAGGATGGCGGAATTCCCGGTGTAGCGGTGAAATGCGTAGAGATCGGGAGGAACATCCGTGGCGAAGGCGGCCATCTGGATCAACACTGACGTTGAGGCACGAAAGCGTGGGGAGCAAACAGG
## 492                                                          ACAGAGGATGCAAGCGTTATCCGGAATCACTGGGCATAAAGCGTCTGTAGGTTGCCTACCAAGTCTGCTGTTAAAGATCAGGGCCTAACCCTGGGAAAGCAGTGGAAACTAGTAGGCTTGAGTGTGGTAGAGGTAGAGGGAATTCCTGGTGTAGCGGTGAAATGCGTAGATATTAGGAAGAACACCAATGGCGAAAGCACTCTACTGGGCCATAACTGACACTGAGAGACGACAGCTAGGGGAGCAAATGGG
## 493                                                          ACGTAGGGTGCAAGCGTTAATCGGAATTACTGGGCGTAAAGCGTGCGCAGGCGGTTATGCAAGACAGAGGTGAAATCCCCGGGCTCAACCTGGGAACTGCCTTTGTGACTGCATAGCTAGAGTACGGCAGAGGGGGATGGAATTCCGCGTGTAGCAGTGAAATGCGTAGATATGCGGAGGAACACCGATGGCGAAGGCAATCCCCTGGGCCTGCACTGACGCTCATGCACGAAAGCGTGGGGAGCAAACAGG
## 494                                                          ACGTAGGGTGCGAGCGTTAATCGGAATTACTGGGCGTAAAGCGTGCGCAGGCGGTTGCGTAAGACAGATGTGAAATCCCCGGGCTCAACCTGGGAACTGCGTTTGTGACTGCGCGACTAGAGTACGGCAGAGGGAGGTGGAATTCCACGTGTAGCAGTGAAATGCGTAGAGATGTGGAGGAACACCGATGGCGAAGGCAGCCTCCTGGGCCAGTACTGACGCTCATGCACGAAAGCGTGGGGAGCAAACAGG
## 495                                                          ACGAGTGCCTCAAGCGTTATCCGGAATCATTGGGCGTAAAGGGTGTGTAGGTGGTCATATTAGTCTTCTGTTAAATTCTTCGGCTTAACCGGGGGCATGCAGGGGAAACGGTATGACTTGAGGATGCGAGGGGTCTGTGGAACTCTAGGTGTAGCGGTGAAATGCGTTGATATCTAGGGGAACACCAAAAGCGAAGGCAGCAGACTGGAGCACTCCTGACACTGAAACACGAAAGCGTGGGTAGCGAATGGG
## 496                                                          ACGTAGGGTGCGAGCGTTAATCGGAATTACTGGGCGTAAAGCGTGCGCAGGCGGTTTCTTAAGCCAGACGTGAAATCCCCGGGCTTAACCTGGGAACTGCGTTTGGAACTGGGAGACTAGAGTGTGTCAGAGGGAGGTGGAATTCCGCGTGTAGCAGTGAAATGCGTAGATATGCGGAGGAACACCGATGGCGAAGGCAGCCTCCTGGGATGACACTGACGCTCATGCACGAAAGCGTGGGGAGCAAACAGG
## 497                                                          ACGGGAGTGGCAAGCGTTATCCGGAATTATTGGGCGTAAAGCGTCCGCAGGCGGTTTTACAAGTCTGTCGTTAAAACGTGGAGCTTAACTCCATTTCAGCGATGGAAACTGTAGAACTAGAGTGTGGTAGGGGCAGAGGGAATTCCCGGTGTAGCGGTGAAATGCGTAGATATCGGGAAGAACACCAGTGGCGAAGGCGCTCTGCTGGGCCATAACTGACGCTCATGGACGAAAGCCAGGGGAGCGAAAGGG
## 498                                                          ACAGAGGGTGCAAGCGTTGTTCGGAATTACTGGGCGTAAAGAGCGCGTAGGCGGTTCTATAAGTCAGACGTGAAAGCCCTGGGCTTAACCCAGGAGGTGCGTTTGAAACTGTAGAGCTAGAGGACGAGAGAGGTGAGTGGAATTCCTAGTGTAGAAGTGAAATTCGTAGATATTAGGAGGAACATCAGTGGCGAAGGCGACTCACTGGCTCGTAACTGACGCTGAGGCGCGAAAGCGTGGGGAGCAAACAGG
## 499                                                          ACAGAGACCTCAAGTGTTATCCGGATTTATTGGGCGTAAAGCGTCCGCAGATTGTTTGACGGGTCAGGGGTTAAAATCCAGTGCTTAACACTGGAATCGCCTTTGAAACTATCAAGCTAGAGGGTGGAAGAGGTAAGCGGAATTGCAGGTGTAGTCGTAATAAGCGTTGATATCTGCAAGAACACCAAAGGCGAAGGCAGCTTACTGGTACACATCTGACGTTCATGGACGAAAGCGTGGGGAGCAAACAGG
## 500                                                          ACGTAGGGTGCAAGCGTTAATCGGAATTACTGGGCGTAAAGCGTGCGCAGGCGGACTTTTAAGCCAGATGTGAAAGCCCCGAGCTTAACTTGGGAATTGCGTTTGGAACTGGGAGTCTAGAGTCTGTCAGAGGGGGATGGAATTCCACGTGTAGCAGTGAAATGCGTAGAGATGTGGAGGAACACCGATGGCGAAGGCAGTCCCCTGGGCCTGCACTGACGCTCATGCACGAAAGCGTGGGGAGCAAACAGG
## 501                                                          ACGGAGGGTGCAAGCGTTGTCCGGAATCATTGGGCGTAAAGAGTTCGTAGGTGGTATGTTAAGTCTGGTGTTAAAGCCCGAAGCTCAACTTCGGTTCGGCACTGGATACTGGCAAACTTGAATGTGGTAGAGGTAAAGGGAATTCCTGGTGTAGCGGTGAAATGCGTAGATATCAGGAGGAACATCGGTGGCGAAAGCGCTTTACTGGGCCATTATTGACACTGAGGAACGAAAGCCGGGGTAGCAAATGGG
## 502                                                          ACGTAGGGTGCAAGCGTTAATCGGAATTACTGGGCGTAAAGCGTGCGCAGGCGGCTTTGCAAGACAGATGTGAAATCCCCGGGCTCAACCTGGGAACTGCATTTGTGACTGCAAGGCTAGAGTACGGTAGAGGGGGATGGAATTCCGCGTGTAGCAGTGAAATGCGTAGATATGCGGAGGAACACCGATGGCGAAGGCAATCCCCTGGACCTGTACTGACGCTCATGCACGAAAGCGTGGGGAGCAAACAGG
## 503                                                          ACGAAGGGGGCTAGCGTTGCTCGGAATGACTGGGCGTAAAGGGCGCGTAGGCGGTTTGATCAGTTAGGTGTGAAATTCCTGGGCTTAACCTGGGGGCTGCACTTAATACGGTCAGGCTAGAGTGTGAAAGAGGGTCGTGGAATTCCCAGTGTAGAGGTGAAATTCGTAGATATTGGGAAGAACACCGGTGGCGAAGGCGGCGACCTGGTTCATTACTGACGCTGAGGCGCGAAAGCGTGGGGAGCAAACAGG
## 504                                                          ACGAAGGGTGCAAGCGTTACTCGGAATTACTGGGCGTAAAGCGTGCGTAGGCGGTTTGTTAAGTCTGCTGTGAAAGCCCTGGGCTCAACCTGGGAACTGCAGTGGATACTGGCGAGCTAGAGTGTGTCAGAGGATGGTGGAATTCCCGGTGTAGCGGTGAAATGCGTAGAGATCGGGAGGAACACCAGTGGCGAAGGCGGCCATCTGGGACAACACTGACGCTGAGGCACGAAAGCGTGGGGAGCAAACAGG
## 505                                                          ACAGAGACCTCAAACGTTGTCCGGATTTATTGGGCGTAAAGGGTCCGCAGGCGGTTGTGTGCGTCACGGGTTAAATTTCAGGGCTCAACCTTGGAACCGCTCGTGATACGACATGACTAGAGACTGGGAGAGGTAAGCGGAATTCACGGTGTAGTAGTAAAATGCGTTAATATCGTGAAGAACACCAAATGCGAAGGCAGCTTACTGGAACAGTTCTGACGCTCAGGGACGAAAGCGTGGGGAGCGAAAGGG
## 506                                                          ACGAAGGGTGCAAGCGTTGCTCGGAATTATTGGGCGTAAAGGGTAGGTAGGTGGTTACGTATGTCGAGAGTGAAATCCCTGGGCTTAACCCAGGAAGTGCTTTGGAAACGGCGTAACTAGAGTTCTGGAGAGGTTCGTGGAATTCCCAGTGTAGCGGTGAAATGCGTAGAGATTGGGAGGAACATCAGAGGCGAAGGCGACGAACTGGACAGATACTGACACTGAACTACGAAAGCGTGGGTAGCAAACAGG
## 508                                                          ACGAAGGTGGCAAGCGTTGTTCGGATTTACTGGGCGTACAGGGAGCGTAGGCGGTTGGGTAAGCCCTCCGTGAAATCTCCGGGCCTAACCCGGAAAGTGCGGAGGGGACTGCTCGGCTAGAGGATGGGAGAGGAGCGCGGAATTCCCGGTGTAGCGGTGAAATGCGTAGAGATCGGGAGGAAGGCCGGTGGCGAAGGCGGCGCTCTGGAACATTTCTGACGCTGAGGCTCGAAAGCGTGGGGAGCAAACAGG
## 510                                                          ACATAGGATGCAAGCGTTATCCGGAGTCACTGGGCGTAAAGGGTTCGCAGCCGGATCTATAAGTCTGTTTTCAAATTCTTTCGCTCAACGAAAGGCAGGGAATAGATACTGTAGATCTAGAGGTTTTTGGAGGGTAATGGAATTCCAGGTGGAGCGGTGAAATGTGTTGATATCTGGAAGAATGCCGAAAGCGAAAGCAGTTACCTACAATTTACCTGACGGTCAGGAACGAAAGCTTGGGGAGCAAATGGG
## 511                                                          ACGGAGGATCCAAGCGTTATCCGGATTTATTGGGTTTAAAGGGTGCGTAGGCGGCGTTGTAAGTCAGTGGTGAAAGTTTGCAGCTTAACTGTAAAATTGCCATTGATACTGCAGTGCTTGAGTACAGATGAGGTGGGCGGAATGTGTCATGTAGCGGTGAAATGCATAGATATGACACAGAACACCGATTGCGAAGGCAGCTCACTAAACTGTAACTGACGCTGAGGCACGAAAGCGTGGGGATCAAACAGG
## 512                                                          ACGTAGGGTGCGAGCGTTAATCGGAATTACTGGGCGTAAAGCGTGCGCAGGCGGTTTGTTAAGACAGTCGTGAAATCCCCGGGCTTAACCTGGGAACTGCGATTGTGACTGGCAAGCTAGAGTACGGCAGAGGGGGGTAGAATTCCTGGTGTAGCAGTGAAATGCGTAGAGATCAGGAGGAATACCGATGGCGAAGGCAGCCCCCTGGGCTAGTACTGACGCTCATGCACGAAAGCGTGGGGAGCAAACAGG
## 513                                                          ACAGAGGGTGCAAGCGTTAATCGGAATTACTGGGCGTAAAGCGCACGCAGGCGGTTGGTTAAGTCAGATGTGAAAGCCCCGGGCTCAACCTGGGAATTGCATTTGAAACTGGCCAACTAGAGTACGTGAGAGGGGGGTAGAATTCCAAGTGTAGCGGTGAAATGCGTAGAGATTTGGAGGAATACCAGTGGCGAAGGCGGCCCCCTGGCACGATACTGACGCTCAGGTGCGAAAGCGTGGGGAGCAAACAGG
## 514                                                          ACAGAGGGTGCAAGCGTTATTCGGAATTACTGGGCATAAAGAGCGCGTAGGCGGTCTAATAAGTCTGATGTGAAATCCCTGGACTCAATTCAGGAAGTGCATCGGATACTGCTAGACTAGAGTGCAGGAGAGGAGAGCGGAATTCCTGGTGTAGAGGTGAAATTCGTAGATATCAGGAGGAACAACAGTGGCGAAGGCGGCTCTCTGGACTGTAACTGACGCTGAGGCGCGAAAGCGTGGGGAGCAAACAGG
## 515                                                          ACGAGTGGTCCAAGCGTTATCCGGAATTATTGGGCGTAAAGAGTGCGTAGGTGGATATACTAGTCGGTTGTTTAACCCACGGGCTCAACCTGTGGATTGCGAGCGAAACGGTATATCTTGAGTATGTTAGGGGTAAGCGGAACTCATAGTGTAGGGGTGAAATCCGTTGATATTATGGGGAACACCAAAAGCGAAGGCAGCTTACTGGAACATTACTGACACTGAGGCACGAAAGCGTGGGTAGCGAATGGG
## 516                                                          ACGTAGGGGGCAAGCGTTATCCGGATTTACTGGGTGTAAAGGGAGCGCAGACGGTTATGCAAGTCTGGAGTGAAACCCCACAGCTCAACTGTGGGCTTGCTTTGGAAACTGTGTAACTAGAGTACTGGAGAGGTAAGCGGAATTCCTAGTGTAGCGGTGAAATGCGTAGATATTAGGAGGAACACCGGTGGCGAAGGCGGCTTACTGGACAGCAACTGACGTTGAGGCTCGAAGGCGTGGGGAGCAAACAGG
## 517                                                          ACAGAGGGTGCGAGCGTTAATCGGAATTACTGGGCGTAAAGGGTGCGTAGGTGGTTATGTAAGTTAGATGTGAAATTTCCGGGCTTAACCTGGGAACTGCGTCTGATACTGCATAGCTAGAGTCCTGTAGAGGAAAGCGGAATTTCCGGTGTAGCAGTGAAATGCGTAGATATCGGAAGGAACATCAGTGGCGAAGGCGGCTTTCTGGACAGAGACTGACACTGAGGCACGAAAGCGTGGGGAGCAAACAGG
## 518                                                          ACGAAGGTGGCGAGCGTTACTCGGATTTACTAGGCGTAAAGCGTGGGCAGGCGGCTCGATAAGTCTCTTGTGAAAGCCTTGGGCTTAACCCAGGGAGGCCAAGAGATACTATCGGGCTTGGATGTGGGAGAGGAGACTGGAATTCCCGGTGTAGCGGTGAAATGCGTTGATATCGGGAGGAACACCAATGGCGAAAGCAAGTCTCTGGACCACCATCGACGCTCATCCACGAAAGCTGGGGGATCAAACAGG
## 519                                                          ACGGGGGATGCAAGTGTTATCCGGATTTACTGGGCGTAAAGCGTCTGCAGGTGGTTTCTTAAGTCTACTGTTAAATCTTGAGGCTCAACCTCAAATCTGCAGTAGAAACTAGGAGACTTGAGTATAGTAGGGGTAGAGGGAATTTCCAGTGGAGCGGTGAAATGCGTAGATATTGGAAAGAACACCGATGGCGAAGGCACTCTACTGGGCTATTACTGACACTCAGAGACGAAAGCTAGGGGAGCAAATGGG
## 520                                                          ACGGAGGGGGCTAGCGTTATTCGGAATTACTGGGCGTAAAGCGCACGTAGGCGGACTGAAAAGTCAGAGGTGAAATCCCAGGGCTCAACCTTGGAACTGCCTTTGAAACTCTCAGTCTTGAGGTCGTGAGAGGTGAGTGGAATTCCGAGTGTAGAGGTGAAATTCGTAGATATTCGGAGGAACACCAGTGGCGAAGGCGGCTCACTGGCACGATACTGACGCTGAGGTGCGAAAGCGTGGGGAGCAAACAGG
## 521                                                          ACGAAGGGTGCGAGCGTTAATCGGAATTACTGGGCGTAAAGCGCGCGTAGGTGGTTTTGTAAGCGGAATGTGAAATCCCCGGGCTCAACCTGGGAACTGCATGCCGAACTGCAAGGCTAGAGTATAGTAGAGGGGTGTGGAATTCCCGGTGTAGCGGTGAAATGCGTAGATATCGGGAGGAACATCAGTGGCGAAGGCGGCACCCTGGACTGATACTGACACTGAGGCGCGAAAGCGTGGGTAGCGAACAGG
## 522                                                          ACATGTGGGGCAAGCGTTGTTCGGAATTACTGGGCATAAAGGGTGCGTAGGCGGCGGTATAAGTCAGCGGTGAAATCCTTCGGCCCAACCGGAGAACTGCCGTTGATACTGTACTGCTTGAGTGCGTGAGGGGTGGCTGGAACGAGTGGTGTAGCGGTGAAATGCGTAGATATCACTCGGAACACCCAAGGCGAAGGCAGGCCACTGGCACGCAACTGACGCTGAGGCACGAAAGCGTGGGGAGCAAACAGG
## 524                                                          ACGTAGGGTGCGAGCGTTAATCGGAATTACTGGGCGTAAAGCGTGCGCAGGCGGTTTGGTAAGTCAGATGTGAAATCCCCGGGCTCAACCTGGGAACTGCATTTGAGACTGCCAAGCTGGAGTTTGGCAGAGGGGGGTGGAATTCCACGTGTAGCAGTGAAATGCGTAGAGATGTGGAGGAACACCGATGGCGAAGGCAGCCCCCTGGGCCAATACTGACGCTCATGCACGAAAGCGTGGGGAGCAAACAGG
## 525                                                          ACGTAGGGTGCGAGCGTTAATCGGAATTACTGGGCGTAAAGCGTGCGCAGGCGGTTTTGTAAGACAGACGTGAAATCCCCGGGCTCAACCTGGGAATGGCGTTTGTGACTGCAAAGCTGGAGTGCGGCAGAGGGGGATGGAATTCCGCGTGTAGCAGTGAAATGCGTAGATATGCGGAGGAACACCGATGGCGAAGGCAATCCCCTGGGCCTGCACTGACGCTCATGCACGAAAGCGTGGGGAGCAAACAGG
## 526                                                          ACGAGGGGTGCAAGCGTTGCTCGGAATTATTGGGCGTAAAGGGTAGGTAGGTGGTCCTGTTTGTCAGAGGTGAAATCCTTGGGCTTAACCCAAGAAGTGCCTCTGAAACGGCAGGACTCGAGTTCTGGAGAGGGTCGTGGAATTCCCGGTGTAGCGGTGAAATGCGTAGAGATCGGGAGGAACACCAGAGGCGAAGGCGGCGACCTGGACAGATACTGACACTCAACTACGAAAGCGTGGGTAGCAAACAGG
## 527                                                          ACGTAGGGTGCAAGCGTTAATCGGAATTACTGGGCGTAAAGCGTGCGCAGGCGGTTATGTAAGACAGAGGTGAAATCCCCGGGCTCAACCTGGGAACTGCCTTTGTGACTGCATAGCTAGAGTACGGTAGAGGGGGATGGAATTCCGCGTGTAGCAGTGAAATGCGTAGATATGCGGAGGAACACCGATGGCGAAGGCAATCCCCTGGGCCTGCACTGACGCTCATGCACGAAAGCGTGGGGAGCAAACAGG
## 528                                                          ACGTAGGGTGCAAGCGTTGTCCGGATTTATTGGGCGTAAAGAGCTCGTAGGCGGCCTGACACGTCGGATGTGAAAACACGGTGCTTAACACCGTGCCTGCATTCGATACGGTCAAGCTAGAGGATGTCAGGGGAAACTGGAATTCCTGGTGTAGCGGTGAAATGCGCAGATATCAGGAGGAACACCGGTGGCGAAGGCGGGTTTCTGGGACATTACTGACGCTGAGGAGCGAAAGCGTGGGGAGCAAACAGG
## 529                                                          ACGGAGGGTGCAAGCGTTACTCGGAATCACTGGGCGTAAAGAGCGTGTAGGCGGGTATATAAGTCAGAAGTGAAATCCAATAGCTTAACTATTGAACTGCTTTTGAAACTGTATACCTAGAATGTGGGAGAGGTAGATGGAATTTCTGGTGTAGGGGTAAAATCCGTAGAGATCAGAAGGAATACCGATTGCGAAGGCGATCTACTGGAACATTATTGACGCTGAGACGCGAAAGCGTGGGGAGCAAACAGG
## 530                                                          ACGGAGGGAGCTAGCGTTGTTCGGAATTACTGGGCGTAAAGCGCACGTAGGCGGCTACTCAAGTCAGAGGTGAAAGCCCAGGGCTCAACCCTGGAACTGCCTTTGAAACTAGGTAGCTAGAATCTTGGAGAGGTCAGTGGAATTCCGAGTGTAGAGGTGAAATTCGTAGATATTCGGAAGAACACCAGTGGCGAAGGCGACTGACTGGACAAGTATTGACGCTGAGGTGCGAAAGCGTGGGGAGCAAACAGG
## 531                                                          ACGAGAGGTGCAAGCGTTATTCGGAATTATTGGGCGTAAAGGGCGCGTAGGCGGCTTTATAAGTTTCTTGTTTAAGACTTCGGCCTAACTGAAGAAAAGCAAGGAATACTGTATAGCTAGAGTATAATAGAGAGAAGTGGAATTTTCGGAGTAGCGGTTAAATGCGTAGATCTCGAAAGGAACGCCAATGGCGAAGGCAGCTTCTTGGGTTATTACTGACGCTGAGGCGCGAAGGCATGGGGAGCAAACAGG
## 532                                                          ACGGAGGGGGCTAGCGTTGTTCGGAATGACTGGGCGTAAAGGGCGCGTAGGCGGTTTTTTAAGTGAGGCGTGAAAGCCCTGGGCTTAACCCAGGAGGTGCGTTTCATACTGGAAGACTTGAGTACGAGAGAGGAAAGCGGAATTCCTAGTGTAGAGGTGAAATTCGTAGATATTAGGAAGAACACCAGTGGCGAAGGCGGCTTTCTGGCTCGTAACTGACGCTGAGGCGCGAAAGCGTGGGGAGCAAACAGG
## 533                                                          ACGAAGGTCCCGAGCGTTGTTCGGAATCACTGGGCGTAAAGGGAGCGTAGGCGGCGTGGTAAGTCAGATGTGAAATCCCGGGGCTCAACCCCGGAACTGCATCCGATACTGCCGTGCTAGAGGATTGGAGAGGTAGCTGGAATTCACGGTGTAGCAGTGAAATGCGTAGATATCGTGAGGAACACTCGTGGCGAAAGCGAGCTACTGGACAACTCCTGACGCTGAGGCTCGAAGGCCAGGGTAGCGAAAGGG
## 534                                                          ACGAAGGGTGCAAGCGTTGCTCGGAATTATTGGGCGTAAAGGGTAGGTAGGTTGTTACGTATGTCTGGGGTGAAATCCCTGAGCTTAACTCAGGAAGTGCCTTGGAAACGGCGTAACTAGAGTGCTAGAGAGGTTCGTGGAATTCCCAGTGTAGCGGTGAAATGCGTAGAGATTGGGAGGAACACCAGAGGCGAAGGCGGCGAACTGGATAGCAACTGACACTGAACTACGAAAGCGTGGGTAGCAAACAGG
## 535                                                          ACGAAGGGGGCTAGCGTTGCTCGGAATTACTGGGCGTAAAGGGCGCGTAGGCGGCTTAGTCAGTCAGGCGTGAAATTCCTGGGCTTAACCTGGGGGCTGCGCTTGATACGGCTAGGCTAGAGGGTAGGAGAGGGTCGCGGAATTCCCAGTGTAGAGGTGAAATTCGTAGATATTGGGAAGAACACCGGTGGCGAAGGCGGCGACCTGGCCTATTACTGACGCTGAGGCGCGACAGCGTGGGGAGCAAACAGG
## 536                                                          ACAGAGGGTGCGAGCGTTGTCCGGAATCACTGGGCGTAAAGGGCGCGTAGGTGGCTTGGTAAGCGTGTGGTGAAAGTCCGGGGCTCAACCCCGGATCTGCCGTGCGAACTGCCGAGCTTGAGCATTGTAGAGGCAGGTGGAATTCCGGGTGTAGCGGTGGAATGCGTAGAGATCCGGAAGAACACCAGTGGCGAAGGCGGCCTGCTGGGCAATTGCTGACACTGAGGCGCGACAGCGTGGGGAGCAAACAGG
## 537                                                          ACAGAGACCTCAAGCATTATCCGGAATTATTGGGCGTAAAGGGTCCGCAGGTGGCTCGGTAAGTCATGCGTCAAATCTTTGGGCTCAACCCAGAGGCTGCGTGTGATACTGCTGAGCTTGAGGCCGGGAGAGGTGCATGGAATTACTGGTGTAGCGGTAAAATGCGTTAAGATCAGTAAGAACACCAAAGGCGAAGGCAATGCACTGGAACGGTCCTGACACTCAGGGACGAAAGCGTGGGGAGCGAAAGGG
## 539                                                          ACGTAGGTCTCGAGCGTTATCCGGAATTACTGGGCGTAAAGCGTCCGCAGCCGGATGTATAAGTCTGTCTTTAAATATCTCGATTCAATCGAGAAAAAGGGATAGATACTGTATGTCTAGAGGACTTTAGAGGTTAGTGGAATTTCCGGTGGAGCGGTGAAATGCGTTGATATCGGAAGGAATGCCAAAAGCGAAAGCAGCTATCTATAAAGTATCTGACGGTCAGGGACGACAGCTTGGGGAGCAAACGGG
## 540                                                          ACAGAGACTGCAAGCGTTATTCGGATTCACTGGGCGTAAAGGGTGCGCAGGCGGTTGCATGTGTTGGGTGTGAAAGCCCGGGGCTCAACCCCGGAATTGCGCCCAAAACTATGCGACTAGAGTACCGGAGAGGATAGCGGAATTCACGGTGTAGCAGTGAAATGCGTAGATATCGTGAAGAACACCAGCGGCGAAGGCGGCTATCTGGAAGGTAACTGACGCTCAGGCACGAAAGCGTGGGGAGCAAAAGGG
## 541                                                          ACGTAGGGTGCGAGCGTTAATCGGAATTACTGGGCGTAAAGCGTGCGCAGGCGGTTTTGTAAGACAGGTGTGAAATCCCCGGGCTTAACCTGGGAACTGCGCTTGTGACTGCAAGACTTGAGTGCGGCAGAGGGGGGTGGAATTCCACGTGTAGCAGTGAAATGCGTAGAGATGTGGAGGAACACCGATGGCGAAGGCAGCCCCCTGGGTCGACACTGACGCTCATGCACGAAAGCGTGGGGAGCAAACAGG
## 542                                                          ACAGGGGATGCAAGTGTTATCCGGAATTATTGGGCGTAAAGCGTCTGCAGGTGGATTTATAAGTCTTTTGTTAAATCTTTGAGCTTAACTCAAAATCGGCAAAAGAAACTATTTATCTTGAGTATGGTAGAGGTAAAGGGAATTTCCAGTGGAGCGGTGAAATGCGTAGATATTGGAAAGAACACCAACAGCGAAGGCACTTTACTGGGCCATTACTGACACTCAGAGACGAAAGCTAGGGGAGCAAACGGG
## 543                                                          ACGGAGGGTGCAAGCGTTATCCGGATTCACTGGGTTTAAAGGGTGCGTAGGCGGGCAGGTAAGTCAGTGGTGAAATCTTCGAGCTTAACTCGGAAACTGCCGTTGATACTATCTGTCTTGAATATAGTGGAGGTAAGCGGAATATGTCATGTAGCGGTGAAATGCTTAGAGATGACATAGAACACCAATTGCGAAGGCAGCTTGCTACACTATTATTGACGCTGAGGCACGAAAGCGTGGGGATCAAACAGG
## 544                                                          ACGTAGGGTGCGAGCGTTAATCGGAATTACTGGGCGTAAAGCGTGCGCAGGCGGTTGTGTAAGCCAGATGTGAAATCCCCGGGCTTAACCTGGGAATGGCATTTGGGACTGCACGGCTGGAGTATGGCAGAGGGGACTGGAATTCCTGGTGTAGCAGTGAAATGCGTAGATATCAGGAGGAATACCGATGGCGAAGGCAGGTCCCTGGGCCAATACTGACGCTCATGCACGAAAGCGTGGGGAGCAAACAGG
## 545                                                          ACAGGGGATGCAAGTGTTATCCGGATTCATTGGGCGTAAAGCGTCTGCAGGTGGATATATAAGTCTGTTGTTAAATCTTTGGGCTTAACCCAAAATCTGCAAAAGAAACTATATATCTTGAGTATGGTAGAGGTAAAGGGAATTTCCGGTGGAGCGGTGAAATGCGTAGATATCGGAAAGAACACCAACTGCGAAGGCACTTTACTGGGCCAGTACTGACACTCAGAGACGAAAGCTAAGGGAGCAAACAGG
## 546                                                          ACGGAGGGGGCTAGCGTTATTCGGAATTACTGGGCGTAAAGCGCACGTAGGCGGACTGAAAAGTCAGAGGTGAAATCCCAGGGCTCAACCTTGGAACTGCCTTTGAAACTCTCAGTCTTGAGGTCGAGAGAGGTGAGTGGAATTCCGAGTGTAGAGGTGAAATTCGTAGATATTCGGAGGAACACCAGTGGCGAAGGCGGCTCACTGGCTCGATACTGACGCTGAGGTGCGAAAGCGTGGGGAGCAAACAGG
## 547                                                          ACGAAGGATCCAAGCGTTGTCCGGATTTACTGGGTTTAAAGGGTGCGTAGGCGGGAGATTAAGTCAGTGGTGAAAGCCCTCAGCTCAACTGAGGAACTGCCATTGAAACTGATTTTCTTGAATATGGTTGAGGTAGATGGAATATAACATGTAGCGGTGAAATGCTTAGATATGTTATAGAACACCGATTGCGAAGGCAGTCTGCTAAGCCATTATTGACGCTGAGGCACGAAAGCGTGGGGAGCAAACAGG
## 548                                                           ACGTAGGTGGCAAGCGTTGTCCGGAATTATTGGGCGTAAAGCGCGCGCAGGCGGGTTTTTAAGTCTGTCTTAAAAGTTCGGGGCTTAACCCCGTGAGGGGATGGAAACTGGAAACCTAGAGTATCGGAGAGGAAAGCGGAATTCCTAGTGTAGCGGTGAAATGCGTAGATATTAGGAAGAACACCAGTGGCGAAGGCGGCTTTCTGGACGAAAACTGACGCTGAGGCGCGAAAGCCAGGGGAGCAAACGGG
## 549                                                         ACGTAGGTGGCGAGCGTTATCCGGATTTATTGGGTTTAAAGGGAGCGTAGGCGGCTGATTAAGTCAGCGGTGAAAGGTAGCAGCTTAACTGTTTTACATGCCGTTGATACTGGTTAGCTTGAGTTGACAGAAGGCAGATAGAATTCCTGGTGTAGCGGTGAAATGCTTAGATACCAGGAGGAATACCGATTGCGAAGGCAGTCTGCTGCAGTCACACTGACGCTGAGGCTCGAAGGTGCGGGGATCAAACAGG
## 550                                                          ACGAAGGTCCCGAGCGTTATTCGGAATCACTGGGCGTAAAGGGAGCGTAGGCGGCGTGGAAAGTCAGATGTGAAATCCCGGGGCTCAACCCCGGAACTGCATCCGATACTACCATGCTAGAGTATTGGAGGGGTAACTGGAATTCTCGGTGTAGCAGTGAAATGCGTGGATATCGAGAGGAACACCTGTGGCGAAAGCGAGTTACTGGACAATTACTGACGCTGAGGCTCGAAGGCTAGGGGAGCAAAAGGG
## 551                                                        ACGTAATCCCTAAGTGGTATCCACGAATATTGGGCCTAAAGCGTTCGTAGCCGGCTGTATAAGTCCTTGGTGAAAGGGGCTTGCTTAACTTGTCCAATGGCTAAGGATACTGTACGGCTGAGAGACCAGTAGAGGTGCATGGAATTACGGGGGTAACGGTGAAATGTTGTGATCCTCGTAGGACCACCGGTAGCGAAGGCGATGCACCAGGATGGTTCTGACGGTGAGGAACGAAAGCGGGGGGAGCGACCCGG
## 552                                                          ACGTAGGTGGCAAGCGTTGTCCGGAATTATTGGGCGTAAAGGGCGTGTAGGTGGATTCTTAAGTCTTGTGTCTAAGTGCGGGGCTCAACCCCGTATGGGCGCAGGAAACTGAGAATCTTGAGTGCAGGAGAGGAAAGTGGAATTCCCAGTGTAGCGGTGAAATGCGTAGATATTGGGAGGAACACCAGTGGCGAAGGCGACTTTCTGGACTGTGTCTGACACTGAGGCGCGAAAGCCAGGGGAGCGAACGGG
## 553                                                          ACGGAGGGTGCGAACGTTATCCGGAATCACTGGGTTTAAAGGGTGCGTAGGCTGTTTAATAAGTCAGTGGTGAAAGGCCGTCGCTTAACGATGGGACTGCCATTGATACTGTTAGACTTGAATCAGGTTGAGGTTGGCGGAATGTGGCATGTAGCGGTGAAATGCATAGATATGCCATGGAATACCGATTGCGAAGGCAGCTGACTGGACCTTGATTGACGCTGAGGCACGAAAGCGTGGGTAGCGAACAGG
## 554                                                          ACGTAGGGTGCAAGCGTTAATCGGAATTACTGGGCGTAAAGCGTGCGCAGGCGGTTTTGTAAGACAGGCGTGAAATCCCCGGGCTTAACCTGGGAATGGCGTTTGTGACTGCAAGACTAGAGTGTGTCAGAGGGGGGTAGAATTCCACGTGTAGCAGTGAAATGCGTAGAGATGTGGAGGAACACCGATGGCGAAGGCAGCCCCCTGGGATAACACTGACGCTCATGCACGAAAGCGTGGGGAGCAAACAGG
## 555                                                         ACGTAGGGGGCGAGCGTTGTCCGGATTTATTGGGCGTAAAGAGCTCGTAGGCGGTTCGGTAAGTCGGGTGTGAAACCTCCAGGCTCAACCTGGAGACGCCACCTGATACTGCTGTGACTAGAGTCCGGTAGGGGAGTGTGGAACTCCTGGTGTAGCGGTGAAATGCGCAGATATCAGGAAGAACACCAGCGGCGAAGGCGGCACTCTGGGCCGGTACTGACGCTGAGGAGCGAAAGCGTGGGTAGCAAACAGG
## 556                                                           ACGAAGGCCCCAAGCGTTATCCGGAATTACTGGGCGTAAAGCGTCTGTAGGTGGTCTGGCAAGTCTCGTGTGAAACTTCAAAGCTCAACTTTGAATTGCACGAGAAACTATCAGACTTGAGTGTGGGAGAGGCAAGCAGAACGGTATGAGTAGGGGTGCAATCCGTTGATACATACCAGAATACCAAAAGCGAAGGCAGCTTGCTGGAACACAACTGACACTGAGAGACGAAAGCGTGGGGAGCGAAAGGG
## 557                                                          ACAGAGGATGCAAGCGTTATCCGGAATGATTGGGCGTAAAGCGTCTGTAGGTGGCTTTCCAAGTCTTTTGTCAAATCCCAGAGCTTAACTTTGGATCGGCACGAGAAACTCGAGAGCTTGAGTACGGTAGGGGCAGAAGGAATTCCCCGTGTAGCGGTGAAATGCGTAGAGATGGGGAAGAACACCGATGGCGAAAGCATTCTGCTGGGCCGTTACTGACACTGAGAGACGAAAGCTAAGGGAGCGAATAGG
## 559                                                          ACGAAGGGTGCTAGCGTTGTTCGGAATCATTGGGCGTAAAGCGTGCGTAGGCGGCTTAGTAAGTCGTTTGTGAAATCCCAAGGCTTAACCTTGGAATTGCAATCGAAACTGCTTTGCTAGAGTATGGTGGGGGATAGTGGAATTCCTAGTGTAGGGGTGAAATCCGTAGAGATTAGGAGGAACATCAGTGGCGAAAGCGACTATCTACGCCAATACTGACGCTGAGGTACGAAAGCGTGGGGAGCAAACAGG
## 560                                                          ACGTAGGGTGCAAGCGTTAATCGGAATTACTGGGCGTAAAGCGTGCGCAGGTGGTTGTGCAAGACAGATGTGAAATCCCCGGGCTTAACCTGGGAACTGCATTTGTGACTGCACAGCTAGAGTGCGGCAGAGGGGGATGGAATTCCGCGTGTAGCAGTGAAATGCGTAGATATGCGGAGGAACACCGATGGCGAAGGCAATCCCCTGGGCCTGCACTGACACTCATGCACGAAAGCGTGGGGAGCAAACAGG
## 561                                                          ACAGAGGGTGCGAGCGTTAATCGGAATGACTGGGCGTAAAGGGTGCGTAGGTGGTTTGTTAAGTTAGCTGTGAAAGCCCCGGGCTTAACCTGGGAATTGCAGCTAATACTGGTGAACTTGAGTAGTTCAGAGGAGTGTGGAATTTCCGGTGTAGCGGTGAAATGCGTAGAGATCGGAAAGAACACCAGTGGCGAAGGCGGCACTCTGGGAACATACTGACACTGAGGCACGAAGGCGTGGGGAGCAAACAGG
## 562                                                          ACGTAGGGTGCGAGCGTTAATCGGAATTACTGGGCGTAAAGCGTGCGCAGGCGGTTATGTAAGACAGATGTGAAATCCCCGGGCTCAACCTGGGACCTGCATTTGTGACTGCATGGCTGGAGTACGGCAGAGGGGGATGGAATTCCGCGTGTAGCAGTGAAATGCGTAGATATGCGGAGGAACACCGATGGCGAAGGCAATCCCCTGGGCCTGTACTGACGCTCATGCACGAAAGCGTGGGGAGCAAACAGG
## 563                                                          ACGAAGGATCCAAGCGTTGTCCGGATTTACTGGGTTTAAAGGGTGCGTAGGCGGACCGATAAGTCAGTGGTGAAAGCCCATAGCTCAACTATGGAACTGCCATTGAAACTGCCGGTCTTGAATTCGGTTGAGGCAGATGGAATATAACATGTAGCGGTGAAATGCTTAGATATGTTATAGAACACCGATTGCGAAGGCAGTCTGCTAAGTCGACATTGACGCTGAGGCACGAAAGCGTGGGGAGCGAACAGG
## 564                                                          ACGTAGGGTGCGAGCGTTAATCGGAATTACTGGGCGTAAAGGGTGCGCAGGCTGGAATACAAGTTAGATGTGAAATACCCGAGCTTAACTTGGGAGGTGCATTTAAGACTGTATACCTAGAGTTAGTCAGAGGATGGGGGAATTCCGAGTGTAGCAGTGAAATGCGCAGATATTCGGAGGAACACCGGTGGCGAAGGCGCCCATCTGGGATATAACTGACGCTCATGCACGAAAGCGTGGGGAGCAAACAGG
## 566                                                          ACGTAGGGTGCGAGCGTTAATCGGAATTACTGGGCGTAAAGCGTGCGCAGGCGGTTTTTTAAGACAGGCGTGAAATCCCCGGGCTTAACCTGGGAACTGCGCTTGTGACTGGAAGGCTAGAGTACGGCAGAGGGGGGTGGAATTCCACGTGTAGCAGTGAAATGCGTAGAGATGTGGAGGAACACCGATGGCGAAGGCAGCCCCCTGGGCCGATACTGACGCTCATGCACGAAAGCGTGGGGAGCAAACAGG
## 567                                                          ACGAAGGGGGCTAGCGTTGTTCGGAATCACTGGGCGTAAAGCGCACGTAGGCGGATCTTTAAGTCAGAGGTGAAATCCCAAGGCTCAACCTTGGAACTGCCTTTGATACTGGGGATCTCGAGTCCGGGAGAGGTGAGTGGAACTGCGAGTGTAGAGGTGAAATTCGTAGATATTCGCAAGAACACCAGTGGCGAAGGCGGCTCACTGGCCCGGTACTGACGCTGAGGTGCGAAAGCGTGGGGAGCAAACAGG
## 569                                                          ACGAGGGGTCCTAGCGTTGTTCGGAATCACTGGGCGTAAAGCGCATGTAGGTGGCTTTTTAAGTCAGGTGTGAAATCCCACGGCTCAACCGTGGAGGTGCACTTGATACTGGGAAGCTTGAGTACGAGAGAGGATGGTAGAATTCCTGGTGTAGTGGTGAAATACGTAGATATCAGGAGGAACACCGGTGGCGAAGGCGGCCATCTGGCTCGATACTGACACTGAGATGCGAAAGCGTGGGGAGCAAACAGG
## 571                                                          ACGAAGGGGGCTAGCGTTGTTCGGAATTACTGGGCGTAAAGCGCACGTAGGCGGGTCGTTAAGTCAGGGGTGAAATCCCGGAGCTCAACTCCGGAACTGCCTTTGATACTGGCGATCTTGAGATCGGAAGAGGTGAGTGGAATTCCCAGTGTAGAGGTGAAATTCGTAGATATTGGGAAGAACACCAGTGGCGAAGGCGGCTCACTGGTCCGATACTGACGCTGAGGTGCGAAAGCGTGGGGAGCAAACAGG
## 572                                                                                                               ACAGACACTGCAAGCATCTTACAACATAACTAGGCAAAGAGCGTTGTTTTGCAAAGTCGAAGCGAAATTATTGCTCAGGAATTTTCAAAGAATCAGTTAAATGAGAGGATATTGAAAGGAACGCCGAAGGTGAAGACAAGAGCATAAATCTTCTAAAGCAACTCAACGAAAGTTCAAGGAGCAAATCGGATTAGATACC
## 574                                                          ACGTAGGGTGCGAGCGTTAATCGGAATTACTGGGCGTAAAGCGTGCGCAGGCGGTTTTGTAAGACAGATGTGAAATCCCCGGGCTTAACCTGGGAACTGCGTTTGTGACTGCAAGGCTAGAGTACGGCAGAGGGGGGTGGAATTCCTGGTGTAGCAGTGAAATGCGTAGAGATCAGGAGGAACACCGATGGCGAAGGCAGCCCCCTGGGCCTGTACTGACGCTCATGCACGAAAGCGTGGGGAGCAAACAGG
## 575                                                          ACGGAGGGTGCGAGCGTTAATCGGAATTACTGGGCGTAAAGCGCGCGTATGTGGTCTGATAAGTCGGATGTGAAAGCCCCGAGCTTAACTTGGGAACGGCATTCGATACTGTCTGACTAGAGTATGGGAGAGGAGAGCGGAATTCCCGGTGTAGCGGTGAAATGCGTAGATATCGGGAGGAACATCAGTGGCGAAGGCGGCTCTCTGGACTAATACTGACACTGAGGCGCGAAAGCGTGGGTAGCAAACAGG
## 576                                                          ACATAGGGTGCAAGCGTTGTCCGGAATTATTGGGCGTAAAGAGCTCGTAGGTCGTTTGTTACGTCGGATGTGAAAACCTGAGGCTCAACCTCAGGCCTGCATTCGATACGGGCAAACTAGAGTTTGGTAGGGGAGACTGGAATTCCTGGTGTAGCGGTGGAATGCGCAGATATCAGGAGGAACACCAATGGCGAAGGCAGGTCTCTGGGCCAATACTGACACTGAGGAGCGAAAGCGTGGGGAGCGAACAGG
## 577                                                          ACGAAGGGTGCTAGCGTTGTTCGGAATCACTGGGCGTAAAGCGCGCGTAGGCGGTTTAACAAGTCAGGGGTGAAAGCCTGGAGCTTAACTCCAGAACTGCCTTTGAAACTGTTAAACTAGAGTGTCGGAGGGGATAGCGGAATTGCTAATGTAGAGGTGAAATTCGTAGATATTAGCAGGAACACCGGTGGCGAAGGCGGCTATCTGGACGACAACTGACGCTGAGGCGCGAAAGCGTGGGGATCAAACAGG
## 578                                                          ACGTAGGGTGCGAGCGTTAATCGGAATTACTGGGCGTAAAGCGTGCGCAGGCGGTTGTGTAAGTCAGATGTGAAATCCCCGGGCTCAACCTGGGAACTGCATTTGAGACTGCACGGCTAGAGTGTGGCAGAGGGGGGTGGAATTCCACGTGTAGCAGTGAAATGCGTAGAGATGTGGAGGAACACCGATGGCGAAGGCAGCCCCCTGGGCCAATACTGACGCTCATGCACGAAAGCGTGGGGAGCAAACAGG
## 579                                                          ACGTAGGGTCCAAGCGTTAATCGGAATTACTGGGCGTAAAGCGTGCGCAGGTGGTTTGTTAAGCACGATGTGAAATCCCCGAGCTCAACTTGGGAATTGCATTGTGAACTGGCTAACTAGAGTACGGCAGAGGGGGGTGGAATTCCACGTGTAGCAGTGAAATGCGTAGAGATGTGGAGGAACACCGATGGCGAAGGCAACCCCCTGGGCTGATACTGACACTCATGCACGAAAGCGTGGGGAGCAAACAGG
## 580                                                          ACGGAGGGTGCAAACGTTGCTCGGAATTATTGGGCGTAAAGCGCATGTAGGCGGTCCGCTAAGTCGGATGTGAAAGCCCTCGGCTTAACCGAGGAAGTGCATCCGAAACTGGCGGGCTAGAGTACGGAAGAGGGTTTCGGAATTCCCGGTGTAGAGGTGAAATTCGTAGATATCGGGAGGAACACCAGTGGCGAAGGCGGAAACCTGGGCCGATACTGACGCTGAGATGCGAAAGCGTGGGGAGCAAACAGG
## 581                                                         ACGAGTGCCTCGAGCGTTATCCGGAATTATTGGGCGTAAAGGGTGTGTAGGTGGTAGTGTTAGTCTTGCGTTAAATCTATTGGCTCAACTGATAGGCTGCGCGGGAAACGGCACAACTGAGAGGATGGTAGGGGTCTGCGGAACTCATAGTGTAGCGGTGAAATGCGTTGATATTATGGGGAACACCAAAAGCGAAGGCAGCAGACTGGACCACTCCTGACACTGAAACACGAAAGCGTGGGTCGCGAATGGG
## 582                                                          ACGTAGGGTGCGAGCGTTAATCGGAATTACTGGGCGTAAAGCGTGCGCAGGCGGTTCTATAAGACAGATGTGAAATCCCCGGGCTTAACCTGGGAACTGCGTTTGTGACTGTAGGACTGGAGTGCGGCAGAGGGGGGTGGAATTCCACGTGTAGCAGTGAAATGCGTAGAGATGTGGAGGAACACCGATGGCGAAGGCAGCCCCCTGGGTCGACACTGACGCTCATGCACGAAAGCGTGGGGAGCAAACAGG
## 583                                                         ACGTAGGGTGCAAGCGTTACCCGGTCTTACTGGGCGTAAAGAGTTCTGTAGGCTGTTGTGTAAGTCAGGGGTTAAATACCTCGGCCCAACCGAGGAAAAGCTTTTGATACTGCATGGCTAGAGGAGGTTAGGGGGCAACGGAACGTATGGTGTAGCAGTGAAATGCGTTGATATCATACGGAACACCAAGGGGGAAGCCAGTTGCCTGGGACCTACCTGACGCTGAGAGACGAAAGCGTGGGGAGCGAAACGG
## 584                                                          ACATAGGTCCCAAGCGTTATCCGGAATTACTGGGCGTAAAGCGTCTGCAGGTGGAAAAGTGTGTGAGATGTGAAAGACCGGGGCTTAACCCCGTGTTTGTGTCTCAAACTGCTTTTCTTGAGTGAGCAAGAGGTATGCGGAATTTATGGAGTAGGAGTGCAATCCGTTGACACCATAAAGAACACCAAAAGCGAAGGCAGCATACTGGGGCTCTACTGACACTCAGAGACGAAAGCGTGGGGAGCGAAAGGG
## 585                                                          ACGGAGGGTGCAAGCGTTATCCGGAATCACTGGGTTTAAAGGGTACGTAGGCGGTTTGATAAGTCAGGTGTGAAATACCGCGGCCTAACTGCGGAACTGCACTTGATACTGTCAGACTTGAGTAGGGTGGAGGTACACGGAATGCATCATGTAGCGGTGAAATGCTTAGATATGATGTAGAACACCGATAGCGAAGGCAGTGTGCCATACCTTAACTGACGCTGAGGTACGAAAGCGTGGGGAGCAAACAGG
## 586                                                          ACGGAGGGTGCAAGCGTTGTTCGGATTTACTGGGCGTAAAGCGTGTCTAGGTGGATTTGCAAGTCAGATGTGAAAGCCCCCGGCTCAACCGGGGAGGGTCATTTGATACTACAGATCTTGAGTTTCGGGGAGGTGAGTGGAATTCCCAGTGTAGAGGTGAAATTCGTAGAGATTGGGAGGAACACCGGAGGCGAAAGCGGCTCACTAACCGAATACTGACACTCAAGCACGAAAGCGTGGGGAGCAAACAGG
## 587                                                          ACGTAGGGTGCAAGCGTTAATCGGAATTACTGGGCGTAAAGCGTGCGCAGGCGGTTATGTAAGACAGAGGTGAAATCCCCGGGCTCAACCTGGGAACTGCGTTCGAAACTGCAAGGCTAGAGTGTGTCAGAGGGGGGTAGAATTCCACGTGTAGCAGTGAAATGCGTAGAGATGTGGAGGAATACCAATGGCGAAGGCAGCCCCCTGGGATAACACTGACGCTCATGCACGAAAGCGTGGGGAGCAAACAGG
## 588                                                          ACGAAGGGTGCTAGCGTTGTTCGGAATGACTGGGCGTAAAGGGCGCGTAGGCGGCCAGATGCGTCTGATGTGAAATCCCTGGGCTTAACCTAGGAACTGCATTGGATACGGTCTGGCTAGAGTCCGCAAGAGGAAGATGGAATTGTGCGTGTAGAGGTGAAATTCGTAGATATGCACAAGAACACCGGTGGCGAAGGCGATCTTCTGGAGCGGTACTGACGCTAAGGCGCGAAAGCGTGGGGAGCAAACAGG
## 589                                                          ACGGGAGTGGCAAGCGTTATCCGGAATTATTGGGCGTAAAGCGTCCGCAGGCGGCCTCGTAAGTCTGTTGTTAAAGCGTGGAGCTTAACTCCATTTCGGCAATGGAAACTGTGAGGCTTGAGTGTGGTAGGGGCAGAGGGAATTCCCGGTGTAGCGGTGAAATGCGTAGATATCGGGAAGAACACCAGTGGCGAAGGCGCTCTGCTGGGCCATAACTGACGCTCATGGACGAAAGCCAGGGGAGCGAAAGGG
## 590                                                          ACGTAGGGTGCGAGCGTTAATCGGAATTACTGGGCGTAAAGCGTGCGCAGGCGGTTTGATAAGACAGGCGTGAAATCCCCGGGCTTAACCTGGGAACTGCGCTTGTGACTGTCTCACTAGAGTACGGCAGAGGGGGGTGGAATTCCACGTGTAGCAGTGAAATGCGTAGAGATGTGGAGGAACACCGATGGCGAAGGCAGCCCCCTGGGCCGATACTGACGCTCATGCACGAAAGCGTGGGGAGCAAACAGG
## 592                                                         ACGTGAGTGGCAAGCGTTATCCGGAATCATTGGGCGTAAAGCGTCCGTAGGGGGGTGTTCAAGTGAAATGTTAAATTCTTCGACCCAATCGAAGGCCTGCGTTTCATACTGTTCACCTAGAAGATGGGAGAGGTAAATAGAATTTCTGGAGTAGGGGTAATATCCGTAGAGACCAGAAGGAATACCAATAGCGAAGGCAATTTACTGGCCCATTCTTGACCCTCAAGTGACGAAAGCGTGGGGAGCAAACAGG
## 594                                                          ACGTAGGGTGCAAGCGTTGTCCGGATTTATTGGGCGTAAAGAGCTCGTAGGCGGTTAGACACGTCGGATGTGAAAACTGGTGGCTCAACCACCAGCCTGCATTCGATACGGTCTGACTTGAATGTGGTAGGGGTAACTGGAATTCCTGGTGTAGCGGTGAAATGCGCAGATATCAGGAGGAACACCGATGGCGAAGGCAGGTTACTGGGCCAATATTGACGCTGAGGAGCGAAAGCGTGGGGAGCAAACAGG
## 595                                                          ACGTAGGGTGCAAGCGTTAATCGGAATTACTGGGCGTAAAGCGTGCGCAGGCGGTTTTGTAAGACAGATGTGAAATCCTCGGGCTCAACCTGAGAACTGCATTTGTGACTGCAAAGCTAGAGTGCGGCAGAGGGGGATGGAATTCCGCGTGTAGCAGTGAAATGCGTAGATATGCGGAGGAACACCGATGGCGAAGGCAATCCCCTGGGCCTGCACTGACGCTCATGCACGAAAGCGTGGGGAGCAAACAGG
## 596                                                          ACGGAGGGTGCAAGCGTTATCCGGATTTATTGGGTTTAAAGGGTCCGTAGGCGGATGATTAAGTCAGTGGTGAAAGCCCACAGCTCAACTGTGGAACTGCCATTGATACTGGTTATCTTGAGTGCGGTTGAAGTTGGCGGAATATGTCATGTAGCGGTGAAATGCTTAGATATGACATAGAACACCTATTGCGAAGGCAGCCAACTAAGCCGTAACTGACGTTGAGGGACGAAAGCGTGGGGAGCGAACAGG
## 597                                                          ACGTAGGTGGCGAGCGTTGTCCGGATTTATTGGGTTTAAAGGGTGCGTAGGCGGCTCATTAAGTCAGTTGTGAAATACGACAGCTTAACTGTCGAGGTGCAATTGATACTGGTGAGCTTGAGTACTGATGAGGTAGGCGGAATGGACGGTGTAGCGGTGAAATGCTTAGAGATCGTCCAGAACACCGATAGCGAAGGCAGCTTACTAAGGAGTAACTGACGCTGAGGCACGAAAGTGTGGGGATCAAACAGG
## 598                                                          ACAGAGGGTGCGAGCGTTAATCGGAATTACTGGGCGTAAAGGGCGCGTAGGTGGTTTATTAAGATTGTTGTGAAATCCCTGGGCTTAACCTAGGAACTGCAACGATGACTGATAAGCTTGAGTATTGTAGAGGGTAGTGGAATTTCCGGTGTAGCGGTGAAATGCGTAGAGATCGGAAGGAACACCAGTGGCGAAGGCGGCTACCTGGACAAATACTGACACTGAGGCGCGAGAGCGTGGGGAGCAAACAGG
## 599                                                          ACGAACCGTGCGAACGTTGTTCGGAATCATTGGGCTTAAAGGGCGCGTAGGCGGGTTGACAAGTCCGGGGTGAAAGCCTCCAGCTCAACTGGAGAAGAGCCCCGGATACTGTCAGTCTGGAGAGGGATAGGGGCACATGGAACTTCCGGTGGAGCGGTGAAATGCGTAGAGATCGGAAGGAACGCCGGTGGCGAAAGCGATGTGCTGGATCTTTTCTGACGCTGAGGCGCGAAAGCTAGGGGAGCAAACGGG
## 600                                                          ACGGAGGGAGCTAGCGTTGTTCGGAATTACTGGGCGTAAAGCGCACGTAGGCGGCTACTCAAGTCAGAGGTGAAAGCCCGGGGCTCAACCCCGGAACTGCCTTTGAAACTAGGTAGCTAGAATCTTGGAGAGGTCAGTGGAATTCCGAGTGTAGAGGTGAAATTCGTAGATATTCGGAAGAACACCAGTGGCGAAGGCGACTGACTGGACAAGTATTGACGCTGAGGTGCGAAAGCGTGGGGAGCAAACAGG
## 601                                                          ACGTAGGGTGCAAGCGTTAATCGGAATTACTGGGCGTAAAGCGTGCGCAGGCGGTTATGTAAGACAGAGGTGAAATCCCCGGGCTCAACCTGGGAACTGCCTTTGTGACTGCATAGCTAGAGTACGGCAGAGGGGGATGGAATTCCGCGTGTAGCAGTGAAATGCGTAGATATGCGGAGGAACACCGATGGCGAAGGCAATCCCCTGGACCTGTACTGACGCTCATGCACGAAAGCGTGGGGAGCAAACAGG
## 602                                                          ACAGAGACTGCAAGCGTTATTCGGATTCACTGGGCGTAAAGGGTGCGCAGGCGGCCAAGTGTGTGAGGTGTGAAAGCCCGGAGCTTAACTCCGGAATTGCACCTCAAACTACTTGGCTAGAGCATTGGAGAGGGTAGCAGAATTCATGGTGTAGCAGTGAAATGCGTAGATATCATGAGGAATACCAGAGGCGAAGGCGGCTACCTGGACAATTGCTGACGCTCAGGCACGAAAGCGTGGGGAGCAAAAGGG
## 603                                                          ACGAAGGGTGCTAGCGTTGTTCGGAATCATTGGGCGTAAAGCGTATGTAGGCTGTTTGGTAAGTCAGTTGTGAAAGCCCCGAGCTTAACTTGGGAATTGCAGCTGAAACTGCCTTGCTTGAATTTGGTGGGGGATAGTGGAATTCTTAGTGTAGGGGTGAAATCCGTAGATATTAAGAGGAACACCGGTGGCGAAGGCGACTATCTACACCAATATTGACGCTGAGATACGAAAGCGTGGGGAGCAAACAGG
## 604                                                          ACGTAGGGTGCAAGCGTTAATCGGAATTACTGGGCGTAAAGCGTGCGCAGGCGGTTTGATAAGCCAGATGTGAAATCCCCGAGCTCAACTTGGGAACTGCGTTTGGAACTGTCAGACTAGAGTGCGTCAGAGGGGGGTGGAATTCCGCGTGTAGCAGTGAAATGCGTAGAGATGCGGAGGAACACCGATGGCGAAGGCAGCCCCCTGGGATGACACTGACGCTCATGCACGAAAGCGTGGGGAGCAAACAGG
## 605                                                          ACGTAGGGGGCAAGCGTTGTCCGGAATTACTGGGCGTAAAGCGCACGCAGGCGGATTAATAAGTCAGTTGTAAAATGCACGGGCTCAACTCGTGTTTGTCGACTGATACTGTTAGTCTAGAGTATGTGAGAGGGAAGTGGAATTCCCGGTGTAGCGGTGAAATGCGTAGATATCGGGAGGAACACCAGTGGCGAAGGCGGCTTCCTGGCACAAAACTGACGCTCATGTGCGAAAGCTAGGGCAGCGAACGGG
## 606                                                          ACGTAGGGTGCGAGCGTTAATCGGAATTACTGGGCGTAAAGCGTGCGCAGGCGGTTGTGCAAGACAGATGTGAAATCCCCGGGCTCAACCTGGGAACTGCATTTGTGACTGCACGACTAGAGTACGGCAGAGGGGGGTAGAATTCCACGTGTAGCAGTGAAATGCGTAGAGATGTGGAGGAATACCGATGGCGAAGGCAGCCCCCTGGGCCGATACTGACGCTCATGCACGAAAGCGTGGGGAGCAAACAGG
## 607                                                          ACGAAGGGGGCTAGCGTTGTTCGGAATCACTGGGCGTAAAGCGCACGTAGGCGGATCTTTAAGTCAGGGGTGAAATCCCGGGGCTCAACCCCGGAACTGCCCTTGATACTGGAGATCTCGAGTCCGGGAGAGGTGAGTGGAACTGCGAGTGTAGAGGTGAAATTCGTAGATATTCGCAAGAACACCAGTGGCGAAGGCGGCTCACTGGCCCGGAACTGACGCTGAGGTGCGAAAGCGTGGGGAGCAAACAGG
## 608                                                                                                               ACAGACACTGCAAGCATCTTACAACATAACTAGGCAAAGAGCGTTGTTTTGCAAAGTCGAAGCGAAATTATTGCTCAGGAATTTTCAAAGAATCAGTTAAATGAGAGGATATTGAAAGGAACGCCGAAGGTGAAGACAAGAGCATAAATCTTCTAAAGCAACTCAACGAAAGTTCAAGGAGCAAATCGGATTAGAAACC
## 609                                                          ACGTAGGGTGCGAGCGTTAATCGGAATTACTGGGCGTAAAGCGTGCGCAGGCGGCGACACAAGACAGATGTGAAATCCCCGGGCTCAACCTGGGAACTGCGTTTGTGACTGTGTTGCTAGAGTGCGGCAGAGGGGGGTGGAATTCCACGTGTAGCAGTGAAATGCGTAGAGATGTGGAGGAACACCGATGGCGAAGGCAGCCCCCTGGGTCGACACTGACGCTCATGCACGAAAGCGTGGGGAGCAAACAGG
## 610                                                          ACGGAGGGTGCTAGCGTTGTTCGGAATTACTGGGCGTAAAGCGTACGTAGGCGGTTTGATAAGTTGGAAGTGAAATCCCAAGGCTTAACCTTGGAATTGCTTTCAAAACTGTCAGACTAGAGAATGGTAGGGGAAAGTGGAATTCCTAGTGTAGAGGTGAAATTCGTAGATATTAGGAGGAACACCAGAGGCGCAAGCGGCTTTCTGGACCATTTCTGACGCTAAGGTACGAAAGCGTGGGGAGCGAACAGG
## 613                                                          ACGAGTGGCCCAAGCGTTATCCGGAATTATTGGGCGTAAAGGATGTGTAGGTGGTTGTATTAGTCGAATGTCTAACCTCTGAGCTCAACTTGGAAACAGCATTCGAAACGGTACGACTAGAGTATGTTAGGGGTAAGCGGAACTAATAGTGTAGGGGTGAAATCCGTTGATATTATTGGGAACACCAAAAGCGAAGGCAGCTTACTGGAACATTACTGACACTGAAACATGAAAGCGTGGGTAGCGAATGGG
## 614                                                          ACGTAGGATCCAAGCGTTGTCCGGATTTACTGGGTATAAAGGGTGCGTAGGCGGCTTTTTGCGTCAGAGGTGAAATCCACGGGCTTAACCCGTGGGGTGCCTTTGAAACGGGAAGGCTTGAGTATGACAGAGGCGGATGGAATTCCTGGTGTAGCAGTGAAATGCGTAGATATCAGGAGGAACACCGGTGGCGAAGGCGGTCCGCTGGGTCATAACTGACGCTGAGGCACGAAAGCGTGGGGATCAAACAGG
## 615                                                          ACAGAGGTGGCAAGCGTTGTTCGGATTTATTGGGTGTAAAGGGCAGGTAGGCTGCTAAGTAAGTCTGTTGTGAAATCCCGGGGCTCAACCCCGGAACTGCAACGGAAACTACTTGGCTGGAGTATTGTAGGGGTGAGGGGAATTCTCGGTGTAAGGGTGAAATCTGTAGATATCGAGAGGAACACCAATGGCGAAGGCACCTCACTGGACAATTACTGACGCTGAGCTGCGAAAGCTGGGGGAGCAAACAGG
## 616                                                          ACGTAGGGTGCAAGCGTTAATCGGAATTACTGGGCGTAAAGCGTGCGCAGGCGGTTTGATAAGACAGAGGTGAAATCCCCGGGCTCAACCTGGGAACTGCCTTTGTGACTGTCAGGCTAGAGTGTGTCAGAGGGGGGTAGAATTCCACGTGTAGCAGTGAAATGCGTAGAGATGTGGAGGAATACCGATGGCGAAGGCAGCCCCCTGGGATAACACTGACGCTCATGCACGAAAGCGTGGGGAGCAAACAGG
## 617                                                          ACGGAGGGTGCAAGCGTTAATCGGAATAACTGGGCGTAAAGCGCACGCAGGCGGTCGGATAAGTCAGATGTGAAAGCCCCGGGCTCAACCTGGGAACTGCATTTGAAACTGTCTGACTAGAGTCTTGTAGAGGGGGGTAGAATTCCAGGTGTAGCGGTGAAATGCGTAGAGATCTGGAGGAATACCGGTGGCGAAGGCGGCCCCCTGGACAAAGACTGACGCTCAGGTGCGAAAGCGTGGGGAGCAAACAGG
## 618                                                          ACGGAGGGTGCAAGCATTGATCGGAATTACTGGGCGTAAAGGGCGCGTAGGCGGCTAGGAAAGTCAGATGTGAAATTCCGGGGCTCAACCCCGGAGCTGCATTTGAAACTTCCTGACTAGAGGTCAGTTAGGGAAAACGGAATTCCACGTGTAGCGGTGAAATGCGTAGATATGTGGAAGAACACCGGTGGTGAAGACGGTTTTCTGGGCTGATACTGACGCTGAGGCGCGAAAGCAAGGGGAGCAAACAGG
## 620                                                          ACAGAGGTGGCAAGCGTTGTTCGGATTCATTGGGTGTAAAGGGCAGGTAGGCTGTTTTGCAAGTTTGGCGTGAAATCCCAGGGCTCAACCCTGGAACTGCGCTGAAAACTGCATGACTTGAGTACTGTAGAGGTGAGGGGAATTCTCGGTGTAAGGGTGAAATCTGTAGATATCGAGAGGAACACCAGTGGCGAAGGCGCCTCACTGGACAGTTACTGACGCTGAGCTGCGAAAGCAAGGGGAGCAAACAGG
## 621                                                          ACATAGGGTGCAAGCGTTGTCCGGAATTATTGGGCGTAAAGAGCTCGTAGGTGGTTCGTCACGTCGGATGTGAAACTCTGGGGCTTAACCCCAGACCTGCATTCGATACGGGCGAGCTTGAGTATGGTAGGGGAGTCTGGAATTCCTGGTGTAGCGGTGGAATGCGCAGATATCAGGAGGAACACCAATGGCGAAGGCAGGTCTCTGGGCCAATACTGACACTGAGGAGCGAAAGTCTGGGGAGCGAACAGG
## 622                                                          ACGAGTGGTCCAAGCGTTATCCGGAATTATTGGGCGTAAAGAGTGCGTAGGTGGATTTATTAGTCAGTTGTTTAACCCATGGGCTCAACCTGTGGATTGCAATTGAAACGGTAAATCTTGAGTATGTTAGGGGTGTGCGGAACTCATAGTGTAGGGGTGAAATCCGTTGATATTATGGGGAACACCAAAAGCGAAGGCAGCACACTGGAACATTACTGACACTGAGGCACGAAAGCGTGGGTAGCGAATGGG
## 623                                                          ACGGAGGATCCAAGCGTTATCCGGATTTATTGGGTTTAAAGGGTGCGTAGGCGGCGTTGTAAGTCAGTGGTGAAAGTTTTCAGCTTAACTGGAAAATTGCCATTGATACTGCAGTGCTTGAGTATAGATGAGGTGGGCGGAATGTGTCATGTAGCGGTGAAATGCATAGATATGACACAGAACACCGATTGCGAAGGCAGCTCACTAAACTATAACTGACGCTGAGGCACGAAAGCGTGGGGATCAAACAGG
## 624                                                          ACGAAGGATCCAAGCGTTGTCCGGATTTACTGGGTTTAAAGGGTGCGTAGGCGGGGAATTAAGTCAGTGGTGAAAGCTGGTAGCTCAACTATCAAATTGCCATTGAAACTGATTCTCTCGAGTACGGTTGAGGTAGCTGGAATGTAACATGTAGCGGTGAAATGCTTAGATATGTTACAGAACACCGATTGCGAAGGCAGGCTACTAAACCGTAACTGACGCTGAGGCACGAAAGCGTGGGGATCAAACAGG
## 625                                                          ACGTAGGGTGCAAGCGTTAATCGGAATTACTGGGCGTAAAGCGTGCGCAGGCGGTTATGTAAGACAGTTGTGAAATCCCCGGGCTCAACCTGGGAACTGCATCTGTGACTGCATAGCTAGAGTACGGCAGAGGGGGATGGAATTCCGCGTGTAGCAGTGAAATGCGTAGATATGCGGAGGAACACCGATGGCGAAGGCAATCCCCTGGGCCTGTACTGACGCTCATGCACGAAAGCGTGGGGAGCAAACAGG
## 626                                                          ACGTAGGGTGCGAGCGTTAATCGGAATTATTGGGCGTAAAGGGTGAGTAGGCGGTTTAGTAAGTTAATAGTGAAAGCATTGGGCTTAACCTGATAAGTGCTATTGATACTGCTAGACTAGAGTTAGTTAGAGGATTGTGGAATTCCAAGTGTAGTAGTGAAATGCGTAGATATTTGGAGGAACCTTAATGGCGAAGGCAGCAATCTGGGATGAGACTGACGCTGAGTCACGAAAGCGTGGGGAGCAAACAGG
## 627                                                          ACGAAGGTGGCGAGCGTTACTCGGAATTACTAGGCGTAAAGCGTGGGTAGGTGGTTCGGCAAGTCTGTTGTGGAAGCTCCTGGCTCAACCGGGAGAGGCCAATGGATACTACCGAGCTTGAATGTGGGAGAGGTCACTGGAATTCCTGGTGTAGCGGTGAAATGCGTAGAGATCAGGAGGAACACCGATGGCGAAAGCAGGTGACTGGACCACTATTGACACTGAGCCACGAAAGCTAGGGGAGCAAACAGG
## 628                                                          ACAGGGGATGCAAGTGTTATCCGGAATTATTGGGCGTAAAGCGTCTGCAGGTTGATATTTAAGTCTTTTGTTAAATTTCTAGGCTTAACCTGGAAGTCGCAAAAGAAACTATTTATCTTGAGTATGGTAGAGGTAAAGGGAATTTCAAGTGGAGCGGTGAAATGCGTAGATATTTGAAAGAACACCAACAGCGAAGGCACTTTACTGGGCCATTACTGACACTCAGAGACGAAAGCTAGGGGAGCAAACAGG
## 629                                                          ACGTAGGTGGCGAGCGTTGTCCGGATTTACTGGGCGTAAAGGGAGCGTAGGCGGACTTTTAAGTGAGATGTGAAATACCCGGGCTCAACTTGGGTGCTGCATTTCAAACTGGAAGTCTAGAGTGCAGGAGAGGAGAATGGAATTCCTAGTGTAGCGGTGAAATGCGTAGAGATTAGGAAGAACACCAGTGGCGAAGGCGATTCTCTGGACTGTAACTGACGCTGAGGCTCGAAAGCGTGGGGAGCAAACAGG
## 630                                                          ACGAAGGTGGCAAGCGTTACTCGGAATTACTAGGCGTAAAGGGCAGGTAGGTGGTTTGATAAGTCTGTTGTGAAAGCTCCTGGCTTAACTGGGAGAGGTCAACAGAAACTATCAGGCTTGAGTATGGGAGAGGGTACTGGAATTCCCGGTGTAGCGGTGAAATGCGCAGAGATCGGGAGGAACACCTATGGCGAAAGCAGGTACCTGGACCATTACTGACACTCAGCTGCGAAAGCTAGGGGAGCAAACAGG
## 631                                                         ACGAGTGCCTCGAGCGTTATCCGGAATTATTGGGCGTAAAGGGTGTGTAGGCGGTAGTGTTAGTCTTGTGTTAAATCTATTGGCTCAACCGATAGGCTGCATGGGAAACGGCACAACTAAGAGGATGGAAGGGGTCTGCGGAACTCATGGTGTAGCGGTGAAATGCGTTGATATCATGGGGAACACCGAAAGCGAAGGCAGCAGACTGGTCCACTCCTGACGCTGAAACACGAAAGCGTGGGTCGCGAATGGG
## 632                                                          ACGGAGGATCCGAGCGTTATCCGGATTTATTGGGTTTAAAGGGAGCGTAGGTGGATTGTTAAGTCAGTTGTGAAAGTTTGCGGCTCAACCGTAAAATTGCAGTTGAAACTGGCAGTCTTGAGTACAGTAGAGGTGGGCGGAATTCGTGGTGTAGCGGTGAAATGCTTAGATATCACGAAGAACTCCGATTGCGAAGGCAGCTCACTAGACTGTCACTGACACTGATGCTCGAAAGTGTGGGTATCAAACAGG
## 633                                                          ACAGAGACCTCGATCGTTGTCCGGATTCATTGGGCGTAAAGGGTCCGTAGGCGGTGGTGCAAGTCATTTGTCAAATCTCAGGGCTTAACTCTGAGGCTGCGAGTGAAACTGCATTACTAGAGGCCGGGAGAGGCAAGCGGAATTACCGGTGTAGTCGTAATAAGCGTTGATATCGGTAAGAACACCAAAGGCGAAGGCAGCTTGCTGGAACGGTTCTGACGCTCAGGGACGAAAGCGTGGGTAGCGAATGGG
## 634                                                          ACGAAGGTGGCGAGCGTTACTCGGAATTACTAGGCGTAAAGCGTGGGCAGGCGGCTCGATAAGTCTTTTGTGAAAGCCCCGGGCTTAACCCGGGGAGGTCAAAGGATACTGTCGGGCTTGGATGTGGGAGAGGTCACTGGAATTCCTGGTGTAGCGGTGAAATGCGTAGATATCAGGAGGAACACCAATGGCGAAGGCAGGTGACTGGACCACTATCGACGCTCATCCACGAAAGCTGGGGGATCAAACAGG
## 635                                                          ACGGAGGGTGCAAGCGTTGTTCGGAATCACTGGGCGTAAAGGGCGCGTAGGCGGTTTGATAAGTCAGATGTGAAAGCCCACGGCTTAACCGTGGAAGTGCATTTGAAACTGTCAGACTTGAGTACCAGAGGGGAAAGTGGAATTCCCGGTGTAGAGGTGAAATTCGTAGATATCGGGAGGAATACCGGTGGCGAAGGCGACTTTCTGGCTGGATACTGACGCTGAGGCGCGAAAGCGTGGGGAGCAAACAGG
## 636                                                          ACGGAGGGTGCGAGCGTTAATCGGAATTACTGGGCGTAAAGCGCGCGTAGGCGGTTTGTTAAGTCAGCTGTGAAAGCCCCGGGCTCAACCTGGGAACTGCAGTTGATACTGGCCGACTAGAGTACGAGAGAGGGAGGTAGAATTCCATGTGTAGCGGTGAAATGCGTAGATATATGGAGGAATACCGGTGGCGAAGGCGGCCTCCTGGCTCGATACTGACGCTGAGGTGCGAAAGCGTGGGGATCAAACAGG
## 637                                                          ACGTAGGGTGCAAGCGTTAATCGGAATTACTGGGCGTAAAGCGTGCGCAGGCGGACTTTTAAGCCAGATGTGAAAGCCCCGAGCTTAACTTGGGAATTGCGTTTGGAACTGAGCATCTAGAGTCTGTCAGAGGGGGATGGAATTCCACGTGTAGCAGTGAAATGCGTAGAGATGTGGAGGAACACCGATGGCGAAGGCAGTCCCCTGGGATAAGACTGACGCTCATGCACGAAAGCGTGGGGAGCAAACAGG
## 639                                                          ACGTAGGGTGCGAGCGTTAATCGGAATTACTGGGCGTAAAGCGTGCGCAGGCGGTTATGTAAGACAGATGTGAAATCCCCGGGCTCAACCTGGGAACTGCGTTTGTGACTGCATAACTAGAGTACGGCAGAGGGAGGTGGAATTCCGCGTGTAGCAGTGAAATGCGTAGATATGCGGAGGAACACCGATGGCGAAGGCAATCCCCTGGGCCTGCACTGACGCTCATGCACGAAAGCGTGGGGAGCAAACAGG
## 640                                                          ACGAAGGATGCAAGCGTTATCCGGATTCATTGGGTTTAAAGGGTGCGTAGGCGGACTTGTAAGTCAGTGGTGAAATCTCTTCGCTTAACGAAGAAACTGCCATTGATACTGCAGGTCTAGAGTATAGATGACGTTGGCGGAATATGACATGTAGTGGTGAAATACTTAGATATGTCATAGAACACCGATTGCGAAGGCAGCTAACGAAACTATAACTGACGCTGAGGCACGAAAGTGCGGGGATCAAACAGG
## 641                                                          ACAGAGACCTCAAGCGTTATCCGGATTTATTGGGCGTAAAGGGTCCGCAGGTGGCTTTGTGCGTCGATGGTTAAATTTTCGGGCTCAACTCGAAAACTGCTATCGATACGGCATGGCTAGAGGCCGGAAGAGGTAAGCGGAATTGCCGGTGTAGTAGTAATATGCGTTAATATCGGCAAGAACACCAAATGCGAAGGCAGCTTACTGGGACGCGCCTGACACTCAGGGACGAAAGCGTGGGGAGCGAAAGGG
## 642                                                          ACGGAAGGTGCGAGCGTTAATCGGAATTACTGGGCGTAAAGCGCACGTAGGTGGCTTGGTAAGTCGGATGTGAAAGCCCCGGGCTCAACCTGGGAACGGCATTCGATACTGCTGAGCTAGAGTACGAAAGAGGGGGGTGGAATTCCAGGTGTAGCGGTGAAATGCGTAGAGATCTGGAGGAACACCGGTGGCGAAGGCGGCCCCCTGGTTCGATACTGACGCTGAGGTGCGAAAGCGTGGGGAGCAAACAGG
## 643                                                          ACGAGGGGAGCTAGCGTTGTTCGGAATTACTGGGCGTAAAGGGCGTGTAGGCGGTACCATAAGTCAGATGTGAAAGCCCTGGGCTTAACCTAGGATGTGCATTTGATACTGTGGAACTAGAGATCGAGAGAGGAAAGTGGAATTACGAGTGTAGAGGTGAAATTCGTAGATATTCGTAAGAACACCAGTGGCGAAGGCGACTTTCTGGCTCGATACTGACGCTAAGGCGCGAAAGCGTGGGGAGCAAACAGG
## 644                                                          ACGTAGGGTGCAAGCGTTGTCCGGAATTATTGGGCGTAAAGAGCTCGTAGGCGGTTTGTCACGTCTGCTGTGAAAATCCAAGGCTCAACCTTGGACTTGCAGTGGGTACGGGCAGGCTAGAGTGCGGTAGGGGAGATGGGAATTCCTGGTGTAGCGGTGGAATGCGCAGATATCAGGAGGAACACCAATGGCGAAGGCACATCTCTGGGCCGTAACTGACGCTGAGGAGCGAAAGCGTGGGGAGCGAACAGG
## 646                                                          ACGGAGGGTGCAAGCGTTATCCGGAATCATTGGGTTTAAAGGGTCCGCAGGCGGGAATGTAAGTCAGAGGTGAAATCCTACAGCTCAACTGTAGAACTGCCTTTGATACTGCGTTTCTTGAATTCGATCGAAGTGGGCGGAATATGACATGTAGCGGTGAAATGCATAGATATGTCATAGAACACCGATAGCGAAGGCAGCTCACTAGGTCTGAATTGACGCTCATGGACGAAAGCGTGGGGAGCAAACAGG
## 647                                                          ACGAAGGGGGCTAGCGTTGTTCGGAATTACTGGGCGTAAAGCGCACGTAGGCGGGTTTGTAAGTAGGGGGTGAAATCCCAGGGCTCAACCCTGGAACTGCCTTCTAAACTGCAAGCCTGGAGGTCAGGAGAGGCGAGTGGAATACCGAGTGTAGAGGTGAAATTCGTAGATATTCGGTGGAACACCAGTGGCGAAGGCGACTCGCTGGACTGATACTGACGCTGAGGTGCGAAAGCGTGGGGAGCAAACAGG
## 648                                                          ACGTAGGGTGCAAGCGTTGTCCGGAATTACTGGGTGTAAAGGGAGCGCAGGCGGACCGGCAAGTTGGAAGTGAAAACTATGGGCTCAACCCATAAATTGCTTTCAAAACTGCTGGCCTTGAGTAGTGCAGAGGTAGGTGGAATTCCCGGTGTAGCGGTGGAATGCGTAGATATCGGGAGGAACACCAGTGGCGAAGGCGACCTACTGGGCACCAACTGACGCTGAGGCTCGAAAGCATGGGTAGCAAACAGG
## 649                                                          ACGTAGGGTGCGAGCGTTAATCGGAATTACTGGGCGTAAAGCGTGCGCAGGCGGTTGTGTAAGACAGGCGTGAAATCCCCGGGCTCAACCTGGGAACTGCGCTTGTGACTGCACAGCTAGAGTACGGCAGAGGGGGGTGGAATTCCACGTGTAGCAGTGAAATGCGTAGAGATGTGGAGGAACACCGATGGCGAAGGCAGCCCCCTGGGCCGATACTGACGCTCATGCACGAAAGCGTGGGTAGCAAACAGG
## 650                                                          ACGGAGGATTCGAGCGTTGTCCGGATTTATTGGGTTTAAAGGGTGCGTAGGCGGATTTTTAAGTCAGTGGTGAAAGCCTGCAGCTCAACTGTAGAATTGCCATTGAAACTGAAAATCTTGAATTTGGTTAAAGTAGGCGGAATGTATCATGTAGCGGTGAAATGCTTAGATATGATACAGAACACCGATAGCGAAGGCAGCTTGCTGAACCAATATTGACGCTGAGGCACGAAAGCGTGGGGAGCAAACAGG
## 651                                                          ACGAAGGGGGCTAGCGTTGTTCGGAATCACTGGGCGTAAAGCGCACGTAGGCGGATCTTTAAGTCAGAGGTGAAAGCCTGGAGCTCAACTCCAGAACTGCCTTTGATACTGGGGATCTCGAGTCCGGGAGAGGTGAGTGGAACTGCGAGTGTAGAGGTGAAATTCGTAGATATTCGCAAGAACACCAGTGGCGAAGGCGGCTCACTGGCCCGGTACTGACGCTGAGGTGCGAAAGCGTGGGGAGCAAACAGG
## 652                                                          ACGTAGGGTGCGAGCGTTAATCGGAATTACTGGGCGTAAAGCGTGCGCAGGCGGTTTCGTAAGACAGACGTGAAATCCCCGGGCTCAACCTGGGAACTGCGTTTGTGACTGCGAGGCTAGAGTATGGCAGAGGGGGGTGGAATTCCACGTGTAGCAGTGAAATGCGTAGATATGCGGAGGAACACCGATGGCGAAGGCAATCCCCTGGGCCTGTACTGACGCTCATGCACGAAAGCGTGGGGAGCAAACAGG
## 653                                                          ACGTAGGGTGCGAGCGTTAATCGGAATTACTGGGCGTAAAGCGTGCGCAGGCGGTTTTGTAAGACAGATGTGAAATCCCCGGGCTCAACCTGGGACCTGCATTTGTGACTGCAAGGCTAGAGTACGGTAGAGGGGGATGGAATTCCGCGTGTAGCAGTGAAATGCGTAGATATGCGGAGGAACACCGATGGCGAAGGCAATCCCCTGGACCTGTACTGACGCTCATGCACGAAAGCGTGGGGAGCAAACAGG
## 655                                                          ACGGAGGGTGCAAACGTTAATCGGAATTACTGGGCGTAAAGCGCGCGTAGGCGGTCCGGCAAGTGGGATGTGAAAGCCCTGGGCTTAACCTGGGAACTGCATTCCAAACTACCGGGCTAGAGTATGGTAGAGGGAGGTAGAATTTCCTGTGTAGCGGTGAAATGCGTAGATATAGGAAGGAATACCAGTGGCGAAGGCGGCCTCCTGGACCAATACTGACGCTGAGGTGCGAAAGCGTGGGGAGCAAACAGG
## 656                                                          ACGGAGGATCCGAGCGTTATCCGGATTTATTGGGTTTAAAGGGAGCGTAGGTGGATTGTTAAGTCAGTTGTGAAAGTTTGCGGCTCAACCGTAAAATTGCAGTTGAAACTGGCAGTCTTGAGTACAGTAGAGGTGGGCGGAATTCGTGGTGTAGCGGTGAAATGCTTAGATATCACGAAGAACTCCGATTGCGAAGGCAGCTCACTAGACTGCAACTGACACTGATGCTCGAAAGTGTGGGTATCAAACAGG
## 658                                                          ACAGAGACTGCAAGCGTTATTCGGATTCACTGGGCGTAAAGGGTGCGCAGGCGGCCGTGTGTGTGAGATGTGAAATCCCGGAGCTTAACTCCGGAACTGCGTCTCAAACTACACGGCTAGAGCATTGGAGAGGGAAGTGGAATTCACGGTGTAGCAGTGAAATGCGTAGATATCGTGAGGAACACCAGAGGCGAAGGCGACTTCCTGGACAATTGCTGACGCTCAGGCACGAAAGCGTGGGGAGCAAAAGGG
## 659                                                          ACGTAGGGTGCGAGCGTTAATCGGAATTACTGGGCGTAAAGCGTGCGCAGGCGGTGATGTAAGACAGATGTGAAATCCCCGGGCTCAACCTGGGAACTGCGTTTGTGACTGCATCACTCGAGTACGGCAGAGGGAGGTGGAATTCCGCGTGTAGCAGTGAAATGCGTAGATATGCGGAGGAACACCGATGGCGAAGGCAGCCTCCTGGGCCAGTACTGACGCTCATGCACGAAAGCGTGGGGAGCAAACAGG
## 662                                                          ACGAAGGCCCCAAGCGTTATCCGGAATTACTGGGCGTAAAGCGTCTGTAGGTGGTTTTTTAAGTCTTTCCGTGAAACTTCGGGGCTCAACCCTGAATTGCGGGAGAAACTGGAAAACTTGAGTGTGGGAGAGGCTAGCAGAACGGTAAGAGTAGGGGTGCAATCCGTTGATACTTACCAGAATACCAAAAGCGAAGGCAGCTAGCTGGAACACTACTGACACTGAGAGACGAAAGCGTGGGGAGCGAAAGGG
## 663                                                          ACGGAGGGTGCAAGCGTTGTCCGGAATCATTGGGCGTAAAGCGTTCGTAGGCGGCATGTCAAGTCTGGTGTTAAAGCCCGGGGCTCAACTCCGGTTCGGCACTGGATACTGGCAAGCTAGAATGTGGTAGAGGTAAAGGGAATTCCTGGTGTAGCGGTGAAATGCGTAGATATCAGGAGGAACACCGGTGGCGTAAGCGCTTTACTGGGCCATAATTGACGCTGAGGAACGAAAGCCGGGGGAGCAAATGGG
## 664                                                          ACGGAGGGTGCAAGCGTTAATCGGAATGACTGGGCGTAAAGCGCACGCAGGCGGGATGTTAAGTCAGATGTGAAATCCCCGGGCTTAACCTGGGAACTGCATTTGAAACTGGCATTCTTGAGTTTCGTAGAGGGGGGTAGAATTCCAGGTGTAGCGGTGAAATGCGTAGATATCTGGAGGAATACCGGTGGCGAAGGCGGCCCCCTGGACGGAGACTGACGCTCAGGTGCGAAAGCGTGGGGATCAAACAGG
## 665                                                          ACATAGGGTGCAAGCGTTGTCCGGAATTATTGGGCGTAAAGAGCTCGTAGGTGGTTAGTTACGTCGGATGTGAAAATCTAGGGCTCAACCCTAGACCTGCATCCGATACGGGCTAGCTTGAGTTTGGTAGGGGAGACTGGAATTCCTGGTGTAGCGGTGGAATGCGCAGATATCAGGAGGAACACCAATGGCGAAGGCAGGTCTCTGGGCCAATACTGACACTGAGGAGCGAAAGTGTGGGGAGCGAACAGG
## 666                                                          ACGGAGGGTGCAAGCGTTGTTCGGATTTATTGGGCGTAAAGCGCGCGCAGGCGGATTTGTAAGTCAGGTGTGAAATCTCGAGGCTCAACCTCGAAACTGCGCCTGAAACTGTGAATCTAGAATATGGTAGGGGGAAGGGGAATTTCACGTGTAGGGGTAAAATCCGTAGAGATGTGAAGGAACACCGGAGGCGAAAGCGCCTTCCTGGGCCATTATTGACGCTGAGGCGCGAAAGCGTGGGGAGCAAACAGG
## 667                                                          ACGAAGGGTGCAAGCGTTGTTCGGAATCATTGGGCGTAAAGCGCGCGCAGGCGGATTGATAAGTCAGATGTGAAATCTCGGGGCTCAACCTCGAAACTGCGTCTGAAACTGTTAGTCTAGAATGTCGGAGGGGGCAGGGGAATTTCACGTGTAGGGGTAAAATCCGTAGAGATGTGAAGGAACACCGGGGGCGAAGGCGCCTGCCTGGACGACTATTGACGCTGAGGCGCGAAAGCGTGGGGAGCAAACAGG
## 668                                                          ACGTAGGGTGCAAGCGTTGTCCGGAATTATTGGGCGTAAAGAGCTCGTAGGCGGTTTGTCACGTCTGCTGTGAAAATCCAAGGCTCAACCTTGGACTTGCAGTGGGTACGGGCAGGCTAGAGTGCGGTAGGGGAGATGGGAATTCCTGGTGTAGCGGTGGAATGCGCAGATATCAGGAGGAACACCAATGGCGAAGGCACATCTCTGGGCCGTAACTGACGCTGAGGAGCGAAAGCGTGGGGAGCAAACAGG
## 669                                                          ACGAGTGCCCCGAGCGTTATCCGGAATTATTGGGCGTAAAGGGTGTGTAGGTGGTCAGGTTAGTCTCACGTCAAAGCCTTCGGCTCAACCGAAGAACTGCGTGGGAAACGGCTTGACTTGAGGGTGCGAGGGGTGAATGGAACTCATGGTGTAGGGGTGAAATCCGTTGATATCATGGGGAACACCAAAAGCGAAGGCAATTCACTGGCGCACATCTGACACTGAAACACGAAAGCGTGGGTAGCGAATGGG
## 670                                                          ACGTAGGGTGCGAGCGTTAATCGGAATTACTGGGCGTAAAGCGTGCGCAGGCGGTTACATAAGCCAGATGTGAAATCCCCGGGCTTAACCTGGGAACTGCATTTGGGACTGTGTGGCTTGAGTATGGCAGAGGGAGGTGGAATTCCACGTGTAGCAGTGAAATGCGTAGAGATGTGGAGGAACACCGATGGCGAAGGCAGCCTCCTGGGCCAATACTGACGCTCATGCACGAAAGCGTGGGGAGCAAACAGG
## 671                                                          ACGAGGGGTCCAAGCGTTGTTCGGAATCATTGGGCGTAAAGCGGGTGTAGGTGGCTCTATAAGTCAGGAGTGAAATCCCCGAGCTTAACTTGGGAAGTGCTTTTGATACTGCAGAGCTTGAATGTGGGAGAGGATCGTAGAATTCCAGGTGTAGTGGTGAAATACGTAGATATCTGGAGGAATACCGGTGGCGAAGGCGGCGATCTGGCCCAACATTGACACTGAGACCCGAAAGCGTGGGGATCAAACAGG
## 672                                                          ACGGAGGGTGCAAGCGTTAATCGGAATTACTGGGCGTAAAGCGCGCGTAGGCGGTTGACTAAGTCTGCTGTGAAAGCCCCGGGCTTAACCTGGGAACTGCAGTGGATACTGGTTGGCTAGAGTATGGTAGAGGGTAGTGGAATTTCCGGTGTAGCAGTGAAATGCGTAGAGATCGGAAGGAACACCAGTGGCGAAGGCGGCTATCTGGACCAATACTGACGCTGAGGTGCGAAAGCGTGGGGAGCAAACAGG
## 674                                                          ACGGAGGGTGCAAGCGTTGCTCGGATTTACTGGGCGTAAAGCGCACGCAGGCGGTCTGGAAGGTCGGGTGTGAAAGCCCGGGGCTCAACCCCGGAACTGCACTCGAAACCGCCAGACTTGAGGTCGAGAGAGGAGGGTAGAATTCCTGGTGTAGAGGTGAAATTCGTTGATATCAGGAGGAATACCGGTGGCGAAGGCGGCCCTCTGGCTCGATTCTGACGCTCATGTGCGAAAGCGTGGGGAGCAAACAGG
## 675                                                          ACGTAGGGTGCAAGCGTTAATCGGAATTACTGGGCGTAAAGCGTGCGCAGGCGGATGTTTAAGCCAGATGTGAAAGCCCCGAGCTTAACTTGGGAATTGCGTTTGGAACTGAGCATCTAGAGTCTGTCAGAGGGGGATGGAATTCCACGTGTAGCAGTGAAATGCGTAGAGATGTGGAGGAACACCGATGGCGAAGGCAGCCCCCTGGGCCAATACTGACGCTCATGCACGAAAGCGTGGGGAGCAAACAGG
## 676                                                          ACGGAGGGTGCGAGCGTTATCCGGATTCACTGGGTTTAAAGGGTGCGTAGGTGGGTTGGTAAGTCAGTGGTGAAATCCTCGAGCTTAACTCGAGAACTGCCGTTGATACTACCAGTCTTGAATTCTGTGGAGGTAAGCGGAATATGTCATGTAGCGGTGAAATGCTTAGATATGACATAGAACACCGATAGCGAAGGCAGCTTACTACGCAGATATTGACACTGAGGCACGAAAGCGTGGGGATCAAACAGG
## 677                                                          ACGGAGGGTGCAAGCGTTATCCGGAATCACTGGGTTTAAAGGGTACGTAGGCGGTTTAATAAGTCAGGTGTGAAATACTGCAGCTTAACTGCGGAACTGCACATGATACTGTTAGACTTGAGTAGGGAGGAGGGACACGGAATGCATCATGTAGCGGTGAAATGCTTAGATATGATGTAGAACACCGATAGCGAAGGCAGTGTTCCATACCTATACTGACGCTGAGGTACGAAAGCGTGGGGAGCGAACAGG
## 678                                                          ACGGAGGGTGCAAGCGTTATCCGGATTCACTGGGTTTAAAGGGTGCGTAGGTGGGCATGTAAGTCAGAGGTGAAATCCCCGAGCTTAACTTGGGAACTGCCTTTGATACTATATGTCTTGAATATCGTGGAGGTAAGCGGAATATGTCATGTAGCGGTGAAATGCATAGAGATGACATAGAACACCAATTGCGAAGGCAGCTTGCTACACGATGATTGACACTGAGGCACGAAAGCGTGGGGAGCAAACAGG
## 679                                                          ACGTAGGTTGCGAGCGTTATCCGGAATTATTGGGCGTAAAGAGTTCGTAGGTGGTATGTTAAGTTGGATGTTAAACCTTTGAGCTCAACTCAAAGAATGTGTCCAAAACTGGCAAACTAGAGACAGGTAGAGGTAAGCGGAATTCTGTATGTAGGGGTAATATCCGTAGATATACAGAGGAACACCAAAAGCGAAGGCAGCTTACTGGACCTGTTCTGACACTCAAGAACGAAAGCGTGGGGAGCAAACGGG
## 680                                                          ACGAAGGGTGCAAGCGTTATTCGGATTGATTGGGCGTAAAGGGTGTGTAGGTGGTGTGCTAAGTCAACTGTTAAATTCCCTGGCTTAACCAGGGGTCAGCAGTAGATACTGGCGCGCTAGAGGATGGAAGAGAGAAGTGGAATTCTCGGAGTAGCGGTAAAATGCGTAGATCTCGAGAAGAACACCGATGGCGAAGGCAGCTTCTTGGTCCATTCCTGACACTGAGGCACGAAAGCGTGGGGAGCAAACAGG
## 681                                                          ACAGGGGGTGCAAGCGTTGCTCGGAATTACTGGGCGTAAAGCGTGCGCAGGCGGTTCAGTAAGTCGGAGGTGAAAGCCCGAGGCTCAACCTCGGAAATGCCTTCGAAACTACTGATCTTGAGTACGGGAGAGGGTCGCGGAATTCTTGGTGTAGAGGTGAAATTCGTAGATATCAGGAAGAACACCAGTGGCGAAAGCGGCGACCTGGTCCGAGACTGACGCTCATGCACGAAAGCGTGGGGAGCAAACAGG
## 682                                                          ACAGAGGGTGCAAGCGTTGTTCGGAATTACTGGGCGTAAAGCGTGCGTAGTCGGTATTGAGAGTCGCGGGTGAAATCCCAGGGCTTAACCCTGGAACTGCCTGCGAGACCTCAGTACTAGAGTGTGAGAGGGGATAGTGGAATACCCAGTGTAGCGGTGAAATGCGTAGAGATTGGGTGGAACACCGGTGGCGAAGGCGGCTATCTGGCTCACAACTGACGATCAGGCACGAAAGCGTGGGGAGCAAACAGG
## 684                                                          ACGGAGGGTGCAAGCGTTAATCGGAATTACTGGGCGTAAAGCGTGCGTAGGCGGCTCGCTAAGTCAGATGTGAAAGCCCCGGGCTCAACCTGGGAACGGCATTTGAAACTGGCAAGCTAGAGTTTAGGAGAGGAGAGTGGAATTTCAGGTGTAGCGGTGAAATGCGTAGAGATCTGAAGGAACACCAGTGGCGAAGGCGACTCTCTGGCCTAAAACTGACGCTGAGGTACGAAAGCGTGGGTAGCAAACAGG
## 685                                                          ACGGAGGGTGCAAGCGTTATCCGGAATCATTGGGTTTAAAGGGTCCGCAGGCGGGCATATAAGTCAGTGGTGAAAGCCTACAGCTTAACTGTAGAACTGCCATTGATACTGTATGCCTTGAATTCGATCGAAGTGGGCGGAATGTGTAGTGTAGCGGTGAAATGCATAGATATTACACAGAACACCGATAGCGAAGGCAGCTCACTAGGTCTGAATTGACGCTCATGGACGAAAGCGTGGGGAGCAAACAGG
## 686                                                          ACGTAGGGAACAAGCGTTGTCCGGATTTACTGGGCGTAAAGGGCACGCAGGCGGTTTTTTAAGTCAGATGTGAAAGGTACCGGCTCAACCGGTGACATGCATTTGAAACTGAAAGACTTGAGTATTGGAGAGGCAAGTGGAATTCCTAGTGTAGCGGTGAAATGCGTAGAGATTAGGAGGAACACCAGTGGCGAAGGCGGCTTGCTGGACAAATACTGACGCTGAGGTGCGAAAGCGTGGGGAGCAAACAGG
## 687                                                          ACGAGGGATCCTAGCGTTGTTCGGAATCATTGGGCGTAAAGCGGGTGTAGGTGGCTCTATAAGTCAGATGTGAAAGCCCTGGGCTTAACCCAGGAAGTGCATTTGATACTGCTGAGCTTGAGTGTGGGAGAGGCTAGTGGAATTCCTGGTGTAGTGGTGAAATACGTAGATATCAGGAGGAACATCGGTGGCGAAGGCGGCTAGCTGGCCCAACACTGACACTGAGACCCGAAAGCGTGGGGATCAAACAGG
## 688                                                         ACAGAGACCACAAGCGTTATCCGGAATTATTGGGCGTAAAGGGTGAGTAGGTGGTATTGTTAGTCGGGTTTGAAAGATCGTGGGCTTAACCCATGATAGGGACCCGAAACGGCAAAACTAGAATATGTAAGAGGTTAGTGGAACTATATGTGTAGGGGTAAAATCCGTTGATATATATGGGAACACCTAAAGCGAAGGCAGCTAACTGGTACATTATTGACACTGAATCACGAAAGCGTGGGTAGCGATACGG
## 689                                                          ACGGAGGATCCAAGCGTTATCCGGAATCATTGGGTTTAAAGGGTCCGTAGGCGGTTTTATAAGTCAGTGGTGAAATCTGGTCGCTCAACGATCAAACGGCCATTGATACTGTAGAACTTGAATTACTTGGAAGTAACTAGAATATGTAGTGTAGCGGTGAAATGCTTAGAGATTACATGGAATACCAATTGCGAAGGCAGGTTACTACAAGTGGATTGACGCTGATGGACGAAAGCGTGGGGAGCGAACAGG
## 690                                                          ACGTAGGGGGCAAGCGTTGTCCGGAATTACTGGGCGTAAAGCGCACGCAGGCGGATTAAAAAGTCAGCCGTAAAATGCACGGGCTCAACCTGTGTTAGTCAGCTGATACTATTAGTCTAGAGTATGTGAGAGGGAAGTGGAATTCCCGGTGTAGCGGTGAAATGCGTAGATATCGGGAGGAACACCAGTGGCGAAGGCGGCTTCCTGGCACAAAACTGACGCTCATGTGCGAAAGCTAGGGCAGCGAACGGG
## 691                                                          ACGTAGGGTGCGAGCGTTAATCGGAATTACTGGGCGTAAAGCGTGCGCAGGCGGTTTTGTAAGACAGATGTGAAATCCCCGGGCTCAACCTGGGACCTGCATTTGTGACTGCAAAGCTGGAGTGCGGCAGAGGGGGATGGAATTCCGCGTGTAGCAGTGAAATGCGTAGATATGCGGAGGAACACCGATGGCGAAGGCAATCCCCTGGGCCTGCACTGACGCTCATGCACGAAAGCGTGGGGAGCAAACAGG
## 692                                                           ACGTAGGTGGCAAGCGTTGTCCGGAATTATTGGGCGTAAAGCGCGCGCAGGCGGGATATCAAGTCTGTCTTAAAAGTGCGAGGCTCAACCTCGTGAGGGGACAGAAACTGGTATTCTTGAGTGTCGGAGAGGAAAGTGGAATTCCAAGTGTAGCGGTGAAATGCGTAGAGATTTGGAAGAACACCAGTGGCGAAGGCGACTTTCTGGACGATGTCTGACGCTGAGGCGCGAAAGCTAGGGGAGCGAACGGG
## 693                                                          ACGGAGGGTGCGAGCGTTAATCGGAATTACTGGGCGTAAAGCGCGCGTAGGCGGTTCGTTGTGTCCGCTGTGAAAGCCCTGGGCTTAACCTGGGAACTGCAGTGGAAACTGGCGGGCTAGAGTATGGTAGAGGAGAGTGGAATTTCCGGTGTAGCAGTGAAATGCGTAGAGATCGGAAGGAACACCAGTGGCGAAGGCGGCTCTCTGGACCAATACTGACGCTGAGGTGCGAAAGCGTGGGGAGCAAACAGG
## 696                                                          ACAGAGGGTGCAAGCGTTGTTCGGAATTATTGGGCGTAAAGCGAGTGTAGGCGGTTATCTAAGTCTGGTGTGAAAGCCCGGGGCTCAACCTCGGAAGTGCACTGGATACTGGGTAACTAGAGGGCGATAGAGGGGAACGGAATTAATGGTGTAGGGGTGAAATCCGTAGATATCATTAGGAACACCGAAAGCGAAGGCTGTTCCCTGGGTCGCAACTGACGCTAAGACTCGAAAGCATGGGTAGCAAACAGG
## 697                                                          ACGAAGGCCCCAAGCGTTATCCGGAATTACTGGGCGTAAAGCGTCTGTAGGTGGTTTGGAAAGTTTCAAGTGAAACGTCAGGGCTCAACCCTGTACTTGCTTGGAAAACTATCAAACTAGAGTGTGGGAGAGGCAAGCAGAACGGTATGAGTAGGGGTGCAATCCGTTGATACATACCAGAATACCAAAAGCGAAGGCAGCTTGCTGGAACACAACTGACACTGAGAGACGAAAGCGTGGGGAGCAAAAGGG
## 698                                                                                                               ACAGATGGAGCGAGCTTCCTGCTATATGATTAGGTAGATAGCGTGTGAAGATGATTCTAGTTTTATTTTAAAGTGGGGGAATTTTCTATGGATTAGTAAAATGATTGTATATAGGAAGGAACGCCAACAGCGAAGGCACTCATTTAAATATAACTCAAATTCAAACACGGAAGTCCGAGGAGCAAAACGGATTAGATAC
## 699                                                          ACGTAGGGTGCAAGCGTTGTCCGGATTTATTGGGCGTAAAGAGCTCGTAGGCGGTTTGTCGCGTCGGATGTGAAAACCTATGGCTTAACCATGGGCCTGCATTCGATACGGGCAAACTAGAGTTCGGTAGGGGAGACTGGAACTCCTGGTGTAGCGGTGAAATGCGCAGATATCAGGAAGAACACCGGTGGCGAAGGCGGGTCTCTGGGCCGATACTGACGCTGAGGAGCGAAAGCGTGGGGAGCAAACAGG
## 700                                                          ACGGAGGGTGCAAGCGTTATCCGGATTCACTGGGTTTAAAGGGTGTGTAGGCGGGCTATTAAGTCAGTGGTGAAATCTCCGGGCTCAACCCGGAAACTGCCATTGATACTATTAGTCTTGAATGTTGTTGAGGTGGGCGGAATATATCATGTAGCGGTGAAATGCATAGATATGATATAGAACACCGATTGCGAAGGCAGCTCACTAAACAACGATTGACGCTGAGGCACGAAAGCGTGGGGATCAAACAGG
## 701                                                          ACGAAGGACCCGAGCGTTATCCGGATTTATTGGGTTTAAAGGGTGCGTAGGCGGATTAGTAAGTCAGTGGTGAAAGCCTCTCGCTCAACGGGAGAACTGCCATTGATACTGCTAGTCTTGAGTATGGTTGAGGTGGGCGGAATGTGTCATGTAGCGGTGAAATGCTTAGATATGACACAGAACACCGATTGCGAAGGCAGCTCGCTAAGCCATAACTGACGCTGAGGCACGAAAGCGTGGGGAGCAAACAGG
## 702                                                          ACAGAGGGGGCAAGCGTTATTCGGAATTATTGGGCGTAAAGGGCGCGTAGGCGGTGATTTAAGTGAGATGTGTAATCCCCGGGCTTAACCTGGGAACTGCATCTCAGACTGGATCGCTAGAGTGCTGGAGAGGGTGGTAGAATTCCACGTGTAGCGGTGAAATGCGTAGAGATGTGGAGGAATACCAGTGGCGAAGGCGGCCACCTGGACAGTAACTGACGCTGAGGCGCGAAAGTGTGGGTAGCAAACAGG
## 704                                                          ACGGAGGGTGCAAGCGTTATCCGGATTTATTGGGTTTAAAGGGTCCGTAGGCGGACCTATAAGTCAGTGGTGAAAGCCCACCGCTCAACGGTGGAACTGCCATTGATACTGTAGGTCTTGAGTATGTATGAAGTTGGCGGAATGTGTGGTGTAGCGGTGAAATGCTTAGATATCACACAGAACACCGATTGCGAAGGCAGCTGACTAATACATAACTGACGCTGAGGGACGAAAGTGTGGGGATCAAACAGG
## 705                                                          ACAGAGGGTGCGAGCGTTAATCGGAATCACTGGGCTTAAAGGGTCCGTAGGCGGAACTGCAGGCGTCGAGTGAAATACCACGGCTCAACCGTGGAACTGCTCGGCGAACCGCAGTTCTTGAGGCGAGTAGAGGCGATCAGAACGATAGGTGGAGCGGTGAAATGCGTAGATATCTATCGGAATGCCGAAGGTGAAGACAGGTCGCTGGGCTCGTCCTGACGCTGAGGGACGAAAGCGTGGGGAGCAAACGGG
## 706                                                          ACGGAGGGGGCTAGCGTTGTTCGGAATTACTGGGCGTAAAGCGCACGTAGGCGGCTTGGTAAGTTAGAGGTGAAAGCCTGGAGCTCAACTCCAGAATTGCCTTTAAGACTGCTTGGCTTGAACACGGGAGAGGTGAGTGGAATTCCGAGTGTAGAGGTGAAATTCGTAGATATTCGGAAGAACACCAGTGGCGAAGGCGGCTCACTGGACCGTTGTTGACGCTGAGGTGCGAAAGCGTGGGGAGCAAACAGG
## 707                                                          ACGTAGGGTGCGAGCGTTAATCGGAATTACTGGGCGTAAAGAGTGCGTAGGCGGTTTTGTAAGACAGATGTGAAATCCCCGGGCTTAACCTGGGAACTGCATTTGTGACTGCAAGACTAGAGTGTGTCAGAGGGAGGTGGAATTCCGCGTGTAGCAGTGAAATGCGTAGAGATGCGGAAGAACACCGATGGCGAAGGCAGCCTCCTGGGATAACACTGACGCTCAGGCACGAAAGCGTGGGGAGCAAACAGG
## 708                                                          ACGAACTGTGCGAACGTTACTCGGAATCACTGGGCTTACAGGGTGCGTAGGTGGCTTGTTAAGCAGGGTGTGAAATCCCTCGGCTCAACCGAGGAACTGCGCTCTGAACTGGCGAGCTGGAGTATGCTAGAGGTGAATGGAACATCCAGTGGAGCGGTGAAATGTGTTGATATTGGATGGAACACCAGAGGCGAAGGCGATTCACTGGGGCATAACTGACACTGAGGCACGAAAGCCAGGGGAGCAAACGGG
## 709                                                          ACGTAGGGTGCAAGCGTTAATCGGAATTACTGGGCGTAAAGCGTGCGCAGGCGGTTATATAAGTCAGATGTGAAATCCCCGGGCTCAACCTGGGAACTGCATTTGAGACTGTATAGCTAGAGTACGGCAGAGGGGGATGGAATTCCGCGTGTAGCAGTGAAATGCGTAGATATGCGGAGGAACACCGATGGCGAAGGCAATCCCCTGGGCCTGCACTGACGCTCATGCACGAAAGCGTGGGGAGCAAACAGG
## 710                                                          ACAGAGGATGCAAGCGTTATCCGGAATCACTGGGCATAAAGCGTCTGTAGGTTGCTTATCAAGTCTGCTGTTAAAGATCAGGGCCTAACCCTGAAAAAGCAGTGGAAACTAGTAGGCTTGAGTGTGGTAGAGGTAGAGGGAATTCCTGGTGTAGCGGTGAAATGCGTAGATATTAGGAAGAACACCAATGGCGAAAGCACTCTACTGGGCCATAACTGACACTGAGAGACGACAGCTAGGGGAGCAAATGGG
## 711                                                          ACGTAGGTGGCAAGCGTTGTCCGGATTTATTGGGTTTAAAGGGTGCGTAGGTGGTTCCATAAGTCAGTGGTGAAATACGGCAGCTTAACTGTCGAGGTGCCATTGATACTGCGGAACTTGAGTACAGACGAGGTAGGCGGAATTGACGGTGTAGCGGTGAAATGCTTAGATATCGTCAAGAACACCGATAGCGAAGGCAGCTTACTAGACTGTAACTGACACTGAGGCACGAAAGTGTGGGGATCAAACAGG
## 712                                                          ACGTATGGTGCAAGCGTTATCCGGATTTACTGGGTGTAAAGGGAGCGCAGGCGGTGCGGCAAGTCTGATGTGAAAGCCCGGGGCTCAACCCCGGTACTGCATTGGAAACTGTCGTACTAGAGTGTCGGAGGGGTAAGCGGAATTCCTAGTGTAGCGGTGAAATGCGTAGATATTAGGAGGAACACCAGTGGCGAAGGCGGCTTACTGGACGATAACTGACGCTGAGGCTCGAAAGCGTGGGGAGCAAACAGG
## 713                                                          ACAGAGGGTGCAAGCGTTGCTCGGAATTATTGGGCGTAAAGGGCAAGTAGGTGGTCTCGTTTGTCTTGGGTGAAATCCCTGGGCTTAACCCAGGAACTGCCCCAGAAACGGCGAGACTAGAGTTCTGGAGAGGGTCGTGGAATTCCCAGTGTAGCGGTGAAATGCGTAGAGATTGGGAGGAACACCAGAGGCGAAAGCGGCGGCCTGGACAGATACTGACACTCAACTGCGAAAGCGTGGGGAGCAAACAGG
## 714                                                          ACGGAGGATCCGAGCGTTATCCGGATTTATTGGGTTTAAAGGGAGCGTAGATGGGTTGTTAAGTCAGTTGTGAAAGTTTGCGGCTCAACCGTAAAATTGCAATTGATACTGGCAGTCTTGAGTACAGTTGAGGTAGGCGGAATTCGTGGTGTAGCGGTGAAATGCTTAGATATCACGAAGAACTCCGATTGCGAAGGCAGCTTACTAACCTGTAACTGACATTGATGCTCGAAAGTGTGGGTATCAAACAGG
## 715                                                          ACGTAGGGGGCGAGCGTTGTTCGGAATTATTGGGCGTAAAGGGCATGTAGGCGGTTTTGTAAGCCTGGTGTGAAATCCTGCAGCTTAACTGTAGAATTGCATTGGGTACTGCGAGACTTGAATCACAGAGGGGAAACTAGAATTCCAGGTGTAGGGGTGGAATCTGTAGATATCTGGAAGAATACCGGTGGCGAAGGCGGGTTTCTAGCTGATGATTGACGCTGAGGTGCGAAAGTGCGGGGAGCAAACAGG
## 716                                                          ACGTAGGGACCAAGCGTTGTTCGGATTTACTGGGCGTAAAGGGCGCGTAGGCGGTGCGGTAAGTCACTTGTGAAATCTCTGAGCTTAACTCAGAACGGCCAAGTGATACTGCAGTACTAGAGTGCAGAAGGGGCAATCAGAATTCTTGGTGTAGCGGTGAAATGCGTAGATATCAAGAGGAATACCTGAGGTGAAGACGGGTTGCTAGGCTGACACTGACGCTGAGGCGCGAAAGCTAGGGGAGCAAACGGG
## 717                                                          ACAGAGGGTGCAAGCGTTGTTCGGAATTACTGGGCGTAAAGCGTGCGTAGTCGGTATTGAGAGTCATAGGTGAAATCCCAGGGCTTAACCCTGGAACTGCCCGTGAGACCTCAGTACTAGAGTGTGAGAGGGGATAGTGGAATACCAAGTGTAGCGGTGAAATGCGTAGAGATTTGGTAGAACACCGGTGGCGAAGGCGGCTATCTGGCTCACAACTGACGATCAGGCACGAAAGCGTGGGGAGCAAACAGG
## 718                                                          ACGGAGGATCCGAGCGTTATCCGGATTTATTGGGTTTAAAGGGAGCGTAGGCGGATTGTTAAGTCAGTTGTGAAAGTTTGCGGCTCAACCGTAAAATTGCAGTTGATACTGGCAGTCTTGAGTGCAGTAGAGGTGGGCGGAATTCGTGGTGTAGCGGTGAAATGCTTAGATATCACGAAGAACTCCGATTGCGAAGGCAGCTCACTGGAGTGTAACTGACGCTGATGCTCGAAAGTGTGGGTATCAAACAGG
## 719                                                          ACGAAGGGGGCTAGCGTTGTTCGGAATCACTGGGCGTAAAGCGCACGTAGGCGGATCTTTAAGTCAGGGGTGAAATCCCGAGGCTCAACCTCGGAACTGCCTTTGATACTGGAGGTCTCGAGTCCGGGAGAGGTGAGTGGAACTGCGAGTGTAGAGGTGAAATTCGTAGATATTCGCAAGAACACCAGTGGCGAAGGCGGCTCACTGGCCCGGTACTGACGCTGAGGTGCGAAAGCGTGGGGAGCAAACAGG
## 720                                                          ACGGAGGGTGCGAGCGTTAATCGGAATTACTGGGCGTAAAGGGTGCGTAGGTGGTTTATATAGTCATGCGTAGAAGCCCCGGGCTCAACCTGGGAATGTCGTGTGATACTGATAGACTGGAGTACAGGAGAGGGCAGTGGAATTTCCGGTGTAGCGGTGAAATGCGTAGATATCGGAAGGAACACCGGTGGCGAAGGCGGCTGCCTGGCCTGATACTGACACTGAGGCACGAAAGCGTGGGGAGCAAACAGG
## 721                                                          ACGGAGGGTGCGAGCGTTATCCGGAATTACTGGGTTTAAAGGGTGCGTAGGCGGCTTTGTAAGTCAGGAGTGAAAGTTTGCGGCTCAACCGTAAAATTGCTTCTGATACTGCATAGCTAGAATTAGGATGAGGTCAGCGGAATGTGGCATGTAGCGGTGAAATGCATAGATATGCCATAGAACACCAATTGCGAAGGCAGCTGGCTAGACCTGGATTGACGCTGAGGCACGAAAGCGTGGGGAGCGAACAGG
## 722                                                          ACGAAGGGGGCTAGCGTTGTTCGGAATTACTGGGCGTAAAGCGCGCGTAGGCGGTGATCTTTGTCAGAGGTGAAAGCCTGGGGCTCAACCCCAGAATTGCCTTTGAAACGGGATCGCTAGAGTCCGAGAGAGGATGGCGGAATTCCTAGTGTAGAGGTGAAATTCGTAGATATTAGGAAGAACACCGGTGGCGAAGGCGGCCATCTGGCTCGGTACTGACGCTCAGGCGCGAAAGCGTGGGGAGCAAACAGG
## 723                                                          ACAGGGGGTGCAAGCGTTGTTCGGAATCATTGGGCGTAAAGCGCGTGTAGGCGGACCGTTAAGTCTGGTGTGAAAGCCCTCGGCTCAACTGAGGAAGTGCATTGGATACTGGCGGCCTTGAGTCCTGGAGAGGGTAGCGGAATTCCCAGTGTAGAGGTGAAATTCGTAGATATTGGGAGGAACACCGGTGGCGAAGGCGGCTACCTGGACAGTGACTGACGCTGAGACGCGAAAGCATGGGTAGCAAACAGG
## 724                                                          ACGTAGGGTGCGAGCGTTAATCGGAATTACTGGGCGTAAAGCGTGCGCAGGCGGTTATGTAAGACAGTTGTGAAATCCCCGGGCTCAACCTGGGAATTGCATCTGTGACTGCATAGCTAGAGTACGGTAGAGGGGGATGGAATTCCGCGTGTAGCAGTGAAATGCGTAGATATGCGGAGGAACACCGATGGCGAAGGCAATCCCCTGGACCTGTACTGACGCTCATGCACGAAAGCGTGGGGAGCAAACAGG
## 725                                                         ACGAAGGGTGCAAGCGTTGTCCGGATTTATTGGGCGTAAAGCGTTCCGCAGGCGCGTGTGTAAGTCATGCTTCAAAGACCGGGGCTCAACCTCGGAAAGGGGTGTGATACTGCACATGTAGAATAATGTAGGGGCTACTGGAACTTTGCATGTAGGAGTGAAATCCGTTGATATGCAAAGGAACACCGAGGGCGTAGGCAGGTAGCTGGACATTTATTGACGCTCAGGGACGACAGCTAGGGTAGCGAAAGGG
## 726                                                          ACGTAGGGTGCAAGCGTTAATCGGAATTACTGGGCGTAAAGCGTGCGCAGGCGGATGTTTAAGCCAGATGTGAAAGCCCCGAGCTTAACTTGGGAATTGCGTTTGGAACTGAGCATCTAGAGTCTGTCAGAGGGGGATGGAATTCCACGTGTAGCAGTGAAATGCGTAGAGATGTGGAGGAACACCGATGGCGAAGGCAATCCCCTGGGCCTGCACTGACGCTCATGCACGAAAGCGTGGGGAGCAAACAGG
## 727                                                          ACAGAGGGTGCAAGCGTTGTTCGGAATTATTGGGCGTAAAGGGCAGGTAGGTTGTCTTTTAAGTCTGCTGTGAAATCCCTGGGCTCAACCTGGGAAGTGCGGTGGATACTGGAAGACTAGAGTACTATAGGGGTATGCGGAATTCCCGGTGTAGCGGTGAAATGCGTAGATATCGGGAGGAACACCAGAGGCGAAGGCGGCATACTGGAGAGTAACTGACACTAAACTGCGAAAGCGTGGGGAGCAAACAGG
## 728                                                         ACGTAGGCACCAAGCGTTGTCCGGATTTATTGGGCGTAAAGAGCTCGTAGGCGGTTCAGCAAGTCGGGTGTGAAAACTATGGGCTTAACCCATAGCCTGCATCCGAAACTGCTGTGACTAGAGTTCGGTAGGGGAGAGGGGAATTCCTAGTGTAGCGGTGAAATGCGCAGATATTAGGAGGAACACCGATGGCGAAGGCACCTCTCTGGGCCGATACTGACGCTGAGGAGCGAAAGCGTGGGTAGCAAACAGG
## 729                                                          ACGAAGGGGGCTAGCGTTGTTCGGAATTACTGGGCGTAAAGGGCGCGTAGGCGGCCCGATAAGTCAGATGTGAAAGCCCCGAGCTTAACTTGGGAACTGCATTTGATACTGTCGGGCTAGAGAGTGAGAGAGGAAAGTGGAATTGCGAGTGTAGAGGTGAAATTCGTAGATATTCGCAAGAACACCAGTGGCGAAGGCGGCTTTCTGGCTCACAACTGACGCTAAGGCGCGAAAGCGTGGGGAGCAAACAGG
## 731                                                          ACGGAGGGTGCAAGCGTTAATCGGAATAACTGGGCGTAAAGCGCACGCAGGCGGTTGGATAAGCGAGATGTGAAAGCCCCGGGCTCAACCTGGGAACTGCATTTCGAACTGTCCGACTAGAGTCTTGTAGAGGGGGGTAGAATTCCAGGTGTAGCGGTGAAATGCGTAGAGATCTGGAGGAATACCGGTGGCGAAGGCGGCCCCCTGGACAAAGACTGACGCTCAGGTGCGAAAGCGTGGGGAGCAAACAGG
## 732                                                          ACAGAGGTCCCGAGCGTTGTTCGGATTCACTGGGCGTAAAGGGTGCGTAGGTGGTGGGGTAAGTCGGATGTGAAATCTCCAAGCTCAACTTGGAAATGGCATTGGAAACTGCCCTGCTAGAGGATCGGAGGGGGGACTGGAATACTTGGTGTAGCAGTGAAATGCGTAGATATCAAGTGGAACACCAGTGGCGAAGGCGAGTCCCTGGACGATACCTGACACTGAGGCACGAAAGCTAGGGGAGCAAACAGG
## 733                                                          ACGAAGGGGGCGAGCGTTATTCGGAATCACTGGGCGTAAAGCGTGCGTAGGCTGCTTGGAAAGTTGGAAGTGAAAGCCCAGGGCTCAACCCTGGAATTGCTTTCAAAACTACCAGGCTCGGATTCGGGAGAGGATAGCGGAATTGTCAGTGTAGCAGTGAAATGCGTAGATATTGACAGGAACACCAGTGGCGCAAGCGGCTATCTGGACCGACATCGACGCTGAGGCACGAAAGCGTGGGGATCAAACAGG
## 734                                                          ACGTAGGGTGCAAGCGTTAATCGGAATTACTGGGCGTAAAGCGTGCGCAGGCGGTTATATAAGTCAGATGTGAAATCCCCGGGCTCAACCTGGGAACTGCATTTGAGACTGTATAGCTAGAGTACGGTAGAGGGGGATGGAATTCCGCGTGTAGCAGTGAAATGCGTAGATATGCGGAGGAACACCGATGGCGAAGGCAGTCCCCTGGGATAAGACTGACGCTCATGCACGAAAGCGTGGGGAGCAAACAGG
## 735                                                          ACGTAGGGTGCAAGCGTTAATCGGAATTACTGGGCGTAAAGCGTGCGCAGGCGGTTATATAAGACAGATGTGAAATCCCCGAGCTTAACTTGGGAACTGCATTTGTGACTGTATAGCTAGAGTACGGCAGAGGGGGATGGAATTCCGCGTGTAGCAGTGAAATGCGTAGATATGCGGAGGAACACCGATGGCGAAGGCAATCCCCTGGGCCTGTACTGACGCTCATGCACGAAAGCGTGGGGAGCAAACAGG
## 736                                                          ACAGAGGGTGCGAGCGTTATCCGGAATTATTGGGCGTAAAGCGTCCGCAGGTGGTTTGTTAAGTCGAGTATTAAAGACAACAGCTCAACTGTTGGAATGTATTCGATACTGGCAAGCTTGAGCCTTGGAGAGGTGAGCAGAATTCTGCATGTAGGGGTAAAATCCGTAGATATGCAGAGGAATACCAAAAGCGAAGGCAGCTCACTGGCCAAGTGCTGACACTCATGGACGAAAGCGTGGGGAGCAAACAGG
## 737                                                          ACAGAGGATGCAAGCGTTATCCGGAATCACTGGGCATAAAGCGTCTGTAGGTGGTTTGGTAAGTCTGCTGTTAAAGACTAGGGCTTAACCCTAGGAAAGCAGTGGAAACTGCTAGACTTGAGTGTGGTAGAGGTAGAGGGAATTCCTAGTGTAGCGGTGAAATGCGTAGATATTAGGAAGAACACCAATGGCGAAAGCACTCTACTGGGCCATAACTGACACTGAGAGACGACAGCTAGGGGAGCAAATGGG
## 740                                                          ACGTAGGGTGCGAGCGTTAATCGGAATTACTGGGCGTAAAGCGTGCGCAGGCGGTTTCGTAAGACAGAGGTGAAATCCCCGGGCTCAACCTGGGAACTGCCTTTGTGACTGCGAGGCTAGAGTATGGCAGAGGGGGGTGGAATTCCACGTGTAGCAGTGAAATGCGTAGAGATGTGGAGGAACACCGATGGCGAAGGCAGTCCCCTGGGATAAGACTGACGCTCATGCACGAAAGCGTGGGGAGCAAACAGG
## 741                                                          ACATAGGGTGCAAGCGTTGTCCGGAATTATTGGGCGTAAAGAGCTCGTAGGTGGTTCGTCACGTCGGATGTGAAACTCTGGGGCTTAACCCCAGACCTGCATTCGATACGGGCGAGCTTGAGTATGGTAGGGGAGTCTGGAATTCCTGGTGTAGCGGTGGAATGCGCAGATATCAGGAGGAACACCAATGGCGAAGGCAGGACTCTGGGCTAACACTGACGCTCATGCACGAAAGCGTGGGGAGCAAACAGG
## 742                                                          ACGGAGGGTGCAAGCGTTATCCGGAATCACTGGGTTTAAAGGGTACGTAGGCGGTTTAATAAGTCAGGTGTGAAATACCGCAGCCTAACTGCGGAACTGCACTTGATACTGTTAGACTTGAGAGGAGTGGAGGTATACGGAATGCATCATGTAGCGGTGAAATGCTTAGATATGATGTAGAACACCGATAGCGAAGGCAGTGTGCCACACTTTATCTGACGCTGAGGTACGAAAGCGTGGGTAGCGAACAGG
## 744                                                          ACGGAGGGTGCGAGCGTTAATCGGAATTACTGGGCGTAAAGCGCACGTAGGCGGTCATGTAAGTGGGATGTGAAATCCCCGGGCTTAACCTGGGACTTGCATCTCAGACTGCATGACTAGAGTATGGGAGAGGGTGGTGGAATTTCCGGTGTAGCGGTGAAATGCGTAGAGATCGGAAGGAACATCAGTGGCGAAGGCGGCCACCTGGCCTAATACTGACGCTGAGGTGCGAAAGCGTGGGGATCAAACAGG
## 745                                                          ACGGGGGGTGCAAGCGTTACTCGGAATCACTGGGCGTAAAGGGAGCGTAGGCGTTCTATTAAGTCTTGAGTGAAAGCCCGCAGCCTAACTGCGGAACTGCTTGGGAAACTGATAGAATTGAGTATGGAAGAGGTCAGTGGAATTTCCGGTGTAGGGGTAAAATCCGTAGATATCGGAAGGAATACCAGAAGCGAAGGCGACTGACTGGGACATTACTGACGCTGAGGCTCGAAAGCGTGGGGAGCAAACAGG
## 747                                                          ACAGAGGCCTCAAGCGTTGTTCGGAATTACTGGGCGTAAAGGGAGTGTAGGTGGTGATGCAAGTCAGATGTGAAAGCCCTGGGCTTAACCCGGGAATTGCATTTGATACTGCGTTGCTGGAGTGCAGGAGAGGTGAGAGGAATTCCTGGTGTAGGGGTGAAATCCGTAGATATCAGGAGGAATACCAATGGCGTAGGCATCTCGCTGGCCTGTAACTGACACTGAGACTCGAAAGCATGGGTAGCAAACAGG
## 748                                                          ACGTAGGGTGCAAGCGTTAATCGGAATTACTGGGCGTAAAGCGTGCGCAGGCGGTTATGTAAGACAGAGGTGAAATCCCCGGGCTCAACCTGGGAACTGCCTTTGTGACTGCATAGCTAGAGTACGGCAGAGGGGGATGGAATTCCGCGTGTAGCAGTGAAATGCGTAGATATGCGGAGGAACACCGATGGCGAAGGCAATCCCCTGGGCCTGCACTGACGCTCATGCACGAAAGCGTGGGGAGCAAACAGG
## 749                                                          ACGGAGGATCCGAGCGTTATCCGGATTTATTGGGTTTAAAGGGAGCGTAGGTGGACAGTTAAGTCAGTTGTGAAAGTTTGCGGCTCAACCGTAAAATTGCAGTTGATACTGGCTGTCTTGAGTACAGTAGAGGTGGGCGGAATTCGTGGTGTAGCGGTGAAATGCTTAGATATCACGAAGAACTCCGATTGCGAAGGCAGCTCACTGGACTGCAACTGACACTGATGCTCGAAAGTGTGGGTATCAAACAGG
## 750                                                          ACGGAGGGTGCAAGCGTTATCCGGATTTATTGGGTTTAAAGGGTCCGTAGGCGGATCGATAAGTCAGTGGTGAAAGCCCGCAGCTCAACTGCGGAACTGCCATTGATACTGTCGATCTTGAGTATAATTGAAGTGGGCGGAATGTGTGGTGTAGCGGTGAAATGCTTAGATATCACACAGAACACCGATTGCGAAGGCAGCTCACTAATTTGTAACTGACGCTGAGGGACGAAAGTGTGGGGATCAAACAGG
## 751                                                          ACAGAGACTGCAAGCGTTATTCGGATTCACTGGGCGTAAAGGGTGCGCAGGCGGCCAAGTGTGTGAGGCGTGAAAGCCCGGGGCTTAACCCCGGAATTGCGCCTCAAACTACATGGCTAGAGCATTGGAGAGGGTAGCAGAATTCATGGTGTAGCAGTGAAATGCGTAGATATCATGAGGAATACCAGAGGCGAAGGCGGCTACCTGGACAATTGCTGACGCTCAGGCACGAAAGCGTGGGGAGCAAAAGGG
## 752                                                          ACGTATGGGGCAAGCGTTGTTCGGAATTATTGGGCGTAAAGGGCTCGCAGGTGGTTTGTTAAGTTGGTGGTTTAATCTCTGGGCTCAACCCAGAGTCAGCCATCAAAACTGGCGAACTTGAGTACGATAGGGGATAGCGGAATTCTCGGTGTAGCGGTGGAATGCGTAGATATCGAGAGGAACACCAATGGCGAAGGCAGCTATCTGGATCGTAACTGACACTCATGAGCGAAAGTGCGGGGAGCAAACAGG
## 753                                                          ACGAAGGGGGCTAGCGTTGCTCGGAATCACTGGGCGTAAAGCGCACGTAGGCGGACTTTTAAGTCAGGGGTGAAATCCTGGAGCTCAACTCCAGAACTGCCTTTGATACTGGAAGTCTCGAGTTCGGGAGAGGTGAGTGGAACTGCGAGTGTAGAGGTGAAATTCGTAGATATTCGCAAGAACACCAGTGGCGAAGGCGGCTCACTGGCCCGATACTGACGCTGAGGTGCGAAAGCGTGGGGAGCAAACAGG
## 755                                                          ACGGAGGATCCAAGCGTTATCCGGAATCATTGGGTTTAAAGGGTCCGTAGGCGGCCTTATAAGTCAGTGGTGAAATCTCCTAGCTCAACTAGGAAACTGCCATTGATACTGTAGGGCTTGAATTTTTGTGAAGTAACTAGAATATGTAGTGTAGCGGTGAAATGCTTAGATATTACATGGAATACCAATTGCGAAGGCAGGTTACTAACAAACGATTGACGCTGATGGACGAAAGCGTGGGGAGCGAACAGG
## 756                                                          ACGTAGGGTGCGAGCGTTAATCGGAATTACTGGGCGTAAAGCGTGCGCAGGCGGTTTCGTAAGACAGAGGTGAAATCCCCGGGCTCAACCTGGGAACTGCCTTTGTGACTGCGAGGCTAGAGTATGGCAGAGGGGGGTGGAATTCCACGTGTAGCAGTGAAATGCGTAGAGATGTGGAGGAACACCGATGGCGAAGGCAATCCCCTGGACCTGTACTGACGCTCATGCACGAAAGCGTGGGGAGCAAACAGG
## 757                                                          ACAGAGGGTGCGAGCGTTAATCGGATTTACTGGGCGTAAAGCGTGCGTAGGCGGCTTTTTAAGTCGGATGTGAAATCCCCGAGCTTAACTTGGGAATTGCATTCGATACTGGGAAGCTAGAGTATGGGAGAGGATGGTAGAATTCCAGGTGTAGCGGTGAAATGCGTAGAGATCTGGAGGAATACCGATGGCGAAGGCAGCCATCTGGCCTAATACTGACGCTGAGGTACGAAAGCATGGGGAGCAAACAGG
## 759                                                          ACGGAGGGTGCAAGCGTTATCCGGAATCATTGGGTTTAAAGGGTCCGCAGGCGGATTTATAAGTCAGTGGTGAAAGCCTACAGCTTAACTGTAGAACTGCCATTGATACTGTAAGTCTTGAATTCGGTCGAAGTGGGCGGAATGTGTAGTGTAGCGGTGAAATGCTTAGATATTACACAGAACACCGATAGCGAAGGCAGCTCACTAGGCCTGGATTGACGCTCAGGGACGAAAGCGTGGGGAGCAAACAGG
## 761                                                          ACGGAGGGTCCGAGCGTTATCCGGATTTATTGGGTTTAAAGGGTGCGTAGGCGGTTTTATAAGTCAGCGGTGAAATTTTCCGGCTCAACCGGGATACTGCCGTTGATACTGTAAGGCTAGGATATGGTTGCTGTGGGTGGAATGTGTGGTGTAGCGGTGAAATGCTTAGAGATCACACAGAATATCGATTGCGAAGGCAGCTCACAAAGCCATTATTGACGCTGAGGCACGAAAGTGTGGGGATCAAACAGG
## 762                                                          ACAGAGGGTGCAAGCGTTAATCGGAATTACTGGGCGTAAAGCGTGCGTAGACGGTTATCTAAGTCGGATGTGAAATCCCCGGGCTCAACCTGGGAACTGCCTTTGTGACTGCGAGGCTAGAGTATGGCAGAGGGGGGTGGAATTCCACGTGTAGCAGTGAAATGCGTAGAGATGTGGAGGAACACCGATGGCGAAGGCAGCCCCCTGGGCCAATACTGACGCTCATGCACGAAAGCGTGGGGAGCAAACAGG
## 763                                                          ACGGAGGATCCAAGCGTTATCCGGATTCATTGGGTTTAAAGGGTGCGTAGGCGGAATGATAAGTCAGTGGTGAAATCTCGCAGCTTAACTGCGAAACTGCCATTGATACTGTCAATCTTGAGTACATTTGATGAGGGCGGAATGTGTCATGTAGCGGTGAAATGCTTAGATATGACACAGAACACCGATTGCGAAGGCAGCTCTCAAAACTGTAACTGACGCTGAGGCACGAAAGCGTGGGTATCAAACAGG
## 764                                                          ACGGAGGGTGCAAGCGTTAATCGGAATTACTGGGCGTAAAGCGCGCGTAGGCGGCTAGATAAGCTAGATGTGAAATCCCCGGGCTTAACCTGGGAACTGCATTTAGAACTGTCTAGCTAGAGTGCAGAAGAGGAGTGTGGAATTTCAGGTGTAGCGGTGAAATGCGTAGAGATCTGAAGGAACATCAGTGGCGAAGGCGACACTCTGGTCTGACACTGACGCTGAGGTGCGAAAGCGTGGGTAGCAAACAGG
## 766                                                          ACGTAGGGTGCAAGCGTTAATCGGAATTACTGGGCGTAAAGCGTGCGCAGGCGGTTTTGTAAGACAGTGGTGAAATCCCCGGGCTCAACCTGGGAACTGCCATTGTGACTGCAAGGCTGGAGTGCGGCAGAGGGGGATGGAATTCCGCGTGTAGCAGTGAAATGCGTAGATATGCGGAGGAACACCGATGGCGAAGGCAATCCCCTGGGCCTGCACTGACGCTCATGCACGAAAGCGTGGGGAGCAAACAGG
## 769                                                          ACGTAGGGTGCAAGCGTTGTCCGGAATTATTGGGCGTAAAGAGCTCGTAGGCGGTTTGTCGCGTCTGCTGTGAAAATCCGAGGCTCAACCTCGGACCTGCAGTGGGTACGGGCAAACTAGAGTGCGGTAGGGGAGAAGGGAATTCCTGGTGTAGCGGTGGAATGCGCAGATATCAGGAGGAACACCGATGGCGAAGGCACTTCTCTGGGCCGTAACTGACGCTGAGGAGCGAAAGCGTGGGGAGCGAACAGG
## 770                                                          ACGAAGGATGCAAGCGTTATCCGGAATTATTGGGTTTAAAGGGTGCGTAGGCGGACTGATAAGTCAGTGGTGAAAGCCTGCAGCTTAACTGCAGAATTGCCATTGATACTGTCGGTCTTGAGTACAGTTGAGGTAGGCGGAATGTGTCATGTAGCGGTGAAATGCTTAGATATGACACAGAACACCGATTGCGAAGGCAGCTTGCTAAGCTGTAACTGACGCTGAGGCACGAAAGCGTGGGGAGCAAACAGG
## 771                                                          ACGTAGGTGGCAAGCGTTGTCCGGATTTATTGGGTTTAAAGGGTGCGTAGGTGGTTTAATAAGTCAGTGGTGAAAGCTGGTTGCTCAACAATCAAGTTGCCATTGATACTGTTAGACTTGAGAGAAGTGGAGGCTGGTGGAATGGATGGTGTAGCGGTGAAATGCATAGATATCATCCAGAACGTCAATTGCGAAGGCAGCTGGCTGTACTTTTTCTGACACTGAGGCACGAAAGTGTGGGGATCAAACAGG
## 772                                                          ACGGAGGATGCAAGTGTTATCCGGAATCACTGGGCGTAAAGCGTCTGTAGGTGGTTTCATAAGTCAACTGTTAAATCTTGAGGCTCAACTTCAAAATCGCAGTCGAAACTATGAGACTGGAGTATAGTAGGGGTAAAGGGAATTTCCAGTGGAGCGGTGAAATGCGTAGAGATTGGAAAGAACACCAATGGCGAAGGCACTTTACTGGGCTATTACTGACACTGAGAGACGAAAGCTAGGGTAGCAAATGGG
## 773                                                          ACGTAGGGTGCGAGCGTTAATCGGAATTACTGGGCGTAAAGCGTGCGCAGGCGGTTTTGTAAGACAGGCGTGAAATCCCCGGGCTCAACCTGGGAACTGCGTTTGTGACTGCAAGGCTAGAGTATGGCAGAGGGGGGTGGAATTCCACGTGTAGCAGTGAAATGCGTAGAGATGTGGAGGAACACCGATGGCGAAGGCAGCCCCCTGGGCCAATACTGACGCTCATGCACGAAAGCGTGGGTAGCAAACAGG
## 775                                                          ACGTAGGTGGCGAGCGTTACTCGGAATTACTAGGCGTAAAGCGTACGTAGGCGGAAGTTTAAGTCTGTTGTGTAATCTCCAAGCTCAACTTGGAAACTGCAACAGAAACTGGGCTTCTTGAGTGAGGCAGAGGAAAACGGAATTCCTGGTGTAGCAGTGAAATGCGTAGATATCAGGAGGAACACCGGTGGCGAAGGCGGTTTTCTGGGCCTTTACTGACGCTAAAGTACGAAAGCTAGGGGAGCAAACGGG
## 778                                                          ACGGAGGGTGCAAGCGTTAATCGGAATAACTGGGCGTAAAGCGCACGCAGGCGGTTAGATAAGTCAGATGTGAAAGCCCCGGGCTCAACCTGGGAACTGCATTTGAAACTGTCTGACTAGAGTCTTGTAGAGGGGGGTAGAATTCCAGGTGTAGCGGTGAAATGCGTAGAGATCTGGAGGAATACCGGTGGCGAAGGCGGCCCCCTGGACAAAGACTGACGCTCATGCACGAAAGCGTGGGGAGCAAACAGG
## 780                                                           ACGAAGGCCCCAAGCGTTATCCGGAATTACTGGGCGTAAAGCGTCTGTAGGTGGTTTCAAAGGTTTCGTGTGAAACTTCGGGGCTCAACCCTGAATTGCACGGAAAACCATGGAACTTGAGTGTGGGAGAGGCTAGCAGAACGGTAAGAGTAGGGGTGCAATCCGTTGATACTTACCAGAATACCAAAAGCGAAGGCAGCTAGCTGGAACATTACTGACACTGAGAGACGAAAGCGTGGGGAGCGAAAGGG
## 781                                                          ACGAGAGCCCCAAGTGTTATCCGGAATTATTGGGCGTAAAGGGTGCGTAGGTGGCTGTATTAGTCTTTCGTTAAATCCTGGGGCTTAACCCCAGAACCGCGAAGGAAACGGTACGGCTAGAGGTTGTGTTGGGTGTATGGAACTCATGGAGTAGGGGTGAAATCCGTTGATATCATGGGGAACACCAAAAGCGAAGGCAGTACACTGGCACAAATCTGACACTGAGGCACGAAACCCTGGGTAGCGAATGGG
## 783                                                          ACGGAGGGTGCAAGCGTTATCCGGAATCATTGGGTTTAAAGGGTCCGCAGGCGGATTTATAAGTCAGTGGTGAAAGCCTACAGCTCAACTGTAGAACTGCCATTGATACTGTAAGTCTTGAATTCGGTCGAAGTGGGCGGAATGTGTAGTGTAGCGGTGAAATGCTTAGATATTACACAGAACACCGATAGCGAAGGCAGCTCACTAGGCCTGAATTGACGCTCATGGACGAAAGCGTGGGGAGCAAACAGG
## 784                                                          ACGTAGGGAGCAAGCGTTATCCGGATTTACTGGGTGTAAAGGGCGCGTAGGCGGGAATGCAAGTCAGATGTGAAATCTGGGGGCTCAACCCTCAAACTGCATTTGAAACTGTATTTCTTGAGTGATGGAGAGGCAAGTGGAATTCCTAGTGTAGCGGTGAAATGCGTAGATATTAGGAGGAACACCAGTGGCGAAGGCGACTTGCTGGACATTAACTGACGCTGAGGCGCGAAAGCGTGGGGAGCAAACAGG
## 785                                                         ACGTAGGATCCAAGCGTTATCCGGAATTACTGGGTGTAAAGAGTTGCGTAGGTGGCAGAGTAAGTAGGGAGTGAAAGCGTTCGGCTCAACCGAATATACATTTCCTAAACTGCTCAGCTAGAGAACGAGAGAGGTAAATGGAATTCCCAGTGTAGGAGTGAAATCCGTAGATATTGGGAGGAACACCGATGGCGTAGGCAGTTTACTGGCTCGTTTCTGACACTAAGGCACGAAAGCGTGGGGAGCAAACAGG
## 786                                                          ACGGAGGGTGCAAGCGTTGTTCGGAATTACTGGGCGTAAAGCGCGTGCAGGCGGTTTTATAAGTTTCGTGTGAAATCCCATGGCTTAACCATGGAACGGCACGGAATACTGCAAGACTTGAATACCGGAGAAGAGGGCGGAATTCCCAGTGTAGAGGTGAAATTCGTAGATATTGGGAGGAACACCGGCGGCGAAAGCGGCTCTCTAGACGGATATTGACGCTCATACGCGAGAGCGTGGGTAGCAAACAGG
## 787                                                          ACGGAGGATCCGAGCGTTATCCGGATTTATTGGGTTTAAAGGGTGCGTAGGCGGAAGAATAAGTCAGTGGTGAAAGTTTGCAGCTCAACTGTAAAATTGCCGTTGAAACTGTTTTTCTTGAGTGTAAATGAGGTAGGCGGAATGTGTTGTGTAGCGGTGAAATGCTTAGATATGACACAGAACACCGATTGCGAAGGCAGCTTACTAAGATACAACTGACGCTGAGGCACGAAAGCGTGGGGATCGAACAGG
## 788                                                          ACGAAGGGTGCTAGCGTTGTTCGGAATCATTGGGCGTAAAGCGTGCGTAGGCGGCTAAGTAAGTCGTTTGTGAAATCCCGAGGCTTAACTTCGGAATTGCAATCGAAACTGCTTAGCTTGAGTATGGTGGGGGATAGTGGAATTCCTAGTGTAGGGGTGAAATCCGTAGAGATTAGGAGGAACATCAGTGGCGAAAGCGACTATCTACGCCAATACTGACGCTGAGGCACGAAAGCGTGGGGAGCAAACAGG
## 789                                                          ACAGAGGATGCAAGCGTTATCCGGAATCACTGGGCATAAAGCGTCTGTAGGTGGTTTGATAAGTCTGCTGTTAAAGACTAGGGCTTAACCCTAGGAAAGCAGTGGAAACTGTCTGACTTGAGTATGGTAGAGGTACAGGGAATTCCTAGTGTAGCGGTGAAATGCGTAGATATTAGGAAGAACACCAATGGCGAAAGCACTGTACTGGGCCACAACTGACACTGAGAGACGACAGCTAGGGGAGCAAATGGG
## 791                                                          ACGAAGGGTGCAAGCGTTAATCGGAATTACTGGGCGTAAAGCGCGCGTAGGTGGTTTGTTAAGTTGGATGTGAAAGCCCCGGGCTCAACCTGGGAACTGCATCCAAAACTGGCAAGCTAGAGTATGGCAGAGGGTGGTGGAATTTCCTGTGTAGCGGTGAAATGCGTAGATATAGGAAGGAACACCAGTGGCGAAGGCGACCACCTGGGCTAATACTGACACTGAGGTGCGAAAGCGTGGGGAGCAAACAGG
## 793                                                          ACGTGGAGTGCGAGCGTTATCCGGATTTACTGGGCGTAAAGAGTTCGTAGGCGTTTGGTAAAGTTTCGTTTGAAAGACCGAGGCTCAACTTCGGAAACGGACGAAATACTTATCAGATTGAGATATCTAGGGGGTACTGGAACTGATAGTGTAGCAGTGAAATGCGTTGATATTATCAAGAACACCAAGGGCGAAGGCAGGTACCTGGGGATATTCTGACGCTGAGGAACGAAAGCTAGGGGAGCGAAAGGG
## 794                                                          ACGTAGGGTGCGAGCGTTAATCGGAATTACTGGGCGTAAAGCGTGCGCAGGCGGCCTTGAAAGTCAGATGTGAAATCCCCGAGCTCAACTTGGGAACTGCGTTTGAAACTCCAAGGCTAGAATATGTCAGAGGGGGGTAGAATTCCACGTGTAGCAGTGAAATGCGTAGAGATGTGGAGGAATACCAATGGCGAAGGCAGCCCCCTGGGATAATATTGACGCTCATGCACGAAAGCGTGGGGAGCAAACAGG
## 795                                                          ACGGAGGGTGCAAGCGTTATCCGGATTCACTGGGTTTAAAGGGTGCGTAGGTGGGTTGGTAAGTCAGTGGTGAAATCTCCGAGCTTAACTTGGAAACTGCCATTGATACTATCAATCTTGAATATTGTGGAGGTTAGCGGAATATGTCATGTAGCGGTGAAATGCTTAGATATGACATAGAACACCAATTGCGAAGGCAGCTGGCTACACATATATTGACACTGAGGCACGAAAGCGTGGGGATCAAACAGG
## 796                                                          ACGAAGGGGGCTAGCGTTGTTCGGAATTACTGGGCGTAAAGCGCACGTAGGCGGGTTTGTAAGTAGAGGGTGAAATCCCAGAGCTCAACTCTGGAACTGCCTTCTAGACTGCAAGCCTGGAGGTCAGGAGAGGCGAGTGGAATACCGAGTGTAGAGGTGAAATTCGTAGATATTCGGTGGAACACCAGTGGCGAAGGCGACTCGCTGGACTGATACTGACGCTGAGGTGCGAAAGCGTGGGGAGCAAACAGG
## 797                                                         ACAGAGGGTGCAAGCGTTATCCGGATTTATTGGGCGTAAAGCGTTTCGTAGGCGGTTTGGAAAGTTATCCTTCAAAGACCACGGCTTAACCGGGGGAAGGGGGGTAATACTGCCAGACTTGAAATATGGTGGGGTATCTGGAACTGATGGTGTAGTAGTGAAATACGTTGATATCATCAGGAACTCCGAGGGCGAAGGCAGGATACTAACCATTTTTTGACGCTGAGGAACGACAGCTAGGGGAGCGAAAGGG
## 798                                                          ACGGAGGGCGCGAGCGTTAATCGGAATCACTGGGCGTAAAGCGCGCGTAGGCGGCCGATCAAGTCAGAGGTGAAAGCCCTCGGCTCAACCGAGGAATTGCCTTTGATACTGGTCGGCTGGAGTGCGGGAGAGGGTGGCGGAATTCCCGGTGTAGGGGTGAAATCCGTAGAGATCGGGAGGAACACCAGTGGCGAAGGCGGCCACCTGGACCGCAACTGACGCTGAGGCGCGAAAGCATGGGGAGCAAACAGG
## 799                                                          ACGAGGGGTGCAAGCGTTATTCGGAATAACTGGGCGTAAAGAGCGCGTAGGCGGCTTGTTAAGTCTGTTGTTAAATTACCTAGCTTAACTGGGAAGAGGCAATAGAAACTGGCGAGCTAGAGGACAAGAGAGAGAAGTGGAATTCTCGGAGTAGCGGTAAAATGCGTAGATCTCGAGAGGAACACCGATGGCGAAGGCAGCTTCTTGGCTTGTATCTGACGCTCAAGTGCGAAAGCGTGGGGAGCAAACAGG
## 800                                                          ACGTAGGGTGCAAGCGTTAATCGGAATTACTGGGCGTAAAGCGTGCGCAGGCGGCTTTGCAAGACAGATGTGAAATCCCCGGGCTCAACCTGGGAACTGCATTTGTGACTGCAAGGCTAGAGTACGGTAGAGGGGGATGGAATTCCGCGTGTAGCAGTGAAATGCGTAGATATGCGGAGGAACACCGATGGCGAAGGCAATCCCCTGGGCCTGCACTGACGCTCATGCACGAAAGCGTGGGGAGCAAACAGG
## 801                                                          ACGGAGGGTGCAAGCGTTGTCCGGAATCATTGGGCGTAAAGAGTTCGTAGGTGGTATGTTAAGTCTGGTGTTAAAGGCAGAAGCTCAACTTCTGTAAGGCACTGGATACTGGCAAACTTGAATGTGGTAGAGGTAAAGGGAATTCCTGGTGTAGCGGTGAAATGCGTAGATATCAGGAGGAACATCGGTGGCGAAAGCGCTTTACTGGGCCATTATTGACACTGAGGAACGAAAGCCGGGGTAGCAAATGGG
## 802                                                          ACGAAGGGGGCTAGCGTTGTTCGGATTTACTGGGCGTAAAGCGCGTGTAGGCGGACTGGAAAGTTGGGGGTGAAATCCCGAGGCTCAACCTCGGAACTGCCTCCAAAACTTCCAGTCTGGACTCAAGCAGAGGCAAGTGGAATTGCGAGTGTAGAGGTGAAATTCGTAGATATTCGCAGGAACACCAGTGGCGAAGGCGACTTGCTGGGCTTGTAGTGACGCTGAGACGCGAAAGCGTGGGGAGCAAACAGG
## 803                                                           ACGGAGAGGGCGAGTGTTACTCATAATGACTGGGCGTAAAGGGTACGTAGGTTGTTAGGTAAGCGGTTAGTGAAATGAGAGGTTAATGACCTTTAAACGCTTTCTGAACTACCTAGCTTGAGTGCTGTTGAGGATACTGGTACTTTTAAAGTAGAGGTGATATTCGATAATATTAAAAGGACAACTTAAGGCGAAGGCAAGTATCTAGGAAGCACTGACACTGAGGTACTGAAGCATGGGTAGCAAATCGG
## 805                                                          ACGGAGGGTGCAAGCGTTACTCGGAATCACTGGGCGTAAAGGATGCGTAGGCTGTAATATAAGTCAGAAGTGAAATCCAACGGCTTAACCGTTGAACTGCTTTTGAAACTGTTTTACTAGAATATGGGAGAGGTAGATGGAATTGGTGGTGTAGGGGTAAAATCCGTAGATATCACCAGGAATACCGATTGCGAAGGCGATCTACTGGAACATTATTGACGCTGAGGCATGAAAGCGTGGGGAGCAAACAGG
## 806                                                          ACATAGGGTGCAAGCGTTGTCCGGAATTATTGGGCGTAAAGAGCTCGTAGGTCGTTTGTTACGTCGGATGTGAAAACCTGAGGCTCAACCTCAGGCCTGCATTCGATACGGGCAAACTAGAGTTTGGTAGGGGAGACTGGAATTCCTGGTGTAGCGGTGGAATGCGCAGATATCAGGAGGAACACCAATGGCGAAGGCAGGACTCTGGGCCATTACTGACACTGAGGAGCGAAAGCGTGGGGAGCGAACAGG
## 807                                                          ACGAAGGGTGCAAGCGTTATCCGGATTCATTGGGTTTAAAGGGTGCGTAGGCGGAGCGTTAAGTCAGTGGTGAAATCCTGCAGCTCAACTGTAGACTTGCCATTGATACTGGCGCTCTTGAGTGCGCTTGAAGTGGGCGGAATGTGCCGTGTAGCGGTGAAATGCTTAGATATGGCACAGAACACCAATTGCGAAGGCAGCTCACTAAGGCGATACTGACGCTGAGGCACGAAAGCGTGGGGATCGAACAGG
## 808                                                          ACGGGAGTGGCAAGCGTTATCCGGAATTATTGGGCGTAAAGCGTCCGCAGGCGGCCTTGAAAGTCTGTCGTTAAAGCGTGGAGCTTAACTCCATTTAAGCGATGGAAACTACAAGGCTAGAGTGTGGTAGGGGCAGAGGGAATTCCCGGTGTAGCGGTGAAATGCGTAGATATCGGGAAGAACACCAGTGGCGAAGGCGCTCTGCTGGGCCATAACTGACGCTCATGGACGAAAGCCAGGGGAGCGAAAGGG
## 809                                                          ACGTAGGGTGCGAGCGTTAATCGGAATTACTGGGCGTAAAGCGTGCGCAGGCGGTTTTGTAAGTCAGATGTGAAAGCCCCGGGCTCAACCTGGGAACTGCGTTTGAAACTACAAGGCTAGAGTGTGTCAGAGGGGGGTAGAATTCCACGTGTAGCAGTGAAATGCGTAGAGATGTGGAGGAATACCGATGGCGAAGGCAGCCCCCTGGGATAACACTGACGCTCATGCACGAAAGCGTGGGGAGCAAACAGG
## 810                                                          ACGGAGGGGGCTAGCGTTGTTCGGAATCACTGGGCGTAAAGCGTGCGTAGGCGGCCATTCAAGTCAGAGGTGAAAGCCCAGGGCTCAACCTTGGAACTGCCTTTGAAACTAGATGGCTCGAACACGGGAGAGGTGAGTGGAATTCCGAGTGTAGAGGTGAAATTCGTAGATATTCGGAAGAACACCAGTGGCGAAGGCGGCTCACTGGACCGTTGTTGACGCTGAGGCACGAAAGCGTGGGGAGCAAACAGG
## 811                                                          ACGGAGGGGGCTAGCGTTATTCGGAATTACTGGGCGTAAAGCGCACGTAGGCGGATTGGAAAGTCAGAGGTGAAATCCCAGAGCTCAACTTTGGAACTGCCTTTGAAACTCCCAGTCTTGAGGTCGAGAGAGGTGAGTGGAATTCCGAGTGTAGAGGTGAAATTCGTAGATATTCGGAGGAACACCAGTGGCGAAGGCGGCTCACTGGCTCGATACTGACGCTGAGGTGCGAAAGCGTGGGGAGCAAACAGG
## 812                                                          ACGGAGGGTGCAAGCGTTAATCGGAATGACTGGGCGTAAAGCGCACGCAGGCGGTCTGTTAAGTTGGATGTGAAATCCCCGGGCTTAACCTGGGAACTGCATTCAAAACTGACAGGCTAGAGTCTCGTAGAGGGGGGTAGAATTCCAGGTGTAGCGGTGAAATGCGTAGAGATCTGGAGGAATACCGGTGGCGAAGGCGGCCCCCTGGACGAAGACTGACGCTCAGGTGCGAAAGCGTGGGGAGCAAACAGG
## 813                                                          ACGGAAGGTCCGGGCGTTATCCGGATTTATTGGGTTTAAAGGGAGCGTAGGCCGGAGATTAAGCGTGTTGTGAAATGTAGATGCTCAACATCTGCACTGCAGCGCGAACTGGTTTCCTTGAGTACGCACAAAGTGGGCGGAATTCGTGGTGTAGCGGTGAAATGCTTAGATATCACGAAGAACTCCGATTGCGAAGGCAGCTCACTGGAGCGCAACTGACGCTGAAGCTCGAAAGTGCGGGTATCGAACAGG
## 814                                                          ACGGAGGGTGCAAGCGTTAATCGGAATTACTGGGCGTAAAGGGTGCGTAGGTGGTTTGCTAAGTTATCTGTGAAATTCCTGGGCTTAACCTGGGCAGGTCAGATAAAACTGGTGAACTCGAGTATGGGAGAGGGTAGTGGAATTTCCGGTGTAGCGGTGAAATGCGTAGAGATCGGAAGGAACACCAGTGGCGAAGGCGGCTACCTGGCCTAATACTGACACTGAGGCACGAAAGCGTGGGGAGCAAACAGG
## 815                                                          ACGTAGGTGGCGAGCGTTACTCGGAGTTACTGGGCGTAAAGGGTCCGTAGGTGGCCATCTAAGTCTGAGGTGAAAGCCTGCAGCTCAACTGCAGAACGGCCTTGGGTACTGGGTGGCTAGAGTGCAGGAGAGGTAAGTGGAATTCGTGGTGTAGCGGTGGAATGCGTAGATATCACGAGGAACACCAGAGGCGAAAGCGGCTTACTGGACTGCAACTGACACTGAGGGACGAAAGCCAGGGGAGCAAAAAGG
## 816                                                          ACGTAGGGTGCAAGCGTTAATCGGAATTACTGGGCGTAAAGCGTGCGCAGGCGGTCTTGCAAGACAGATGTGAAATCCCCGGGCTTAACCTGGGAACTGCATTTGTGACTGCAAGGCTGGAGTGCGGCAGAGGGGGATGGAATTCCGCGTGTAGCAGTGAAATGCGTAGATATGCGGAGGAACACCGATGGCGAAGGCAATCCCCTGGGCCTGCACTGACGCTCATGCACGAAAGCGTGGGGAGCAAACAGG
## 817                                                          ACGAAGGGACCTAGCGTAGTTCGGAATTACTGGGCTTAAAGAGTTCGTAGGTGGTTAAAAAAGTTGATGGTGAAATCCCAAGGCTCAACCTTGGAACTGCCATCAAAACTTTTTAGCTAGAGTGTGATAGAGGAAAGTGGAATTTCTAGTGTAGAGGTGAAATTCGTAGATATTAGAAAGAACATCAAAAGCGAAGGCAACTTTCTGGATCATTACTGACACTGAGGAACGAAAGCATGGGTAGCGAAGAGG
## 818                                                          ACAGAGGGTGCAAACGTTGCTCGGAATTATTGGGCGTAAAGCGCGTGTAGGCGGTCTCGTAAGTCGACTGTGAAAGCCCTTGGCTCAACCAAGGAAGTGCAGTCGAAACTGCGAGACTAGAGTGCTGAAGAGGATAGCGGAATGGCCAGTGTAGAGGTGAAATTCGTAGATATTGGCTAGAACACCGGTGGCGAAGGCGGCTATCTGGGAAGTTACTGACGCTGAGACGCGAAAGCGTGGGGAGCAAACAGG
## 821                                                          ACGGGAGTGGCAAGCGTTATCCGGAATTATTGGGCGTAAAGCGTCCGCAGGCGGTCTTTTAAGTCTGCTGTTAAAGCGTGGAGCTTAACTCCATTTCGGCAGTGGAAACTGGAAGACTAGAGTGTGGTAGGGGCAGAGGGAATTCCCGGTGTAGCGGTGAAATGCGTAGATATCGGGAAGAACACCAGTGGCGAAGGCGCTCTGCTGGGCCATAACTGACGCTCATGGACGAAAGCCAGGGGAGCGAAAGGG
## 822                                                          ACGGAGGGTGCAAGCGTTATCCGGATTCACTGGGTTTAAAGGGTGCGTAGGTGGGCAGGTAAGTCAGTGGTGAAATCTTCGAGCTTAACTCGGAAACTGCCATTGATACTATCTGTCTTGAATATCCTGGAGGTGAGCGGAATATGTCATGTAGCGGTGAAATGCTTAGATATGACATAGAACACCAATTGCGAAGGCAGCTCACTACGGGATGATTGACACTGAGGCACGAAAGCGTGGGGATCAAACAGG
## 823                                                          ACGAGGGGTCCAAGCGTTGTTCGGAATTATTGGGCGTAAAGCGGGTGTAGGTGGATCTATAAGTCAGGAGTGAAAGCCCCGAGCTTAACTTGGGAAGTGCTTTTGATACTGTAGATCTTGAATGTGGGAGAGGATCGTGGAATTCCAGGTGTAGTGGTGAAATACGTAGATATCTGGAGGAACACCGGTGGCGAAGGCGGCGATCTGGCCCAACATTGACACTGAGACCCGAAAGCATGGGGATCAAACAGG
## 824                                                          ACGGAGGGTGCAAGCGTTATCCGGATTCACTGGGTTTAAAGGGTGCGTAGGTGGGCAGGTAAGTCAGTGGTGAAATCCCCGAGCTCAACTTGGGAACTGCCATTGATACTATCTGTCTTGAATACCGTGGAGGTTAGCGGAATATGTCATGTAGCGGTGAAATGCTTAGATATGACATAGAACACCGATTGCGAAGGCAGCTGGCTACACGAATATTGACACTGAGGCACGAAAGCGTGGGGATCAAACAGG
## 825                                                          ACGTAGGTGGCAAGCGTTGTCCGGATTTACTGGGCGTAAAGGGAGCGTAGGTGGATATTTAAGTGGGATGTGAAATACTCGGGCTTAACCTGGGTGCTGCATTCCAAACTGGATATCTAGAGTGCAGGAGAGGAAAGTAGAATTCCTAGTGTAGCGGTGAAATGCGTAGAGATTAGGAAGAATACCAGTGGCGAAGGCGACTTTCTGGACTGTAACTGACACTGAGGCTCGAAAGCGTGGGGAGCAAACAGG
## 827                                                          ACGGAGGGTGCAAGCGTTAATCGGAATTACTGGGCGTAAAGCGCACGCAGGCGGTTGGATAAGTTAGATGTGAAAGCCCCGGGCTCAACCTGGGAATTGCATTTAAAACTGTCCAGCTAGAGTCTTGTAGAGGGGGGTAGAATTCCAGGTGTAGCGGTGAAATGCGTAGAGATCTGGAGGAATACCGGTGGCGAAGGCGGCCCCCTGGACCTGTACTGACGCTCATGCACGAAAGCGTGGGGAGCAAACAGG
## 828                                                          ACGTAGGGTGCAAGCGTTAATCGGAATTACTGGGCGTAAAGCGTGCGCAGGCGGTTATATAAGACAGATGTGAAATCCCCGGGCTCAACCTGGGAACTGCATTTGTGACTGTATAGCTAGAGTACGGTAGAGGGGGATGGAATTCCGCGTGTAGCAGTGAAATGCGTAGATATGCGGAGGAACACCGATGGCGAAGGCAATCCCCTGGGCCTGTACTGACGCTCATGCACGAAAGCGTGGGGAGCAAACAGG
## 830                                                      ACAGAGACCTCAAGCGTTATCCGGAATCATTGGGCGTAAAGCGTACCGATAGGTGGTTTAGAAAGTCAGAAGTGAAATCTCCAAGCTCAACTTGGAGTCTGTCTTTTGAAACTTCTAAACTAGAGGGGCAAAGAGGAAGCTGGAACGAACGGTGTAGTAGTGAAATGCGTTGATATCGTTCGGAACACCAATAGCGAAGGCAAGCTTCTGGGTGCCACCTGACACTGCTAGGACGAGAGCGTGGGGAGCGAATGGG
## 831                                                          ACGTAGGGTGCAAGCGTTGTCCGGATTTATTGGGCGTAAAGAGCTCGTAGGCGGTTTGTTACGTCGGATGTGAAATCCTGGAGCTTAACTCCGGGCCTGCATTCGATACGGGCAGACTAGAGTGTTGTAGGGGAGACTGGAACTCCTGGTGTAGCGGTGAAATGCGCAGATATCAGGAAGAACACCGATGGCGAAGGCAGGTCTCTGGGCAACTACTGACGCTGAGGAGCGAAAGCGTGGGGAGCAAACAGG
## 832                                                          ACGGAGGGTGCAAGCGTTGTTCGGAATTACTGGGCGTAAAGCGCGTGCAGGTGGTTTTTTAAGTCTCGTGTGAAATCCCTCGGCTTAACCGAGGAACTGCGCGAGATACTGGAGAACTTGAATACGGGAGAAGAGGGCGGAATTCCCAGTGTAGAGGTGAAATTCGTAGATATTGGGAGGAACACCGGCGGCGAAAGCGGCTCTCTAGACCGATATTGACGCTCATACGCGAGAGCGTGGGTAGCAAACAGG
## 833                                                          ACGGAGGGTGCAAGCGTTATCCGGAATCATTGGGTTTAAAGGGTCCGCAGGCGGATTTATAAGTCAGTGGTGAAATCCTATCGCTTAACGATAGAACTGCCATTGATACTGTAAGTCTTGAATTCGGTCGGAGTGGGCGGAATGTGTAGTGTAGCGGTGAAATGCATAGATATTACACAGAACACCGATAGCGAAGGCAGCTCACTAGGCCTGAATTGACGCTCATGGACGAAAGCGTGGGGAGCAAACAGG
## 834                                                          ACAGAGGGTGCAAGCGTTGTTCGGATTTACTGGGCGTAAAGCGTGCGTAGGTTGTATTGAGAGTCGCGGGTGAAATCCCAGGGCTTAACCCTGGAACTGCCTGCGAGACCTCAGTACTGGAGTATGAAAGAGGATAGTGGAATTCCTGGTGTAGCGGTGAAATGCGTAGAGATCAGGAGGAACACCTGTGGCGAAGGCGGCTATCTGGTTCATTACTGACACTGAGGCACGAAAGCGTGGGTAGCAAACAGG
## 835                                                          ACGAAGGGGGCTAGCGTTGCTCGGAATCACTGGGCGTAAAGCGCACGTAGGCGGATCTTTAAGTCAGAGGTGAAATCCCAAGGCTCAACCTTGGAACTGCCTTTGATACTGGGGATCTCGAGTCCGGAAGAGGTGAGTGGAACTCCGAGTGTAGAGGTGAAATTCGTAGATATTCGGAAGAACACCAGTGGCGAAGGCGGCTCACTGGTCCGGTACTGACGCTGAGGTGCGAAAGCGTGGGGAGCAAACAGG
## 836                                                          ACGGAGGGTGCAAGCGTTACTCGGAATCACTGGGCGTAAAGAGCGTGTAGGCGGATTGATAAGTTTGAAGTGAAATCCTATAGCTTAACTATAGAACTGCTTTGAAAACTGTTAATCTAGAATGTGGGAGAGGTAGATGGAATTTCTGGTGTAGGGGTAAAATCCGTAGAGATCAGAAGGAATACCGATTGCGAAGGCGATCTACTGGAACATTATTGACGCTGAGATGCGAAAGCGTGGGGAGCAAACAGG
## 837                                                          ACGTAGGTGGCAAGCGTTGTCCGGATTTACTGGGCGTAAAGGATGCGTAGGCGGATTTTTAAGTCAGATGTGAAATACCCGAGCTTAACTTGGGTGCTGCATTTGAAACTGGAAGTCTAGAGTGCGGGAGAGGAGAGTGGAATTCCTAGTGTAGCGGTGAAATGCGTAGAGATTAGGAAGAACACCAGTGGCGAAGGCGACTCTCTGGACCGTAACTGACGCTGAGGCATGAAAGCGTGGGTAGCAAACAGG
## 839                                                          ACATAGGGTGCAAGCGTTGTCCGGAATTATTGGGCGTAAAGAGCTCGTAGGTGGTCGATCACGTCGGATGTGAAAATCTGAGGCTTAACCTCAGACCTGCATTCGATACGGGTTGACTAGAGTGTGGTAGGGGAGACTGGAATTCCTGGTGTAGCGGTGGAATGCGCAGATATCAGGAGGAACACCGATGGCGAAGGCAGGTCTCTGGGCCATTACTGACACTGAGGAGCGAAAGCGTGGGGAGCGAACAGG
## 840                                                          ACGTAGGGTGCGAGCGTTAATCGGAATTACTGGGCGTAAAGGGTGCGCAGGCGGCTATGTAAGTCAGATGTGAAATCCCCGGGCTTAACCTGGGAATTGCGTTTGAAACTACATGACTAGAGTGTAACAGAGGGAAGTGGAATTCCATGTGTAGCAGTGAAATGCGTAGAGATATGGAGGAACATCGATGGCGAAGGCAGCTTCCTGGGTTAACACTGACGCTCATGCACGAAAGCGTGGGGAGCAAACAGG
## 843                                                           ACAGAGGGTGCAAGCGTTATTCGGAATTACTGGGCGTAAAGCGCGCGCGGGCTGTCTAGCAAGTCTGATGTGAAATCCCCTGCAAAAACAGGGAAGTGCATTAGATACTACTAGACTAGAGTATAGGAGGGGAGAGTGGAATTCCTGGTGTAGAGGTGAAATTCGCAGATATCAGGAGGAACAACGGTGGCGAAGGCGACTCTCTGGACTATTACTGACGCTGAGGCGCGAAAGCGTGGGGAGCAAACAGG
## 844                                                          ACGTAGGGTGCGAGCGTTAATCGGAATTACTGGGCGTAAAGCGTGCGCAGGCGGTTTTGTAAGTCAGATGTGAAATCCCCGGGCTCAACCTGGGAACTGCATTTGAGACTGCAAGGCTAGAGTGTAGCAGAGGGGGGTAGAATTCCACGTGTAGCAGTGAAATGCGTAGAGATGTGGAGGAATACCGATGGCGAAGGCAGCCCCCTGGGTTACTACTGACGCTCATGCACGAAAGCGTGGGGAGCAAACAGG
## 845                                                          ACGAAGGTGGCAAGCGTTACTCGGAATTACTAGGTGTAAAGCGCAGGTAGGTGGTTAGATTAGTCTGTAATGAAATCTCCCGGCTTAACTGGGAGAGGTTTATGGATACTGTCTAGCTCGAGTGTGGGAGAGGGTGCTGGAATTCCCGGTGTAGCGGTGAAATGCGTAGATATCGGGAGGAACACCAATGGCGAAAGCAGGCACCTGGCCCAATACTGACACTGAGCTGCGAAAGCTAGGGGAGCAAACAGG
## 846                                                          ACGAGGGGTCCAAGCGTTGTTCGGAATTATTGGGCGTAAAGCGGGTGTAGGTGGCTCTGTAAGTCAGGAGTGAAAGCCCCGAGCTTAACTTGGGAAGTGCTTTTGATACTGCAGAGCTTGAGTGTGGGAGAGGATCGTGGAATTCCAGGTGTAGTGGTGAAATACGTAGATATCTGGAGGAACACCGGTGGCGAAGGCGGCGATCTGGCCCAACACTGACACTGAGACCCGAAAGCGTGGGGATCAAACAGG
## 847                                                          ACAGAGACCTCAAACGTTATCCGGATTTATTGGGCGTAAAGGGTCCGCAGGTGGTTTGAAAAGTCCGTGGTTAAACCCCGATGCTTAACATCGGGACTGCTGTGGATACTGTCAAACTTGAGGCTGGGAGAGGCAAGCGGAACTATCGGTGTAGTCGTAATAAGCGCTGATATCGATGGGAACACCAAAGGCGAAGGCAGCTTGCTGGAACATGCCTGACACTCAGGGACGAAAGCGTGGGGAGCGAAGGGG
## 848                                                          ACGAGGGGTGCAAGCGTTGTTCGGAATTACTGGGCGTAAAGCGCGTGTAGGTGGTCAAGCAAGTCGGATGTGAAAGCCCTCGGCTTAACCGAGGAAGTGCGTCCGAAACTATTTGGCTTGAGTGTGGGAGAGGAGGGTGGAATTCCCAGTGTAGAGGTGAAATTCGTAGATATTGGGAGGAACACCGGTGACGAAGGTGACTCTCTGGACCACAACTGACACTAAGACGCGAAAGCGTGGGTAGCAAACAGG
## 849                                                          ACAGAGACTGCAAGCGTTATTCGGATTCACTGGGCGTAAAGGGTGCGCAGGCGGTTGGGTGTGTCAGATGTGAAATCCCGAGGCTTAACCTCGGAACTGCGTCTGAAACTACTCGACTAGAGTACTGGAGAGGGAAACGGAATTCACGGTGTAGCAGTGAAATGCGTAGATATCGTGAGGAACACCAGAGGCGAAGGCGGTTTCCTGGACAGTTACTGACGCTCAGGCACGAAAGCATGGGGAGCAAAAGGG
## 850                                                          ACGGGGGGTGCAAGCGTTACTCGGAATCACTGGGCGTAAAGAGCATGTAGGCGGATTAATAAGTTTGAAGTGAAATCCTATAGCTTAACTATAGAACTGCTTTGAAAACTGTTAATCTAGAATGTGGGAGAGGTAGATGGAATTTCTGGTGTAGGGGTAAAATCCGTAGAGATCAGAAGGAATACCGATTGCGAAGGCGATCTACTGGAACATTATTGACGCTGAGGCATGAAAGCGTGGGGAGCAAACAGG
## 851                                                          ACGTAGGGTGCAAGCGTTGTCCGGAATTATTGGGCGTAAAGAGCTCGTAGGCGGTTGGTCGCGTCTGCTGTGAAAACCCGAGGCTCAACCTCGGGCCTGCAGTGGGTACGGGCCAACTAGAGTGCGGTAGGGGAGATCGGAATTCCTGGTGTAGCGGTGGAATGCGCAGATATCAGGAGGAACACCAATGGCGAAGGCAGATCTCTGGGCCGTAACTGACGCTGAGGAGCGAAAGCGTGGGGAGCGAACAGG
## 852                                                          ACGGAGGGTGCAAGCGTTATCCGGAATCACTGGGTTTAAAGGGTGCGTAGGCGGCTTAGTAAGTCAGTGGTGAAAGGCTGTGGCTTAACCATGGAATTGCCATTGATACTGCAGAGCTTGAATGAGGTTGAGGTTGGCGGAATGTGACATGTAGCGGTGAAATGCTTAGATATGTCATGGAACACCGATTGCGAAGGCAGCTGACTGGACCTATATTGACGCTGAGGCACGAAAGCGTGGGGAGCGAACAGG
## 853                                                          ACAGGGGATGCAAGTGTTATCCGGATTCATTGGGCGTAAAGCGTCTGCAGGTTGATATTTAAGTCTTTTGTTAAAACTTCGGGCTTAACCCGAAATCTGCAAAAGAAACTATGTATCTTGAGTATGGTAGAGGTAAAGGGAATTTCCAGTGGAGCGGTGAAATGCGTAGAGATTGGAAAGAACACCAACAGCGAAGGCACTTTACTGGGCCAGTACTGACACTGAGAGACGAAAGCTAGGGGAGCAAACGGG
## 854                                                          ACGAAGGATCCAAGCGTTGTCCGGATTTACTGGGTTTAAAGGGTGCGTAGGCGGGCTATTAAGTCAGTGGTGAAAGCTTGCAGCTTAACTGTAAAATTGCCATTGAAACTGATAGCCTCGAATATGGTTGAGGTGGTTGGAATGTATCATGTAGCGGTGAAATGCTTAGATATGATACAGAACACCGATTGCGAAGGCAGATCACTAAGCCATTATTGACGCTGAGGCACGAAAGCGTGGGGATCAAACAGG
## 857                                                          ACAGAGGTGGCGAGCGTTGTTCGGATTTACTGGGCGTAAAGGGCGCGTAGGCGGCGCGGTGTGTCGGGTGTGAAATCCATGGGCTCAACCCATGAAGTGCGCCCGAAACTGCCGCGCTCGAGTTCGGGAGGGGGGATCGGAATGCAGGGTGTAGCGGTGAAATGCGTTGATATCCTGCAGAACACCGGAGGCGAAGGCGGATCCCTGGAACGACACTGACGCTGAGGCGCGAAAGCAGGGGGAGCAAACAGG
## 858                                                          ACGTAGGGTGCAAGCGTTAATCGGAATTACTGGGCGTAAAGCGTGCGCAGGCGGTTCCATAAGACAGATGTGAAATCCCCGGGCTCAACCTGGGAACTGCATTTGTGACTGTGGGGCTAGAGTACGGTAGAGGGGGATGGAATTCCGCGTGTAGCAGTGAAATGCGTAGATATGCGGAGGAACACCGATGGCGAAGGCAATCCCCTGGACCTGTACTGACGCTCATGCACGAAAGCGTGGGGAGCAAACAGG
## 859                                                          ACGTAGGATGCAAGCGTTGTCCGGATTTATTGGGCGTAAAGAGTTCGTAGGCGGTTATTTAAGTCTGGTGTTAAATACTGTGGCCCAACCACAGAACTGCATTGGATACTGGATGACTTGAGTGCGGTAGAGGCGAGTGGAATTCCCAGTGTAGCGGTGAAATGCGTAGATATTGGGAAGAACACCAGTGGCGTAAGCGACTCGCTGGCCCGTAACTGACGCTGAGGAACGAAAGCCAGGGGAGCGAATGGG
## 860                                                          ACGTAGGTGGCAAGCGTTGTCCGGATTTATTGGGTTTAAAGGGTGCGTAGGCGGTCCATTAAGTCAGTGGTGAAATACGGCAGCTTAACTGTCGAGGTGCCATTGATACTGGAGGACTTGAGTACAGACGAGGTAGGCGGAATTGACGGTGTAGCGGTGAAATGCTTAGATATCGTCAAGAACACCGATAGCGAAGGCAGCTTACTAGACTGTAACTGACGCTGAGGCACGAAAGTGTGGGGATCAAACAGG
## 864                                                          ACAGAGGGTGCAAGCGTTGCTCGGAATTATTGGGCGTAAAGGGCAGGTAGGCGGTCTTATTTGTCAGAGGTGAAAGCCTTGGGCTTAACCCAAGAAGTGCCTCTGAAACGGTAAGACTAGAGTACTAGATAGGGTCGCGGAATTCCTGGTGTAGCGGTGAAATGCGTAGAGATCAGGAGGAACATCGGAGGCGAAGGCGGCGACCTGGAGAGTGACTGACGCTCAACTGCGAAAGCGTGGGTAGCAAACAGG
## 865                                                          ACGAAGGTGGCTAGCGTTGTTCGGATTTACTGGGCGTAAAGGGAGCGTAGGCGGTGAAACAAGTCAGATGTGAAAGCCCAAGGCTTAACCTTGGAATAGCATTTGAAACTGTTTTGCTAGAGTATGGAAGAGGATAGCGGAATTCCTAGTGTAGAGGTGAAATTCGTAGATATTAGGAAGAACACCAGTGGCGAAGGCGGCTATCTGGTCCATTACTGACGCTGAAGTTCGAAAGTGTGGGGATCAAACAGG
## 866                                                          ACGAAGGGGGCTAGCGTTGTTCGGAATTACTGGGCGTAAAGCGCGCGCAGGCGGCTATCCAAGTCAGTGGTGAAAGCCCGGAGCTCAACTCCGGAACTGCCATTGAAACTGTTTAGCTTGAGGACGAGAGAGGTGAGTGGAATTCCCAGTGTAGAGGTGAAATTCGTAGATATTGGGAAGAACACCGGTGGCGAAGGCGGCTCACTGGCTCGTATCTGACGCTCAGGCGCGATAGCGTGGGGATCAAACAGG
## 868                                                           ACGTAGGTGGCAAGCGTTGTCCGGAATTATTGGGCGTAAAGGGAGCGCAGGCGGGAAGGTAAGTCGGTCTTAAAAGTGCGGGGCTCAACCCCGTGATGGGATCGAAACTATCTTTCTTGAGTGCAGGAGAGGAAAGTGGAATTCCTAGTGTAGCGGTGAAATGCGTAGATATTAGGAGGAACACCAGTGGCGAAGGCGACTTTCTGGACTGTAACTGACGCTGAGGCTCGAAAGCCAGGGGAGCGAACGGG
## 869                                                          ACGAGGGACCCTAGCGTTGTTCGGAATCATTGGGCGTAAAGCGGGTGCAGGTGGCTTTGTAAGTCAGGTGTGAAAGCCCAGGGCTCAACCCTGGAAGTGCATTTGATACTGCGAAGCTTGAGTGCTGGAGAGGTTACTAGAATTCCAGGTGTAGTGGTGAAATACGTAGATATCTGGAGGAATACCGGAGGCGAAGGCGGGTAACTGGCCAGACACTGACACTCAGACCCGAAAGCGTGGGGATCAAACAGG
## 871                                                           ACGTAGGTGGCAAGCGTTGTCCGGAATTATTGGGCGTAAAGGGCGTGCAGGCGGCCAGGCAAGTCTGTCTTAAAAGTGCGGGGCTTAACCCCGTGATGGGATGGAAACTGTCAGGCTAGAGTGTCGGAGAGGAAAGCGGAATTCCTAGTGTAGCGGTGAAATGCGTAGATATTAGGAGGAACACCAGTGGCGAAAGCGGCTTTCTGGACGACAACTGACGCTGAGGCGCGAAAGCCAGGGGAGCAAACGGG
## 872                                                          ACGGAGGATCCAAGCGTTATCCGGAATCATTGGGTTTAAAGGGTCCGTAGGCGGTTTTATAAGTCAGTGGTGAAATCCGGCAGCTCAACTGTCGAACTGCCATTGATACTGTAGAACTTGAATTACTTGGAAGTAACTAGAATATGTAGTGTAGCGGTGAAATGCTTAGAGATTACATGGAATACCAATTGCGAAGGCAGGTTACTACGAGTATATTGACGCTGATGGACGAAAGCGTGGGGAGCGAACAGG
## 875                                                          ACGTAGGATGCAAGCGTTGTCCGGAATCATTGGGCGTAAAGCGTTCGTAGGCGGCATATTAAGTCTGATGTTAAAGCCCGGGGCTCAACTCCGGTTCGGCATTGGATACTGATAAGCTAGAGTGCAATAGAGGTAGGGGGAATTCCTAGTGTAGCGGTGAAATGCGTAGATATTAGGAGGAACACCGGTGGCGAAAGCGCCCTACTGGGTTGTAACTGACGCTGAGGAACGAAAGCCAGGGTAGCAAATGGG
## 876                                                          ACGGAGGATGCAAGTGTTATCCGGAATCACTGGGCGTAAAGCGTCTGTAGGTGGTCTAATAAGTCAACTGTTAAATCTTGAGGCTCAACTTCAAAACCGCAGTCGAAACTATTAGACTAGAGTATAGTAGAGGTAAAGGGAATTTCCAGTGGAGCGGTGAAATGCGTAGATATTGGAAAGAACACCGATGGCGAAAGCACTTTACTGGGCTATTACTAACACTCAGAGACGAAAGCTAGGGTAGCAAATGGG
## 877                                                          ACAGAGACTGCAAGCGTTACTCGGATTCACTGGGCGTAAAGGGAGCGCAGGCGGACTGGTGTGTCAGGCGTGAAATCCCGGGGCTCAACCCCGGGGCTGCGTTTGAAACTACCAGTCTAGAGATTCGGAGGGGTAAGCGGAATTCGTGGTGGAGCAGTGAAATGCGTAGATATCACGAGGAACACCGACGGCGAAGGCAGCTTACTGGACGAAATCTGACGCTCAGGCTCGAAAGCATGGGGAGCAAAAGGG
## 881                                                          ACGTAGGACCCGAGCGTTGTCCGGATTTACTGGGTATAAAGGGTGCGTAGGCGGCCTTGTGCGTCAGAGGTGAAATATCCGGGCTTAACCCGGAGGGTGCCTTTGATACGGCGAGGCTTGAGTGCGAGAGAGGATGATGGAATTCCTGGTGTAGCGGTGAAATGCGTAGATATCAGGAGGAACACCGGTGGCGAAGGCGGTCATCTGGCTCGTAACTGACGCTGAGGCACGAAAGCGCGGGGATCAAACAGG
## 882                                                          ACGTAGGGTGCAAGCGTTAATCGGAATTACTGGGCGTAAAGCGTGCGCAGGCGGTTATATAAGACAGATGTGAAATCCCCGGGCTCAACCTGGGAACTGCATTTGTGACTGTATAGCTAGAGTACGGTAGAGGGGGATGGAATTCCGCGTGTAGCAGTGAAATGCGTAGATATGCGGAGGAACACCGATGGCGAAGGCAATCCCCTGGGCCTGCACTGACGCTCATGCACGAAAGCGTGGGGAGCAAACAGG
## 883                                                          ACGGAGGGTGCAAGCGTTAATCGGAATTACTGGGCGTAAAGGGTTCGTAGGTGGTTAATTAAGTTATCTGTGAAATCCCCGGGCTCAACCTGGGCAGGTCAGATAATACTGGTTAACTCGAGTATGGGAGAGGGTAGTGGAATTTCCGGTGTAGCGGTGAAATGCGTAGAGATCGGAAGGAACACCAGTGGCGAAGGCGGCTACCTGGCCTAATACTGACACTGAGGAACGAAAGCGTGGGGAGCAAACAGG
## 884                                                          ACAGAGGGTGCAAGCGTTGCTCGGAATTATTGGGCGTAAAGGGCAAGTAGGTGGTCTGATTTGTCAGGGGTGAAAGCCTTGGGCTTAACCTGAGAAGTGCCCCTGAAACGGTCAGACTTGAGTCCTGGAGAGGGTCGTGGAATTCCCGGTGTAGCGGTGAAATGCGTAGAGATCGGGAGGAACACCTGAGGCGAAGGCGGCGACCTGGACAGGAACTGACACTCAACTGCGAAAGCATGGGTAGCAAACAGG
## 885                                                          ACAGAGGGTGCAAGCGTTAATCGGAATTACTGGGCGTAAAGCGCGCGTAGGCGGCTGATTAAGTCGGATGTGAAATCCCCGGGCTTAACCTGGGAACTGCGTTCGATACTGATCGGCTAGAGTATGAGAGAGGGAGGTAGAATTCCACGTGTAGCGGTGAAATGCGTAGATATGTGGAGGAATACCGGTGGCGAAGGCGGCCTCCTGGCTTAATACTGACGCTGAGGTGCGAAAGCGTGGGGAGCAAACAGG
## 886                                                          ACGTAGGGGGCAAGCGTTATCCGGATTTACTGGGTGTAAAGGGAGTGTAGACGGTCTGGTAAGTCAGATGTGAAAGCCCGGGGCTCAACCCCGGGACTGCATTTGAAACTATCAGACTAGAGTGCAGGAGAGGTAAGTGGAATTCCTAGTGTAGCGGTGAAATGCGTAGATATTAGGAGGAACACCAGTGGCGAAGGCGGCTTACTGGACTGTAACTGACGTTGAGGCTCGAAAGCGTGGGGAGCAAACAGG
## 887                                                          ACGAATGTGGCAAGCGTTGTTCGGAATCACTGGGCGTAAAGAATGCGTAGGCGGCTTGGCAAGTCAGAAGTGAAATCCCAGGGCTTAACCCTGGAACTGCTTTTGAAACTGTTAGGCTAGTGTCCCGGAGGGGTTGATGGAATTCCGAGTGTAGAGGTGAAATTCGCAGATATTCGGAGGAACACCAGTGGCGAAAGCGATCAACTGGACGGGTACAGACGCTGAGGCATGAAAGTGTGGGGATCAAACAGG
## 888                                                          ACGTAGGGTGCGAGCGTTAATCGGAATTACTGGGCGTAAAGCGTGCGCAGGCGGCTTTGTAAGACAGGTGTGAAATCCCCGGGCTTAACCTGGGAACTGCGCTTGTGACTGCAAGGCTCGAGTGCGGCAGAGGGGGGTGGAATTCCACGTGTAGCAGTGAAATGCGTAGAGATGTGGAGGAACACCGATGGCGAAGGCAGCCCCCTGGGTCGACACTGACGCTCATGCACGAAAGCGTGGGTAGCAAACAGG
## 890                                                          ACGGAAAGTGCGAGCGTTGTTCGGATTGACTGGGCGTAAAGAGCACGTAGGCGGGTCTGTAAGTCGATTGTGAAATCCCCGGGCTCAACCCGGGAACTGCAGTCGATACTGTGGATCTTGAATGTTCGAGGGGTTGGTGGAATTCCAGGTGTAGAAGTGAAATTCGTAGATATCTGGAGGAACACCGGAGGCGAAGGCGGCCGACTGGCGAAACATTGACGCTGAGGTGCGAAAGCGTGGGGAGCAAACAGG
## 891                                                          ACGTGAGGTGCAAACATTACCCGGAATCATTGGGCGTAAAGCGTCCGCAGGCGGCGCAGTAAGTCAGGTATTAAATCTTAAGGCTCAACTTTAAGGCTGTACTTGAAACTACTGTGCTAGAGTTTTGGAGAGGTGAGCGGAATTCTACATGTAGGGGTAAAATCCGTAGATATGTAGAGGAACACCAAAAGCGAAGGCAGCTCACTGGCCAAATACTGACGCTCATGGACGAAAGCGTGGGTAGCAAACGGG
## 892                                                          ACGAAGGGTGCAAGCGTTATTCGGAATGACTGGGTGTAAAGGGCGTGTAGGCGGTTGATTAAGTTAATGGTAAAAGGCCATAGCCTAACTGTGGTATAGCCGTTAAAACTAATCAGCTAGAGTTAGGAAGAGAGAAGTGGAATTCTCGGAGTAGCGGTAAAATGCGTAGATCTCGAGAGGAACACCGATGGCGAAGGCAGCTTCTTGGTCCTATACTGACGCTGAGGCGCGAAAGCGTGGGGAGCAAACAGG
## 895                                                         ACGTAGGCTCCAAGCGTTGTCCGGATTTATTGGGCGTAAAGAGCTCGTAGGCGGTTCAGCAAGTCGGGTGTGAAAACTATGGGCTTAACCCATAGCCTGCACCCGAAACTGCTGTGACTAGAGTTTGGTAGGGGAGCGGGGAATTCCTGGTGTAGCGGTGAAATGCGCAGATATCAGGAGGAACACCGGTGGCGAAGGCGCCGCTCTGGGCCAATACTGACGCTGAGGAGCGAAAGCGTGGGTAGCAAACAGG
## 896                                                           ACGTATGTCGCAAGCGTTATCCGGATTTATTGGGCGTAAAGCGCGTCTAGGCGGAAAAATAAGTCTGATGTTAAAATGCGGGGCTCAACTCCGTATTGCGTTGGAAACTGTTTTTCTAGAGTACTGGAGAGGTGGGCGGAACTACAAGTGTAGAGGTGAAATTCGTAGATATTTGTAGGAATGCCGATGGAGAAGTCAGCTCACTGGACAGATACTGACGCTAAAGCGCGAAAGCGTGGGGAGCAAACAGG
## 897                                                          ACGAAGGTCCCGAGCGTTGTTCGGAATCACTGGGCGTAAAGGGAGCGTAGGCGGCGCGGTAAGTCAGATGTGAAATCCCGGGGCTCAACCCCGGAACTGCATCCGATACTGCCGTGCTAGAGTAATGGAGAGGTAGCTGGAATTATCGGTGTAGCAGTGAAATGCGTAGATATCGATAGGAACACTCGTGGCGAAAGCGAGCTACTGGACATTTACTGACGCTGAGGCTCGAAGGCTAGGGGAGCGAAAGGG
## 898                                                          ACAGAGGGTGCAAGCGTTAATCGGAATTACTGGGCGTAAAGCGCGCGTAGGCGGTTCGTTAAGTCCGTTGTGAAATCCCTGGGCTCAACCTGGGAATGGCGATGGATACTGGCGAGCTAGAGTATGAGAGAGGTGGGTAGAATTCCAGGTGTAGCGGTGAAATGCGTAGATATCTGGAGGAATACCGGTGGCGAAGGCGGCCCACTGGCTTAATACTGACGCTGAGGTGCGAAAGCGTGGGGAGCAAACAGG
## 899                                                          ACGGAGGGGGCGAGCGTTGTTCGGAATCACTGGGCGTAAAGCGTGCGTAGGCGGTTATACAAGTCAGGAGTGAAATCCCAGGGCTCAACCCTGGAATTGCTTTTGAGACTGTATGGCTAGAGATCTGGAGAGGTTGGGGGAATTCCGAGTGTAGCAGTGAAATGCGCAGATATTCGGAGGAACACCAGTGGCGTAGGCGCCCAACTGGCCAGATACTGACGCTGAGGCACGAAAGCGTGGGGAGCAAACAGG
## 900                                                          ACAGAGGGTGCAAGCGTTGCTCGGAATTATTGGGCGTAAAGGGCAGGTAGGTGGTCTCGTTTGTCTGTGGTGAAATCCCTGGGCTCAACCCAGGAAGTGCCTCAGAAACGGCGGGACTAGAGTGCTAGAGAGGGTCGTGGAATTCCCGGTGTAGCGGTGAAATGCGTAGAGATCGGGAGGAACACCAGAGGCGAAGGCGGCGACCTGGATAGCAACTGACACTCAACTGCGAAAGCGTGGGGAGCAAACAGG
## 902                                                          ACGGGGGATGCAAGTGTTATCCGGAATCACTGGGCGTAAAGCGTCTGTAGGTGGTTAAATAAGTCAACTGTTAAATCTTGAGGCTCAACCTCAAAATCGCAGTCGAAACTGTTTGACTAGAGTATAGTAGGGGTAAAGGGAATTTCCAGTGGAGCGGTGAAATGCGTAGATATTGGAAAGAACACCGATGGCGAAGGCACTTTACTGGGCTATTACTAACACTGAGAGACGAAAGCTAGGGTAGCAAATGGG
## 903                                                         ACGTAGGATCCAAGCGTTATCCGGAATTACTGGGCGTAAAGCGTCTGTAGGCGGCTTTTTAAGTCGGGTGCGAAATCTTGCGGCTCAACCGTATAGACTGTGCTCGAAACTATGAAGCTAGAGGTAGGTAGAGGCAAGTGGAATTTCTGGTGTAGGAGTGACATCCGTAGATATCAGAAGGAACACCAATGGCGAAGGCAGCTTGCTGGGCCTTACCTGACGCTCAGAGACGAAAGCGTGGGGAGCGAACGGG
## 904                                                           ACGTAGGTGGCAAGCGTTGTCCGGAATTATTGGGCGTAAAGCGCGCGCAGGTGGGATCTTAAGTCCATTTTGAAAGTTCGGAGCTCAACTCCGTGATGGGATGGAAACTGGGATTCTTGAGTACCGGAGAGGAAAGCGGAATTCCCCGTGTAGCGGTGAAATGCGTAGATATGGGGAGGAACACCAGTGGCGAAGGCGGCTTTCTGGACGGTGTCTGACACTGAGGCGCGAAAGCCAGGGGAGCGAACGGG
## 905                                                          ACGTAGGTGGCAAGCGTTGTCCGGAATTACTGGGTGTAAAGGGCGTGTAGGCGGGAATGCAAGTCAGATGTGAAATTCCGGGGCTCAACCCCGGCGCTGCATCTGAAACTGTGTTTCTTGAGTGCTGGAGAGGAAAGCGGAATTCCTAGTGTAGCGGTGAAATGCGTAGATATTAGGAGGAACACCAGTGGCGAAGGCGGCTTTCTGGACAGTAACTGACGCTGAGGCGCGAAAGCGTGGGGAGCAAACAGG
## 906                                                          ACGAACCGTCCAAACGTTATTCGGAATCACTGGGCTTAAAGGGTGCGTAGGCGGCGCGGAAAGTTGGGTGTGAAATCCCTCGGCTCAACCGAGGAACTGCGCCCAAAACTACCGTGCTCGAGGGAGATAGAGGTGAGCGGAACTTAGGGTGGAGCGGTGAAATGCGTTGATATCCTAAGGAACACCGGTGGCGAAAGCGGCTCACTGGATCTCTTCTGACGCTGAGGCACGAAAGCTAGGGTAGCGAACGGG
## 908                                                          ACAGAGACTGCAAGCGTTATTCGGATTCACTGGGCGTAAAGGGTGCGCAGGCGGCAATGTGTGTGAGACGTGAAAGCCCGGAGCTTAACTCCGGAATTGCGCCTCAAACTACATTGCTAGAGCATTGGAGAGGGTAGCAGAATTCACGGTGTAGCAGTGAAATGCGTAGATATCGTGAGGAATACCAGAGGCGAAGGCGGCTACCTGGACAATTGCTGACGCTCAGGCACGAAAGCGTGGGGAGCAAAAGGG
## 909                                                          ACAGAGGGTGCGAGCGTTAATCGGAATTACTGGGCGTAAAGCGCGCGTAGGCGGTTTGTTAAGTCGGATGTGAAATCCCCGGGCTCAACCTGGGAACTGCATTCGATACTGGCAGACTAGAGTATGGGAGAGGATGGTAGAATTCCAGGTGTAGCGGTGAAATGCGTAGAGATCTGGAGGAATACCGATGGCGAAGGCAGCCATCTGGCCTAATACTGACGCTGAGGTGCGAAAGCATGGGGAGCAAACAGG
## 910                                                          ACGGAGGGTGCAAGCGTTGTTCGGAATCATTGGGCGTAAAGGGCGTGTAGGTGGTCTGCCAAGTCAGGTGTGAAATCCCGCGGCTCAACCGCGGAACTGCACTTGAAACTGGTGGACTCGAGTGTCGGAGGGGGCAGCGGAATTCCCGGTGTAGAGGTGAAATTCGTAGATATCGGGAGGAACACCAGTGGCGAAGGCGGCTGCCTGGCCGATGACTGACACTGAGACGCGAAAGCGTGGGGAGCAAACAGG
## 913                                                          ACAGGGGGTGCTAGCGTTGTTCGGAATTACTGGGCGTAAAGGGCGCGTAGGCGGTCTTGCAAGTCAGAGGTGAAATCCCGGAGCTTAACTCCGGAATTGCTTTTGATACTGCAAGGCTAGAGTATGTTAGAGGAAATCGGAATTCCTAGTGTAGAAGTGAAATTCGTAGATATTAGGAAGAACACCGGTGGCGAAGGCGGATTTCTGGGACATAACTGACGCTGAGGCGCGAAAGCGTGGGGATCAAACAGG
## 914                                                          ACGTAGGGTGCAAGCGTTATCCGGAATCATTGGGCGTAAAGCGTTCGTAGGTGGTTTAGAAAGTTTGTGGTTAAATCCGATCGCTCAACGATCGAATCGCTACAAAAACTCCTAAACTAGAGAATGATAGAGGCAACTGGAATTTCGCATGTAGGGGTAAAATCCGTTGATATGCGAAGGAACACCAAAAGCGAAGGCAGGTTGCTGGGTCATTTCTGACACTCAAGAACGAAAGCGTGGGGAGCAAACAGG
## 915                                                        ACGTACTCCCTGAGTGGTATCCACTGATATTGGGCCTAAAGCGTTCGTAGCCCGTGAAGAAAGTCCTTGGTGAAAGTGGCACGCTCAACGTGTCAAATGGCTAAGGATACTACTTCGCTTTGAGGTGATACGAGGTCAAGAGAATTGCAGGGGTAACGGTGAAATGTTGTGATCCTTGCAGGACTACCAGTAGCGAAGGCGCTTGACCAGGATCATCCTGACGGTGAGGAACGAAAGCTGGGGGAGCGACCCGG
## 916                                                          ACGTAGGGTGCAAGCGTTAATCGGAATTACTGGGCGTAAAGCGTGCGCAGGCGGATATGTAAGTTAGAAGTGAAATCCCCGGGCTCAACCTGGGAATGGCTTTTAAGACTGCGTATCTAGAGTTTGTCAGAGGGGGGTGGAATTCCAAGTGTAGCAGTGAAATGCGTAGAGATTTGGAGGAACACCAATGGCGAAGGCAACCCCCTGGGATAAAACTGACGCTCATGCACGAAAGCGTGGGTAGCAAACAGG
## 919                                                          ACGAGTGGTGCAAGCGTTATCCGGAATTATTGGGCGTAAAGGATGTGTAGGTGGCAATGTTAGTCGAGTGTTTAACCCATGGGCTCAACCTGTGGATCGCATTCGAAACGGCATAGCTAGAAGGTATTAGGGGTATGCGGAACTATATGTGTAGGGGTGAAATCCGTTGATATATATGGGAACACCAAAAGCGAAGGCAGCATACTGGAATATTCTTGACACTGAGACATGAAAGCGTGGGTAGCGAATGGG
## 920                                                          ACAGAGGGTGCGAGCGTTGTCCGGAATCACTGGGCGTAAAGGGCGCGTAGGTGGCTTGGTAAGCGTGTGGTGAAAGTCCGGGGCTCAACCCCGGATCTGCCGTGCGAACTGCTGAGCTTGAGCATTGTAGAGGCAGGTGGAATTCCGGGTGTAGCGGTGGAATGCGTAGAGATCCGGAAGAACACCAGTGGCGAAGGCGGCCTGCTGGGCAATAGCTGACACTGAGGCGCGACAGCGTGGGGAGCAAACAGG
## 921                                                          ACAGAGAGGGCAAGCGTTGTCCGGAATGACTGGGCGTAAAGAGCGCGTAGGCGGTTTTTTAAGTCGGATGTGAAATCCCTGGGCTTAACCTAGGAACTGCATCCGATACTGGAGAACTAGAGGACGAGAGAGGAAAGTGGAATGACGAGTGTAGAGGTGAAATTCGTAGATATTCGTCGGAACACCAGAGGCGAAGGCGGCTTTCTGGCTCGTACCTGACGCTGAGGCGCGAAAGCGTGGGGAGCAAACAGG
## 922                                                          ACGAAGGATCCAAGCGTTATCCGGATTCATTGGGTTTAAAGGGTGCGTAGGCGGGTTTGTAAGTCAGTGGTGAAAGCCGGTCGCTCAACGATCGAATTGCCATTGATACTGCGGACCTAGAATATAGATGATGTTGGCGGAATATGACATGTAGCGGTGAAATGCATAGATATGTCATAGAACACCTATTGCGAAGGCAGCTGACAAAACTATTATTGACGCTGATGCACGAAAGTGCGGGGATCAAACAGG
## 923                                                          ACGAATGGGGCAAGCGTTGTTCGGAATCACTGGGCTTAAAGCGTGCGTAGGCGGTCTTGCAAGTCAGAAGTGAAAGCCTGGGGCTCAACCCCAGAATTGCTTTTGAAACTGCATGACTAGAATACCGGAGAGGTTGGGAGAATTCCTAGTGTAGAGGTGAAATTCGCAGATATTAGGAGGAATACCAGTGGCGTAGGCGCCCAACTGGACGGTTATTGACGCTCAGGCACGAAAGCGTGGGGAGCAAACAGG
## 924                                                          ACGGAGGATGCGAGCGTTATCCGGATTTATTGGGTTTAAAGGGTGCGTAGGTGGTGATTTAAGTCAGCGGTGAAAGTTTGTGGCTCAACCATAAAATTGCCGTTGAAACTGGGTTACTTGAGTGTGTTTGAGGTAGGCGGAATGCGTGGTGTAGCGGTGAAATGCATAGATATCACGCAGAACTCCGATTGCGAAGGCAGCTTACTAAACCATAACTGACACTGAAGCACGAAAGCGTGGGGATCAAACAGG
## 925                                                          ACGAGGGATCCTAGCGTTGTTCGGAATCATTGGGCGTAAAGCGGGTGTAGGTGGCTCTGTAAGTCAAGTGTGAAAGCCCCGGGCTCAACCTGGGAAGTGCATTTGATACTGCGGAGCTTGAGTGCTAGAGAGGTCATTAGAATACCTGGTGTAGTGGTGAAATACGTAGATATCAGGTGGAATACCAGAGGCGAAGGCGGATGGCTGGCTAGACACTGACACTCAGACCCGAAAGCGTGGGGATCAAACAGG
## 927                                                          ACGGAGGATCCAAGCGTTATCCGGAATCATTGGGTTTAAAGGGTCCGTAGGCGGTTTTATAAGTCAGTGGTGAAATCTGGTCGCTCAACGATCAAACGGCCATTGATACTGTAGAACTTGAATTACTTGGAAGTAACTAGAATATGTAGTGTAGCGGTGAAATGCTTAGAGATTACATGGAATACCAATTGCGAAGGCAGGTTACTACGAGTATATTGACGCTGATGGACGAAAGCGTGGGGAGCGAACAGG
## 928                                                          ACGTAGACCACAAGCGTTATCCGGATTTATTGGGCGTAAAGGGTGTGTAGGAGGTTGTGTGCGTCTTTGGTTAAATCCCACTGCCTAACAGTGGACCTGCCAGAGATACGACACGGCTAGAGGAAGTTAGAGGTTCATGGAGCGCACGGTGTAGGGGTGAAATCCGTTGATATCGTGCGAAACACCAAAGGCGAAGGCAATGAACTGGGACTCTCCTGACTCTGAAACACGAAAGCGTGGGGAGCAAAAAGG
## 929                                                          ACGAGGGATCCTAGCGTTGTTCGGAATCATTGGGCGTAAAGCGGGTGTAGGTGGCTACATAAGTCAGATGTGAAAGCCTTGGGCTTAACCCAAGAAGTGCATTTGATACTGTGTAGCTTGAGTGTGGTAGAGGCTACTAGAATTCCTGGTGTAGTGGTGAAATACGTAGATATCAGGAGGAATACCGGAGGCGAAGGCGGGTAGCTGGGCCAACACTGACACTGAGACCCGAAAGCGTGGGGATCAAACAGG
## 930                                                          ACGGGAGTGGCAAGCGTTATCCGGAATTATTGGGCGTAAAGCGTCCGCAGGCGGCCTTTTAAGTCTGTTGTTAAAGCGTGGAGCTTAACTCCATTTCAGCAATGGAAACTGGAAGGCTTGAGTGTGGTAGGGGCAGAGGGAATTCCCGGTGTAGCGGTGAAATGCGTAGATATCGGGAAGAACACCAGTGGCGAAGGCGCTCTGCTGGGCCATAACTGACGCTCATGGACGAAAGCCAGGGGAGCGAAAGGG
## 931                                                          ACGTAGGGTGCAAGCGTTAATCGGAATTACTGGGCGTAAAGCGTGCGCAGGCGGTTATATAAGTCAGATGTGAAATCCCCGGGCTCAACCTGGGAACTGCATTTGAGACTGTATAGCTAGAGTACGGCAGAGGGGGATGGAATTCCGCGTGTAGCAGTGAAATGCGTAGATATGCGGAGGAACACCGATGGCGAAGGCAATCCCCTGGACCTGTACTGACGCTCATGCACGAAAGCGTGGGGAGCAAACAGG
## 932                                                          ACAGAGGTCTCAAGCGTTGTTCGGAATCACTGGGCGTAAAGCGTGCGTAGGCTGTTTCGTAAGTCGTGTGTGAAAGGCGCGGGCTCAACCCGCGGACGGCACATGATACTGCGAGACTAGAGTAATGGAGGGGGAACCGGAATTCTCGGTGTAGCAGTGAAATGCGTAGATATCGAGAGGAACACTCGTGGCGAAGGCGGGTTCCTGGACATTAACTGACGCTGAGGCACGAAGGCCAGGGGAGCGAAAGGG
## 933                                                          ACGGGAGTGGCAAGCGTTATCCGGAATTATTGGGCGTAAAGCGTCCGCAGGCGGTCTTACAAGTCTGTCGTTAAAGCGTGGAGCTTAACTCCATTTCGGCGATGGAAACTGTAAGACTAGAGTGTGGTAGGGGCAGAGGGAATTCCCGGTGTAGCGGTGAAATGCGTAGATATCGGGAAGAACACCAGTGGCGAAGGCGCTCTGCTGGGCCATAACTGACGCTCATGGACGAAAGCCAGGGGAGCGAAAGGG
## 934                                                          ACGAAGGGTGCAAGCGTTAATCGGAATTACTGGGCGTAAAGCGCGCGTAGGCGGTTTGATAAGTTGGATGTGAAAGCCCCGGGCTCAACCTGGGAATTGCATCCAAGACTGTCAGGCTAGAGTACGGGAGAGGGGGGTAGAATTCCACGTGTAGCGGTGAAATGCGTAGAGATGTGGAGGAATACCGGTGGCGAAGGCGGCCCCCTGGATCGATACTGACGCTGAGGTGCGAAAGCGTGGGGAGCAAACAGG
## 936                                                          ACGGAGGGTGCAAGCGTTATCCGGATTCACTGGGTTTAAAGGGTGCGTAGGCGGGCAGGTAAGTCAGTGGTGAAATCTTCGAGCTTAACTCGGAAACTGCCGTTGATACTATCTGTCTTGAATTTAGTGGAGGTAAGCGGAATATGTCATGTAGCGGTGAAATGCTTAGATATGACATAGAACACCAATTGCGAAGGCAGCTTGCTACACTATTATTGACGCTGAGGCACGAAAGCGTGGGGATCAAACAGG
## 937                                                          ACGAAGGTGGCAAGCGTTACTCGGAATTACTAGGCGTAAAGGGCAGGTAGGTGGTTTTGTAAGTCTGTTGTGAAAGCTCCTGGCTTAACTGGGAGAGGTCAACGGATACTGCAGGACTTGAGTATAGGAGAGGTTACTGGAATTCCCGGTGTAGCGGTGAAATGCGCAGATATCGGGAGGAACACCAATGGCGAAAGCAGGTAACTGGACTATTACTGACACTCAGCTGCGAAAGCTAGGGGAGCAAACAGG
## 938                                                          ACAGAGACCTCAAACGTTATCCGGATTCATTGGGCGTAAAGCGTCCGCAGGTGGTTTGATAAGTGAGTGGTTAAATCCCGGGGCTTAACTTCGGGGCTGCCGTTCATACTGTCAAGCTAGAGGATGGAAGAGGTAAGCGGAATTCTCGGTGTAGTCGTAATAAGCGCTGATATCGAGAAGAACACCAAAGGCGAAGGCAGCTTACTGGTACATTCCTGACACTCATGGACGAAAGCGTGGGGAGCGAATAGG
## 941                                                         ACATGTGGGGCGAGCGTTGTTCGGAATTACTGGGCATAAAGGGTGCGTAGGCGGTCTTGTACGTCTGGGGTGAAAACCTGCGGCCTAACCGTAGGACGGCCTTGGATACGGCTCGACTTGAGTGCGTGAGGGGTGATCGGAACGCGTGGTGTAGCGGTGAAATGCGTAGATATCACGCGGAAGGCCAACGGCGAAGGCAGATCACTGGTGCGCAACTGACGCTGAGTGCACGAAAGCGTGGGGAGCAAACAGG
## 942                                                          ACGAAGGGGGCAAGCGTTGTTCGGAATTACTGGGCGTAAAGGGCGCGTAGGCGGCTTATCAAGTCAGGCGTGAAATTCCCGGGCTCAACCTGGGGGCTGCGCTTGATACTGATGAGCTTGAATGCGGGAGAGGATAGTGGAATTCCCAGTGTAGAGGTGAAATTCGTAGATATTGGGAAGAACACCGGTGGCGAAGGCGGCTATCTGGCCCGTAATTGACGCTGAGGCGCGAAAGCGTGGGGAGCAAACAGG
## 943                                                          ACGGAGGGTGCAAGCGTTATCCGGAATCACTGGGTTTAAAGGGTGCGTAGGCGGGGGATTAAGTCAGTGGTGAAAGCCTGTAGCTTAACTACAGAATTGCCATTGATACTGATTTTCTTGAATTGGGTTGAGGTTAGCGGAATGTGACATGTAACGGTGAAATGTTTAGATATGTCATGGAACACCGATTGCGAAGGCAGCTAGCTGGGCCTTGATTGACGCTGAGGCACGAAAGCGTGGGTAGCGAACAGG
## 944                                                          ACGGAGGGTGCAAGCGTTAATCGGAATCACTGGGCGTAAAGCGCACGTAGGCGGCTTGGTAAGTCAGGGGTGAAATCCCACAGCCCAACTGTGGAACTGCCTTTGATACTGCCAGGCTTGAGTACCGGAGAGGGTGGCGGAATTCCAGGTGTAGGAGTGAAATCCGTAGATATCTGGAGGAACACCGGTGGCGAAGGCGGCCACCTGGACGGTAACTGACGCTGAGGTGCGAAAGCGTGGGTAGCAAACAGG
## 945                                                          ACGAAGGGTGCAAGCGTTGTTCGGAATTACTGGGCGTAAAGGGCGCGTAGGTTGGTTTGTAAGTCTAATGTGAAATCCCCGGGCTTAACCCGGGACGTGCATTGGAAACTGCAGATCTTGAATGCTTAAGAGGGTGGTGGAATTCCAGGAGTAGAAGTGAAATTCGTAGATATCTGGAGGAACACCTGAGGCGAAGGCGGCCACCTGGTAAGACATTGACGCTGAGGCGCGAAAGTGCGGGGAGCAAACAGG
## 946                                                         ACGGAGGATGCGAGCGTTATCCGGATTTATTGGGTTTAAAGGGTTCGTAGGCGGCTTTGTAAGTCAGCGGTGAAATACCAACGCTCAACGTTGGGGCTGCCGTTGATACTGCTGGGCTTGAGATGGATCGACGATGGCGGAATGTGACAAGTAGCGGTGAAATGCATAGATATGTCACAGAACACCGATAGCGAAGGCAGCTGTCGAGGTCATCATCTGACGCTGAGGAACGAAAGCGTGGGGATCAAACAGG
## 947                                                         ACAGAGGGTGCAAGCGTTAATCGGAATTACTGGGCGTAAAGGGCTTGTAGGTGGTTAGATAAGTTGGACGTGAAAACCCTGGGCTCAACCTAGGAATGGCGACCAATACTGTTTAGCTAGAGTATAGTAGAGGGTATAGGAATTTCCGGTGTAGCGGTGAAATGCGTAGATATCGGAAGGAACATCAGTGGCGAAGGCGTATACCTGGACTAAGAACTGACACTGAGGAGCGAAAGCGTGGGGATCAAACAGG
## 949                                                          ACGGAGGATTCAAGCGTTATCCGGATTTATTGGGTTTAAAGGGTGCGTAGGCGGTTTGATAAGTTAGAGGTGAAATTTCGGGGCTCAACCCTGAACGTGCCTCTAATACTGTTGAGCTAGAGAGTAGTTGCGGTAGGCGGAATGTATGGTGTAGCGGTGAAATGCTTAGAGATCATACAGAACACCGATTGCGAAGGCAGCTTACCAAACTATATCTGACGTTGAGGCACGAAAGCGTGGGGAGCAAACAGG
## 952                                                          ACGGAGGATGCGAGCGTTATCCGGATTCATTGGGTTTAAAGGGTGCGCAGGCGGATATATAAGTCGGAGGTGAAATTCTGATGCTCAACATCAGCCGTGCCTTCGATACTGTATATCTAGAGTCAAGTTGGAGTAGGCGGAATGTGTTGTGTAGCGGTGAAATGCTTAGATATAACACAGAACACCGATCGCGAAGGCAGCTTACTAAACTTTGACTGACGCTCATGCACGAAAGCGTGGGGATCAAACAGG
## 953                                                         ACGGAGGATGCGAGCGTTATCCGGATTTATTGGGTTTAAAGGGTTCGTAGGCGGCTGAATAAGTCAGCGGTGAAATACCAGAGCTTAACTCTGGGGCTGCCGTTGATACTGTTTAGCTAGAGTATGATCGACGTTGGCGGAATGTGACAAGTAGCGGTGAAATGCATAGATATGTCACAGAACACCGATAGCGAAGGCAGCTGACGAGGTCACATACTGACGCTGAGGAACGAAAGCGTGGGGATCAAACAGG
## 954                                                          ACGAAGGGTGCAAGCGTTGTTCGGAATCATTGGGCGTAAAGCGTGCGTAGGCGGCTTAGTAAGTCGTTTGTGAAATCCCAGGGCTTAACCTTGGAATTGCAATCGAAACTGCTTTGCTAGAGTTTAGTGGGGGATAGTGGAATTCCTAGTGTAGGGGTGAAATCCGTAGAGATTAGGGGGAACATCAGTGGCGAAAGCGACTATCTACGCTAATACTGACGCTGAGGTACGAAAGCGTGGGGAGCAAACAGG
## 955                                                          ACGAAGGTCCCAAGCGTTGTTCGGAATCATTGGGCGTAAAGCGGATGTAGGCGGTTCCATAAGTCAGAGGTGAAATCCCGGGGCTTAACCCCGGAAGTGCCTTTGAGACTGTGGAACTTGAATGTAGGAGAGGGTGCTAGAATTCCTGGTGTAGTGGTGAAATACGTAGAGATCAGGAGGAATTCCAGATGCGAAGGCGGGTGCCTGGCCTAACATTGACGCTGAGATCCGAAAGCGTGGGGATCAAACAGG
## 957                                                          ACGAAGGGTGCAAGCGTTACTCGGAATTACTGGGCGTAAAGCGTGCGTAGGCGGTTTGTTAAGTCTGATGTGAAATCCCCGGGCTCAACCTGGGACGTGCATTGGAAACTGGCAATCTAGAGTACGGTAGAGGGTGGTGGAATTCCCGGTGTAGCAGTGAAATGCGTAGAGATCGGGAGGAACACTCGTGGCGAAGGCGGCCACCTGGACCAGTACTGACGCTGAGGCACGAAAGCGTGGGGAGCAAACAGG
## 958                                                          ACGTAGGGTGCAAGCGTTAATCGGAATTACTGGGCGTAAAGCGTGCGCAGGCGGTTATATAAGTCAGATGTGAAATCCCCGGGCTCAACCTGGGAACTGCATTTGAGACTGTATAGCTAGAGTACGGTAGAGGGGGATGGAATTCCGCGTGTAGCAGTGAAATGCGTAGATATGCGGAGGAACACCGATGGCGAAGGCAGCCTCCTGGGCCAGTACTGACGCTCATGCACGAAAGCGTGGGGAGCAAACAGG
## 959                                                          ACGGAGGGTGCAAGCGTTATCCGGATTCACTGGGTTTAAAGGGTGCGTAGGCGGGCTAGTAAGTCAGTGGTGAAATCCCCGAGCTTAACTTGGGAACTGCCGTTGATACTATTAGTCTTGAATATCGTGGAGGTAAGCGGAATATGTCATGTAGCGGTGAAATGCTTAGATATGACATAGAACACCAATTGCGAAGGCAGCTTGCTACACGGTCATTGACACTGAGGCACGAAAGCGTGGGGAGCAAACAGG
## 960                                                          ACAGAGGGTGCGAGCGTTAATCGGAATTACTGGGCGTAAAGCGTGTGTAGGTGGTTAGATAAGTTAGATGTGAAATCCCTGGGCTTAACCTGGGCACTGCGTTTAAGACTGTTTAGCTAGAGTACTGTAGAGGATAGTGGAATTTCCAGTGTAGCGGTGAAATGCGTAGATATTGGAAGGAACACCAGTGGCGAAGGCGACTATCTGGACAGATACTGACACTGAGACACGAAAGCGTGGGGAGCAAACAGG
## 961                                                          ACGGAGGGTGCAAGCGTTATCCGGATTTACTGGGTTTAAAGGGTGTGTAGGCGGGCTATTAAGTCAGTGGTGAAATCTCCGAGCTTAACTTGGAAACTGCCATTGATACTATTAGTCTTGAATTTTGTTGAGGTAGGCGGAATAAGTCATGTAGCGGTGAAATGCATAGATATGACTTAGAACACCAATTGCGAAGGCAGCTTGCTAAACAAACATTGACGCTGAGGCACGAAAGCGTGGGGATCAAACAGG
## 962                                                          ACAGAGGGTGCAAGCGTTGCTCGGAATTATTGGGCGTAAAGGGCAGGTAGGTGGTCTCGTTTGTCCCGGGTGAAATCCCTGAGCTTAACTCAGGAAGTGCCCTGGAAACGGCGAGACTAGAGTTCTGGAGAAGGTCGTGGAATTCCCAGTGTAGCGGTGAAATGCGTAGAGATTGGGAGGAACACCAGAGGCGAAAGCGGCGACCTAGACAGATACTGACACTCAACTGCGAAAGCGTGGGGAGCAAACAGG
## 964                                                          ACGGAGGGAGCTAGCGTTGTTCGGAATTACTGGGCGTAAAGCGTGCGTAGGCGGTGACTCAAGTCAGAGGTGAAAGCCTGGAGCTCAACTCCAGAACTGCCTTTGAAACTAGGTCGCTAGAATCTGGGAGAGGTGAGTGGAATTCCGAGTGTAGAGGTGAAATTCGTAGATATTCGGAAGAACACCAGTGGCGAAGGCGGCTCACTGGACCAGAATTGACGCTGAGGCACGAAAGCGTGGGGAGCAAACAGG
## 966                                                          ACGAGGGGAGCTAGCGTTGTTCGGAATTACTGGGCGTAAAGCGCGCGTAGGCGGCTTTTTAAGTCAGAGGTGAAAGCCCGGGGCTCAACCCCGGAATAGCCTTTGAAACTGGAAAGCTAGAATCTTGGAGAGGTCAGTGGAATTCCGAGTGTAGAGGTGAAATTCGTAGATATTCGGAAGAACACCAGTGGCGAAGGCGACTGACTGGACAAGTATTGACGCTGAGGTGCGAAAGCGTGGGGAGCAAACAGG
## 969                                                          ACGAGTGCCCCGAGCGTTATCCGGAATTATTGGGCGTAAAGGGTGTGTAGGCGGTCGTGTTAGTCCTCCGTTAAATCTCTCGGCTCAACCGAGAACCAGCGGGGGAGACGGCACGACTCGAGGGTATGAGAGGTGCATGGAACTCACAGTGTAGGGGTGAAATCCGTTGATATTGTGGGGAACACCAAATGCGAAGGCAGTGCACTGGCATACTTCTGACGCTGAGACACGAAAGCGTGGGTCGCGAATGGG
## 972                                                          ACAGAGAGTGCAAGCGTTAATCGGAATTATTGGGCGTAAAGGGTGTGTAGGTTGTTAAAAAAGTTAGATGTGAAATCCCTGGGCGCAACCTAGGAATTGCATTTAATACTTTTTATCTAGAGTATTGTAGGGGGAAGTGGAATTTCCGGTGTAGCGGTGAAATGCGTAGATATCGGAAGGAACACCAGTGGCGAAGGCGACTTCCTGGACAAATACTGACACTGAGACACGAAAGCGTGGGGAGCAAACAGG
## 973                                                          ACGGAGGATCCAAGCGTTATCCGGAATCATTGGGTTTAAAGGGTCCGTAGGCGGTTTGGTAAGTCAGTGGTGAAAGCCCTTCGCTCAACGGAGGAACGGCCATTGATACTGCCAGACTTGAATTACTGGGAAGTAACTAGAATATGTAGTGTAGCGGTGAAATGCTTAGATATTACATGGAATACCAATTGCGAAGGCAGGTTACTACCGGTGTATTGACGCTGATGGACGAAAGCGTGGGGAGCGAACAGG
## 975                                                          ACGGGAGTGGCAAGCGTTATCCGGAATTATTGGGCGTAAAGCGTCCGCAGGCGGCCTTGTAAGTCTGTCGTCAAAGCGTGGAGCTTAACTCCATTTAAGCGATGGAAACTGCAAGGCTAGAGTGTGGTAGGGGCAGAGGGAATTCCCGGTGTAGCGGTGAAATGCGTAGATATCGGGAAGAACACCAGTGGCGAAGGCGCTCTGCTGGGCCATAACTGACGCTCATGGACGAAAGCCAGGGGAGCGAAAGGG
## 976                                                          ACGTAGGTGGCAAGCGTTATCCGGAATTATTGGGCGTAAAGCGCGCGTAGGCGGTTTTTTAAGTCTGATGTGAAAGCCCACGGCTCAACCGTGGAGGGTCATTGGAAACTGGAAAACTTGAGTGCAGAAGAGGAAAGTGGAATTCCATGTGTAGCGGTGAAATGCGCAGAGATATGGAGGAACACCAGTGGCGAAGGCGACTTTCTGGTCTGTAACTGACGCTGATGTGCGAAAGCGTGGGGATCAAACAGG
## 977                                                          ACATAGGGTGCAAGCGTTGTCCGGAATTATTGGGCGTAAAGAGCTCGTAGGTGGTTCGTCACGTCGGATGTGAAACTCTGGGGCTTAACCCCAGACCTGCATTCGATACGGGCGAGCTTGAGTATGGTAGGGGAGTCTGGAATTCCTGGTGTAGCGGTGGAATGCGCAGATATCAGGAGGAACACCAATGGCGAAGGCACTCTACTGGACCATAACTGACACTGAGAGACGACAGCTAGGGGAGCAAATGGG
## 978                                                          ACGTAGGGTGCAAGCGTTGTTCGGAATCATTGGGCGTAAAGCGCGTGTAGGCGGTTAGTTAAGTCAGGTGTGAAATCCCTCGGCTCAACCGAGGAAGTGCATTTGAAACTAGCTAGCTCGAAGATGGTAGAGGAAGGTGGAATTCCAAGTGTAGAGGTGAAATTCGTAGATATTTGGAGGAATACCTGTGGCGAAGGCGGCCTTCTGGGCCATTCTTGACGCTGAGACGCGAAAGCGTGGGTAGCAAACAGG
## 980                                                          ACGGAGGGTGCAAGCGTTATCCGGATTCACTGGGTTTAAAGGGAGCGTAGGCGGACTTGTAAGTCCGTGGTGAAAGCCCTCAGCTTAACTGAGGAACTGCCATGGATACTATAAGTCTTGAATGTCGTTGAGGTTTGCGGAATAGGTCATGTAGCGGTGAAATGCTTAGATATGACCTAGAACACCTATTGCGAAGGCAGCAGGCTAAACGAATATTGACGCTGAGGCTCGAAAGCGTGGGGATCAAACAGG
## 981                                                          ACGGAGGATGCAAGTGTTATCCGGAATCACTGGGCGTAAAGCGTCTGTAGGTGGTCAAATAAGTCAACTGTTAAATCTTGAGGCTCAACCTCAAAATCGCAGTCGAAACTATTAGACTAGAGTATAGTAGGGGTAAAGGGAATTTCCAGTGGAGCGGTGAAATGCGTAGAGATTGGAAAGAACACCGATGGCGAAGGCACTTTACTGGGCTATTACTGACACTCAGAGACGAAAGCTAGGGTAGCAAATGGG
## 982                                                         ACGGGGGCTCCAAGCGTTGTCCGGTATCATTGGGCGTAAAGAGTTATGTAGGCGGTTATAAAAGTTACGGGTTAAATATTTGGGCTCAACCCAAAATCTGCTTGTAATACTGTATGACTAGAGAAAATCAGGGGAAGGTAGAACGCATGGTGTAGCGGTAAAATGCTTTGATATCATGCGGAATACTAAAGGCGTAAGCAGCCTTCTGGGATTTTTCTGACGCTGAGATACGAAAGCGTGGGTAGCAAAACGG
## 983                                                          ACGAAGGGTGCAAGCGTTGTTCGGAATCACTGGGCGTAAAGAATGCGTAGGCGGCTTAACAAGTCGAGAGTGAAATCCCAGGGCTCAACCCTGGAACTGCTTTCGAAACTGTTTTGCTAGAGTCCCGGAGGGGTTAGCGGAATTCCGAGTGTAGAGGTGAAATTCGCAGATATTCGGAGGAACACCAGTGGCGAAGGCGGCTAACTGGACGGGAACTGACGCTGAGGCATGAAAGTGTGGGGATCAAACAGG
## 984                                                          ACGTAGGTGGCAAGCGTTGTCCGGATTTATTGGGCGTAAAGCGAGCGCAGGCGGTTCCTTAAGTCTGATGTGAAAGCCCACGGCTCAACCGTGGAAGGTCATTGGAAACTGGGGAACTTGAGTGCAGAAGAGGAGAGTGGAATTCCATGTGTAGCGGTGAAATGCGTAGATATATGGAGGAACACCAGTGGCGAAGGCGACTCTCTGGTCTGTAACTGACGCTGAGGCTCGAAAGCGTGGGGAGCAAACAGG
## 985                                                          ACGGAGGGTGCGAGCGTTAATCGGAATTACTGGGCGTAAAGCGCGCGTAGGTGGCTTGGCACGCCGGTTGTGAAAGCCCCGGGCTCAACCTGGGAACGGCATCCGGAACGGCCAGGCTAGAGTGCAGGAGAGGAAGGTAGAATTCCCGGTGTAGCGGTGAAATGCGTAGAGATCGGGAGGAATACCAGTGGCGAAGGCGGCCTTCTGGCCTGACACTGACACTGAGGTGCGAAAGCGTGGGTAGCAAACAGG
## 986                                                          ACGAAGGGTGCAAGCGTTGTTCGGATTTATTGGGCGTAAAGCGCGCGTAGGCGGACCTGCAAGTCAGATGTGAAATCTCGGGGCTCAACCCCGAAACTGCGTCTGAAACTACAGGTCTAGAATCTTGGAGGGGGAAGGGGAATTTCGCATGTAGGGGTAAAATCCGTAGAGATGCGAAGGAACACCAGAGGCGAAGGCGCCTTCCTGGACAAGTATTGACGCTGAGGCGCGAAAGCGTGGGGATCAAACAGG
## 988                                                          ACGGAGGGTGCAAGCGTTAATCGGAATTACTGGGCGTAAAGCGCACGCAGGCGGTCTGTTAAGTCAGATGTGAAATCCCCGGGCTCAACCTGGGAACTGCATTTGAAACTGGCAGGCTTGAGTCTCGTAGAGGGAGGTAGAATTCCAGGTGTAGCGGTGAAATGCGTAGAGATCTGGAGGAATACCGGTGGCGAAGGCGGCCTCCTGGACGAAGACTGACGCTCAGGTGCGAAAGCGTGGGGAGCAAACAGG
## 989                                                          ACGAAAGGTGCAAGCGTTACTCGGAATCACTGGGCGTAAAGCGCACGTAGGCGGACTCGTAGGTCAATTGTGAAATACCTGGGCTCAACCCGGGAACTGCAGTTGAAACCGCTTGTCTTGAATACCTGAGAGGGTGGGGGAATACCTGGTGGAGAAGTGAAATTCGTAGAGATCAGGTGGAACACCGGAGGCGAAGGCGCCTGCCTGGCAGTGTATTGACGCTGAGGTGCGAAAGTGCGGGGAGCAAACAGG
## 992                                                          ACGGAGGGTGCAAGCGTTATCCGGAATCATTGGGTTTAAAGGGTCCGCAGGCGGATTTATAAGTCAGTGGTGAAATCCTGCAGCTCAACTGCAGAACTGCCATTGATACTGTAAGTCTTGAATTTGATCGGAGTGGGCGGAATGTGTAGTGTAGCGGTGAAATGCATAGATATTACACAGAACACCGATAGCGAAGGCAGCTCACTAGGTTATAATTGACGCTCATGGACGAAAGCGTGGGGATCAAACAGG
## 993                                                          ACAGAGGTCCCAAGCGTTGTTCGGATTCACTGGGCGTAAAGGGTGCGTAGGCGGTCGGGTAAGTCTGACGTGAAATCTTCAAGCTCAACTTGGAAACTGCGTCGGATACTATTCGGCTAGAGGAATGGAGGGGAGACTGGAATACTTGGTGTAGCAGTGAAATGCGTAGATATCAAGTGGAACACCAGTGGCGAAGGCGAGTCTCTGGACATTTCCTGACGCTGAGGCACGAAAGCCAGGGGAGCAAACGGG
## 994                                                          ACGAACCGTACGAACGTTATTCGGAATTATTGGGCTTAAAGGGTGCGTAGGCTGTGCAGAAAGTTGGGTGTGAAAGCCCTCGGCTCAACCGAGGAATTGCGCCCAAAACTACTGTGCTCGAGGGAGACAGAGGTGAGCGGAACTCAAGGTGGAGCGGTGAAATGCGTTGATATCTTGAGGAACACCGGTGGCGAAAGCGGCTCACTGGGTCTCTACTGACGCTGAGGCACGAAAGCCAAGGTAGCAAACGGG
## 995                                                          ACGAAGGTGGCAAGCGTTACTCGGAATTACTAGGCGTAAAGGGCAGGTAGGTGGTTTGATAAGTCTGTTGTGTAAGCTCTCGGCTTAACCGGGAGAGGTCAACAGAAACTGTCAGGCTTGAGTATAGGAGAGGGTACTGGAATTCCCGGTGTAGCGGTGAAATGCGCAGAGATCGGGAGGAACACCAATGGCGAAAGCAGGTACCTGGACTATTACTGACACTAAGCTGCGAAAGCTAGGGGAGCAAACAGG
## 996                                                          ACGGAGGGTGCAAGCGTTAATCGGATTTATTGGGCGTAAAGGGCGCGTAGGCTGATGAATAAGTCGGGTGTGAAATCCTGGGGCTTAACCCCAGGGCTGCACCTGAAACTGTTTGTCTAGAGGGTAGGCGGAGAAAACGGAATTCCACAAGTAGCGGTGAAATGCGTAGATATGTGGAAGAACACCGGTGGCGAAGGCGGTTTTCTAGCTTACTCCTGACGCTGAGGCGCGAAAGTTAGGGGATCAAACAGG
## 998                                                          ACGAAGGACCCAAGCGTTATCCGGATTTATTGGGTTTAAAGGGTGCGTAGGCGGGTTGTTAAGTCAGTGGTGAAAGCCTAGAGCTTAACTCTAGAATTGCCATTGATACTGACGATCTTGAGGATAGTTGAGGTGGGCGGAATGTATCATGTAGCGGTGAAATGCTTAGAGATGATACAGAACACCGATTGCGAAGGCAGCTCACTAAGCTATTTCTGACGCTGAGGCACGAAAGCGTGGGGATCAAACAGG
## 999                                                          ACGTAGGGTGCAAGCGTTAATCGGAATTACTGGGCGTAAAGCGTGCGCAGGCGGTTATATAAGTCAGATGTGAAATCCCCAGGCTTAACCTGGGACGTGCATTTGAAACTGTGATTCTTGAGTATTGGAGAGGGTAGTGGAATTGTAGGTGTAGGAGTGACATCCGTAGAGATCTGCAGGAACATCAGAGGCGAAGGCGACTACCTGGCCGATTACTGACACTGAGGAACGAAAGCGTGGGGAGCAAACAGG
## 1000                                                         ACAGAGGGTGCAAGCGTTGTTCGGAATTACTGGGCGTAAAGGGCGTGTAGGCGGCTTAGCAAGTCAGATGTGAAATCCCGAGGCTTAACTTCGGAAGTGCATTTGAAACTGCTAGGCTAGAGTGATGTAGAGGTTGGTGGAATTCCCAGTGTAGAGGTGAAATTCGTAGATATTGGGAGGAACACCGGTGGCGAAGGCGGCCAACTGGGCATTAACTGACGCTGAGGCGCGAAAGCGTGGGGAGCAAACAGG
## 1001                                                         ACGAGTGGCCCTAGCGTTAGTCGGAATTACTGGGCTTAAAGGGTGCGTAGGCGGACCTGTAAGTGCTTTGTGAAAGCCCCTCGCTCAACGAGGGAACCGCAGCGCATACTGCAGGTCTTGAAGAAGTTAGAGGCCGGTGGAACGATAGGTGGAGCGGTGGAATGCGTAGAGATCTATCGGAACGCCAATGGTGAAGACAACCGGCTGGGACTTTCTTGACGCTGAGGCACGAAAGCGTGGGGAGCAAACAGG
## 1003                                                         ACGGAGGGTGCGAGCGTTATCCGGAATTACTGGGTTTAAAGGGTGCGTAGGCGGCTTTGTAAGTCAGGAGTGAAAGTTTGCGGCTCAACCGTAAAATTGCTTTTGATACTGCAGAGCTAGAATCAGGATGAGGTTAGCGGAATGTGGCATGTAGCGGTGAAATGCATAGATATGCCATAGAACACCAATTGCGAAGGCAGCTGGCTAGACCTGCATTGACGCTGAGGCACGAAAGCGTGGGGAGCGAACAGG
## 1005                                                         ACAGAGACCCCGAGCGTTATCCGGATTTATTGGGCGTAAAGGGCAGGTAGGCGGACATGTTAGTCGGATGTCAAATATCCCGGCTCAACCGGGAAAAGGCATTCGAAACGGCATGTCTCGAGAATGTGAGAGATCGGTGGAATTCACGGTGTAGTAGTGAAATGCGTTGATATCGTGAGGAACACCCAAGGCGAAGGCAACCGATTGGCACATTTCTGACGCTGAGCTGCGAAAGCGTGGGGAGCGAATGGG
## 1006                                                        ACGAGAGGAGCGAGCATTATTCGAAATGATTAGGCGTAAAGGGTTTGTAGGTTGTTTTTTAAGTTGAAAAAAAAATATGAAAGCTTAACTTTCTTTCTTTTTTCAAAACTGATAAACTAGAGTATAAATAGAGGATAATAGAATTCTTATTGTAAGGATAAAATCTTATAATAATAAGAGGAATTTCAAAGGCGAAGGCAATTATCTGGGTATATACTGACACTGAGGAACGAAAGCTTGGGTAGCGAACGGG
## 1007                                                         ACGTAGGGTGCTAGAGTTGTCCGGATTTACTGGGTGTAAAGGGTGCTCAGGTGGGTTTGTAAGTCAGAGGTGAAATCCCAAAGCTTAACTTTGGAGCTGCCTTTGATACTGCAAGTCTGGAGTTTGAGAGAGGGCAATGGAATATCTGGTGTAGCAGTGAAATGCGTAGATATCAGATAGAACACCAATGGCGAAGGCAGTTGCCTGGCTCAAAACTGACGCTAAAGCACGAAAGTGTGGGGAGCAAACAGG
## 1009                                                         ACGAGCTCCCCAAACGTTATTCGGTATCACTGGGCTTAAAGAGTTCGTAGGCGGCGAAGTAGGTGAGGTGTGAAAGCCCTCGGCTCAACCGAGGAACTGCGCTTCAAACCACTTTGCTAGAGGGAGATAGGGGTAAGCGGAACAGATGGTGGAGCGGTGAAATGCATTGATATCATCTGGAACACCGGTGGCGAAAGCGGCTTACTGGGTCTTTTCTGACGCTGAGGAACGAAAGCTAGGGTAGCGAACGGG
## 1010                                                         ACGAAGGTGGCGAGCGTTACTCGGAATTACTAGGCGTAAAGGGCAGGTAGGCGGTTTGATAAGTCTGTTGTGTCAGCTCCTGGCTTAACTGGGAGAGGTCAACGGAAACTATCAGGCTTGAGTGTAGGAGAGGATACTGGAATTCCTGGTGTAGCGGTGAAATGCGCAGAGATCAGGAGGAACACCAATGGCGAAAGCAAGTATCTGGACTATTACTGACGCTAAGCTGCGAAAGCTAGGGGAGCAAACAGG
## 1012                                                         ACGTAGGGTGCGAGCGTTAATCGGAATTACTGGGCGTAAAGCGTGCGCAGGCGGATTGTTAAGCAAGACGTGAAATCCCCGGGCTTAACCTGGGAATGGCGTTTTGAACTGGCAGTCTAGAGTGTGTCAGAGGGGGGTGGAATTCCACGTGTAGCAGTGAAATGCGTAGAGATGTGGAGGAACACCAATGGCGAAGGCAGCCCCCTGGGATAACACTGACGCTCATGTACGAAAGCGTGGGTAGCAAACAGG
## 1013                                                         ACAGAGGTCTCAAGCGTTGTTCGGAATCACTGGGCGTAAAGGGTGCGTAGGTGGCGTGGTAAGTCAGATGTGAAAGCCCGGGGCTCAACCCCGGAATTGCATCCGATACTACCATGCTGGAGTACTGAAGAGGTGACTAGAATTCTAGGTGTAGCAGTGAAATGCGTAGATATCTAGAGGAATACCAAAGGCGAAGGCAGGTCACTGGGCAGTTACTGACACTGAGGCACGAAGGCCAGGGTAGCGAACGGG
## 1014                                                        ACGTAGGTGGCGAGCGTTATCCGGATTTATTGGGTTTAAAGGGTGCGTAGGCGGCTTAATAAGTCGGTGGTTAAAAGTAGCAGCTTAACTGTTTTACATGCCATCGATACTGTTGAGCTTGAGATATTTGGAGGCAGCTAGAATTTCCGGTGTAGCGGTGAAATGCATAGATACCGGAAGGAATACCAATTGCGAAGGCAGGTTGCTACGAATAAACTGACGCTGATGCACGAAAGCGTGGGGATCAAACAGG
## 1015                                                         ACGTAGGGTGCGAGCGTTAATCGGAATGACTGGGCGTAAAGAGTGCGTAGGCGGTTACGCGAGCTTGATGTGAAATCCCCGGGCTTAACCTGGGAATGGCATGGAGGACAGCGTAGCTAGAGTCTGTCAGAGGTGGGTAGAATTCCCTGTGTAGCAGTGAAATGCGTAGAGATGGGGAGGAATACCGATGGCGAAGGCAGCCTGCTGGGATAAGACTGACGCTGAGGCACGAAGGCGTGGGGAGCAAACAGG
## 1016                                                         ACGGAGGGTGCAAGCGTTATCCGGAATCACTGGGTTTAAAGGGTGCGTAGGCGGTTGTATAAGTCAGTGGTGAAAGGCCGTAGCTTAACTATGGGATTGCCATTGATACTGTATGACTTGAATGAGGTTGAGGTTGGCGGAATGTGGCATGTAGCGGTGAAATGCTTAGATATGCCATGGAACATCGATTGCGAAGGCAGCTGACTGGACCTGAATTGACGCTGAGGCACGAAAGCGTGGGGAGCGAACAGG
## 1017                                                         ACGTAGGGTGCGAGCGTTAATCGGAATTACTGGGCGTAAAGCGTGCGCAGGCGGTTTCGTAAGACAGATGTGAAATCCCCGGGCTTAACCTGGGAACTGCATTTGTGACTGCGAGACTAGAGTGTAGCAGAGGGGGGTAGAATTCCACGTGTAGCAGTGAAATGCGTAGAGATGTGGAGGAATACCGATGGCGAAGGCAGCCCCCTGGGCTAACACTGACGCTCATGCACGAAAGCGTGGGGAGCAAACAGG
## 1018                                                         ACGAGTGCCCCAAGCGTTATCCGGAATTACTGGGCGTAAAGGGTGCGTAGGTGGCGTGGTTAGTCTCTTGTTAAAGCTCCCGGCTTAACCGGGAAAGTGCGAGAGATACGGCCAGGCTCGAGGTGGTGAGAGGTCTATGGAACTCATGGTGTAGGGGTGAAATCCGTTGATATCATGGGGAACACCAAATGCGAAGGCAATAGGCTGGCACCATACTGACACTGAAGCACGAAAGCGTGGGGATCAAACGGG
## 1019                                                         ACAGAGGGTGCAAGCGTTGTTCGGAATTATTGGGCGTAAAGCGCGTGTAGGCGGCTTTCTAAGTCTGGTGTGAAAGCCCGGGGCTTAACCTCGGAAGTGCACTGGATACTGGATCGCTCGAGGACGGGAGAGGGTAGTGGAATTCCTGGTGTACCGGTGAAATGGGTAGATATCAGGAAGAACATCGGTGGCGAAGGCGGCTACCTGGCCCGTATCTGACGCTGAGACGCGAAAGCGTGGGGAGCAAACAGG
## 1020                                                         ACGTAGGGTGCAAGCGTTAATCGGAATTACTGGGCGTAAAGCGTGCGCAGGCGGTTATATAAGACAGATGTGAAATCCCCGGGCTCAACCTGGGAACTGCATTTGTGACTGTATAGCTAGAGTACGGCAGAGGGGGATGGAATTCCGCGTGTAGCAGTGAAATGCGTAGATATGCGGAGGAACACCGATGGCGAAGGCAATCCCCTGGACCTGTACTGACGCTCATGCACGAAAGCGTGGGGAGCAAACAGG
## 1021                                                         ACGTAGGGTGCAAGCGTTAATCGGAATTACTGGGCGTAAAGGGTGCGCAGGCGGTTTTGTAAGTCAGATGTGAAATCCCCGGGCTTAACCTGGGAATTGCGTTTGAAACTACAAAACTGGAGTGTAGCAGAGGGAGGTGGAATTTCATGTGTAGCAGTGAAATGCGTAGAGATATGGAAGAACATCGATGGCGAAGGCAGCCTCCTGGGTTAACACTGACGCTCATGCACGAAAGCGTGGGGAGCAAACAGG
## 1022                                                         ACGAAGGGGGCTAGCGTTGCTCGGATTTACTGGGCGTAAAGGGCGCGTAGGCGGACATTTAAGTCAGGGGTGAAATCCCGGGGCTCAACCTCGGAACTGCCTTTGATACTGGGTGTCTTGAGTATGAGAGAGGTGTGTGGAACTCCGAGTGTAGAGGTGAAATTCGTAGATATTCGGAAGAACACCAGTGGCGAAGGCGACACACTGGCTCATTACTGACGCTGAGGCGCGAAAGCGTGGGGAGCAAACAGG
## 1024                                                         ACGTAGGGGGCAAGCGTTATCCGGATTTACTGGGTGTAAAGGGAGCGCAGACGGAGAGGTAAGTCAGATGTGGAAGCCCCGGGCATAACCCGGGGAGTGCATTTGAAACTATCACTCTTGAGTACTGGAGAGGTAAGCGGAATTCCTGGTGTAGCGGTGAAATGCGTAGATATCAGGAGGAACACCGGTGGCGAAGGCGGCTTACTGGACAGAAACTGACGTTGAGGCTCGAAAGCGTGGGTAGCAAACAGG
## 1026                                                         ACGGAGGGTGCAAGCGTTGTTCGGAATTACTGGGCGTAAAGTGTGTCTAGGTGGGCATGTAAGTCAGATGTGAAATTACCCGGCTCAACCGGGGAGGGTCATTTGATACTGCATGTCTTGAGTCCCGAAGAGGTGGGTGGTATTCCCAGTGTAGAGGTGAAATTCGTAGATATTGGGAGGAACACCGGAGGCGAAAGCGACTCACTGGTCGGGCACTGACACTAAAGCACGAAAGCGTGGGTAGCAAACAGG
## 1027                                                          ACGGGAGGGGCAAGCGTTATCTGAAATAACTGGGCGTAAAGAGTCCGTAGACTGTAAAGTAAGTTACTTGTTAAATTTTTAAGCTTAACTTTAAGTTAGCTTGTAATACTGCTTTACTTGAGTTTTATACAGAAAAGTAGAATTTTATACGAAAGGGTAAAATCTTAAGATATATAAAGGAATACCAGCAGCGAAGGCGACTTTTTAGTAAAAACTGACGTTGAGGGACGAAAGTGTGGGTAGCAAACAGG
## 1028                                                         ACGGAGGGGGCTAGCGTTGTTCGGAATTACTGGGCGTAAAGCGCACGTAGGCGGATCAGAAAGTCAGAGGTGAAATCCCGGGGCTCAACCCCGGAACTGCCTTTGAAACTCCTGATCTTGAGGTCGAGAGAGGTGAGTGGAATTCCGAGTGTAGAGGTGAAATTCGTAGATATTCGGAGGAACACCAGTGGCGAAGGCGGCTCACTGGCTCGATACTGACGCTGAGGTGCGAAAGCGTGGGGAGCAAACAGG
## 1029                                                         ACAGAGGCCTCAAGCGTTGTTCGGAATTACTGGGCGTAAAGGGAATGTAGGTGGCGATGTAAGTCAGATGTGAAAGCCCTGGGCTTAACCTGGGAACGGCATTTGATACTGCATTGCTCGAGTGTGGAAGAGGAGAGTGGAATTCCCGGTGTAGGAGTGAAATCCGTAGATATCGGGAGGAATTCCGATGGCGAAGGCAACTCTCTGGTCCATTACTGACACTGAGATTCGAAAGCATGGGTAGCAAACAGG
## 1030                                                         ACGAAGGTGGCTAGCGTTGTTCGGATTTATTGGGCGTAAAGGGTCCGCAGGCGGTTCGGTAAGTCAGATGTGAAATCCCACAGCTTAACTGTGGACACGCATTTGAAACTGCCAGACTAGAGTATGAGAGGGGTAAGAGGAATTTCCAGTGTAGGGGTGAAATCCGTAGATATTGGAAAGAACACCGGCCGCGAAGGCGTCTTACTGGCTCATTACTGACGCTCAGGGACGAAAGCCAGGGGAGCAAACGGG
## 1031                                                        ACGAGGACCCCGAGCGTTATCCGGATTTATTGGGCGTAAAGAGTGCGCAGGAGGTCTGATGCGTCTTGTGTTAAATGTCACTGCTTAACAGTGTCGCCGCATGAGATACGGTCAGACTAGAGCATGTTAGAGGTTACTAGAACTTATGGTGTAGGGGTGAAATCCGTTGATATCATAGGGAATACCAAAGGCGAAGGCATGTAACTGGAACATTGCTGACTCTCATTGCACGAAAGCGTGGGGAGCAAAAAGG
## 1033                                                         ACGTAGGGTGCGAGCGTTAATCGGAATTACTGGGCGTAAAGCGTGCGCAGGCGGTTTTGTAAGACAGATGTGAAATCCCCGGGCTTAACCTGGGAACTGCGTTTGTGACTGCAAGACTCGAGTGTGGCAGAGGGGGGTGGAATTCCACGTGTAGCAGTGAAATGCGTAGAGATGTGGAGGAACACCGATGGCGAAGGCAGCCCCCTGGGTCAACACTGACGCTCATGCACGAAAGCGTGGGTAGCAAACAGG
## 1034                                                         ACGGAGGATCCAAGCGTTATCCGGATTTATTGGGTTTAAAGGGTGCGTAGGCGGCCTATTAAGTCAGGGGTGAAAGACGGTGGCTCAACCATCGCAGTGCCCTTGATACTGATGGGCTTGATTACACTTGAGGTAGGCGGAATGTGACAAGTAGCGGTGAAATGCATAGATATGTCACAGAACACCAATTGCGAAGGCAGCTTACTAAGGTGTTAATGACGCTGAGGCACGAAAGCGTGGGGATCAAACAGG
## 1035                                                         ACGAGTGGCCCAAGCGTTATCCGGAATTATTGGGCGTAAAGGATGTGTAGGTGGTTATGTTAGTCTAGTGTCTAATCTCTCTGCTCAACAGGGAGCCAGCACAGGAAACGGCATGACTAGAGTATGTTAGGGGTAAGCGGAACTCATAGTGTAGGGGTGAAATCCGTTGATATTATGGGGAACACCAAAAGCGAAGGCAGCTTACTGGAACATTACTGACACTGAAACATGAAAGCGTGGGTAGCGAATGGG
## 1038                                                         ACAGGGGGTGCAAGCGTTGTTCGGAATTACTGGGCGTAAAGGGAGCGTAGGCGGGATTACAAGACAGGGGTTAAATCTCCGGGCTCAACCTGGAACTTGCCTTTGTGACTGTAGTCCTTGAATATGGTAGAGGTCAGTGGAATTTCCGGTGTAGCGGTGGAATGCGTAGAGATCGGAAAGAACACCAGAGGCGAAGGCGACTGACTGGACCAATATTGACGCTGAGGCTCGAAAGCGTGGGGAGCAAACAGG
## 1041                                                         ACGAGTGCCTCAAGCGTTATCCGGAATTATTGGGCGTAAAGGTTGTGTAGGTGGTAATGTTAGTCTCTTGTAAAATTCTTCGGCTTAACCGGGGGCTCGCAAGGGAAACGGCATTACTAGAGGATGCGAGGGGTTTGCGGAACTCATGGTGTAGCGGTGAAATGCGTTGATATCATGGGGAACACCAAAAGCGAAGGCAGCAAACTGGAGCACTCCTGACACTGAAACAAGAAAGCGTGGGTCGCGAATGGG
## 1042                                                         ACGTAGGGTGCAAGCGTTAATCGGAATTACTGGGCGTAAAGCGTGCGCAGGCGGCTTTGCAAGACAGATGTGAAATCCCCGGGCTCAACCTGGGAACTGCGTTCGAAACTGCAAGGCTAGAGTGTGTCAGAGGGGGGTAGAATTCCACGTGTAGCAGTGAAATGCGTAGAGATGTGGAGGAATACCAATGGCGAAGGCAGCCCCCTGGGATAACACTGACGCTCATGCACGAAAGCGTGGGGAGCAAACAGG
## 1043                                                         ACGGAGGGGGCTAGCGTTGTTCGGAATTACTGGGCGTAAAGCGCACGTAGGCGGACCTGAAAGTCAGAGGTGAAATCCCGGGGCTCAACCCCGGAACGGCCTTTGAAACTCCTGGTCTTGAGGTCGAGAGAGGTGAGTGGAATTCCGAGTGTAGAGGTGAAATTCGTAGATATTCGGAGGAACACCAGTGGCGAAGGCGGCTCACTGGCTCGATACTGACGCTGAGGTGCGAAAGCGTGGGGAGCAAACAGG
## 1044                                                         ACGTAGGGTGCAAGCGTTATCCGGAATTATTGGGCGTAAAGAGCTCGTAGGCGGTTTGTCGCGTCTGCTGTGAAAATCTGAGGCTCAACCTCGGACTTGCAGTGGGTACGGGCAGACTAGAGTGCGGTAGGGGAGAAGGGAATTCCTGGTGTAGCGGTGGAATGCGCAGATATCAGGAGGAACACCAATGGCGAAGGCACTTCTCTGGGCCGCTACTGACGCTGAGGAGCGAAAGCGTGGGGAGCGAACAGG
## 1045                                                         ACGAGTGCCTCAAGCGTTATCCGGAATCATTGGGCGTAAAGGTTGTGTAGGTGGTTATATTAGTCTTTTGTTAAATTCTTCGGCTTAACCGGGGGCATGCAGAGGAAACGGTATAACTAGAGGATGCGAGGGGTTTCTGGAACTCATAGTGTAGCGGTAAAATGCGTTGATATTATGGGGAACACCAAAAGCGAAGGCATGAAACTAGAGCACTCCTGACACTGAAACAAGAAAGCGTGGGTCGCGAATGGG
## 1047                                                         ACAGAGGGTGCAAGCGTTGTTCGGAATTATTGGGCGTAAAGCGCGTGTAGGCGGCTTGGCAAGTCAGATGTGAAAGCCCATGGCTTAACCATGGAAGTGCGTCTGAAACTGCTGAGCTTGAGTGCCGGAGAGGATGGCGGAATTCCGCAAGTAGAGGTGAAATTCGTAGATATGCGGAGGAACACCGGTGGCGAAGGCGGCCATCTGGACGGCAACTGACGCTGAGACGCGAAAGCGTGGGGAGCAAACAGG
## 1048                                                         ACGGAGGATGCAAGCGTTATCCGGAATCATTGGGTTTAAAGGGTCCGTAGGCGGTTTAATAAGTCAGTGGTGAAAGCCCATCGCTCAACGATGGAACGGCCATTGATACTGTTAAACTTGAATTATTAGGAAGTAACTAGAATATGTAGTGTAGCGGTGAAATGCTTAGAGATTACATGGAATACCAATTGCGAAGGCAGGTTACTACTAATATATTGACGCTGATGGACGAAAGCGTGGGGAGCGAACAGG
## 1049                                                          ACGTAGGGGGCGAGCGTTATCCGGATTTATTGGGCATAAAGAGCATGTAGGCGGTAATTTAAGTCAGGCGTGAAATATTGTGGCCTAACCATAATATGCATTTGATACTGGATTACTAGAGATCGGGAGAGGAAAGTGGAACTCATGGTGTAGCGGTGAAATGCGTAGATATCATGAGGAACACCAGTGGCGAAGGCGGCTTTCTGGACCGATTCTGACGCTGAGATGCGAAAGCGTGGGGAGCAAACAGG
## 1050                                                         ACAGAGGGTGCAAGCGTTAATCAGAATGACTGGGCGTAAAGGGCGTGTAGGTGGTTGACTAGGTTTGATGTGAAATCCCCGGGCTTAACCTGGGAATTGCGTCGAAAACGGGTCGACTCGAGTGAGATAGAGGGTTGTGGAATTTCCGGTGTAGCGGTGAAATGCGTAGATATCGGAAAGAACATCAGTGGCGAAGGCGGCAACCTGGATCTTAACTGACACTGAGGCGCGAAGGCGTGGGGAGCAAACAGG
## 1051                                                         ACGAAGGTCCCTAGCGTTGTTCGGAATCATTGGGCGTAAAGCGCACGTAGGTGGCTTTGCAAGTCAGGTGTGAAAGCCTAGGGCTTAACTCTAGAAGTGCACTTGAAACTGCTTAGCTTGAGTACGGGAGAGGTGAGTGGAATTCCTGGTGTAGTGGTGAAATACGTAGATATCAGGAGGAACACCGGTGGCGAAGGCGGCTCACTGGCCCAGTACTGACACTGAGGTGCGAAAGCATGGGGATCAAACAGG
## 1052                                                         ACAGAGGGTGCAAGCGTTGTTCGGAATTATTGGGCGTAAAGCGCGTGTAGGCGGCTTGGCAAGTCAGATGTGAAAGCCCTCGGCTTAACCGAGGAAGTGCGTCTGAAACTGCTGAGCTTGAGTCTCGGAGAGGATGGCGGAATTCCGCAAGTAGAGGTGAAATTCGTAGATATGCGGAGGAACACCGGTGGCGAAGGCGGCCATCTGGACGAGTACTGACGCTGAGACGCGAAAGCGTGGGGAGCAAACAGG
## 1053                                                         ACGGAGGGTGCAAGCGTTATCCGGATTTATTGGGTTTAAAGGGTCCGTAGGCGGACGATTAAGTCAGCGGTGAAAGCCCACAGCTCAACTGTGGAACTGCCATTGATACTGGTTGTCTTGAGTACGGTTGATGTTGGCGGAATAAGTCATGTAGCGGTGAAATGCATAGATATGACTTGGAACCCCGATTGCGAAGGCAGCCAACAAAGCCGAAACTGACGCTGAGGGACGAAAGCGTGGGTAGCGAACAGG
## 1055                                                        ACAGAGACCCCAAGCGTTATCCGGAATTATTGGGCGTAAAGGGTGAGTAGGTGGTATTATTAGTCGGGTGTTAAAGTTCTCGGGCTCAACCTGAGAAATGCATCCGAAACGGTAAAACTAGAATATTGGAGAGGTGAGTGGAACTCTATGTGTAGGGGTAAAATCCGTTGATATATAGGGGAACACCAAAAGCGAAGGCAGCTCACTGGCCAATTATTGACACTGAATCACGAAAGCGTGGGTAGCGATACGG
## 1056                                                         ACGGAGGGTGCGAGCGTTGTCCGGAATCACTGGGCGTAAAGGGCGCGTAGGTGGTCTGCTAAGCGTGCGGTGAAAGCCTGGGGCTCAACCCCAGGTCGGCCGTGCGAACTGGTGGACTGGAGCACTGTAGAGGCAGGTGGAATTCCGGGTGTAGCGGTGGAATGCGTAGAGATCCGGAAGAACACCGGTGGCGAAGGCGGCCTGCTGGGCAGTTGCTGACACTGAGGCGCGACAGCGTGGGGAGCAAACAGG
## 1057                                                         ACGTAGGGTCCAAGCGTTAATCGGAATTACTGGGCGTAAAGCGTCCGCAGGTGGCTTGATAAGATAGACGTGAAATCCCTGGGCTCAACCTAGGAATTGCGTTTATGACTGTCTCGCTAGAGTATGGGAGAGGGGGGTGGAATTCCACGTGTAGCAGTGAAATGCGTAGAGATGTGGAGGAACACCGATGGCGAAGGCAACCCCCTGGCCTAATACTGACACTCATGGACGAAAGCGTGGGGAGCAAACAGG
## 1058                                                         ACGAGGGGTGCAAGCGTTGCTCGGAATTATTGGGCGTAAAGGGCAGGTAGGTGGTCTTATTTGTCAGGGGTGAAAGCCTTGGGCTTAACCTGAGAAGTGCCCCTGAAACGGTAAGACTAGAGTCTTGGAGAGGGTTGCGGAATTCCCGGTGTAGCGGTGAAATGCGTAGAGATCGGGAGGAACACCAGAGGCGAAAGCGGCAGCCTGGACAAGCACTGACACTCAACTGCGAAAGCGTGGGGAGCAAACAGG
## 1059                                                        ACAGGGGATGCAAGCGTTATCCGGAATTATTGGGCGTAAAGCGTCCTGAGGTGGCTTACCCAGTCGACTGTGAAAGCTCAAGGCTTAACCTTGAAACGGCAGTCGAAACTTGTAAGCTTGAGTACGGTAGGGGCAGAGGGAATTCCCGGAGGAGAGGTGAAATTCGTAGATATCGGGAGGAAGGCCAAAGGCGAAAGCACTCTGCTGGGCCGATTACTGACACTGACAGACGAAAGCTAGGGGAGCGAAAGGG
## 1060                                                         ACGAAGGGTGCAAGCGTTGCTCGGAATTATTGGGCGTAAAGGGCAAGTAGGTGGTCATATAAGTCTGTGGTGAAATCCCTGAGCTCAACTCAGGACGTGCCATGGAAACTATATGACTAGAGTATCGGAGAGGTTCGTGGAATTCCCGGTGTAGCGGTGAAATGCGTAGAGATCGGGAGGAACACCAGAGGCGAAAGCGACGAACTGGACGATGACTGACACTCAATTGCGAAAGCGTGGGTAGCAAACAGG
## 1062                                                         ACGAAGGGGGCTAGCGTTGTTCGGAATTACTGGGCGTAAAGCGCACGCAGGCGGTTCGGTCAGTCAGAAGTGAAAGCCCCGGGCTTAACCTGGGAACTGCTTTTGATACTGCCGAGCTTGAATCACGGAGAGGGTAGTGGAATTCCGAGTGTAGAGGTGAAATTCGTAGATATTCGGAAGAACACCAGTGGCGAAGGCGACTACCTGGCCGTCGATTGACGCTCATGTGCGAAAGCGTGGGGAGCAAACAGG
## 1063                                                         ACGAGGGATCCTAGCGTTGTTCGGAATCATTGGGCGTAAAGCGGGTGTAGGCGGATTTCTAAGTCAGGTGTGAAATCCCAGGGCTCAACCCTGGAAGTGCATTTGATACTGGAAATCTTGAGTGTGGTAGAGGCTACTGGAATTGTTGGTGTAGTGGTGAAATACGTAGATATCAACAGGAATACCGGAGGCGAAGGCGAGTAGCTGGGCCAACACTGACGCTGAGACCCGAAAGCGTGGGGATCAAACAGG
## 1064                                                         ACGGAGGGTGCAAGCGTTGTTCGGAATTACTGGGCGTAAAGCGCGTGCAGGTGGTTTTTTAAGTCTCGTGTGAAATCCCTCGGCTTAACCGAGGAACTGCGCGAGATACTGAGAGACTTGAATACGGGAGAAGAGGGCGGAATTCCCAGTGTAGAGGTGAAATTCGTAGATATTGGGAGGAACACCGGCGGCGAAAGCGGCTCTCTAGACCGATATTGACGCTCATACGCGAGAGCGTGGGTAGCAAACAGG
## 1065                                                         ACGAGTGCTCCAAGCGTTATCCGGAATTATTGGGCGTAAAGGGTGTGTAGGCGGTTTTGTTAGTCGGTTGTTAAAACTCCTCGCTTAACGGGGAAGGTGCAATCGAAACGGCAAATCTAGAGGATGCGAAAGGTAAACAGAACTCATAGTGTAGGGGTGAAATCCGTTGATATTATGGGGAATACCAAATGCGAAGGCAGTTTACTAGCGCACTCCTGACGTTGAAACACGAAAGCGTGGGTAGCGAATGGG
## 1066                                                         ACGAAGGGGGCTAGCGTTGTTCGGAATTACTGGGCGTAAAGCGCACGTAGGCTGTGCGATAAGTCAGGGGTGAAATCCCGGAGCTCAACTCCGGAACTGCCTTTGAAACTATCGTGCTAGAATCTCAAAGGGGATAGCGGAATTCCAAATGTAGGGGTGAAATCCGTAGATATTTGGAGGAACACCGGTGGCGAAGGCGGCTATCTGGGTGAGTATTGACGCTGAGGTGCGAAAGCGTGGGGATCAAACAGG
## 1068                                                         ACGAAGGGGGCTAGCGTTGTTCGGAATTACTGGGCGTAAAGCGCACGTAGGCTGACATTTAAGTCAGGGGTGAAATCCCGAGGCTCAACCTCGGAACTGCCCTTGATACTGGGTGTCTCGAGTCCGGAAGAGGTTAGTGGAATTCCCAGTGTAGAGGTGAAATTCGTAGATATTGGGAAGAACACCAGTGGCGAAGGCGGCTAACTGGTCCGGTACTGACGCTGAGGTGCGAAAGCGTGGGGAGCAAACAGG
## 1069                                                          ACGTATGTCGCAAGCGTTATCCGGAATTATTGGGCATAAAGCGCGTCTAGGCGGTTTACTAAGTCTGATGTGAAAATGCGGAGCTCAACTCCGTATTGCGTTGGAAACTGGTAGACTAGAGTATTGGAGAGGTAGGCGGAACTACAAGTGTAGAGGTGAAATTCGTAGATATTTGTAGGAATGCCAATGACGAAGGTAGCTTACTGGACAAATACTGACGCTGAAGCGCGAAAGCTAGGGGAGCAAACAGG
## 1070                                                         ACGTAGGGTGCGAGCGTTAATCGGAATTACTGGGCGTAAAGCGTGCGCAGGCGGTTCGTTAAGTTTGCGGTGAAAGCCCCGGGCTCAACCTGGGAATGGCCGTGGAAACTGGCGGGCTGGAGTGCGGCAGAGGGGGGTGGAATTCCGCGTGTAGCAGTGAAATGCGTAGAGATGCGGAGGAACACCGATGGCGAAGGCAGCCCCCTGGGCCGACACTGACGCTCAGGCACGAAAGCGTGGGGAGCAAACAGG
## 1071                                                         ACAGAGGGAGCGAGCGTTAACCGGAATGATTGGGCGTAAAGCGTCTGTAGGTGGCTTTTTAAGTCTACTGTCAAAACCCAGAGCTTAACTTTGGATCGGCAGTGGAAACTGAAAAGCTTGAGTGCGGTAGAGGCAGAGGGAATTCCTAGTGGAGCAGTGAAATGCGTAGAGATTAGGAAGAACACCCGTGGCGAAGGCGCTCTGCTGGGCCGTAACTGACACTGAGAGACGAAAGCTAGGGGAGCAAATGGG
## 1072                                                         ACGGAGGATCCGAGCGTTATCCGGATTTATTGGGTTTAAAGGGAGCGTAGGCGGGTTGTTAAGTCAGTTGTGAAAGTTTGCGGCTCAACCGTAAAATTGCAGTTGATACTGGCGACCTTGAGTGCAACAGAGGTAGGCGGAATTCGTGGTGTAGCGGTGAAATGCTTAGATATCACGAAGAACTCCGATTGCGAAGGCAGCTTACTGGATTGTAACTGACGCTGATGCTCGAAAGTGTGGGTATCAAACAGG
## 1073                                                         ACGAGGGGTCCTAGCGTTGTTCGGAATCATTGGGCGTAAAGCGTATGTAGGTGGCCTTATAAGTCAGGTGTGAAAGCCCCGGGCTCAACCCGGGAAGTGCATTTGATACTGTTTGGCTTGAGTACCGAAGAGGATAGTGGAATTCCAGGTGTAGTGGTGAAATACGTAGATATCTGGAGGAACACCGGTGGCGAAGGCGGCTATCTGGTCGAGTACTGACACTGAGATACGAAAGCGTGGGGATCAAACAGG
## 1075                                                         ACGGAGGGTGCAAGCGTTAATCGGAATTACTGGGCGTAAAGCGCACGTAGGCGGTTTTCTAAGTCAGATGTGAAATCCCCGGGCTTAACCTGGGAACTGCACCTGATACTGGAAGACTAGAATGTGGGAGAGGAGAGTGGAATTTCCGGTGTAGCGGTGAAATGCATAGAGATCGGAAGGAACACCAGTGGCGAAGGCGGCTCTCTGGACCAACATTGACGCTGAGGTGCGAAAGCGTGGGTAGCAAACAGG
## 1076                                                         ACGGAGGGTGCAAGCGTTATCCGGATTCACTGGGTTTAAAGGGAGCGTAGGTGGGCAGGTAAGTCAGTGGTGAAATCTCCGGGCTTAACCCGGAAACTGCCATTGATACTATTTGTCTTGAATATTCTGGAGGTAAGCGGAATATGTCATGTAGCGGTGAAATGCTTAGATATGACATAGAACACCCATTGCGAAGGCAGCTTACTACGGAATTATTGACACTGAGGCTCGAAAGCGTGGGGATCAAACAGG
## 1077                                                         ACGGAGGATCCGAGCGTTATCCGGATTTATTGGGTTTAAAGGGAGCGTAGGCGGATGTTTAAGTCAGTTGTGAAAGTTTAAGGCTCAACCTTGAAATTGCAGTTGATACTGGACGTCTTGAGTGCATTAAATGTGGGCGGAATTCGTGGTGTAGCGGTGAAATGCTTAGATATCACGAAGAACTCCAATTGCGAAGGCAGCTCACAGGACTGTAACTGACGCTGATGCTCGAAAGTGTGGGTATCAAACAGG
## 1080                                                         ACGGAGGATCCAAGCGTTATCCGGATTTATTGGGTTTAAAGGGTGCGTAGGCGGGAGATTAAGTCAGTGGTGAAAGTTTGCAGCTCAACTGTAAAATTGCCATTGAAACTGATTTTCTTGAGTGCAAATGAGGTAGGCGGAATGTGTTGTGTAGCGGTGAAATGCTTAGATATAACACAGAACCCCGATTGCGAAGGCAGCTTACTGGGATGCAACTGACGCTGAGGCACGAAAGCGTGGGGATCAAACAGG
## 1082                                                        ACGAAGGGAGCTAGCGTTATCCAAAATGATTGGGCGTAAAGAGTTTGTAGGCGGTATTTGTTGTTATATGTGAAATATTAATTTTATTTTTTAATAATGCATATAATACTCTTTTACTTGAGTATAACAGAGGATAACAGAATTTCATACGGAAGAGTAAAATCTAATGATTTATGAAGGAATGTCAGAGGCGAAAGCGGTTATCTGGGTTATTTACTGACGCTGAGGAACGAAAGCATGGGGAGCAAACAGG
## 1083                                                         ACGAAGGGTGCTAGCGTTGTTCGGAATCACTGGGCGTAAAGCGTACGTAGGCGGTTTAGTAAGTTGATTGTGAAAGCCCCAGGCTTAACTTGGGAACTGCAATCAATACTGCTTTTCTAGAATATGGTGGGGGATAGTGGAATTCCTAGTGTAGGGGTGAAATCCGTAGATATTAGGAGGAATACCGGTGGCGAAAGCGACTATCTATGCCATTATTGACGCTAAGGTACGAAAGCGTGGGGAGCAAACAGG
## 1085                                                         ACAGAGGCCCCGAGCGTTGTTCGGAATTACTGGGCGTAAAGGGAATGTAGGTGGCTATATAAGTCAGATGTGAAATCCCCGGGCTTAACCCGGGAACTGCATTTGATACTGCATAGCTCGAGTGCGGAAGAGGAGAGCGGAATTCCCAGTGTAGGAGTGAAATCCGTAGATATTGGGAGGAATACCGATGGCGAAGGCAGCTCTCTGGTCCGTAACTGACACTGAGATTCGAAAGCGTGGGTAGCAAACAGG
## 1086                                                          ACGTAGGGGGCAAGCGTTGTCCGGAATTATTGGGCGTAAAGCGCGCGCAGGCGGCTTGTTAAGTCTGATGTGAAAGGCAAGGGCTCAACCCTTGTTAGCATTGGAAACTGACAGACTAGAGTGCAGCAGAGGAAAGTGGAATTCCACGTGTAGCGGTGAAATGCGTAGAGATGTGGAGGAACACCAGTGGCGAAGGCGACTTTCTGGGCTGTAACTGACGCTGAGGCGCGAAAGCGTGGGGAGCAAACAGG
## 1088                                                         ACGGAGGGTGCAAGCGTTATCCGGAATCATTGGGTTTAAAGGGTCCGCAGGCGGGCTTATAAGTCAGTGGTGAAAGCCCATCGCTTAACGATGGAACTGCCATTGATACTGTAAGTCTTGAATTCGGTCGAAGTGGGCGGAATGTGACATGTAGCGGTGAAATGCTTAGATATGTCACAGAACACCGATAGCGAAGGCAGCTCACTAGGCCTGGATTGACGCTCATGGACGAAAGCGTGGGGAGCAAACAGG
## 1089                                                         ACAGAGGTCTCAAGCGTTGTTCGGATTCATTGGGCGTAAAGGGTGCGTAGGCGGCGTGGTAAGTTGAGTGTGAAATCCTAGGGCTTAACCTTAGAACTGCACTCAATACTGCCATGCTTGAGGAATGTAGAGGAGAGTGGAATTCACGGTGTAGCAGTGAAATGCGTAGATATCGTGAGGAAGACCAGTTGCGAAGGCGACTCTCTGGGCATTTCCTGACGCTGAGGCACGAAGGCCAGGGGAGCAAACGGG
## 1091                                                         ACGTAGGGTGCAAGCGTTGTCCGGAATTACTGGGTGTAAAGGGAGCGCAGGCGGGAAGACAAGTTGGAAGTGAAAACCATGGGCTCAACCCATGAATTGCTTTCAAAACTGTTTTTCTTGAGTAGTGCAGAGGTAGATGGAATTCCCGGTGTAGCGGTGGAATGCGTAGATATCGGGAGGAACACCAGTGGCGAAGGCGGTCTACTGGGCACCAACTGACGCTGAGGCTCGAAAGCATGGGTAGCAAACAGG
## 1092                                                         ACATAGGGTGCAAGCGTTGTCCGGAATTATTGGGCGTAAAGAGCTCGTAGGTGGTTCGTCACGTCGGATGTGAAACTCTGGGGCTTAACCCCAGACCTGCATTCGATACGGGCGAGCTTGAGTATGGTAGGGGAGTCTGGAATTCCTGGTGTAGCGGTGGAATGCGCAGATATCAGGAGGAACACCAATGGCGAAGGCAGGTCTCTGGGCCATAACTGACACTGAGGAGCGAAAGTGCGGGGAGCGAACAGG
## 1093                                                         ACAGAGGATGCAAGTGTTATCCGGAATTACTGGGCGTAAAGCGTCTGTAGGTGGTTTGATAAGTCAATTGTTAAATCTTGATGCTTAACTTCAAAACTGCAATTGAAACTATTTGACTTGAGTATGGTAGAGGTAAAAGGAATTTTCAGTGGAGCGGTGAAATGCGTAGATATTGGAAAGAACACCGATGGCGAAAGCATTTTACTGGGCCATCACTGACACTCAGAGACGAAAGCTAGGGTAGCAAATGGG
## 1094                                                         ACGGAGGGTGCAAGCGTTATTCGGAGTTACTGGGCGTAAAGGGCGCGTAGGCGGTCTGTTATGTCTGATGTGAAAGCCCTGGACTTAATTCAGGAAGTGCATTGGAAACTGGCAGGCTAGAGTATAGGAGGGGAGAGCGGAATTCCTGGTGTAGAGGTGAAATTCGTAGATATCAGGAGGAACAACAGTGGCGAAGGCGGCTCTCTGGACTATAACTGACGCTGAGGCGCGAAAGCGTGGGTAGCAAACAGG
## 1096                                                         ACGAAGGGTGCAAGCGTTGTTCGGAATCATTGGGCGTAAAGCGCGCGCAGGCGGATTTGCAAGTCAGATGTGAAATCTCGAAGCTCAACTTCGAAACTGCGTCTGAAACTGCTAGTCTAGAATGTCGGAGGGGGCAGGGGAATTTCACGTGTAGGGGTAAAATCCGTAGAGATGTGAAGGAACACCGGAGGCGAAGGCGCCTGCCTGGACGACTATTGACGCTGAGGCGCGAAAGCGTGGGGAGCAAACAGG
## 1097                                                         ACAGGGGGTGCAAGCGTTGTTCGGAATCACTGGGCGTAAAGGGAGCGTAGGCGGAATTGCAAGACAGAGGTTAAATGTCCGGGCTCAACCTGGAACCTGCCTTTGTGACTGCAGTTCTTGAGTATGGCAGAGGCTAGTGGAATTACAGGTGTAGCGGTGGAATGCGTAGATATCTGTAAGAACACCAGAGGCGAAGGCGACTAGCTGGGCCAATACTGACGCTGAGGCTCGAAAGTGTGGGGAGCAAACAGG
## 1099                                                         ACAGGGGATGCAAGTGTTATCCGGAATTATTGGGCGTAAAGCGTCTGCAGGTTGATAATTAAGTCTTTTGTTAAACCTCTGGGCTTAACCCAGAATCTGCAAAAGAAACTATTTATCTTGAGTATGGTAGAGGTAAAGGGAATTTCCAGTGGAGCGGTGAAATGCGTAGAGATTGGAAAGAACACCAACAGCGAAGGCACTTTACTGGGCCAGTACTGACACTCAGAGACGAAAGCTAGGGGAGCAAACAGG
## 1100                                                         ACGAAGGGGGCAAGCGTTGTTCGGAATCACTGGGCGTAAAGCGTGCGTAGGCGGTCTAACAAGTCAGGAGTGAAATCCCCGGGCCTAACCCGGGAACTGCTTTTGAAACTGTTAGACTTGAATCCGGGAGAGGTTGGCGGAATTCTCAGTGTAGAGGTGAAATTCGCAGATATTGAGAGGAACACCAGTGGCGTAAGCGGCCAACTGGACCGGTTTTGACGCTGAGGCACGAAAGCGTGGGGATCAAACAGG
## 1101                                                        ACGAAGGTGGCAAGCGTTGTCCGGTCTTATTGGGCGTAAAGAGTTCTGTAGGCGGAATTATAAGTTATATGTTAAAGGCTACGGCCCAACCGTAGTTTCGCGTGTAATACTGTAGTTCTTGAGTAATGTAGAGGTTAGCAGAACGCACGGTGTAGTAGTGAAATGCGTTGATATCGTGCGGAATACCAAAGGCGTAGGCAGCTAACTGGACATTTACTGACGCTGAGAGACGAAAGCTTGGGGAGCAAAACGG
## 1102                                                         ACGGGGGGTGCAAGCGTTATTCGGAATTATTGGGCGTAAAGGGTGCGTAGGCGGTTTTGTAAGTCAGATGTGAAATACCGGGGCTTAACTTCGGAACTGCATTTGATACTGCGAGACTTGAGTATGGGAGAGGAGAGCGGAATTCCTGGTGTAGAGGTGAAATTCGTAGATATCAGGAGGAACACCGGTGGCGAAGGCGGCTCTCTGGACCAATACTGACGCTGAGGCACGAAGGCGTGGGGAGCAAACAGG
## 1103                                                         ACGAAGGGTGCTAGCGTTGTTCGGAATCACTGGGCGTAAAGCGCGCGTAGGCGGTTATTCAAGTCAGGGGTGAAATCCTGAGGCTCAACCTCAGAATTGCCTTTGAAACTGTATGACTGGAGTGTCGGAGGGGATAGCGGAATTGCTAATGTAGCGGTGAAATGCGTAGATATTAGCAGGAACACCGGTGGCGAAGGCGGCTATCTGGACGACAACTGACGCTGAGGCGCGAAAGCGTGGGTATCAAACAGG
## 1104                                                         ACGGAGGGTGCAAGCGTTGTTCGGAATTACTGGGCGTAAAGCGCGCGTAGGCGGTTTGCTAAGTCAGATGTGAAAGCCCTCGGCTTAACCGGGGACGTGCATTTGAAACTGGCAGACTTGAGTACTGGAGGGGGTGGTGGAATTCCCGGTGTAGAGGTGAAATTCGTAGATATCGGGAGGAATACCGGTGGCGAAGGCGACCACCTGGCCAGATACTGACGCTGAGGTGCGAAAGCGTGGGGAGCAAACAGG
## 1105                                                         ACGTAGGGTGCGAGCGTTAATCGGAATTACTGGGCGTAAAGGGTGCGCAGGCGGTTGAGTAAGACAGATGTGAAATCCCCGAGCTTAACTCGGGAATGGCATATGTGACTGCTCGACTAGAGTGTGTCAGAGGGAGGTGGAATTCCACGTGTAGCAGTGAAATGCGTAGATATGTGGAAGAACACCGATGGCGAAGGCAGCCTCCTGGGACATAACTGACGCTCAGGCACGAAAGCGTGGGGAGCAAACAGG
## 1107                                                         ACAGAGGTCTCAAGCGTTGTTCGGATTCATTGGGCGTAAAGGGTGCGTAGGTGGCGATGTAAGTCGGATGTGAAATCTCCAAGCTTAACTTGGAAACTGCATTCGATACTGCGTTGCTAGAGGACTGTAGAGGGCATTGGAATTCACGGTGTAGCAGTGAAATGCGTAGATATCGTGAGGAAGACCAGTGGCGAAGGCGAATGCCTGGGCAGTTCCTGACACTGAGGCACGAAGGCCAGGGGAGCAAACGGG
## 1109                                                         ACGAAGGGTGCTAGCGTTGTTCGGAATCATTGGGCGTAAAGCGTGCGTAGGCGGCTTAGTAAGTCGTTTGTGAAATACCTAGGCTTAACTTAGGAACTGCAATCGAAACTACTTTGCTAGAGTTTGGTGGGGGATAGTGGAACTCCTAGTGTAGGGGTGAAATCCGTAGAGATTAGGAGGAATATCAGTGGCGAAGGCGACTATCTACACCAATACTGACGCTGAGGTACGAAAGCGTGGGGAGCAAACAGG
## 1110                                                         ACATAGGGTGCAAGCGTTGTCCGGAATTATTGGGCGTAAAGAGCTCGTAGGTGGTTCGTCACGTCGGATGTGAAACTCTGGGGCTTAACCCCAGACCTGCATTCGATACGGGCGAGCTTGAGTATGGTAGGGGAGTCTGGAATTCCTGGTGTAGCGGTGGAATGCGCAGATATCAGGAGGAACACCAATGGCGAAGGCACATCTCTGGGCCGTAACTGACGCTGAGGAGCGAAAGCGTGGGGAGCGAACAGG
## 1111                                                         ACGAAGGATGCAAGCGTTATCCGGATTCATTGGGTTTAAAGGGAGCGTAGGCGGACTTATAAGTCAGTGGTGAAATCTTTGGGCTTAACCCAGAAACTGCCATTGATACTGTAAGTCTAGAGTACAGTTGCCGTTGGCGGAATATGACATGTAGTGGTGAAATACTTAGATATGTCATAGAACACCGATTGCGAAGGCAGCTAACGAAGCTGTAACTGACGCTGAGGCTCGAAAGTGCGGGGATCAAACAGG
## 1113                                                         ACAGGGGATGCAAGTGTTATCCGGAATTATTGGGCGTAAAGCGTCTGTAGGTTGGTAATTAAGTCTTTTGTTAAACCTTCGGGCTCAACCCGAAATCTGCAAAAGAAACTATTTATCTTGAGTATAGTAGGGGTAAAGGGAATTTCCAGTGGAGCGGTGAAATGCGTAGAGATTGGAAAGAACACCAAAAGCGAAGGCACTTTACTGGGCTATAACTGACACTGAGAGACGAAAGCTAGGGGAGCAAACGGG
## 1114                                                         ACGGAGGGGGCTAGCGTTGTTCGGAATCACTGGGCGTAAAGCGCGCGTAGGCGGCTTGATAAGTTAGAAGTGAAATCCCTGGGCTTAACCTAGGAATTGCTTTTAAAACTGTCGAGCTAGAGAATGGTAGAGGAGAGTGGAATTCCTAGTGTAGAGGTGAAATTCGTAGATATTAGGAGGAACACCAGAGGCGAAGGCGGCTCTCTGGGCCATTTCTGACGCTAAGGCGCGAAAGCGTGGGGAGCAAACAGG
## 1116                                                        ACGAACCAAGCGAACGTTATTCGGAATTACTGGGCTTAAAGCGCGTGTAGGCGGATGGCCACGTCGGTTGTTGAAATCCCCCGGCTTAACCGGGGAACAGGCACCGATACGAGTCGTCTAGAGGTGGGTAGGGGAGACTGGAACTTCCGGTGGAGCGGTGAAATGCGTTGAGATCGGAAGGAACGCCTGCGGCGAAAGCGAGTCTCTGGCCCCATACTGACGCTGAGACGCGAAAGCCAGGGGAGCAAACGGG
## 1117                                                         ACGTATGGGGCGAGCGTTGTTCGGAATTATTGGGCGTAAAGGGTGCGCAGGCGGTTGTACAAGTCTGGTGTTAAAGCCCGCAGCTTAACTGCGGAAAGGCATTGGAAACTGTACGACTAGAGTATTGGAGGGGTAGCTGGAATTCCAGGTGTAGGGGTGAAATCTGTAGATATCTGGAAGAACATCAATGGCGAAGGCAAGCTACTAGCCGCATACTGACGCTGAGGCACGAAAGCGTGGGGAGCAAACAGG
## 1118                                                         ACGTAGGGTGCGAGCGTTAATCGGAATTACTGGGCGTAAAGCGTGCGCAGGCGGTTATACAAGACAGGCGTGAAATCCCCGGGCTTAACCTGGGAATGGCGCCTGTGACTGTATAGCTAGAGTACGGTAGAGGGGGATGGAATTCCGCGTGTAGCAGTGAAATGCGTAGATATGCGGAGGAACACCGATGGCGAAGGCAATCCCCTGGACCTGTACTGACGCTCATGCACGAAAGCGTGGGGAGCAAACAGG
## 1121                                                         ACGTAGGATCCAAGCGTTGTCCGGATTTATTGGGTTTAAAGGGTGCGTAGGCGGAAAGATAAGTCAGTGGTGAAAGCCGGTCGCTCAACGATCGAATTGCCATTGATACTGTTTTTCTAGAATACGGTTGAGGTAGGCGGAATGTGTAGTGTAGCGGTGAAATGCATAGATATTACACAGAACTCCGATTGCGAAGGCAGCTTACTAAGCCGTTATTGACGCTGAGGCACGAAAGCGTGGGGATCAAACAGG
## 1122                                                         ACGAAGGATGCAAGCGTTATCCGGATTCATTGGGTTTAAAGGGTGCGTAGGCGGATTAGTAAGTCAGTGGTGAAATACCATCGCTTAACGATGGGACTGCCATTGATACTGCTGATCTTGAGTACATATGACGTTGGCGGAATATGACATGTAGCGGTGAAATGCATAGATATGTCATAGAACACCGATTGCGAAGGCAGCTAACGAAAATGTAACTGACGCTGATGCACGAAAGTGCGGGGATCAAACAGG
## 1123                                                         ACGGAGGATCCAAGCGTTATCCGGATTTATTGGGTTTAAAGGGTGCGTAGGCGGACTGTTAAGTCAGTGGTGAAAGGTTGACGCTCAACGTTAACATTGCCATTGAAACTGGTAGTCTTGAGTGCAAATGAGGCAGGCGGAATGTGTTGTGTAGCGGTGAAATGCTTAGATATGACACAGAACACCAATTGCGAAGGCAGCTTGCTGGAATGTAACTGACGCTGAGGCACGAAAGCGTGGGGATCAAACAGG
## 1124                                                         ACGTAGGGTGCAAGCGTTATTCGGAATTATTGGGCGTAAAGCGCGTGTAGGCGGTCTAGTTAGTCTAATGTGAAATCCCTCGACTCAATCGAGGAACTGCATCGGATACTGCTAGACTAGAGTACAGGAGGGGTGAGCGGAATTCCTGGTGTAGAGGTGAAATTCGTAGATATCAGGAGGAACAACAGCGGCGAAGGCGGCTCACTGGACTGTAACTGACGCTCAGACGCGAAAGCGTGGGGAGCAAACAGG
## 1125                                                         ACGGAGGGTGCGAGCGTTATCCGGAATTACTGGGTTTAAAGGGTGCGTAGGCGGATGATTAAGTCTAAGGTGAAAGTTTGTCGCTTAACGATAAAATTGCCTTGGATACTGGTTATCTTGAATAAGGATGAGGATGGCGGAATGTGGCATGTAGCGGTGAAATGCATAGATATGCCATAGAACACCAATTGCGAAGGCAGCTATCTGGTCTTTTATTGACGCTGAGGCACGAAAGCGTGGGGATCGAACAGG
## 1126                                                         ACGAAGGGGGCAAGCGTTGTTCGGAATTACTGGGCGTAAAGGGAGTGTAGGCGGTTATGTAAGATAGTGGTGAAATCCCAGAGCTTAACTTTGGAATTGCCATTATGACTATGTGGCTAGAATTACAGAGAGGATAGTGGAATACCCAGTGTAGAGGTGAAATTCGTAGATATTGGGTAGAACACCAGTGGCGAAGGCGACTATCTGGCTGTATATTGACGCTGAGGCTCGAAAGCATGGGGATCAAACAGG
## 1127                                                          ACGTAGGCGGCAAGCGTTGTCCGGAATTATTGGGCGTAAAGGGAGCGCAGGCGGGAAACTAAGCGGATCTTAAAAGTGCGGGGCTCAACCCCGTGATGGGGTCCGAACTGGTTTTCTTGAGTGCAGGAGAGGAAAGCGGAATTCCCAGTGTAGCGGTGAAATGCGTAGATATTGGGAAGAACACCAGTGGCGAAGGCGGCTTTCTGGACTGTAACTGACGCTGAGGCTCGAAAGCTAGGGTAGCGAACGGG
## 1128                                                         ACATAAGGTGCGAGCGTTATCCGGAATCACTGGGCGTAAAGGGCATCTAGGCGGTTTATTAAGTCAGGGGTGAAAAGCCGAAGCTCAACTTTGGTCTTGCCTTTGAAACTGATAGACTAGAATACGGGATAGGAGGGCGGAACTACAAGTGTAGAGGTGAAATTCGTAGATATTTGTAGGAATGCCGATGACGAAGGTAGCTCTCTGGACCGATATTGACGCTGAAGTGCGAAAGCTAGGGTAGCAAACAGG
## 1130                                                         ACGAGGGATCCTAGCGTTGTTCGGAATCATTGGGCGTAAAGCGGGTGTAGGTGGCTTTTTAAGTCAGGTGTGAAATCCTAGGGCTTAACCCTAGAAGTGCATTTGATACTATTAAGCTTGAGTGTGGAAGAGGCTATTAGAATTCCTGGTGTAGTGGTGAAATACGTAGATATCAGGAGGAATACCGGTTGCGAAGGCGGATAGCTGGTCCAACACTGACACTAAGACCCGAAAGCGTGGGGATCAAACAGG
## 1133                                                         ACGGAGGATGCAAGCGTTATCCGGATTCATTGGGTTTAAAGGGTGCGTAGGCGGCTTTTTAAGTCAGTGGTGAAATCCTGCCGCTTAACGGTAGAATTGCCATTGATACTGAAAAGCTTGAGTACAGTTGAGGTAGGCGGAATGTGTAGTGTAGCGGTGAAATGCTTAGATATTACACAGAACACCGATTGCGAAGGCAGCTTACTAAACTGTTACTGACGCTGATGCACGAAAGCGTGGGGAGCGAACAGG
## 1134                                                        ACGAACCAAGCGAACGTTATTCGGAATTACTGGGCTTAAAGCGCGTGTAGGCGGATTGCCACGTCGGCTGTTGAAATCCCCCGGCTCAACCGGGGAACAGGCACCGATACGAGTGATCTTGAGGTGGGTAGGGGAGACTGGAACTTCCGGTGGAGCGGTGAAATGCGTTGAGATCGGAAGGAACGCCCGCGGCGAAAGCGAGTCTCTGGCCCCATACTGACGCTGAGACGCGAAAGCCAGGGGAGCAAACGGG
## 1136                                                         ACGGAGGATCCGAGCGTTATCCGGATTTATTGGGTTTAAAGGGAGCGTAGGTGGACTGGTAAGTCAGTTGTGAAAGTTTGCGGCTCAACCGTAAAATTGCAGTTGATACTGTCAGTCTTGAGTACAGTAGAGGTGGGCGGAATTCGTGGTGTAGCGGTGAAATGCTTAGATATCACGAAGAACTCCGATTGCGAAGGCAGCTCACTAGACTGCAACTGACACTGATGCTCGAAAGTGTGGGTATCAAACAGG
## 1137                                                          ACATATGTCGCAAGCGTTATCCGGATTTATTGGGCGTAAAGCGCGTCTAGGCGGTTTATTAAGTCTGATGTGAAAATGCGGGGCTCAACTCCGTATTGCGTTGGAAACTGGTAGACTAGAGTACTGGAGAGGTGGGCGGAACTACAAGTGTAGAGGTGGAATTCGTAGATATTTGTAGGAATGCCGATGGAGAAGTCAGCTCACTGGACAGATACTGACGCTAAAGCGCGAAAGCGTGGGGAGCAAACAGG
## 1138                                                         ACGTAGGTCCCAAGCGTTATCCGGATTTACTGGGTGTAAAGAGTCTGTAGCAGGTTCTATGAGTCTATTTTTAAAGACCTCGGCTCAACTGAGGGAAGGGAATGGATACTGTAGAACTGGAGGTTATGAGGGGTTAGTAGAATTCCCGGTGGAGGGGTGAAATCCGTTGATATTGGGAGGAATGCCGAAAGCGAAAGCAGCTAACTATTATTATCCTGACTGTGAGAGACGAAAGCATGGGTAGCAAACGGG
## 1139                                                         ACAGAGACCTCAAGCATTATCCGGATTTATTGGGCGTAAAGGGTCCGCAGGTGGTTTGGTGCGTCATGGGTTAAATCTACGAGCTCAACTCGTGGACCGCTCGTGATACGACCAAACTCGAGGCCGGGAGAGGTAAGCGGAATTACGGGTGTAGCAGTAAAATGCGTTAATATCCGTAAGAACACCAAATGCGAAGGCAGCTTACTGGAACGGTTCTGACACTCAGGGACGAAAGCGTGGGTAGCGAAAGGG
## 1141                                                         ACATGTGGGGCAAGCGTTGTTCGGAATTACTGGGCATAAAGGGTGCGTAGGCGGTCTGATAAGTCGGGCGTGAAATCCGGTGGCCCAACCATCGAACTGCGCTTGATACTGTTGGACTTGAGTGCGTGAGGGGAGACTGGAACGAGTGGTGTAGCGGTGAAATGCGTAGATATCACTCGGAACGCCAACGGCGAAGGCAGGTCTCTGGTACGCAACTGACGCTGAGGCACGAAAGCGTGGGGAGCAAACAGG
## 1142                                                         ACGTAGGGGGCGAGCGTTGTTCGGATTCACTGGGTGTAAAGGGCGTGTAGGTGGTTATTTAAGTCGAATGTGAAATCCCTTGGCTCAACCAAGGAAGTGCATTCGAAACTAAGTGACTAGAGTTTGGGAGAGGAGAGCGGAATTCTCGGTGTAGCGGTGAAATGCGTAGAGATCGAGAAGAACACCAGCGGCGAAAGCGGCTCTCTGGACCAATACTGACACTGAGGCGCGAAAGCGTGGGGAGCAAACGGG
## 1143                                                         ACAGAGGGTGCGAGCGTTAATCGGAATTACTGGGCGTAAAGCGCGCGTAGGCGGCTTGTTAAGTCGGATGTGAAATCCCTGGGCTCAACCTAGGAACTGCATTCGATACTGGCAAGCTAGAGTGTGGGAGAGGAAGGTAGAATTCCAGGTGTAGCGGTGAAATGCGTAGAGATCTGGAGGAATACCGATGGCGAAGGCAGCCTTCTGGCCTAACACTGACGCTGAGGTGCGAAAGCATGGGGAGCAAACAGG
## 1144                                                         ACGAAGGGTGCAAGCGTTGTTCGGATTTATTGGGCGTAAAGCGCACGTAGGTGGACTTATAAGTCAGATGTGAAATCTCGGGGCTCAACCTCGAAACTGCGTCTGAAACTGTGAGTCTAGAATCTTGGAGGGGGTAGGGGAATTTCACATGTAGGGGTAAAATCCGTAGATATGTGAAGGAACACCAGAGGCGAAGGCGCCTGCCTGGACAAGTATTGACACTGAGGTGCGAAAGCGTGGGGATCAAACAGG
## 1145                                                         ACGGAGGGTGCAAGCGTTATCCGGAATCATTGGGTTTAAAGGGTCCGTAGGCGGATAATTAAGTCAGAGGTGAAATCCCACAGCTTAACTGTGGAACTGCCTTTGATACTGGTTGTCTTGAATTATAGTGAAGTAGATAGAATGTGTAGTGTAGCGGTGAAATGCTTAGATATTACACAGAATACCGATTGCGAAGGCAGTTTACTAACTATATATTGACGCTAATGGACGAAAGCGTGGGGAGCGAACAGG
## 1146                                                         ACGGAGGGTGCAAGCGTTACTCGGAATCACTGGGCGTAAAGGATGCGTAGGCTGTAATATAAGTCAGGAGTGAAATCCAACGGCTCAACCGTTGAACTGCTCTTGAAACTGTTTTACTAGAATATGGGAGAGGTAGATGGAATTGGTGGTGTAGGGGTAAAATCCGTAGATATCACCAGGAATACCGATTGCGAAGGCGATCTACTGGAACATTATTGACGCTGAGGCATGAAAGCGTGGGGAGCAAACAGG
## 1147                                                         ACGTAGGGGGCAAGCGTTATCCGGATTTACTGGGTGTAAAGGGAGCGTAGACGGCGATGCAAGTCCGATGTGAAAACCCATGGCTCAACCATGGGATTGCATTAGAAACTGTATTGCTGGAGTGCAGGAGAGGTAAGCGGAATTCCTGGTGTAGCGGTGAAATGCGTAGATATCAGGAGGAACACCGGTGGCGAAGGCGGCTTACTGGACTGTAACTGACGTTGAGGCTCGAAGGCGTGGGGAGCAAACAGG
## 1148                                                         ACGGAGGATCCAAGCGTTATCCGGATTTATTGGGTTTAAAGGGTCCGTAGGCGGGTCTTTAAGTCAGTGGTGAAAGCCTGCAGCTTAACTGTAGAATTGCCATTGATACTGGAGACCTTGAGTGTAGTAGAAGTAGGCGGAATAAGGCATGTAGCGGTGAAATGCATAGATATGCCTTAGAACACCGATTGCGAAGGCAGCTTACTATGTTACAACTGACGCTGAGGGACGAAAGCGTGGGGAGCGAACAGG
## 1150                                                         ACGAAGGCCCCAAGCGTTATCCGGAATTACTGGGCGTAAAGCGTCTGTAGGTGGTTTGGAAAGTCTCAAGTGAAAGGTCAGGGCTCAACCCTGTACTTGCTTGGGAAACTATCGAACTAGAGTGTGGGAGAGGCAAGCAGAACGGTATGAGTAGGGGTGCAATCCGTTGATACATACCAGAATACCAAAAGCGAAGGCAGCTTGCTGGAACATTACTGACACTGAGAGACGAAAGCGTGGGGAGCAAAAGGG
## 1151                                                         ACGGGGGGTGCAAGCGTTACTCGGAATCACTGGGCGTAAAGAGCATGTAGGCGGATTAATAAGTTTGAAGTGAAATCCTATAGCTTAACTATAGAACTGCTTTGAAAACTGTTAATCTAGAATGTGGGAGAGGTAGATGGAATTTCTGGTGTAGGGGTAAAATCCGTAGAGATCAGAAGGAATACCGATTGCGAAGGCGATCTACTGGAACAATATTGACGCTGAGACGCGAAAGCGTGGGGAGCAAACAGG
## 1152                                                         ACGGAGGGTGCAAGCGTTATCCGGAATCATTGGGTTTAAAGGGTCCGCAGGCGGGCGTATAAGTCAGTGGTGAAAGCCTATTGCTTAACAATAGAACTGCCATTGATACTGTACGTCTTGAATTCGGTCGAAGTGGGCGGAATATGACATGTAGCGGTGAAATGCTTAGATATGTCATAGAACACCGATAGCGAAGGCAGCTCACTAGGCCTGAATTGACGCTCATGGACGAAAGCGTGGGGAGCAAACAGG
## 1153                                                         ACGTAGGGTGCGAGCGTTAATCGGAATTACTGGGCGTAAAGGGTGCGTAGGCGGGAGTATAAGTTAGATGTGAAATCCCTGAGCTTAACTTAGGAATAGCATTTAAGACTGTACACCTAGAGTTTATCAGAGGGGGGTAGAATTCCAAGTGTAGCAGTGAAATGCGTAGAGATTTGGAGGAATACCGATGGCGCAGGCAGCCCCCTGGGATGAGACTGACGCTGAGGCACGAAAGCGTGGGTAGCAAACAGG
## 1154                                                         ACGGAGGGTGCAAGCGTTATTCGGAGTTACTGGGCGTAAAGGGCGCGTAGGCGGCTCGTTAAGTCTGATGTGAAAGCCCTGGACTCAATCCAGGAAGTGCATTGGATACTGGCGAGCTAGAGTGCAGGAGGGGAGAGCGGAATTCCTGGTGTAGAGGTGAAATTCGTAGATATCAGGAGGAACAACAGTAGCGAAGGCGGCTCTCTGGACTGTAACTGACGCTGAGGCGCGAAAGCGTGGGTAGCAAACAGG
## 1155                                                         ACGAGGGGTGCAAGCGTTATTCGGAATAACTGGGCGTAAAGAGCGCGTAGGCGGCTTGTTAAGTCTGTTGTTAAATTGTCTGGCCTAACCAGATGTCGGCAATAGAAACTGTCAAGCTAGAGGACAAGAGAGAGGGGTGGAATTCTCGGAGTAGCGGTAAAATGCGTAGATCTCGAGAGGAACACCAATGGCGAAGGCAGCCCCTTGGCTTGTATCTGACGCTCAAGTGCGAAAGCGTGGGGAGCAAACAGG
## 1157                                                         ACGTAGGGTGCAAGCGTTAATCGGAATTACTGGGCGTAAAGGGTGCGCAGGCGGTATTGTAAGCCAGATGTGAAATCCCCGGGCTCAACCTGGGAACTGCGTTTGGAACTGCAATGCTAGAGTGTGTCAGAGGGGGGTGGAATTCCACGTGTAGCAGTGAAATGCGTAGAGATGTGGAGGAACACCGATGGCGAAGGCAGTCCCCTGGGATAAGACTGACGCTCATGCACGAAAGCGTGGGGAGCAAACAGG
## 1158                                                         ACGGAGGGTGCAAGCGTTAATCGGAATCACTGGGCGTAAAGCGCGCGTAGGCCGCCTTCTAAGTCGGACGTGAAAGCCCTCGGCTCAACCGAGGAACTGCGTTCGAAACTGGGAGGCTTGAGTCCTGGAGAGGGTGGCGGAATTCCGGGTGTAGGAGTGAAATCCGTAGATATCCGGAGGAACACCGGTGGCGAAGGCGGCCACCTGGACAGGTACTGACGCTGAGGCGCGAAAGCGTGGGGAGCAAACAGG
## 1161                                                         ACGAAGGGTGCAAGCGTTGTTCGGAATTATTGGGCGTAAAGGGTTCGTAGGTTGGAGAGTAAGTCAAGTGTGAAATCCCCGAGCTTAACTTGGGACGTGCACCTGAAACTGCTTTTCTTGAATGTTAGAGAGGGTCGTAGAATTGCTGGTGTAGGAGTGAAATCCGTAGATATCAGCAGGAATACCAGAGGCGAAGGCGGCGACCTGGCTAAACATTGACACTGAGGAACGAAAGCGTGGGGAGCAAACAGG
## 1163                                                         ACGGAGGGTGCGAGCGTTAATCGGAATTACTGGGCGTAAAGCGCGCGTAGGCGGTTTGATCAGTCCGTCGTGAAAGCCCCGGGCTTAACCTGGGAACTGCGGTGGATACTGTCGGGCTAGAGTGTGGTAGAGGGGAGTGGAATTTCCGGTGTAGCAGTGAAATGCGTAGAGATCGGAAGGAACACCAGTGGCGAAGGCGGCTCCCTGGACTAACACTGACGCTGAGGTGCGAAAGCGTGGGGAGCAAACAGG
## 1164                                                         ACGAAGGGTGCAAGCGTTGTTCGGAATTATTGGGCGTAAAGCGCGCGCGGGCGGATAGGCAAGTCAGATGTGAAATCTCGAGGCTCAACTTCGAAACTGCGTCTGAAACTACCAATCTAGAATGTCGGAGGGGGAAGGGGAATATCGCATGTAGGGGTAAAATCCGTAGAGATGTGATGGAACACCAGAGGGGAAGCCGCCTTCCTGGACGAATATTGACGCTTAGGCGCGAAAGCGTGGGGAGCAAACAGG
## 1168                                                         ACGAAGGGGGCTAGCGTTGTTCGGAATTACTGGGCGTAAAGCGAGCGTAGGTTGCTCGATAAGTCAGTGGTGAAAGCCCAGAGCTCAACTCTGGAACTGCCATTGAAACTGTCGAGCTGGAATCTCAGAGGGGGTAGCGGAATTCCAAATGTAGGGGTGAAATCCGTAGATATTTGGAGGAACACCGGTGGCGAAGGCGGCTACCTGGATGAGTATTGACACTGAGGCTCGAAAGCGTGGGGATCAAACAGG
## 1169                                                         ACGTAGGGTGCAAGCGTTGTCCGGATTTATTGGGCGTAAAGAGCTCGTAGGCGGTTAGACACGTCGGATGTGAAAACTGGGGGCTCAACTCCCAGCCTGCATTCGATACGGTCTGACTAGAGTGTGGTAGGGGTAACTGGAATTCCTGGTGTAGCGGTGAAATGCGCAGATATCAGGAGGAACACCGATGGCGAAGGCAGGTTACTGGGCCACCACTGACGCTGAGGAGCGAAAGCGTGGGGAGCAAACAGG
## 1171                                                         ACATAGGGTGCAAGCGTTGTCCGGATTTATTGGGCGTAAAGAGTTCGTAGGCGGTTTCTCGCGTCGGATGTGAAAACTCAGGGCTCAACCCTGAGCCTGCATTCGATACGGGGAAACTTGAGGACGGCAGGGGAGACTGGAACTCCTGGTGTAGCGGTGGAATGCGCAGATATCAGGAAGAACACCAATGGCGAAGGCAGGTCTCTGGGCCGATTCTGACGCTGAGGAACGAAAGCGTGGGGAGCAAACAGG
## 1173                                                         ACAGAGGTCTCAAGCGTTGTTCGGAATCACTGGGCGTAAAGGGTGCGTAGGCGGCTTGGTAAGTCAGATGTGAAAGCTCAGGGCTCAACCCTGAAATTGCATCCGATACTGCCAGGCTAGAGGACTGGAGAGGTGTCTAGAATTCTCGGTGTAGCAGTGAAATGCGTAGATATCGAGAGGAATACCAATGGCGAAGGCAGGACACTGGACAGTATCTGACGCTGAGGCACGAAGGCTAGGGTAGCGAACGGG
## 1174                                                         ACGGGAGTGGCAAGCGTTATCCGGAATTATTGGGCGTAAAGCGTCCGCAGGCGGCCTTGTAAGTCTGCTGTCAAAGCGTGGAGCTTAACTCCATTTCGGCAGTGGAAACTACAAGGCTTGAGTGTGGTAGGGGCAGAGGGAATTCCCGGTGTAGCGGTGAAATGCGTAGATATCGGGAAGAACACCAGTGGCGAAGGCGCTCTGCTGGGCCATAACTGACGCTCATGGACGAAAGCCAGGGGAGCGAAAGGG
## 1176                                                         ACGAAGGTGGCAAGCGTTACTCGGAATTACTAGGCGTAAAGGGCAGGTAGGTGGTTTAATAAGTCTGTTGTGTCAGCTCCTGGCTTAACTGGGAGAGGTCAACGGAAACTATTAGGCTTGAGTATGGGAGAGGGTGCTGGAATTCCTGGTGTAGCGGTGAAATGCGCAGATATCAGGAGGAACACCAATGGCGAAAGCAAGCACCTGGACCATTACTGACGCTAAGCTGCGAAAGCTAGGGGAGCAAACAGG
## 1177                                                         ACGTAGGGTGCGAGCGTTAATCGGAATTACTGGGCGTAAAGCGTGCGCAGGCGGTTTCGTAAGACAGAGGTGAAATCCCCGGGCTCAACCTGGGAACTGCCTTTGTGACTGCGAGGCTAGAGTATGGCAGAGGGGGGTGGAATTCCACGTGTAGCAGTGAAATGCGTAGAGATGTGGAGGAACACCGATGGCGAAGGCAGCCCCCTGGGTCGACACTGACGCTCATGCACGAAAGCGTGGGGAGCAAACAGG
## 1178                                                        ACGTAGGCACCAAGCGTTGTCCGGATTTATTGGGCGTAAAGAGCTCGTAGGCGGTTCAGTAAGTCGGGTGTGAAAACTCTGGGCTCAACCCAGAGACGCCACTCGATACTGCTGTGACTAGAGTGCGGTAGGGGAGCGGGGAATTCCTAGTGTAGCGGTGAAATGCGCAGATATTAGGAGGAACACCAGTGGCGAAGGCGCCGCTCTGGGCCGTAACTGACGCTGAGGAGCGAAAGCGTGGGGAGCAAACAGG
## 1180                                                        ACAGAGACCCCAAGCGTTATCCGGAATTATTGGGCGTAAAGGGTGAGTAGGTGGTTTTATTAGTCTTGTTTTAAAGATTTTGGGCTTAACCTAAAATATGAATAAGAAACGGTAGAACTAGAATTTTGGAGAGGTTAGTGGAACTCTATGTGTAGGGGTAAAATCCGTTGATATATAGGGGAACACCTAAAGCGAAGGCAGCTAACTGGCCAAACATTGACACTGAATCACGAAAGCGTGGGTAGCGATACGG
## 1181                                                         ACAGAGGTCTCAAGCGTTGTTCGGAATCACTGGGCGTAAAGGGTGCGTAGGTGGCGTGGTAAGTCAGATGTGAAAGCCCGGGGCTCAACTCCGGAACTGCATCCGATACTGCCGTGCTAGAGCACTGAAGAGGTGACCGGAATTCTCGGTGTAGCAGTGAAATGCGTGGATATCGAGAGGAACACCAAAGGCGAAGGCAGGTCACTGGGCAGTTGCTGACACTGAGGCACGAAGGCCAGGGTAGCGAACGGG
## 1182                                                         ACGTAGGGTGCGAGCGTTAATCGGAATTACTGGGCGTAAAGCGTGCGCAGGCGGTTTCGTAAGACAGAGGTGAAATCCCCGGGCTCAACCTGGGAACTGCCTTTGTGACTGCGAGGCTAGAGTATGGCAGAGGGGGGTGGAATTCCACGTGTAGCAGTGAAATGCGTAGAGATGTGGAGGAACACCGATGGCGAAGGCAGCCCCCTGGGCCAATACTGACACTGAGGAGCGAAAGCGTGGGGAGCGAACAGG
## 1185                                                         ACGTAGGGTGCAAGCGTTAATCGGAATTACTGGGCGTAAAGCGTGCGTAGGCGGCTATTCAAGTCAGAGGTGAAAGCCTGGAGCTCAACTCCAGAACTGCCTTTGAAACTAGATAGCTAGAATCTTGGAGAGGTGAGTGGAATTCCGAGTGTAGAGGTGAAATTCGTAGATATTCGGAAGAACACCAGTGGCGAAGGCGACTCACTGGACAAGTATTGACGCTGAGGTACGAAAGCGTGGGGAGCAAACAGG
## 1186                                                         ACGGGGGGTGCAAGCGTTACTCGGAATCACTGGGCGTAAAGAGCATGTAGGCGGGTAAATAAGTTGGAAGTGAAATCCTATAGCTTAACTATAGAACTGCTTCCAAAACTGTTTACCTAGAATATGGGAGAGGTAGATGGAATTTCTGGTGTAGGGGTAAAATCCGTAGAGATCAGAAGGAATACCGATTGCGAAGGCGATCTACTGGAACATTATTGACGCTCAGATGCGAAAGCGTGGGGAGCAAACAGG
## 1187                                                         ACGGAAGGTCCGGGCGTTATCCGGATTTATTGGGTTTAAAGGGAGCGTAGGCCGCCTTATAAGCGAGCAGTGAAATGTAGAGGCCCAACCTCTGAATTGCTGTTCGAACTGTAGGGCTTGAGTACGCACGAGGTAGGCGGAATTTGTGGTGTAGCGGTGAAATGCTTAGATATCACAAAGAACCCCGATTGCGAAGGCAGCTTACCGGAGCGCAACTGACGCTGAAGCTCGAAAGTGCGGGTATCGAACAGG
## 1188                                                         ACGTGGGGTGCAAGCATTAATCGGATTTATTGGGCGTAAAGGGTGCGTAGGCGGATCAGAAAGTGGGATGTGAAATCCCGGGGCTTAACCCCGGAACTGCATTCCAAACTCCTGATCTTGAGGGTAGACGGGGAGAATAGAATTCCACATGTAGCGGTGAAATGCGTAGATATGTGGAAGAATACCGGTGGCGTAGGCGATTCTCTAGTTTATACCTGACGCTGAGGCACGAAAGCTGGGGGAGCAAACAGG
## 1189                                                         ACGTAGGGTGCGAGCGTTAATCGGAATTACTGGGCGTAAAGCGTGCGCAGGCGGTTTCGTAAGACAGACGTGAAATCCCCGGGCTCAACCTGGGAACTGCGTTTGTGACTGCGAGGCTAGAGTATGGCAGAGGGGGGTGGAATTCCACGTGTAGCAGTGAAATGCGTAGAGATGTGGAGGAACACCGATGGCGAAGGCAATCCCCTGGACCTGTACTGACGCTCATGCACGAAAGCGTGGGGAGCAAACAGG
## 1190                                                         ACAGAGGATGCAAGTGTTATCCGGAATTACTGGGCGTAAAGCGTCTGTAGGTGGTTTACTAAGTCGATTGTTAAAGCTTGAGGCTTAACTTCAAAACAGCAATTGAAACTAGTAGACTTGAGTATGGTAGAGGTAAAAGGAATTTTCAGTGGAGCGGTGAAATGCGTAGATATTGGAAAGAACACCGATGGCGAAAGCATTTTACTGGACCATAACTGACACTCAGAGACGAAAGCTAGGGTAGCAAATGGG
## 1192                                                         ACGAACTGTGCAAACGTTATTCGGAATCACTGGGCTTGAAGAGTGCGTAGGCGGTTTTGTAAGTAGGGTGTGAAAGCCCCCAGCTCAACTGGGGAATTGCGCCCTAAACTACAAGGCTGGAGTGAGGTAGGGGTGTGTGGAACTTCCAGTGGAGCGGTGAAATGTGTTGATATTGGAAGGAACGCCGGTGGCGAAAGCGACACACTGGGCCTTGTCTGACGCTGAGGCACGAAAGCCAGGGGAGCAAACGGG
## 1195                                                         ACGAACCGTGCGAACGTTGTTCGGAATCATTGGGCTTAAAGGGCGCGTAGGCGGGTGTTCAAGTCCGGGGTGAAAGCCTCCAGCTCAACTGGAGAAGAGCCCCGGATACTGTTCATCTGGAGAGGGATAGGGGCACATGGAACTTCCGGTGGAGCGGTGAAATGCGTAGAGATCGGAAGGAACGCCGGTGGCGAAAGCGATGTGCTGGATCTTTTCTGACGCTGAGGCGCGAAAGCTAGGGGAGCAAACGGG
## 1196                                                         ACGAAGGGTGCAAACGTTGCTCGGAATCATTGGGCGTAAAGCGCACGTAGGCGGCTTTCTAAGTCGGATGTGAAATCCCTCGGCTTAACCGAGGACGTGCATCCGAAACTGGGAGGCTTGAGTATGGAAGAGGGTCGCAGAATTCCCGGTGTAGAGGTGAAATTCGTAGATATCGGGAGGAATACCAGTGGCGAAGGCGGCGACCTGGGCCAATACTGACGCTGAGGTGCGAAAGCGTGGGGAGCAAACAGG
## 1198                                                         ACGGAAGGTCCGGGCGTTATCCGGATTTATTGGGTTTAAAGGGAGCGCAGGCCGGAAGCAAAGCGTGCCGTGAAATGTAGATGCTCAACATCTGAATTGCGGCGCGAACTGGTTTTCTTGAGTGCATACGAGGTAGGCGGAATTTGTGGTGTAGCGGTGAAATGCTTAGATATCACGAAGAACTCCGATTGCGAAGGCAGCCTACCAGGATGTAACTGACGCTAAAGCTCGAAAGTGCGGGTATCGAACAGG
## 1199                                                         ACGGAGGATCCGAGCGTTATCCGGATTTATTGGGTTTAAAGGGTGCGTAGGCGGAATAGCAAGTCAGTGGTGAAAGTTTGCAGCTTAACTGTAAAATTGCCATTGAAACTGCTGTTCTTGAGTACAGTAGAGGTAGGCGGAATGTGTAGTGTAGCGGTGAAATGCATAGATATTACACAGAACACCGATTGCGTAGGCAGCTTACTGGAGTGTAACTGACGCTGAGGCACGATAGCGTGGGGATCAAACAGG
## 1200                                                         ACAGAGGGTGCAAGCGTTGCTCGGAATCATTGGGCGTAAAGCGCGTGTAGGCGGTTTATTAAGTCTGGTGTGAAAGCCCGGGGCTCAACCCCGGAAGTGCACTGGATACTGGTAAACTAGAGGACGAGAGAGGATGGTGGAATTCCTGGTGTAGGGGTGAAATCCGTAGATATCAGGAGGAACATCGGTGGCGAAGGCGGCCATCTGGCTCGTTTCTGACGCTCAGACGCGAAAGCGTGGGTAGCAAACAGG
## 1202                                                        ACGTAGGGTGCAAGCATTATCCGGATTTACTGGGCGTAAAGAGTTCTGTAGGCGGTAATATAAGTTCAGGGTTAAATCTTCTCGCTCAACGAGAAAACTGCTTTGAATACTGTATTACTAGAGGCATTTAGGGGACAGCGGAACGTATGGTGAAGCAGTGAAATGCGTTGATATCATACGGAACATCAAGGGGGAAGCCAGCTGTCTGGGAATGATCTGACGCTGAGAGACGAAAGCGTGGGTAGCGAAAAGG
## 1206                                                         ACAGAGGGTGCAAACGTTGTTCGGAATTACTGGGCGTAAAGCGCGTGTAGGCGGCCCGACAAGTCGGATGTGAAAGCCCCGGGCTCAACCCGGGAAGTGCATTCGATACTGTCAGGCTGGAGTCCCGGAGAGGATGGTGGAACTCTCGGTGTAGAGGTGAAATTCGTAGATATCGAGAAGAACACCGGCGGCGAAGGCGGCCATCTGGACGGTGACTGACGCTGAGACGCGAAAGCGTGGGGAGCAAACAGG
## 1208                                                         ACAGAGGGTGCAAGCGTTAATCGGAATTACTGGGCGTAAAGGGCGCGTAGGCGGTCAGGTCAGTCGGATGTGAAATCCCCGGGCTCAACCTGGGAATTGCATTCGATACTGCCTGGCTAGAGTGTGGAAGAGGGAAGCGGAATTTCCGGTGTAGCGGTGAAATGCGTAGATATCGGAAGGAACACCAGTGGCGAAGGCGGCTTCCTGGTCCAACACTGACGCTGAGGCGCGAAAGCGTGGGGAGCGAACAGG
## 1209                                                         ACGAAGGATCCAAGCGTTGTCCGGATTTACTGGGTTTAAAGGGTGCGTAGGCGGGCTATTAAGTCAGTGGTGAAAGCCTGTAGCTTAACTATAGAATTGCCATTGAAACTGATAGTCTTGAGTATGTTTGAGGTTACTGGAATATAACATGTAGCGGTGAAATGCTTAGATATGTTATAGAACACCAATTGCGAAGGCAGGTAACTAAGTCATAACTGACGCTGAGGCACGAAAGCGTGGGGAGCAAACAGG
## 1210                                                         ACAGAGACTGCAAGCGTTACTCGGATTCACTGGGCGTAAAGGGAGCGCAGGCGGATTGGTGTGTCAGGCGTGAAATCCCGGGGCTCAACCCCGGGGCTGCGCCCGAAACTACCAATCTAGAGACTTGGAGGGGTAAGCGGAATTCTTGGTGGAGCAGTGAAATGCGTAGATATCAAGAGGAACACCAACGGCGAAGGCAGCTTACTGGACAAGATCTGACGCTCAGGCTCGAAAGCGTGGGGAGCAAAAGGG
## 1211                                                         ACAGAGGCCCCAAGCATTATCCGGAATCACTGGGCGTAAAGGGTGTCAAGGCGGCTATATTAGTCGTTCGTTAAATCCGTGGGCTCAACCTACGGTCTGCGAGCGAAACGGTATAGCTAGAGGTTGGAAGAGGTGCGTGGAACTCACGGTGTAGGGGTGAAATCCGTTGATATCGTGGGGAACACCAAAGGCGAAGGCAGCGCACTGGTCCAAATCTGACGCTCACACACGAAAGCCAGGGTAGCGAACGGG
## 1212                                                         ACAGAGGGTGCAAACGTTGTTCGGAATTACTGGGCGTAAAGCGTGTGTAGGCGGCTAAGTAAGTCAGATGTGAAAGCCCCGGGCTCAACCCGGGAAGTGCATTTGATACTGCCTAGCTTGAGTATCGGAGAGGTTGGTGGAATTCTTGGTGTAGAGGTGAAATTCGTAGATATCAAGAGGAACACCGGTGGCGAAGGCGGCCAACTGGACGAATACTGACGCTGAGACACGAAAGCGTGGGGAGCAAACAGG
## 1213                                                         ACATAGGGTGCAAGCGTTGTCCGGAATTATTGGGCGTAAAGAGCTCGTAGGTGGTTCGTCACGTCGGATGTGAAACTCTGGGGCTTAACCCCAGACCTGCATTCGATACGGGCGAGCTTGAGTATGGTAGGGGAGTCTGGAATTCCTGGTGTAGCGGTGGAATGCGCAGATATCAGGAGGAACACCAATGGCGAAGGCAGGACTCTGGGCCATTACTGACACTGAGGAGCGAAAGTCTGGGGAGCGAACAGG
## 1214                                                        ACGAAGGGGGCTAGCGTTGTTCGGAATTACTGGGCGTAAAGCGCGCGTAGGCGGTTTTACAAGCCAGAGGTGAAATGCCCGAGCTTAACTTGGGAATTGCCTTTGGAACTGTATGGCTTTGAGGACGAGAGAGGTGAGTGGAATTCCCAGTGTAGAGGTGAAATTCGTAGATATTGGGAAGAACACCGGTGGCGAAGGCGGCTCACTGGCTCGTTTCTGACGCTGTAGCGCGAAAGCGTGGGGATCAAACAGG
## 1217                                                         ACAGAGACCTCAAGCGTTATCCGGATTTATTGGGCGTAAAGCGTCCGCAGGTGGTTTGGCAAGTCAGGGGTTAAAACTTAACGCTTAACGTTAAGGCTGCTCTTGATACTACCAAACTAGAGATCGGAAGAGGTAAGCGGAATTCTCGGTGTAGTCGTAATAAGCGTTGATATCGAGAAGAACACCAAATGCGAAGGCAGCTTACTGGTACGTTTCTGACACTCATGGACGAAAGCGTGGGGAGCAAACAGG
## 1218                                                         ACAGAGGATGCAAGCGTTATCCGGAATCACTGGGCATAAAGCGTCTGTAGGTGGTTTGGTAAGTCTGCTGTTAAAGACTGGGGCTCAACCCCAGAAAAGCAGTGGAAACTGCTAGACTTGAGTGTGGTAGAGGTAGAGGGAATTCCTAGTGTAGCGGTGAAATGCGTAGATATTAGGAAGAACACCAATGGCGAAGGCAATCCCCTGGACCTGTACTGACGCTCATGCACGAAAGCGTGGGGAGCAAACAGG
## 1219                                                         ACGTAGGGTGTTAGCGTTGTTCGGAATCATTGGGCGTAAAGCGCGTGTAGGTGGCTGTGTAAGTCGAATGTGAAATCCCTGGGCTCAACCGAGGAAGTGCATTCGAAACTACATAGCTCGAGGACGGTAGAGGAAGGTGGAATTCCAAAAGTAGAGGTGAAATTCGTAGATATTTGGAGGAATACCGGCGGCGAAGGCGGCCTTCTGGGCCGTTCCTGACACTGAGACGCGAAAGCGTGGGTAGCAAACAGG
## 1220                                                         ACGAAGGGTGCAAGCGTTACTCGGAATTACTGGGCGTAAAGCGTGCGTAGGTGGTTCGTTAAGTCTGTCGTGAAATCCCCGGGCTCAACCTGGGAATGGCGATGGATACTGGCGGGCTAGAGTGCGGTAGAGGAGAGTGGAATTCCCGGTGTAGCAGTGAAATGCGTAGAGATCGGGAGGAACATCAGTTGCGAAGGCGGCTCTCTGGACCAGCACTGACACTGAGGCACGAAAGCGTGGGGAGCAAACAGG
## 1221                                                         ACAGAGGGTGCAAACGTTGTTCGGAATTACTGGGCGTAAAGCGCGTGTAGGCGGCTTGGCAAGTCGGGTGTGAAAGCCCTGGGCTCAACCCAGGAAGTGCACTCGAAACTGCTTCGCTAGAGTACCGGAGAGGATAGCGGAATTCTCGGTGTAGAGGTGAAATTCGTAGATATCGAGAGGAACACCGGTGGCGAAGGCGGCTATCTGGACGGTTACTGACGCTGAGACGCGAAAGCGTGGGGAGCAAACAGG
## 1222                                                         ACAGAGGATGCAAGCGTTATCCGGAATGATTGGGCGTAAAGCGTCTGTAGGTGGCTTAAAAAGTCTCCTGTCAAAGATCAGGGCTTAACCCTGGGCCGGCAGGAGAAACTCTTAGGCTAGAGTTTGGTAGGGGCAGAGGGAATTCCCGGTGGAGCGGTGAAATGCGTAGAGATCGGGAGGAACACCAAAGGCGAAAGCACTCTGCTGGGCCATAACTGACACTGAGAGACGAAAGCGAGGGGAGCAAAAGGG
## 1224                                                         ACAGAGACCTCAAGCGTTATCCGGATTAATTGGGCGTAAAGCGTAGGTAGGCGGTCTTGTTAGTCAGATGTCAAAGCCCGAGGCTCAACCTCGGAAAGGCATTTGAAACGGCAAGACTCGAGAGTGTGAGAGATCAGTGGAATTCATGGTGTAGTAGTGAAATGCGTTGATATCATGAGGAACATCCAAGGCGAAGGCAACTGATTGGCGCACTTCTGACGCTCAGCTACGAAAGCGTGGGTAGCGAATGGG
## 1225                                                          ACAGAGGTCCCGAGCGTTGTTCGGAATTACTGGGCGTAAAGGGCGCGTAGGCGGTAATACAAGTTGCGGGTAAAATTTCACGGCTCAACCGTGTCAGGCTCGCAAGACTGTGTTGCTAGAGGGCTGTAAGGGGGGTTGGAATTCTCGGTGTAGCAGTGAAATGCGTAGATATCGAGAGGAACATCAGTGGCGAAAGCGAACCCCTGGGCAGATCCAGACGCTGAGGCGCGAAAGCCAAGGTAGCAAACGGG
## 1226                                                         ACGGAGGATGCAAGCGTTATCCGGATTCATTGGGTTTAAAGGGTGCGCAGGCGGAATAATAAGTCAGCGGTGAAAGCCTACAGCTTAACTGTAGAACTGCCATTGATACTGTTATTCTTGAGTGTTGTTGAGGTAGGCGGAATGTGTCATGTAGCGGTGAAATGCTTAGATATGACACAGAACACCGATCGCGAAGGCAGCTTACCAAGCAACAACTGACGCTCATGCACGAAAGCGTGGGGATCAAACAGG
## 1228                                                         ACATAGGGTGCAAGCGTTGTCCGGAATTATTGGGCGTAAAGAGCTCGTAGGTTGTTTGTTACGTCGGATGTGAAAACCTGAGGCTCAACCTCAGGCCTGCATTCGATACGGGCAAACTAGAGTTTGGTAGGGGAGACTGGAATTCCTGGTGTAGCGGTGGAATGCGCAGATATCAGGAGGAACACCAATGGCGAAGGCAGGACTCTGGGCCATTACTGACACTGAGGAGCGAAAGCGTGGGGAGCGAACAGG
## 1230                                                         ACGGAGGGTGCAAGCGTTATTCGGAATTACTGGGCGTAAAGCGCGCGTAGGCTGCTTTGTAAGTCAGGGGTGAAATCCCACGGCTCAACCGTGGAACTGCCTTTGAAACTGCGAAGCTTGAATCCTGGAGAGGGTGGCGGAATTCCTGGTGTAGGAGTGAAATCCGTAGATATCAGGAGGAACACCGGTGGCGAAGGCGGCCACCTGGACAGGTATTGACGCTGAGGTGCGAAAGTGTGGGGAGCAAACAGG
## 1231                                                         ACGGAGGGGGCTAGCGTTGTTCGGAATTACTGGGCGTAAAGCGCACGTAGGCGGACCGGAAAGTCAGAGGTGAAATCCCAGGGCTCAACCTTGGAACTGCCTTTGAAACTCCTGGTCTTGAGGTCGAGAGAGGTGAGTGGAATTCCGAGTGTAGAGGTGAAATTCGTAGATATTCGGAGGAACACCAGTGGCGAAGGCGGCTCACTGGCTCGATACTGACGCTGAGGTGCGAAAGCGTGGGGAGCAAACAGG
## 1232                                                         ACGGAGGGTGCGAGTGTTATTCGGAATCACTGGGCGTAAAGCGAACGTAGGCGGATAGGCAAGTCAGATGTGAAATCCCGGGGCTCAACCTCGGAACTGCATTTGATACTGCTTGTCTTGAGTATGGTAGAGGTAGGTAGAATTCCCGGTGTAGAGGTGAAATTCGTAGATATCGGGAGGAATACCAGAGGCGAAGGCGACCTACTGGGCCATTACTGACGCTGAGGTTCGAAAGCGTGGGTAGCAAACAGG
## 1234                                                         ACGGAGGGTGCAAGCGTTGTTCGGAATCATTGGGCGTAAAGCGCGCGCAGGCGGATCAGCAAGTCATGTGTGAAATCTCGGGGCTCAACTCCGAAATTGCGCCTGAAACTGCTAGTCTAGAATATCATAGGGGATAGGGGTATTTCACGTGTAGGGGTAAAATCCGTAGAGATGTGAAGGAACACCGGAGGCGAAGGCGCCTATCTGGATGATTATTGACGCTGAGGCGCGAAAGCGTGGGGAGCAAACAGG
## 1235                                                         ACGTAGGATGCGAGCGTTGTCCGGAATTATTGGGCGTAAAGCGTACGTAGGCGGTTTGTTAAGTTCGATCTTAAAGACTGAGGCTCAACCTTAGGAGTGGATTGAAGACTGGCAGGCTAGAGGACATCAGAGGGAAGTGGAATTCCGAGTGTAGCGGTGAAATGCGTAGATATTCGGAGGAACACCAGAGGCGAAAGCGGCTTCCTGGGGTGGACCTGACGCTGAGGTACGAAAGCTTGGGGAGCAAACAGG
## 1236                                                         ACGGAGGGTGCAAGCGTTATCCGGATTTATTGGGTTTAAAGGGTGCGTAGGCGGGTTTGTAAGTCAGTGGTGAAAGCCTACAGCTTAACTGTAGAATTGCCATTGATACTGCAGACCTTGAGTATAGTTGAAGTAGGCGGAATGTGTCATGTAGCGGTGAAATGCTTAGATATGCCACAGAACACCGATTGCGAAGGCAGCTTGCTAAGCTATAACTGACGCTGAGGCACGAAAGCGTGGGGAGCAAACAGG
## 1239                                                         ACGAGTGGTGCAAACGTTATTCGGAATCACTGGGCTTAAAGGGTGCGTAGGCGGCTATGAAGGTCTGATGTGAAATACCACGGCTCAACCGTGGAATGGCATTGGAAACCATATGGCTGGAGTGAGACAGAGGTGAGCGGAACAGATGGTGGAGCGGTGAAATGCGTTGATATCATCTGGAACACCGGTGGCGAAAGCGGCTCACTGGGTCTTTACTGACGCTGAGGCACGAAAGCTAGGGGAGCAAACGGG
## 1240                                                         ACGTACTGTGCGAACGTTATTCGGAATCACTGGGCTTAAAGGGTGCGTAGGCGGCGATTCAAGTCAGATGTGAAATGCCTGAGCTCAACTGAGGCATTGCGTTTGAAACTGAATTGCTAGAGTATTTCAGGGGTGTGCGGAACTGCCAGTGGAACGGTGAAATGTGTTGATATTGGCAGGAACACCGGAGGCGAAAGCGGCACACTGGGGAATAACTGACGCTGAGGCACGAAAGCCAAGGGAGCGAACGGG
## 1241                                                        ACAGAGACCCCAAGCGTTATCCGGAATCATTGGGCGTAAAGGGTGAGTAGGTGGTATTATTAGTCGTGTGTTAAAGTCCAAGGGCTCAACCTTTGGAATGCATACGAAACGGTAAAACTAGAATATGTAAGAGGTGAGTGGAACTCTATGTGTAGGGGTAAAATCCGTTGATATATAGGGGAACACCAAAAGCGAAGGCAGCTCACTGGTACATTATTGACACTGAATCACGAAAGCGTGGGTAGCGATACGG
## 1242                                                        ACAGAGGGTGCAAGCGTTAATCGGAATCACTTGGGCGTAAAGGGTGCGTAGGTGGAATGATAAGTTTCGTGTGAAATCCCTGGGCTTAACCTGGGAACTGCACGGGAGACTGTTGTTCTAGAGTACAGGAGAGGTGAGTGGAATTTCCGGTGTAGCAGTGAAATGCGTAGAGATCGGAAGGAACACCAGTGGCGAAGGCGACTCACTGGCCTGATACTGACACTCAAGCACGAAAGCGTGGGGAGCAAACAGG
## 1243                                                         ACGTAGGCACCAAGCGTTGTCCGGATTTATTGGGCGTAAAGAGCTCGTAGGTGGTTCGTCACGTCGGATGTGAAACTCTGGGGCTTAACCCCAGACCTGCATTCGATACGGGCGAGCTTGAGTATGGTAGGGGAGTCTGGAATTCCTGGTGTAGCGGTGGAATGCGCAGATATCAGGAGGAACACCAATGGCGAAGGCAGGACTCTGGGCCATTACTGACACTGAGGAGCGAAAGCGTGGGGAGCGAACAGG
## 1245                                                         ACGAAGGGGGCTAGCGTTGTTCGGAATTACTGGGCGTAAAGCGCACGTAGGCGGACTTTTAAGTCAGGTGTGAAATCCCGGGGCTCAACCTCGGAACTGCATTTGAAACTGGAAGTCTTGAGACCAGGAGAGGTTAGCGGAATACCGAGTGTAGAGGTGAAATTCGTAGATATTCGGTGGAACACCAGTGGCGAAGGCGGCTAACTGGACTGGTACTGACGCTGAGGTGCGAAAGTGTGGGGAGCAAACAGG
## 1246                                                         ACGAAGGGTGCGAGCGTTAATCGGAATTACTGGGCGTAAAGCGCGCGTAGGTGGTTAAGTAAGCGATCTGTGAAAGCCCCGGGCTTAACCTGGGAACTGCAGGTCGAACTGCTTGGCTAGAGTATGGTAGAGGGGTGTGGAATTTCCTGTGTAGCGGTGAAATGCGTAGATATAGGAAGGAACATCAGTGGCGAAGGCGACACCCTGGACCGATACTGACACTGAGGCGCGAAAGCGTGGGTAGCGAACAGG
## 1247                                                         ACGAAGGGTGCAAGCGTTGTTCGGAATTATTGGGCGTAAAGGGTTCGTAGGCTGGAATTTAAGTCAAGTGTGAAATCCTCGGGCTTAACCTGAGACGTGCATTTGAAACTGAGTTTCTTGAGTACCAGAGAGGGTAGTGGAATTGCTGGTGTAGGAGTGACATCCGTAGAGATCAGCAGGAACACCGGAGGCGAAGGCGACTGCCTGGCTGGTAACTGACGCTGAGGAACGAAAGCGTGGGGAGCAAACAGG
## 1248                                                         ACGTAGGGTGCAAGCGTTATCCGGAATTACTGGGTTTAAAGGGTCTGTAGGCGGGTTGATAAGTCAGTGGTGAAATATCTGGGCTTAACTTGGAGGGTGCCATTGATACTATTAATCTTGAGAGCGGATGAAGTGGGCGGAATGTGCCGTGTAGCGGTGAAATGCTTAGATATGGCACAGAACACCGATTGCGAAGGCAGCTCACTAAACCGCGTCTGACGCTGAGGGACGAAAGCGTGGGGAGCAAACAGG
## 1249                                                         ACGGAGGGTGCAAGCGTTGTCCGGAATCATTGGGCGTAAAGAGTTCGTAGGCGGTTTATTAAGTCTGGTGTTAAAGCCCGAAGCTCAACTTCGGTTCGGCACTGGATACTGGTAGACTAGAATGTGGTAGAGGTAAAGGGAATTCCTGGTGTAGCGGTGAAATGCGTAGATATCAGGAGGAACATCGGTGGCGTAAGCGCTTTACTGGGCCATAATTGACGCTGAGGAACGAAAGCCGGGGTAGCGAATGGG
## 1250                                                         ACGAAGGGTGCTAGCGTTGTTCGGAATCACTGGGCGTAAAGCGCGCGTAGGCGGCGATACAAGTCAGAGGTGAAAGCCTGGAGCTCAACTCCAGAACTGCCTTTGAAACTGTATTGCTAGAGTGTCGGAGGGGATAGCGGAATTGCTAATGTAGAGGTGAAATTCGTAGATATTAGCAGGAACACCGGTGGCGAAGGCGGCTATCTGGACGACAACTGACGCTGAGGCGCGAAAGCGTGGGGATCAAACAGG
## 1251                                                         ACGAACCGTGCAAACGTTATTCGGAATCACTGGGCTTAAAGGGCGCGTAGGCGGGTAATCAAGTCAATGGTGAAATCCTCCAGCTTAACTGGAGAAGTGCCTTTGATACTGGTTGTCTAGAGGGAGGTAGGGGCATGTGGAACTTCAGGTGGAGCGGTGAAATGCGTAGATATCTGAAGGAACGCCAGTGGCGAAAGCGATGTGCTGGACCTCTTCTGACGCTGAGGCGCGAAAGCTAGGGGATCAAACGGG
## 1254                                                         ACGTAGGGTGCAAGCGTTAATCGGAATTACTGGGCGTAAAGCGTGCGCAGGCGGTTGTGTAAGACAGGCGTGAAATCCCCGGGCTCAACCTGGGAATGGCGCTTGTGACTGCACAGCTCGAGTGCGGCAGAGGGGGATGGAATTCCGCGTGTAGCAGTGAAATGCGTAGATATGCGGAGGAACACCAATGGCGAAGGCAATCCCCTGGGCCTGCACTGACGCTCATGCACGAAAGCGTGGGGAGCAAACAGG
## 1255                                                         ACGGAGGGTGCAAGCGTTATCCGGATTTACTGGGTTTAAAGGGTGCGTAGGCGGGCCTTTAAGTCAGTGGTGAAATCTCCCGGCTTAACCGGGAAACTGCCATTGATACTATTGGTCTTGAATTCAGTTGAGGTGGGCGGAATGTGTCGTGTAGCGGTGAAATGCTTAGATATGACACAGAACACCAATTGCGAAGGCAGCTCGCTAAACTGACATTGACGCTGAGGCACGAAAGCGTGGGGATCAAACAGG
## 1256                                                         ACAGAGGGTGCAAGCGTTAATCGGAATTACTGGGCGTAAAGAGTGCGTAGGCGGATAATTAAGTCATATGTGAAAGCCCCGGGCTTAACCTGGGAATTGCATCTGATACTGGTTATCTAGAGTATGGTAGAGGATAGTGGAATTTCTGGTGTAGCGGTGAAATGCGTAGATATCAGAAGGAACACCAGTGGCGAAGGCGACTGTCTGGGCCAATACTGACGCTGAGGTACGAAAGCGTGGGTAGCGAACAGG
## 1258                                                         ACGTATGTCGCAAGCGTTATCCGGAATTATTGGGCGTAAAGGGCATCTAGGCGGTTAATCAAGTCGAAGGTGAAAACTTGTGGCTCAACCATAAGCTTGCCTACGAAACTGAATAACTAGAGTACTGGAGAGGTGGACGGAACTACACGAGTAGAGGTGAAATTCGTAGATATGTGTAGGAATGCCGATGATGAAGATAGTTCACTGGACAGAAACTGACGCTGAAGTGCGAAAGCTAGGGGAGCAAACAGG
## 1259                                                         ACGTAGGTGGCAAGCGTTGTCCGGAATTACTGGGCGTAAAGGGCGCGTAGGCGGCACCTTAAGTCAGATGTGAAAACCCAGGGCTTAACCTTGGGACTGCATTTGAAACTGAGGAGCTAGAGTGCCGGAGAGGAAAGCGGAATTCCTAGTGTAGCGGTGAAATGCGTAGATATTAGGAGGAACACCAGTGGCGAAGGCGGCTTTCTGGACGGTAACTGACGCTGAGGCGCGAAAGCGTGGGGAGCAAACAGG
## 1261                                                       ACGTAGGGTGCAAGCATTATCCGGAGTGACTGGGCGTAAAGAGTTGCGTAGGTGGACAAGTAAGCGAATAGTGAAATCTGGGGGCTCAACCTCACAGACTATTATTCGAACTGCTTGTCTCGAGAATGGTAGAGGTAACTGGAATTTCTAGTGTAGGAGTGAAATCCGTAGATATTAGAAGGAACACCAATGGCGTAGGCAGGTTACTGGACCATTTCTGACACTGAGGCACGAAAGCGTGGGGAGCGAACCGG
## 1265                                                         ACGGAGGGGGCTAGCGTTGTTCGGAATGACTGGGCGTAAAGGGCGCGTAGGCGGTGTCTTAAGTGAGGCGTGAAAGCCCCGGGCTTAACCCGGGAGGTGCGTTTCATACTGAGATGCTAGAGTGCGAGAGAGGAAAGTGGAATTCCTAGTGTAGAGGTGAAATTCGTAGATATTAGGAAGAACACCAGAGGCGAAGGCGGCTTTCTGGCTCGTAACTGACGCTGAGGCGCGAAAGCGTGGGGAGCAAACAGG
## 1266                                                         ACGGAGGGTGCAAGCGTTATCCGGATTCACTGGGTTTAAAGGGTGCGTAGGTGGCTTTGTAAGTCAGTGGTGAAAGCCCGGAGCTCAACTCCGGAACTGCCATTGATACTGCTTAGCTTGAATCAACTTGAGGTGGATGGAATATTACATGTAGCGGTGAAATGCTTAGATATGTAATAGAACACCGATTGCGAAGGCAGTTCACTAAGGTTGTATTGACACTGAGGCACGAAAGCGTGGGGATCAAACAGG
## 1268                                                         ACGGAGGGTGCAAGCGTTAATCGGAATTACTGGGCGTAAAGCGCACGCAGGCGGTTTGTTAAGTCAGATGTGAAATCCCCGCGCTTAACGTGGGAACTGCATTTGAAACTGGCAAGCTAGAGTCTTGTAGAGGGGGGTAGAATTCCAGGTGTAGCGGTGAAATGCGTAGAGATCTGGAGGAATACCGGTGGCGAAGGCGGCCCCCTGGACAAAGACTGACGCTCAGGTGCGAAAGCGTGGGGAGCAAACAGG
## 1270                                                        ACGGAGGGTGCAAGCGTTATCCGGAATCACTGGGTTTAAAGGGTGCGTAGGCGGTTTGATAAGTCAGTTGTGAAATTCCCTCGCTCAACGAGTGGGACTGCGATTGATACTGTTGAACTTGAATCAGGTTGAGGTAGGCGGAATGTGGCATGTAGCGGTGAAATGCATAGATATGCCATAGAACACCAATTGCGAAGGCAGCTTGCTGGGCTTTGATTGACGCTGAGGCACGAAAGCGTGGGGAGCAAACAGG
## 1272                                                         ACGGAGGGTGCAAGCGTTATCCGGATTTATTGGGTTTAAAGGGTCCGTAGGCGGACCTATAAGTCAGTGGTGAAATCTTCCGGCTCAACCGGGAAACTGCCATTGATACTGTAGGTCTTGAGTATGTATGAAGTTGGCGGAATGTGTGGTGTAGCGGTGAAATGCTTAGATATCACACAGAACACCGATTGCGAAGGCAGCTGACTAATACATAACTGACGCTGAGGGACGAAAGTGTGGGGATCAAACAGG
## 1276                                                         ACGAAGGGGGCTAGCGTTGTTCGGAATTACTGGGCGTAAAGCGCACGTAGGCGGATATTTAAGTCAGGGGTGAAATCCCGGGGCTCAACCCCGGAACTGCCTTTGATACTGGGTATCTAGAGTATGGAAGAGGTGAGTGGAATTCCGAGTGTAGAGGTGAAATTCGTAGATATTCGGAGGAACACCAGTGGCGAAGGCGGCTCACTGGTCCATTACTGACGCTGAGGTGCGAAAGCGTGGGGAGCAAACAGG
## 1277                                                         ACGGAGGGGGCTAGCGTTGTTCGGAATTACTGGGCGTAAAGAGCGCGTAGGCGGTTTGGTAAGTTGGAAGTGAAATCCCGGGGCTTAACCTCGGAATTGCTTTCAAAACTGCCAATCTAGAGTGTAGTAGGGGATGATGGAATTCCTAGTGTAGAGGTGAAATTCTTAGATATTAGGAGGAACACCGGTGGCGAAGGCGGTCATCTGGGCTACAACTGACGCTGATGCGCGAAAGCGTGGGGAGCAAACAGG
## 1278                                                         ACATAGGGTGCAAGCGTTGTCCGGAATTATTGGGCGTAAAGAGCTCGTAGGTTGTTTGTTACGTCGGATGTGAAAACCTGAGGCTCAACCTCAGGCCTGCATTCGATACGGGCAAACTAGAGTTTGGTAGGGGAGACTGGAATTCCTGGTGTAGCGGTGGAATGCGCAGATATCAGGAGGAACACCAATGGCGAAGGCAGGTCTCTGGGCCAATACTGACACTGAGGAGCGAAAGCGTGGGGAGCGAACAGG
## 1279                                                         ACGTAGGGTGCAAGCGTTAATCGGAATTACTGGGCGTAAAGCGTGCGCAGGCGGTTATGTAAGACAGAGGTGAAATCCCCGGGCTCAACCTGGGAACTGCCTTTGTGACTGCATAGCTAGAGTACGGTAGAGGGGGATGGAATTCCGCGTGTAGCAGTGAAATGCGTAGATATGCGGAGGAACACCGATGGCGAAGGCAGTCCCCTGGGATAAGACTGACGCTCATGCACGAAAGCGTGGGGAGCAAACAGG
## 1280                                                         ACGAAGGGTGCGAGCGTTGTTCGGAATGACTGGGCGTAAAGGGTTCGTAGGCGGGAATACAAGTCAAGTGTGAAATCCCCAGGCTCAACCTGGGACGTGCACTTGAAACTGTATTTCTTGAGTCGAGGAGAGGGTAATGGAATTGCTGGTGTAGGAGTGACATCCGTAGAGATCAGCAGGAACACCGGAGGCGAAGGCGATTACCTGGCCGAAGACTGACGCTGAGGAACGAAAGCGTGGGGAGCAAACAGG
## 1281                                                         ACGGAGGGTGCAAGCGTTAATCGGAATAACTGGGCGTAAAGCGCACGCAGGCGGTTAGATAAGTCAGATGTGAAATCCCCGGGCTCAACCTGGGAACTGCATTTGAAACTGTCTGACTAGAGTCTTGTAGAGGGGGGTAGAATTCCAGGTGTAGCGGTGAAATGCGTAGAGATCTGGAGGAATACCGGTGGCGAAGGCGGCCCCCTGGACAAAGACTGACGCTCATGCACGAAAGCGTGGGGAGCAAACAGG
## 1282                                                         ACGGAGGGTGCTAGCGTTGTTCGGAATCACTGGGCGTAAAGGGCGTGTAGGCGGTCTGCTAAGTCTGATGTGAAATCCCCTCGCTCAACGAGGGAACTGCGTCGGATACTGGTAGACTTGAGTACCGGAGAGGAAGGTGGAATTCCCGGTGTAGCGGTGAAATGCGTAGATATCGGGAGGAACATCGGTGGCGAAGGCGGCCTTCTGGACGGATACTGACGCTGAGACGCGAAAGCGTGGGGAGCAAACAGG
## 1283                                                         ACGGAGGGTGCAAGCGTTAATCGGAATAACTGGGCGTAAAGCGCACGCAGGCGGTTTGCTAAGCTAGATGTGAAAGCCCCGGGCTCAACCTGGGAACTGCATTTAGAACTGGCGAACTAGAGTCTTGTAGAGGGGGGTAGAATTCCAGGTGTAGCGGTGAAATGCGTAGAGATCTGGAGGAATACCGGTGGCGAAGGCGGCCCCCTGGACAGAGACTGACGCTCAGGTGCGAAAGCGTGGGGAGCAAACAGG
## 1284                                                         ACGTAGGTGGCAAGCGTTGTCCGGAATTATTGGGCGTAAAGCGCGCGCAGGCGGTCCTTTAAGTCTGATGTGAAAGCCCACGGCTCAACCGTGGAGGGTCATTGGAAACTGGGGGACTTGAGTGCAGAAGAGGAAAGTGGAATTCCACGTGTAGCGGTGAAATGCGTAGAGATGTGGAGGAACACCAGTGGCGAAGGCGACTTTCTGGTCTGTAACTGACGCTGAGGCGCGAAAGCGTGGGGAGCAAACAGG
## 1285                                                         ACGAAGGGGGCTAGCGTTACTCGGAATTACTGGGCGTAAAGGGCGCGTAGGCGGCTCTGTAAGTCAGGCGTGAAATTCCTGGGCTCAACCTGGGGGCTGCGCTTGAGACTGTGGGGCTAGAGGATGGAAGAGGGTCGTGGAATTCCCAGTGTAGAGGTGAAATTCGTAGATATTGGGAAGAACACCGGTGGCGAAGGCGGCGACCTGGTCCATTACTGACGCTGAGGCGCGACAGCGTGGGGAGCAAACAGG
## 1286                                                       ACAGAGGGTGCAAGCGTTATCCGGATTTATTGGGCGTAAAGCGTTTCGTAGGTGGTTTTGTAAGTTAAACTTCAAAGACTCGGGCTCAACTCGGGGAAGGGGTTTAATACTGCAGGACTTAGAGAGTTTTAGGGGCAATCGGAACTGTTGGTGTAGGGGTGAAATCCGTTGATATCAACAGGAACGTCAAGGGCGTAGGCAGATTGCTGGGAAATTTCTGACACTGAGGAACGAAAGCTAGGGGAGCGAAAGGG
## 1289                                                         ACGAAGGACCCGAGCGTTATCCGGATTTACTGGGTTTAAAGGGTGCGTAGGTGGACTTTTAAGTCAGTGGTGAAAGCCTCCCGCTCAACGGGAGAACTGCCATTGATACTGATAGTCTTGAGTATAGTTGAGGTAGGCGGAATATGTCATGTAGTGGTGAAATACTTAGATATGACATAGAACACCAATTGCGAAGGCAGCTTGCTAAGCTATAACTGACACTGAGGCACGAAAGCGTGGGGAGCAAACAGG
## 1291                                                         ACAGAGGGTGCAAGCGTTGCTCGGATTTACTGGGCGTAAAGCGTACGCAGACGGCATCAAAGGTCGGGTGTGAAAGCCCGGGGCTCAACCCCGGAATTGCATCCGAAACCGTGAAGCTAGAGTATGGGAGAGGGAAGCGGAATTCCTGGTGTAGAGGTGAAATTCGTAGATATCAGGAGGAACATCTGTGGCGAAGGCGGCTTCCTGGCCCATTACTGACGTTCAGGTACGAAAGCGTGGGTAGCAAACAGG
## 1292                                                         ACGGGAGTGGCAAGCGTTATCCGGAATTATTGGGCGTAAAGCGTCCGCAGGCGGCCGTACAAGTCTGTCGTTAAAGCGTGGAGCTTAACTCCATTTAAGCGATGGAAACTGTAAGGCTAGAGTGTGGTAGGGGCAGAGGGAATTCCCGGTGTAGCGGTGAAATGCGTAGATATCGGGAAGAACACCAGTGGCGAAGGCGCTCTGCTGGGCCATAACTGACGCTCATGGACGAAAGCCAGGGGAGCGAAAGGG
## 1293                                                         ACGTAGGGGGCGAGCGTTGTCCGGAATTACTGGGCGTAAAGCGCACGCAGGCGGGTTGTTAAGTCGGCTGTGAAAAGCACGGGCTCAACCTGTGTCTTGCGGTCGATACTGGCAGCCTCGAGTACGAGAGAGGGAAGTGGAATTCCCGGTGTAGCGGTGAAATGCGTAGATATCGGGAGGAACACCAGTGGCGAAGGCGGCTTCCTGGCTCGTGACTGACGCTCATGTGCGAAAGCTGGGGGAGCGAACGGG
## 1294                                                         ACGAGGGGTCCTAGCGTTGTTCGGAATCATTGGGCGTAAAGCGCTCGTAGGTGGCTTAGTAAGTCGGGTGTGAAAGCCCCGGGCTCAACCCGGGAAGTGCACTCGAAACTGCATAGCTTGAGTACGGTAGAGGTGAGTGGAATTCCAGGTGTAGTGGTGAAATACGTAGATATCTGGAGGAACATCGGTGGCGAAGGCGGCTCACTGGACCAGTACTGACACTGAGGAGCGAAAGCATGGGGATCAAACAGG
## 1295                                                         ACGAGAGGTGCAAGCGTTATTCGGAATTATTGGGCGTAAAGGGCGCGTAGGCGGCGCTATAAGTTTTTCGTTTAATTCTCCAGCCTAACTGGAGCCATGCGAAAAAGACTGTAGTGCTAGAGTATAATAGAGAGAAGTGGAATTTTCGGAGTAGCGGTTAAATGCGTAGATCTCGAAAGGAACGCCAATGGCGAAGGCAGCTTCTTGGGTTATTACTGACGCTGAGGCGCGAAGGCATGGGGAGCAAACAGG
## 1297                                                         ACGAGGGATCCTAGCGTTGTTCGGAATCATTGGGCGTAAAGCGGATGTAGGTGGCTTTGTAAGTCAGGTGTGAAAGCCCCGGGCTCAACCCGGGAAGTGCATTTGATACTGCGAAGCTTGAGTGCTGGAGAGGTTACTAGAATACCTGGTGTAGTGGTGAAATACGTAGATATCAGGTGGAATACCGGAGGCGAAGGCGGGTAACCAGCCAGACACTGACACTCAGATCCGAAAGTGTGGGGATCAAACAGG
## 1298                                                         ACAGAGGGTGCAAACGTTGCTCGGAATTACTGGGCGTAAAGGGCGTGTAGGCGGGAAGGTAAGTCGGGCGTGAAATCCCCAAGCTCAACTTGGGAACTGCACTCGAAACTACCTTTCTTGAGTGCCGGAGAGGAAAGCGGAATTCCTGGTGTAGAGGTGAAATTCGTAGATATCAGGAGGAACACCAGTGGCGAAGGCGGCTTTCTGGACGGTGACTGACGCTGAGACGCGAAAGCGTGGGGAGCAAACAGG
## 1299                                                         ACGGAGGGTGCAAGCGTTACTCGGAATTACTGGGCGTAAAGCGCGTGTAGGTGGTTTTGTAAGTCTGGTGTGAAATCCCGGGGCTCAACCCCGGAAGTGCATTGGATACTGCAAGACTAGAGTGCAGGAGAGGATGGTGGAATTCCAGGTGTAGAGGTGAAATTCGTAGATATCTGGAGGAACATCGGAGGCGAAGGCGACCATCTGGACTGCAACTGACACTGAGACGCGAAAGCGTGGGGAGCAAACAGG
## 1300                                                         ACATAGGGTGCAAGCGTTGTCCGGAATTATTGGGCGTAAAGAGCTCGTAGGTGGTTCGATACGTCGGATGTGAAAATCAGGGGCTCAACCCCTGACCTGCATTCGATACGGTCGAGCTAGAGTTTGGTAGGGGAGACTGGAATTCCTGGTGTAGCGGTGGAATGCGCAGATATCAGGAGGAACACCGATGGCGAAGGCAGTCCCCTGGGATAAGACTGACGCTCATGCACGAAAGCGTGGGGAGCAAACAGG
## 1301                                                         ACGAGGGGTCCTAGCGTTGTTCGGAATCACTGGGCGTAAAGCGCATGTAGGCGGCTTTGTAAGTCAGTTGTGAAAGCCCTGGGCTTAACCCAGGAAGTGCATCTGATACTGCAAAGCTTGAGTATGGGAGAGGGTAGTAGAATTCCTGGTGTAGTGGTGAAATACGTAGATATCAGGAGGAATACCGGTGGCGAAGGCGGCTACCTGGCCCAATACTGACGCTGAGATGCGAAAGCGTGGGGATCAAACAGG
## 1302                                                         ACGTAGGGTGCGAGCGTTAATCGGAATTACTGGGCGTAAAGGGTGCGTAGGCGGATATATAAGTTAGATGTGAAAAGCCTGGGCTTAACCTAGGTCGTGCATTTAAGACTGTATGTCTAGAGTTTGTAAGAGGGGGGTAGAATTCCAAGTGTAGCAGTGAAATGCGTAGAGATTTGGAGGAATACCGATGGCGAAGGCAGCCCCCTGGTATAGAACTGACGCTGAGGCACGAAAGCGTGGGTAGCAAACAGG
## 1304                                                         ACAGAGGGGGCAAGCGTTATTCGGAATTATTGGGCGTAAAGGGCGCGTAGGCGGCCTTGTAAGTCTTGGGTGAAATCCCTCAGCTCAACTGAGGAACTGCCTCGGATACTGCAGGGCTTGAGGCCGGGAGAGGGTAGTGGAATTCCCAGTGTAGCGGTGAAATGCGTAGATATTGGGAGGAACACCGGTGGCGAAGGCGGCTACCTGGACCGGTTCTGACGCTGATGCGCGAAAGCGTGGGGAGCAAACAGG
## 1306                                                       ACGGGAGGAACAAGCGTTACTCATGGTAACTTGGTGTTTAGGGTGCCGCGGCGGTTTGTGTGGAAAGAAAGGGAGAAAGCACAGCAGGAATGGTGGAAGGTCCTTGGTAATCCGCAAACTAGAGTTTGAATAGGGGATATGGTTATGGGTAGGGAAGCAATGAAATGCCGAGAAACTACTCGGAGCTCCAGCGGCGCAGGCGCGTATCCATCTGAAACTGACGCTAACGCACTAAAGCGTGGGGTTCAAACAGG
## 1308                                                         ACGTAGGTGGCGAGCGTTACTCGGAATTACTAGGCGTAAAGCGCGTGTAGGCGGGAGGTTAAGTCTGCTGTTAAATCTCATGGCTCAACCATGAAATGCCGGCGGATACTGGCCTTCTTGAATTCGGTAGAGGTGACTGGAATTCCGTGTGTAGCGGTGAAATGCGTAGATATACGGAAGAACGCCAAAGGCGAAGGCAGGTCACTGGGCCGCAATTGACGCTGAGACGCGAAAGCTAGGGGAGCAAACAGG
## 1310                                                         ACATAGGCTTCAAGCGTTGTCCGGATTTATTGGGCGTAAAGAGTTCGTAGGCGGTCGAGTAAGTCGGGTGTGAAAATTCTGGGCTCAACCCAGAGACGCCACCCGATACTGCTTAACTTGAGTTCGATAGGGGAGTGGGGAATTCCTAGTGTAGCGGTGAAATGCGCAGATATTAGGAGGAACACCGGTGGCGAAGGCGCCACTCTGGATCGACACTGACGCTGAGGAACGAAAGCATGGGTAGCAAACAGG
## 1311                                                         ACAGAGGGTGCAAGCGTTGCTCGGAATCATTGGGCGTAAAGGGCAAGTAGGTGGTCTCGTTTGTCTGAGGTGAAATCCCTGGGCTTAACCTAGGAACTGCCTCAGAAACGGCGAGACTGGAGTCTCGGAGAGGGTCGCGGAATTCCCGGTGTAGCGGTGAAATGCGTAGAGATCGGGAGGAATACCAGAGGCGAAAGCGGCGACCTGGACGAGTACTGACACTCAACTGCGAAAGCGTGGGGAGCAAACAGG
## 1313                                                        ACAGAGGGTGCAAGCGTTATCCGGATTTATTGGGCGTAAAGCGTTTCGTAGGTTGTTTTGTAAGTTATGTTTTAAAGACCACGGCTTAACCGGGGGAAGGGACATAATACTGCAAGACTTGAGAATTTTAGGGGCAATCGGAACTGTTGGTGTAGGGGTGAAATCCGTTGATATCAACAGGAACGTCAAGGGCGTAGGCAGATTGCTGGGAAATATCTGACACTGAGGAACGACAGCTAAGGGAGCGAAAGGG
## 1314                                                       ACGTGGGGTGCAAGCGTTATCCGGATTTACTGGGCGTAAAGCGTTCTGTAGGCGGTTTTGTAAGTATATGGTTAAAGGCTGCGGCTCAACCGTAGACATGCTGTATAAACTACAATGACTAGATGGTTGTCGGGAGTATCGGAATTCTCGGTGGAGGGGTGAAATCCGTTGATATCGAGAGGAACACCAAAGGCGAAGGCAGATACTTAGGCAATTTATGACGCTGATAGACGACAGCTAGGGGAGCGAAAGGG
## 1315                                                         ACGAAGGATGCTAGCGTTACTCGGAATTACTGGGCGTAAAGAATATGTAGGCTGACTGTTAAGTTGATAGTGAAATCCCCGAGCCTAACTCGGGAACTGCTCTCAAAACTGATAGTCTAGAATATGGTAGAGAGTAAGGGAATTCTTGGTGTAGGGGTAAAATCCGTAGATATCAGGAGGAAGACCGGAAGCGAAAGCGCTTACTTGGGCCATCATTGACGCTGAGATATGAAAGCGTGGGGATCGAACAGG
## 1316                                                         ACGAAGGGTGCAAGCGTTACTCGGAATTACTGGGCGTAAAGCGTGCGTAGGTGGTTGTTTAAGTCTGATGTGAAAGCCCTGGGCTCAACCTGGGAATGGCATTGGATACTGGATAGCTAGAGTGCGGTAGAGGGTAGTGGAATTCCCGGTGTAGCAGTGAAATGCGTAGAGATCGGGAGGAACATCCGTGGCGAAGGCGACTACCTGGACCAGCACTGACACTGAGGCACGAAAGCGTGGGGAGCAAACAGG
## 1317                                                         ACGAAGGACCCAAGCGTTATCCGGATTCATTGGGTTTAAAGGGTGCGTAGGCGGATTGATAAGTCAGTGGTGAAAGCCCGCAGCTTAACTGCGGAACTGCCATTGATACTGTTGATCTTGAATGTGTTTGATGTGGGCGGAATATGACATGTAGCGGTGAAATGCTTAGATATGTCATAGAACACCGATTGCGAAGGCAGCTCACAAAGCCACTATTGACGCTGAGGCACGAAAGCGTGGGGATCAAACAGG
## 1320                                                         ACGTAGGTGGCGAGCGTTATTCGGAATCACTGGGCGTAAAGGGTCCGTAGGCGGTCTGTCAAGTCTGATGTGAAATCTTGGGGCTCAACCCCAAACGTGCATTGGGTACTGACGGACTGGAGTGCATCAGAGGCAAGCGGAATTTGTGGTGTAGCGGTGGAATGCGTAGATATCACAAGGAACGCCAAAGGCGAAAGCAGCTTGCTGGGATGTCACTGACGCTGAGGGACGAAGGTCAGGGGAGCAAAAAGG
## 1321                                                         ACAGAGGGTGCAAGCGTTATTCGGAATTACTGGGCGTAAAGAGCGCGTAGGCTGTTTGATAAGTTTGATGTGAAAGCCCTGGACTCAATTCAGGAAGTGCATCGAATACTGTCAGACTAGAGTATAGGAGAGGAGAACGGAATTCCTGGTGTAGAGGTGAAATTCGTAGATATCAGGAGGAACAACAGTGGCGAAGGCGGTTCTCTGGACTATTACTGACGCTGAGGCGCGAAAGTGTGGGGAGCAAACAGG
## 1323                                                         ACGTAGGGAGCGAGCGTTGTCCGGAATTACTGGGTGTAAAGGGAGTGTAGGCGGGAAAACAAGTCAGATGTGAAAACTATGGGCTCAACCTGTAGACTGCATTTGAAACTATTTTTCTTGAGTGAAGTAGAGGTAAGCGGAATTCCTAGTGTAGCGGTGAAATGCGTAGATATTAGGAGGAACATCAGTGGCGAAGGCGGCTTACTGGGCTTTTACTGACGCTGAGGCTCGAAAGCGTGGGGAGCAAACAGG
## 1326                                                         ACGAACTGTGCGAACGTTATTCGGAATCACTGGGCTTACAGGGTGCGTAGGCGGTTTTCCAAGTCAGGTGTGAAATCCCCCAGCTCAACTGGGGAACTGCGCTTGAAACTGGATCACTTGAGGATGCTAGAGGTAGACGGAACTTCCGGTGGAGCGGTGAAATGTGTTGAGATCGGAAGGAACACCAGAGGCGAAGGCGGTCTACTGGGGCATATCTGACGCTGAGGCACGAAAGCCAGGGGAGCAAACGGG
## 1328                                                        ACGTAGGATCCAAGCGTTATCCGGATTTATTGGGCGTAAAGAGTTGCGTAGGCGGCAGAGTAAGCAGGTAGTGAAAGAGTATGGCTTAACCATATAAACATTACTTGAACTGCTCAGCTAGAGAGCGGAAGAGGTAGCTGGAATTATTAGTGGAGGAGTAATATCCGTAGATATTAGTAAGAACACCAATGGCGAAGGCAGGCTACTGGTCCGTTTCTGACGCTCAGGCACGAAAGCGTAGGTAGCAAACAGG
## 1330                                                         ACGAAGGGGGCAAGCGTTGTTCGGAATTACTGGGCGTAAAGCGCGCGTAGGCGGCGTCGTCAGTCAGAGGTGAAATCCCAGGGCTCAACCTTGGAATTGCCTTTGATACTGCGATGCTTGAGTTCGAGAGAGGGTGGCGGAATACCCAGTGTAGAGGTGAAATTCGTAGATATTGGGTAGAACACCAGTGGCGAAGGCGGCCACCTGGCTCGATACTGACGCTGAGGTGCGAAAGCGTGGGGAGCAAACAGG
## 1331                                                         ACGAAGGTCCCAAGCGTTGTTCGGAATCATTGGGCGTAAAGCGAGTGCGGGTGGCTCTTTAAGTCAGAAGTGAAAGCCCAGAGCTTAACTCTGGAAGTGCTTTTGATACTGGAGAGCTTGAATGTGGCAGAGGGTGCTAGAATACCTAATGTAGTGGTGAAATACGTAGATATTAGGTGGAACACCGGTGGCGAAGGCGGGCACCTGGGCCAACATTGACACTTAGACTCGAAAGCGTGGGGATCAAACAGG
## 1332                                                         ACGGAGGGTGCAAGCGTTATCCGGAATCATTGGGTTTAAAGGGTCCGCAGGCGGATTTATAAGTCAGTGGTGAAAGCCTACAGCTTAACTGTAGAACTGCCATTGATACTGTAAGTCTTGAATTCGGTCGAAGTGGGCGGAATGTGTAGTGTAGCGGTGAAATGCTTAGATATTACACAGAACACCGATAGCGAAGGCAGCTCACTAGGCCTGAATTGACGCTCATGGACGAAAGCGTGGGGAGCAAACAGG
## 1335                                                         ACGTAGGTCTCGAGCGTTATCCGGAATTACTGGGCGTAAAGCGTCTGTAGCCGGGTATATAAGTCTGTCTTTAAATATCCTGATTCAATCGGGAAAAGGGGATAGATACTGTATACCTAGAGGATTTTAGAGGTTAGTGGAATTTCCGGTGGAGCGGTGAAATGTGTTGATATCGGAAGGAATGCCGAAAGCGAAAGCAGCTAACTATAAAATATCTGACGGTCAGAGACGAAAGTTTGGGTAGCAAACGGG
## 1336                                                           ACGAAAGGGGCAAGCGTTATCCGGAATTATTGGGCGTAAAGGGTTTGTAGGCGGTCCAAGAAGTATGCAATGAAAGCACTTTAAAAAGGAGTAACTGTTGCATAGCCCTTAGGACTTGAGGTCAAAAGACGAGAATCGAATTTCGGGTGGAGCGATAAAATGCAAAGATATCCGAAGGAAGACCAAAAGCGAAGGCAGTTCTCGAGGTTGAACCTGACGTTGAAAAACGAAAGCGTGGGGAGCAAACGGG
## 1337                                                         ACGTAGGGTGCAAGCGTTAATCGGAATTACTGGGCGTAAAGCGTGCGCAGGCGGTTTCGTAAGCTTGATGTGAAATCCCCGGGCTCAACCTGGGAACTGCATTGAGGACTGCGAGACTAGAATACGGCAGAGGGGGGTGGAATTCCACGTGTAGCAGTGAAATGCGTAGAGATGTGGAGGAACACCGATGGCGAAGGCAGCCCCCTGGGCCTGTATTGACGCTCATGCACGAAAGCGTGGGGAGCAAACAGG
## 1338                                                         ACGTAGGGGGCAAGCGTTATCCGGATTTACTGGGTGTAAAGGGAGCGTAGGCGGCAAGGCAAGTCTGATGTGAAAACCCGGGGCTCAACCCCGGGACTGCATTGGAAACTGCTTAGCTAGAGTGTCGGAGAGGTAAGTGGAATTCCTAGTGTAGCGGTGAAATGCGTAGATATTAGGAGGAACACCAGTGGCGAAGGCGACTTACTGGACGATAACTGACGCTGAGGCTCGAAAGCGTGGGGAGCAAACAGG
## 1339                                                         ACGAAGGGGGCGAGCGTTGTTCGGAATCACTGGGCGTAAAGGGCACGTAGGCGGCCGTGCAAGTTGGGCGTGAAAGCCCCGGGCTCAACCCGGGATGTGCGCTCAAGACTGCATGGCTCGAGTTCGGAAGAGGGTGGCGGAATTCCCAGTGTAGAGGTGAAATTCGTAGATATTGGGAAGAACACCGGTGGCGAAGGCGGCTACCTGGTCCGATACTGACGCTGAGGTGCGAAAGCGTGGGGAGCAAACAGG
## 1340                                                        CCCAGCGCCACGAGTGGTTATCACGTTTATTGGGCCTAAAGCGTTCGTAGCCGGTTTGGTACATCTCTTGTGAAATTGTTCTGCTTAACAGGACAGCGTGCAGGAGAGACGGCCAGACTCGAGACTGGGAGGGGTTAGAAGTATGTCGTGGGGACTGGTAAAATGGGATAATCCACGATAGACTACCGATGGCGAAGGCATCTAACCAGAACAGTTCTGACGGTGAGGAACGAAAGCCAGGGGAGCGAACCGG
## 1341                                                         ACGTAGGGTGCAAGCGTTAATCGGAATTACTGGGCGTAAAGCGTGCGCAGGCGGTTATGTAAGACAGAGGTGAAATCCCCGGGCTCAACCTGGGAACTGCCTTTGTGACTGCATAGCTAGAGTACGGTAGAGGGGGATGGAATTCCGCGTGTAGCAGTGAAATGCGTAGATATGCGGAGGAACACCGATGGCGAAGGCAGCCTCCTGGGCCAGTACTGACGCTCATGCACGAAAGCGTGGGGAGCAAACAGG
## 1343                                                         ACGGAGGATGCGAGCGTTATCCGGAATCATTGGGTTTAAAGGGTCCGTAGGCGGTTTAATAAGTCAGTGGTGAAATCTGGTCGCTCAACGATCAAACTGCCATTGATACTGTTAGACTTGAATAATTGTGAAGTAACTAGAATATGTAGTGTAGCGGTGAAATGCTTAGATATTACATGGAATACCAATTGCGAAGGCAGGTTACTAACAATTTATTGACGCTGATGGACGAAAGCGTGGGGAGCGAACAGG
## 1344                                                         ACGTAGGTGGCGAGCGTTATCCGGAATTATTGGGCGTAAAGAGCGCGCAGGTGGTTGATTAAGTCTGATGTGAAAGCCCACGGCTTAACCGTGGAGGGTCATTGGAAACTGGTCGACTTGAGTGCAGAAGAGGGAAGTGGAATTCCATGTGTAGCGGTGAAATGCGTAGAGATATGGAGGAACACCAGTGGCGAAGGCGGCTTCCTGGTCTGTAACTGACACTGAGGCGCGAAAGCGTGGGGAGCAAACAGG
## 1345                                                         ACGAAGGGTGCAAGCGTTGCTCGGAATTATTGGGCGTAAAGGGTAGGTAGGTGGTGATACAAGTCTGGAGTGAAATCCCTGAGCTCAACTCAGGACGTGCTTTGGAAACTGTATCACTAGAGTGCTAGAGAGGTTCGTGGAATTCCTGGTGTAGCGGTGAAATGCGTAGAGATCAGGAGGAACATCTGCGGCGAAGGCGGCGAACTGGATAGTTACTGACACTCAACTACGAAAGCGTGGGGAGCAAACAGG
## 1346                                                         ACGGAGGGTGCGAGCGTTAATCGGAATTACTGGGCGTAAAGGGTGCGTAGGTGGTTTGATAAGTTATCTGTGAAAGCCCTGGGCTCAACCTGGGAACTGCAGATAAGACTGTTAGACTCGAGTATAGGAGAGGGTAGTGGAATTTCCGGTGTAGCGGTGAAATGCGTAGAGATCGGAAGGAACACCAGTGGCGAAGGCGGCTACCTGGCCTAATACTGACACTGAGGCACGAAAGCGTGGGGAGCAAACAGG
## 1347                                                         ACGGAGGGAGCTAGCGTTGTTCGGAATTACTGGGCGTAAAGCGCACGTAGGCGGCGACTCAAGTCAGAGGTGAAAGCCTGGAGCTCAACTCCAGAACTGCCTTTGAAACTAGGTCGCTAGAATCTTGGAGAGGCGAGTGGAATTCCGAGTGTAGAGGTGAAATTCGTAGATATTCGGAAGAACACCAGTGGCGAAGGCGACTCGCTGGACAAGTATTGACGCTGAGGTGCGAAAGCGTGGGGAGCAAACAGG
## 1350                                                         ACAGAGAGTGCAAACGTTGTTCGGAATTACTGGGCTTAAAGCGCACGTAGGCTGCGTCGCAAGTGTCAGGTGAAATCCCACGGCTTACCCGTGGAACTGCCTGGCAGACTGCGATGCTAGAGTGTGGTAGGGGAGAGCGGAACTCCTGGTGGAGCGGTGAAATGCGTAGATATCAGGAGGAACACCGGTGGCGAAGGCGGCTCTCTGGTCCACAACTGACGCTGAGGTGCGAAAGCTAGGGGAGCAAACGGG
## 1352                                                         ACGAAGGGGGCTAGCGTTGCTCGGAATTACTGGGCGTAAAGGGAGCGTAGGCGGACTGTTTAGTCAGAGGTGAAAGCCCAGGGCTCAACCTTGGAATTGCCTTTGATACTGGCAGTCTTGAGTACGGAAGAGGTATGTGGAACTCCGAGTGTAGAGGTGAAATTCGTAGATATTCGGAAGAACACCAGTGGCGAAGGCGACATACTGGTCCGTTACTGACGCTGAGGCTCGAAAGCGTGGGGAGCAAACAGG
## 1353                                                         ACGAAGGGTGCAAGCGTTACTCGGAATTACTGGGCGTAAAGGGTGCGTAGGTGGTTTTTTAAGTCTGCTGTGAAATCCCCGGGCTCAACCTGGGAATGGCAGTGGATACTGGAAAGCTGGAGTGCGGTAGAGGGTAGTGGAATTCCCGGTGTAGCAGTGAAATGCGTAGAGATCGGGAGGAACACCAGTGGCGAAGGCGGCTACCTGGACCAGCACTGACACTGAGGCACGAAAGCGTGGGGAGCAAACAGG
## 1354                                                         ACGTAGGGTGCAAGCGTTAATCGGAATTACTGGGCGTAAAGCGTGCGCAGGTGGTTCGTTAAGACAGTTGTGAAATCCCCGGGCTTAACCTGGGAACTGCAATTGTGACTGGCGGACTAGAGTTTGGCAGAGGGGGGTGGAATTCCTGGTGTAGCAGTGAAATGCGTAGATATGCGGAGGAACACCGATGGCGAAGGCAATCCCCTGGGCCTGTACTGACGCTCATGCACGAAAGCGTGGGGAGCAAACAGG
## 1355                                                         ACAGAGGTCTCAAGCGTTGTTCGGAATCACTGGGCGTAAAGGGTGCGTAGGTGGCGTGGCAAGTCAGATGTGAAAGCCCGGGGCTCAACCTCGGAATTGCATCCGATACTGCCATGCTAGAGTACTGAAGAGGTGACTAGAATTCTAGGTGTAGCAGTGAAATGCGCAGATATCTAGAGGAATACCAACGGCGAAGGCAGGTCACTGGGCAGTTACTGACACTGAGGCACGAAGGCCAGGGGAGCAAACGGG
## 1356                                                         ACGTATGGAGCAAGCGTTATCCGGATTTACTGGGTGTAAAGGGAGTGTAGGTGGCCAGGCAAGTCAGAAGTGAAAGCCCGGGGCTCAACCCCGGGACTGCTTTTGAAACTGCAGGGCTAGAGTGCAGGAGGGGCAAGTGGAATTCCTAGTGTAGCGGTGAAATGCGTAGATATTAGGAGGAACACCAGTGGCGAAGGCGGCTTGCTGGACTGTAACTGACACTGAGGCTCGAAAGCGTGGGGAGCAAACAGG
## 1357                                                         ACGAGTACCCCGAGCATTATCCGGAATTATTGGGCGTAAAGGGTGAGTAGGCGGTGCAATTAGTCGTGTGTTAAATCCCAGAGCTTAACTTTGGATTCGCACACGAAACGGTTGCACTAGAGGGTGTGAGAGGTGAATGGAACTCATAGTGTAGGGGTGAAATCCGTTGATATTATGGGGAACACCAAATGCGAAGGCAATTCACTGGCACATTTCTGACGCTCAATCACGAAAGCGTAGGTAGCGAATGGG
## 1364                                                         ACGTAGGGTGCGAGCGTTGTCCGGAATTACTGGGCGTAAAGAGCTCGTAGGTGGTTTGTCGCGTTGTTCGTGAAATCTCACGGCTTAACTGTGAGCGTGCGGGCGATACGGGCAGACTGGAGTACTGCAGGGGAGACTGGAATTCCTGGTGTAGCGGTGGAATGCGCAGATATCAGGAGGAACACCGGTGGCGAAGGCGGGTCTCTGGGCAGTAACTGACGCTGAGGAGCGAAAGCGTGGGGAGCGAACAGG
## 1365                                                         ACGAAGGTGGCGAGCGTTACCCGGATTGACTGGGCGTAAAGCGTCCGCAGACGGACAACCAAGTCTTTGGTTAAATCCTGAGGCTCAACCTCAGGACCGCTAAAGATACTGTTTGTCTAGAGGCTGGGAGAGGCAAGCGGAATTACTGGTGTAGTAGTAAAATGCGTTAATATCAGTAAGAACACCAAAAGCGAAGGCAGCTTGCTAGAACAGTCCTGACGTTCAGGGACGAAAGCGTGGGGAGCGAATGGG
## 1368                                                         ACGGAGGGTGCAAGCGTTATCCGGAATCACTGGGTTTAAAGGGTGCGTAGGCGGATTATTAAGTCAGTGGTGAAATCCTGTCGCTTAACGATAGAATTGCCATTGATACTGATAATCTTGAATTGGGTTGAGGTTAGCGGAATGTGACATGTAGCGGTGAAATGCATAGATATGTCATGGAACACCAATTGCGAAGGCAGCTAGCTGGGCCTTGATTGACGCTGAGGCACGAAAGCGTGGGTAGCGAACAGG
## 1369                                                         ACAGAGGGGGCTAGCGTTGTTCGGAATGACTGGGCGTAAAGGGCGCGTAGGTGGTATGACAAGTCAGGTGTGAAAGACCCAGGCTTAACCTGGGAACGGCATTTGAGACTGTCATGCTTGAGTCTGGGAGAGGATAGCGGAATTCCTAGTGTAGAGGTGAAATTCGTAGATATTAGGAAGAACACCGGTGGCGAAGGCGGCTATCTGGAACAGAACTGACACTGAGGCGCGAAAGCGTGGGGAGCAAACAGG
## 1370                                                         ACGGAGGGTGCAAGCGTTGTTCGGAATTACTGGGCGTAAAGCGTACGTAGGCGGTTGAATAAGTCAAATGTTAAAGCCCGGGGCTCAACCCCGAGGATGCATTTGATACTGTTCGACTAGAGGGCGATAGAGGAAAGCGGAATTCCGAGTGTAGCGGTGAAATGCGTAGATATTCGGAGGAACATCGGTAGCGAAAGCGGCTTTCTGGGTCGCTCCTGACGCTGAGGTACGAAAGCGTGGGTAGCAAACGGG
## 1371                                                         ACAGAGGCCACGAGCGTTAGTCGGAATCACTGGGCTTAAAGGGTGCGTAGGCGGGCCCGCAAGCGTCTTGTGAAATCCCATGGCTCAACCATGGAATCGCAGGGCGAACTGCGGGTCTTGAGGCAGGTAGGGGCCGGAAGAACAGTAGGTGGAGCGGTGAAATGCGTAGATATCTACTGGAATGCCGATGGGGAAGCCGTCCGGCTGGGCCTGTCCTGACGCTGAGGCACGAAAGCGTGGGGAGCAAACAGG
## 1372                                                         ACGGAGGGTGCAAACGTTGCTCGGAATTATTGGGCGTAAAGCGCACGTAGGCGGTTCGTTATGTCGGATGTGAAAGCCCTGGGCTTAACCCAGGAAGTGCATCCGAAACTGGCGAGCTTGAGTACGGAAGAGGGTCTCGGAATTCCCGGTGTAGAGGTGAAATTCGTAGATATCGGGAGGAACACCAGTGGCGAAGGCGGAGACCTGGGCCGATACTGACGCTGAGGTGCGAAAGCGTGGGGAGCAAACAGG
## 1374                                                        ACAGAGACCCCAAGCGTTATCCGGAATTATTGGGCGTAAAGGGTGAGTAGGTGGTTTTATTAGTCAGATGTTAAATTTCTCGGGCTTAACCTGAGAACTGCATTTGAAACGGTAGAACTAGAATTTTGGAGAGGTCTGTGGAACTATATGTGTAGGGGTAAAATCCGTTGATATATATGGGAACACCTAAAGCGAAGGCAGCAGACTGGCCAAACATTGACACTGAATCACGAAAGCGTGGGTAGCGATACGG
## 1376                                                        ACGGGGGGTGCAAGCGTTGTCCGGAATGATTGGGCGTAAAGCATCTGTAGGCGGTTTCTAAAGTCAACTGTTAAAGCCCAGGGCTTAACTTTGGAAAAGCGGTTGAGTACTTAGGAACTTGAGTTTGGTAGGGGTAAAGGGAATTCCTAGTGGAGCGGTGAAATGCACAGAGATTAGGAAGAACACCGATGGCGAAAGCACTTTACTGGGCCAAAACTGACGCTGAGAGATGAAAGTTGGGGGAGCGAATAGG
## 1377                                                         ACGTAGGGTGCGAGCGTTAATCGGAATTACTGGGCGTAAAGCGTGCGCAGGCGGCGTCGCAAGTCAGGCGTGAAATCCCCGAGCTTAACTTGGGAATTGCGCTTGAAACTACGATGCTTGAGTGTGGCAGAGGGAGGTGGAATTCCACGTGTAGCGGTGAAATGCGTAGATATGTGGAGGAACACCGATGGCGAAGGCAGCCTCCTGGGCCAACACTGACGCTCATGCACGAAAGCGTGGGGAGCAAACAGG
## 1378                                                         ACAGAGACTGCAAGCGTTACTCGGATTCACTGGGCGTAAAGGGAGCGCAGGCGGGCGGGTGTGTCATATGTGAAATCCCGGGGCTCAACCCCGGGGCTGCATCTGAAACTACCTGTCTAGAGATTCGGAGGGGTAAGCGGAATTCCTGGTGGAGCAGTGAAATGCGTAGATATCAGGAGGAACACCAACGGCGAAGGCAGCTTACTGGACGAAATCTGACGCTCAGGCTCGAAAGCATGGGGAGCAAACAGG
## 1380                                                         ACAGAGACCTCGAGCGTTATCCGGAATTACTGGGCGTAAAGCGCGAGCAGCCGGTTGAGTTAGTCCGATGTTAAATCCCAAGGCTTAACTTTGGAACCGCATTGGAAACGGCTCAACTCGAGTGGGTAAGAGGCACGCGGAACGTGCAGTGTAGGGGTGAAATCCATTGATATTGCACGGAACACCAAAAGCGAAGGCAGCGTGCTAGTGCTCAACTGACGGTCATTCGCGAAAGCGTGGGGATCAAAAAGG
## 1381                                                         ACGTAGGGGGCAAGCGTTATCCGGATTTACTGGGTGTAAAGGGAGCGTAGACGGTGTGGCAAGTCTGATGTGAAAGGCATGGGCTCAACCTGTGGACTGCATTGGAAACTGTCATACTTGAGTGCCGGAGGGGTAAGCGGAATTCCTAGTGTAGCGGTGAAATGCGTAGATATTAGGAGGAACACCAGTGGCGAAGGCGGCTTACTGGACGGTAACTGACGTTGAGGCTCGAAAGCGTGGGGAGCAAACAGG
## 1382                                                        ACGAACCGTCCAAACGTTATTCGGAATCACTGGGCTTAAAGGGCGCGTAGGCGGCCGAGCGGGTCGTGGGTGAAATCCCCCAGCTTAACTGGGGAACTGCCCTTGATACCACTGCGGCTCGAGGAAGGAAGGGGCAATCGGAACTGTCGGTGGAGCGGTGAAATGCGTTGATATCGACAGGAACGCCGGTGGCGAAAGCGGATTGCTGGTCCTTTTCTGACGCTGAGGCGCGAAAGCCAGGGGAGCAAACGGG
## 1384                                                         ACGAAGGGGGCTAGCGTTGTTCGGATTTACTGGGCGTAAAGCGCTCGTAGGTGGTTTGACAAGTCAGTGGTGAAATCCCAAGGCTCAACCTTGGAACTGCCATTGAAACTGTCAGACTGGAATCTCAGAGGGGGTGGCGGAATTGCTAATGTAGGGGTGAAATCCGTAGATATTAGCAGGAACACCGATGGCGAAGGCAGCCACCTGGATGAGTATTGACACTGAGGAGCGAAAGCGTGGGGATCAAACAGG
## 1385                                                         ACGGAGGATCCAAGCGTTATCCGGATTTATTGGGTTTAAAGGGTGCGTAGGCGGAATAGTAAGTCAGTGGTGAAAGCCTGTCGCTTAACGACAGAACTGCCATTGATACTGCTAATCTTGAGTATAATTGAAGTGGGCGGAATGTGTCGTGTAGCGGTGAAATGCTTAGATATGACACAGAACACCAATTGCGAAGGCAGCTCACTAAACTATAACTGACGCTGAGGCACGAAAGCGTGGGGATCAAACAGG
## 1387                                                         ACAGAGGTCTCAAGCGTTGTTCGGAATCACTGGGCGTAAAGGGTGCGTAGGTGGCGTGGTAAGTCAGATGTGAAAGCCCGGGGCTCAACCTCGGAACTGCATCCGATACTACCATGCTAGAGTACTGAAGAGGAAACTAGAATTCTCGGTGTAGCAGTGAAATGCGTAGATATCGAGAGGAATACCAAAGGCGAAGGCAGGTTTCTTGGCAGTTACTGACACTGAGGCACGAAGGCCAGGGGAGCGAACGGG
## 1388                                                         ACGTAGGGGGCAAGCGTTATCCGGATTTACTGGGTGTAAAGGGAGCGTAGACGGCAAGGCAAGTCTGATGTGAAAACCCAGGGCTTAACCCTGGGACTGCATTGGAAACTGTCTGGCTCGAGTGCCGGAGAGGTAAGCGGAATTCCTAGTGTAGCGGTGAAATGCGTAGATATTAGGAAGAACACCAGTGGCGAAGGCGGCTTACTGGACGGTAACTGACGTTGAGGCTCGAAAGCGTGGGGAGCAAACAGG
## 1389                                                         ACGTAGGATGCAAGCGTTGTCCGGATTTATTGGGCGTAAAGAGTTCGTAGGCGGTTTGTTAAGTCTGATGTTAAAGACCGGGGCTCAACCTCGGAAATGCATTGGATACTGGCAGACTGGAGTGCAGTAGAGGCTAGTGGAATTCCCAGTGTAGCGGTGAAATGCGTAGATATTGGGAAGAACACCGGTGGCGTAGGCGACTAGCTGGGCTGTAACTGACGCTGAGGAACGAAAGCCAGGGGAGCGAATGGG
## 1391                                                         ACGTAGGGTGCAAGCGTTGTCCGGAATTATTGGGCGTAAAGAGCTCGTAGGCGGTTTGTCACGTCTGCTGTGAAATTTCGAGGCTCAACCTCGAACTTGCAGTGGGTACGGGCAGGCTAGAGTGCGGTAGGGGAGATGGGAATTCCTGGTGTAGCGGTGGAATGCGCAGATATCAGGAGGAACACCAATGGCGAAGGCACATCTCTGGGCCGTAACTGACGCTGAGGAGCGAAAGCGTGGGGAGCAAACAGG
## 1394                                                         ACGAAGGGGGCTAGCGTTGCTCGGAATTACTGGGCGTAAAGGGCGCGTAGGCGGATAGTTTAGTCAGAGGTGAAAGCCCAGGGCTCAACCTTGGAATTGCCTTTGATACTGGCTATCTTGAGTATGGAAGAGGTATGTGGAACTCCGAGTGTAGAGGTGAAATTCGTAGATATTCGGAAGAACACCAGTGGCGAAGGCGACATACTGGTCCATTACTGACGCTGAGGCGCGAAAGCGTGGGGAGCAAACAGG
## 1396                                                         ACAGAGGGTGCGAACGTTGCTCGGAATTACTGGGCGTAAAGCGCGTGTAGGCGGGCTAGCAAGTCAGATGTGAAATCCCCGGGCTCAACCCGGGAACTGCATCTGAAACTGCTGGCCTTGAGTGCCGGAGAGGGTGGCGGAATTCCTGGTGTAGAGGTGAAATTCGTAGATATCAGGAGGAACATCAGTGGCGAAGGCGGCCACCTGGACGGCAACTGACGCTGAGACGCGAAAGCGTGGGGAGCAAACAGG
## 1397                                                         ACGTAGGGTGCAAGCGTTAATCGGAATTACTGGGCGTAAAGCGTGCGCAGGCGGTTATATAAGACAGATGTGAAATCCCCGGGCTCAACCTGGGAACTGCATTTGTGACTGTATAGCTAGAGTACGGTAGAGGGGGATGGAATTCCGCGTGTAGCAGTGAAATGCGTAGATATGCGGAGGAACACCGATGGCGAAGGCAGTCCCCTGGGATAAGACTGACGCTCATGCACGAAAGCGTGGGGAGCAAACAGG
## 1400                                                         ACGGAGGGTGCGAGCGTTATCCGGAATCACTGGGTTTAAAGGGTGCGTAGGCGGCTAGGTAAGTCAGGGGTGAAAGCTTCCCGCTCAACGGGAGAACTGCCTTTGATACTGCCTGGCTCGAATTGGGTTGAGGCTGGCGGAATGTGGCATGTAGCGGTGAAATGCTTAGATATGCCATAGAACACCGATTGCGAAGGCAGCCAGCCAAGCCTTGATTGACGCTGAGGCACGAAAGCGTGGGGAGCGAACAGG
## 1401                                                          ACGTAGGTCCCGAGCGTTGTCCGGATTTATTGGGCGTAAGGCGAGCGCAGGCGGTTTGATAAGTCTGAAGTTAAAGGCTGTGGCTCAACCATAGTTCGCTTTGGAAACTGTCAAACTTGAGTGCAGAAGGGGAGAGTGGAATTCCATGTGTAGCGGTGAAATGCGTAGATATATGGAGGAACACCGGTGGCGAAAGCGGCTCTCTGGTCTGTAACTGACGCTGAGGCTCGAAAGCGTGGGGAGCAAACAGG
## 1403                                                        CCCAGCGGTCCAAGTCGCAGCCACATTTGTTGGGTCTAAAACATCCGTAGCTTGCTTTTCAAGTCTTTTGTGAAATCGGGACTCTTAAGGTTCCGGCGTGCAAAAGATACTGTTAAGCTAGAGACCGGGAGGCGTAAGAAGTACGTATAGGGTAGCGGTAAAATGCGTTGATCCTATGTGGACTCACAGTCGCGAAGGCGTCTTACGAGAACGGTTCTGACAGTGAGGGATGAAGGCTAGGGGCGCAAAACGG
## 1404                                                        ACGAGGGGAGCAAGTGTTATACAGTCAAACTAGGCGTAAAGGGTTCTGCGGCGGAGTAATGAGAGATCTTCAGAGATACGGTGGTTGTTACCCATCTGCATGAAGATGACCATTTCTCTTAGTGTTATGTAGGGGTTGGAAGGACTCCGCGCAAAGCGTTGCAATGCGATGACCCGCGGGGGCCAACCAGTAGAGAAGTCGTCCAACTAGACATGACAGACGCTTAGGAACAAAAGCATGGGAAGCAAATGGG
## 1406                                                         ACGTAGGGTGCAAGCGTTAATCGGAATTACTGGGCGTAAAGCGTGCGCAGGCGGTTATGCAAGACAGATGTGAAATCCCCGGGCTCAACCTGGGAACTGCATTTGTGACTGCATAGCTAGAGTACGGTAGAGGGGGATGGAATTCCGCGTGTAGCAGTGAAATGCGTAGATATGCGGAGGAACACCGATGGCGAAGGCAATCCCCTGGGCCTGTACTGACGCTCATGCACGAAAGCGTGGGGAGCAAACAGG
## 1408                                                         ACGAACCGTGCGAACGTTGTTCGGAATCACTGGGCTTAAAGGGTGCGTAGGCGGAATGGCAAGTCGGAGGTGAAAGCCCCCAGCTTAACTGGGGAAGTGCCCTCGATACTGCTGTTCTCGAGAGAGGTAGGGGCGTGTGGAACTTCCGGTGGAGCGGTGAAATGCGTAGATATCGGAAGGAACGCCGGTGGCGAAAGCGATGCGCTGGACCTTTTCTGACGCTGAGGCACGAAAGCTAGGGGAGCGAACGGG
## 1409                                                         ACGAGGGGTGCAAGCGTTGTTCGGAATCACTGGGCGTAAAGCGCGTCTAGGCGGGACAGAAAGTCTGATGTGAAAGTTCTTGGCTCAACCAAGAAAATGCGTCGGAGACTTCTGATCTTGAGTCATGGAGGGATGGGCGGAATTCCGGGTGTAGCAGTGAAATGCGTAGAGATCCGGAGGAACATCAGAGGCGAAGGCGGCTCATTGGCCATGTACTGACGCTAAAGCGCGAAAGCGTGGGTAGCAAACAGG
## 1410                                                         ACGGAGGGTGCAAGCGTTATCCGGATTCACTGGGTTTAAAGGGTGCGTAGGTGGATCAGTAAGTCAGTGGTGAAAGCCCCGGGCTCAACTTGGGAATTGCCATTGATACTGCTGGTCTTGAGTTCGGTTGAGGTAGGCGGAATAAGTCATGTAGCGGTGAAATGCTTAGATATGACTTAGAACACCGATTGCGAAGGCAGCTTACCAAGCCGACACTGACACTGAGGCACGAAAGCGTGGGGATCAAACAGG
## 1412                                                         ACGTAGGGTGCGAGCGTTAATCGGAATTACTGGGCGTAAAGCGTGCGCAGGTGGTTGTGTAAGTCAGATGTGAAATCCCCGGGCTCAACCTGGGAACTGCATTTGAGACTGCACGGCTAGAGGATGGCAGAGGGGGGTAGAATTCCACGTGTAGCAGTGAAATGCGTAGAGATGTGGAGGAATACCGATGGCGAAGGCAGCCCCCTGGGCTATTTCTGACACTCATGCACGAAAGCGTGGGGAGCAAACAGG
## 1417                                                         ACGAAGGTGGCAAGCGTTACTCGGAATTACTAGGCGTAAAGGGCAGGTAGGTGGTTCGGCAAGTCTGTTGTGGAAGCTCCCGGCTTAACTGGGAGAGGCCAACGGAAACTGCCGGACTTGAGTGTAGGAGAGGGTATTGGAATTCCCGGTGTAGCGGTGAAATGCGCAGAGATCGGGAGGAACACCAATGGCGAAAGCAGATACCTGGACTATTACTGACACTAAGCTGCGAAAGCTAGGGGAGCAAACAGG
## 1418                                                         ACGTAGGGTGCGAGCGTTAATCGGAATTACTGGGCGTAAAGCGTGCGCAGGCGGTTGTATAAGTCAGATGTGAAATCCCCGGGCTCAACCTGGGAACTGCATTTGAGACTGTACGGCTAGAGTGTGGCAGAGGGGGGTGGAATTCCACGTGTAGCAGTGAAATGCGTAGAGATGTGGAGGAACACCGATGGCGAAGGCAGCCCCCTGGGCCAATACTGACGCTCATGCACGAAAGCGTGGGGAGCAAACAGG
## 1419                                                        ACGTAGGACCCAAGCGTTATCCGGAATTACTGGGCGTAAAGAGTTGCGTAGGCGGCTTGTTAAGCAAGATGTGAAATCGTGAGGCTCAACCTTATACACATATTTTGAACTGGCAAGCTAGAGTATAGAAGAGGTCACTGGAATTCCAAGTGTAGGAGTGAAATCCGTAGATATTTGGAGGAACACCGATGGCGTAGGCAGGTGACTGGTCTATTACTGACGCTAAGGCACGAAAGCGTGGGGAGCAAACGGG
## 1421                                                         ACGAAGGTGGCGAGCGTTACTCGGAATTACTAGGCGTAAAGCGTGGGCAGGCGGTTTGGTAAGTCTGCTGTGAAAGCCCCGGGCTCAACCCGGGGAGGTCAGTGGATACTGCCGGACTTGGATGTGGGAGAGGTGACTGGAATTCCTGGTGTAGCGGTGAAATGCGTAGATATCAGGAGGAACACCGATGGCGAAAGCAGGTCACTGGACCACTATCGACGCTCATCCACGAAAGCTGGGGGATCAAACAGG
## 1423                                                         ACGGAGGGTGCGAGCGTTTATCGGAATTACTGGGCGTAAAGCGCGCGTAGGCGGCTTGATAAGTCGGATGTGAAATCCCCGGGCTCAACCTGGGAACTGCATCCGATACTGTTGAGCTGGAGTACGATAGAGGATGGTAGAATTTCTAGTGTAGCGGTGAAATGCGTAGATATTAGAAGGAATACCGGTGGCGAAGGCGGCCATCTGGATCGATACTGACGCTGAGGTGCGAAAGCGTGGGGATCAAACAGG
## 1425                                                         ACAGAGAGTGCGAACGTTGTTCGGAATCACTGGGCATAAAGCGCACGTAGGCGGCACCATAAGTGCGAGGTGAAATCCCACGGCTTACCCGTGGAACTGCTTTGCAGACTGTGGTGCTCGAGGATGGTAGGGGAGAGCGGAACTCTTGGTGGAGCGGTGAAATGCGTAGAGATCAAGAGGAACACCGGTGGCGAAGGCGGCTCTCTGGTCCATATCTGACGCTGAGGTGCGAAAGCTAGGGGAGCAAACGGG
## 1427                                                         ACGTAGGTGGCAAGCGTTATCCGGATTTACTGGGTGTAAAGGGCGCGTAGGCGGGAAGGCAAGTCAGATGTGAAATCCAGGGGCTCAACCCCTGAACTGCATTTGAAACTGCTTTTCTTGAGTGTCGGAGAGGTAGATGGAATTCCTAGTGTAGCGGTGAAATGCGTAGATATTAGGAGGAACACCAGTGGCGAAGGCGATCTACTGGACGATAACTGACGCTGAGGCGCGAAAGCGTGGGGAGCAAACAGG
## 1428                                                         ACAGAGAGTGCAAGCGTTAATCGGAATTATTGGGTGTAAAGGGTGTGTAGGTGGATAAAAAAGTTATATGTGAAATCCCTAAGCTCAACTTAGGAACTGCATGTAATACTTTTTATCTAGAGTATTACAGAGGAAAGTAGAATTTCCGGTGTAGCGGTGAAATGCGTAGATATCGGAAGGAATACCAGAGGCGAAGGCGACTTTCTGGGTAATTACTGACACTGATACACGAAAGCGTGGGGAGCAAACAGG
## 1429                                                         ACGAAGGGTGCAAGCGTTGCTCGGAATTATTGGGCGTAAAGGGTAGGTAGGTGGTCTTATTTGTCTAGGGTGAAATCCCAGGGCTTAACCCTGGAAGTGCCTTAGAAACGGTAGGACTAGAGTTCTAGAGAGGGTCGTGGAATTCCCGGTGTAGCGGTGAAATGCGTAGAGATCGGGAGGAACACCAGAAGCGAAGGCGACGACCTGGATAGATACTGACACTCAACTACGAAAGCGTGGGGAGCAAACAGG
## 1430                                                         ACGGAGGATGCGAGCGTTATCCGGATTTATTGGGTTTAAAGGGTGCGTAGGCCGACTTTTAAGTCAGCGGTGAAATTTTCCGGCTTAACCGGGACCTTGCCGTTGAAACTGATAGTCTTGAGTGTAAATGAGGTAGGCGGAATGTGTGGTGTAGCGGTGAAATGCTTAGATATCACACAGAACTCCGATTGCGAAGGCAGCTTACTGGCATACAACTGACGCTGAGGCACGAAAGTGTGGGTATCAAACAGG
## 1432                                                         ACGTAGGGTGCGAGCGTTAATCGGAATTACTGGGCGTAAAGCGTGCGCAGGCGGTTTCGTAAGACAGACGTGAAATCCCCGGGCTCAACCTGGGAACTGCGTTTGTGACTGCGAGGCTAGAGTATGGCAGAGGGGGGTGGAATTCCACGTGTAGCAGTGAAATGCGTAGAGATGTGGAGGAACACCGATGGCGAAGGCAGTCCCCTGGGATAAGACTGACGCTCATGCACGAAAGCGTGGGGAGCAAACAGG
## 1434                                                         ACGGAGGTGGCAAGCGTTACTCGGAATTATTGGGTGTAAAGGGCAAGTAGGCGGTTTATTAAGTGGGTGGTGAAATACTCCGGCTCAATCGGAGAACAGCCTTCCAAACTGATAGGCTTGAGTGTGACAGAGGAAATTGGAATTCTCGGTGTAAGGGTGAAATCTGTAGATATCGAGAAGAACACCAGCAGCGAAGGCGAATTTCTGGGTCACTACTGACGCTAAATTGCGAAAGCTAGGGGAGCAAACAGA
## 1436                                                         ACGGAGGGTGCAAGCGTTATCCGGATTCACTGGGTTTAAAGGGTGCGTAGGTGGGTTGGTAAGTCAGTGGTGAAATCCCCGAGCTTAACTTGGGAACTGCCATTGATACTATTAATCTTGAATACCGTGGAGGTCGGCGGAATATGTCATGTAGCGGTGAAATGCTTAGATATGACATAGAACACCAATTGCGAAGGCAGCTGGCTACACGAGTATTGACACTGAGGCACGAAAGCGTGGGGATCAAACAGG
## 1443                                                         ACGAAGGGTGCTAGCGTTGTTCGGAATTACTGGGCGTAAAGGGCGCGTAGACGGTTAGATAAGTTAGATGTGAAAGCCCTAGGCTTAACCTAGGATGTGCATTTAATACTGTTTAGCTAGAGTACAAGAGGGGAAAATGGAATTGCGTGTGTAGAGGTGAAATTCGTAGATATACGCAGGAACACCGGTGGCGAAGGCGATTTTCTGGCTTGATACTGACGTTAAGGCGCGAAAGCGTGGGGAGCAAACAGG
## 1444                                                         ACGTAGGGTGCGAGCGTTAATCGGAATCACTGGGCGTAAAGCGCTCGTAGGCGGTTCGGTCAGTCAGATGTGAAAGCCCTCGGCTTAACCGGGGAACTGCATTTGATACTGCCAGACTTGAGTGTCGGAGAGGGTGGCGGAATTCCGGGTGTAGGAGTGAAATCCGTAGATATCCGGAGGAACACCAGTGGCGAAGGCGGCCACCTGGACGACAACTGACGCTGAGGAGCGAAAGCGTGGGGAGCAAACAGG
## 1449                                                                                                              ACAGAAGGGGCAAGCATCTTGCAGAATGACTCGGCGAAGAGCGTGGTTTTGCAAAGTCGAATGAGACAAATTGTTTTGGAATTCTCAAAGAATGAGTCAAATCATTGGAGATTGGGAGGAACGCCAAAGGTGAAGACAGAAACACACAACATTCAAAAGCAGACCTACGGAAGCCCAGGGAGCAAAACGGATTAGATAC
## 1452                                                         ACGTAGGGTGCAAGCGTTGTCCGGAATTATTGGGCGTAAAGAGCTCGTAGGCGGTTTGTCACGTCTGCTGTGAAAATCCGAGGCTCAACCTCGGACTTGCAGTGGGTACGGGCAGGCTAGAGTGCGGTAGGGGAGACGGGAATTCCTGGTGTAGCGGTGGAATGCGCAGATATCAGGAGGAACACCGATGGCGAAGGCACGTCTCTGGGCCGTAACTGACGCTGAGGAGCGAAAGCGTGGGGAGCGAACAGG
## 1453                                                        ACGTAGAGTACAAGCGTTATCCGGATTTATTGGGCGTAAAGCGTATCGCAGGTGGAAATGTGCGTGGCGGGTGTAAGCCTTGGGCTTAACTCAAGAACCGCTCGTCATACGGCATTTCTGGAGTTTGTTGGAGGGAACTGGAATTCTTCATGGAGGGGTGAAATCCGTAGATATGAAGAGGAACGCCAAGGGCGAAGGCAAGTTCCTACGACAATACTGACACTCATGAACGACAGCTAGGGGAGCAAAAGGG
## 1454                                                         ACGAAGGGTGCTAGCGTTGTTCGGAATCATTGGGCGTAAAGCGTACGTAGGCGGTTTAGTAAGTCTGTCGTGAAATACCCGAGCTTAACTTGGGAATTGCGATAGAAACTGCTTTGCTGGAGTATAGTGGGGGATAGTGGAATTCCTAGTGTAGGGGTGAAATCCGTAGAGATTAGGAGGAACATCAGTGGCGAAGGCGACTATCTACGCTATTACTGACGCTAAGGTACGAAAGCGTGGGGAGCAAACAGG
## 1456                                                        ACGGAGGATCCAAGCGTTATCCGGATTTACTGGGCGTAAAGAGTTGCGTAGGTGGTTTGTTAAGTGAATAGTGAAATCTGGTGGCTCAACCATCAACACATTATTCAAACTGACAAACTCGAGAATGAGAGAGGTCACTGGAATTCCTTGTGTAGGAGTGAAATCCGTAGATATAAGGAGGAACACCAATGGCGTAGGCAGGTGACTGGCTCATTTCTGACACTGAGGCACGAAAGCGTGGGGAGCGAACGGG
## 1457                                                         ACGAAGGGTGCAAGCGTTAATCGGAATTACTGGGCGTAAAGCGCGCGTAGGTGGTTCAGCAAGTTGGATGTGAAAGCCCTGGGCTCAACCTGGGAACTGCATCCAAAACTACTGAGCTAGAGTACGGTAGAGGGTAGTGGAATTTCCTGTGTAGCGGTGAAATGCGTAGATATAGGAAGGAACACCAGTGGCGAAGGCGACTACCTGGACTGATACTGACACTGAGGTGCGAAAGCGTGGGGAGCAAACAGG
## 1460                                                         ACGTAGGGTGCAAGCGTTAATCGGAATTACTGGGCGTAAAGCGAGCGCAGGCGGTTATGTAAGACAGATGTGAAATCCCAGGGCTTAACCTTGGAACTGCATTTGTGACTGCATGGCTAGAGTGTGTCAGAGGGGGGTGGAACTCCACGTGTAGCAGTGAAATGCGTAGAGATGTGGAAGAACACCGATGGCGAAGGCAGCCCCCTGGGATAACACTGACGCTCAGGCTCGAAAGCGTGGGGAGCAAACAGG
## 1462                                                         ACGTAGGTGACAAGCGTTATCCGGAATTACTGGGCGTAAAGCGCACGCAGGCGGTGTTATAAGTCTGATGTGAAACCTTCTGGCTTAACCAGAAGACGTCGTTGGATACTGTAACACTTGAGGCGATGAGAGGAATGCGGAATTCTCGGTGTAATGGTGGAATATGTAGATATCGAGAGGAACATCTGTGGCGAAAGCGGCATTCTAGCATTAGCCTGACGCTCATGTGCGAAAGCGTGGGGAGCAAACGGG
## 1464                                                         ACGAAGGTCCCAAGCGTTATCCGGATTTATTGGGTTTAAAGGGTGCGTAGGCGGACTATTAAGTCAGTGGTGAAAGCCCCAAGCTTAACTTGGGAACTGCCATTGATACTGATGGTCTTGAGTACATTTGAGGTGGGCGGAATGTGTCATGTAGCGGTGAAATGCTTAGATATGACACAGAACACCGATTGCGAAGGCAGCTCACTAAACTGTAACTGACGCTGAGGCACGAAAGCGTGGGGAGCAAACAGG
## 1466                                                         ACGGAGGGGGCTAGCGTTGTTCGGAATTACTGGGCGTAAAGCGTACGTAGGCGGTGTCTCAAGTCAGAGGTGAAAGCCTGGAGCTCAACTCCAGAACTGCCTTTGAAACTAGGACGCTAGAATTGCGGAGAGGTGAGTGGAATTCCGAGTGTAGAGGTGAAATTCGTAGATATTCGGAAGAACACCAGTGGCGAAGGCGGCTCACTGGACGCATATTGACGCTGAGGTACGAAAGCGTGGGGAGCAAACAGG
## 1467                                                        ACGGAGGATCCAAGCGTTATCCGGAATTACTGGGCGTAAAGAGTTGCGTAGGTGGCATTGTAAGTTAGTAGTGAAAGCGTGTGGCTCAACCATACTCACATTATTAAAACTGCAAAGCTAGAGTATATGAGAGGTAGCTGGAATTTCTAGTGTAGGAGTGAAATCCGTAGATATTAGAAGGAACACCGATGGCGTAGGCAGGCTACTGGCGTATTACTGACACTAAGGCACGAAAGCGTGGGGAGCGAACAGG
## 1470                                                         ACGAAGGATCCAAGCGTTGTCCGGATTTACTGGGTTTAAAGGGAGCGTAGGCGGTCTCATAAGTCAGTGGTGAAAGCCTTCAGCTTAACTGAAGAATTGCCATTGAAACTGCGAGACTCGAGTATGGTTGAGGTCACTGGAATATAACATGTAGCGGTGAAATGCTTAGATATGTTATAGAACACCAATTGCGAAGGCAGGTGACTAAGCCATAACTGACGCTGAGGCTCGAAAGCGTGGGGAGCAAACAGG
## 1471                                                         ACAGAGGGTGCAAACGTTGCTCGGAATTATTGGGCGTAAAGCGCGTGTAGGCGGCCCGTTAAGTCGGTTGTGAAAGCCCATGGCTCAACCATGGAAGTGCGACTGAAACTGGCGGGCTGGAGTCCTGAAGAGGGTGGCGGAATTCCCAGTGTAGAGGTGAAATTCGTAGATATTGGGAGGAACACCGGTGGCGAAGGCGGCCACCTGGGCAGGCACTGACGCTGAGACGCGAAAGCGTGGGGAGCAAACAGG
## 1472                                                         ACGGAGGGTGCAAGCGTTAATCGGAATTACTGGGCGTAAAGCGAGTGTAGGCGGCTTCGTAAGTCGGATGTGAAAGCCCTGGGCTTAACCTGGGAATTGCATTCGATACTGCGCGGCTAGAGTCTGATAGAGGGAAGTGGAATTCCAGGTGTAGCGGTGAAATGCGTAGATATCTGGAGGAACACCAGTGGCGAAGGCGGCTTCCTGGATCAAGACTGACGCTGAGGCTCGAAAGCGTGGGGATCAAACAGG
## 1473                                                         ACGTAGGATCCGAGCGTTGTCCGGAATTACTGGGTGTAAAGGGTGCGTAGGCGGACTTGTGCGTCAGAGGTGAAATCCACGGGCTTAACCCGTGGGGTGCCTTTGATACGGCAAGTCTTGAGTGCGAGAGAGGTGGGTGGAATTCCTGGTGTAGCAGTGAAATGCGTAGATATCAGGAGGAACACCGATGGCGAAGGCAGCCCACCGGCTCGCAACTGACGCTGAGGCACGAAAGCGTGGGGATCAAACAGG
## 1476                                                         ACGAGAGGTGCAAGCGTTATTCGGAATTATTGGGCGTAAAGGGTGCGTAGACGGTTTTTTAAGTCTCTTGTTTAATTCTCCGGCCTAACTGGGGGTCAGCGAGAGATACTGAATAACTAGAGAATGAAAGAGGGAAGTGGAATTCTCGGAGTAGCGGTTAAATGCGTAGATCTCGAGAGGAACACCGATGGCGAAAGCAGCTTCCTGGTTCATATCTGACGTTCAAGCACGAAAGCGTGGGGAGCAAACAGG
## 1479                                                          CAGAGGTGGCAAGCGTTGTTCGGAATTACTGGGCGTAAAGGGCGCGTAGGCGGCCTTGTAAGTCTCGTGTGAAAGCCCTCGGCTTAACTGAGGATCTGCACGGGATACTGCGAGGCTTGAGTTCGGGAGAGGAAAGCGGAATTCCGGGTGTAGCGGTGAAATGCGTAGATATCCGGAGGAACACCAGTGGCGAAGGCGGCTTTCTGGACCGACACTGACGCTGAGGCGCGAAAGCTAGGGGAGCAAACGGG
## 1484                                                          ACGTAGGTGGCAAGCGTTGTCCGGAATTATTGGGCGTAAAGGGAGCGCAGGTGGGAATATCAGTCTGTCTTAAAAGTTCGGGGCTCAACCCCGTGATGGGATGGAAACTATATTTCTTGAGTGCAGGAGAGGAAAGTGGAATTCCTAGTGTAGCGGTGAAATGCGTAGATATTAGGAGGAACACCAGTGGCGAAGGCGACTTTCTGGACTGTAACTGACACTGAGGCTCGAAAGCCAGGGTAGTGAACGGG
## 1487                                                       ACCAGCGCCTCGAGTGGTACCCACGAATATTGGGCCTAGAGTGTCCGTAGCCGGACTATTAAGTCCGCTGTGAAATGTTGGGGCTCAACCCTAACGCGTGCAGTGGATACTGTTAAGTCTTGGGAATGGGGGAGGGTCGGAGTACTTCCAGGGTAGGGGTAAAATCCGTTGAGCCTGGAAGGACTACCAGTGGCGAAGGCGCCGACCCAAAACATATCCGACGGTGAGGGACGAAAGCTGGGGGAGCGAACGGG
## 1488                                                         ACAGAGACTGCAAGCGTTATTCGGATTCACTGGGCGTAAAGGGTGCGCAGGCGGTCGCGTGTGTTAGGTGTGAAAGCCCGAGGCTCAACCTCGGAATTGCGCCTAAAACTACGTGACTAGAGCATTGGAGAGGGTAGCGGAATTCATGGTGTAGCAGTGAAATGCGTAGATATCATGAGGAACACCAGAGGCGAAGGCGGCTACCTGGACAATTGCTGACGCTCAGGCACGAAAGCGTGGGGAGCAAAAGGG
## 1490                                                         ACGGAGGGTGCAAGCGTTAATCGGAATTACTGGGCGTAAAGCGCACGTAGGCGGTCTGATAAGTCTGATGTGAAATCCCCGGGCTCAACCTGGGAATGGCATTGGATACTGTCCGACTAGAGTATGGTAGAGGGTAGTGGAATTTCCGGTGTAGCAGTGAAATGCGTAGAGATCGGAAGGAACACCAGTGGCGAAGGCGGCTACCTGGACCAATACTGACGCTGAGGTGCGAAAGCGTGGGGAGCAAACAGG
## 1491                                                         ACGTAGGACCCAAGCGTTGTCCGGAATTACTGGGTGTAAAGGGTGCGCAGGCGGGCTTGTGCGTCGAGGGTTAAAGACCCCAGCTTAACTGGGGAAGTGCCCTCGATACGGCAAGTCTTGAGTGTGAGAGAGGAAAGTGGAATTCTTGGTGTAGCAGTGAAATGCGTAGATATCAAGAGGAACACCGGTGGCGAAGGCGGCTTTCTGGCTCACAACTGACGCTCAGGCACGAAAGCGTGGGGAGCGAACAGG
## 1493                                                        ACGAGAGCCCCAAGCGTTATCCGGAATTATTGGGCGTAAAGGGTGCGTAGGCGGTTGTGTTAGTCAGGTGTTAAAGACCCGAGCTTAACTTGGGGAATGCACTCGAAACGGCACGACTTAGAATGAAGAAGAGGTGTGCGGAACTCATGGTGTAGGGGTGAAATCCGTTGATATCATGGGGAACACCAAATGCGAAAGCAGCACACTGGTCTTTTATTGACGCTGAGGCACGAAAGCGTGGGTAGCGAATGGG
## 1494                                                         ACGAAGGGGGCGAGCGTTATCCGGAATGACTGGGCGTAAAGGGTCCGTAGGCAGCATTTTTTGTTAGAGGTAAAAGAAAAAAGCTCAACTTTTTTTGGCTTCTAAAACGTAAATGCTTTGAGTTTGAAAGGGGATAGCAGAATTTCACATGGAGGGGTGATATCCGTAGATATGTGAAGGAATACCAGAGGCGAAGGCGGCTATCCAGTTCAATACTGACGTTGAGGGACGAAAGCATGGGGAGCAAACAGG
## 1495                                                         ACGGAGGGGGCTAGCGTTGTTCGGAATGACTGGGCGTAAAGGGCGTGTAGGCGGTTGATCAAGTCAGACGTGAAAGCCTTGGGCTTAACCCAAGAACTGCGTTTGAGACTGGTTGACTTGAGGACGAGAGAGGAAAGTAGAATTCCGAGTGTAGAGGTGAAATTCGCAGATATTCGGAGGAATACCAGAGGCGAAGGCGTCTTTCTGGCTCGTTTCTGACGCTGAGGCGCGAAAGCGTGGGGAGCAAACAGG
## 1496                                                         ACGAAGGATGCAAGCGTTATCCGGATTCATTGGGTTTAAAGGGAGCGTAGGTGGATTTGTAAGTCAGTGGTGAAAGCCGGCAGCTTAACTGTCGAACTGCCATTGATACTGCAGATCTTGAGTATGGTTGCTGTGGGCGGAATATGACATGTAGAGGTGAAATTCATAGAGATGTCATAGAACACCGATTGCGAAGGCAGCTCACAAAACCATTACTGACACTGAGGCTCGAAAGTGCGGGGATCAAACAGG
## 1497                                                         ACGAGGGATCCTAGCGTTGTTCGGAATCATTGGGCGTAAAGCGGGTGTAGGCGGCTTCGTAAGTCAGGTGTGAAAGCCCAGGGCTCAACCCTGGAAGTGCATTTGATACTGCGGAGCTTGAGTGCTAGAGAGGTCATTAGAATACCTGGTGTAGTGGTGAAATACGTAGATATCAGGTGGAATACCGGAGGCGAAGGCGGATGACTGGCTAGACACTGACGCTCAGACCCGAAAGTGTGGGGATCAAACAGG
## 1498                                                         ACAGGGGATGCAAGTGTTATCCGGAATTATTGGGCGTAAAGCGTCTGCAGGTTGATAAATAAGTCTTTTGTCAAATCTCTGAGCTTAACTCAGATCTTGCAAAAGAAACTATATATCTTGAGTTTGATAGAAGTAAAGGGAATTTTTAGTGGAGCGGTGAAATGCGTAGATATTAGAAAGAACACCAAGAGCGAAGGCACTTTACTGGGTCATAACTGACACTCAGAGACGAAAGCTAGGGTAGCAAACGGG
## 1501                                                         ACGGAGGGTGCAAGCGTTGTTCGGATTTATTGGGCGTAAAGCGCGCGCAGGCGGGTTTGTAAGTCAGATGTGAAATCTCGGGGCTCAACTCCGAAACTGCGTCTGAAACTACAAATCTAGAATCTCATAGGGGGTAGGGGAATTTCACGTGTAGGGGTAAAATCCGTAGAGATGTGAAGGAACACCTGAGGCGAAGGCGCCTACCTGGATGAGTATTGACGCTGAGGCGCGAAAGCGTGGGGAGCAAACAGG
## 1503                                                         ACGGAGAGTGCAAGCATTAATCGGAATTATTGGGCGTAAAGGGCGCGTAGGCGGAAAAGAAAGTCAGATGTGAAATTCCGAAGCTCAACTTCGGAGCTGCATTTGAAACTCCTTTTCTAGAGGGTTGACGGAGAAAACGGAATTCCACGTGTAGCGGTGAAATGCGTAGATATGTGGAAGAACACCGGTGGCGAAAGCGGTTTTCTAGTTAATACCTGACGCTGAGGCGCGAGAGCAAGGGGAGCAAACAGG
## 1504                                                         ACGAGGGATGCAAGCGTTATCCGGAATTATTGGGCGTAAAGCGTCCTGAGGTGGCTTAAAAAGTCGTTCGTGAAAGCTCAGGGCTTAACCTTGAAATTGCGGCCGAAACTTCTAAGCTTGAGTATAGTAGGGGCAGAGGGAATTTCCGGAGGAATGGTGAAATATGTAGATATCGGAAAGAAGGCCAAAGGCGAAAGCACTCTGCTGGGCTAATACTGACACTGACAGACGAAAGCTAGGGGAGCGAAAGGG
## 1506                                                         ACAGAGGGTGCAAGCGTTGTTCGGAATTATTGGGCGTAAAGGGAGTGTAGGCTGTCCCCTAAGTCTTCTGTGAAATCTACGGGCTCAGCCTGTAACGTGCAGAGGAAACTGGGGGGCTTGAGTGTCGGAGAGGATAGTGGAATTCCCGGTGTAGAAGTGAAATTCGTAGAGATCGGGAGGAACACCAGTGGCGAAGGCGACTGTCTGGCTGACAACTGACGCTGAGACTCGAAAGCATGGGGAGCAAACAGG
## 1508                                                        ACAGAGACCACAAGCGTTATCCGGAATCATTGGGCGTAAAGGGTGAGTAGGTGGTATTGTTAGTCGGGTTTAAAAGATTTGGGGCTTAACCCCAGATAGGGACTCGAAACGGCAAAACTAGAATATGTGAGAGGTTAGTGGAACTCTATGTGTAGGGGTAAAATCCGTTGATATATAGGGGAACACCTAAAGCGAAGGCAGCTAACTGGCACATTATTGACACTGAATCACGAAAGCGTGGGTAGCGATACGG
## 1510                                                         ACGTAGGGTGCGAGCGTTAATCGGAATTACTGGGCGTAAAGGGTGCGCAGGCGGTTATGTAAGTCAGATGTGAAATCCCCGGGCTTAACCTGGGAATTGCGTTTGAAACTACATGACTAGAGTGTAGCAGAGGGAAGTGGAATTCCATGTGTAGCAGTGAAATGCGTAGAGATATGGAGGAACATCAATGGCGAAGGCAGCTTCCTGGGTTAACACTGACGCTCATGTACGAAAGCGTGGGGAGCAAACAGG
## 1511                                                         ACGTAGGTGGCAAGCGTTATCCGGAATCACTGGGCGTAAAGGGCATCTAGGCGGTTTATTAAGTCAGAAGTGAAATACCGGGGCTCAACCCCGGGACTGCTTTTGAAACTGGTAGACTAGAATACGGGATAGGAGGGCGGAACTACAAGTGTAGAGGTGAAATTCGTAGATATTTGTAGGAATGCCGATGACGAAGGTAGCTCTCTGGACCGATATTGACGCTGAAGTGCGAAAGCTAGGGTAGCAAACAGG
## 1512                                                         ACGAAGGGAACTAGCGTTGTCCGGATTTACTGGGCGTAAAGGGCGCGTAGGCGGTTACATAAGTTAGAAGTGAAAGCCCCGGGCTTAACCTGGGAATTGCTTTTAATACTGTGTAACTAGAGTTCGGCAGAGGATAGTAGAATTTCCAGTGTAGAGGTGAAATTCGTAGATATTGGAAGGAATACCAGTGGCGAAGGCGGCTATCTGGACCGAAACTGACGCTAATGCGCGAAAGCGTGGGGAGCAAACAGG
## 1515                                                         ACGAAGGCCCCAAGCGTTATCCGGAATTACTGGGTGTAAAGCGTCTGTAGGTGGTTCGGAAAGTTTCAAGTGAAAGGTCAGGGCTCAACCCTGTACTTGCTTGGAAAACTATCGAACTTGAGTATGGGAGAGGCAAGCAGAATAATAAGAGTAGGGGTGCAATCCGTTGATACTTATTAGAATACCAAAAGCGAAGGCAGCTTGCTGGAACATTACTGACACTGAGAGACGAAAGCGTGGGGAGCAAAAGGG
## 1522                                                          ACGTAGGTGGCGAGCGTTGTCCGGAATTATTGGGCGTAAAGAGCATGTAGGCGGCTTAATAAGTCGAGCGTGAAAATGCGGGGCTCAACCCCGTATGGCGCTGGAAACTGTTAGGCTTGAGTGCAGGAGAGGAAAGGGGAATTCCCAGTGTAGCGGTGAAATGCGTAGATATTGGGAGGAACACCAGTGGCGAAGGCGCCTTTCTGGACTGTGTCTGACGCTGAGATGCGAAAGCCAGGGTAGCGAACGGG
## 1523                                                        ACGTAGGGGGCGAGCGTTGTCCGGATTTATTGGGCGTAAAGAGCTCGTAGGCGGTTTGGTAAGTCAAATGTGAAATCTCCACGCTCAACGTGGAGGGGCCATTCGATACTGCCATGACTTGAGTTCGGTAGGGGAATGTGGAATTCCTGGTGTAGCGGTGAAATGCGCAGATATCAGGAGGAACACCCGTAGCGAAGGCGGCATTCTGGGCCGATACTGACGCTGAGGAGCGAAAGCGTGGGGAGCGAACAGG
## 1527                                                          ACGTAGGTGGCGAGCGTTGTCCGGAATTATTGGGCGTAAAGAGCATGTAGGCGGCTTTTTAAGTCGGTCGTGAAAATGCGGGGCTTAACCCCGTATTGCGGTCGAAACTGGAGGGCTTGAGTGCAGGAGAGGAAAGGGGAATTCCCAGTGTAGCGGTGAAATGCGTAGATATTGGGAGGAACACCAGTGGCGAAGGCGCCTTTCTGGACTGTGTCTGACGCTGAGATGCGAAAGCCAGGGTAGCAAACGGG
## 1528                                                         ACGAGAGCCTCAAGCGTTATCCGGAATCATTGGGCGTAAAGGGTGCGTAGGTGGTTTGATTAGTCTTCTGTTAAAGACTCGGGCTTAACCCGAGAAGAGCAGAGGATACGGTCAGGCTAGAGTATGTGAGGGGTGTGTGGAACTCATGGTGTAGGGGTGAAATCCGTTGATATCATGGGGAACACCAAAAGCGAAGGCAGCACACTGGCACATCACTGACACTGAAGCACGAAACCCTGGGTAGCGAATGGG
## 1530                                                         ACGTAGGATGCAGACGTTATCCGGAATTATTGGGCGTAAAGAGTTCGTAGGTGGTTTGTTAAGTAGAGCGTTAAAGACTTGAGCTCAACTCAAGACAGGTGCTCTATACTGGCTAACTAGAGTAAGGCAGAGGCAGGTGGAATTCTGTGTGTAGGGGTAATATCCGTAGATATACAGAGGAACACCAAAAGCGAAGGCAGCCTGCTGGGCCCTTACTGACACTCAAGAACGAAAGCGTGGGGAGCAAACAGG
## 1531                                                         ACGAGGGATCCTAGCGTTGTTCGGAATCATTGGGCGTAAAGCGGGTGTAGGTGGCTTTGTAAGTCAGGTGTGAAATCCCAGGGCTTAACCCTGGAAGTGCATTTGATACTGCGAAGCTTGAGTGCCGGAGAGGTTATTAGAATACCTGGTGTAGTGGTGAAATACGTAGATATCAGGTGGAACACCGGAGGCGAAGGCGGATAACTGGCCGGACACTGACACTCAGACCCGAAAGCGTGGGGATCAAACAGG
## 1534                                                          ACGTAGGTGGCGAGCGTTGTCCGGAATTATTGGGCGTAAAGGGAGCGCAGGCGGGAGATCAAGTCTATCTTAAAAGTGCGGGGCTCAACCCCGTGAGGGGATGGAAACTGGTCTTCTTGAGTGCAGGAGAGGAAAGCGGAATTCCTAGTGTAGCGGTGAAATGCGTAGATATTAGGAGGAACACCAGTGGCGAAGGCGGCTTTCTGGACTGTAACTGACGCTGAGGCTCGAAAGCGTGGGGAGCGAACAGG
## 1537                                                        ACAGAGACCCCAAGCGTTATCCGGAATTATTGGGCGTAAAGGGTGAGTAGGTTGTTTTATTAGTCAGATGTCAAAGCTCTAGGGCTCAACCTTAGAATCGCATTTGAAACGGTAAAACTAGAATATTGGAGAGGTGAGTGGAACTCTATGTGTAGGGGTAAAATCCGTTGATATATAGGGGAACACCAAAAGCGAAGGCAGCTCACTGGCCAATTATTGACACTGAATCACGAAAGCGTGGGTAGCGATACGG
## 1545                                                         ACGGAGGGTGCAAGCGTTATCCGGATTCACTGGGTTTAAAGGGTGCGTAGGTGGCTTTATAAGTCAGTGGTGAAAGCCCGGAGCTTAACTCCGGAACTGCCATTGATACTGTTTAGCTTGAATCAAGTAGAGGTGGATGGAATATTACATGTAGCGGTGAAATGCTTAGAGATGTAATAGAACACCAATTGCGAAGGCAGTTCACTATGCTTGCATTGACACTGAGGCACGAAAGCGTGGGGATCAAACAGG
## 1547                                                         ACGTAGGGTGCGAGCGTTAATCGGAATTACTGGGCGTAAAGCGTGCGCAGGCGGTTCGTTAAGACAGATGTGAAATCCCCGGGCTCAACCTGGGAACTGCATTTGTGACTGGCGGACTAGAGTTCGGTAGAGGGGGGTGGAATTCCACGTGTAGCAGTGAAATGCGTAGAGATGTGGAGGAACACCAATGGCGAAGGCAGCCCCCTGGGCCTGAACTGACGCTCATGCACGAAAGCGTGGGGAGCAAACAGG
## 1550                                                         ACGTAGGGTGCAAGCGTTAATCGGAATTACTGGGCGTAAAGCGTCCGCAGGCGGTCTTGTAAGTCTGTTGTTAAAGCGTGGAGCTTAACTCCATTTCAGCAATGGAAACTGTAAGACTAGAGTGTGGTAGGGGCAGAGGGAATTCCCGGTGTAGCGGTGAAATGCGTAGATATCGGGAAGAACACCAGTGGCGAAGGCGCTCTGCTGGGCCATAACTGACGCTCATGGACGAAAGCCAGGGGAGCGAAAGGG
## 1551                                                        ACGGAGGATCCAAGCGTTATCCGGAATTACTGGGCGTAAAGAGTTGCGTAGGCGGCATTGTAAGCAGGTAGTGAAAGCGTGTGGCTCAACCATACTCACATTACCTGAACTGCAAAGCTAGAGGACAAGAGAGGTCACTGGAATTCCTAGTGTAGGAGTGAAATCCGTAGATATTAGGAGGAACACCGATGGCGTAGGCAGGTGACTGGCTTGTTCCTGACGCTCAGGCACGAAAGCGTGGGTAGCAAACGGG
## 1553                                                        ACGTAGGCACCAAGCGTTGTCCGGATTTATTGGGCGTAAAGAGCTCGTAGGCGGTTTAGTAAGTCGGGTGTGAAAACTTTGGGCTTAACCCAAAGCCTGCATTCGATACTGCTATGACTTGAGTTCGGTAGGGGAGCGGGGAATTCCTAGTGTAGCGGTGAAATGCGCAGATATTAGGAGGAACACCGGTGGCGAAGGCGCCGCTCTGGGCCGATACTGACGCTGAGGAGCGAAAGCGTGGGTAGCAAACAGG
## 1554                                                         ACGTAGGGTGCAAGCGTTAATCGGAATTACTGGGCGTAAAGCGTACGCAGGCGGCTTTGCAAGACAGGTGTGAAATCCCCGGGCCTAACCTGGGAACTGCATTTGTGACTGCATGGCTAGAGTCTGTCAGAGGGGGGTGGAATTCCACGTGTAGCAGTGAAATGCGTAGAGATGTGGAAGAACACCGATGGCGAAGGCAGCCCCCTGGGATGAGACTGACGCTCATGTACGAAAGCGTGGGGAGCAAACAGG
## 1557                                                         ACGTAGGGAGCGAGCGTTGTCCGGAATCATTGGGCGTAAAGAGCACGTAGGTGGATCAATTAGTCTGCTGTGAAAGTCGAAGGCTCAACCTTCGAAAGCCGGTGGATACTGTTGATCTAGAGTACGGAAGAGGCGAGTGGAATTCCTGGTGTAGCGGTGGAATGCGCAGATATCAGGAGGAACACCAATAGCGAAGGCAGCTCGCTGGGACGTAACTGACACTGAGGTGCGAAAGCGTGGGGAGCAAACAGG
## 1558                                                         ACGAAGGGGGCTAGCGTTGTTCGGAATTACTGGGCGTAAAGCGAGCGTAGGTCGCTTAGCAAGTCAGTGGTGAAAGCCCAAGGCTCAACCTTGGAACTGCCATTGAAACTGCTAAGCTGGAATCTCAGAGGGGATAGCGGAATTCCAAATGTAGAGGTGAAATTCGTAGATATTTGGAGGAACACCGGTGGCGAAGGCGGCTATCTGGATGAGTATTGACACTGAGGCTCGAAAGTGCGGGGATCAAACAGG
## 1561                                                         ACAGAGGGTGCAAGCGTTGTTCGGAATTACTGGGCGTAAAGCGCGTGTAGGCTGCGATGTAAGTCGGATGTGAAATCCCTCAGCTCAACTGAGGAAGTGCATCCGATACTGCGTCGCTTGAGTGTCAGAGGGGATGGCGGAATTCCCGGTGTAGAGGTGAAATTCGTAGATATCGGGAGGAACACCTGTGGCGAAGGCGGCCATCTGGATGACGACTGACGCTGAGACGCGAAAGCGTGGGTAGCAAACAGG
## 1565                                                         ACGAAGGGTGCAAGCGTTGTTCGGAATTATTGGGCGTAAAGCGCGCGCAGGCGGGTCAGCAAGTCAGATGTGAAATCTCGGGGCTTAACCCCGAAACTGCGTCTGAAACTACTAACCTAGAATATCGGAGAGGAAGATGGAATTTCGCATGTAGGGGTAAAATCCGTAGAGATGCGAAGGAACACCAGAGGCGAAAGCGATTTTCTGGACGATTATTGACGCTGAGGCGCGAAAGCGTGGGGAGCAAACAGG
## 1569                                                         ACAGAGGGTGCAAGCGTTAATCGGAATTACTGGGCGTAAAGCGCACGTAGGTGGCTGCTTAAGTTGGATGTGAAAGCCCCGGGCTCAACCTGGGAATTGCATTCAATACTGTGTGGCTAGAGTATGATAGAGGAGTGTGGAATTTCCGGTGTAGCGGTGAAATGCGTAGATATCGGAAGGAACATCAGTGGCGAAGGCGACACTCTGGATTAATACTGACACTGAGGAGCGAAAGCGTGGGGATCAAACAGG
## 1570                                                         ACGAAGGGTGCAAGCGTTGTTCGGATTTATTGGGCGTAAAGCGCGCGCAGGCGGGTAATTAAGTCAGGTGTGAAATCTCAGGGCTCAACCCTGAAACTGCGCCTGAAACTGGTTATCTAGAATGTCGGAGAGGGAAGGGGAATTTCGCATGTAGGGGTAAAATCCGTAGATATGCGAAGGAATACCAGAGGCGAAGGCGCCTTCCTGGACGATGATTGACGCTGAGGCGCGAAAGCGTGGGGAGCAAACAGG
## 1571                                                         ACGAGGGATCCTAGCGTTGTTCGGAATTATTGGGCGTAAAGCGGGTGCAGGTGGCCTTGTAAGTCGAGTGTGAAAGCCTGGGGCTTAACCCCAGAAGTGCATTCGATACTGCGAGGCTTGAGTGTTGGAGAGGCTATTAGAATTCCTGGTGTAGTGGTGAAATACGTAGATATCAGGAGGAATACCGGAGGCGAAGGCGGATAGCTGGCCAAACACTGACGCTCAGACCCGAAAGCGTGGGGATCAAACAGG
## 1574                                                         ACAGAGGGTGCTAGCGTTGTTCGGAATGACTGGGCGTAAAGGGTGAGTAGGTGGTCTACTAAGTAAGGTGTGAAATCCCTGGGCTTAACTCAGGAACGGCACTTTATACTGGTTGGCTAGAGTACGGTAGAGGAAGTAGGAATTTCTAGTGGAGAGGTAATATTCATAGATATTAGAAGGAACACCAAAGGCGAAGGCAAACTTCTGGGCCGTCACTGACACTGAGTCACGAAAGCGTGGGGAGCAAACAGG
## 1576                                                        ACGGAGGATCCAAGCGTTATCCGGAATTACTGGGCGTAAAGAGTTGCGTAGGTGGCAGAGTAAGCAAATAGTGAAAGCGTGAGGCTCAACCTCATACACATTATTTGAACTGCTCAGCTTGAGTATGGGAGAGGTAGATGGAATTCCCTGTGTAGGAGTGAAATCCGTAGATATAGGGAGGAACACCGATGGCGTAGGCAGTCTACTGGCCCATTACTGACACTAAGGCACGAAAGCGTGGGGAGCAAACGGG
## 1577                                                         ACGTAGGGGGCAAGCGTTATCCGGATTTACTGGGTGTAAAGGGAGCGTAGACGGCACGGCAAGCCAGATGTGAAAGCCCGGGGCTCAACCCCGGGACTGCATTTGGAACTGCTGAGCTAGAGTGTCGGAGAGGCAAGTGGAATTCCTAGTGTAGCGGTGAAATGCGTAGATATTAGGAGGAACACCAGTGGCGAAGGCGGCTTGCTGGACGATGACTGACGTTGAGGCTCGAAAGCGTGGGGAGCAAACAGG
## 1579                                                         ACGAAGGGAGCTAGCGTTGTTCGGAATGACTGGGCGTAAAGGGCGCGTAGGCGGTTTTATAAGTTTGATGTGAAAGCCCTGGGCTTAACCTAGGAATAGCATTGGATACTGTAAGACTAGAGTCCGAGAGGGGGAAATGGAATTGCGAGTGTAGAGGTGAAATTCGTAGATATTCGCAAGAACACCAGTGGCGAAGGCGATTTCCTGGCTCGGTACTGACGCTAAGGCGCGAAAGCGTGGGGAGCAAACAGG
## 1581                                                         ACGAAGGGTGCGAACGTTATTCGGAATAATTGGGCGTACAGGGTGCGCAGGCGGTGCGTTAAGTCAACTGTTAAATCTCCCGGCCTAACCGGGAATCAGCGGTAGATACTGGCGCGCTAGAGGGTGGAAGAGAGAAGTGGAATTCTTGGAGTAGCGGTAAAATGCGTAGATCTCAAGAGGAACACCGATGGCGAAGGCAGCTTCTTGGTCCACATCTGACGCTCATGCACGAAAGCGTGGGGAGCAAACAGG
## 1582                                                         ACGTAGGGTGCGAGCGTTAATCGGAATTACTGGGCGTAAAGCGTGCGCAGGCGGTTTCGTAAGACAGACGTGAAATCCCCGGGCTCAACCTGGGAACTGCGTTTGTGACTGCGAGGCTAGAGTATGGCAGAGGGGGGTGGAATTCCACGTGTAGCAGTGAAATGCGTAGAGATGTGGAGGAACACCGATGGCGAAGGCAGCCCCCTGGGTCGACACTGACGCTCATGCACGAAAGCGTGGGGAGCAAACAGG
## 1584                                                         ACGGAGAGTGCGAGCGTTAATCGGAATTACTGGGCGTAAAGCGCACGTAGGCGGTTTGATAAGCGAGATGTGAAATCCCTGGGCTTAACCTGGGAATGGCATTTCGAACTGTCAGGCTAGAGTACGGTAGAGGATGGTGGAATTCCCGGTGTAGCGGTGAAATGCGTAGATATCGGGAGGAACATCGGTGGCGAAGGCGGCCATCTGGACTGATACTGACGCTGAGGTGCGAAAGCGTGGGGAGCAAACAGG
## 1585                                                        ACGAAGGGTGCTAACGTTATTCGGAGTTATTGGGCGTAAAGAGCGTGTAGGCGGTCTGTTAAGTTGACTGTTAAATATCCCGGCCTAACCGGGAAAATGCAGTCAAAACTGGCAAGACTAGAGGGCAGGAGAGAGAAGTGGAATTCTCGGAGTAGCGGTAAAATGCGTAGATCTCGAGAGGAACACCGATTGCGAAGGCAGCTTCTTGGCCTGTACCTGACGCTGAGACGCGAAAGCGTGGGGAGCAAACAGG
## 1586                                                       ACGGAGGAGGCAAGCGTTATCCGGATTTACTGGGCGTAAAGAGTTGCGTAGGCGGTTTTGTAAGCAAATAGTGAAATCATGTGGCTCAACCATACTGACTATTATTTGAACTGCAGAACTTGAGAGCGAGAGGGGTGATTGGAATTCCCGGTGTAGGAGTGAAATCCGTAGATATCGGGAGGAACACCAATGGCGTAGGCAGATCACTGGCTCGTTTCTGACGCTAAGGCACGAAAGCGTGGGTAGCAAACGGG
## 1587                                                         ACAGAGGGTGCAAGCGTTGTTCGGAATTACTGGGCGTAAAGCGCGTGTAGGCGGTCCTTCAAGTCGGACGTGAAATCCCTGGGCTTAACCCAGGAACTGCGTTCGAAACTGCTGGACTAGAGTATCGGAGGGGATGGTAGAATTCCTGGTGTAGAGGTGAAATTCGTAGATATCAGGAGGAATACCTGTGGCGAAGGCGACCATCTGGAAGATTACTGACGCTGAGACGCGAAAGCGTGGGGAGCAAACAGG
## 1588                                                         ACAGAGGTTGCAAACGTTGTTCGGAATTACTGGGCATAAAGCGCACGTAGGTGGTCTGCTAAGTCGCGTGTGAAATGTCCCGGCTCAACCGGGGAATTGCGCGTGATACTGGCAGGCTAGAGGTCGGGAGGGGTGAGCGGAACTCCTGGTGGAGCGGTGAAATGCGTAGATATCAGGAGGAACACCGGTGGCGAAAGCGGCTCACTGGCCCGAATCTGACACTGAGGTGCGAAAGCTAGGGTAGCAAACGGG
## 1589                                                         ACGTAGGTGGCAAGCGTTGTCCGGATTTATTGGGTTTAAAGGGTGCGTAGGCGGCTTAATAAGTCAGTGGTGAAAGCCGGTAGCTTAACTATCGAAATGCCATTGATACTGCTAAGCTTGAGTACAGTTGAGGTAGGCGGAATTGACAGTGTAGCGGTGAAATGCTTAGATATTGTCAAGAACACCAATTGCGAAGGCAGCTTACTAAAGTGTAACTGACGCTGAGGCACGAAAGTGCGGGGATCAAACAGG
## 1592                                                         ACGTAGGGTGCGAGCGTTAATCGGAATTACTGGGCGTAAAGCGTGCGCAGGCGGTTTTGTAAGACAGCTGTGAAATCCCCGGGCTTAACCTGGGAACTGCGGTTGTGACTGCAAGACTGGAGTCAGGCAGAGGGGGGTGGAATTCCGCGTGTAGCAGTGAAATGCGTAGAGATGCGGAGGAACACCGATGGCGAAGGCAGCCCCCTGGGCTATGACTGACGCTCATGCACGAAAGCGTGGGGAGCAAACAGG
## 1593                                                         ACGGGGGGGGCTAGCGTTGTCCGGAATTATTGGGCGTAAAGGGCGCGTAGGCGGAATTGTAAGTCAGTGGTTAAATTTCGGGACTTAACTCCGAGTCTGCCTTTGATACTACAATTCTTGAGTGCGAAAGAGGAAAGCGGAATTCCCGGTGTAGCGGTGAAATGCGTAGATATCGGGAAGAACACCAGTAGCGAAGGCGGCTTTCTGGTTCGCAACTGACGCTGAGGCGCGAAAGCGTGGGTAGCAAACAGG
## 1595                                                         ACGGGGGATGCAAGCGTTATCCGGATTTATTGGGTTTAAAGGGTGCGTAGGCGGTCTGTCAAGTCGGCGGTGAAAACAGCATGCTAAACATGGTGCTTGCCGTCGAGACTGGTGGGCTTGAATGTGGATGATGCCGGCGGAATGTGTAAAGTAGCGGTGAAATGCATAGATATTACACAGAAGGTCGATTGCGAAAGCAGCTGGCAAAGGCACTATTGACGCTGAGGCACGAAAGTGCGGGGATCGAACAGG
## 1598                                                         ACAGAGGGTGCGAGCGTTAATCGGAATTACTGGGCGTAAAGCGCACGTAGTCGGCTTGCCAAGTCGGGTGTGAAATCCCCGGGCTCAACCTGGGAACTGCATCCGAAACTGGCTTGCTAGAGTATGGGAGAGGGAGGCGGAATTTCTGGTGTAGCAGTGAAATGCGTAGATATCAGAAGGAACATCAGTGGCGAAGGCGGCCTCCTGGACCAATACTGACGATCAGGTGCGAAAGCGTGGGGAGCAAACAGG
## 1599                                                         ACGGAGGGTGCAAGCGTTAATCGGAATCACTGGGCGTAAAGCGCACGTAGGCCGTTTGGTAAGTCAGAGGTGAAAGCCCGTAGCTCAACTACGGAATTGCCTTTGATACTGCTGGACTTGAGTTCGGGAGAGGGTAGTGGAATTCCAGGTGTAGGAGTGAAATCCGTAGATATCTGGAGGAACATCAGTGGCGAAGGCGACTACCTGGACCGATACTGACGCTGAGGTGCGAAAGCGTGGGGAGCAAACAGG
## 1600                                                         ACGAAGGGTGCAAGCGTTAATCGGAATTACTGGGCGTAAAGCGTGCGTAGGCGGTTCGCTAAGTCTGCCGTGAAAGCCCCGGGCTCAACCTGGGAATTGCGGTGGATACTGGCGAGCTGGAGTGTGGTAGAGGAGGGTGGAATTCCCGGTGTAGCGGTGAAATGCGTAGAGATCGGGAGGAACATCAGTGGCGAAGGCGGCCCTCTGGACCAACACTGACGCTGAGGCACGAAAGCGTGGGGAGCAAACAGG
## 1601                                                        ACGTAGACCTCGAGCGTTATCCGGATTTATTGGGCGTAAAGCGTGTGTAGGGGGTTGTGTGCGTCTTCGGGTTAAATCCCACCGCCTAACGGTGGACCTGCCCAAGATACGACACAACTAGAGGAAGTTAGAGGTGCATGGAGCGCACGGTGTAGGGGTGAAATCCGTTGATATCGTGCGAAACACCAAAGGCGAAGGCAGTGCACTGGGACTTTCCTGACCCTGAGACACGAAAGCGTGGGGAGCAAAAAGG
## 1603                                                         ACGTAGGGTGCGAGCGTTAATCGGAATTACTGGGCGTAAAGCGTGCGCAGGCGGAAATATAAGTTAGAAGTGAAATCCCCGAGCTCAACTTGGGAATGGCTTTTAAGACTGTATATCTAGAGTTTGTCAGAGGGTGGTGGAATTCCAAGTGTAGCAGTGAAATGCGTAGAGATTTGGAGGAACACCGATGGCGAAGGCAGCCACCTGGGATAGAACTGACGCTGAGGCACGAAAGCGTGGGTAGCAAACAGG
## 1604                                                         ACGAGGGGTGCTAGCGTTATTCGGAATCATTGGGCGTAAAGGGTATGTAGGCGGTTTATTAAGTCTGTTGTTAAATTTCTCAGCCTAACTGGGAGTCGGCGATAGAAACTGGTTTACTAGAGTGTGGGAGAGAGAGGTGGAATTCTCGGAGTAGCGGTAAAATGCGTAGATCTCGAGAGGAACACCGATGGCGAAGGCAGCCTCTTGGCCCACTACTGACGCTGAGATACGAAAGCGTGGGGAGCAAACAGG
## 1606                                                         ACGAAGGATGCTAGCGTTACTCGGAATTACTGGGCGTAAAGAATGCGTAGGCGGATTATTAAGTTGATAGTGTAATCCCCAAGCTCAACTTGGGAATTGCTATCAAAACTAATAATCTAGAATACATGAGAGAGTAAGGGAATTCTTGGTGTAGGGGTAAAATCCGTAGATATCAGGAGGAAGACCGGAAGCGAAAGCGCTTACTTGGACTGTTATTGACGCTGAGGCATGAGAGCGTGGGGAGCAAACAGG
## 1607                                                         ACGAAGGACCCGAGCGTTATCCGGATTTATTGGGTTTAAAGGGTGCGTAGGCGGACTTTCAAGTCAGTGGTGAAAGCCCATCGCTCAACGATGGAATTGCCATTGATACTGTTGGTCTTGAGTATAGTTGAGGTGGGCGGAATGTGTCATGTAGCGGTGAAATGCTTAGATATGACACAGAACACCTATTGCGAAGGCAGCTCGCTAAGCTATAACTGACGCTGAGGCACGAAAGTGTGGGTAGCAAACAGG
## 1612                                                         ACGGAGGATTCGAGCGTTATCCGGATTTATTGGGTTTAAAGGGTGCGTAGGCGGACCAATAAGTCAGTGGTGAAATACATCAGCTTAACTGATGCACTGCCATTGATACTGTTGGACTAGAATATAAATGGTGCAGGCGGAATGAATCATGTAGAGGTGACATTCATAGATATGATTCAGAACACCGATCGCGAAGGCAGCTTGCAAAGATATTATTGACGCTGATGCACGAAAGCGTGGGTAGCGAACAGG
## 1613                                                         ACGAAGGGGGCTAGCGTTGTTCGGAATTACTGGGCGTAAAGCGAGTGTAGGTGGTTGTCATAGTCAGGTGTGAAAGCCTTGAGCTCAACTCAAGAAATGCACTTGATACTGGGCGACTAGAGGACCGGAGAGGATAGTGGAATTCCCAGTGTAGTGGTGAAATACGTAGAGATTGGGAAGAACACCAGTGGCGAAGGCGGCTATCTGGACGGTTACTGACACTAAGACTCGAAAGCGTGGGGAGCAAACAGG
## 1614                                                       ACGTATGGGGCAAGCGTTGTCCGGATTTACTGGGCGTAAAGAGCTCGTAGGCGGTTATGTAAGTCGATAAGTGAAATCTCCAGGCTCAACCTGGAGCGGCCTTACGATACTACATTGACTTGAGTACGGTAGAGGAGTGTGGAATTTCTGGTGTAGCGGTGAAATGCGCAGATATCAGAAGGAACACCTATGGCGAAGGCAGCACTCTGGGCCGAAACTGACGCTGAGGAGCGAAAGCGTGGGGAGCGAACAGG
## 1615                                                         ACGGAGGATGCAAGCGTTATCCGGATTTATTGGGTTTAAAGGGTGCGTAGGCGGCTTCTTAAGTCAGCGGTGAAACTCACGGGCTCAACTCGTGACCTGCCGTTGATACTGGGAGGCTTGAATACGGATGAAGCGGGAGGAATGTATGGTGTAGCGGTGAAATGCTTAGATATCATACAGAAGGCCTAATGCGAAGGCATCTCGCTAATCCGTCATTGACGCTGAGGCACGAAAGCGTGGGGATCAAACAGG
## 1618                                                         ACGGAGGATCCAAGCGTTATCCGGATTCATTGGGTTTAAAGGGTGCGTAGGCGGATTTGTAAGTCAGCGGTGAAATCCTACAGCTTAACTGTAGAACTGCCGTTGATACTGCAAGTCTTGAGTACATTTGATGTGGGCGGAATGTGTCATGTAGCGGTGAAATGCTTAGATATGACACGGAACACCAATTGCGAAGGCAGCTCACAAAACTGTAACTGACGCTGAGGCACGACAGCGTGGGGATCAAACAGG
## 1619                                                         ACAGAGGCCTCAAGCATTATCCGGAATTATTGGGCGTAAAGGGTGTGTAGGCGGATATATTAGTCTCCTGTTAAAGATCCCGGCTTAACCGGGAAACAGCAGGGGAAACGGTATGTCTAGAGATTGTGAGAGGTGAACAGAACTCATGGTGTAGGAGTGAAATCCGTTGATATCATGGGGAATACCAAATGCGAAGGCAGTTCACTGGCACATTTCTGACGCTGAAACACGAAAGCGTGGGTAGCGAATGGG
## 1620                                                        ACGAAGGGGGCTAGCGTTGTTCGGAATTACTGGGCGTAAAGCGCGCGTAGGCGGTTTTGCAAGCCAGAGGTGAAATGCCTGAGCTCAACTTAGGAATTGCCTTTGGGACTGCGAGGCTTTGAGGACGGAAGAGGTGAGTGGAATTCCCAGTGTAGAGGTGAAATTCGTAGATATTGGGAAGAACACCGGTGGCGAAGGCGGCTCACTGGTCCGTTTCTGACGCTGTAGCGCGAAAGCGTGGGGATCAAACAGG
## 1621                                                         ACAGAGGCCCCAAGCGTTACCCGGATTTATTGGGCGTAAAGGGTGTGTAGGTGGTTGTATTAGTCGTTTGTGAAATCCCAGGGCTTAACTCTGGACCCGCAAGCGAAACGGTACAACTAGAGGGTGCGAGAGGTATGTGGAACTCATGGTGTAGGGGTGAAATCCGTTGATATCATGGGGAACACCGAAAGCGAAGGCAGCATACTGGCGCATTCCTGACACTGAAACACGAAAGCGTGGGTAGCGAACGGG
## 1622                                                         ACGGAGGGGGCTAGCGTTATTCGGAATTACTGGGCGTAAAGCGCACGTAGGCGGACCAGAAAGTCAGATGTGAAATCCCCGGGCTCAACCTGGGAACTGCATTTGAGACTGTATAGCTAGAGTACGGTAGAGGGGGATGGAATTCCGCGTGTAGCAGTGAAATGCGTAGATATGCGGAGGAACACCGATGGCGAAGGCAATCCCCTGGACCTGTACTGACGCTCATGCACGAAAGCGTGGGGAGCAAACAGG
## 1626                                                        ACGAAGGGTGCGAGCGTTATTCGGAATAACTGGGCGTAAAGGGCGCGTAGGCGGTTTTCTAAGTTGACGGTTAAAAGCCATGGCTTAACCATGGAGATGCCGGCGATACTGGAATGACTAGAGGGTAGTAGAGAGGAGTGGAATTCTCGGAGTAGCGGTAAAATGCGTAGATCTCGAGAGGAACACCGATGGCGAAGGCAGCTCCTTGGGCTACAACTGACGCTAAGGCGCGAAAGCGTGGGGAGCAAACAGG
## 1628                                                         ACGTAGGTGGCGAGCGTTGTCCGGAATTACTGGGTGTAAAGGGCGTGTAGGCGGGAATGTAAGTCAGATGTGAAATCCCACGGCTTAACTGTGGATCTGCATCTGAAACTGCGTTTCTTGAGTGCTGGAGAGGAAAGCGGAATTCCTAGTGTAGCGGTGAAATGCGTAGATATTAGGAGGAACACCAGTGGCGAAGGCGGCTTTCTGGACAGTAACTGACGCTGAGGCGCGAAAGCGTGGGGAGCAAACAGG
## 1629                                                         ACGAAGGGAGCTAGCGTTGTTCGGAATTACTGGGCGTAAAGGGCGCGTAGGCGGCGCGACAAGTCAGATGTGAAAGCCCCGGGCTTAACCTGGGAATTGCATTTGATACTGTCGTGCTAGAGGACGAGAGAGGAAAGTGGAATTGCGAGTGTAGAGGTGAAATTCGTAGATATTCGCAAGAACACCGGTGGCGAAGGCGACTTTCTGGCTCGTATCTGACGCTAAGGCGCGAAAGCGTGGGGAGCAAACAGG
## 1632                                                        ACAGGGGATGCAAGCGTTACTCGGAATTACTGGGCGTAAAGCGTCTGTAGGTGTCTCTCCACGTCTGGCGTAAAAGCACTGGGGCTCAACCCCGTGTACGTGTCGGAAACGAGAGAGATCGAGTCACTCAGAGGTATCTGGAATGCCGTGTGTAGGGGTAAAATCCGTAGATCCACGGTGGAACGCCAAAAGCGAAAGCAGGATACTGGGGGTGTACTGACACTCAGAGACGAAAGCGTGGGGAGCAAAGGGG
## 1633                                                         ACGTAGGGACCAAGCGTTGTTCGGATTTACTGGGCGTAAAGGGCGCGTAGGCGGCTTTGTAAGTCACTTGTGAAATCTCTGAGCTTAACTCAGAACGGCCAAGTGATACTGCAATGCTAGAGTGCAGAAGGGGCAATCGGAATTCTTGGTGTAGCGGTGAAATGCGTAGATATCAAGAGGAACACCTGAGGTGAAGACGGGTTGCTGGGCTGACACTGACGCTGATGCGCGAAAGCCAGGGGAGCAAACGGG
## 1634                                                         ACGAGGGGTGCAAGCGTTATTCGGAATTACTGGGCGTAAAGCACGCGTAGGCGGTCTACTAAGTTGGTTGTTAAATCCCTCGGCTTAACCGAGGACCAGCGACCAATACTGGTAAGCTAGAGAGCAAGAGAGAGAAGTGGAATTCTCGGAGTAGCGGTGGAATGCGTAGATCTCGAGAGGAACACCGATGGCGAAGGCAGCTTCTTGGCTTGTATCTGACGCTGAGGCGTGAAAGCGTGGGGAGCAAACAGG
## 1635                                                         ACGAGAGCTTCGAGCGTTATCCGGAATTATTGGGCGTAAAGGGTGCGTAGGTTGCAATGTTAGTCTTTTGTCAAATCCTTGTGCTTAACATAAGGGCTGCGAGAGAAACGGCATAGCTTGAAAGTGTGAGAGGCGTACGGAACTCATGGTGTAGGGGTGAAATCCGTTGATATCATGGGGAACACCAAATGCGAAGGCAGTACGCTGGCGCATATTTGACACTGAAGCACGAAAGCGTGGGTAGCGAATGGG
## 1636                                                         ACGAGTGCTCCAAGCGTTATCCGGAATTATTGGGCGTAAAGGGTGTGTAGGTGGTAGTATTAGTCTTCTGTTAAATTCTTCGGCTTAACCGGGGTCCTGCAGAAGATACGGTACAACTAGAGGGTGCAAGAGGCATATGGAACTCATGGAGTAGGGGTGAAATCCGTTGATATCATGGGGAACACCAAAAGCGAAGGCAATATGCTGGTGCATTCCTGACACTGAAACACGAAAGCGTAGGTAGCGAATGGG
## 1638                                                         ACATAGGGTGCAAGCGTTGTCCGGAATTATTGGGCGTAAAGAGCTCGTAGGTCGTTTGTCGCGTCGATTGTGAAAATCTGAGGCTCAACCTCAGACCTGCAGTCGATACGGGCAAACTAGAGTGTGGTAGGGGAGACTGGAATTCCTGGTGTAGCGGTGGAATGCGCAGATATCAGGAGGAACACCAATGGCGAAGGCAGGTCTCTGGGCCATTACTGACACTGAGGAGCGAAAGCGCGGGGAGCGAACAGG
## 1639                                                         ACGGAGGATCCGAGCGTTATCCGGATTTATTGGGTTTAAAGGGTGCGTAGGCGGGATATTAAGTCAGTGGTGAAAGTTTGACGCTCAACGTTAAAATTGCCATTGATACTGGTTTTCTTGAGTACAGTTGAGGTAGGCGGAATGTGGTGTGTAGCGGTGAAATGCATAGATATACCACAGAACACCGATTGCGAAGGCAGCTTACTAAGTTGTAACTGACGCTGAAGCACGAAAGCATGGGGATCAAACAGG
## 1641                                                         ACGAACCACACAAACGTTATTCGGAATTACTGGGCTTAAAGGGTGCGTAGGCTGCGCAGTAAGTTGGGTGTGAAATACCTCGGCTCAACCGAGGAACTGCGCCCAAAACTACTGTGCTTGAGGAAGATAGAGGTAAGCGGAACTTACGGTGGAGCGGTGAAATGCGTTGATATCGTAAGGAACACCTGTGGCGAAAGCGGCTTACTGGGTCTTTTCTGACGCTGATGCACGAAAGCTAGGGGAGCAAACGGG
## 1642                                                         ACGGAGGGGGCGAGCGTTGTTCGGAATTACTGGGCGTAAAGCGTATGTAGGCGGCTTTGTAAGTCGGGTGTGAAATCCCGGGGCTTAACCCCGGAACTGCATTCGAGACTGCAAGGCTTGAGTACCGGAGAGGTTGGCGGAATTCCCAGTGTAGAGGTGAAATTCGCAGATATTGGGAGGAACACCGATGGCGCAAGCAGCCAACTGGACGGTCACTGACGCTGAGATACGAAAGCGTGGGGAGCAAACAGG
## 1643                                                         ACGGAGGGTGCAAGCGTTATCCGGATTCACTGGGTTTAAAGGGTGTGTAGGCGGGCATTTAAGTCAGTGGTGAAATCTCCGAGCTCAACTCGGAAACTGCCATTGATACTATTTGTCTTGAATATCGTTGAGGTTGGCGGAATATATCATGTAGCGGTGAAATGCTTAGATATGATATAGAACACCGATTGCGAAGGCAGCTAACTAAGCGATGATTGACGCTGAGGCACGAAAGCGTGGGGATCAAACAGG
## 1644                                                         ACGAAAGGTGCAAGCGTTACTCGGAATTACTGGGCGTAAAGCGCTCGTAGGCGGATTAGTATGTCAATTGTGAAAGCCCAGAGCTCAACTCTGGAATCGCAGTTGAAACTGCTTTTCTTGAATACCTGAGAGGGTGGGGGAATACCTAGTGGAGAAGTGAAATTCGTAGAGATTAGGTGGAACACCGGAGGCGAAGGCGCCTGCCTGGCAGTGTATTGACGCTGAGGAGCGAAAGTGTGGGGAGCAAACAGG
## 1647                                                         ACGAGTGCCTCAAGCGTTACCCGGAATCACTGGGCGTAAAGGTTGTGTAGGTTGTTTTATTAGTCTTTTGTTAAATCCTGTGGCTTAACCACAGAACTGCAAGAGAAACGGTATAACTAGAGGATGTGAGAGGTCTGCGGAACCCATAGTGTAGCGGTAAAATGCGTTGATATTATGGGGAACACCAAAAGCGAAGGCAGCAGACTGGCACACTCCTGACACTGAAACAAGAAAGCGTGGGTAGCGAATGGG
## 1650                                                         ACGTAGGGGGCTAGCGTTGTTCGGATTTACTGGGCGTAAAGAGCGCGTAGGTGGTTATATAAGTTAGGAGTGAAATCCCGGAGCTTAACTTCGGAACTGCTTCTAAAACTATATAGCTTGAGTGTGGTAGGGGGTAATAGAATTCCTAGTGTAGAGGTGAAATTCTTAGATATTAGGAGGAATACCGATGGCGAAGGCGATTACCTGGGCCACTACTGACACTGAGGCGCGAAAGCGTGGGGAGCAAACAGG
## 1651                                                         ACAGAGGTGGCGAGCGTTACTCGGATTTATTGGGTGTAAAGGGCAAGTAGGCGTCTTGACAAGTTGGGGGTGAAATCCTGCAGCTTAACTGCAGAACTGCTCTCAAAACTGCCAAGATTGAGACTGGGAGAGGAAAGCGGAATTCTCGGTGTAAGAGTGAAATCTGTAGATATCGAGAGGAACACCAGTGGCGAAGGCGGCTTTCTGGTCCAGTACTGACGCTGAATTGCGAAAGCTAGGGGAGCAAACAGG
## 1654                                                         ACAGAGGCCTCAAGCGTTATCCGGAATTACTGGGCGTAAAGGGTGTGTAGGTGGTTTAGTTAGTCTTTCGTTAAATTCTTCGGCTTAACCGGGGGCATGCGGAGGAAACGGCTAGACTTGAGGGTACGAGGGGTGCATGGAACTCATGGAGTAGGGGTGAAATCCGTTGATATCATGAGGAACACCAAAAGCGAAGGCAGTGCACTAGCGTATTCCTGACACTCAAACACGAAAGCGTGGGTAGCGAATGGG
## 1656                                                         ACAGAGGGTGCAAACGTTGTTCGGAATTACTGGGCGTAAAGCGCGTGTAGGCTGCGATGCAAGTCGGATGTGAAAGCCCTGGGCTCAACCTAGGAAGTGCATTCGAAACTGCGTTGCTTGAGTTCTGGAGAGGAAGGCGGAATTCTCGGTGTAGAGGTGAAATTCGTAGATATCGAGAGGAACACCGGTGGCGAAGGCGGCCTTCTGGACAGCAACTGACGCTGAGACGCGAAAGCGTGGGGAGCAAACAGG
## 1658                                                         ACAGAGACCACAAGCGTTATCCGAATTTATTGGGCGTAAAGGGTCCGCAGGCGGCAGTGTGCGTCATTGGTTAAACCTCGAGGCTCAACTTCGAGACTGCCCGTGATACGACACAGCTAGAGATCGGGAGAGGTAAGCGGAATTGCCGGTGTAGTAGTAAAATGCGTTAATATCGGCAAGAACACCAAATGCGAAGGCAGCTTACTGGAACGAATCTGACGCTCAGGGACGAAAGCGTGGGTAGCGAAAGGG
## 1660                                                         ACGAAGGGTGCAAGCGTTGCTCGGAATTATTGGGCGTAAAGGGTAGGTAGGTGGTCTTATTTGTCATAGGTGAAATCCTTGGGCTCAACCCGAGACGTGCCTATGAAACGGTAAGACTAGAGTAATAGAGGGGTTTGTGGAATTCCCGGTGTAGCGGTGAAATGCGTAGAGATCGGGAGGAACACCAGCGGCGAAGGCGGCAAACTGGATATTAACTGACACTCAACTACGAAAGCGTGGGGATCAAACAGG
## 1661                                                        ACGTAAGTGGCGAGCGTTGTCCGGAATTACTGGGCGTAAAGTGTGCGTAGGCGGCTTTTTAAGTTAGCATTGAAATCTTCGGGCTTAACCTGAAGTCTGGTGTTAAGACTGGAAAGCTGAGAGAATCAGAGGGGAAGGTAGAATTTCACGTGTAGCGGTGAAATGCGTAGATATGTGAGAGAATACCAGTGGCGAAGGCGGCCTTCTGGCTGATTACTGACGCTGAAGCACGAAAGCGTGGGGAGCAAACAGG
## 1664                                                         ACGTAGGATGCGAGCGTTGTTCGGAATTATTGGGCGTAAAGAGTACGTAGGCGGTTTGGTAAGTGAGAGGTTAAAGACCAGGGCTTAACCCTGGGAGTGCCCCTCAAACTGCTGAACTAGAGGTCCATAGGGGAAGGTGGAATTCCGAGTGTAGCGGTGAAATGCGTAGATATTCGGAGGAACACCGGTGGCGAAAGCGACCTTCTGGGTGGATTCTGACGCTGAGGTACGAAAGCATGGGTAGCAAACGGG
## 1666                                                        ACGTAGGTGGCGAGCGTTATCCGGATTTATTGGGTTTAAAGGGTGCGTAGGCGGCTTGATAAGTCAGTGGTTAAAGGTAGCAGCTTAACTGTTTTACATGCCATTGATACTGTTTAGCTTGAGTTATCAGAAGGTAGATGGAATTCCTGGTGTAGCGGTGAAATGCTTAGATACCAGGAGGAACACCGATTGCGAAGGCATTCTACTGCAGATAAACTGACGCTGAGGCACGAAAGCGTGGGGATCAAACAGG
## 1669                                                        ACGGAGGGTGCAAGCGTTATCCGGAATCACTGGGTTTAAAGGGTGCGTAGGCGGAGAGGTAAGTCAGTCGTGAAATTTCTTGGCTTAACCAAAGAGACTGCGATTGATACTGCTTCCCTTGAATAAGGATGAGGTAGGCGGAATGTGGCATGTAGCGGTGAAATGCTTAGATATGCCATAGAACACCGATTGCGAAGGCAGCTTGCTGGACCTCTATTGACGCTGAGGCACGAAAGCGTGGGGAGCAAACAGG
## 1672                                                         ACAGAGAGTGCAAACGTTGTTCGGAATTACTGGGCTTAAAGAGCACGTAGGCTGTGTCGTAAGTGTCTGGTGAAATCCCACGGCTTACCCGTGGAACTGCCGGGCATACTGCGGCGCTAGAGTGTGGTAGGGGAGAGCGGAACTCCAGGTGGAGCGGTGAAATGCGTAGATATCTGGAGGAACACCGGTGGCGAAGGCGGCTCTCTGGTCCACAACTGACGCTGAGGTGCGAAAGCTAGGGGAGCAAACGGG
## 1674                                                         ACGAAGGGGGCTAGCGTTGTTCGGAATGACTGGGCGTAAAGGGTGTGTAGGCGGTTAAACACGTCAGATGTGAAATCCCCGGGCTTAACCTGGGAATGGCATTTGATACGGTATAGCTAGAGATTGGTAGAGGGTAGAGGAATTGCGAGTGTAGAGGTGAAATTCGTAGAGATTCGCAGGAACACCGGTGGCGAAGGCGGCTACCTGGGCCAATACTGACGCTGAGGCACGAGAGCGTGGGGAGCAAACAGG
## 1675                                                         ACGAGGGATCCTAGCGTTGTTCGGAATCATTGGGCGTAAAGCGGGTGTAGGTGGCTATGTAAGTCAGGTGTGAAAGCCTAGGGCTCAACCCTAGAAGTGCATCTGATACTGCGTAGCTTGAGTGCTAGAGAGGTTACTAGAATTGTTGGTGTAGTGGTGAAATACGTAGATATCAACAGGAATACCGGTGGCGAAGGCGGGTAACTGGCTAGACACTGACACTCAGACCCGAAAGCGTGGGGATCAAACAGG
## 1679                                                         ACAGAGGGTGCGAGCGTTAATCGGAATTACTGGGCGTAAAGGGCGCGTAGGCGGCGCGATAAGTGAGCTGTGAAAATCATGGGCTTAACCTGTGAAAGAAAGCTCATACTGTCGAGCTAGAGTACAGGAGAGGGAAGTGGAATTTCCGGTGTAGCGGTGAAATGCGTAGATATCGGAAGGAACACTCGAGGCGAAGGCGACTTTCTGGCCTGGAACTGACGCTGAGGCGCGAAAGCGTGGGTAGCGAACAGG
## 1680                                                         ACAGGGGATGCAAGCGTTATCCGGAATGATTGGGCGTAAAGCGTCTGTAGGTGGCTTCGCAAGTCGGCTGTCAAAAATCAGGGCTTAACCTTGAGTCGGCAGTTGAAACTATGAAGCTTGAGTGAGGTAGGGGCAGAGGGAATTCCTAGTGTAGCGGTGAAATGCGTAGATATTAGGAAGAACACCAACGGCGAAAGCACTCTGCTGGGCCATGACTGACACTGAGAGACGAAAGCTAGGGGAGCAAATGGG
## 1681                                                         ACGGAGGATGCAAGTGTTATCCGGAATCACTGGGCGTAAAGCGTCTGTAGGTGGTTCGAAAAGTCCACTGTTAAAAATTGAGGCTTAACCTCAAAGATGCCGTGGAAACTTTCAAACTAGAGTATGGTAGGGGTAGAGGGAATTTCCAGTGGAGCGGTGAAATGCGTAGATATTGGAAAGACCACCGATGGCGAAGGCACTCTACTGGGCCATTACTGACACTGAGAGACGAAAGCTAGGGGAGCAAATGGG
## 1682                                                         ACAGAGGGTGCGAACGTTGCTCGGAATTACTGGGCGTAAAGCGCGTGTAGGCGGACGGGTAAGTCAGATGTGAAATCCCCGAGCTCAACTCGGGAACTGCATTTGAAACTGCTCGTCTTGAGTATCGGAGAGGGTGGCGGAATTCCTGGTGTAGAGGTGAAATTCGTAGATATCAGGAGGAACACCGGTGGCGAAGGCGGCCACCTGGACGGTCACTGACGCTGAGACGCGAAAGCGTGGGGAGCAAACAGG
## 1683                                                         ACGTAGGATGCGAGCGTTGTTCGGAATTACTGGGCGTAAAGAGTACGTAGGCGGCAATGAAAGTCTGATATTAAAGACTGGGGCTTAACCTCAGGAGTGTATCGGAAACTTCATAGCTAGAGGACAGTAGAGGAAGTCGGAATTCCCAGTGTAGCGGTGAAATGCGTAGATATTGGGAGGAACACCAGTGGCGAAAGCGGACTTCTGGGCTGTTACTGACGCTGAGGTACGAAAGTGTGGGGAGCAAACAGG
## 1685                                                         ACGAAGGGTGCTAGCGTTGTTCGGAATCACTGGGCGTAAAGCGCGCGTAGGCGGCTTTGCAAGTCAGAGGTGAAATCCCCAAGCTTAACTTGGGAACTGCCTTTGAAACTGCAAAGCTGGAGTGTCGGAGGGGATGGCGGAATTGCTAATGTAGAGGTGAAATTCGTAGATATTAGCAGGAACACCGGTGGCGAAGGCGGCCATCTGGACGACAACTGACGCTGAGGCGCGAAAGCGTGGGGATCAAACAGG
## 1686                                                         ACGAAGGGTGCAAGCGTTGCTCGGAATTATTGGGCGTAAAGGGTAGGTAGGTGGTTGCGTATGTCTGGGGTGAAATCCCTGAGCTTAACTCAGGAAGTGCCTTGGAAACGGCGTAACTAGAGTACTAGAGAGGTTCGTAGAATTCCCGGTGTAGCGGTGAAATGCGTAGAGATCGGGAGGAATACCAGAGGCGAAGGCGACGAACTGGATAGTTACTGACACTCAACTACGAAAGCGTGGGGAGCAAACAGG
## 1689                                                         ACGAAGGTCCCTAGCGTTGTTCGGAATCATTGGGCGTAAAGAGCTCGTAGGTGGTTTTGTAAGTCAGGTGTGAAAGCCCCGAGCTTAACTCGGGAAGTGCACTTGAAACTGCTTAACTTGAGTACGGAAGAGGTGAGTGGAATTCCTGGTGTAGTGGTGAAATACGTAGATATCAGGAGGAACATCGGTGGCGAAGGCGGCTCACTGGTCCAGTACTGACACTGAGGAGCGAAAGCATGGGGATCAAACAGG
## 1690                                                         ACGGAGGATCCGAGCGTTATCCGGATTTATTGGGTTTAAAGGGTGCGTAGGCGGCCTGTCAAGTCGGCGGTGAAACCCCCGGGCTCAACCCGGGGACTGCCGTCGAAACTGACGGGCTTGAGTACGTTGGGAGTGCGCGGAATGTGTGGTGTAGCGGTGAAATGCTTAGATATCACACAGAACACCGATCGCGAAGGCAGCGCACTACGCCGTAACTGACGCTGAGGCACGAAGGTGCGGGGATCGAACAGG
## 1692                                                         ACGGAGGGTGCAAGCGTTATCCGGATTTACTGGGTTTAAAGGGTGCGTAGGCGGGAGAATAAGTCAGTGGTGAAAGCACCGGGCTCAACCCGGTAACTGCCATTGATACTGTTTTTCTTGAATGCGATTGAGGTAAGCGGAATGTTGCATGTAGCGGTGAAATGCTTAGATATGTTACAGAACACCAATTGTGAAGACAGCTTACTAAGTCGTGATTGACGCTGAGGCACGAAAGCGTGGGGAGCAAACAGG
## 1694                                                         ACGAATGGGGCAAGCGTTGTTCGGAATCACTGGGCTTAAAGCGTGCGTAGGCGGATTTGCAAGTCAGAAGTGAAATCCCAGGGCTTAACCCTGGAATTGCTTTTGAAACTGCAAATCTAGAAGATTGAAGAGGTTAGCAGAATTCCGAGTGTAGAGGTGAAATTCGCAGATATTCGGAGGAATACCAATGGCGTAGGCAGCTAACTGGGCAATTTTTGACGCTGAGGCACGAAAGCGTGGGGAGCAAACAGG
## 1697                                                         ACGGAGGGTGCAAGCGTTATCCGGAATCATTGGGTTTAAAGGGTCCGCAGGCGGATTTATAAGTCAGTGGTGAAATCCTATCGCTCAACGATAGAACTGCCATTGATACTGTAAGTCTTGAATTCGGTCGAAGTGGGCGGAATATGACATGTAGCGGTGAAATGCTTAGATATGTCATAGAACACCGATAGCGAAGGCAGCTCACTAGGCCTGGATTGACGCTCAGGGACGAAAGCGTGGGGAGCAAACAGG
## 1698                                                         ACGGAGGGTGCAAGCGTTATCCGGAATCATTGGGCGTAAAGAGTCCGCAGGCGGCATAGTAAGTTAGGTATTAAAGACGGCAGCTCAACTGTCGCACTGTACCTGATACTGCTAAGCTAGAGCCTTGGAGAGGTGCGTAGAATTCTACATGTAGGGGTAAAATCCGTAGATATGTAGAGGAATACCAAAAGCGAAGGCAGCGCACTGGCCAAGAGCCGACGCTCATGGACGAAAGCGTGGGGATCAAACAGG
## 1704                                                         ACGAGGGGGGCAAGCGTTGTTCGGAATTACTGGGCGTAAAGCGCACGCAGGCGGTCTAATAAGTCAGAGGTGAAATCCCGGGGCTCAACTTCGGAATTGCCTTTGAAACTATTAGACTAGAGTACGGTAGAGGGTGGTGGAATTCCTAGTGTAGAGGTGAAATTCGTAGATATTAGGAGGAACACCAGTGGCGAAGGCGGCCACCTGGACCGTTACTGACGCTCATGTGCGAAAGCGTGGGGAGCAAACAGG
## 1705                                                         ACGAGGGATCCTAGCGTTGTTCGGAATCATTGGGCGTAAAGCGGGTGTAGGCGGGCTTATAAGTCAGGTGTGAAATCCCAGGGCTCAACCCTGGAAGTGCATTTGATACTGTAAGTCTTGAGTGTGGTAGAGGTTACTGGAATTCCTGGTGTAGTGGTGAAATACGTAGATATCAGGAGGAATACCGGAGGCGAAGGCGAGTAACTGGGCCAACACTGACGCTGAGATCCGAAAGCGTGGGGATCAAACAGG
## 1706                                                         ACGAAGGGTGCTAGCGTTGTTCGGAATCACTGGGCGTAAAGCGTGCGTAGGCGGCTTTGTAAGTCGATTGTGAAATCCCCAGGCTTAACTTGGGAACTGCAATCGAAACTGCTTAGCTTGAATATGGTCGAGGGTAGTAGAATTCCTAGTGTAGGGGTGAAATCCGTAGATATTAGGAGGAATACCGGTGGCGAAAGCGACTACCTAGGCCATTATTGACGCTGAGGCACGAAAGCGTGGGGAGCAAACAGG
## 1707                                                       ACGGAGGATCCAAGCGTTATCCGGAATTACTGGGCGTAAAGAGTTGCGTAGGCGGCCAAGTTAGCAGAGCATGAAATAGTGTGGCTCAACCATACTGCTATGTTCTGAACTGCTTCGGCTAGAGGATGAGAGAGGTTATTGGAATTCCCAGTGTAGGAGTGAAATCCGTAGATATTGGGAGGAACACCGATGGCGTAGGCAGATAACTGGCTCATTCCTGACGCTCAGGCACGAAAGCGTGGGGAGCAAACGGG
## 1709                                                         ACGTATGTCTCGAGCGTTATCCGGAATTACTGGGTGTAAAGGGTTCGCAGGCGGTTTGTTAAGTTGGGTATGAAAGACCGGGGCTCAACCCCGAGTTTGTGCTCAAAACTGGCAGACTAGAATTAGGGAGAGGTAAGCGGAATTCTAAGTGTAGGGGTGCAATCCGTAGATACTTAGAAGAACACCAAAAGCGAAGGCAGCTTACTGGAACTATATTGACGCTAATGAACGAAAGCGTGGGGAGCAAAAAGG
## 1711                                                         ACGGAGGTGGCGAGCGTTACTCGGAATTATTGGGTGTAAAGGGCAAGTAGGCGGTTTGTTAAGTGAGGGGTGAAATCTTCCGACTCAATCGGAAAACTGCCTTTCAAACTGACAGACTTGAGTGTGGAAGGGGGAAATGGAATTCGCGGTGTAAGGGTGAAATCTGTAGATATCGCGAAGAACACCAGCGGTGAAGACGGTTTCCTGGTCCATATCTGACGCTAAATTGCGAAAGCTAGGGGAGCAAACAGA
## 1713                                                         ACGTAGGGTGCGAGCGTTAATCGGAATTACTGGGCGTAAAGCGTGCGCAGGCGGTTTCGTAAGACAGACGTGAAATCCCCGGGCTCAACCTGGGAACTGCGTTTGTGACTGCGAGGCTAGAGTATGGCAGAGGGGGGTGGAATTCCACGTGTAGCAGTGAAATGCGTAGAGATGTGGAGGAACACCGATGGCGAAGGCAGCCCCCTGGGCCAATACTGACACTGAGGAGCGAAAGCGTGGGGAGCGAACAGG
## 1714                                                         ACGAAGGGGGCTAGCGTTGTTCGGAATTACTGGGCGTAAAGGGCGCGTAGGCGGCCGTTTAAGTTAGGTGTGAAAGTCCTGGGCTCAACCTGGGGACTGCACTTAATACTGGATGGCTTGAATTCGGGAGAGGATAGCGGAATTCCCAGTGTAGAGGTGAAATTCGTAGATATTGGGAAGAACACCGGTGGCGAAGGCGGCTATCTGGACCGACATTGACGCTGAGGCGCGAAAGCGTGGGGAGCAAACAGG
## 1716                                                         ACGGAGGGTGCAAGCGTTAATCGGAATTACTGGGCGTAAAGCGCGCGTAGGCGGCTGATTAAGTCGGATGTGAAAGCCCCGGGCTCAACCTGGGAACTGCATACGATACTGATCGGCTAGAGTATGAGAGAGGGAGGTAGAATTCCACGTGTAGCGGTGAAATGCGTAGATATGTGGAGGAATACCGGTGGCGAAGGCGGCCTCCTGGCTTAATACTGACGCTCATGCACGAAAGCGTGGGGAGCAAACAGG
## 1718                                                         ACGTAGGGTGCAAGCGTTAATCGGAATTACTGGGCGTAAAGCGTGCGCAGGCGGTTATATAAGACAGATGTGAAATCCCCGGGCTCAACCTGGGAACTGCATTTGTGACTGTATAGCTAGAGTACGGCAGAGGGGGATGGAATTCCGCGTGTAGCAGTGAAATGCGTAGATATGCGGAGGAACACCGATGGCGAAGGCAATCCCCTGGGCCTGCACTGACGCTCATGCACGAAAGCGTGGGGAGCAAACAGG
## 1720                                                         ACAGAGCCCACAAGCGTTATCCGGATTCATTGGGCGTAAAGCGTCCGCAGGTGTCCCAATAAGTTTTCTGTTAAACCTTGATGCTTAACATCAAGACTGCAGAAAAAACTATTGGGATTGAGACCGGGAGAGGCAAGCGGAACTATCGGTGTAGTCGTAATAAGCGCTGATATCGATAGGAACACCAAAGGCGAAAGCAGCTTGCTAGAACGTGTCTGACACTCATGGACGAAAGCGTGGGTAGCGAATGGG
## 1722                                                         ACGTGGGGTGCAAGCGTTGTTCGGAATTACTGGGCGTAAAGCGCGTGTAGGTGGCCAAGCGGGTCGGATGTGAAAGCCCTCAGCTTAACTGAGGAAGTGCGTCCGAAACTATTTGGCTCGAGTATGGGAGAGGAAGATGGAATTCCCAGTGTAGAGGTGAAATTCGTAGATATTGGGAAGAACACCTGTGGCGAAAGCGATTTTCTGGACCATAACTGACACTGAGACGCGAAAGCGTGGGGATCAAACAGG
## 1724                                                         ACGAGGGATCCTAGCGTTGTTCGGAATCATTGGGCGTAAAGCGGGTGTAGGTGGCTTGGTAAGTCGGGTGTGAAAGCCCAGGGCTCAACCCTGGAAGTGCATTCGATACTGCTGAGCTTGAGTGTTGGAGAGGTCATCGGAATACCTGGTGTAGTGGTGAAATACGTAGATATCAGGTGGAACACCGGAGGCGAAGGCGGGTGACTGGCCAAACACTGACACTGAGACCCGAAAGTGTGGGGATCAAACAGG
## 1725                                                         ACAGAGAGTGCAAACGTTGTTCGGAATTACTGGGCTTGAAGCGCACGTAGGCGGCATCGTAAGTGTCGGGTGAAATCCCACGGCTCACCCGTGGAACTGCCCGGCATACTGCGATGCTCGAGTGTGGTAGGGAAGAGCGGAACTCCTGGTGGAGCGGTGAAATGCGTAGATATCAGGAGGAACACCGGTGGCGAAGGCGGCTCTTTGGTCCACAACTGACGCTGAGGTGCGAAAGCCAGGGGAGCAAACGGG
## 1728                                                         ACGTAGGGTGCGAGCGTTAATCGGAATTACTGGGCGTAAAGCGTGCGCAGGCGGTTTGGTAAGACAGGCGTGAAATCCCCGGGCTCAACCTGGGAACTGCGCTTGTGACTGCCTCACTAGAGTACGGCAGAGGGGGGTGGAATTCCACGTGTAGCAGTGAAATGCGTAGAGATGTGGAGGAACACCGATGGCGAAGGCAGCCCCCTGGGCCGATACTGACGCTCATGCACGAAAGCGTGGGGAGCAAACAGG
## 1730                                                         ACGAAGGGGGCTAGCGTTGTTCGGAATTACTGGGCGTAAAGCGCACGCAGGCGGTGGTCATAGTCAGAAGTGAAAGCCCTGGGCTCAACCCGGGAATTGCTTTTGATACTGGACCGCTAGAATCACGGAGAGGGTAGTGGAATTCCGAGTGTAGAGGTGAAATTCGTAGATATTCGGAAGAACACCAGTGGCGAAGGCGACTACCTGGCCGTCGATTGACGCTCATGTGCGAAAGCGTGGGGAGCAAACAGG
## 1731                                                         ACGGAGGATGCAAGCGTTATCCGGATTTATTGGGTTTAAAGGGTCCGCAGGCGGAAAGGTAAGTCAGTGGTGAAAGCCCGTAGCTCAACTACGGAACTGCCATTGATACTGTTTTTCTTGAGTATGGTTGCCGTTGGCGGAATATAACATGTAGCGGTGAAATGCTTAGATATGTTATAGAACACCGATTGCGAAGGCAGCTAACGAAACCATTACTGACGCTCAGGGACGAAAGCGTGGGGAGCAAACAGG
## 1732                                                         CCCAGCGGTCCAAGTCGCAACCACAATTATTGGGTCTAAAACATCCGTAGCTTGCTTGGTAAGTCTCTTGTGAAATCCGGCCTCTTAAGGGTCGAAGAGCAAGAGATACTGCTGGGCTAGAGACCGGGAGAGGTAAGAAGTATGGTCGGGGTACCGGTAAAATGGGTTGATCCTGTCTAGACTAACAACAGCGAAGGCATCTTACCAGCACGGATCTGACAGTAAGGGATGAAGGCTAGGGGCGCAAAAGGG
## 1734                                                         ACGGAGGATCCAAGCGTTATCCGGAATCATTGGGTTTAAAGGGTTCGTAGGCGGCTTTGTAAGTCAGTGGTGAAATCTTTCAGCTTAACTGAGAAACTGCCATTGATACTGCAGTGCTTGAATTACTTTGAAGTAACTAGAATATGTAGTGTAGCGGTGAAATGCTTAGATATTACATGGAATACCAATTGCGAAGGCAGGTTACTAAGAGTTTATTGACGCTGATGAACGAAAGCGTGGGGAGCGAACAGG
## 1737                                                         ACGGAGGGTGCAAGCATTAATCGGATTTATTGGGCGTAAAGGGCGCGTAGGCGGGAAGATAAGTCAGATGTGAAATCCTGGGGCTCAACCTCAGAACTGCATTTGAAACTATCTATCTAGAGGGTAGGAGGAGAAAACGGAATTCCACAAGTAGCGGTGAAATGCGTAGATATGTGGAAGAACACCGGTGGCGAAGGCGGTTTTCTATCTTATTCCTGACGCTGAGGCGCGAAAGCAAGGGGATCAAACAGG
## 1739                                                         ACGGGGGGTCCTAGCGTTGTTCGGAATCATTGGGCGTAAAGCGTGTGCAGGTGGCTCTGCAAGTTGGATGTGAAAGCCCCGGGCTTAACTCGGGAAGTGCATTCAAAACTGCTGAGCTTGAGTATGGAAGAGGGTCGTGGAATTCCAGGTGTAGTGGTGAAATACGTAGATATCTGGAGGAACACCGGCGGCGAAGGCGGCGACCTGGTCCTATACTGACACTCATACACGAAAGCGTGGGGATCAAACAGG
## 1740                                                         ACGTAGGATGCGAGCGTTGTCCGGAATTATTGGGCGTAAAGAGTTCGTAGGTGGTATGTCAAGTCTGGTGTTAAATCCCGAAGCTTAACTTCGGATCGGCACTGGATACTGACAAACTAGAGTGCAGTAGAGGCAAGGGGAATTCCCAGTGTAGCGGTGAAATGCGTAGATATTGGGAGGAACACCGGTGGCGTAAGCGCCTTGCTGGGCTGTAACTGACGCTGAGGAACGAAAGCCAGGGGAGCAAATGGG
## 1743                                                         ACAGAAGGTGCAAGCGTTACTCGGAATTACTGGGCGTAAAGCGTCTGTAGGTGTCCTTCCACGTCTGGATTTAAATTGCGGGGCTTAACCCCGCATCTGATCTGGAAACGAGAGGGATCGAGTTATTCAGAGGCATCTGGAATGTCGTGTGTAGGGGTAAAATCCGTAGATACACGATGGAACGCCAAAAGCGAAGGCAGGATGCTAGGAATATACTGACACTCAGAGACGAAAGCGTGGGGAGCAAAGGGG
## 1752                                                         ACGGAGGGTGCAAGCGTTACTCGGAGTTACTGGGCGTAAAGCGCACGTAGGTGGCTTGGTAAGTTTGGTGTGAAAGCCCGAGGCTCAACTTCGGAAGTGCACTGAATACTGCCATGCTAGAGTATAGGAGGGGAGAGCGGAATTCCTGGTGTAGAGGTGAAATTCGTAGATATCAGGAGGAACAACAGTGGCGAAAGCGGCTCTCTGGACTATAACTGACACTGAGGTGCGAAAGCGTGGGGAGCAAACAGG
## 1753                                                         ACGGAGGGTGCAAGCGTTATCCGGATTTACTGGGTTTAAAGGGTGCGTAGGCGGACTTATAAGTCAGTGGTGAAAGCTCGGAGCTTAACTCCGGAACTGCCATTGATACTGTTAGTCTTGAATTTGGTAGAGGTAGGCGGAATGTAACATGTAGCGGTGAAATGCATAGATATGTTACAGAACACCGATTGCGAAGGCAGCTTGCTATGCCAATATTGACGCTGATGCACGAAAGCGTGGGTAGCGAACAGG
## 1755                                                        ACGGAGGATGCAAGCGTTATCCGGACTGATTGGGTGTAAAGCATCTGTAGGCGGTTTTGGAAGTCTACGGTTAAATCCCAGAGCTTAACTTTGGTTCAGCCGTAGAGTACTTCAAAGCTTGAGTATGGTAGGGGTAGAGGGAACTTCTCGTGGAGCGGTGAAATGCACAGAGATGAGAAGGAACACCAGTGGCGAAAGCGCTCTACTGGGCCATGACTGACGCTGAGAGATGACAGCTGAGGGAGCGAATAGG
## 1757                                                          ACGGGAGGGGCAAGCGTTATCCAACTTGACTGGGCGTAAAGGGTCCGTAGACTGTAAAATAAGTTACTAGTTAAATTTTAAAGCCCAACTTTAAAATAGCTTGTAATACTGTTTTACTTGAGTTTTGTACAGAAGAGTAGAATTTTATACGAAGGGGTGAAATCCGGAGATATATAAAGGAATACCAGTAGCGAAGGCGACTCTTTGGTAAAAACTGACGTTGAGGGACGAAAGTATAGGTAGCAAACAGG
## 1759                                                         ACGAGGGGTGCAAGCGTTGTTCGGAATCACTGGGCGTAAAGCGAGTCTAGGCGGATTGGTCAGTCGGATGTGAAAGCCCCAGGCTTAACCTGGGATGTGCATTCGATACTGCCAGTCTTGAATCATGGAGAGATGGACGGAATTCCGAGTGTAGCAGTGAAATGCGTAGATATTCGGAGGAACATCAGAGGCGAAAGCGGTTCATTGGCCACGTATTGACGCTAAAGCTCGAAAGCGTGGGTAGCAAACAGG
## 1760                                                         ACAGAGGATGCAAGCGTTATCCGGAATCACTGGGCATAAAGCGTCTGTAGGTGGTTTGGTAAGTCTGCTGTTAAAGACTGGGGCTCAACCCCAGAAAAGCAGTGGAAACTGCTAGACTTGAGTGTGGTAGAGGTAGAGGGAATTCCTAGTGTAGCGGTGAAATGCGTAGAGATCTGGAGGAATACCGGTGGCGAAGGCGGCCCCCTGGACAAAGACTGACGCTCAGGTGCGAAAGCGTGGGGAGCAAACAGG
## 1761                                                         ACAGAGGCCCCGAGCGTTATCCGGAATTATTGGGCGTAAAGGGTGTGTAGGTGGCGTTGTTAGTCGTTTGTGAAAGATCCGGGCTTAACCTGGAAAACGCGAACGAAACGGCAATGCTTGAATATGTGAGAGGTACGCAGAACTCATGGTGTAGGGGTGAAATCCGTTGATATCATGGGGAATACCAAAAGCGAAGGCAGCGTACTGGCACATTATTGACACTGAAACACGAAAGCGTGGGTAGCGAATGGG
## 1762                                                         ACAGAGGGTGCAAACGTTGCTCGGAATTACTGGGCGTAAAGCGCGTGTAGGCGGTTTCTTAAGTCAGGTGTGAAAGCCCGAGGCTCAACCTCGGAAGTGCACATGATACTGGGAAGCTTGAGTACTGAAGAGGAGAGCGGAATTCTCGGTGTAGAGGTGAAATTCGTAGATATCGGGAGGAACACCAGTGGCGAAGGCGGCTCTCTGGGCAGATACTGACGCTGAGACGCGAAAGCATGGGGAGCAAACAGG
## 1763                                                         ACGAGGGATCCTAGCGTTGTTCGGAATTATTGGGCGTAAAGCGGGTGTAGGTGGCTTTGTAAGTCAGGTGTGAAAGCCCAGGGCTCAACCCTGGAAGTGCATTTGATACTGCGAAGCTTGAATGTGGGAGAGGTGAGTAGAATTCCAGGTGTAGTGGTGAAATACGTAGATATCTGGAGGAATACCGGAGGCGAAGGCGGCTCACTGGCCCAACATTGACACTCAGACCCGAAAGCGTGGGGATCAAACAGG
## 1764                                                         ACGGAGGGTGCAAGCGTTAATCGGAATTACTGGGCGTAAAGCGCGCGTAGGTGGTTTGGCAAGTCGGGTGTGAAAGCCCTGAGCTTAACTTGGGAATTGCATTCGATACTGCCTGGCTAGAGTATGGTAGAGGGAAGCGGAATTCCGCATGTAGCGGTGAAATGCGTAGATATGCGGAGGAACACCGGTGGCGAAGGCGGCTTCCTGGACCAATACTGACACTGAGGCGCGAAAGCGTGGGGAGCAAACAGG
## 1770                                                     ACGTGGGGTGCAAGCGTTATCCGGATTTACTGGGCGTAAAGAGTCTGTAGGCGGATTACCAAGTTTCGTTTGAAATACCGAGGCTCAACTTCGGGAGCGGGCGGAAAACTGGTAGTCTTTTGAGAACATGTCAGGGGGTACTAGAATTCACGGTGTAGCAGTGAAATGCGTTGATATCGTGAAGAATACCAAGGGCGAAGGCAAGTACCTGGGGCATTGTTGACGCTGAGAGACGAAAGCTAGGGGAGCGAAACGG
## 1771                                                         ACGGAGGGTGCAAGCGTTATCCGGATTCACTGGGTTTAAAGGGTGCGTAGGTGGATATGTAAGTCAGAGGTGAAATCCCCGAGCTTAACTTGGGAACTGCCTTTGATACTATATATCTTGAATGTTGCGGAGGTAAGCGGAATATGTCATGTAGCGGTGAAATGCTTAGATATGACATAGAACACCGATTGCGAAGGCAGCTTGCTACACAATTATTGACACTGAGGCACGAAAGCGTGGGGAGCAAACAGG
## 1773                                                         ACGAAGGGTGCAAGCGTTATTCGGAATCACTGGGCGTAAAGAGCTCGTAGGCTGTTAATTAAGTCATATGTGAAATCCCTGGGCTCAAACCAGGACGTGCATACGATACTGATTAACTGGAGTGCAGGAGAGGAAAGCGGAATTCCAGGTGTAGAGGTGAAATTCGTAGATATCTGGAGGAACAACAGAAGCGAAGGCGGCTTTCTGGACTGTCACTGACGCTGAGGAGCGAAAGCGTGGGGAGCAAACAGG
## 1774                                                         ACGTAGGTGGCAAGCGTTGTCCGGATTTACTGGGTTTAAAGGGTGCGCAGGTGGCTATTCAAGTCCGTGTTGAAAGAGCTAGGCTTAACCTAGTAAACGGCATGGATACTGATTGGCTTGAGACTAGAGGAGGTAAGCGGAATGTAGGGTGTAGCGGTGAAATGCTTAGATACCCTACAGAACTCCTATTACGAAGGCAGCTTACTACACTATGTCTGACGCTCAGGCACGAAAGCGTGGGGAGCAAACAGG
## 1775                                                         ACGTAGGGGGCAAGCGTTGTCCGGATTTACTGGGTGTAAAGGGCGCGCAGGCGGGATAGCAAGTCAGAGGTGAAATCCTACAGCTTAACTGTAGAACTGCCTTTGATACTGTTATTCTTGAGTTCGGGAGAGAGAGACGGAATTCCAGGTGTAGTGGTGAAATACGTAGATATCTGGAAGAACACCAGTTGCGAAGGCGGTCTCTTGGTCCGATACTGACGCTGAGGCGCGAAAGCGTGGGGAGCAAACAGG
## 1776                                                         ACGAAGGGGGCTAGCGTTGTTCGGAATGACTGGGCGTAAAGCGCACGTAGGCGGGTATTTAAGTCAGGGGTGAAATCCCGCAGCTCAACTGCGGAACTGCCTTTGATACTGGGTATCTTGAGTATGGAAGAGGTAAGTGGAATTGCGAGTGTAGAGGTGAAATTCGTAGATATTCGCAGGAACACCAGTGGCGAAGGCGGCTTACTGGTCCATTACTGACGCTGAGGTGCGAAAGCGTGGGGAGCAAACAGG
## 1778                                                         ACAGAGGGTGCAAGCGTTGTTCGGAATTATTGGGCGTAAAGGGAGTGTAGGCGGTGAGTTAAGTCTTCTGTGAAATCTACGGGCTTAGCCTGTAACGTGCAGAGGAAACTGGCTGACTTGAGTATCGGAGAGGGCAGGGGAATACCCAGTGTAGAGGTGAAATTCGTAGAGATTGGGTGGAACACCAGTGGCGAAGGCGCCTGCCTGGCTGATGACTGACGCTGAGACTCGAAAGCATGGGGAGCAAACAGG
## 1779                                                         ACGGAGGGTGCAAGCATTAATCGGATTTATTGGGCGTAAAGGGCGCGTAGGCGGGAAAGTTAGTCAAATGTGAAATCCCGGAGCTCAACTCCGGAACTGCATCTGAAACTACTTTTCTTGAGGGTAGGCGGAGAAAATGGAATTCCACGTGTAGCGGTGAAATGCGTAGATATGTGGAAGAACACCTGTGGCGAAAGCGGTTTTCTAGCTTATTCCTGACGCTGAGGCGCGAAAGCAAGGGGATCAAACAGG
## 1782                                                         ACGAAGGGGGCAAGCGTTGTTCGGAATCATTGGGCGTAAAGCGCGTGTAGGCTGTCTATTAAGTCGGATGTGAAAGCCCTCGGCTCAACCGAGGAAGTGCATTCGAAACTGGTAGACTAGAAGATGGTAGAGAAAGGTGGAATTCCTAGTGTAGAGGTGAAATTCGTAGATATTAGGAGGAATACCGGTGGCGAAGGCGGCCTTTTGGGCCATTCTTGACGCTGAGACGCGAAAGCGTGGGTAGCAAACAGG
## 1787                                                         ACGGAGGGAGCGAGCGTTGTTCGGATTTACTGGGCGTAAAGCGTATGTAGGCGGCTTCGTAAGACAGGTGTGAAATCCCAGAGCTTAACTTTGGAACTGCATTTGTGACTGCGAGGCTTGAGTATCGGAGAGGTTGGCGGAATTCCCAGTGTAGAGGTGAAATTCGCAGATATTGGGAGGAACACCGATGGCGTAAGCAGCCAACTGGACGATTACTGACGCTGAGATACGAAAGCGTGGGGAGCAAACAGG
## 1788                                                         ACGTAGGTGGCAAGCGTTGTCCGGATTTATTGGGTTTAAAGGGTGCGTAGGCGGTTTATTAAGTCAGTGGTGAAAGACGATAGCTTAACTATCGAGGTGCCATTGATACTGATGAACTTGAGTGCACACGAGGTAAGCGGAATGGACGGTGTAGCGGTGAAATGCATAGATATCGTCCAGAACACCGATAGTGAAGACAGCTTACCAGAGTGCAACTGACGCTGAGGCACGAAAGTGTGGGGATCAAACAGG
## 1790                                                         ACAGAGGGACCTAGCGTTGTTCGGAATGACTGGGCGTAAAGGGTGCGTAGGTGGTTGTTTAAGTAGAGTGTTAAATCCTTGGGCTCAACCCAAGAATCGCATTCTATACTGGATGACTGGAGTATGGGAGAGGAGATTGGAATTCCTAGTGTAGAGGTGAAATTCGTAGATATTAGGAGGAACACCAGAGGCGAAGGCGAATCTCTGGCCCATTACTGACACTGAGGCACGAAAGCGTGGGGAGCAAACAGG
## 1791                                                        ACAGAGGGTGCAAGCGTTAACCGGAATGATTGGGTGTAAAGGGTCTGTAGGTGGTTTCGGAAGTCGAGTGTCAAATCCCAGGGCTCAACCCTGGACCGGCATTCGAGAACTCCGAGACTTGAGTTCGGTAGGGGCAGAGGGAATTTCTTGTTTAGAGGTGAAATTCATAGATATAAGAAAGAACACCAGTGGCGAAGGCGCTCTGCTGGGCCGATACTGACACTGAGAGACGAAAGCCAAGGGAGCAAATAGG
## 1798                                                         ACAGAGGGTGCAAACGTTGTTCGGAATCACTGGGCATGAAGCGTACGTAGGCGGCCTCGTAAGTGTCTGATGAAATCCCACGGCTTACCCGTGGAACGGTCAGGCATACTGCGAGGCTCGAGGATGGTAGAGGAGAGCGGAACTCTTGGTGGAGCGGTGAAATGCGTAGATATCAAGAGGAACACCGGTGGCGAAGGCGGCTCTCTGGTCCATGACTGACGCTGAGGTACGAAAGCCAGGGGAGCAAACGGG
## 1800                                                         ACGAATGGGGCAAGCGTTGTTCGGAATCACTGGGCTTAAAGCGTGCGTAGGCGGACTCACAAGTCAGAAGTGAAATCCCCGGGCTCAACCCGGGAATTGCTTTTGAAACTGTGAGTCTAGAATCCCGGAGAGGTTAGGAGAATTCCGAGTGTAGAGGTGAAATTCGCAGATATTCGGAGGAATACCAGTGGCGTAGGCGCCTAACTGGACGGGCATTGACGCTGAGGCACGAAAGCGTGGGTAGCAAACAGG
## 1802                                                          ACGTAGGGGGCTAGCGTTATCCGGATTTACTGGGCGTAAAGGGTGCGTAGGCGGTCTTTTAAGTCAGGAGTGAAAGGCTACGGCTCAACCGTAGTAAGCTCTTGAAACTGGAGGACTTGAGTGCAGGAGAGGAGAGTGGAATTCCTAGTGTAGCGGTGAAATGCGTAGATATTAGGAGGAACACCAGTAGCGAAGGCGGCTCTCTGGACTGTAACTGACGCTGAGGCACGAAAGCGTGGGGAGCAAACAGG
## 1803                                                         ACGAGAGCCTCAAGCGTTATCCGGAATTATTGGGCGTAAAGGGTGTGTAGGTGGTTGTGTTAGTCGAATGTTAAAAATCCAAGCTCAACTTGGAAGCAGCGTTCGAAACGGCACGACTAGAGTAGAGGAGAGGTGTACGGAACTCATGGTGTAGGGGTGAAATCCGTTGATATCATGGGGAACACCGAAAGCGAAGGCAGTACACTGGCCTCTTACTGACACTGAAACACGAAAGCGTGGGTAGCGAATGGG
## 1804                                                        ACGAAGGGGGCGAGCGTTATTCGGAATTACTGGGCGTAAAGGACGCGTAGGCGGTCTATTAAGTCAGCTGTTAAATTCCAGGGCCTAACCCTGGGTATGCAGATGATACTGATTGGACTAGAGTATGGAAGAGAGGGGTGGAATTCTCGGAGTAGCGGTAAAATGCGTAGATCTCGAGAGGAACACCGATGGCGAAGGCAGCCCCTTGGTCCAATACTGACGCTGAGGCGTGAAAGCGTGGGGAGCAAACAGG
## 1806                                                         ACAGAGACCTCAAGCGTTATCCGGAATTACTGGGCGTAAAGCGCGAGCAGCCGGTTGTATTAGTCCGATGTTAAATCACGGGGCTTAACTCCGTGGCAGCATCGGAAACGGTACGACTCGAGCGGGTAAGAGGTATGCGGAACGTGCAGTGTAGGGGTGAAATCCACTGATATTGCACGGAACACCAAAAGCGAAGGCAGCATACTAGTGCTCTGCTGACGGTCATTCGCGAAAGCGTGGGGATCAAAAAGG
## 1812                                                         ACGGAGGGTGCGAGCGTTAATCGGAATGACTGGGCGTAAAGGGTGCGTAGGCGGCGAGATCAGTCAGTCGTGAAATCCCTGGGCTTAACCTGGGAATGGCGATTGAAACGGTAGCGCTAGAGTGGGTCAGAGGATCGTGGAATTCCGGGTGTAGCGGTGAAATGCGTAGATATCCGGAGGAACACCGATGGCGAAGGCAGCGGTCTGGAGCCACACTGACGCTGAGGCACGAAAGCGTGGGGAGCAAACAGG
## 1813                                                      ACGGGGGGGGCAAGCGTTATTCGTCTTTATTGGGCGTAAAGGGTCCGTAGGTGACATTTTACTGTTTAGATTTACAAATGGCGGTTTAGCTGCATTTTGGTCTTTTCAAAAATTATTTGTTTGGGTTTAATGGGGGATAGCGGTACTCTCGATGGAGAGGTGAAATTCGTGGATATCGATGGGGACCACCAGAGGCGAAGGCGGCTATCTATTTTAAACTGACGCTGAGGGACGAAAGCGTGGGGAGCAAACGGG
## 1815                                                          ACATATGGTGCAAGCGTTATCCGGATTTATTGGGCGTAAAGCGCGCGTAGGCGGTTTGGTAAGTTAGAAGTTAAATTCCAGGGCTCAACCCTGGTTCGCTTTTAAAACTGCCAAGCTAGAGTTTTGGAGAGGGAAGGGGAACTAACAGAGTAGCGGTGAAATGCGTAGATACTGTTAGGAACACCAGTGGCGAAGGCGCCTTCCTGGACAATAACTGACGTTAAAGTGCGAAAGCGTGGGGAGCAAACAGG
## 1817                                                        ACATGTGGGGCGAGCGTTGTTCGGAATCACTGGGCATAAAGGGTGCGTAGGCGGTGTCGTACGTCTGGGGTGAAATCCTTCGGCCCAACCGGAGGTCTGCCTTGGATACGGCGATGCTTGAGTGCGTGAGGGGTAACCGGAACGCGTGGTGTAGCGGTGAAATGCGTAGATATCACGCGGAAGGCCAACGGCGAAGGCAGGTTACTGGCGCGCAACTGACGCTGAGTGCACGAAAGCGTGGGGAGCAAACAGG
## 1818                                                        ACGAGGGGGGCAAGCGTTGTTCGGATTTACTGGGCGTAAAGGGTGCGTAGGCGGCATACTAAGTTAGTTGTTTAATCTTGCAGCTCAACTGCAAATCTGCGACTAAGACTGGTTGTGCTAGAGTTTGGAAGAGAGAAGTGGAATTCTCGGAGTAGCGGTTAAATGCGTAGATCTCGAGAGGAACACCGGTGGCGAAGGCGGCTTCTTGGTCCATGACTGACGCTAAAGCACGAAAGCGTGGGGAGCAAACAGG
## 1819                                                         ACAGAGGGTGCGAGCGTTAATCGGAATTACTGGGCGTAAAGCGCACGTAGGCGGCTTCGTAAGTCGGATGTGAAATCCCCGGGCTCAACCTGGGAACTGCATTCGAAACTGCGATGCTTGAGTATGGGAGAGGACAGCGGAATTCCGGGTGTAGCGGTGAAATGCGTAGATATCCGGAGGAACATCAGTGGCGAAGGCGGCTGTCTGGCCCAATACTGACGCTCAGGTGCGAAAGCGTGGGGAGCAAACAGG
## 1820                                                         ACAGAGGCCTCAAGCGTTGTTCGGAATTACTGGGCGTAAAGCGAATGTAGGTGGTGACGTAAGTCAGACGTGAAAGCCCTGGGCTCAACCCAGGAAGTGCGTTTGATACTGCGTTGCTGGAGTACGATAGAGGAAAGTGGAATTTCTGGTGTAGGGGTGAAATCCGTAGATATCAGAAGGAATACCAATTGTGAAGACAGCTTTCTGGGTCGTAACTGACACTGAGATTCGAAAGCGTGGGGAGCAAACAGG
## 1821                                                         ACGGAGGGTGCAAGCGTTATCCGGATTTATTGGGTTTAAAGGGTCCGTAGGCGGACCATTAAGTCAGTGGTGAAAGCCCATAGCTCAACTATGGAATTGCCATTGATACTGGTGGTCTTGAGTACAGTTGAAGTTGGCGGAATGTGTGGTGTAGCGGTGAAATGCTTAGATATCACACAGAACACCGATTGCGAAGGCAGCTAACTAAGTTGTAACTGACGCTGAGGGACGAAAGTGTGGGGATCAAACAGG
## 1823                                                         ACAGAGGGTGCAAGCGTTAATCGGAATTACTGGGCGTAAAGGGTCTGTAGGCGGATTATTAAGTCGGATGTGAAATCCCCGGGCTCAACCTGGGAACGGCATTCGATACTGGTAGTCTAGAGTGTGCAAGAGGATGGGGGAATTCCTGGTGTAGCAGTGAAATGCGTAGAGATCAGGAGGAACATCAATGGCGAAGGCACCTTTCTGGGGCAACACTGACGCTGAGAGACGAAAGCGTGGGGAGCAAACAGG
## 1824                                                         ACGAAGGTCCCGAGCGTTATCCGGATTTATTGGGTTTAAAGGGTGCGTAGGCGGAATGATAAGTCAGTGGTGAAATCTTACAGCTTAACTGTAAAACTGCCATTGATACTGTCGTTCTTGAGTACAAATGATGTGGGCGGAATGTGCCATGTAGTGGTGAAATACTTAGATATGGCACAGAACACCGATTGCGAAGGCAGCTCACAAAATTGTAACTGACGCTGAGGCACGAAAGCGTGGGGATCAAACAGG
## 1825                                                         ACGTAGGGTGCAAGCGTTAATCGGAATTACTGGGCGTAAAGGGTGCGTAGGCGGCACGGAAAGTTCGGGGTGAAATCTCCAGGCTCAACTTGGAAACTGCCCTGAATACTTCCGTGCTCGAGGGTCGCAGAGGAGATTGGAATTCTCGGTGTAGCAGTGAAATGCGTAGATATCGAGAGGAACATCAGTGGCGAAAGCGAATCTCTGGGCGGCATCTGACGCTGAGGCACGAAAGCCAGGGGAGCAAACGGG
## 1827                                                         ACAGAGGGTGCAAGCGTTGTTCGGAATTATTGGGCGTAAAGGGAGTGTAGGCGGTTTTCTAAGTCTTCTGTGAAATCTACGGGCTTAGCCTGTAACGTGCAGAGGAAACTGGAAGGCTTGAGTGTCGGAGAGGATAGGGGAATACCCAGTGTAGAGGTGAAATTCGTAGAGATTGGGTGGAACACCAGTGGCGAAGGCGCCTATCTGGCTGACAACTGACGCTGAGACTCGAAAGCATGGGGAGCAAACAGG
## 1829                                                         ACAGAGGGTGCAAGCGTTGTTCGGAATTATTGGGCGTAAAGGATGCGTAGGCGGCTTTATAAGTCTGTTGTGAAATACCAAAGCTTAACTTTGGAACTGCAATAGATACTGTAAGGCTTGAGATCGGGAGAGGTAAATGGAATTTCTAGTGTAGGGGTCAAATCCGTAGATACTAGGAAGAACACCAGAGGCGAAGGCGATTTACTGGAACGATTCTGACGCTGAGGCATGAAAGCGTGGGGAGCAAACAGG
## 1831                                                         ACGTAGGAGGCGAGCGTTATCCGGAGTTACTGGGCGTAAAGCGTGCGCAGGCGGTGCCATAAGTCAGGTGTGAAAGGTTCCGGCTCAACCGGGAAAACGCGCTTGAGACTGTGGGACTAGAGTGGGATAGAGGGACGTGGAATTCCGGGTGGAGCGGTGAAATGTGTAGAGATCCGGAGGAACACCAGAGGCGCAAGCGGCGTCCTGGGTCCCAACTGACGCTGAGGCACGAAAGCTTGGGGAGCGAACGGG
## 1832                                                         ACGTAGGGTGCAAGCGTTAATCGGAATTACTGGGCGTAAAGCGTGCGCAGGCGGACTTTTAAGCCAGATGTGAAAGCCCCGAGCTTAACTTGGGAATTGCATTTGTGACTGCACGGCTAGAGTGTGTCAGAGGGGGGTAGAATTCCACGTGTAGCAGTGAAATGCGTAGATATGTGGAGGAATACCGATGGCGAAGGCAGCCCCCTGGGATAACACTGACGCTCATGCACGAAAGCGTGGGGAGCAAACAGG
## 1837                                                         ACGAAGGGTGCGAGCGTTGTTCGGAATTACTGGGCGTAAAGGGTTCGTAGGCGGGAATGCAAGTCAAGTGTGAAATCCCCGGGCTCAACCTGGGAACTGCCTTTGTGACTGCATAGCTGGAGTGCGGCAGAGGGGGATGGAATTCCGCGTGTAGCAGTGAAATGCGTAGATATGCGGAGGAACACCGATGGCGAAGGCAATCCCCTGGGCCTGCACTGACGCTCATGCACGAAAGCGTGGGGAGCAAACAGG
## 1840                                                         ACGAGGGGGGCAAGCGTTATTCGGAATTACTGGGCGTAAAGAGCGTGTAGGTGGCTTGTTAAGTCGGATGTTAAATCTCACAGCCTAACTGTGAGGCGGCATTGGATACTGATGAGCTAGAGGATAAGAGAGAGAAGTGGAATTCTCGGAGTAGCGGTTAAATGCGTGGATCTCGAGAGGAACACCGATGGCGAAGGCAGCTTCTTGGCTTATTTCTGACACTAAAATGCGAAGGCGTGGGGAGCAAACAGG
## 1843                                                         ACAGAGGCCTCGAGCGTTGTTCGGAATTACTGGGCGTAAAGGGAATGTAGGTGGTCTTGTAAGTCAGATGTGAAAGCCCTGGGCTCAACCCGGGAACTGCATTTGATACTGCAAGGCTTGAGTGCAGGAGAGAAGAGCGGAATTTCTAGTGTAGGAGTGAAATCCGTAGATATTAGAAGGAATACCGATGGCGAAGGCAGCTCTTTGGCCTGTAACTGACACTGAGATTCGAAAGCATGGGTAGCAAACAGG
## 1844                                                         ACGAGGGATCCTAGCGTTGTTCGGAATCATTGGGCGTAAAGCGGGTGTAGGTGGCTATGTAAGTCAGATGTGAAAGCCCCGGGCTCAACCTGGGAACTGCATTTGAAACTGTCTGACTAGAGTCTTGTAGAGGGGGGTAGAATTCCAGGTGTAGCGGTGAAATGCGTAGAGATCTGGAGGAATACCGGTGGCGAAGGCGGCCCCCTGGACAAAGACTGACGCTCAGGTGCGAAAGCGTGGGGAGCAAACAGG
## 1845                                                          ACGTAGGGGGCAAGCGTTATCCGGAATTACTGGGCGTAAAGGGTGCGTAGGTGGCCTGACAAGTCAGGGGTCAAAGGCAACGGCTCAACCGTTGTAAGCCCTTGAAACTGTCGGGCTTGAGTTCAGGAGAGGAAAGCGGAATTCCTAGTGTAGCGGTGAAATGCGTAGATATTAGGAGGAACACCAGTGGCGAAGGCGGCTTTCTGGACTGATACTGACACTGAGGCACGAAAGCGTGGGGAGCAAACAGG
## 1848                                                        ACAGAGGGTGCAAGCGTTGTCCGGAATGATTGGGCGTAAAGCATCTGTAGGTGGTCATTAAAGTCAACTGTTAAATCCCAGAGCTCAACTTTGGACAAGCAGTTGAGTACTTAGTGACTTGAGTATGGTAGGGGCAGAGGGAATTCCTAGTGGAGCGGTGAAATGCACAGAGATTAGGAAGAACACCAGTGGCGAAAGCGCTCTGCTGGGCCATGACTGACACTGAGAGATGAAAGTTGGGGGAGCGAAAAGG
## 1851                                                        ACAGAGGGAGCAAGCGTTATCCGGAATGATTGGGCGTAAAGCGTTTGTAGGTGGCTTGTAAAGTCTGCTGTCAAATATCAGGGCTTAACCTTGAGAAGGCAGCGGAGTACTCACAAGCTTGAGTGTGGTAGAGGTAGAGGGAATTCCATGTGGAGCGGTGAAATGCGTAGAGATATGGAAGAACACCAGCGGCGAAGGCGCTCTACTGGGCCAAGACTGACACTGAGAAACGAAAGCTAGGGGAGCGAATAGG
## 1856                                                           ACGTAAGGTACAAGTGTTACTCGGATTTATTGGGTTTAAAGCGTGCGTAGATGGATTTTTAAGTTAGTAGTAAAATATTAGAGCTCAACTCTATTAAGCTATTAATACTGATTATCTTGAGTTTTTTTTAGGTAGGCATAATTTGCAGTGTAGTGGTGGAATACTTAAATATTGTAAGGAATATCTTTAGCGTAGGCAGCTTACTAGGGAAAACTGACATTGATAGCACGAAAGGCGGGTAGCAAACAGG
## 1857                                                         ACGGAGGGGGCTAGCGTTGTTCGGAATCACTGGGCGTAAAGGGCGCGTAGGCGGCTTTGCAAGTCAGGTGTGAAATCCCGGGGCTTAACCCCGGAACTGCACTTGAAACTGCATGGCTGGAGTCTCGGAGAGGATGGTGGAATTCCTAGTGTAGAGGTGAAATTCGTAGATATTAGGAGGAACACCGATGGCGAAGGCAGCCATCTGGACGAGTACTGACGCTGAGGCGCGAAAGCGTGGGGAGCAAACAGG
## 1858                                                         ACAGAGGATGCGAACGTTGCTCGGAATTACTGGGCGTAAAGCGCGCGCAGGTTGCCTCGTAAGTCGTGTGTGAAATCCCTCTGCTCAACGGAGGAAGTGCACGGGATACTGCGAGGCTGGAGTACAGAAGAGGACAGCGGAATTCCCGGTGTAGAGGTGAAATTCGTAGATATCGGGAGGAACACCAGTGGCGAAGGCGGCTGTCTGGGCTGATACTGACACTGAGGCGCGAAAGCGTGGGAAGCAAACAGG
## 1859                                                         ACGTAGGATCCGAGCGTTGTCCGGAATTACTGGGCATAAAGGGAGCGTAGGCGGATTTGTGCGTCAGAGGTGAAATCCCACAGCTTAACTGTGGAACTGCCTTTGATACGGCGAATCTTGAGTATGAGAGAGGCGAGTAGAATTCCAGGTGTAGCGGTGAAATGCGTAGATATCTGGAGGAATACCAGTGGAGAAGTCGGCTCGCTGGCTCATAACTGACGCTGAGGCTCGAAAGCGTGGGGATCAAACAGG
## 1861                                                         ACGTGGGGTGCAAGCGTTATCCGGATTTATTGGGCGTAAAGAGTTCGTAGGCGTCTAGTTAAGTTTTGTTTGAAATCTCGGGGCTTAACCCCGAAATTGGGCGAAATACTGGCTAGATCGAGTCTGTTAGGGGGTACTGGAACTGTCAGTGTAGCAGTGAAATGCGTTGATATTGACAGGAACACCAAGGGCGAAGGCAGGTACCTGGGACATGACTGACGCTGAGGAACGAAAGCTAGGGGAGCGAAAGGG
## 1862                                                         ACGTAGGGGGCAAGCGTTGTCCGGAATTACTGGGCGTAAAGCGCACGCAGGCTGAATTATAAGTCGGTAGTAAAAGGCGGGGGCTCAACTTCCGTATATCTATCGATACTGTAAATCTAGAGTATGTGAGAGGGAAGCGGAATTCCCGGTGTAGCGGTGAAATGCGTAGATATCGGGAGGAACACCAGTGGCGAAGGCGGCTTCCTGGCACAATACTGACGCTCATGTGCGAAAGCTAGGGCAGCGAACGGG
## 1864                                                         ACAGAGAGTGCAAACGTTGTTCGGAATTACTGGGCTTAAAGCGCACGTAGGCGGCTCCGCAAGTGTCGGGTGAAATCCCACGGCTTACCCGTGGAACTGCCCGGCAGACTGCGGAGCTAGAGGATGGTAGGGGAGAGCGGAACTCCAGGTGGAGCGGTGAAATGCGTAGATATCTGGAGGAACACCGGTGGCGAAGGCGGCTCTCTGGTCCATAACTGACGCTGAGGTGCGAAAGCTAGGGGAGCAAACGGG
## 1866                                                         ACGGAGGGTGCAAGCGTTACTCGGAATTACTGGGCGTAAAGGGCGCGTAGGCGGCTTGTTAAGTTTGATGTGAAAGCCCCGGACTCAATTCGGGAAGTGCATTGAATACTGGCGAGCTGGAGTTTGGGAGAGGAAAGCGGAATTCCTGGTGTAGAGGTAAAATTCGCAGATATCAGGAGGAACAACAGTAGCGAAGGCGGCTTTCTGGACCAATACTGACGCTGAGGCGCGAAAGCGTGGGGAGCAAACAGG
## 1867                                                         ACGTAGGTGGCAAGCGTTGTCCGGATTTATTGGGTTTAAAGGGTGCGTAGGCGGCCCTGTAAGTCAGTGGTGAAATACTCCAGCTTAACTGGAGGGGTGCCATTGATACTGCAGGGCTTGAGTAAAGTTGAAGTTGGCGGAATTGACGGTGTAGCGGTGAAATGCTTAGATATCGTCAAGAACACCGATAGTGAAGACAGCTAACTAAGCTTTAACTGACGCTGAGGCACGAAAGTGTGGGGATCAAACAGG
## 1868                                                         ACAGAGGACCCGAGCGTTAATCGGAATCACTGGGCTTAAAGGGTGCGTAGGCGGACCCGAAGGTGCATCGTGAAATCCCTTGGCTCAACCAAGGAATTGCGGTGCATACCACGGGTCTTGAGGCAAGTAAGGGCGGTCGGAACGCTCGGTGGAGCGGTGAAATGCGTAGATATCGAGCGGAACGCCGAAGGTGAAGACAGGCCGCTGGGCTTGACCTGACGCTGAGGCACGAAAGCGTGGGGATCAAACAGG
## 1869                                                         ACAGAGGGTGCGAGCGTTAATCGGAATAACTGGGCGTAAAGGGTGCGTAGGCGGAACATTAAGTCAGATGTGAAATCCCCGGGCTTAACCTGGGAACTGCATTTGAAACTGATGTTCTAGAGTAGAGGAGAGGAGAGTGGAATTTCAGGTGTAGCGGTGAAATGCGTAGATATCTGAAGGAACACCAGTGGCGAAGGCGGCTCTCTGGCCTCATACTGACGCTGAGGCACGAAAGCGTGGGGAGCAAACAGG
## 1872                                                         ACAGGGGATGCAAGTGTTATCCGGAATTATTGGGCGTAAAGCGTTTGTAGGTCGATTTTTAAGTCTTTTGTAAAATTTTTAGGCTCAACCTAAAGCATGCAAAAGAAACTAAATATCTAGAGTTTAATAAAGGTAAAGGGAATTTCCGGTGGAGCGGTGAAATGTGTTGAGATCGGAAGGAACACCAAAGGCGAAGGCACTTTACTTGGTTAAAACTGACACTGAGAAACGAAAGCTAGGGGAGCAAATGGG
## 1874                                                         ACGTAGGGAGCAAGCGTTGTCCGGATTTATTGGGCGTAAAGAGCTCGTAGGCGGTTCCGTAAGTCGAGTGTGAAAAATCTGGGCTCAACCCAGTGAAGCACTCGATACTGCGGTGACTAGAGTACGGTAGAGGAGTGTGGAATTTGTGGTGTAGCGGTGAAATGCGCAGATATCACAAGGAACACCAACGGCGAAGGCAGCACTCTGGGCCGGTACTGACGCTGAGGAGCGAAAGCGTGGGGAGCAAACAGG
## 1878                                                         ACGAAGGCCCCAAGCGTTATCCGGAATTACTGGGCGTAAAGCGTCTGTAGGTGGTTTCTTAAGTCTTTCCGTGAAACTTCAGGGCTCAACCCTGAATTGCGGGAGAAACTGGGAGACTAGAGTGTGGGAGGGGGAAGCAGAACGGTAAGAGTAGGGGTGCAATCCGTTGATACTTACCAGAATACCAAAAGCGAAGGCAGCTTCCTGGAACACTACTGACACTGAGAGACGAAAGCGTGGGGAGCAAAAGGG
## 1880                                                         ACGTGAGTGGCAAGCGTTATCCGGAATTACTGGGCGTAAAGAGCGCGTAGGCGTCTCGATAAGTCAGGGGTGAAAAGTCAGGGCTTAACCTTGGCCTTGCCTTTGAAACTATCGAGATAGAGTATTGGATAGGAGGGCGGAACTACAAGTGTAGCGGTGGAATGCGTAGATATTTGTAGGAATGCCGATGACGAAGGTAGCTCTCTGGACAAATACTGACGCTGAGGTGCGAGAGCTAGGGGAGCAAACAGG
## 1882                                                         ACAGAGGTGGCGAGCGTTGTTCGGAATTACTGGGCTTAAAGGGCGAGTAGGCGGTGGAGCAAGTCTGAGGTGGAATCCTACGGCTTAACCGTAGAACTGCCTTAGAAACTGCTTTACTTGAGTCAGTGAGGGGAGGGCGGAATTCCTGGTGTAGCGGTGAAATGCGTAGATATCAGGAGGAAGGCCTGTGGCGAAAGTGGCCCTCTGGCACTGTACTGACGCTGAGGCGCGAAAGCGTGGGGAGCAAACAGG
## 1883                                                         ACAGAAGCCCCGAGCATTACCCGGATTTACTGGGCGTAAAGGATGTGTAGGTGGTGTAATTAGTCGGGCGTTAAATCCTGGGGCTCAACCCCAGGCTCGCGTTCGAAACGGTTACACTAGAGGAAGTGAGAGGTGTGTGGAACTCAAGGTGTAGGGGTGAAATCCGTTGATATCTTGGGGAACACCAAAAGCGAAGGCAACACACTGGCACTCTCCTGACACTGAAACATGAAAGCGTGGGTAGCGAATCGG
## 1885                                                         ACGGAGGGTGCAAGCATTAATCGGATTTATTGGGCGTAAAGGGCGCGTAGGCTGGAAGGAAAGTCAGATGTGAAATGCCGGGGCTCAACCCCGGAACTGCATTTGAAACTTCCTTTCTTGAGGGTAGGCGGAGATAATGGAATTCCACAAGTAGCGGTGAAATGCGTAGATATGTGGAAGAACACCGGTGGCGAAGGCGATTATCTAGCTTATTCCTGACGCTGAGGCGCGAAAGCTAGGGGAGCAAACAGG
## 1886                                                         CAGAGGGAGCTAGCGTTTTTCGGATTTATTGGGCGTAAAGAGCGCGTAGGCGGTTAGCCAAGTTGATAGTAAAAGCCCAGGGCTTAACCCTGGAACGTCTATCAAGACTGGTTGACTAGAGTTCGGTAGAGGATGATGGAATTCTTGGTGTAGGGGTGAAATTCTTAGATATCAGGAGGAACACCGGTGGCGAAAGCGATCATCTGGGCCGTAACTGACGCTAATAGCGCGAAAGCGTGGGGAGCAAACAGG
## 1888                                                         ACGTAGGAAGCAAGCGTTGTCCGGAATTACTGGGCGTAAAGAGCTCGTAGGCGGGATCGTAAGTCTTAGAGTAAATCTCACGGCTCAACCGTGGGGCCTTCTGAGATACTGCGATTCTTGAGGATGTCAGAGGAAAGTGGAATTCCCGGTGTAGCGGTGAAATGCGTAGATATCGGGAGGAACACCAGTGGCGAAGGCGACTTTCTGGGACATTCCTGACGCTGAGGAGCGAAAGCGTGGGGAGCAAACGGG
## 1890                                                         ACGAAGGGGGCTAGCGTTGTTCGGAATGACTGGGCGTAAAGGGCGCGTAGGCGGTGAGGTACGTTTGATGTGAAAGTCCTGGGCTTAACCTGGGGATTGCATTGAATACGGCTTTGCTAGAGTGCGAGAGGGGAAGATGGAATTGCTGGTGTAGAGGTGAAATTCGTAGATATCAGCAAGAACACCGGAGGCGAAGGCGATCTTCTGGCTCGCAACTGACGCTGAGGCGCGAAAGCGTGGGGAGCAAACAGG
## 1893                                                         ACGTAGGGTGCAAGCGTTAATCGGAATTACTGGGCGTAAAGCGTGCGCAGGCGGTTATGTAAGACAGAGGTGAAATCCCCGGGCTCAACCTGGGAACGGCCTTTGTGACTGCATAGCTAGAGTACGGTAGAGGGGGATGGAATTCCGCGTGTAGCAGTGAAATGCGTAGATATGCGGAGGAACACCGATGGCGAAGGCAATCCCCTGGGCCTGCACTGACGCTCATGCACGAAAGCGTGGGGAGCAAACAGG
## 1895                                                         ACGAGGGGTGCAAGCGTTGCTCGGAATCATTGGGCGTAAAGGGTAGGTAGGTGGTCTCATTTGTCTGAGGTGAAAGCCTTGGGCTCAACTCAAGAAGTGCCTCGGAAACGGTGAGACTAGAGTGCTGGAGGGGGTCGTGGAATTCCCGGTGTAGCGGTGAAATGCGTAGAGATCGGGAGGAACACCAGAAGCGAAGGCGGCGACCTGGACAGCAACTGACACTCAACTACGAAAGCGTGGGTAGCAAACAGG
## 1897                                                         ACGTAGGAGACAAACGTTATCCGGAGTTACTGGGCGTAAAGGGCGTGCAGGTGGTTCGTTAAGTCTGGCGTGAAAGCCTCCGGCTCAACCGGGGAAAGTCGTCGGAGACTGACGAACTAGAGGGCAGTAGAGGCGTGTGGAATTCCGGGTGTAGTGGTGAAATGCGTAGAGATCCGGAGGAACACCAGTGGCGAAGGCGACACGCTGGGCTGCACCTGACACTGAGACGCGACAGCATGGGGAGCGAACGGG
## 1900                                                         ACGAAGGGTGCAAGCGTTATTCGGATTTATTGGGCGTAAAGGGTTCGTAGGTGGGAAAGCAAGTTAATTGTGAAATCCCCGGGCTTAACCTGGGACGTGCAGTTAAAACTGCTTTTCTTGAGTGTGGAAGAGGGTAGTGGAATTGCTGGTGTAGGAGTGAAATCCGTAGATATCAGCAGGAACATCAGAGGCGAAGGCGACTACCTGGTCCTTCACTGACACTGAGGAACGAAAGCGTGGGGAGCAAACAGG
## 1902                                                        ACAGAGACCCCAAGCGTTATCCGGAATTATTGGGCGTAAAGGGTGAGTAGGTTGTTTTATTAGTCAGATGTTAAAGTTCTCGGGCTCAACCTGGGAAATGCATTTGAAACGGTAAAACTAGAATATGTAAGAGGTGAGTGGAACTCTATGTGTAGGGGTAAAATCCGTTGATATATAGGGGAACACCAAAAGCGAAGGCAGCTCACTGGTACATTATTGACACTGAATCACGAAAGCGTGGGTAGCGATACGG
## 1905                                                         ACGAAGGGTGCAAGCGTTGTTCGGATTTATTGGGCGTAAAGCGCGCGCAGGCGGAAAAGCAAGTCAGATGTGAAATCTCGGGGCTTAACCCCGAACGTGCGTCTGAAACTACTTTTCTAGAGTAGCGAAGGGGGAAGGGGAATTTCACATGTAGGGGTAAAATCCGTAGAGATGTGAAGGAACACCAGAGGCGAAGGCGCCTTCCTGGTCGCTAACTGACGCTGAGGCGCGAAAGCGTGGGGATCAAACAGG
## 1911                                                         ACGAGAGCCCCGAGCGTTATCCGGAATTACTGGGCGTAAAGGGTGAGTAGGTGGCGTTATTAGTCGAGTGTTAAATATCTGTGCTTAACATGGATGCTGCATTCGAAACGGTAATGCTAGAGGGTGTGAGAGGTCTGTGGAACTCATGGTGTAGGGGTGAAATCCGTTGATATCATGGGGAACACCAAAAGCGAAGGCAACAGACTGGCACATTCCTGACACTGAATCACGAAAGCGTAGGTAGCGAATGGG
## 1912                                                         ACAGAGGATGCAAGCGTTATCCGGAATCACTGGGCATAAAGCGTCTGTAGGTTGCCTACCAAGTCTGCTGTTAAAGATCAGGGCCTAACCCTGGGAAAGCAGTGGAAACTAGTAGGCTTGAGTGTGGTAGAGGTAGAGGGAATTCCTGGTGTAGCGGTGAAATGCGTAGATATTAGGAAGAACACCAATGGCGAAAGCACTCTACTGGGCCACAACTGACACTGAGAGACGACAGCTAGGGGAGCAAATGGG
## 1914                                                         ACGTAGGCCTCGAGCGTTATCCGGATTTATTGGGCGTAAAGAGCATGTAGGAGGCTTTGCAAGTCTTTGGTTAAAGACCACCGCCTAACGGTGGGAGTGCCGAGGAAACTGCGGAGCTGGAGGAGGTTAGAGGTGCATAGAACGCATGGTGTAGGGGTGAAATCCGTTGATATCATGCGGAATACCAAAGGCGAAGGCAGTGCACTGGGACCTTCCTGACTCTGAGATGCGAAAGCGTGGGGAGCAAAAAGG
## 1917                                                         ACAGAGGGTGCAAGCGTTGTTCGGAATTACTGGGTGTAAAGGGCGCGTAGGCGGCTATTTAAGTCAGATGTGAAAACCCGAAGCTCAACTTCGGACGTGCATTTGAAACTATTTAGCTAGAGTGCAGTAGAGGATGGTGGAATTCCCAGTGTAGAGGTGAAATTCGTAGAGATTGGGAGGAACACCGGTGGCGAAGGCGGCCATCTGGGCTGCAACTGACGCTGAGGCGCGAAAGCGTGGGGATCAAACAGG
## 1919                                                         ACGGAGGATGCAAGCGTTATCCGGAATTATTTGGCGTAAAGCGTCTGTAGGTGGTTTGGTAAGTCCTGTGTTAAATTTTAGAGCTTAACTCTATAAAGGCATCGGAAACTGCTAGACTTGAGTCCGGTGGAAGTAAAGGGAATTCCCAGTGTAGCGGTGAAATGCGTAGATATTGGGAGGAACACCAAAGGCGAAAGCACTTTACTAGGCCGATACTAACACTCAGAGACGAAAGCTAAGGTAGCGAACGGG
## 1920                                                         ACGGAGGTTCCAAGCGTTGTTCGGAATTACTGGGCGTAAAGGGAGCGCAGGCGGACGGGTCAGTTTAAAGTGGAATCCCAGGGCTCAACCCTGGACGTGCTTTGGATACTGCCTGTCTTGAGTTCGTGAGAGGAAAGTGGAATTCAGGGTGTAGCGGTGAAATGCTTAGATATCCTGAGGAACACCAGTGGCGAAAGCGACCTTCTGGCACGAAACTGACGCTGAGGCTCGAAAGCATGGGTAGGGAACGGG
## 1925                                                           ACGAGAAGACCTAGCGTTAATTGGAATTATTGGGCGTAAAACGTAAATAGACTGTTATTGTTTGTTAAAAATTAAATTTTAAACTGTATTTATAACGTTTTTAATTACAAATTTCTAGAGTGAAAGAAAAGATTATAGAACTTTAAAAGGAAAAATAAAATTTGTAGATTTTTGAAGGAATGCTAAAAGTGAAAACAATAATCTTGCTTTATACTGACGTTGTTTTACTAAAGCATAGGTAGCAAAAAGG
## 1926                                                          CGTAGGTGGCGAGCGTTGTTCGGATTTATTGGGCGTAAAGGGTCCGCAGGGGGTTGATTAAGTCTGACGTGAAATCTCTGAGCTCAACTCAGAAATTGCGTTGGAGACTGGTTGACTAGAGTGTTGGAGAGGTAATTGGAATTCCAGGTGTAGCGGTGGAATGCGTAGATATCTGGAAGAACATCGAAAGCGAAGGCAGATTACTGGACAACAACTGACCCTCAGGGACGAAAGCATGGGGAGCAAACAGG
## 1930                                                         ACGGGGGGAGCGAACGTTGTTCGGAATCACTGGGCGTAAAGGGCGTGCAGGCGGCTCGGTAAGTGGAAGGTGAAATGCCCTGGCTCAACCAGGGAACTGCCTTCCAGACTGCCGAGCTTGAGTGCGGGAGAGGATGGGAGAATTCCCGGTGTAGCGGTGGAATGCATTGATATCGGGAGGAATACCAGTGGCGAAGGCGCCCATCTGGCCCGACACTGACGCTCAGGCGCGAAAGCTAGGGGAGCAAACAGG
## 1933                                                          ACGTAGGTGGCGAGCGTTATCCGGAATTATTGGGCGTAAAGAGGGAGCAGGCGGCACTAAGGGTCTGTGGTGAAAGATCGAAGCTTAACTTCGGTAAGCCATGGAAACCGTAGAGCTAGAGTGTGTGAGAGGATCGTGGAATTCCATGTGTAGCGGTGAAATGCGTAGATATATGGAGGAACACCAGTGGCGAAGGCGACGATCTGGCGCATAACTGACGCTCAGTCCCGAAAGCGTGGGGAGCAAATAGG
## 1934                                                         ACGGAGGATCCGAGCGTTATCCGGATTTATTGGGTTTAAAGGGTGCGTAGGTGGATTGTTAAGTCAGTGGTGAAAGTTTTCAGCTCAACTGAAAAATTGCCGTTGAAACTGTCGATCTTGAGTGCAAATGAGGTAGGCGGAATGTGTTGTGTAGCGGTGAAATGCTTAGATATAACACAGAACTCCGATTGCGAAGGCAGCTTACTGGGATGCAACTGACACTGATGCACGAAAGCGTGGGTATCAAACAGG
## 1936                                                         ACGTAGGTGGCGAGCGTTACTCGGAATTACTAGGCGTAAAGCGCGTGTAGGCGGGAGCTTAAGTCTGTCGTGAAAGCTCAGGGCTCAACCCTGAAATGCCGGCAGAAACTGGGCTTCTTGAGTGCGGTAGAGGAGATCGGAATTCCAGGTGTAGCGGTGAAATGCGTAGATATCTGGAAGAACGCCAAAGGCGAAGGCAGATCTCTGGGCCGATACTGACGCTGAGACGCGAAAGCTAGGGGAGCAAACAGG
## 1938                                                         ACAGAGACTGCAAGCGTTATCCGGATTTATTGGGTTTAAAGGGTGCGCAGGCGGGCTTGTAAGTCAGTGGTGAAAGACTGCCGCTTAACGGTAGCATTGCCATTGATACTGCAAGTCTTGAGTACATTTGAAGTGGGCGGAATGTGTAGTGTAGCGGTGAAATGCTTAGATATTACACAGAACACCGATAGCGTAGGCAGCTCACTAAGATGCAACTGACGCTCAGGCACGAAAGCGTGGGGATCAAACAGG
## 1939                                                         ACGGGGGGGGCAAGCGTTGTTCGGAATTACTGGGCGTAAAGGGCGCGTAGGCGGCACGGATAGTTGGGTGTGAAAGCCCGGAGCTTAACTCCGGAATGTCACTCAATACTTCCGAGCTAGAGTGTGGAAGGGGCAATCGGAATTCTTGGTGTAGCGGTGAAATGCGTAGATATCAGGAAGAACACCTGTGGCGAAGGCGGATTGCTGGGCTAACACTGACGCTGAGGCGCGAAAGCTAGGGGAGCGAACGGG
## 1942                                                   ACGAAGAGGGTTAGTATTATTCAGAATTATTGGGCGTAAAACGTTTGTAGGGAATTATTAGTATGTAAAAACAAAAAACCTAAATTACACGTTTTTAGTAAAGTTTTTAAATAAATAATTTGAGTATAACTTTGATACGATCATAGAATTTCAGGTGGAGAAATAAAATTCTTAGATCTCTGAAGGAATGCCAAAGGCTAATGCAATGATCTTGATTTATACTGACCTTGATAAACTAAAGCATAGGGAGCAAAGAGG
## 1944                                                         ACGAAGGGAGCTAGCGTTACTCGGAATCATTGGGCGTAAAGCGTGCGTAGGCGGTTTTTTAAGTCGGAAGTGAAAGCCTGGGGCTCAACCTCAGAATTGCTTTCGATACTGGAGAACTAGAGTATCAGAGGGGATGGTGGAATTCCTAGTGTAGAGGTGGAATTCGTAGATATTAGGAGGAACACCGGTGGCGAAGGCGACCATCTGGCTGAATACTGACGCTGTTGCACGAAAGCGTGGGGAGCAAACAGG
## 1946                                                         ACATAGGCCCCAAGCGTTATCCGGAATTACTGGGCGTAAAGAGTTCGTAGGCGGAGAAACAAGTCAGATGTGAAAGATACGGGCTCAACCCGTTATTAGCATCCGAAACTGTTTCCCTAGAATTAGGGAGAGGTAAGCAGAACTGTATGAGTAGGGGTGCAATCCGTTGATACATACAAGAATACCAAAAGCGAAGGCAGCTTACTGGAACTATATTGACGCTGAGGAACGAAAGCGTGGGGAGCGAAAAGG
## 1947                                                         ACGGAGGGTGCAAGCGTTGTTCGGATTTACTGGGCGTAAAGCGTGTCTAGGTGGACTTGTAAGTCAGATGTGAAATCCCCCGGCTTAACTGGGGAGGGTCATTTGAAACTACAAGCCTTGAGTTTCGGGGAGGTGGGTGGAATTCCCAGTGTAGAGGTGAAATTCGTAGAGATTGGGAGGAACATCGGAGGCGAAAGCGGCTCACTAACCGAACACTGACACTGACACACGAAAGCGTGGGTAGCAAACAGG
## 1948                                                      ACGAGAGGGGCAAGCGTTATTCGGAATGATTGGGCGTAAAGAGCACGTAGGTGGTTTTTTTATTTTTTATTTGAAATTTCAAAGCTTAACTTTGAAACAGAATAAATTAAAATAAAAAACTTGAGGAAGTCAAAAGAAAATAGAATTCCTAGAGGAGAGGTAAAATTTGTTGATTTTAGGAGGAATACCAAAAGCGAAAGCAGTTTTCTTGGACTTTCCTGACATTGAGGTGCGAAAGCGTGGGGATCAAAAGGG
## 1951                                                          ACGTAGGTCCCGAGCGTTGTCCGGATTTATTGGGCGTAAAGCGAGCGCAGGTGGTTTATTAAGTCTGGTGTAAAAGGCAGTGGCTCAACCATTGTATGCATTGGAAACTGGTAGACTTGAGTGCAGGAGAGGAGAGTGGAATTCCATGTGTAGCGGTGAAATGCGTAGATATATGGAGGAACACCGGTGGCGAAAGCGGCTCTCTGGCCTGTAACTGACACTGAGGCTCGAAAGCGTGGGGAGCAAACAGG
## 1954                                                         ACGGAGGGTGCAAGCGTTATCCGGATTTACTGGGTTTAAAGGGTGTGTAGGCGGGTTGGTAAGTCAGTGGTGAAAGCCCCGAGCTTAACTTGGGAACTGCCATTGATACTGCCAGTCTTGAATACGGTTGAGGTAGGCGGAATATAACATGTAGCGGTGAAATGCTTAGATATGTTATAGAACACCGATTGCGAAGGCAGCTTACTAAGCCGTGATTGACGCTGAGACACGAAAGCGTGGGGAGCAAACAGG
## 1956                                                         ACGAAGGGTGCAAGCGTTAATCGGAATTACTGGGCGTAAAGCGCGTGTAGGCGGTTGGCTAAGTCAGATGTGAAAGCCCCGGGCTCAACCTGGGAATTGCATTTGATACTGGCTGGCTAGAATGTGGTAGAGGGAGGCGGAATTCCCGGTGTAGCGGTGAAATGCGTAGATATCGGGAAGAACACCAATGGCGAAGGCAGCCTCCTGGGCCAACATTGACGCTGAGACGCGAAAGCGTGGGGAGCAAACAGG
## 1963                                                         ACGGAGGGTGCAAGCGTTGTTCGGAATCATTGGGCGTAAAGCGCGTGTAGGCGGTCTTATAAGTTTGGTGTGAAAGCCCGGGGCTCAACCCCGGAAGTGCACTGAATACTGTTTGACTAGAGGACTGGAGAGGGAAGTGGAATTCCTGGTGTAGGGGTGAAATCCGTAGATATCAGGAGGAACATCGGAAGCGAAGGCGGCTTCCTGGCCAGTAACTGACGCTGAGACGCGAAAGCGTGGGGAGCAAACAGG
## 1965                                                         ACGAGGGGGGTTAGCGTTATCCTAAATGATTGGGTATAAAGAGTTTGTAGGCTGTATTTGTTGTTACTTATAAAATATTTGGATAATCTTTAAATAAAATATGTAATACTTATATACTAGAGTATAACAGGGGAAGACAGAATTTTATAAGGAGGAGTAGAATCCAGTGATTTTTAAAGGAATGTCAGTAGCGAAAGCGGTCTTCTAGGTTTTTACTGACGCTGAGGAACGAAGGCATAGGTAGCAAACAGG
## 1967                                                         ACGAAGGTGACAACCGTTGCTCGGAATCACTGGGCATAAAGCGCGCGTAGGTGGCTGTGTAAGTCCATGGTGAAATCCCTCGGCTCAACCGAGGAACAGCCGTGGATACTGCATAGCTTGAGAACGGGAGGGGTGAGTGGAACTCTGGGTGTAGCGGTGAAATGCGTAGATATCCAGAGGAACGCCGGTGGCGAAAGCGACTCACTGGACCGTTACTGACACTGAGGCGCGAGAGCTAGGGGAGCAAACAGG
## 1968                                                        ACGAAGGGAGTAAGCGTTATCCAAAATGATTGGGCGTAAAGGGTTCGTAGGTTGTAATTGTTGTCGTATGTGAAATCTTAAAATTTTCTTTTAAATGTGCGTACGAAACTCAATTACTTGAGTATATCAGAGGATAACAGAATTTTATATGGAGGAGTAAAATCCGATGATCTGTAAAGGAATGCCAGTGGCGAAAGCGGTTATCTGGGATATATACTGACGCTGAGGAACGAAAGCGTGGGGAGCAAACAGG
## 1969                                                         ACGGAGGGTGCAAGCGTTAATCGGAATAACTGGGCGTAAAGCGCACGCAGGCGGTTGGATAAGTCAGATGTGAAAGCCCCGGGCTCAACCTGGGAACTGCATTTGAAACTGTCTGACTAGAGTCTTGTAGAGGGGGGTAGAATTCCAGGTGTAGCGGTGAAATGCGTAGAGATCTGGAGGAATACCGGTGGCGAAGGCGGCCCCCTGGACAAAGACTGACGCTCATGCACGAAAGCGTGGGGAGCAAACAGG
## 1975                                                         ACAGAGGTCGCAAGCGTTGTTCGGATTTACTGGGCGTAAAGGGCGCGTAGGCGGCAGGGCATGTCAAGTGTGAAATCTCCGGGCTCAACCCGGAAAGTGCGCTTGAAACTACCTTGCTTGAGTCTGGGAGAGGGAATCGGAATATAGGGTGTAGCGGTGAAATGCGTTGATATCCTATAGAACACCGGAGGCGAAGGCGGATTCCTTGAACAGTACTGACGCTGAGGCGCGAAAGTCGGGGGAGCAAACAGG
## 1976                                                         ACGAGAGGTGCGAGCGTTATTCGGAATCATTGGGCGTAAAGGGTGCGTAGACGGTATTGTAAGTCTGTTGTTAAAATTCTTGGCCTAACTGAGAGTCTGCAATGGATACTGCTTTGCTAGAGGGTGAGAGAGAGAAGTGGAATTCTCGGAGTAGCGGTCAAATGCGTAGATCTCGAGAGGAACACCGATGGCGAAGGCAGCTTCTTGGCTCATTTCTGACGTTGATGCACGAAAGCGTGGGGAGCAAACAGG
## 1978                                                         ACGGAGGGGGCTAGCGTTGTTCGGAATTACTGGGCGTAAAGAGCACGTAGGTGGTTAATTTAGTTAGGTGTGAAATCCCTGGGCTTAACCTAGGAACTGCGCTTAATACTGATTAACTTGAGTCCGATAGGGGAAAGTGGAATTCCTAGTGTAGAGGTGAAATTCGTAGATATTAGGAGGAACACCAGAGGCGAAGGCGTCTTTCTGGATCGGTACTGACACTGAGGTGCGAAAGCGTGGGGATCAAACAGG
## 1979                                                         ACGGAGGGTGCAAGCGTTAATCGGAATAACTGGGCGTAAAGGGCATGCAGGCGGTTCATCAAGTAGGATGTGAAATCCCCGGGCTCAACCTGGGAACAGCATACTAAACTGGTGGACTAGAGTATTGCAGGGGGAGACGGAATTCCAGGTGTAGCGGTGGAATGCGTAGATATCTGGAAGAACACCAAAGGCGAAGGCAGTCTCCTGGGCAAATACTGACGCTCATATGCGAAAGCGTGGGTAGCAAACAGG
## 1984                                                         ACGAAGGACCCGAGCGTTATCCGGATTCATTGGGTTTAAAGGGTGCGTAGGCGGAATGATAAGTCAGTAGTGAAATCCTGCAGCTTAACTGCAGAACTGCCATTGATACTGTCGTTCTTGAATGCAACTGATGTGGGCGGAATGTGCCATGTAGTGGTGAAATACTTAGATATGGCACAGAACACCGATTGCGAAGGCAGCTCACAAAATTGTTATTGACGCTGATGCACGAAAGCGTGGGGATCAAACAGG
## 1985                                                         ACAGAGGGTGCAAGTGTTGTTCGGAATCATTGGGCGTAAAGGGCGCGTAGGTGGTCCGGTAAGTCAGATGTGAAAGCCCTGGGCTTAACCCAGGAACTGCATTTGAGACTGCCGGACTAGAGTTCAGTAGAGGATAGTGGAATTCCCGGTGTAGAGGTGAAATTCGTAGAGATCGGGAGGAACACCAGTGGCGAAAGCGGCTATCTGGGCTGCAACTGACACTGAGGCGCGAAAGCGTGGGGAGCAAACAGG
## 1986                                                        CCCGGCGCCACAAGTGGCAATCACGATTATTGGGTCTAAAGCGTCCGTAGCCGGTTATGTAAATCTTTTGTGAAATCGTTGGACTTAACCTAACGGCTTGCAGAAGACACTGCATGACTTGGGACCGGAAGGAGTGTGAAGTATGGAATAGGGAACGGTAAAATGTGATAATCTATTCTAGACTACCAATGGCGAAGGCATCACACTAGGACGGATCCGACGGTGAGGGACGAAAGCTAGGGGAGCGAACCGG
## 1988                                                         ACAGAGACCTCAAACATTGTCCGGATTCATTGGGCGTAAAGCGTCCGCAGGTGTTCTGATAAGTTTTCTGCTAAATCCCAAGGCTTAACCTTGGGTCTGCGGAGAATACTATCAGAATCGAGGTTGGGAGAGGCACACGGAACTGCCGGTGTAGTAGTAAAATGCGTTGATATCGGCAGGAACACCAAAGGCGAAAGCAGTGTGCTGGAACAATTCTGACACTAAGGGACGAAAGCGTGGGGAGCGAATGGG
## 1989                                                         ACGAGAGGTGCAAGCGTTGTTCGGAATGACTGGGCGTAAAGAGCACGTAGGCGGGTTTATAAGTCGGATGTGAAAACCCTGGGCTTAACCTGGGACGAGCATACGAAACTGTAGATCTTGAATGTCTGAGAGATTGGTGGAATTCCAGGTGTAGGAGTGAAATCCGTAGATATCTGGAGGAACATCGGAGGCGAAGGCGGCCGATTGGCAGAGCATTGACGCTGAGGTGCGAAAGCGTGGGGAGCAAACAGG
## 1991                                                         ACAGAGGGTGCGAGCGTTAATCGGAATTACTGGGCGTAAAGGGCGCGTAGGCGGTTTTTTAAGTTAGATGTGAAATCCCTGGGCTTAACCTAGGAATGGCATTTAAAACTGAAGAGCTAGAGTAGAGTAGAGGTAAGTAGAATTTCCGGTGTAGCGGTGAAATGCGTAGATATCGGAAAGAATACCAGTGGCGAAGGCGGCTTTCTGGACTCATACTGACGCTGAGGCGCGAAAGCATGGGTAGCAAACAGG
## 1996                                                         ACGAATGTGGCAAGCGTTGTTCGGAATCACTGGGCGTAAAGAATGCGTAGGCGGCTTGACAAGTCAGGAGTGAAATCCCCGGGCTCAACCCGGGAACTGCTTTTGAAACTGTCTTGCTAGAGTCCCGGAGGGGTCAATGGAATTCCGAGTGTAGAGGTGAAATTCGCAGATATTCGGAGGAACACCAGTGGCGAAGGCGATTGACTGGACGGGAACTGACGCTGAGGCATGAAAGTGTGGGGAGCAAACAGG
## 1998                                                        ACGTAGGGTGCAAGCGTTGCCCGGATTTACTGGGCGTAAAGCGTTTCGTAGGCGGTTTGTCGCATCTTCTTTCAAAGACCGAGGCTCAACCTCGGGAAGGGAGAAGAGATGGGCAAACTTGAAATATTTCGGGGCTGATGGAACTATCGGTGTAGGGGTGAAATCCGTTGATATCGATAGGAACTCCAAGGGCGAAGGCAGTCAGCTAGGAATTTTTTGACGCTGAGGAACGAAAGCTAGGGGAGCGAAAGGG
## 1999                                                         ACAGAGGGTGCAAACGTTGCTCGGATTTACTGGGCGTAAAGCGCATGTAGGCGGGACGCTAAGTCGGGTGTGAAATCCCTGGGCTCAACCTAGGAACTGCATCCGAAACTGGCGTCCTGGAGTAGTGGAGAGGAAAGCGGAATTCCGGGTGTAGAGGTGAAATTCGTAGATATCCGGAGGAACACCAGTGGCGAAGGCGGCTTTCTGGACACTAACTGACGCTGAGATGCGAAAGCGTGGGGAGCAAACAGG
## 2001                                                        ACGGAGGGTGCAAGCGTTAATCGGAATCACTGGGCGTAAAGCGCGCGTAGGCGGCTTTGTAAGTCAGAGGTGAAAGCCCACGGCTCAACCGTGGAACTGCCTTTGATACTGCAAGGCTACGAGTTTGTGAGAGGATAGTGGAATTCCTGGTGTAGGAGTGAAATCCGTAGAGATCAGGAGGAACATCAGTGGCGAGGGCGACTGTCTGGCACAACACTGACGCTGAGGTGCGAAAGCGTGGGTATCAAACAGG
## 2003                                                        ACAGGTGGTACAAGTAGTATCCACGAATATTGAGTCTAAAGCGCTCGTAGCAGGCTTGGACAGTTTTCTGTGAAATCTTCCGGCTCAACCGGAAGGCGCGCAGGGAATACTACCAGGCTTGGGAGTGGGGGAGGCCAAGGGTACTTGAGGGGGAGGAGTAAAATCCTGTAATCCTTCAAGGACCTTCAGTAGCGAAGGCGCTTGGCCAAAACACGTCCGACTGTGAGGAGCGAAGGCTAGGGGAGCAAATCGG
## 2009                                                         ACGAAGGGTGCAAGCGTTAATCGGAATTACTGGGCGTAAAGCGCGCGTAGGCGGTTTGGTTAGTTGGATGTGAAATCCCCGGGCTCAACCTGGGAATTGCATTTGTGACTGCACGGCTAGAGTGTGTCAGAGGGGGGTAGAATTCCACGTGTAGCAGTGAAATGCGTAGATATGTGGAGGAATACCGATGGCGAAGGCAGCCCCCTGGGATAACACTGACGCTCATGCACGAAAGCGTGGGGAGCAAACAGG
## 2010                                                         ACGAGAGGTGCAAACGTTACCCGGAATCATTGGGCGTAAAGCGTCCGCAGGCGGCGCAGTAAGTCTGGTATTAAATCTCGAGGCTCAACTTTGAGGCTGTACTAGATACTACTGTGCTAGGGACTTGAAGAGGTGAATGGAATTCTGCATGTAGGGGTAAAATCCGTAGATATGCAGAGGAACACCAAAAGCGAAGGCAGTTCACTGGTCAAGTTCTGACGCTCATGGACGAAAGCGTGGGTAGCAAACGGG
## 2011                                                         ACGAACCGGACGAACGTTATTCGGAATCACTGGGCTTAAAGAGTGCGTAGGCGGCCTTGTAGGTTGGGTGTGAAAGCCCACGGCTCAACCGTGGAACTGCGCTCAAAACCACAAGGCTTGAGGAAGGCAGAGGTGAGCGGAACTGATGGTGGAGCGGTGAAATGCGTTGATATCATCAGGAACACCAGTGGCGAAGGCGGCTCACTGGGCCTTTACTGACGCTGAGGCACGAAAGCTAGGGGAGCGAACGGG
## 2016                                                         CCCAGCGGTCCAAGTCGCAACCACATTTGTTGGGTCTAAAACATTCGTAGCTTGCTTAGTAAGTCGCTTGTGAAATCCCCTCTCATGAGGAGGGAACGGCACGCGATACTGCTGAGCTTGAGACCGGAAGACGCATTGAGTACCTACAGAGTACCGGTAAAATGGGTTGATCCTGTAGGGACTACCACCAGCGAAGGCACGATGCGAGTACGGATCTGACAGTGAGGAATGAAGGCTAGGGGCGCAAAGTGG
## 2017                                                        ACGGAGGGTGCAAGCGTTATCCGGAATCACTGGGTTTAAAGGGTGCGTAGGCGGTCTTGTAAGTCAGTCGTGAAATCGCTGGGCTCAACCCATGCAATTGCGATTGATACTGCGAGACTTGAATGGGGTTAAGGCAGGCGGAATGTGGCATGTAGCGGTGAAATGCATAGATATGCCATAGAACACCGATTGCGAAGGCAGCCTGCTGGGCCTTTATTGACGCTGAGGCACGAAAGCGTGGGGAGCAAACAGG
## 2018                                                        ACAGAGACCCCAAGCGTTATCCGGAATTATTGGGCGTAAAGGGTGAGTAGGTTGTTTTATTAGTCGGGTGTCAAAGCTCCAGGGCTCAACCTTGGAATCGCATCCGAAACGGTAAAACTAGAATATGTAAGAGGTTAGTGGAACTCTATGTGTAGGGGTAAAATCCGTTGATATATAGGGGAACACCTAAAGCGAAGGCAGCTAACTGGTACATTATTGACACTGAATCACGAAAGCGTGGGTAGCGAATGAG
## 2020                                                         ACAGAGACTGCAAGCGTTACTCGGATTCACTGGGCGTAAAGGGAGCGCAGGCGGGCGGGTGTGTCATATGTGAAATCCCGGGGCTCAACCCCGGGGCTGCATGTGAAACTACCTGTCTAGAGATTCGGAGGGGTAAGCGGAATTCCTGGTGGAGCAGTGAAATGCGTAGATATCAGGAGGAACACCAACGGCGAAGGCAGCTTACTGGACGAAATCTGACGCTCAGGCTCGAAAGCATGGGGAGCAAACAGG
## 2022                                                        ACCGGCGGCTCGAGTGGTGGCCACTATTACTGGGCTTAAAGGGTCCGTAGCTGGATATACAAGTCTCTTGAGAAATCCGCCGGCTTAACCGGTGGGCGTTCAGGGGAAACTGTATTTCTAGGGACCGGGAGAGGTGAGAGGTACTGCCGGGGTAGGAGTGAAATCCTGTAATCCCGGTGGGACCACCTATGGCGAAGGCATCTCACCAGAACGGGTCCGACAGTGAGGGACGAAAGCTGGGGGAGCAAACCGG
## 2024                                                          ACGTAGGTCTCGAGCGTTATCCGGATTTACTGGGCGTAAAGAGTCCGTAGCAGGATTTGTAAGTCTGTTTTCAAATATTCTGACTTAATCAGAAAAAGGGAGCAGATACTGCAGATCTGGAGGTTTTTAGAGGATAGTGGAATTTCCAGTGGAGAGGTGAAATTTGTTGATATTGGAAGGAACGCCGAAAGCGAAAGCAGCTATCTATAACAACCTGACTGTGAGGGACGAAAGCTTGGGTAGCAAACGGG
## 2026                                                         ACAGAGACCTCAAGCATTATCCGGATTTATTGGGCGTAAAGCGTCCGCAGGTTGTTTGGCAAGTCTGACGTCAAAACTCACCGCTCAACGGTGATCCGGCGTTTGATACTACCGAACTAGAGGGCGGGAGAGGTGAGCGGAACTCTCGGTGTAGTCGTAATAAGCGTTGATATCGAGAAGAACACCAAATGCGAAGGCAGCTCACTGGAACGCCTCTGACACTCATGGACGAAAGCGTGGGGAGCAAACAGG
## 2030                                                          ACGAAGGTCGCAAGCGTTATCCGGAGTTACTGGGCGTAAAGAGTCCGCATTTGGTGTAGTAAGTCTGATTTCAAAGACTCTGTCTCAAGCAGAGGAAGGGGTCGGATACTGCTACGCTGGAGATTACTAGGGGTTAGTGGAATTACCGGTGGAGGGGTGAAATCCGTTGATATCGGTAGGAACGCCAAAAGCGAAAGCAGCTAACTATAGTTTTCTGACAATCAAGGACGACAGCTAGGGGATCAAACGGG
## 2034                                                         ACGGAGGGTGCAAGCGTTATCCGGATTTATTGGGTTTAAAGGGTCCGTAGGCGGACTCGTAAGTCAGTGGTGAAATCTCATAGCTTAACTATGAAACTGCCATTGATACTGCGGGTCTTGAGTAAAGTAGAAGTGGCTGGAATAAGTAGTGTAGCGGTGAAATGCATAGATATTACTTAGAACACCAATTGCGAAGGCAGGTCACTATGTTTTAACTGACGCTGATGGACGAAAGCGTGGGGAGCGAACAGG
## 2036                                                         ACGTAGGGGGCAAGCGTTATCCGGAATCACTGGGCGTAAAGGGCGTCTAGGCGGCAAGGCAAGTCTCGCGTAAAAGCTCCCGGCTCAACTGGGAGAGCGCGCGGGAAACTGCCTGGCTAGAGACTCGGAGAGGGGTGTGGAATTCCCGGTGGAGTGGTGAAATGCGTTGAGATCGGGAGGAACACCAGTAGCGAAGGCGGCACCCTGGCCGAGTTCTGACGCTGAAGCGCGACAGCGTGGGGAGCAAACGGG
## 2037                                                        ACAGAAGGTGCAAGCGTTTATCGGAATGATTGGGCGTAAAGCGTCTGTAGGTGGTTCTGGAAGTCTACTGTCAAATATCAAGGCTCAACCTTGACCAGGCAGTGGAGTACTCCAGGACTAGAGTATGGTAGGGGTAGAGGGAATTCCTTGTGAAGCGGTGAAATGCATAGATATAAGGAAGAACACCAGTGGCGAAAGCGCTCTACTGGGCCAAAACTGACACTGAGAGACGAAAGCTAAGGGAGCGAATAGG
## 2038                                                         ACGGAGGGTGCAAGCGTTGTTCGGAATTACTGGGCGTAAAGCGTACGTAGGCGGTGGATTAAGTCGAATGTTAAAGCCTGGGGCTCAACCCCAGAGATGCATTCGATACTGGTTCACTAGAGGGTGATAGAGGAAAGCGGAATTCCGAGTGTAGCGGTGAAATGCGTAGATATTCGGAGGAACATCAGTAGCGAAGGCGGCTTTCTGGGTCACACCTGACGCTGAGGTACGAAAGCGTGGGGAGCAAACGGG
## 2039                                                          ACGAGAGGGACGAGTGTTACACCGTCAAACTGGGCGTAAAGCGTTCTGTCGCGGACGAGTGCCTGCCTTCCTGTTTTTCCAGACCGTACGGCTGGATCAAGGAGGGAACGGCTCGTCCAGAGTGACAAGGACGGGGGATGGATTCGCAGACTAGGGTTTGCCTCCGTAGACGCTGCGACGACAACCGCGCGCGCAAGCCTCCTTCGACTTGTTACTGACGCCGAGGAACGAAAGCGTGGGGAGCCAACGGG
## 2045                                                         ACGAAGGGTGCAAGCGTTATTCGGAATAATTGGGCGTAAAGGGTTCGTAGGTTGGATTGTAAGTCAAGTGTGAAATCCCCAGGCTTAACCTGGGACGTGCATTTGAGACTGTGTTTCTTGAGTTTCGGAGAGGGTGGTGGAATTGCTGGTGTAGGAGTGACATCCGTAGAGATCAGCAGGAACACCGGAGGCGAAGGCGACCACCTGGCCGGATACTGACGCTGAGGAACGAAAGCGTGGGGAGCAAACAGG
## 2052                                                         ACGAAGGGGGCTAGCGTTGTTCGGAATAACTGGGCGTAAAGGGCGCGTAGGCGGTTAGATACGTCTGATGTGAAAGTCCTGGGCTTAACCTAGGGACTGCATTGGATACGGTTTAGCTAGAGTGCGAGAGGGGAAAATGGAATTGCGCGTGTAGAGGTGAAATTCGTAGAGATGCGCAAGAACACCAGTGGCGAAGGCGATTTTCTGGCTCGCAACTGACGCTGAGGCGCGAAAGCGTGGGGAGCAAACAGG
## 2053                                                         ACGGAAGGTCCGGGCGTTATCCGGATTTATTGGGTTTAAAGGGAGCGTAGGCCGGTTTGTAAGCCAGCAGTGAAATGTAGATGCTCAACATCTGAGTTGCTGTTGGAACTGCGGACCTTGAGTGCGCGCAAGGGCGGCGGAATTTGTGGTGTAGCGGTGAAATGCTTAGATATCACGAAGAACCCCGATTGCGAAGGCAGCTGCCCGGAGCGCAACTGACGCTGAAGCTCGAAGGTGCGGGTATCGAACAGG
## 2056                                                        ACGTAGGTGGCAAGCGTTACTCGGAATTACTAGGCGTAAAACGCAGGTAGGTGGTTGGGTAAGTCTGTAATGAAATCTCCCGGCTTAACTGGGAGGGGTTTATAGATACTGCCTGACTTGGAGTGTGGGAGAGGTAACTGGAATTCCTGGTGTAGCGGTGAAATGCGTAGATATGCGGAGGAACACCGATGGCGAAGGCAATCCCCTGGACCTGTACTGACGCTCATGCACGAAAGCGTGGGGAGCAAACAGG
## 2058                                                        TACGTAGGGTGCAAGCGTTGTTCGGAATCACTGGGCGTAAAGGGCGCGTAGGTTGTTTGGTTTGTCTGTTGTGAAATCCCATGGCTCAACCATGGAACGGCAATGGAAACTGCCAAACTAGAGTACAAGAGGGGATGGCGGAATTCCCGGTGTAGAAGTGAAATTCGTAGATACCGGGAGGAACACCTGTGGCGAAAGCGGCCATCTGGATTGTAACTGACACTGAGGCGCGAAAGCATGGGGAGCAAACAGG
## 2060                                                                                                              ACAGATGGAGCAAGTATCTTGTAGTATGATTAGGTTTACAGCGTGTGAATACGATTATAATGGTTATATTTATTTAAAGAGATAGAATTCCTTATTTAGTGGTTAAATATGTAAAAATGAGTGAGGAATGCCAACAGCGAAAGCAATCTCTTAAAAACCATTAAAGTTTAAACACGGAAGTCCGAGTATCGAAACGGAT
## 2061                                                          ACGTAGGGGACGAGCGTTGTCCGGATTTACTGGGCGTAAAGGGCGCGTAGGCGGTCATTTAAGTCAGCTGTGAAATGGTTTGGCTTAACCGGATCATGCGGTTGAAACTGGATGACTGGAGTGCTGGAGAGGCAAGTGGAATTCCCAGTGTAGCGGTGAAATGCGTAGATATTGGGAAGAACACCGGTGGCGAAGGCGACTTGCTGGCCAGCAACTGACGCTGAGGCGCGAAAGCCAGGGGAGCAAACGGG
## 2063                                                         ACAGAGGATGCAAGCGTTATCCGGAATGATTGGGCGTAAAGCGTCTGTAGGTGGCCTGACAAGTCTTCTGTCAAAGATCAGGGCTTAACCCTGGACAGGCAGGAGAAACTATCAGGCTAGAGTTCGGTAGAGGCAGAGGGAATTCCCAGTGAAACGGTGAAATGTGTAGATATTGGGAGGAACACCAACGGCGAAAGCACTCTGCTGGGCCGAGACTGACACTGAGAGACGAAAGCGAGGGGAGCAAATGGG
## 2065 ACAGATAGGGCAGCGTTATGAGAAATAAACAGGTCTTAAGTGTACTAAGATGGTTAAACCCTATTAAGGTAATGACTAGAATAATGTATAAGTAATAAGAATTTAAAGAGTAAGGATAGAATCTGTCGATACTTTAAGGAAAACCAAATTTTATTTAATATTAAACATTTTATGTTTTAATTAGTTAACTCGAGCACTATCTCAAGTTAATTAAGTATATTAAGATAATAATGATTTAAATATATAAAACACTATAAATTACTGTACATTATTGACATTGAAGTACGAAAGTTTAGGTATCAAATAGG
## 2066                                                         ACGTAGGGGGCAAGCGTTATCCGGATTTACTGGGTGTAAAGGGAGCGCAGACGGCGATGCAAGTCTGGAGTGAAAGCCCGGGGCTCAACCCCGGGACTGCTTTGGAAACTGTATGGCTAGAGTGCTGGAGAGGCAAGCGGAATTCCTAGTGTAGCGGTGAAATGCGTAGATATTAGGAAGAACACCAGTGGCGAAGGCGGCTTGCTGGACAGTAACTGACGTTCAGGCTCGAAAGCGTGGGGAGCAAACAGG
## 2067                                                         ACGGAGGGAGCTAGCGTTGTTCGGAATTACTGGGCGTAAAGCGCACGTAGGCGGTTTTTCAAGTTAGGGGTGAAAGCCCGGAGCTCAACTCCGGAATTGCCCTTGAAACTGGATGACTGGAATCTTGGAGAGGTCAGTGGAATTCCGAGTGTAGAGGTGAAATTCGTAGATATTCGGAAGAACACCAGTGGCGAAGGCGACTGACTGGACAAGTATTGACGCTGAGGTGCGAAAGCGTGGGGAGCAAACAGG
## 2069                                                         ACGGGGGGTGCAAGCGTTATTCGGAATTATTGGGCGTAAAGAGCGCGTAGGCGGTCTCTTAAGTCAGATGTGAAAGCCCTGGGCTTAACCCTGGAAGTGCATTTGAAACTAAGGGACTTGAGTATGGGAGAGGGAAGTGGAATTCCTGGTGTAGAGGTGAAATCCGTAGATATCAGGAGGAACACCGGTGGCGAAGGCGACTTCCTGGACCAATACTGACGCTGAGGCGCGAAGGCGTGGGGAGCAAACAGG
## 2076                                                         ACAGAGGAGGCAAGCGTTGTTCGGAATCATTGGGCTTAAAGGGCGTGTAGGCGGCGGCGCAAGTGTCGAGTGAAATCCCACTGCTCAACGGTGGAACCGCTCGGCAAACTGCGTCGCTTGAGGTCGGTAGAGGTGCTCGGAACTCTAGGTGGAGCGGTGAAATGCGTAGATATCTAGAGGAACGCCTGAGGCGAAAGCGGGGCACTGGGCCGAATCTGACGCTGAGACGCGAAAGCGTGGGGATCAAACAGG
## 2078                                                         ACAGAGGCCCCGAGCGTTATCCGGAATTATTGGGCGTAAAGGGTGCGTAGGTGGCTATATTAGTCTTTCGTTAAATCCATGGGCCTAACCTGTGGTCTGCGAAGGAAACGGTATAGCTTGAGAGCGAGAGAGGTGAATGGAACTAATAGTGTAGGGGTGAAATCCGTTGATATTATTGGGAACACCAAAAGCGAAGGCAATTCACTGGCTCGTTTCTGACACTGATGCACGAAAGCGTGGGTAGCGAATGGG
## 2083                                                         ACGGAGGGCGCAAGCATTAATCAGATTTACTGGGTGTAAAGGGTGCGTAGGCCGTTAGGCAAGTCAGACGTGAAATTCCGGTACTCAATACCGGAGCTGCGTTTGAAACTACCTTTCTGGAGGATAGGCGGAGAAAACGGAATTCCACATGTAGCGGTGAAATGCGTAGATATGTGGAAGAACACCTGTGGCGAAAGCGGTTTTCTAGCTTATTTCTGACGCTGAGGCACGAGAGCGTGGGGAGCAAACAGG
## 2086                                                         ACGTAGGGTGCAAGCGTTAATCGGAATTACTGGGCGTAAAGCGTGCGCAGGCGGTTATATAAGACAGATGTGAAATCCCCGGGCTCAACCTGGGAACTGCATTTGTGACTGTATAGCTGGAGTGCGGCAGAGGGGGATGGAATTCCGCGTGTAGCAGTGAAATGCGTAGATATGCGGAGGAACACCGATGGCGAAGGCAATCCCCTGGACCTGTACTGACGCTCATGCACGAAAGCGTGGGGAGCAAACAGG
## 2089                                                         ACGAGGGGAACGAGCGTTATACAGTCAAACTGGGCGTAAAGGGTTCTACGGCGGAAGGTTGATAGGAAACGATGAAAGTTTCGAGAGCAATCCGAAATTATCTTTTCGACGAATTTTCTTGAATGAAATAGGGGTATGTGTTATTCCATCCAGAGAGTTAAAATGCGTAGACGGATGGCCGAGGGCCATGGGCGAAGGCCGCATACCAGGTTTCATTGACGCTTAGGAACAAAAGCATGGGGAGCGATCGGG
## 2092                                                         ACGGAGGGTGCAAGCGTTGTCCGGATTTATTGGGTTTAAAGGGTGCGCAGGTGGTTTATTAAGTCAGTGGTGAAAGACGGTCGCTCAACGATCAAACGGCCATTGATACTGATAGACTTGAATTACTGGGAAGTAACTAGAATATGTAGTGTAGCGGTGAAATGCTTAGATATTACATGGAATACCAATTGCGAAGGCAGGTTACTACCAGTGGATTGACGCTGATGGACGAAAGCGTGGGGAGCGAACAGG
## 2093                                                         ACGTAGGGTGCGAGCGTTAATCGGAATTACTGGGCGTAAAGCGTGCGCAGGCGGTTATATAAGACAGATGTGAAATCCCCGGGCTTAACCTGGGACCTGCATTTGTGACTGTATAGCTAGAGTACGGTAGAGGGGGATGGAATTCCGCGTGTAGCAGTGAAATGCGTAGATATGCGGAGGAACACCGATGGCGAAGGCAATCCCCTGGGCCTGCACTGACGCTCATGCACGAAAGCGTGGGGAGCAAACAGG
## 2096                                                         ACGTAGGTGGCGAACGTTATCCGGAATTATTGGGCGTAAAGCGTTCTTAGGTGGTTGTATAAGTTGGATGCGAAATCTACAGGCCCAACCTGTAAATTGTATCCGAAACTGTACGGCTGGAGCTTGTTAGAGGCCAGTGGAATTTCTGGAGTAGGGGTAAAATCCGTTGATACCAGAAGGAACACCAATTGCGAAGGCAGCTGGCTGGGGCATTGCTGACACTTATGAACGAAAGCGTGGGGAGCGAAACGG
## 2107                                                         ACAGAGGGTGCAAGCGTTGTTCGGAATCACTGGGCGTAAAGCGCGTGTAGGCTGCTGATTGTGTCGAGTGTGAAAGCCCTCGGCTTAACCGAGGAAGTGCACTCGAAACTGGTCAGCTTGAGTACCGGAGAGGAAGGTGGAATTCCCGGTGTAGAGGTGAAATTCGTAGATATCGGGAGGAACACCAGTGGCGAAAGCGACCTTCTGGACGGATACTGACGCTGAGACGCGAAAGCGTGGGGAGCAAACAGG
## 2108                                                         ACGGAGGATCCAAGCGTTATCCGGATTTATTGGGTTTAAAGGGAGCGTAGGTTGACATATAAGTCAGCTGTGAAAGTTTACGGCTCAACCGTGAAATTGCAGTTGATACTGTATGTCTTGAGTGTACAAGAGGTGGGCGGAATTCGTGGTGTAGCGGTGAAATGCTTAGATATCACGAAGAACTCCAATTGCGAAGGCAGGACTCTGGGCCATTACTGACACTGAGGAGCGAAAGCGTGGGGAGCGAACAGG
## 2112                                                         ACAGAGACTACAAGCGTTACTCGGATTCACTGGGCGTAAAGGGTGCGTAGGCGGTTGTGTGTGTTAAATGTGAAATCCTGGGGCTCAACTCCAGAACTGCATTTAAAACTACACATCTCGAGTTTTAGAGAGGTAAGCGGAATTCTTAGTGGAGCAGTGAAATGCATAGATATTAAGAGGAACACCAATTGCGAAGGCAGCTTACTGGATAAAAACTGACGCTGAGGCACGAAAGCGTGGGGAGCAAAAGGG
## 2113                                                         ACGGAGGGTGCAAGCGTTATCCGGATTTACTGGGTTTAAAGGGTGCGTAGGCGGACAGGTAAGTCAGTGGTGAAATCCCCGAGCTTAACTCGGGAACTGCCGTTGATACTATCTGTCTTGAATATCGTGGAGGTAAGCGGAATATGTCATGTAGCGGTGAAATGCTTAGATATGACATAGAACACCAATTGCGAAGGCAGCTTGCTACACGATTATTGACGCTGAGGCACGAAAGCGTGGGGATCAAACAGG
## 2114                                                         ACGTAGGGGGCGAGCGTTGTTCGGAATTACTGGGCGTAAAGGGCATGTAGGCGGCCTGATAAGCCTGGCGTGAAAGTCCACGGCTCAACCGTGGGATTGCGTCGGGAACTGTCGGGCTTGAGTCACGGAGGGGAAGGTAGAATTCCTGGTGTAGGGGTGAAATCTGTAGAGATCAGGAAGAATACCAGTGGCGAAGGCGACCTTCTGGCCGATGACTGACGCTGAGGTGCGAAAGTGTGGGGATCAAACAGG
## 2117                                                         ACGAACCGTGCGAACGTTATTCGGAATCACTGGGCTTAAAGGGTGCGTAGGCGGGTATATAAGTCGAGGGTGAAAGCCTCTAGCTCAACTAGAGAAGTGCCCTCGAAACTATATGTCTTGAGAGAGATAGGGGCATACGGAACTTCCGGTGGAGCGGTGAAATGCGTAGATATCGGAAGGAACGCCGGTGGCGAAAGCGGTATGCTGGATCTTTTCTGACGCTGAGGCACGAAAGCTAGGGTAGCGAACGGG
## 2122                                                         ACGAGTGCCTCAAGCGTTATCCGGAATCATTGGGCGTAAAGGGTGTGTAGGTGGTTGTGTTAGTCTTGCGTAAAATGCTCCGGCTCAACCGGAGTTCCGCGCAGGAAACGGCACGACTAGAGGACGGTAGAGGTTTCTGGAACTCATAGTGTAGCGGTGAAATGCGTTGATATTATGGGGAACACCAAAAGCGAAGGCAAGAAACTGGACCGCTCCTGACACTGAAACACGAAAGCGTGGGTCGCGAATGGG
## 2126                                                         ACAGAGGGTGCAAACGTTGCTCGGAATTATTGGGCGTAAAGCGCACGTAGGCGGCATCGTAAGTCCGATGTGAAAGCCCCAGGCTCAACCTGGGAAGTGCATTGGATACTGCGGAGCTTGAATCTGGAAGAGGGTCGCGGAATTCCCGGTGTAGAGGTGAAATTCGTAGATATCGGGAGGAACACCAGTGGCGAAGGCGGCGACCTGGGCCAAGATTGACGCTGAGGTGCGAAAGCGTGGGGAGCAAACAGG
## 2131                                                         ACAGAGGGTACAAACGTTGTTCGGAATCACTGGGCTTAAAGCGCGCGTAGGTGGTTCGGTAAGTGCGATGTGAAAGCCCCCGGCCTAACCGGGGAACTGCATCGCAGACTGCCGAGCTTGAGAGCGGTAGGGGAGACTAGAACTCCTGGTGGAGCGGTGAAATGCGTAGATATCAGGAGGAATACCGGTGGCGAAGGCGAGTCTCTGGTCCGCTTCTGACACTTAGGCGCGAAAGCTAGGGGAGCAAACGGG
## 2133                                                         ACAGAGACTGCAAGCGTTACTCGGAATCACTGGGCGTAAAGGGAGCGCAGGCGGAAAGATGTGTCAGGTGTGAAACCCCGGAGCTCAACTGCGGGACTGCGCCTGAAACTGTCTTTCTAGGGACTCGGAGGGGCTGGTGGAATTCCCGGTGGAGCAGTGAAATGCGTAGATATCGGGAGGAACACCGACGGCGAAGGCAGCCAGCTGGACGAGATCCGACGCTCAGGCTCGAAAGTATGGGGAGCAAAAAGG
## 2135                                                         ACGAAGGGTGCAAGCGTTGTTCGGAATTACTGGGCGTAAAGCGCGCGTAGGCGGATCTGTAAGTTGATTGTGAAATCCCTGGGCTTAACCTGGGACGTGCAGTCAAAACTGCAGATCTTGAATGTCTGAGAGGATGGCGGAATTCCCGGTGTAGTAGTGAAATACGTAGATATCGGGAAGAACACCTGAGGCGAAGGCGGCTATCTGGCGGAACATTGACGCTAAGGCGCGAAAGCGTGGGGAGCAAACAGG
## 2139                                                         ACAGAGGGTGCTAGCGTTGTTCGGATTGACTGGGCGTAAAGGGCGCGTAGGCGGTTATATTAGTCAGGCGTGAAAGCCCCGGGCTTAACCCGGGAATTGCGTTTGAAACGGTATAACTAGAGACCGGAAGAGGAAAGCAGAATTCCTAGTGTAGAGGTGAAATTCGTAGATATTAGGAGGAATACCAGAGGCGAAGGCGGCTTTCTGGTCCGGTTCTGACGCTGAGGCGCGAAAGCGTGGGGAGCAAACAGG
## 2140                                                         ACGGAGGGTGCAAGCGTTGTTCGGAATTATTGGGCGTAAAGGGCAGGTAGGTGGTTTAGGAAGTTCACTGTGAAAGCCCCGGGCTTAACCTGGGAAGTGCAGTGAATACTCCATGACTAGAGTGCCGGAGGGGTACGTGGAATTCCCGGTGTAGCGGTGAAATGCGTAGATATCGGGAGGAACACCAGAGGCGAAGGCGGCGTACTGGACGGCAACTGACACTCAACTGCGAAAGCGTGGGTAGCAAACAGG
## 2143                                                         ACGAAGGGTGCAAGCGTTGTTCGGATTTATTGGGCGTAAAGCGCGCGCAGGCGGATCTAAAAGTCAGGTGTGAAATCTCGGAGCTTAACTCCGAAAGTGCGCCTGAAACTCTAGATCTAGAATATCGGAGAGGGAAGGGGAATTTCTCATGTAGGGGTAAAATCCGTAGATATGAGAAGGAACACCAGAGGCGAAGGCGCCTTCCTGGACGATGATTGACGCTGAGGCGCGAAAGCGTGGGGAGCAAACAGG
## 2144                                                         ACGTAGGGTGCAAGCGTTAATCGGAATTACTGGGCGTAAAGCGTGCGCAGGCGGTTATGTAAGACAGAGGTGAAATCCCCGGGCTCAACCTGGGAACTGCCTTTGTGACTGCATAGCTAGAGTACGGTAGAGGGGGATGGAATTCCGCGTGTAGCAGTGAAATGCGTAGATATGCGGAGGAACACCGATGGCGAAGGCAGCTTACTACGCATATATTGACGCTGAGGCACGAAAGCGTGGGGATCAAACAGG
## 2147                                                         ACGTAGGGGGCAAGCGTTGTTCGGAATTACTGGGCGTAAAGGGCGCGTAGGCGGCATGGCAAGTCGGGAGTGAAAGCCCGGGGCTCAACCCCGGAACTGCTTTCGAAACTGCCAAGCTAGAGTACGGAAGGGGCAACTGGAATTCCCAGTGTAGCGGTGAAATGCGTAGATATTGGGAAGAACACCGGAGGCGAAGGCGGGTTGCTGGGCCGATACTGACGCTGAGGCGCGAAAGCCAGGGGAGCAAACGGG
## 2150                                                         ACGAAGGGGGCGAGCGTTGTTCGGAATCACTGGGCGTAAAGCGTGCGTAGGCTGTTTTGCAAGTTGGAAGTGAAAGCCCAGGGCTCAACCCTGGAATTGCTTTCAAAACTGCAAGACTCGAATCTCGGAGAGGATAGCGGAATTCCCAGTGTAGTAGTGAAATACGTAGATATTGGGAGGAACACCGGTGGCGTAAGCGGCTATCTGGACGAGTATTGACGCTGAGGCACGAAAGCGTGGGGATCAAACAGG
## 2152                                                         ACGAGGGGAACGAGCGTTATACAGTCAAACTGGGCGTAAAGGGTCCTGCGGCGGGAGATAGTTCGTAGGAGATCAAAGTTTTAAGAGTAATCTGAAAGTTTCTCCTATACGTATCTTCTGGAATGAAATAGGGGTATGTGTTAGTCCATCCAGAGAGTTAAAATGCGTAGACGGATGGCCGAGGACCAATGGCGAAGGCCACATACTAGTTTTCATTGACGCTTAGGGACAAAAGCATGGGGAGCGATCGGG
## 2154                                                         ACGTGAGTTCCAAGCGTTACCCGGAATTATTGGGTGTAAAGGGTGCGTAGGTGGATTTTTAAGTCAGGTATGAAAGACCGGAGCTCAACTCCGTGTTTGTGCTTGAAACTGGAAATCTGGAATCAGGGAGAGGTTAGCGGAATTCTAAGTGTAGGGGTGCAATCCGTAGATACTTAGAAGAACACCAAAAGCGAAGGCAGCTAACTGGAACTGTATTGACACTGAGGCACGAAAGCGTGGGGAGCAAAAAGG
## 2155                                                         ACGTAGGGTGCAAGCGTTAATCGGAATTACTGGGCGTAAAGCGTGCGCAGGCGGTTTCGTAAGACAGAGGTGAAATCCCCGGGCTCAACCTGGGAACTGCCTTTGTGACTGCGAGGCTAGAGTATGGCAGAGGGGGGTGGAATTCCACGTGTAGCAGTGAAATGCGTAGAGATGTGGAGGAACACCGATGGCGAAGGCAGTCCCCTGGGATAAGACTGACGCTCATGCACGAAAGCGTGGGGAGCAAACAGG
## 2156                                                         ACGAACCGTCCAAACGTTATTCGGTATCACTGGGCTTAAAGAGTTCGTAGGCGGCTAAACAGGTGAGGTGTGAAAGCCCTCGGCTCAACCGAGGAATTGCGCTTCAAACCGTTTAGCTAGAGGAAGACAGAGGTGAGCGGAACTGATGGTGGAGCGGTGAAATGCGTTGATATCATCAGGAACACCGGTGGCGAAAGCGGCTCACTGGGTCTTTTCTGACGCTGAGGAACGAAAGCTAGGGGAGCGAACGGG
## 2159                                                         ACGTAGGGTGCGAGCGTTGTCCGGAATTACTGGGCGTAAAGAGCTCGTAGGCGGTTTGTCGCGTCGTCTGTGAAATTCTGCAACTCAATTGTAGGCGTGCAGGCGATACGGGCAGACTTGAGTACTACAGGGGAGACTGGAATTCCTGGTGTAGCGGTGAAATGCGCAGATATCAGGAGGAACACCGGTGGCGAAGGCGGGTCTCTGGGTAGTAACTGACGCTGAGGAGCGAAAGCGTGGGTAGCGAACAGG
## 2161                                                        ACAGAGGGTGCGAGCGTTATCCGGAATCATTGGGCGTAAAGCGTATGTAGGCGGATTTGTAAGTAAGGGATAAAATCTCTTCGCTCAACGAAGAATTTGTTCTTTAAACTGCAAGTCTTCGAAGCATGCAGAGGCAGGTGGAATTCTGAGTGTAGGGGTAAAATCCGTAGATACTCGGAGGAACGCCAAAAGCGAAAGCAGCCTGCTGGGCATGTCTTGACGCTGAGATACGAAAGCGTGGGGAGCAAACAGG
## 2164                                                         ACGGGAGATGCAAGCGTTATCCGGAATCACTGGGCGTAAAGCGTCTGTAGGTGGTTTGTTAAGTCTGTTGTTAAAAACTGAAGCTTAACTTCAGGAGTGCGGCAGAAACTGGCAAACTTGAGTATGGTAGGGGTAGGAGGAATTTCTAGTGTAGCGGTGAAATGCGTAGATATTAGGAAGAACACCATTAGCGAAGGCGTCCTACTGGGCCATAACTGACACTCAGAGACGAAAGCTAGGGGAGCAAACGGG
## 2173                                                         ACGGAGGACCCGAGCGTTATCCGGATTCATTGGGTTTAAAGGGTGCGTAGGCGGATTTGTAAGTCAGTGGTGAAATCTCCCGGCTCAACTGGGAAACTGCCATTGATACTGTGAATCTTGAATACAATTGATGTAGGCGGAATATGACATGTAGTGGTGAAATACTTAGATATGTCATAGAACACCGATTGCGAAGGCAGCTTACAAAGTTGTTATTGACGCTGAGGCACGAAAGCGTGGGGAGCAAACAGG
## 2174                                                         ACGTAGGGTGCAAGCGTTAATCGGAATTACTGGGCGTAAAGCGTGCGCAGGCGGTTATATAAGACAGATGTGAAATCCCCGGGCTCAACCTGGGAACTGCATTTGTGACTGTATAGCTAGAGTACGGTAGAGGGGGATGGAATTCCGCGTGTAGCAGTGAAATGCGTAGATATGCGGAGGAACACCGATGGCGAAGGCAGCCTCCTGGGCCAGTACTGACGCTCATGCACGAAAGCGTGGGGAGCAAACAGG
## 2175                                                         ACGGAGGATCCAAGCGTTATCCGGATTTATTGGGTTTAAAGGGTGCGTAGGCGGTTTGGTAAGTTAGAGGTGAAATCTCTGTGCTCAACACAGAAATTGCCTCTAATACTGTTGAACTAGAGAGTGGTTGCTGTAGGCGGAATGTGTGGTGTAGCGGTGAAATGCTTAGAGATCACACAGAACACCGATTGCGAAGGCAGCTTACAAAGCCACATCTGACGTTGAGGCACGAAAGCGTGGGGAGCAAACAGG
## 2177                                                         ACGGAGGATCCGAGCGTTATCCGGATTTATTGGGTTTAAAGGGTGCGTAGGCGGGCTTATAAGTCAGCGGTGAAAGTTTGCGGCTCAACCGTAAAATTGCCGTTGATACTGTGAGTCTTGAGTGTAGATGAGGTGGGTGGAATTCGTGGTGTAGCGGTGAAATGCATAGATATCACGAAGAACTCCAATTGCGTAGGCAGCTCACTGGGCTACAACTGACGCTGAGGCACGAAGGTGCGGGGATCAAACAGG
## 2179                                                         ACGAAGGGGGCAAGCGTTGTTCGGAATTACTGGGCGTAAAGGGAGTGTAGGCGGTTTAATAAGGTAGCGGTGAAATACCTGGGCTCAACCTAGGAACTGCCGTTATGACTATTAAACTAGAGTAATAGAGAGGATAGTGGAATACCCAGTGTAGAGGTGAAATTCGTAGATATTGGGTAGAACACCGGTGGCGAAGGCGACTATCTGGCTATTAACTGACGCTGAGGCTCGAAAGCATGGGGATCAAACAGG
## 2180                                                         ACGTAGGGGGCGAGCGTTGTCCGGAATCATTGGGCGTAAAGCGCGTGTAGGCGGTTTCGTAAGTCTGCTGTGAAAGTCGGAGGCTCAACCTTCGAATGCCAGTGGATACTGCGAGACTTGAGTCCGGAAGAGGAGAGTGGAATTCCTGGTGTAGCGGTGAAATGCGCAGATATCAGGAGGAACACCAACGGCGAAGGCAGCTCTCTGGGACGGTACTGACGCTGAGACGCGAAAGCGTGGGGAGCAAACAGG
## 2186                                                         ACGTAGGGTGCGAGCGTTAATCGGAATTACTGGGCGTAAAGCGTGCGCAGGCGGTTATGTAAGACAGATGTGAAATCCCCGGGCTCAACCTGGGACCTGCATTTGTGACTGCATAGCTAGAGTACGGTAGAGGGGGATGGAATTCCGCGTGTAGCAGTGAAATGCGTAGATATGCGGAGGAACACCGATGGCGAAGGCAATCCCCTGGGCCTGTACTGACGCTCATGCACGAAAGCGTGGGGAGCAAACAGG
## 2188                                                         ACGAAGGACCCAAGCGTTATCCGGATTTATTGGGTTTAAAGGGTGCGTAGGTGGACTTATAAGTCAGTGGTGAAAGCCTGTTGCTTAACAACAGAACTGCCATTGATACTGTAGGTCTTGAGTACAGTTGAGGTGGGCGGAATGTGTCAAGTAGCGGTGAAATGCTTAGATATGACACAGAACACCAATTGCGAAGGCAGCTCGCTAAGCTGTAACTGACACTGAGGCACGAAAGCGTGGGTAGCAAACAGG
## 2190                                                         ACGAGTGCCTCGAGCGTTATCCGGAATCATTGGGCGTAAAGGGTGTGCGGGCGGTCTTGTTAGTCTTCTGTTAAATTTCTCGGCTTAACCGGGAACCTGCAGGGGAAACGGCAAGACTTGAGGATGCGAGAGGTTAACGGAACTCATAGTGTAGCGGTGAAATGCGTTGATATTATGGGGAACACCAAATGCGAAGGCAGTTAACTGGAGCACTCCTGACGCTGAAACACGAAAGCGTGGGTAGCGAATGGG
## 2191                                                         ACAGAGGGTGCGAGCGTTAATCGGAATTACTGGGCGTAAAGCGCATGCAGATGGTTTGACAAGTCGGATGTGAAAGCCCCGGGCTTAACCTGGGAACGGCACTCGAAACTGTCAGACTAGAATGTGGAAGAGGGAAGTGGAATTCCGGGTGTAGCAGTGAAATGCGTAGATATCCGGAGGAACATCAGTGGCGAAGGCGGCTTACTGGGCTTTTACTGACGCTGAGGCTCGAAAGCGTGGGGAGCAAACAGG
## 2193                                                         ACGGGAGTGGCAAGCGTTATCCGGAATTATTGGGCGTAAAGCGTCCGCAGGCGGTTTTACAAGTCTGTCGTTAAAGCGTGGAGCTTAACTCCATTTCAGCGATGGAAACTGTAAGACTAGAGTGTGGTAGGGGCAGAGGGAATTCCCGGTGTAGCGGTGAAATGCGTAGATATCGGGAAGAACACCAGTGGCGAAGGCGCTCTGCTGGGCCATAACTGACGCTCATGGACGAAAGCGTGGGGAGCAAACAGG
## 2199                                                         ACGGAGGGTGCAAGCGTTATCCGGATTCACTGGGTTTAAAGGGAGCGCAGGCGGGCATGTAAGTCCGTGGTGAAATACCTCAGCTTAACTGAGGAACTGCCATGGATACTATATGTCTTGAATGTTGTGGAGGTTTGCGGAATAGGTCATGTAGCGGTGAAATGCATAGATATGACCTAGAACACCAATTGCGAAGGCAGCAGGCTACACAAATATTGACGCTGAGGCTCGAAAGCGTGGGGATCAAACAGG
## 2203                                                         ACGGAGGATGCGAGCGTTATCCGGATTTATTGGGTTTAAAGGGTGCGTAGGTGGCTTTTTAAGTCAGCGGTGAAAACTGGCGGCTCAACCGCAAGAGTGCCGTTGAAACTGGGGAGCTAGAGTATGAACGAGGTAGGCGGAATGTGTGGTGTAGCGGTGAAATGCATAGATATCACGCAGAACTCCGATAGCGAAGGCAGCTTACCAGGCCATAACAGACACTGAAGCACGAAAGCGTGGGGATCGAACAGG
## 2205                                                         ACAGAGACCTCAAGTGTTATCCGGATTTATTGGGCGTAAAGCGTCCGCAGATGGTTTGGCGGGTAAGGGGTTAAAACCTAGTGCTTAACATTAGGATGGCCTTTTAAACTACCAAACTAGAGGATGGAAGAGGCAAGCGGAATTGCAGGTGTAGTCGTAATAAGCGTTGATATCTGCAAGAACACCAAAGGCGAAGGCAGCTTGCTAGTACATTTCTGACATTCATGGACGAAAGCGTGGGGAGCAAACAGG
## 2206                                                          ACGTAGGGGGCAAGCGTTGTCCGGAATTATTGGGCGTAAAGAGCGCGTAGGCGGTAAATTAAGTTGGGTGTAAAAGGCATTGGCTCAACCAATGTAAGCATCCAAAACTGGTTTACTTGAGTGCAGGAGAGGAGAGTGGAATTCCCGGTGTAGCGGTGGAATGCGTAGATATCGGGAGGAACACCAGTGGCGAAGGCGACTCTCTGGACTGTAACTGACGCTGAGGCGCGAAAGCGTGGGGAGCAAACAGG
## 2207                                                         ACAGAGACCTCAAGCGTTATCCGGATTTATTGGGCGTAAAGCGTAGGTAGGTTGTCTAGTTAGTCGGATGTCAAAGCTCGGGGCTCAACCCCGAAAAGGCATTCGAAACGGCTAGGCTAGAGGATGTGAGAGATCAGTGGAATTGATGGTGTAGTAGTGAAATGCGTTGATATCATCAAGAACACCCAAGGCGAAGGCAACTGATTGGCACATATCTGACACTGAGCTACGAAAGCGTGGGGAGCGAATGGG
## 2208                                                         ACGGAGGATCCAAGCGTTGTCCGGATTTATTGGGTTTAAAGGGTGCGTAGGCGGATCTTTAAGTCAGTGGTGAAAGCCTGCAGCTCAACTGTAGAATTGCCATTGAAACTGAAGATCTTGAATTAGGTTAAAGTAGGCGGAATGTATCATGTAGCGGTGAAATGCTTAGATATGATACAGAACACCGATAGCGAAGGCAGCTTGCTGAACCTACATTGACGCTGAGGCACGAAAGCGTGGGGAGCGAACAGG
## 2210                                                         ACGGAGGATGCGAGCGTTATCCGGATTTATTGGGTTTAAAGGGTGCGTAGGCGGCACGCCAAGTCAGCGGTGAAATTTTCGGGCTCAACCCGGACTGTGCCGTTGAAACTGGCGAGCTAGAGTGCACAAGAGGCAGGCGGAATGCGTGGTGTAGCGGTGAAATGCATAGATATCACGCAGAACCCCGATTGCGAAGGCAGCCTGCTAGGGTGAAACAGACGCTGAGGCACGAAAGCGTGGGTATCGAACAGG
## 2216                                                         ACAGGGGGCCCGAGCGTTACGCGGAATCACTGGGCTTAAAGGGTGCGTAGGCGGTTCGGAAAGCATTACGTGAAAGCCCCCGGCTCAACCGGGGAACTGCGCGATGTACTACCGAACTTGAGACAGGTAAGCGACGAGGGAACAGTGGGTGGAGCGGTGAAATGCGTAGAGATCCACTGGAACGCCGATGGTGAAGATGCTCGTCTGGGCCTGTTCTGACGCTGAGGCACGAAAGCGTGGGGAGCAAACAGG
## 2217                                                         ACGAGAGGTCCAAGCGTTAGTCGGAATTACTGGGCTTAAAGGGTGCGTAGGCGGGTCTGTAAGTGCCTTGTGAAATCCCCGGGCTCAACCTGGGAATTGCAGGGCATACTGCGGACCTTGAGGAAGCTAGAGGCCGGCAGAACGATAGGTGGAGCGGTGGAATGCGTAGATATCTATCGGAATGCCAAAGGTGAAGACAGCCGGCTGGGGCTTTCCTGACGCTGAGGCACGAAAGCGTGGGGATCAAACAGG
## 2219                                                         ACAGAGGTCCCAAGCGTTGTTCGGAATCACTGGGCGTAAAGGGTGCGTAGGCGGTATAGTAAGACTTGGGTGAAATCTCCATGCTTAACGTGGAGGGAGCCTGGGAGACTGCTGTGCTAGAGGATTGGAAGGGAGACTGGAATTCTTGGAGTAGCAGTGAAATGCGTAGATATCAAGAGGAACACCAGTGGCGAAGGCGAGTCTCTGGACAAGACCTGACGCTGAGGCACGAAGGCCAGGGGATCAAACGGG
## 2220                                                         ACGGGGGGGGCAAGCGTTGTTCGGAATTACTGGGCGTAAAGGGCTCGTAGGTGGCCACCTAAGTCTGACGTGAAATCCCTCGGCTTAACCGGGGAACTGCGTCGGATACTGGGTGGCTTGAATCCGGGAGAGGGACGTGGAATTCCAGGTGTAGCGGTGAAATGCGTAGATATCTGGAGGGACACCGGTGGCGAAGGCGGCGTCCTGGACCGGAATTGACACTGAGGAGCGAAAGCCAGGGGAGCAAACGGG
## 2221                                                         ACGGAGGGTGCAAACGTTGTTCGGAATCATTGGGCGTAAAGCGCACGCAGGCGGTTTATCAGGTCGGATGTGAAATGCCGGGGCTCAACCCCGGAACGGCGTTCGAAACCGGTAAGCTGGAGTCACGGAGGGAGAAGGGGAATTCCTGGTGTAGGGGTGAAATCCGTAGAGATCAGGAGGAACACCGGCGGCGAAAGCGCCTTCTTGGACGATGACTGACACTGAGGTGCGAAAGCGTGGGGAGCAAACAGG
## 2224                                                          ACGAAGAGGGCAAGTGTTATTCGTCTTGACTGGGCGTAAAGCGTGCGTCGGTTGTTACTTAACGTTCTGTTTAAAGCATAGAAATTATTTTTATACAAACATTACAAGTGAGTACCTAGAGTATATGGGAGTTTAGCAGTACGCTGTGTGTAGAGGTAGAATTCTTTGATCCACAGAGGACTGTCATTCGACGACGGTTGCTAATCAACATATACTGACACTTAAGCACAAAAGCATAGGGAGCAAACAGG
## 2229                                                         ACAGAGGATGCAAGCGTTATCCGGAATCACTGGGCATAAAGCGTCTGTAGGTGGTTTGGTAAGTCTGCTGTTAAAGACTGGGGCTCAACCCCAGAAAAGCAGTGGAAACTGCCAGACTTGAGTGTGGTAGAGGTAAAGGGAATTCCTAGTGTAGCGGTGAAATGCGTAGATATTAGGAAGAACACCAATGGCGAAGGCAGGTCTCTGGGCCATAACTGACACTGAGGAGCGAAAGCGCGGGGAGCGAACAGG
## 2230                                                        ACGTAGGGTGCAAGCGTTAATCGGAATTACTGGGCGTAAAGAGCGCGTAGGCGGCTCGTTAAGTGTGAAGTGAAATCTCCAGTGCTCAACACGGAAACTGCTTTACATACTGGCGAGCTTGAGGAAAGCAGAGGCAACTGGAATTCCTGGTGTAGCGGTGAAATGCGTTGATATCAGGAGGAACACCCATGGCGAAGGCAGGTTGCTGGGCTTTATCTGACGCTGAGGCGCGAAAGCGTGGGTAGCAAACAGG
## 2234                                                         ACATAGGGTGCAAGCGTTGTCCGGAATTATTGGGCGTAAAGAGCTCGTAGGTGGTTCGATACGTCGGATGTGAAAATCAGGGGCTCAACCCCTGACCTGCATCCGATACGGTCGAGCTAGAGTTTGGTAGGGGAGACTGGAATTCCTGGTGTAGCGGTGGAATGCGCAGATATCAGGAGGAACACCGATGGCGAAGGCAGTCCCCTGGGATAAGACTGACGCTCATGCACGAAAGCGTGGGGAGCAAACAGG
## 2236                                                         ACAGGGGGCACAAGCGTTGCTCGGAATCACTGGGCGTAAAGCGTACGCAGGCGGACAGTCAAGTCGGGTGTGAAAGCCCGGGGCTCAACCCCGGAATTGCATTCGATACTAGCTGTCTTGAATACGGGAGAGGTGAGTGGAATTCCTGGTGTAGAGGTGAAATTCGTAGATATCAGGAGGAACACCGGTGGCGAAAGCGGCTCACTGGCCCGATATTGACGCTCATGTACGAAAGCGTGGGAAGCAAACAGG
## 2238                                                         ACAGAGAGTGCAAGCGTTAATCGGAATTATTGGGCGTAAAGGGTGTGTAGGTGGTTAAAATAGTTAAATGTGAAATCCCAAAGCTTAACTTTGGAACTGCGTTTAATACTTTTTGACTAGAGTATTGTAGGGGAAAGTAGAATTTTCGGTGTAGCGGTGAAATGCGTAGATATCGAAAGGAATACCAGAGGCGAAGGCGACTTTCTGGGCAAATACTGACACTGAAACACGAAAGCGTGGGGAGCAAACAGG
## 2239                                                         ACGTAGGGGGCAAGCGTTGTCCGGAATTATTGGGCGTAAAGCGCGTGTAGGCGGCTCGATAAGTCCGCTCTGAAAGCCTAGGGCTCAACCCTAGGAGGCGGGTGGATACTGTCGAGCTTGAGTCTGGAAGAGGCGAGTGGAATTCCCGGTGTAGCGGTGAAATGCGCAGATATCGGGAGGAACACCAATTGCGAAGGCAGCTCGCTGGGACATGACTGACGCTGAGACGCGAAAGCGTGGGGAGCAAACAGG
## 2241                                                         ACGGAGGGTGCAAGCGTTATCCGGAATTACTGGGTTTAAAGGGTGCGTAGGCGGCGCTGTAAGTCAGGGGTGAAAGGTTCCGGCTCAACCGGAGAATTGCCTTTGATACTGCAGTGCTAGGATTGGGTTGAGGTTGGCGGAATGTGACATGTAGCGGTGAAATGCATAGATATGTCATAGAACACCGATTGCGAAGGCAGCCAACTGGACCTTGATTGACGCTGAGGCACGAAAGCGTGGGGAGCAAACAGG
## 2242                                                         ACGTAGGGTGCGAGCGTTAATCGGAATTACTGGGCGTAAAGCGTGCGCAGGCGGTTATGTAAGACAGTTGTGAAATCCCCGGGCTCAACCTGGGAATTGCATCTGTGACTGCATAGCTAGAGTACGGTAGAGGGGGATGGAATTCCGCGTGTAGCAGTGAAATGCGTAGATATGCGGAGGAACACCGATGGCGAAGGCAGTCCCCTGGGATAAGACTGACGCTCATGCACGAAAGCGTGGGGAGCAAACAGG
## 2244                                                         ACAGAGGGTGCAAACGTTGTTCGGAATCACTGGGCGTAAAGCGCGTGTAGGCGGCTTCTCAAGTCGGGTGTGAAATCCCAGGGCTCAACCCTGGAAGTGCATTCGAAACTGGGTAGCTAGAGTACTGGAGAGGAAGGTGGAATGCTTGGTGTAGAGGTGAAATTCGTAGATATCAAGCGGAACACCGGTGGCGAAGGCGGCCTTCTGGACAGATACTGACGCTGAGACGCGAAAGCGTGGGGAGCAAACAGG
## 2246                                                         ACGTAGGGTGCAAGCGTTACTCGGAATTATTGGGCGTAAAGCGCGTGTAGGTGGTCATGTAAGTCTGATGTGAAATCCCTGGACTCAATTCAGGAACTGCATTGGATACTGCATGGCTAGAGTATGGGAGAGGTAGGCGGAATTCCTGGTGTAGAGGTGAAATTCGTAGATATCAGGAGGAACAACAGTGGCGAAGGCGGCTTACTGGACCATAACTGACACTGAGACGCGACAGTGTGGGGAGCAAACAGG
## 2248                                                         ACGTAGGGGGCGAGCGTTGTCCGGAATGATTGGGCGTAAAGGGCGCGTAGGCGGCTTTGTAAGTCTGGAGTGAAAGTCCTCTTTTCAAGGGAGGAATTGCTTTGGAAACTATAGAGCTTGAGTGCAGGAGAGGTTATCGGAATTCCCGGTGTAGCGGTGAAATGCGTAGATATCGGGAGGAACATCGGTGGCGAAGGCGGGTAACTGGACTGCAACTGACGCTGAGGCGCGAAAGTGTGGGGAGCAAACAGG
## 2250                                                         ACGGAGGATGCAAGCGTTATCCGGAATCATTGGGTTTAAAGGGTCCGTAGGCGGTTTAATAAGTCAGTGGTGAAAGCCCATCGCTCAACGATGGAACGGCCATTGATACTGTTAAACTTGAATTATTAGGAAGTAACTAGAATATGTAGTGTAGCGGTGAAATGCTTAGAGATTACATGGAATACCAATTGCGAAGGCAGGTTACTACTAATGGATTGACGCTGATGGACGAAAGCGTGGGTAGCGAACAGG
## 2257                                                         ACGGAGGGGACGAGTGTTACTCATAATAACTGGGCGTAAAGGGTATGTAGATGGTTCATTTTGTTTATTATTAAAGAATAAATTAATAGTTTATTATAATAATTAATACGTTTAAACTAGAGTATAATAGAGGAATATGGAATTTTTAGTGGAAGGGTGAAATTTTACAATAATAAAAAGACGATCAAAAGCGAAGGCAATATTCTGGATTATTACTGACATTGATATACTGAAGCATGGGGAGCAAATCGG
## 2258                                                        ACCAGCGCCACGAGTGGGAGCCACGATTATTGGGCCTAAAGCGTCCGTAGCTGATCGAGTAAATTTCTTGTGAAATTGTCCTGCTTAACAGGTCAACGTGCAGGAAACACTGCTCGACTTGGGACTGGGAGGGGTTAGGAGTATCTTAGGGGTAGCGGTAAAATGTTATAATCCTTAAGAGACTACCGGTGGCGAAGGCGCCTAACCAGAACAGATCCGACAGTGAGGGACGAAAGCTAAGGGAGCAAATCGG
## 2263                                                         ACAGAGGGTGCAAGCGTTAATCGGAATTACTGGGCGTAAAGGGCCTGTAGGTGGTTACTTAAGTTAGATGTGAAAGCCCTGGGCTCAACCTGGGAACTGCGTCTAAAACTGGGTGACTAGAGTATGGTAGAGGGAAGTGGAATTTCTAGTGTAGCGGTGAAATGCGTAGATATTAGAAAGAACACCAGTGGCGAAGGCGACTTCCTGGACCAATACTGACACTGAGGGGCGAAAGCGTGGGGAGCAAACAGG
## 2265                                                         ACAGGGGATGCGAGTGTTATCCGGAATTATTGGGCGTAAAGCGTCTGTAGATGGATTTATAAGTCTTTTATTAAATCTTTAAGCTCAACTTAAAAAAAGTGAAAGATACTATAAATCTTGAGTTTAGTAGAGGTATGGAGAATTTCAAGTGGAGCGGTGAAATGTGTAGATATTTGAAGGAATACCAATGGCGAAAGCACTGTACTGGTCTAAAACTGACATTGAGAGACGAAAGCTAGGGGAGCAAACAGG
## 2271                                                          ACGTAGGGTGCAAGCGTTATCCGGAATTATTGGGCGTAAAGCGTCTGTAGGCGGTATGTTAAGTAGGAGATAAAATCTTCGGGCCCAACCCGAAACATGTTTCCTAAACTGGCGTACTAGAGGTATGGAGAGGTAACTGGAATTTCTGGTGTAGGGGTAAAATCCGTTGATATCAGAAGGAACGCTAAAAGCGAAGGCAGGTTACTGGCCATTTCTGACGCTGAGAGACGAAAGCGTGGGTAGCAAACAGG
## 2273                                                         ACGAGGGGTGCGAGCGTTGTTCGGAATCACTGGGCGTAAAGGGCGCGTAGGCGGCCTTCTAAGTCGGGAGTCTAATCACGGGGCCTAACCCCGTGGCCGCTCCCGATACTGGAAGGCTTGGATCCCTCAGGGGCTGTCGGAACACCTGGTGTAGCGGTGGAATGCGTAGATATCAGGTGGAACACCAGTGGCGAAGGCGGACAGCTGGGAGGAGATCGACGCTGAGGCGCGAAAGCATGGGGAGCGAACAGG
## 2279                                                         ACAGAGGGTGCAAGCGTTAATCGGAATTACTGGGCGTAAAGCGTGCGTAGACGGTTATCTAAGTCGGATGTGAAATCCCCGGGCTCAACCTGGGAACTGCGTTTGTGACTGTGTTGCTAGAGTGCGGCAGAGGGGGGTGGAATTCCACGTGTAGCAGTGAAATGCGTAGAGATGTGGAGGAACACCGATGGCGAAGGCAGCCCCCTGGGTCGACACTGACGCTCATGCACGAAAGCGTGGGGAGCAAACAGG
## 2280                                                         ACGTAGGGTGCAAGCGTTAATCGGAATTACTGGGCGTAAAGCGTGCGCAGGCGGTTATGTAAGACAGATGTGAAATCCCCGGGCTCAACCTGGGAACTGCATTTGTGACTGCATAGCTGGAGTGCGGCAGAGGGGGATGGAATTCCGCGTGTAGCAGTGAAATGCGTAGATATGCGGAGGAACACCGATGGCGAAGGCAATCCCCTGGACCTGTACTGACGCTCATGCACGAAAGCGTGGGGAGCAAACAGG
## 2281                                                         ACGTAGGGTGCGAGCGTTAATCGGAATTACTGGGCGTAAAGGGTGCGTAGGCGGGTAGATAAGTTAGATGTGAAATCCCTGAGCTTAACTTAGGAAAGGCATTTAAGACTGTTTATCTAGAGTATATCAGAGGGTGGTAGAATTCCAAGTGTAGCAGTGAAATGCGTAGATATTTGGAGGAATACCAGTGGCGAAGGCGGCCATCTGGGGTAATACTGACGCTGAGGCACGAAAGCGTGGGGAGCAAACAGG
## 2283                                                         ACGTAGGTCACGAACGTTATTCGGAATCACTGGGCTTAAAGGGCGCGTAGGCGGCTTCGTAAGTCAAGGTTGAAATCCGGCGGCTCACCCGTCGAACTGGCCTTGATACTGCGGAGCTTGAGGTGCGGAGGGATGCGTGGAACTTGTGGTGGAGCGGTGAAATGCCCAGATATCACAGGGAACACCGGTGGCGAAGGCGACGCATTGGCCGCAATCTGACGCTGAGGCGCGAAAGCATGGGGAGCGAACGGG
## 2285                                                         ACGTAGGGTGCGAGCGTTAATCGGAATTACTGGGCGTAAAGCGTGCGCAGGCGGTTATGTAAGACAGAGGTGAAATCCCCGGGCTCAACCTGGGAACTGCCTTTGTGACTGCATAGCTGGAGTGCGGCAGAGGGGGATGGAATTCCGCGTGTAGCAGTGAAATGCGTAGATATGCGGAGGAACACCGATGGCGAAGGCAATCCCCTGGACCTGTACTGACGCTCATGCACGAAAGCGTGGGGAGCAAACAGG
## 2288                                                         ACAGGGGATGCAAGCGTTACTCGGAATTACTGGGCGTAAAGCGTCTGCAGGTGTCTTGCCACGTTTGGTGTCAAATGATGCGGCTCAACCGCGTCAACGTGCCGAAAACGAGCGAGATCGAGTCATTCAGAGGTACCTGGAATGTCGTGTGTAGGGGTAAAATCCGTAGATCCACGATGGAACGCCAAAAGCGAAGGCAGGGTACTGGGAATGTACTGACACTCAGAGACGAAAGCGTGGGGAGCAAAGGGG
## 2292                                                         ACGGAGGGTGCAAGCGTTATTCGGATTTATTGGGCGTAAAGAGCGTGTAGGCGGCCGGTCAAGCCAGATGTGAAAGCCCAGGGCTTAACCCTGGAAGTGCATTTGGAACTGGTTGGCTTGAGTTCGGGAGAGGGAAGTGGAATTCCGGGTGTAGAGGTGAAATTCGTAGATATCCGGAGGAACACCGGTGGCGAAGGCGACTTCCTGGACCGAAACTGACGCTGAGACGCGAAAGCGTGGGGAGCAAACAGG
## 2301                                                           ACGGAGAGGGCAAGTGTTATTCGTATTTACTGGGCGTAAAGCGTATGGAGGTTGTTTATTAAGTTATAGATGAAAATAGAAAGTTATAATTCTAAGATTTATAATACTGATTTACTTGAGTATTTTAGAGTATAGTAGTACTTTAAGTGGAAAGGTAACATTTTTCAATATTTAAAGGACTAACATTCGACGACGGTTACTATTCATAAAATACTGACACTCAAGTACGAAAGCATGGGGAGCAAACAGG
## 2306                                                         ACGTAGGGTGCAAGCGTTAATCGGAATTACTGGGCGTAAAGCGTGCGCAGGCGGTTTCGTAAGACAGAGGTGAAATCCCCGGGCTCAACCTGGGAACTGCCTTTGTGACTGCGAGGCTAGAGTATGGCAGAGGGGGGTGGAATTCCACGTGTAGCAGTGAAATGCGTAGAGATGTGGAGGAACACCGATGGCGAAGGCAGCCCCCTGGGTCGACACTGACGCTCATGCACGAAAGCGTGGGGAGCAAACAGG
## 2324                                                        ACAGAGACCTCAAGCATTATCCGGATTTATTGGGCGTAAAGGGTCCGCAGGTGGCCCGGCACGTCATCGGTCAAATCTTCGGGCTCAACCCGAAAACCGCCGATGATACGGCCAAGGCTAGAGAGTGGGAGAGGCGAGCGGAATTACCGGTGTAGTAGTAAAATGCGTTAATATCGGTAAGAACACCAAATGCGAAGGCAGCTCGCTGGAACACTTCTGACACTCAGGGACGAAAGCGTGGGGAGCGAAAGGG
## 2327                                                          ACGTAGGGGGCGAGCGTTGTCCGGAATTACTGGGCGTAAAGGGCATGCAGGCTGTGAATCAAGTCAAGTGTTAAAGGATGCGGCTCAACCGTGTTATGCACTTGAAACTGATTTACTGGAGTGTTGGAGAGGCAAGTGGAATTCCCAGTGTAGCGGTGAAATGCGTAGATATTGGGAAGAACATCGGTGGCGAAGGCGACTTGCTGGCCGACAACTGACGCTGATATGCGAAAGCCAGGGGAGCAAACGGG
## 2329                                                        ACGGAGGGTGCAAGCGTTATCCGGAATCACTGGGTTTAAAGGGTGCGTAGGCGGTGATGTAAGTCAGTCGTGAAATTCCCCGGCTTAACCGGTGGGACTGCGATTGATACTGTGTTACTTGAATCGGGTTGAGGCAGGCGGAATGTGACATGTAGCGGTGAAATGCTTAGATATGTCATAGAACACCGATTGCGAAGGCAGCCTGCTGGGCCTTGATTGACGCTGAGGCACGAAAGCGTGGGGAGCAAACAGG
## 2331                                                         ACGAGAGCCCCGAGCGTTATCCGGAATTATTGGGCGTAAAGGGTGCGTAGGTGGTTGTGTTAGTCGTTTGTCAAATCCTTGGGCTTAACCCAAGAACCGCGAACGAAACGGCACAACTTGAGAATGTAAGGGGTATAGGGAACTCATGGTGTAGGGGTGAAATCCGTTGATATCATGGGGAACACCAAAAGCGAAGGCACTATACTGGTACATTTCTGACACTCAAGCACGAAAGCGTGGGTAGCGAATGGG
## 2341                                                         ACGTAGGTGGCAAGCGTTGTCCGGATTTATTGGGTTTAAAGGGTGCGTAGGCGGTTTGATAAGTCAGTCGTTAAAGCCATCAGCTTAACTGATGCAAGGCGATTGATACTGTCAGACTTGAATTCTTTGGAGGTAGCCGGAATGCGTAGTGTAGCGGTGAAATGCTCAGATATTACGCAGAACACCGATTGCGGAGGCAGGTTGCTACAAAGAGATTGACGCTGAGGCACGAAAGTGTGGGGAGCAAACAGG
## 2343                                                         ACGTAGGGTGCTAGCGTTACTCGGATTTACTGGGCGTAAAGCACATGTAGGCTGGTTGGTAAGACAGAAGTGCAATCCCTGAGCTTAACTCAGGAACTGCTTTTGTTACTGCCAATCTGGAGTATGATATGGGATAATGGAATTCCTAGTGTAGGGGTAAAATCCGTAGATATTAGGAGGAACACCAAAAGCGAAAGCAATTATCTGGGTCATTACTGACGCTGAGATGTGAAAGCGTGGGGAGCAAACAGG
## 2345                                                         ACGTAGGGTGCAAGCGTTAATCGGAATTACTGGGCGTAAAGCGTGCGCAGGCGGTTATGTAAGACAGATGTGAAATCCCCGGGCTCAACCTGGGAACTGCATTTGTGACTGCATAGCTAGAGTACGGTAGAGGGGGATGGAATTCCGCGTGTAGCAGTGAAATGCGTAGATATGCGGAGGAACACCGATGGCGAAGGCAATCCCCTGGGCCTGTACTGACGCTCATGCACGAAAGCGTGGGGAGCAAACAGG
## 2351                                                         ACAGAGGGCTCAAGCGTTAATCGGAATCACTGGGCTTAAAGCGAGCGTAGGCGGATCGGTAGGCACGTTGTGAAAGCCCACGGCTCAACCGTGGAATTGCAGCGTGAACCCCCGATCTTGAGGCGAGTAGAGGTAGCCAGAACGATAGGTGGAGCGGTGAAATGCGTAGAGATCTATCGGAATGCCGAAGGTGAAGACAGGCTACTGGGCTCGAACTGACGCTGAGGCTCGAAAGCGTGGGGATCAAACGGG
## 2359                                                        ACCGGCGCCACAAGTGGAGATCATGTTTATTGGGTCTAAAGTGTTCGTAGCTGGCTTTGTAAGTTTCTTGTGAAAGTTCTGTGGCTTAACCCAGAAATTGCAAGGAATACTGCAAGGCTTGAGATTGGGAGAGGTATGAGGTATTGTTTGGGTAGCGGTAAAATGTTATAATCCTTACAAGACCAACTAAGGCGAAGGCATCATACTTGAACAAATCTGACAGTGAGGAACGAGAGCTGGGGGAGCGAACCGG
## 2365                                                         ACGGAGGGTGCAAGCGTTATCCGGATTCACTGGGTTTAAAGGGTGCGTAGGCGGGCTTATAAGTCAGTGGTGAAAGCCCGGAGCTTAACTTCGGAACTGCCATTGATACTGTAAGTCTTGAGTACACTTGAGGTGGGCGGAATACATCATGTAGTGGTGAAATACTTAGATATGATGTAGAACACCGATTGCGAAGGCAGCTCACTAAATTGTAACTGACGCTGAGGCACGAAAGCGTGGGGATCAAACAGG
## 2370                                                         ACGGAGGGTGCAAGCGTTGTCCGGATTTATTGGGTTTAAAGGGTGCGTAGGCGGCCATTTAAGTCAGCGGTGAAAGACTTCAGCTCAACTGAAGTACTGCCGTTGATACTGAGTGGCTTGAGTATTGACGAAGAGGGCAGAATTGAAGGTGTAGCGGTGAAATGCATAGATATCTTCAAGAATACCGATTGCGAAGGCAGCTCTCTAGGGAATAACTGACGCTGAGGCACGAAAGCGTGGGTAGCAAACAGG
## 2371                                                         ACAGAGACCTCAAGCGTTATCCGGAATTATTGGGCGTAAAGCGTCCGCAGGTGGTTTGGTAAGTCAGGGGTTAAAACTTCGCGCTTAACGTGAAGATTGCTTCTGAAACTACCATTCTAGAGTATGGGAGAGGTAAGCGGAATTCTCGGTGTAGTCGTAATAAGCGTTGATATCGAGAAGAACACCAAATGCGAAGGCAGCTTACTGGAACATTACTGACACTCATGGACGAAAGCGTGGGGAGCAAACAGG
## 2375                                                         ACAGAGGCCCCGAGCGTTACCCGGAATCACTGGGCGTAAAGCGTGTGTCGGTGGTCGTATTAGTCGTCCGTTAAAGCCCCGGGCTCAACCCGGGAAGTGCGGACGAAACGGTACGACTCGAGGGTGTGAGAGGTGAGTGGAACTCATGGTGTAGGGGTGAAATCCGTTGATATCATGGGGAACACCGAAAGCGAAGGCAACTCACTGGCGCATTCCTGACACTTAAACACGAAAGCGTAGGTAGCGAACGGG
## 2377                                                         ACAGAGGATGCAAGCGTTATCCGGAATCACTGGGCATAAAGCGTCTGTAGGTTGCTTGCCAAGTCTGCTGTTAAAGATCAGGGCTTAACCCTGGGAAAGCAGTGGAAACTAGTAGGCTTGAGTGTGGTAGAGGTAGAGGGAATTCCTGGTGTAGCGGTGAAATGCGTAGATATTAGGAAGAACACCAATGGCGAAAGCACTCTACTGGGCCATAACTGACACTGAGAGACGACAGCTAGGGGAGCAAATGGG
## 2382                                                         ACGTAGGGTGCAAGCGTTAATCGGAATTACTGGGCGTAAAGCGTGCGCAGGCGGTTATGTAAGACAGATGTGAAATCCCCGGGCTCAACCTGGGAACTGCATTTGTGACTGCATAGCTAGAGTACGGTAGAGGGGGATGGAATTCCGCGTGTAGCAGTGAAATGCGTAGATATGCGGAGGAACACCGATGGCGAAGGCAATCCCCTGGGCCTGCACTGACGCTCATGCACGAAAGCGTGGGGAGCAAACAGG
## 2385                                                         ACGTAGGGTGCGAGCGTTAATCGGAATTACTGGGCGTAAAGCGTGCGCAGGCGGTTTTGTAAGACAGATGTGAAATCCCCGGGCTCAACCTGGGACCTGCATTTGTGACTGCAAGGCTAGAGTACGGTAGAGGGGGATGGAATTCCGCGTGTAGCAGTGAAATGCGTAGATATGCGGAGGAACACCGATGGCGAAGGCAATCCCCTGGGCCTGCACTGACGCTCATGCACGAAAGCGTGGGGAGCAAACAGG
## 2388                                                      CCCAGCAGTCCAAGTTGCAGCCAACTTTATTGGGTCTAAAACATCCGTAGCTTGCTTAACAAGTTCTTTGTGAAATCCGGCATCTTAAGTGTCGGGCGCGCAAAGAATACTGTTAGGCTAGAGACCGGTAGATGTAAAGAGTATGTCTAAGGTAGTGGTAAAATACGTTAATCTTAGGCAGACTAACAATCGAGCGAAGGCACTTTACAAGGACGGATCTGACAGTGAGGGATGAAGGCTAGGGGCGCAAAACGG
## 2392                                                        ACCGGCGGCTCGAGTGGTAACCGTTATTATTGGGTCTAAAGGGTCTGTAGCCGGCCGGGTAAGTCTCTTGGGAAATCCAGCAGCTTAACTGTTGGGCTTTCAGGAGATACTGCCTGGCTCGAGGCCGGGAGAGGTGAGAGGTACTTCAGGGGTAGGGGTGAAATCTTGTAATCCTTGAAGGACCACCAGTGGCGAAGGCGTCTCACCAGAACGGACCTGACGGCAAGGGACGAAAGCTAGGGGCACGAACCGG
## 2394                                                         ACGTAGGGTGCAAGCGTTAATCGGAATTACTGGGCGTAAAGCGTGCGCAGGCGGTTATGTAAGACAGAGGTGAAATCCCCGGGCTCAACCTGGGAACGGCCTTTGTGACTGCATAGCTAGAGTACGGTAGAGGGGGATGGAATTCCGCGTGTAGCAGTGAAATGCGTAGATATGCGGAGGAACACCGATGGCGAAGGCAATCCCCTGGGCCTGTACTGACGCTCATGCACGAAAGCGTGGGGAGCAAACAGG
## 2398                                                        ACGAGTGCCCCAAGCGTTATCCGGAATTATTGGGCGTAAAGGGTGTGTAGGTGGCACTGTTAGTCTTTCGTGAAAGCTCTCCGGCTTAACTGGGGAAACGCGGGGGAAACGGCAGCGCTAGAGGAGGCGAGGGGTCTATGGAACTCATGGTGTAGGGGTGAAATCCGTTGATATCATGGGGAACACCAAAAGCGAAGGCAGTAGACTGGCGCCTAACTGACACTGAAACACGAAAGCGTGGGGATCAAACGGG
## 2402                                                          ACGGGGGGGCAAGCGTTGTTCGGAATTACTGGGCGTAAAGCGCGTGTAGGCGGCTTTTCAAGTCAGACGTGAAAGCCCTGGGCTTAACCCAGGAATTGCGTTTGATACTGTAAGGCTAGAGTGTGGAAGAGGCTACTGGAATTCCGTGTGTAGCGGTGAAATGCGTAGATATACGGAGGAACACCTGCGGCGAAGGCGGGTAGCTGGGCCAACACTGACGCTGAGACGCGAAAGCTAGGGTAGCAAACGGG
## 2403                                                         ACGTACCGGACAAACGTTATTCGGAATCACTGGGCTTAAAGAGTGCGTAGGCGGCCTAGCAGGCGAGGTGTGAAATCCCACGGCTCAACCGTGGAATTGCGCTTCGAACCGCTAGGCTCGAGGGAGACAGGGGTGAGTGGAACTGATGGTGGAGCGGTGAAATGCGTTGATATCATCAGGAACACCAGTGGCGAAGGCGACTCACTGGGTCTCTACTGACGCTGAGGCACGAAAGCTAGGGGAGCAAACGGG
## 2406                                                         ACGTAGGGGGCAAGCGTTGTCCGGAATTACTGGGCGTAAAGCGCACGCAGGTTGAATTGTAAGTCAGCTGTAAAAGACCACGGCTCAACTGTGGCAAGTCGGCTGATACTGCAAATCTAGAGTATGTGAGAGGGAAGCGGAATTCCCGGTGTAGCGGTGAAATGCGTAGATATCGGGAGGAACACCAGTGGCGAAGGCGGCTTCCTGGCACAATACTGACGCTCATGTGCGAAAGCTAGGGCAGCGAACGGG
## 2409                                                         ACGGAGGGTGCAAGCGTTGCTCGGAATTACTGGGCGTAAAGGGCGTGTAGGCGGTTTCGTATGTCTGGCGTGAAAGCCCCGGGCTCACCTCGGGAAGTGCGTCGGAAACTGCGAGACTAGAGTCCCGGAGAGGGAGGTGGAATTCCAGGTGTAGCGGTGAAATGCGTAGATATCTGGAAGAACACCGGTGGCGAAAGCGGCCTCCTGGACGGTGACTGACGCTGAGACGCGAAAGCGTGGGGAGCAAACAGG
## 2415                                                         ACAGAGGATGCAAACGTTGCTCGGAATTACTGGGCGTAAAGCGCGCGCAGGTGGTTCGATAAGTCGGGTGTGAAATCCCGGGGCTCAACCTCGGACGTGCACTCGATACTGTCAAACTTGAGTACTGGAGAGGATCGCGGAATTCCCGGTGTAGAGGTGAAATTCGTAGATATCGGGAGGAACACCAGTGGCGAAGGCGGCGATCTGGACAGATACTGACACTGAGGCGCGAAAGCGTGGGAAGCAAACAGG
## 2421                                                       ACGAAGGGGGCGAGCGTTATCCAGAATGATTGGGCGTAAAGTGTTCGTAGATGGTAATTTTTGTTATATGTAAAATATTAAAAGTTTTTTTTAACGATGCATATAAAACGTTTTTACTTGAGTATTATAGGGGATACACAGAATTCTATGTATAGGGGTAAAATCCAGTTAATCATAGAGGAATGCCAGTGGCGAAGGTGGTTATCTATATTTTTTACTGACATTGAGGAACGAAAGCATGGGGAGCAAACAGG
## 2423                                                         ACCAGCACCCCGAGTGGTCGGGACGATTATTGGGCCTAAAGCATCCGTAGCCGGTTTTACAAGTCCTCCGTTAAATCCAACTGCTTAACAGATGGGCCGCGGAGGATACTATAAGACTAGGAGGCAGGAGAGGCAAGCGGTACTCAGTGGGTAGGGGTAAAATCCGTTGATCCATTGAAGACCACCAGTGGCGAAGGCGGCTTGCCAGAATGCGCTCGACGGTGAGGGATGAAAGCTGGGGGAGCAAACCGG
## 2424                                                         ACGGAGGGTGCAAGCGTTATCCGGAATCATTGGGTTTAAAGGGTCCGCAGGCGGGCTTATAAGTCAGTGGTGAAAGCCTACAGCTTAACTGTAGAACTGCCATTGATACTGTAAGTCTTGAATTCGGTCGAAGTGGGCGGAATGTGTAGTGTAGCGGTGAAATGCTTAGATATTACACAGAACACCGATAGCGAAGGCAGCTCACTAGGCCTGGATTGACGCTCAGGGACGAAAGCGTGGGGAGCAAACAGG
## 2425                                                         ACGAGGGATCCTAGCGTTGTTCGGAATCATTGGGCGTAAAGCGGGTGTAGGCGGTCTTATAAGTCAGGTGTGAAATCCCGGGGCTCAACCCCGGAAGTGCACTTGATACTGTAAGTCTTGAATGTGGTAGAGGCTACTGGAATTCTTGGTGTAGTGGTGAAATACGTAGATATCAAGAGGAATACCGGAGGCGAAGGCGGGTAGCTGGGCCAACATTGACGCTGAGACCCGAAAGTGTGGGGATCAAACAGG
## 2426                                                            AAGGGGGCGAGCGTTGTTCGGAATTACTGGGCTTAAAGCGTGCGTAGGCGGACAAGAAGGTCAGAAGTGAAATCCCGGGGCTTAACCCCGGAACTGCTTTTGAAACTCCTAGTCTAGAATTCGGAAGAGGTTGGCGGAATACCCAGTGTAGAGGTGAAATTCGCAGATATTGGGTAGAACACCGGTGGCGAAGGCGGCCAACTGGTCCGACATTGACGCTGAGGCACGAAAGCGTGGGGAGCAAACAGG
## 2431                                                         ACATAGGGTGCAAGCGTTGTCCGGAATTATTGGGCGTAAAGAGCTCGTAGGTTGTTTGTCGCGTCGATTGTGAAAATCTGAGGCTCAACCTCAGACCTGCAGTCGATACGGGCAAACTAGAGTGTGGTAGGGGAGACTGGAATTCCTGGTGTAGCGGTGGAATGCGCAGATATCAGGAGGAACACCAATGGCGAAGGCAGGTCTCTGGGCCATTACTGACACTGAGGAGCGAAAGCGCGGGGAGCGAACAGG
## 2435                                                         ACGTAGGTGGCAAGCGTTGTCCGGATTCATTGGGTTTAAAGGGTGCGTAGGCGGCTATTTAAGTCAGTGCTGAAATACCCCGGCTTAACCGGGGGGGTGGCATTGATACTGGGTAGCTTGAGTGCTAGCGAGGCAGGCGGAATTGACGGTGTAGCGGTGAAATGCATAGATATCGTCAAGAACACCGATAGTGTAGACAGCTTGCCAGGGAGCAACTGACGCTGAGGCACGAAAGTGTGGGGATCAAACAGG
## 2439                                                         ACAGAGGGTGCAAGCATTAATCGGATTTATTGGGCGTAAAGAGCGCGTAGGCGGGTTTTTAAGTCAGATGTGAAATCCCGGGGCTCAACCCCGGAACAGCATTTGAAACTGAAAATCTTGAGGGTAGTCGGAGAAAATGGAATTCCACGTGTAGCGGTGAAATGCGTAGATATGTGGAAGAACACCGGTGGCGAAGGCGATTTTCTAGATTATTCCTGACGCTGATGCGCGAAAGCAAGGGGATCAAACAGG
## 2440                                                       ACAGGAGGGGCAGGCGTTATTCATAATTATTGAGCGTAAAGGGTAAGTCAACGGTTTTTTTAATACTTTATAAAAGTCTGGGTTTAATACTTAGAGCAAGTAATTTTGTAAAATAGAACTTGAGTTATTTAATAAGATAGTAGAATTTCTAAGGTAAAAGTATAATTTTATGATCTTAGAAGGAATACCAATAGCGAAAGCAACTATCAATTAATAACTGACGTTTAGGTACGTAGGTATAGGGAGCAAACGGG
## 2441                                                         ACAGAGGCCTCAAGCGTTGTTCGGAATTACTGGGCGTAAAGGGTGTGTAGGCGGTAAAGTAAGTCGAACTATTAAATACCATCGCTCAACGATGGGGTCTAGCCGATACTATTTTACTAGAGGGTCTCAAAGAAGATGGGAATTCACGGTGTAGCAGTGAAATGCGTGGATATCGTGAGGAACACCAGAGGCGAAGGCGCATCTTTAGGGGATTTCTGACGCTGAAACACGAAGGTTAGGGGAGCAAATGGG
## 2452                                                         ACAGAGGTCTCAAGCGTTGTTCGGATTTACTGGGCGTAAAGGGTGCGTAGGCTGTTTTGTAAGTCTGATGTGAAATTTTGTGGCTCAACTACAAAAGCGCATTGGAAACTACAAAGCTAGAGGACTGGAGGGGAAGTCGGAATTCTCGGTGTAGCAGTGAAATGCGTAGATATCGAGAGGAACACCAGTGGCGAAGGCGGACTTCTAGACAGTTCCTGACGCTGAAGCACGAAAGCTAGGGGAGCAAACAGG
## 2454                                                         ACAGAGACCTCAAGCGTTATCCGGATTCATTGGGCGTAAAGCGTCCGCAGGTGTCTCGACAAGTTTTCTGTTAAACTCTTGGGCTTAACTCAAGAACTGCGGAAAAAACTGTCGGGATTGAGGCCGGGAGAGGTAAGCGGAACGCATGGTGTAGTCGTAATAAGCGCTGATATCATGCGGAACACCAAAGGCAAAGGCAGCTTACTAGAACGGTTCTGACACTCATGGACGAAAGCGTGGGTAGCGAATGGG
## 2457                                                         ACGGAGGGTCCAAGCGTTGTCCGGATTTATTGGGTTTAAAGGGTACGTAGGCGGGACTTTAAGTCAGCGGTGAAAGCCTCCAGCTTAACTGGAGAATTGCCATTGAAACTGAAGTTCTTGAGTATGGATGAGGTGGTTGGAATATAACATGTAGCGGTGAAATGCTTAGATATGTTATAGAACACCAATTGCGAAGGCAGATCACTAAACCATTACTGACGCTGAGGTACGAAAGCGTGGGGAGCGAACAGG
## 2464                                                         ACGGAGGGTGCAAGCGTTGTTCGGAATTACTGGGCGTAAAGCGCGTGCAGGTGGTCTTTTAAGTTTCGTGTGAAAGCCCTCGGCTTAACCGAGGAACAGCACGGAAAACTGAAAGGCTAGAGTACGGGAGAAGAGGGCGGAATTCCCAGTGTAGAGGTGAAATTCGTAGATATTGGGAGGAACACCGGTGGCGAAAGCGGCTCTCTAGACCGATACTGACGCTCATACGCGAGAGCGTGGGGAGCAAACAGG
## 2465                                                         ACGAGTGCCCCGAGCGTTATCCGGAATTATTGGGTGTAAAGGGTGTGTAGGCGGCTTTGTTAGTCTTTTGTCAAAGCTCCCGGCTTAACCGGGAAAAGGCAAAGGAAACGGCAAGACTTGAGAGTGCGAGGGGTGAACAGAACTCATAGTGTAGGGGTGAAATCCGTTGATATTATGGGGAATACCAAAAGCGAAGGCAGTTCACTGGCGCATTTCTGACGCTGAGACACGAAAGCGTGGGTAGCGAATGGG
## 2467                                                          CGAAGGGTGCAAGCGTTGTTCGGATTTATTGGGCGTAAAGCGCGCGCAGGCGGATGTTTAAGTCAGGTGTGAAATACCCGAGCACAACTTGGGGGGTGCGCCTGAAACTGGAGATCTAGAATAGTGTAGGGGGAAGGAGAATTTCGCATGTAGGGGTAAAATCCGTAGAGATGCGAAGGAATACCGGAGGCAAAAGCGCCTTCCTGGTCACTAATTGACGCTGAGGCGCGAAAGCGTGGGGAGCAAACAGG
## 2468                                                         ACAGAGGATGCAAGCGTTATTCGGAATTACTGGGCGTAAAGCGTGCGTAGGTTGTTTTGTAAGTCATTAGTGAAAACTCGGAGCTCAACTCCGAGCCTGCTTTTGATACTACAAGACTTGAGATTGGAAGAGGTAAGTGGAATTCCTAGTGTAGAGGTAAAATTCGTGGATATTAGGAGGAACACCAACGGCGAAGGCAGCTTACTGGGACATTTCTGACACTGAGGCACTAAAGCGTGGGGAGCAAACAGG
## 2470                                                         ACGTAGGGTGCAAGCGTTAATCGGAATTACTGGGCGTAAAGCGTCCGCAGGCGGTCTTGTAAGTCTGTTGTTAAAGCGTGGAGCTTAACTCCATTTCAGCAATGGAAACTGTAAGACTAGAGTGTGGTAGGGGCAGAGGGAATTCCCGGTGTAGCGGTGAAATGCGTAGATATCGGGAAGAACACCAGTGGCGAAGGCGCTCTGCTGGGCCATAACTGACGCTCATGGACGAAAGCGTGGGGAGCAAACAGG
## 2472                                                         ACGAGGGATGCGAGCGTTGTTCGGAATCACTGGGCGTAAAGAGAGCGTAGGCGGGGAAATAAGTTTGGTGTTTAAAGCAAGGGCCTAACCCTTGTGTGGCACTGAATACTGTTTCTCTTGAGTAGGTTAGGGGTTATCGGAATTCCTGGTGTAGCGGTGGAATGCGTAGATATCAGGAAGAACACCAATGGCGAAGGCAGATAACTGGGACCATACTGACGCTGAGGCTCGAAAGTGCGGGGAGCAAACAGG
## 2473                                                        ACGTAGGGTCCAAGCGTTATCCGGAATTACTGGGCGTAAAGCGTACGCAGATGGTTCAATAGGTGAGATGCGAAATCTTGTGGCTCAACCACATAGACTGTATCTCAAACCATTGAACTAGAGGTTGGTAGAGGCAAGCGGAATTTCTGGTGTAGCAGTGACATGCGTAGATATCAGAAGGAACACCAATGGCGAAGGCAGCTTGCTGGGCCACACCTGACGTTAAGGTACGAAAGCGTGGGGAGCGAACGGG
## 2476                                                         ACGTAAGGTGCAAGCGTTGTTCGGAATTATTGGGCGTAAAGGGCATGTAGGCGGTTCTCTAAGCTTGGTGTGAAATCCCGGGGCTTAACCCCGGAACTGCATTGAGAACTGGAGAACTAGAGTGACTGAGGGGCAGCCAGAATTCCAGGTGTAGGGGTGAAATCTGTAGATATCTGGAAGAATACCAATGGCGAAGGCAGGCTGCCAGCAGATGACTGACGCTGAGGTGCGAAGGTGCGGGGAGCAAACAGG
## 2479                                                         ACGAAGGTGGCGACCGTTGTTCGGAATCACTGGGCATAAAGAGCTCGTAGGCGGTGAGAAAAGTAGGGTGTGAAATCCCTCGGCCCAACCGAGGAACTGCGCTCTAAACTGTCTTGCTTGAGGACCGAAGGGGACCACGGAATTCCTGGTGGAGCGGTGAAATGCGTTGATATCAGGAGGAACACCGGCGGCGAAGGCGGTGGTCTGGTCGGTTTCTGACGCTGAGGAGCGAAAGCTAGGGGAGCAAACGGG
## 2481                                                         ACGTAGGTGGCGAATGTTGTTCGGAATTACTGGGCATAAAGAGTTTGTAGGCTGTATGATAAGTCAGGTGTGAAATCCCGGGGCTCAACTTCGGAATGGCATTTGATACTGTTGTACTAGAATAACGGAGAGGTGAGTAGAATTCATGGTGTAGCGGTGAAATGCGTTGATATCATGGAGAATACCAATGGCGAAGGCAGCTCACTGGCCGATTATTGACGCTGAGAAACGAAAGCTAGGGTAGCGAAAGGG
## 2482                                                         ACGTAGGTGGCAAGCGTTGTCCGGATTTACTGGGCGTAAAGCGCGCGCAGGCGGTGAATGAAGTCGAAAGTGAAAGCCCCGGGCTCAACTCGGGAACAGCTTTCGAAACCAGTTCACTTGAAGGTGGAAGAGGGAAGTGGAATTCCGAGTGTAGCGGTGGAATGCGCAGATATTCGGAGGAACACCAGAGGCGAAGGCGACTTCCTGGTCCAATCTTGACGCTGAGGCGCGAAAGCTAGGGGAGCAAACAGG
## 2484                                                         ACAGGGGGTGCAAGCGTTACTCGGAATCATTGGGCGTAAAGAGCGTGTAGGCGGTTTGTTAAGTCTGGTGTGAAAGCCCTGGGCTCAATCCAGGAAGTGCATTGGATACTGACAGACTAGAGTGTAGGAGAGGATGGTGGAATCCCTGGTGTAGAGGTGAAATTCGTAGATATCAGGGGGAACAACAGTGGCGAAGGCGACCATCTGGACTATTACTGACGCTAAGGCGCGAAAGCGTGGGGAGCAAACAGG
## 2486                                                         ACGTAGGGTGCGAGCGTTGTCCGGAGTTACTGGGCGTAAAGAGCGTGTAGGCGGTTGAAAAAGTCAGGTGTGAAATTCTTCGGCTCAACCGAGGGACTGCGCCTGAAACTTTTCGACTGGAGTTCGGGAGGGGCAACTGGAATTCCTGGTGTAACGGTGAAATGTGTAGATATCAGGAGGAACACCCGTGGCGAAGGCGGGTTGCTGGACCGAAACTGACGCTGAGACGCGAAAGCGTGGGTAGCAAACAGG
## 2488                                                         ACGAAGGGTGCAAACGTTGTTCGGAATGACTGGGCGTAAAGCGTATGTAGGCGGCTTTGCAGGTCGGATGTGAAATCCCTTGGCTCAACCAAGGAAGTGCATTCGAAACCGCAGAGCTTGAGTGCCTGAGAGGAAGGTGGAATTCTCGGTGTAGAGGTGAAATTCGTAGATATCGAGAGGAACACCCGTGGCGAAAGCGGCCTTCTGGAAGAGCACTGACGCTGAGATACGAAAGCATGGGGAGCAAACAGG
## 2489                                                         ACAGAGGGTGCGAGCGTTAATCGGAATTACTGGGCGTAAAGGGTGCGTAGGCGGTCTGATAAGTCGGATGTGAAATCCCTGGGCTTAACCTAGGAATTGCATACGATACTGCCAGACTTGAGTACAGTAGAGGATAGTGGAATTCCACGTGTAGCGGTGAAATGCGTAGATATGTGGAGGAACACCAGTGGCGAAGGCGGCTATCTGGACTGATACTGACGCTGAGGCACGAAAGCGTGGGGAGCAAACAGG
## 2490                                                        ACGGAGGATCCAAGCGTTATCCGGAATGACTGGGCGTAAAGAGTTGCGTAGGTGGTTTATTAAGCGGTGTGTGAAATCTTGTGGCTCAACCATAAGGCTATGCACTGAACTGATAGACTTGAGTATGAGAGAGGTTATTGGAATTCCCAGTGTAGGAGTGAAATCCGTAGATATTGGGAGGAACACCGATGGCGTAGGCAGATAACTGGCTCATTACTGACACTAAGGCACGAAAGCGTGGGGAGCAAACGGG
## 2491                                                         ACAGGTGTTGCAAGCGTTACTCGGATTCACTGGGTGTAAAGGGTCAGCAGGCGGCTTTGTAAGTCTTAGGTGAAAGCCAACGGCTCAACTGTTGAATTGCCTAAGATACTGCATGGCTAGAGTGTGATAGAGGAGAGTGGAACAGCTAGTGTAAAGGTGAAATTTGTGGATATTAGCTGGAACGCCAACGGTGAAGACAGCTCTCTGGATCATAACTGACGCTCAGGGACGAAAGCGTAGGGAGCAAACAGG
## 2497                                                         ACGAAGGGTGCAAGCGTTGCTCGGAGTTATTGGGCGTAAAGGGTGAGTAGGTGGTCTGATTTGTCTGGGGTGAAAGCCTTAAGCTTAACTTAAGAAGTGCCTTAGAAACGGTCGGACTAGAGTGCTGGAGAGGATCGCAGAATTCCTGGTGTAGCGGTGAAATGCGTAGAGATCAGGAGGAATACCTGTGGCGAAAGCGGCGATCTGGACAGTAACTGACACTCAATCACGAAAGCGTGGGGAGCAAACAGG
## 2498                                                        ACGGAGGATCCAAGCGTTATCCGGAATTACTGGGCGTAAAGAGTTGCGTAGGTGGCATTGTAAGTCTGTAGTGAAAGCGTGGGGCTCAACCCCATACACATTATGGAAACTGCAAAGCTAGAGGACAAGAGAGGTTATTGGAATTCCTAGTGTAGGAGTGAAATCCGTAGATATTAGGAGGAACACCGATGGCGTAGGCAGCTTCTTGGTCCATATCTGACGTTGAAGTACGAAAGCGTGGGGAGCAAACAGG
## 2501                                                         ACGTAGGGTGCGAGCGTTAATCGGAATTACTGGGCGTAAAGCGTGCGCAGGCGGTTATACAAGACAGGCGTGAAATCCCCGAGCTTAACTTGGGAACTGCCTTTGATACTATATGTCTTGAATATCGTGGAGGTAAGCGGAATATGTCATGTAGCGGTGAAATGCATAGAGATGACATAGAACACCAATTGCGAAGGCAGCTTGCTACACGATGATTGACACTGAGGCACGAAAGCGTGGGGAGCAAACAGG
## 2502                                                        ACCAGCGCCTCGAGTGGGGATCGCGATTATTGGGTCTAAAGCGTCCGTAGCAGGCTTTGTAAATGCTTTGTGAAATTGTCTGGCTTAACCAGACAGCGTGCAGAGTAGACTGCAGAGCTCGAGACCGGGAGAGGTTAGCAGTATTCCTAGGGTACCGGTAAAATGGGTTAATCCTAGGAGGACTACCAATGGCGAAGGCAGCTAACCAGAACGGTTCTGACTGTGAGGGACGAAAGCTAGGGGAGCAAAACGG
## 2503                                                         ACGTAGGTGACAAGCGTTATCCGGAATTACTGGGCGTAAAGCGCGTGCAGGTGGCTTCGCAAGTCGGGCATGAAATCTCTCGGCCTAACCGAGAGACGCTGTTCGATACTGCGGAGCTTGAGGGCAGTAAAAGATGGCGGAATTCCGCATGTAGTGGTGAAATGCGTAGAGATGCGGAGGAACACCAGTGGCGAGGGCGGCCATCTGGGCTGTCCCTGACACTGAAGCGCGAAAGCTAGGGTAGCGAACGGG
## 2509                                                       ACGGGAGGGGCAAGCGTTAACCGGAATGATTGGGCGTAAGGGGTTCGTAGATGGTATTGTGGATGTTACTGATTAAAGACCAAGGCTTAACCTTGGGAAGGTTAGTAAAACACCTATACTAGAGTTAAGTAGAGGATAGTAGAATTCCGTGTGAAGGGGCGGAATCCATTGAAATACGGAGGAATGCCAAAGGCGAAAGCAACTATCTGGACTTATACTGACGTTGAGGAACGAAAGCGTGGGGAGCAAATAGG
## 2511                                                         ACGAACTGTGCAAACGTTATTCGGAATCACTGGGCTTAAAGGGTGCGTAGGCGGCCATTTAAGTAGGGTGTGAAAGCCCCCGGCTCAACCGGGGAATTGCGTCCTAAACTGGGTGGCTCGAGTGAGATAGAGGTGTGTGGAACTTCCGGTGGAGCGGTGAAATGTGTTGATATCGGAAGGAACGCCGGTGGCGAAAGCGACACACTGGGTCTTAACTGACGCTGAGGCACGAAAGCCAGGGGAGCGAACGGG
## 2515                                                         ACGAAGGATCCAAGCGTTATCCGGATTCATTGGGTTTAAAGGGTGCGTAGGCGGATAGATAAGTCAGTGGTGAAAGCCGGTCGCTCAACGATCGAATTGCCATTGATACTGTTTATCTAGAATATAGATGATGTTGGCGGAATATGACATGTAGCGGTGAAATGCATAGATATGTCATAGAACACCTATTGCGAAGGCAGCTTACTACGCATATATTGACGCTGAGGCACGAAAGCGTGGGGATCAAACAGG
## 2516                                                         ACGTAGGATGCGAGCGTTGTCCGGATTTACTGGGTTTAAAGGGTGCGTAGGCGGTTTATTAAGTCAGTGGTGAAATTTTGTAGCTTAACTATAAGTGTGCCATTGATACTGATAGACTTGAGTTTTTATGAGGTGGGCGGAATTGGTAGTGTAGCGGTGAAATGCATAGATATTACCAGGAACGTCGATAGCGAAGGCAGCCCACTAATAAATAACTGACGCTGAGGCACGAAAGTGCGGGGATCAAACAGG
## 2522                                                         ACAGAGGTCTCAAGCGTTGTTCGGAATCACTGGGCGTAAAGGGTCCGTAGGCGGACTTATAAGTCAGTGGTGAAAGCCTGTCGCTTAACGATAGAACTGCCATTGATACTGTAAGTCTTGAGTATATTTGAGGTAGCTGGAATAAGTAGTGTAGCGGTGAAATGCATAGATATTACTTAGAACACCAATTGCGAAGGCAGGTTACCAAGATATAACTGACGCTGAGGGACGAAAGCGTGGGGAGCGAACAGG
## 2523                                                         ACGAAGGGGGCTAGCGTTGTTCGGAATTACTGGGCGTAAAGCGCACGTAGGCGGACTGTTAAGTCGGGGGTGAAAGCCCAGGGCTCAACCCTGGAACGGCCTTCGATACTGGCAGTCTTGAGAATGGGAGAGGTGAGTGGAATTCCTAGTGTAGAGGTGAAATTCGTAGATATTAGGAAGAACACCAGTGGCGAAGGCGGCTCACTGGCCCATTTCTGACGCTGAGGTGCGAAAGCGTGGGGAGCAAACAGG
## 2525                                                         ACAGAGGATGCAAGCGTTATCCGGAATCACTGGGCATAAAGCGTCTGTAGGCGGCTGATCAAGTCGGGTGTGAAAGCCCGGGGCTTACCCCCGGAAGTGCGCTCGAAACTGGTCAGCTAGAGTATGGGAGGGGGAAGTGGAATTCCTGGTGTAGCGGTGAAATGCGTAGATATCAGGAGGAACACCGGCGGCGAAGGCGGCTTCCTGGACCAATACTGACGCTGAGACGCGAAAGCGTGGGGAGCAAACAGG
## 2527                                                       ACAGACGGGGCAAGTATTTGGTGTCTTGACTGGGTTTAAAACGCAGGTAGGTGGTGAACGTCGAAAGAAGATTCTTGCCAACAACTGTTGGCAAATCTTGCACTTGAAGACGTTGCTAGAGTTTATTCAGAGGTTAAGAGTATGTCTCATGTATGGGTTAAATCAATTTCATATGAGACGGAATGCCTAAGGCGAAGGCGCTTTTCTGCTATCTAAACTGACGCTGAGCTGCGGAAGCGTGGGGATCAAACGGG
## 2530                                                      ACGGGAGGAACAAGTGTTATTCATAGTGATTTGGCGTATAGGGTATCATGGTGGTATTTTAGAAGAATTTCTTGGAAATCCGTCTAGGCAATAGGCGGGGGTAAGATTTCAATTTTTAATACTTGAGTTTATAAAAGATAAGGGTATGTTAAGGGGAGCAATGAAATGCTGTGATACTTTTCGGAACACCATAGGCGAAGGCGCTTATCTAGTTAAAACTGACACTAGCGTACTAAAGCATGGGGATCAAACGGG
## 2531                                                         ACGTAGGCCCCGAGCGTTATCCGGATTTACTGGGCGTAAAGAGTCTGTAGGTGGAAAGGCAAGTCAAACATGAAATGCCGGGGCTCAACCCCGCGCCCGTGTTCGAAACTATCTTTCTTGAACGGCTAAGAGGGAAGCGGAATGGTGGGAGTAGGGGTGCAATCCGTTGACACCCACCAGAACACCAAAAGCGAAGGCAGCTTCCTGGGAGCCTGTTGACACTGAGAGACGAAAGCGTGGGGAGCGAAAAGG
## 2532                                                         ACGAAGGATGCAAGCGTTATCCGGATTTACTGGGTTTAAAGGGTGTGCAGGTGGGATAATAAGTCAGTGGTGAAAGCCCCCGACTCAATCGGGGAACTGCCATCGATACTGTTATTCTTGAGTGTAGTTGAGGCGGGCGGAATATGACATGTAGCGGTGAAATGCGTAGATATGTCATAGAACACCGATCGCGAAGGCAGCTCGCCAAACTATTACTGACACTGAAACACGAAAGTGCGGGGATCAAACAGG
## 2534                                                        ACGAAGGGAGCAAGCGTTATCCGGAATAATTGGGCGTAAAGCGTCTGTAGGTGGTTTAGAAAGTCTAATGTTAAATCCCAGGGCTCAACCTTGGTTAAGCATTGGAGTACTTTTAAACTTGAGTGTGGTAGAGGTAGAGGGAATCCCTAGCGTAACAGTGAAATGTGTAGATTTTAGGGAGAACACCAGCGGCGAAGGCGCTCTACTGGGCCAATACTGACACTGAGAGACGAAAGCTACGGGAGCGAAAAGG
## 2535                                                         ACGGGGGATGCGAGCGTTATCCGGATTTATTGGGTTTAAAGGGTGCGTAGGCTGCGCGACAAGTCGGCGGTGAAACCCCGGGGCTCAACCCCGGGCCTGCCGTCGAAACTGGCGCGCTGGAGTTCGGATGCCGTGGGAGGAATGAGTGGTGTAGCGGTGAAATGCATAGATATCACTCAGAACACCGATTGCGAAGGCATCTCACGAATCCGAGACTGACGCTGAGGCACGAAAGCGTGGGGATCAAACAGG
## 2538                                                         ACGTATGGAGCGAGCGTTACTCGGAATCACTGGGCGTAAAGAGTTCGTAGGCGGATTCGTAAGTCAGAAGTGAAATTTTGGGGCTCAACCCCGAAACTGCTTTTGAAACTGCGTATCTTGAGTATGTGAGAGGAAGATGGAATTCCTGGTGTAGCGGTGGAATGCGTAGATATCAGGAAGAACACCGGTGGCGAAGGCGGTCTTCTGGCACAATACTGACGCTGAGGAACGAAAGCGTGGGGAGCAAACAGG
## 2539                                                         ACAGAGGGTGCAAACGTTGCTCGGATTTACTGGGCGTAAAGCGCATGTAGGCGGGACGCTAAGTCGGGTGTGAAATCCCCGGGCTCAACCTGGGAACTGCCTTTGTGACTGCATAGCTAGAGTACGGCAGAGGGGGATGGAATTCCGCGTGTAGCAGTGAAATGCGTAGATATGCGGAGGAACACCGATGGCGAAGGCAATCCCCTGGGCCTGTACTGACGCTCATGCACGAAAGCGTGGGGAGCAAACAGG
## 2541                                                         ACGGAGGGTGCAAGCGTTGTTCGGATTTACTGGGCGTAAAGGGCGCGTAGGCGGCTTTATAAGCCAGATGTGAAATCCCACGGCTCAACCGTGGAAGTGCATCTGGAACTGTGAGGCTTGAGTATGGGAGAGGAAAGTGGAATTCCCGGTGTAGAGGTGAAATTCGTAGATATCGGGAGGAATACCGGTGGCGAAGGCGACTTTCTGGACCAATACTGACGCTGAGGCGCGAAAGCGTGGGTAGCAAACAGG
## 2549                                                         ACGAAGGGGGCAAGCGTTGCTCGGATTTATTGGGCGTAAAGGGTGTGTAGGTGGTCCTACAAGTTAGATGTGAAAGCCCCGGGCTCAACCTGGGAACTGCATTTGAGACTGTATAGCTAGAGTACGGTAGAGGGGGATGGAATTCCGCGTGTAGCAGTGAAATGCGTAGATATGCGGAGGAACACCGATGGCGAAGGCAATCCCCTGGACCTGTACTGACGCTCATGCACGAAAGCGTGGGGAGCAAACAGG
## 2555                                                        ACGAGAGCCCCGAGCGTTATCCGGAATTATTGGGCGTAAAGGGTGTGTAGGTGGCGTTATTAGTCTTTTGTGAAAGACCCAAGCCTAACTTGGGGAACGCATTGGAAACGGTAATGCTTTGAGAGTGTGAGAGGTGCACGGAACTTATGGTGTAGGGGTGAAATCCGTTGATATCATAGGGAACACCAAATGCGAAGGCAGTGCACTGGCACATATCTGACACTGAGACACGAAAGCGTGGGTAGCGAATGGG
## 2558                                                         ACAGAGGTCCCAAGCGTTGTTCGGATTCACTGGGCGTAAAGGGTGCGTAGGCGGTGGGGTAAGTTTGAGGTGAAATCTCCGGGCCTAACCCGGAAAGTGCGGAGGGGACTGCTCGGCTAGAGGATGGGAGAGGAGCGCGGAATTCCTGGTGTAGCAGTGAAATGCGTAGATATCAGGAGGAACACCGGTGGCGAAGGCGGTTTTCTGGGCCTTTACTGACGCTAAAGTACGAAAGCTAGGGGAGCAAACGGG
## 2560                                                        ACCGGCAGCTCAAGTGGTAGCCATTTTTATTGGGCCTAAAGCGTTCGTAGCCGGTTTGATAAGTCTCTGGTGAAATCCCACAGCTTAACTGTGGGAATTGCTGGAGATACTATTAGACTTGAGGTCGGGAGAGGTTAGCGGTACTCCCAGGGTAGGGGTGAAATCCTGTAATCCTGGGAGGACCACCTGTGGCGAAGGCGGCTAACTGGAACGAACCTGACGGTGAGGGACGAAAGCCAGGGGCGCGAACCGG
## 2568                                                         ACGGAGGATCCAAGCGTTATCCGGATTTACTGGGTTTAAAGGGTGCGTAGGTGGACTTTTAAGTCAGTGGTGAAATCTGATAGCTTAACTATCAAAGTGCCATTGATACTGATTGTCTTGAGTATGGTTGGAGTGAATGGAATGTGTTGTGTAGCGGTGAAATGCTTAGATATAACACAGAACACCAATCGCGAAGGCAGTTCACTAAGCCTTTACTGACACTGATGCACGAAAGCTTGGGGAGCAAACAGG
## 2574                                                         ACGAGAGGGGCGTGCGTTATTCATACGAACTTGGCGTAAAGCGTTTTTAGGTGGTTTAACGTAGTTAGCATCTAAAGCTCAGAACGTAATTCTGGAATGTGTGCTAAGACGGTTATTCTTGAGTTGTTTTAGGGTTAGTAGTATTTCTTCTCGAGGGTTAAAATTCATAGAAAGAAGAAAGAGGGTCTAGGGTGAAGACAACTAACTAAATTCAACTGACACCAAAGAACTACAGCGTAGGGAGCGATCGGG
## 2575                                                          ACGTAGGATCCGAGCGTTATCCGGAATTATTGGGCGTAAAGGAAGCGTAGGTGGTTTAGTAAGTAAGATGCGAAAGCTCTCCGCTCAACGGAGAAACTGTATCTTATACTGCTAAACTAGAGGTTGGTAGAGGTAAGCGGAATTAGTAGTGTAGCGGTGAAATGCGTAGATATTACTAAGAACTCCAATGGCGAAGGCAGCTTACTGGGCCACACCTGACACTGAGGCTTGTAAGCGTGGGTAGCGATCGG
## 2591                                                         ACGTAGGGTGCGAGCGTTAATCGGAATTACTGGGCGTAAAGCGTGCGCAGGCGGTTATGCAAGACAGAGGTGAAATCCCCGGGCTCAACCTGGGAACTGCCTTTGTGACTGCATAGCTAGAGTACGGCAGAGGGGGATGGAATTCCGCGTGTAGCAGTGAAATGCGTAGATATGCGGAGGAACACCGATGGCGAAGGCAATCCCCTGGGCCTGCACTGACGCTCATGCACGAAAGCGTGGGGAGCAAACAGG
## 2593                                                         ACGTAGGATGCAAGCGTTGTTCGGAATTACTGGGCGTAAAGCGTATGTAGGCGGTTATGTAAGTCTTTTATTAAAGACCGAGGCTCAACCTCGGGAGTGTAAAGGAAACTGTATAACTAGAGGTCAGAAGGGGAAATCGGAATTCTTAGTGTAGCGGTGAAATGCGTAGATATTAGGAAGAACACCGGTGGCGAAAGCGGATTTCTGGTCTGAAACTGACGCTGAGATACGAAGGTGTGGGGAGCAAACGGG
## 2596                                                         ACGTAGGGTGCAAGCGTTAATCGGAATTACTGGGCGTAAAGCGTGCGCAGGCGGCTTTGCAAGACAGATGTGAAATCCCCGGGCTCAACCTGGGAACTGCATTTGAGACTGTATAGCTAGAGTACGGTAGAGGGGGATGGAATTCCGCGTGTAGCAGTGAAATGCGTAGATATGCGGAGGAACACCGATGGCGAAGGCAATCCCCTGGGCCTGCACTGACGCTCATGCACGAAAGCGTGGGGAGCAAACAGG
## 2600                                                          CGTAGGTGGCGAGCGTTATCCGGAATGATTGGGCGTAAAGGGTGCGTAGGTGGCAGATCAAGTCTGGAGTAAAAGGTATGGGCTCAACCCGTACTGGCTCTGGAAACTGATCAGCTAGAGAACAGAAGAGGACGGCGGAACTCCATGTGTAGCGGTAAAATGCGTAGATATATGGAAGAACACCGGTGGCGAAGGCGGCCGTCTGGTCTGGATTCTGACACTGAAGCACGAAAGCGTGGGGAGCAAATAGG
## 2601                                                         ACAGAGGGTGCAAGCGTTAATCGGAATTACTGGGCGTAAAGGGCACGTAGGCGGTGGATGAAGTTGAAGGTGAAAGACAGCGGCTTAACCGCTGGAGGTCCTTCAATACTCATTGACTAGAGTTGGGAAGAGGAGAGAGGAATTTCCGGAGTAGCAGTGAAATGCGTAGATATCGGAAGGAACATCAGTGGCGAAGGCGAGTCTCTGGTCCCGAACTGACGCTGAGGCGCGAAAGCGTGGGGAGCAAACAGG
## 2602                                                         ACAGAGGCCTCAAGCGTTGTTCGGAATCACTGGGCGTAAAGGGTGCGTAGGTCGTGTCGCAAGTCAATTTGTGAAATACATGCGCTCAACGCATGTCTGCATTTGATACTGCGAGACTAGAGATATTTTGGGGAGATGGGAATTCACGGTGTAGCAGTGAAATGCGTAGATATCGTGAGGAACACCAAAGGCGTAAGCACATCTCTAAGAATTTTCTGACACTGAGGCACGAAGGCTAGGGTAGCAAATGGG
## 2604                                                                 ACCTGAGTGATTAGCATTATTCGGCGTGATTGGGCGTAAAGGATACGTAGGCGGGGTTTGTAAAACCAACTGGATTCCTAAAAGATGTAAAGTTTTTTTAAATGAACCCTTGAGTGAATATTAAGATGGGGGGTTTTCCTGGGTAACGATAAAATGTAGCGACCCACGACTGCCTGCCAGTGTACTAATGTGCCTGTCTAATATTTTCTGACGCTGAGGTATTAAAGTAGGGGGTGTAAACAGG
## 2605                                                         ACGTAGGGTGCAAGCGTTAATCGGAATTACTGGGCGTAAAGCGTGCGCAGGTGGTTGTATAAGACAGATGTGAAATCCCCGGGCTCAACCTGGGAACTGCATTTGTGACTGTACGGCTAGAGTGTGTCAGAGGGAGGTGGAATTCCACGTGTAGCGGTAAAATGCGTAGATCTCGACAGGAACACCGGTTGCGAAGGCGGCTTCTTGGCTTGTAACTGACGCTAAAGCGCGAAAGCGTGGGGAGCAAACAGG
## 2630                                                         ACAGAGGTGTCAAGCGTTGCTCGGTGTCACTGGGCTTAAAGGGCGCGTAGGCTGTGCGGTACGGTTCGGGTGAAATCCCTCGGCTCAACCGAGGAACTGCCCGGGCGACGGCCGTGCTGGACTGCATCAGGGGCGGATGGAACTCCCGGTGGAGCGGTGAAATGCGTAGATATCGGGAGGAACGCCGGTGGCGAAAGCGATCCGCTGGGGTGCATGTGACGCTGAGGCGCGAAAGCCAGGGGAGCAAACGGG
## 2634                                                        ACGAAGGGGGCGAGCGTTATCCGGAATGATTGGGCGTAAAGAGTTCGTAGGTGGTATTTGTTGTTATCTGTAAAATACTAAATTTATTATTTAGAAATGCATTTAATACTCTTTTACTTGAGTCTAGTAGAGGATAATAGAATTTCATACATAGGAGTGAAATCCGATTATTTATGAAGGAATGTCAGTGGCGAAAGCGATTATCTGGACTAGTTACTGACACTGAGGAACGAAAGCATAGGGAGCAAACAGG
## 2636                                                         ACGTAGGGTGCAAGCGTTACTCGGAATCATTGGGCGTAAAGGGTGTGTAGGCGGAACAGTAAGTCAGATGTGAAAGCCTTGGGCTCAACTCAAGAAGTGCATTTGATACTGCTGTTCTAGAATTCAGGAGAGGAAGGAAGAATTCCCGGTGTAGAGGTGAAATCCGTAGATATCGGGAGGAATACCGGAGGCGAAGGCGTCCTTCTGGACTGACATTGACGCTGAGACACGAAAGCGTGGGTAGCAAACAGG
## 2637                                                         ACAGGAGATGCAAGCGTTATCCGGAATTATTGGGCGTAAAGCGTCTGTAGATGGTTTAAAAAGTTTGGTGTTAAATATCAGAGCTCAACTTTGAAATGGCATCGAAAACTCTTAAACTTGAGTGCAATAGGGGCAAGGGGAATTTTCAGTGTAGCGGTGAAATGCGTAGAGATTGAAAAGAACACCAATGGCGAAAGCACCTTGCTAGGTTGATACTGACATTCAGAGACGAAAGCTAGGGGAGCAAAAAGG
## 2647                                                         ACGTAGGATCCGAGCGTTGTCCGGAGTTACTGGGTGTAAAGGGTGCGTAGGCTGTCCCTTAGGTCAATTGCGAAATTCTGGAGCTCAACTCCAGCGCTGTAATTGATACTATTGGACTAGAATGTGGAAGAGGTGAATGGAATTTTCAGTGTAAGAGTGAAATCTTTAGATATTGAAAGGAACACCAGAGGCGAAGGCGATTCACTGGGACATTATTGACGCTGAGGCACGAAAGCGTGGGGAGCAAACAGG
## 2660                                                         ACGTAGGACCCGAGCGTTGTCCGGATTTACTGGGTGTAAAGGGAGCGCAGGCGGATTATTAAGTCGATTGTGAAATCTTGCGGCTCAACCGCAAAACAGCAGTCGATACTGATGATCTTGAGTGTGGAAGAGAGTGGCGGAATTCATGGTGTAGCAGTGAAATGCGTAGATATCATGAAGAACACCGGTTGCGAAGGCGGCCACTTAGTCCATTACTGACGCTCATGCTCGAAAGCGTGGGGATCAAACAGG
## 2665                                                         ACGGAGGATGCGAGCGTTATCCGGATTTATTGGGTTTAAAGGGTCCGTAGGCGGTAGATTAAGTCAGTGGTGAAATCCTACAGCTTAACTGTAGAATTGCCATTGATACTGGTTTACTTGAATTAGGTTGAAGTAGCTAGAATGTGTAGTGTAGCGGTGAAATGCATAGATATTACACAGAATACCGATAGCGAAGGCAGGTTACTAAGCCTATATTGACGCTGAGGGACGAAAGCGTGGGGAGCAAACAGG
## 2669                                                       ACGAGAGGGGCAAGCGTTATTCGGAATGATTGGGCGTAAAGCGCACGTAGACTGTTTAGACACTTAAAAAGACAAATTTCAAAGCTTCACTTTGACACGTCTTTTCATAAGTACTAAACTGGAGTGAGTAAGAGGAGAATGGAATTCCTGGTGTAGAGGTAAAATCTGTTGATATCAGGAGGAACACCAAAAGCGTAAGCAATTCTCTAGTACTCGACTGACGTTGAGGTGCGAAAGCGTGGGGATCAAAAGGG
## 2675                                                         ACGAAGGGAGCGAGCGTTGTCCGGAATTACTGGGCGTAAAGAGCACGTAGGCGGCTTTATTAGTTGGGTGTGAAATCCCAGGGCTCAACCCTGGAACTGCATCCAATACTGTAAAGCTAGAGACCGAGAGAGGAAAATGGAATTACGAGTGTAGAGGTGAAATTCGTAGATATTCGTAGGAACACCAGAGGCGAAGGCGATTTTCTGGCTCGGTACTGACGCTCAGGTGCGAAAGCGTGGGGAGCAAACAGG
## 2677                                                         ACGTAGGGTGCGAGCGTTAATCGGAATTACTGGGCGTAAAGCGTGCGCAGGCGGTTATATAAGACAGATGTGAAATCCCCGGGCTTAACCTGGGACCTGCATTTGTGACTGTATAGCTAGAGTACGGTAGAGGGGGATGGAATTCCGCGTGTAGCAGTGAAATGCGTAGATATGCGGAGGAACACCGATGGCGAAGGCAATCCCCTGGGCCTGTACTGACGCTCATGCACGAAAGCGTGGGGAGCAAACAGG
## 2696                                                         ACAGAGGGTGCAAGCGTTATCCGGAATCATTGGGCGTAAAGCGTCCGCAGGCCGTTTTTTAAGTCTGGTATTAAATCTTGAAGCCCAACTTCAAGGCCGTACTGGAAACTGGAAAACTAGAGTCTTTCAGAGGTACGTGGAATTCTGTGTGTAGGGGTAAAATCCGTAGATACACAGAGGAACACCAAAAGCGAAGGCAGCGTACTGGGAAAGTACTGACGCTCATGGACGAAAGCGTGGGGAGCGAACAGG
## 2704                                                         ACGGGAGTGGCAAGCGTTATCCGGAATTATTGGGCGTAAAGCGTCCGCAGGCGGTCTTACAAGTCTGTCGTTAAAGCGTGGAGCTTAACTCCATTTCGGCGATGGAAACTGTAAGACTAGAGTGTGGTAGGGGCAGAGGGAATTCCCGGTGTAGCGGTGAAATGCGTAGATATCGGGAAGAACACCAGTGGCGAAGGCGCTCTGCTGGGCCATAACTGACGCTCATGGACGAAAGCGTGGGGAGCAAACAGG
## 2706                                                         ACGGAGGGTGCGAGCGTTGTCCGGATTTATTGGGTTTAAAGGGTGCGTAGGTGGTATGGTAAGTCTGGTTTGAAAGCTGGTCGCTTAACGATCAGATGTGGCTGGAAACTGCGGTACTTGAATGCGGTGGCGGTAGCTGGAACGGGTCATGTAGCGGTGAAATGCATAGATATGACCCAGAACCCCGATTGCGAAGGCAGGCTACTACGCCGTGATTGACACTGAGGCACGAGAGCATGGGTAGCGAACAGG
## 2709                                                         ACGAGGGGAGCAAGCGTTATACAGTCAAACTGGGCGTAAAGGGTCTTTCAGTGAAAAAATGTAAGAAAATAATGAAAAATTAAAAGGTAACTTTAATGGTTTATTTTTACTATTTTTTTAGAATCTTATTAGGGTTAACAGTATTTCAACAAAAGAGTTAAAATGCTTAGACTGTTGAAGGAAAGCCAAGGGCGAAAGCAGTTAACTGTATAAGATTGACACTTAGAGACAAAAGCATGGGGAGCAATCGGG
## 2712                                                         ACGTAGGATCCGAGCGTTGTCCGGAGTTACTGGGTGTAAAGGGTGCGTAGGCTGTTCTGTGCGTCAGAGGTGAAATCCACGGGCTTAACTCGTGGGGTGCCTTTGATACGGCAGGGCTTGAGTACGAGAGAGGTGCGTGGAATTCCTGGTGTAGCAGTGAAATGCGTAGATATCAGGAGGAACACCGGTGGCGAAGGCGGCGCACTGGCTCGTAACTGACGCTCATGTGCGAAAGCTAGGGCAGCGAACGGG
## 2716                                                         ACGAGGGGGGCAAATGTTGTTCGGAATCACTGGGCGTAAAGGGAGCGTAGGCGGGTTCTTAAGTAAGGTGTTTAATCCCAGGGCCCAACCCTGGACCGGCACTTTAAACTGGGAATCTTGAATAGTGGAGAGGTCAGTGGAATTGGAGGTGTAGCGGTGGAATGCGTAGAGATCTCCAAGAACACCAGTTGCGAAGGCGGCTGACTGGACACATATTGACGCTGAGGCTCGAAAGCATGGGGAGCAAACAGG
## 2719                                                         ACGGGGGGAACGAGCGTTGTTCGGATTTACTGGGCGTAAAGGGCACGCAGGCGGGGAGACAAGTCAGATGTGAAAACCCAAAGCTTAACTCTGGGCCTGCATTTGATACTGTTTCTCTTGAGTACGGGAGAGGATAGCGGAATTCTTGGTGTAGCGGTGAAATGCGTAGATATCAAGAAGAACACCGGTGGCGAAGGCGGCTATCTGGCCCGATACTGACGCTCATGTGCGAAGGCAAGGGGAGCAAACAGG
## 2723                                                         ACGTAGGCCCCGAGCGTTATCCGGAATTATTGGGCGTAAAGAGTTCGTAGGCGGGCCATTAGGTCCAGCATGAAAGACCGGGGCTTAACCCCGTGTTTGTGCCGGATACCGGTGGTCTTGAATCCGGGAGAGGTAAGCAGAACTGTATGAGTAGGGGTGCAATCCGTTGATACATACAAGAATACCAAAAGCGAAGGCAGCTTACTGGAACGGCATTGACGCTGAGGAACGAAAGCGTGGGGAGCGAAAAGG
## 2727                                                         ACGAAGGGTGCAAACGTTGTCCGGAATTACTGGGCGTAAAGAGCGCGTAGGTGGTTTGGAAAGTCGTTCATGAAAGACTAGGGCTTAACCCTGGGACCGTGGGCGAAACTTCCATACTTGAGTATGGCAGGGGCTAGTAGAACACTCGGTGGAGGGGTGACATCCGTAGATATCGAGTGGAATACCAAAGGCGAAGGCAGCTAGCTAGGCCATTACTGACACTGAGGCGCGAAAGCGTGGGGAGCGATCGGG
## 2736                                                         ACCCGGGGCCCGAGTGATGGTCGCGATTATTTGGTTTAAAGCATTCGTAGGCTGTTTGATAAGTCTCTTGTGAAATCCTGTAGCAAACTACAGAAAGTGCAAGAGATACTATCAGACTTGAGACTGGGAGAGGACAACGATACTTCTTGTGTAGGGGTTAAATCCTTTAATCCAAGAAGGATCACCTGTGGCGAAGGCGGTTGTCTAGAACAGGTCTGACGCTGAGGGATGAAAGCTAGGGGAGCGATCGGG
## 2740                                                         ACGTAGGTGGCAAGCGTTGTCCGGATTTACTGGGTTTAAAGGGTGCGTAGGCGGGATTATAAGTCAGTGGTGAAATTCGGTTGCTTAACGATACGAGTGCCATTGATACTGTAGTTCTAGAGTAAAGTTAAGGTATGTGGAACGACTGGTGTAGCGGTGAAATGCATAGATATCAGTCAGAACACCGATTGCGAAGGCAGCGTACTGAGCTTTTACTGACGCTGAGGCACGAAAGTGCGGGTATCAAACAGG
## 2744                                                        CCCGACGCTCCAAGTCGCAGCCACAATTATTGGGTCTAAAACATTCGTAGCTTGTTCTGTAAGTTCTTTGTGAAATTCTAAGTCTTAAATTTAGAGCGTGCAAGGAATACTGCAGAACTTGAGACCGGAAGAAGTAAGAAGTATGACGAAGGTAGCGGTAAAATGTGTTAATCTTTGTCAGACTAACAACAGCGAAGGCATCTTACTAGTACGGATCTGACAGTGAGGAATGAAGGCTAGGGTCGCGAAACGG
## 2753                                                        ACGAGAGGTGCAAGCGTTATCCGGAATTACTGGGCGTAAAGCGTCCGCAGGTTGATAGGAAAGTCTACGAGTTAAATTATAAAGCTCAACTTTATAGCTGCTTGGGAAACTATCTATCTAGAGGGTGGTAGAGGTCTGTGGAATTGTACAAGTAGGGGTAAAATCCGTAGATATGTACAGGAACACCAAAAGCGAAGGCAGCAGACTGGACCAACCCTGACACTCATGGACGAAAGCGTGGGGAGCGAAACGG
## 2754                                                         ACAGAGGGTGCAAGCGTTAATCGGAATTACTGGGCGTAAAGGGCGCGTAGGTTGCTATGTAAGTTGGATGTGAAATGTTCGGGCTTAACCTGAAACGGCCATCCAATACTGCATAGCTAGAGTACAGTAGAGGGAAGGGGAATTTCCGGTGTAGCGGTGAAATGCGTAGATATCGGAAGGAACATCAGTGGCGAAGGCGCCTTCCTGGACTGATACTGACACTGAGGCGCGACAGCGTGGGGAGCGAACAGG
## 2767                                                         ACGTAGGGTGCAAGCGTTGTCCGGAATTATTGGGCGTAAAGAGCTCGTAGGCGGTTTGTCACGTCTGCTGTGAAAATCCGAGGCTCAACCTCGGACCTGCAGTGGGTACGGGCAGACTAGAGTGCGGTAGGGGAGAAGGGAATTCCTGGTGTAGCGGTGGAATGCGCAGATATCAGGAGGAACACCAATGGCGAAGGCACTTCTCTGGGCCGCTACTGACGCTGAGGAGCGAAAGCGTGGGGAGCGAACAGG
## 2770                                                         ACGTAGGGTGCAAGCGTTGCTCGGAATCATTGGGCGTAAAGAGTAGGTAGGTGGAATAGTAAGTCAGACGTGAAAGCCTGGGGCTCAACCTCAGAAGTGCGTTTGATACTGCCATTCTAGATTTCGGAAGAGGGGAACAGAATTCCTAGTGTAGGAGTGAAATCCGTAGATATTAGGAGGAATACCAGAGGCGAAGGCGGTTCCCTGGGCCGATAATGACACTGTGCTACGAAAGCGTGGGTAGCAAACAGG
## 2775                                                         ACGAACCGTGCGAACGTTGTTCGGATTCACTGGGCTTAAAGGGCGCGTAGGCGGGCGTCCAAGTCAGGGGTGAAATCCTCCGGCTCAACCGGAGAACAGCTTCTGATACTGGATGTCTCGAGGGAGGTAGGGGCAGATGGAACTTCCGGTGGAGCGGTGAAGTGCGTAGATATCGGAAGGAACGCCGGTGGCGAAAGCGATCTGCTGGACCTCTTCTGACGCTGAGGCGCGAAAGCCAGGGGAGCAAACGGG
## 2778                                                        CCCAGCGCCACAAGTGGGAACCACGTTTATTGGGTCTAAAGCATCCGTAGTCGGTTTGGCAAATGCCCTGTGAAATCGCATCGCTCAACGGTGCGGCGTGCAGGGTAGACTACCAAACTAGGGACCAGGAAAGGAAAGCGGTATTTCTGAGGGAGCGGTAAAATGTTATAATCTCAGAAGGACCACCAATGGCGAAGGCAGCTTTCTAGAATGGATCCGACGATGAGGGATGAAAGCTAGGGGAGCGATCCGG
## 2790                                                         ACAGAGGTGGCAAGCGTTGTTCGGATTTACTGGGCGTAAAGGGTGCGTAGGTGGTATTGTATGTCGGATGTGAAATCTCACCGCTCAACGGTGAAATGGCATTCGAAACTGCGATACTTGAGTGCAGGAGAGGAGAGCGGAATTCTTGGTGTAGCGGTGGAATGCGTTGAGATCAAGAAGAACACCGGTGGCGTAGGCGGCTCTCTGGTATGCAACTGACACTGAGGCACGAAAGTTGGGGGAGCAAACAGG
## 2796                                                        ACGGGGGTCCCAAACGTTGTCCGGTCTTATTGGGCGTAAAGAGTTCTGTAGGCGGTAGTGTAAGTTACGGGTTAAATCCTTCGGCTCAACCGGAGAACTGCTTGTAATACTGCATTACTAGAAATATACAGGGGAGGGCGGAACGCATGGTGTAGCGGTAAAATGCGTTGATATCATGCGGAACACCGAAGGCGTAGGCAGCCCTCTGGGTATTATTTGACGCTAATAGACGAAAGCGTGGGGAGCAAAACGG
## 2801                                                         ACAGAGGGCTCGAGCGTTAATCGGAATCACTGGGCTTAAAGCGTGCGTAGGCGGGTGTGCAGGCACCTTGTGAAATCCCCGGGCTCAACCTGGGAACTGCCTTTGTGACTGCATAGCTAGAGTACGGCAGAGGGGGATGGAATTCCGCGTGTAGCAGTGAAATGCGTAGATATGCGGAGGAACACCGATGGCGAAGGCAATCCCCTGGGCCTGTACTGACGCTCATGCACGAAAGCGTGGGGAGCAAACAGG
## 2802                                                          ACGAGGGGGGCGAGCGTTATTCGGAATCACTGGGCGTAAAGCGTGCGTAGGCGGTTATGAAAGTTAGGAGTGAAATCCCAGGGCTTAACCTTGGAATTGCTTTTAAAACTACATGGCTCGAATTCGATAGAGGATGGCGGAATTATCAGTGTAGCAGTGAAATGCGTAGATATTGATAGGAACACCGGTGGCGTAGGCGGCCATCTAGATCGACATTGACGCTGAAGTACGAAAGCGTGGGTAGCAAACAG
## 2812                                                         ACGTAGGATCCGAGCGTTGTCCGGAGTTACTGGGTGTAAAGGGTGCGTAGGCTGTTCTGTGCGTCAGAGGTGAAATCCCCGGGCTCAACCTGGGAACTGCCTTTGTGACTGTCAGGCTAGAGTGTGTCAGAGGGGGGTAGAATTCCACGTGTAGCAGTGAAATGCGTAGAGATGTGGAGGAATACCGATGGCGAAGGCAGCCCCCTGGGATAACACTGACGCTCATGCACGAAAGCGTGGGGAGCAAACAGG
## 2816                                                         ACAGAGACCTCAAGCGTTATCCGGATTTATTGGGCGTAAAGCGTCCGCAGGTTCCTTGGTAAGTCCTTGGTCAAAACTCAGAGCTCAACTCTGAACCGGCTGAGGATACTGTCGAGGTAGAGAATGGGAGAGGCAAGCGGAATTCTTGGTGTAGCAGTAAAATGCGTTGATATCAAGAAGAACACCAAATGCGAAGGCAGCTTGCTGGAACATTTCTGACACTCATGGACGAAAGCGTGGGGAGCAAACAGG
## 2822                                                         ACGGAGGGTGCAAGCGTTGTTCGGAATTATTGGGCGTAAAGCGCGTGTAGGCGGCTATCTAAGTCTGGTGTGAAATCCCTCGGCTTAACCGAGGAAGTGCATTGGATACTGAATAGCTAGAGGATGAGAGAGGAGAGTAGAATTCCCGGTGTAGGGGTGAAATCCGTAGAGATCGGGAGGAATACCAGAGGCGAAGGCGGCTCTCTGGCTCATATCTGACGCTGAGACGCGAAAGCGTGGGGAGCAAACAGG
## 2827                                                         ACGAAGGGGACTAGCGTTGTTCGGAATGACTGGGCGTAAAGGGCGCGTAGGCGGTTTTGTTAGTTTGTTGTGAAAGCCTTGGGCTTAACCTAAGATGTGCAATGAATACTGCAAGACTAGAGTATGGAAGGGGAGAACGGAATTGCGTGTGGAGAGGTGAAATTCGTAGATATACGCAGGAACACCGGCGGTGAAGACGGTTCTCTGGTACAATACTGACGCTAAGGCGCGAAAGCGTGGGGAGCAAACAGG
## 2839                                                         ACGTATGTCCCGAGCGTTATCCGGATTTATTGGGCGTAAAGCGAGCGCAGACGGTTGATTAAGTCTGATGTGAAAGCCCGGAGCTCAACTCCGGAATGGCATTGGAAACTGGTTAACTTGAGTGCAGTAGAGGTAAGTGGAACTCCATGTGTAGCGGTGGAATGCGTAGATATATGGAAGAACACCAGTGGCGAAGGCGGCTTACTGGACTGTAACTGACGTTGAGGCTCGAAAGTGTGGGTAGCAAACAGG
## 2842                                                        ACGAGAGCTCCAAGCGTTATCCGGAATTACTGGGCGTAAAGCGTATGTAGGCGGCGATGTTAGTCTTCCGTGAAATTCCTCGGCTTAACCGGGGCGCTGCGGAGGAAACGGCATTGCTTCGAATCCGTAAGAGGTCTGTGGAACTCATCATGTAGGGGTGAAATCCGTTGATATGATGGGGAACACCAAAGGCGAAGGCAACAGGCTGGTACGGTATTGACGCTGAAATACGAAAGCGTGGGTAGCGAATGGG
## 2846                                                         ACGGAGGTGGCGAGCGTTGTTCGGAATTACTGGGCTTAAAGGGAGCGTAGGCGGTATGCTAAGTCGCAGGTGGAATCCCGGGGCTCAACCCCGGTATGTCCTGCGAAACTGGTGTACTTGAGTTGTTCAGGGGATGGCGGAATTCCTGGTGTAGCGGTGAAATGCGTAGATATCAGGAGGAAGGCCTGTGCCGAAGGGGGCCATCTGGGGACAAACTGACGCTGAGGCTCGAAAGCGTGGGGAGCAAACAGG
## 2861                                                         ACGTAGGGTGCGAGCGTTAATCGGAATTACTGGGCGTAAAGCGTGCGCAGGCGGTTTTGTAAGACAGATGTGAAATCCCCGGGCTCAACCTGGGACCTGCATTTGTGACTGCAAGGCTGGAGTACGGCAGAGGGGGATGGAATTCCGCGTGTAGCAGTGAAATGCGTAGATATGCGGAGGAACACCGATGGCGAAGGCAGGTCTCTGGGCCAATACTGACACTGAGGAGCGAAAGCGTGGGGAGCGAACAGG
## 2864                                                      ACGGGGGGGGCAAGCGTTACTCTTCGTTACTGGGTGTAAAGCGTAGTAAGGTGGTTTAATAAGTTTTTAGTGAAATCTTAAAATTTTGTTTTAAAAATATTAATAAATACTGTTAAACTTAGAGTATATTAGAAAATAGTTGTATGCTTAGTGTAGGGTTAGAATCCGTAAATATTAAGAAGACATTCAATAGGTGAAAACTACTATTTATAATATTACTGACACTGATATACAAAAGCGTGAGGAGCAAACAGG
## 2869                                                        ACGTAGGGTGCAAGCGTTAATCGGAATTACTGGGCGTAAAGCGTGCGCAGGCGGTTTTATAAGTCGGGTGCGAAATCTTGCGGCTCAACCGTATAGACTGTGCTCGAAACTATGAAGCTAGAGGTAGGTAGAGGCAAGTGGAATTTCTGGTGTAGGAGTGACATCCGTAGATATCAGAAGGAACACCAATGGCGAAGGCAGCTTGCTGGGCCTTACCTGACGCTCAGAGACGAAAGCGTGGGGAGCGAACGGG
## 2870                                                         ACAGAGGGTGCGAGCGTTAATCGGAGTGACTGGGCGTAAAGGGCACGTAGGCGGTAAGATTAGTCTGATGTGAAAGACCTGGGCTTAACCTGGGATGTGCATTGGAAACGGTTTTGCTAGAGTTTGGGAGAGGGAAGTGGAATTTCCGGTGTAGCGGTGAAATGCGTAGAGATCGGAAAGAACACTCGTGGCGAAGGCGGCTTTCTGGCCCAAAACTGACGCTGAGGTGCGAAAGCGTGGGGAGCAAACAGG
## 2873                                                      ACGGAGGGGGTAAGTGTTACTCATCAAAACTGGGCGTAAAGGGTATGTAGGCGGTTTATAATGTGGAAATAAAAGTCTAAAGAACAACTTTATATAATTATTTTTCAGACATATAAACTAAGAGTATAATAGAAGTGTGTTGCACTTTCAGTGGAGAGGTAAAATACGTAAATATTGTAGAGACAATCAATAGGTGAAAACGACTAACTATATTATAACTGACGCTGAGATACAAAAGCATGGGTATCAAACAGG
## 2875                                                         ACGAAGGGTGCTAGCGTTGTTCGGAATAACTGGGCGTAAAGGGCGTGTAGGCGGTCTTATGCGTCAGATGTGAAAGTCCTGGGCTCAACCTGGGGACGGCATTTGATACGGTAAGACTAGAGGTTATGAGAGGAAAGCGGAATTGTGCGTGTAGGAGTGAAATCCGTAGAGATGCACAGGAACACCAGTTGCGAAGGCGGCTTTCTGGTATTAACCTGACGCTAAGGCGCGAAAGCGTGGGGAGCAAACAGG
## 2878                                                         ACGAAGGTGGCAAGCGTTACTCGGAATTACTAGGCGTAAAGCGTGGGCAGGTGGTTCGTAAAGTCTGTCGTGAAAGCTCCCGGCTCAACCGGGAGAGGCCGATGGATACTCACGGACTTGAATCGGAGAGGGGTGACTGGAATTCCCGGTGTAGCGGTGAAATGCGTAGATATCGGGAGGAACGCCAAAGGCGAAAGCAGGTCACTGGATCCGTATTGACACTCATCCACGAAAGCTGGGGGATCAAACAGG
## 2890                                                         ACGAAGGGTGCAAGCGTTGTTCGGATTTATTGGGCGTAAAGGGCGCGTAGGCGGGTCATTAAGTCAGGTGTGAAATCTCGGGGCTCAACCCCGAAATTGCGCCTGAAACTGATAATCTAGAATGTCAAAAGGAGAAAGGGAATTTCGCATGTAGGAGTAAAATCCGTAGAGATGCGAAGGAACACCAGAGGCGAAGGCGCTTTCTTGGTTGACTATTGACGCTGATGCGCGAAAGCGTGGGGAGCAAACAGG
## 2899                                                         ACGGAGGGTGCGAGCGTTATCCGGAATCACTGGGTTTAAAGGGTGCGTAGGCGGTTTATTAAGTCAGTAGTGAAAGACTGTCGCTTAACGACGGATGTGCTAATGATACTGGTAGACTTGAAAGAGGTTAAGGTTGTCGGAATGAGTCATGTAGCGGTGAAATGCATAGATATGACTTGGAACACCGATAGCGAAGGCAGGCAACTGAGCCTTATTTGACGCTGAGGCACGAAAGTGTGGGTAGCGAACAGG
## 2903                                                         ACGTAGGGTGCAAGCGTTAATCGGAATTACTGGGCGTAAAGCGTGCGCAGGCGGTTTCGTAAGACAGAGGTGAAATCCCCGGGCTCAACCTGGGAACTGCCTTTGTGACTGCGAGGCTAGAGTATGGCAGAGGGGGGTGGAATTCCACGTGTAGCAGTGAAATGCGTAGAGATGTGGAGGAACACCGATGGCGAAGGCAATCCCCTGGACCTGTACTGACGCTCATGCACGAAAGCGTGGGGAGCAAACAGG
## 2921                                                          CGGAGGGTGCGAGCGTTGTCCGGAATCACTGGGCGTAAAGCGCGCGTAGGCGGCTTTGTCAATCAGGGGTGAAATCCCGGGGCTCAACCCCGGAACTGCCTTTGATACTGCAAGGCTTGAGTCCGTGAGAGGATGGTGGAATACCCAGTGTAGAGGTGAAATTCGTAGATATTGGGTGGAACACCGGTGGCGAAGGCGACCATCTGGCACGGTACTGACGCTGAGGTGCGAAAGCGTGGGGAGCAAACAGG
## 2922                                                         ACGAAGGGTGCAAGCGTTACTCGGAATTACTGGGCGTAAAGCGTGCGTAGGTGGTGGGTTAAGTCGGATGTGAAAGCCCTGGGCTCAACCTGGGAATGGCATTCGATACTGGCCGACTTGAGTTCGAGAGAGGGTGGTGGAATTCCCGGTGTAGCGGTGAAATGCGTAGAGATCGGGAGGAACATCAGTGGCGAAGGCGACCGCCTGGCTCAGAACTGACACTAAGGCACGAAAGCGTGGGGAGCAAACAGG
## 2926                                                         ACGAAGGTGGCAAGCGTTGTTCGGATTTACTGGGCGTACAGGGAGCGTAGGCGGTTGGGTAAGCCCTCCGTGAAATCTCCGGGCCTAACCCGGAAAGTGCGGAGGGGACTGCTCGGCTAGAGGATGGGAGAGGAGCGCGGAATTCCCGGTGTAGCGGTGAAATGCGTAGAGATCGGGAGGAAGGCCGGTGGCGAAGGCGCCTTCCTGGTCTAAGACTGACGCTGAGGTGCGAAAGCGTGGGGAGCAAACGGG
## 2928                                                         ACGTAGGGTGCGAGCGTTAATCGGAATTACTGGGCGTAAAGCGTGCGCAGGCGGTTATGTAAGACAGATGTGAAATCCCCGGGCTCAACCTGGGACCTGCATTTGTGACTGCATAGCTAGAGTACGGTAGAGGGGGATGGAATTCCGCGTGTAGCAGTGAAATGCGTAGATATGCGGAGGAACACCGATGGCGAAGGCAATCCCCTGGGCCTGCACTGACGCTCATGCACGAAAGCGTGGGGAGCAAACAGG
## 2937                                                         CCGGCGGTCCAAGTCGCAGCCACAATTATTGGGTCTAAAACATCCGTAGCTTGTTTATTAAGTTTCTTGTGAAATCTTATCTCCCAAGGATAAGGCGTGCAGGAAATACTAATAAACTTGAGACCGGAAGAAGTAAGAAGTACAATTGAGGTAGTGGTAAAATATGTTAATCTCAATTGGACTAACAATAGCGAAGGCATCTTACTAGTACGGATCTGACAGTGAGGGATGAAGGCTAGGGTCGCAAAACGG
## 2941                                                  ACGGGGGGAACCGGCGTTATTCGTAAGAATTTGGCGTAAAGGGTACGTAGACATGCCTGAGAACATTTTAGTCGAAAAGCCACGTCCTAAAACGTGGAATAGACATTATAACCTCGGGGATAGAGTGGTGGCCGCGTTACCGGTATTCTCGGTGTAAATCGGGGTCAAATCCAACCATACTTAGAGGAATGCCTAAGGTGAAGACGGGTTTCTGGGCTGCCACTGACGTTCAGGTACGGAGGTGTGAGTATCAAACGGG
## 2950                                                         ACGGAGGGGGCTAGCGTTGTTCGGAATTACTGGGCGTAAAGGGCGCGTAGGCGGTTATGTGTGTCAGATGTGAAATCCCAAGGCTTAACCTTGGAACTGCATTTGAAACTGCATGACTAGAGTGAGTAAGAGGCGGACGGAATTCCGAGTGTAGCAGTGAAATGCGCAGATATTCGGAGGAACATCGGTGGCGAAGGCGGTTCGCTGGTACTCTACTGACGCTGAGGCGCGAAAGCGTGGGGAGCAAACAGG
## 2961                                                         ACAGAGGGTGCAAGCGTTGTTCGGAATCATTGGGCGTAAAGGGCGCGTAGGCGGTTTAGTAAGTCAGATGTGAAATCCCTGAGCCTAACTCAGGACGTGCATTTGAAACTGCTAGACTCGAGTGCGGTAGAGGTTGGTGGAATTCCCAGTGTAGAGGTGAAATTCGTAGATATTGGGAGGAACACCGATGGCGAAGGCAGCCAACTGGGCCGCAACTGACGCTGAGGCGCGAAAGCGTGGGGAGCAAACAGG
## 2963                                                         ACGGGAGTGGCAAGCGTTATCCGGAATTATTGGGCGTAAAGCGTCCGCAGGCGGTCTTGTAAGTCTGTTGTTAAAGCGTGGAGCTTAACTCCATTTCAGCAATGGAAACTGTAAGACTAGAGTGTGGTAGGGGCAGAGGGAATTCCCGGTGTAGCGGTGAAATGCGTAGATATCGGGAAGAACACCAGTGGCGAAGGCGCTCTGCTGGGCCATAACTGACGCTCATGGACGAAAGCGTGGGGAGCAAACAGG
## 2970                                                         ACGGGAGTGGCAAGCGTTATCCGGAATTATTGGGCGTAAAGCGTCCGCAGGCGGTCTTGTAAGTCTGTCGTTAAAGCGTGGAGCTTAACTCCATTTCAGCGATGGAAACTGCAAGACTAGAGTGTGGTAGGGGCAGAGGGAATTCCCGGTGTAGCGGTGAAATGCGTAGATATCGGGAAGAACACCAGTGGCGAAGGCGCTCTGCTGGGCCATAACTGACGCTCATGGACGAAAGCGTGGGGAGCAAACAGG
## 2981                                                         ACGAGGGGAACGAGCGTTATACAGTCAAACTGGGCGTAAAGGGTTCTAAGGTGGAAAATTGGACGTAGAAGATCAAAACTTTAAGAGTAATCTAAAGATATCTTTTAAACTAATTTTCTTGAATAAGATAGGGGTATGCTTTAGTTCATCCAGAGAGTTAAAATGCGTAGACGGATGATCGAGGGCCATTGGCGAAAGCCGCATACCAGGTTTTATTGACACTTAGGGACAAAAGCATGGGGAGCGATCGGG
## 2982                                                         ACAGAGGTCCCAAGCGTTAATCGGAATTACTGGGCTTAAAGGGTCCGCAGGCGGCTCTGCAAGTATCTTGTGAAATACCGTGCCTCAATCACGGAACTGCTTGGTAAACTGCAGAGCTTGAACATAACAGAGGTAGTCGGAACTGTAGGTGGAGCGGTGAAATGCGTAGATATCTACAGGAACGCCGAAGGTGAAGACAGGCTACTGGGTTATAGTTGACGCTCATGGACGAAAGCGTGGGTAGCGAACGGG
## 2983                                                         ACGAGTGGTCCAAGCGTTATCCGGAATTATTGGGCGTAAAGCGTGCGTAGGTGGATTTGTTAGTCGGGTGTTTAAACCATGGGCTCAACCTGTGGATTGCATTCGAAACGTCAAATCTTGAAGATGTGAGGGGTGAATGGAACTATATGTGTAGGGGTGAAATCCGTTGATATATATGGGAACACCAAAAGCGAAGGCAATTCACTGGCACATTCTTGACACTGAGGCACGAAAGCGTGGGTAGCGAATGGG
## 2992                                                         ACGAGGGGTGCGAGCGTTGTTCGGAATCACTGGGCGTAAAGGGCGCGTAGGCGGGCCGTTAAGTCGGGGGTTTAATTTCGGGGCCCAACCCCGGAGCCGCCCCCGATACTGACGGCCTTAGGTAGGTCAGGGGCTGTGGGAATATCTGGTGTAGCGGTGGAATGCGTAGATATCAGATAGAACACCAGTGGCGAAGGCGCATAGCTGGGGCCATACCGACGCTGAGGCGCGAAAGCGTGGGGAGCGAACAGG
## 2999                                                         ACGGAGGTGGCAAGCGTTGTTCGGAATTACTGGGCGTAAAGCGCGCGTAGGCGGTTAATCAAGTCGGGAGTGAAAGCCACGGGCTCAACCTGTGAACTGCTCTCGAAACTGGTTGACTGGAGTCCGGCAGAGGATGGCGGAATTCCCCGTGTAGAGGTGAAATTCGTAGATATGGGGAGGAACACCAGAGGCGAAGGCGGCCATCTGGACCGGTACTGACGCTGAGGCGCGAAAGCGTGGGGAGCGAACAGG
## 3000                                                         ACATAGGTGGCGAGCGTTGTCCGGAATGACTGGGCGTAAAGAGAGTGTAGGCGGTAGTGCAAGTTATATGTGAAAGCCCTTGGCTCAACTGAGGAATAGCATATGATACTGTAAAACTAGAGTACGAGAGAGGGAGATGGAATTTCCAGTGTAGCGGTTAAATGCTTAGATATTGGAAAGAACACCAGAGGCGAAGGCGGTCTCCTGGCTCGATACTGACGCTGAGACTCGAAAGCGTGGGGAGCAAACAGG
## 3005                                                         ACGTAGGGTGCGAGCGTTAATCGGAATTACTGGGCGTAAAGAATTTGTAGGTTGTTGTGTAAGTTAGTTGTGAAATCCCTGGGCTTAACCTAGGAATTGCGACTAAGACTACACGGCTAGAGTATGTAAGAGGCAAGTGGAATTCCTAGTGTAGCAGTGAAATGCGTAGATATTAGGAGGAACACCGGTGGCGAAGGCGACTTGCTGGGACAATACTGACACTGAGAAATGAAAGCGTGGGGAGCAAACAGG
## 3007                                                         ACAGAGGGTGCGAGCGTTAATCGGAGTTACTGGGCGTAAAGGGCGCGTAGGCGGTTTAATGGGTGTGATGTGAAAGCCCCCGGCTCAACCGGGGAATTGCGTCCTAAACTGGGTGGCTCGAGTGAGATAGAGGTGTGTGGAACTTCCGGTGGAGCGGTGAAATGTGTTGATATCGGAAGGAACGCCGGTGGCGAAAGCGACACACTGGGTCTTAACTGACGCTGAGGCACGAAAGCCAGGGGAGCGAACGGG
## 3010                                                         ACGAAGGATCCAAGCGTTGTCCGGATTTACTGGGTTTAAAGGGTGCGTAGGCGGTTGATTAAGTCAGTGGTGAAAGCCTTCAGCTTAACTGAAGAATTGCCATTGAAACTGATCAACTCGAGTATGGTTGAGGTTACCGGAATGTAACATGTAGCGGTGAAATGCTTAGATATGTTACAGAACACCGATTGCGAAGGCAGGTAACTAAGCCATAACTGACGCTGAGGCACGAAAGCGTGGGGAGCAAACAGG
## 3014                                                        ACGGAGGATGCAAGCGTTATCCGGATTTATTGGGTTTAAAGGGTGCGCAGGCGGGCCAATAAGTCAGTGGTGAAATCCTGCCGCTTAACGGTAGGACTGCCATTGATACTGTTGCGCCTTGAGTATGGTTGAGGTAGGCGGAATGTGCAGTGTAGCGGTGAAATGCATAGATATTGCACAGAACGCCGATTGCGGAGGCAGCTTACTAAGCCATCACTGACGCTGAGGCACGAAAGCGTGGGTAGCGAACAGG
## 3018                                                      ACAGAGACCTCAAGCGTTATCCGGAATCATTGGGCGTAAAGCGTACCGATAGGTGGCTTTACAAGTCAGAAGTGAAATCCTAAGGCTTAACTTTGGGACTGTCTTTTGAAACTGTAAAGCTAGAGGGGCAAAGAGGAAGCTGGAACGAACGGTGTAGTAGTGAAATGCGTTGATATCGTTCGGAACACCAATAGCGTAGGCAGCAGACTGGAGCACTCCTGACACTGAAACACGAAAGCGTGGGTAGCGAATGGG
## 3021                                                         ACGGAGGGGGCTAGCGTTGTTCGGAATTACTGGGCGTAAAGAGCGCGTAGGCGGATTAGTAAGTTGGGAGTGAAATCCCGGGGCTTAACCTCGGAACTGCTTCCAAAACTGCTAGTCTAGAGTATAGTAGGGGGTGATGGAATTTCTAGTGTAGAGGTGAAATTCTTAGATATTAGAAGGAACACCGATGGCGAAGGCATTCACCTGGGCTATAACTGACGCTGATGCGCGAAAGCGTGGGGAGCAAACAGG
## 3035                                                         ACGAAGGGTGCAAGCGTTGTTCGGAATTACTGGGCGTAAAGGGTGTGTAGGCGGTTTGATAAGATAGGGGTGAAATCCCGGGGCTTAACCTCGGAACTGCCTTTATAACTATTAGACTTGAATAATGGAGAGGACAGTGGAATACCCAGTGTAGAGGTGGAATTCGTAGATATTGGGTAGAACACCAGAGGCGAAGGCGACTGTCTGGCCATTAATTGACGCTGAGGCACGAAAGTGTGGGGAGCAAACAGG
## 3037                                                          ACATAGGTGGCAAGCGTTATCCGGAATTATTGGGCGTAAAGAGTGAGTAGGTGGCCATTTAAGTCACAAGTCAAATGCAGTAGCTCAACTACTGTTCGCTTGTGATACTGAAAGGCTAGAGTGCGAGAGGGGTAAGTGGAATTCCATGTGTAGCGGTGAAATGCGTAGATATATGGAGGAACACCAGTGGCGAAGGCGGCTTACTGGCTCGTAACTGACACTCAGTCACGAAAGCGTGGGGAGCAAATAGG
## 3038                                                          ACGGAGGGAGCTAGCGTTGTCCGGATTTATTGGGCGTAAAGGGCATGTAGGTTGGATGATCAGTCAGTTGTAAAATCTTATGGCTTAACCATATTAAGCAATTGATACTGTCATTCTTGAGTTTATGAGGGGTCGATGGAATACCTAGTGTAGCGGTGGAATGCGTAGATATTAGGTAGAACACCAGTGGAGAAGTCGGTCGACTGGCATAATACTGACACTGAGATGCGAAAGCGTGGGTAGCAAACAGG
## 3039                                                         ACGAAGGGGGCGAGCGTTGTTCGGAATCACTGGGCTTAAAGAGTGCGTAGGTGGATTTGTAAGTCAGGAGTGAAATCCCGGGGCTCAACCTCGGAACTGCTTTTGAAACTGCAAGTCTGGAGTCCCGGAGGGGTTAGGAGAATTCCGAGTGTAGAGGTGAAATTCGCAGATATTCGGAGGAATACCAGTGGCGTAGGCGCCTAACTGGACGGGTACTGACACTGAGGCACGAAAGCGTGGGGATCAAACAGG
## 3042                                                         ACGAAAGGTGCTAGCGTTACTCGGATTTACTGGGCGTAAAGCGCGTGTAGGTTGATTTGTAAGTCAGTTGTTAAAAACCGAAGCTTAACTTCGGCGCTGCAATTGAAACTGCTTATCTTGAATATGGTAGGGGATAGCGGAATTTCTGGTGTAGGGGTAAAATCCGTAGATATCAGAAGGAACACCAAAGGCGAAGGCAGCTATCTGGGCCATTATTGACACTGAGACGCGAAAGCGTGGGGAGCAAACAGG
## 3045                                                          ACGTAGGGGGCGAGCGTTATCCGGATTTATTGGGCGTAAAGCGTGTGTAGGTGGTTCGTAAGGTCAGGCGTAAAATATCACAGCTTAACTGTGTAGGGCGTTTGAAACCGACGGACTTGAATACGGTAGAGGCAGGTGGAATTCCATGTGTAGCGGTGGAATGCGTAGATATATGGAGGAACACCAATGGCGAAGGCAACCTGCTGGGCCGGAATTGACACTGAGACACGAAAGCGTGGGGAGCAAACAGG
## 3048                                                         ACGGAGGGGGCTAGCGTTGTTCGGAATTACTGGGCGTAAAGCGCGCGCAGGCGGCATAATAAGTTGGAAGTGAAAGCCCGGGGCTTAACCTCGGAATGGCTTCCAAAACTGTTATTCTAGAGTGTAGTAGGGGATGATGGAATTCCTAGTGTAGTAGTGAAATACGCAGATATTAGGAGGAACACCGGTGGCGAAAGCGATCATCTAGGCTACAACTGACGCTGAGGCGCGAAAGCATGGGGAGCAAACAGG
## 3049                                                         ACGAAGGGTGCAAACGTTGCTCGGAATTATTGGGCGTAAAGCGCACGTAGGCGGCTTAGCAAGTCGGAAGTGAAATCCCACGGCTCAACCATGGAAGTGCTTTCGAAACTGCTGAGCTTGAGTACGAAAGAGGATCGCGGAATTCCCGGTGTAGAGGTGAAATGCGTAGATATCGGAAGGAACATCAGTGGCGAAAGCGGCTTCCTGGTCCAGCACCGACGTTCAGGCACGAAAGCGTGGGGAGCAAACAGG
## 3050                                                         ACGGAGGGGGCTAGCGTTGTTCGGAATTATTGGGCGTAAAGGGTGCGTAGGCGGACTAAAAAGTTAGAAGTGAAATCCTGGGGCTCAACCTCAGAACTGCTTTTAAAACTCTTAGTCTTGAGTCTGGCGGGGGGTAACGGAATTCCTAGTGTAGGGGTAAAATCCTTAGATATTAGGAAGAACACCGATGGCGAAGGCAGTTACCTATACCATGACTGACGCTGAGGCACGAAAGCGTGGGGAGCAAACAGG
## 3064                                                         ACGTAGGTGGCAAGCGTTGTCCGGATTTATTGGGTTTAAAGGGTGCGTAGGCGGTTTGATCAGTCAGTCGTTAAAGGCAGCAGCTTAACTGTTGTAAGGCGATTGATACTGTCGAACTTGAATTCGTTTGAGGTACTTGGAATGCGTAGTGTAGCGGTGAAATGCATAGATATTACGCGGAACGCCAATTGCGGAGGCAGAGTACTAGGACGAGATTGACGCTGAGGCACGAAAGTGTGGGTAGCAAACAGG
## 3079                                                         ACGGAGGATGCAAGCGTTATCCGGATTTATTAGGTTTAAAGGGTGTGTAGGCGGAAAACTAAGTCAGCGGTGAAATCCTACGGCTTAACTGTAGAACTGCCATTGATACTGGTTTTCTTGAATGCATTTGAAGTGGGCGGAATGTGTAATGTAGCGGTGAAATGCTTAGATATTACACAGAACACCAATTGCGAAGGCAGCTCACTAAATTGTGATTGACGCTGAGGCACGAAAGCGTGGGGAGCAAACAGG
## 3085                                                         ACAGAGGGTGCAAGCGTTGTTCGGAATCACTGGGCGTAAAGCGCGCGCAGGCGGATGGTTGCGTCCGGAGTGAAATTTCGCCGCTCAACGGCGAAGCTGCTTTGGATACGGACCGTCTAGGGACCGGTAGAGGCTGGTAGAATTGCAGGTGTAGCGGTGGAATGCGTAGAGATCTGCAAGAATACCCGTGGCGAAGGCGGCCAGCTGGGCCGGATCCGACGCTGAGGCGCGAAAGCGTGGGGAGCAAACAGG
## 3090                                                          ACGTAGGTGGCGAGCGTTATCCGGATTTATTGGGCGTAAAGGGTGCGTAGGCGGTATGTTAAGTATAAAATAAAAGCCCGAAGCTTAACTTCGGTTCGTTTTATAAACTGGCAAACTAGAGTACAGTAGGGGCAAATGGAATTCCTAGTGTAGTGGTAAAATGCGTAGATATTAGGAGGAACACCAGTGGCGAAGGCGATTTGCTGGGCTGTAACTGACGCTGAGGCACGAAAGCGTGGGGAGCAAATAGG
## 3091                                                         ACGTAGGGTGCAAGCGTTAATCGGAATTACTGGGCGTAAAGCGTGCGCAGGCGGTTATGTAAGACAGATGTGAAATCCCCGGGCTCAACCTGGGAACTGCATTTGTGACTGCATAGCTAGAGTACGGTAGAGGGGGATGGAATTCCGCGTGTAGCAGTGAAATGCGTAGATATGCGGAGGAACACCGATGGCGAAGGCAGTCCCCTGGGATAAGACTGACGCTCATGCACGAAAGCGTGGGGAGCAAACAGG
## 3094                                                         ACATAGGGTGCAAGCGTTGTCCGGAATTATTGGGCGTAAAGAGCTCGTAGGTGGTTCGATACGTCGGATGTGAAAATCAGGGGCTCAACCCCTGACCTGCATTCGATACGGTCGAGCTAGAGTTTGGTAGGGGAGACTGGAATTCCTGGTGTAGCGGTGGAATGCGCAGATATCAGGAGGAACACCGATGGCGAAGGCAGGTCTCTGGGCCAATACTGACACTGAAACAAGAAAGCGTAGGTAGCGAATGGG
## 3097                                                         ACGTAGGATGCAAGCGTTGTCCGGAATTATTGGGTTTAAAGGGTGTGTAGGTTGTTACATAAGTCAGCGGTGAAATACATTTGCTTAACAGATAGTAAGACGTTGATACTGTGTGGCTAGAGTGCGATATAGGTGGGCGGAATTTGTAATGTAGCGGTGAAATGCATAGATATTACAAGGAACGTCGATAGCGAAGGCAGCTCACTGTGTCTGCACTGACACTGAGGCACGAAAGCGTGGGTATCGAACAGG
## 3104                                                         ACGGAGGGTGCAAGCGTTAATCGGAATCACTGGGCGTAAAGCGCATGTAGGCTGTCTTGTAAGTCGGGCGTGAAATCCCACGGCTCAACCGTGGAACTGCGTTCGATACTGCCAGACTTGAGTGCAGGAGAGGGTGGCGGAATTCCCGGTGTAGGAGTGAAATCCGTAGATATCGGGAGGAACACCAGTGGCGAAGGCGGCCACCTGGACTGTAACTGACGCTGAGATGCGAAAGTGCGGGTAGCAAACAGG
## 3107                                                         ACGGAGGGAGCTAGCGTTATTCGGAATTACTGGGCGTAAAGAGCGCGTAGGCGGTATAGTAAGTTGGAAGTGAAAGCCCAGGGCTCAACCCTGGAACTGCTTTCAAAACTACTAAACTAGAGATTGATAGGGGATAGCGGAATTCCTAGTGTAGAGGTGAAATTCTTAGATATTAGGAGGAACACCGGTGGCGAAAGCGGCTATCTGGCTCAAATCTGACGCTGAGGTGCGAAAGCGTGGGGAGCAAACAGG
## 3108                                                         ACGGAGGGTGCAAGCGTTATCCGGATTTATTGGGTTTAAAGGGAGCGTAGGCGGCCCGATAAGTCAGTGGTGAAATCTCCTGGCTTAACTAGGAAATTGCCATTGATACTGTTGGGCTTGAGTACAGTTGCCGTTGGCGGAATATGACATGTAGTGGTGAAATACGTAGAGATGTCATAGAACACCGATGGCGAAGGCAATCCCCTGGGCCTGCACTGACGCTCATGCACGAAAGCGTGGGGAGCAAACAGG
## 3110                                                        ACGGAGGATCCAAGCGTTATCCGGAATTACTGGGCGTAAAGAGTTGCGTAGGTGGCATAGTAAGCAGGTAGTGAAATTGCGGGGCTCAACCCCGTACCCATTACCTGAACTGCTAAGCTAGAGAATGAGAGAGGTAGATGGAATTTCTAGTGTAGGGGTAAAATCCGTTGATATTAGAAGGAACACCAATGGCGTAGGCAGTCTACTGGCTCATTTCTGACACTCAGGCACGAAAGCGTGGGGAGCAAACAGG
## 3112                                                         ACGGAGGGTGCGAGCGTTGTTCGGATTTACTGGGCGTAAAGGGTGTGTAGGCGGATATAAAGGTCAGGTGTGAAATCCCAAGGCTCAACCTTGGAACTGCATTTGAAACCGTATGTCTAGAGTGTGACAGAGGATGATGGAATTTCTAGTGTAGCAGTGAAATGCGTAGATATTAGAAGGAACACCGGAGGCGAAGGCGATCATCTGGGTCATAACTGACGCTGAGGCACGAAAGTGTGGGTAGCAAACAGG
## 3114                                                         ACGTAGGATGCGAGCGTTGTTCGGAATCACTGGGCGTAAAGCGTACGTAGGCGGTAAATTGTGTCTGCTATTAAAGACCGGGGCTCAACTCCGGGAATGTAGCAGAAACTGATATGCTAGAGGTTAGTAGAGGGAATCGGAATTCTCAGTGTAGCGGTGAAATGCGTAGATATTGGGAAGAACACCTGTGGCGAAAGCGGATTTCTGGGCTAATACTGACGCTGAGGTACGAAAGCGTGGGGAGCAAACGGG
## 3119                                                         ACAGAGGGACCTAGCGTTGTTCGGAATGACTGGGCGTAAAGGGTGCGTAGGTGGTTTGATAAGTCAGCGGTGAAAGTTTGCAGCTTAACTGTAAAAATGCCGTTGAAACTGTCGGACTTGAGTGTAAATGAGGTAGGCGGAATGCGTGGTGTAGCGGTGAAATGCATAGATATCACGCAGAACTCCGATTGCGAAGGCAGCTTACTAAGCTACAACTGACACTGAAGCACGAAAGCGTGGGGATCAAACAGG
## 3124                                                        CCCGACGCTCCGAGTCGCAGCCACAATTATTGGGTCTAAAACATTCGTAGCTTGTTTACTAAGTTTCTTGTGAAATCCGGCATCTTAAGTGTCGGGCGCGCAAGAAATACTGGTAGACTTGAGACCGGAAGAAGTAAGGAGTACAGCCGAGGTAGTAGTAAAATACGTTAATCTTGGTTGGACTAACAACAGCGAAGGCACCTTACTAGTACGGATCTGACAGTGAGGGATGAAGGCTAGGGTCGCGAAAGGG
## 3163                                                         ACGGAGGATCCAAGCGTTATCCGGAATCATTGGGTTTAAAGGGTCCGTAGGCGGTTTAATAAGTCAGTGGTGAAAGCCCATCGCTCAACGGTGGAACGGCCATTGATACTGTTAAACTTGAATTATTAGGAAGTAACTAGAATATGTAGTGTAGCGGTGAAATGCTTAGAGATTACATGGAATACCAATTGCGAAGGCAGGTTACTACTAATATATTGACGCTGATGGACGAAAGCGTGGGGAGCGAACAGG
## 3179                                                       ACGAAGGGGGCAAGCGTTATTCGAAATGATTGGGCGTAAAGGGTTCGTAGGTGGTACTTGTTACTATTAGTAAAAAAATTAAAAATGTTCTTTTTAATTGTGCTAATACTATAGTTGTACTTGAGTATAGCATAGAATAATAGAATTTCTACATAGGGGTAAAATCCATTTATTTATGAAGGAATGTCTTTGGTGAAAACGATTATTTAGGCTATTACTGACATTGAGGAACGAAAGCGTAGGGAGCAAACAGG
## 3182                                                         ACGTAGGGTGCAAGCGTTAATCGGAATTACTGGGCGTAAAGCGTGCGCAGGCGGTTATATAAGACAGATGTGAAATCCCCGAGCTTAACTTGGGAACTGCATTTGTGACTGTATAGCTAGAGTACGGCAGAGGGGGATGGAATTCCGCGTGTAGCAGTGAAATGCGTAGATATGCGGAGGAACACCGATGGCGAAGGCAATCCCCTGGACCTGTACTGACGCTCATGCACGAAAGCGTGGGGAGCAAACAGG
## 3229                                                        CCCAGCGCCACGAGTGGGAACCACAATTATTTGGCCTAAAGCGTCCGTAGCTGGCTTTGTAAATCTTTTGTGAAATCGGAAAGCTTAACTTTCCGGCGTGCAGAAGACACTGCATTGCTTGAGACCGGGAGGGGTAAGGAGTATTTCCAGGGAACCGGTAAAATGGGATAATCCTGGAAGGACTACCTGTGGCGAAGGCGCCTTACCAGAACGGGTCTGACAGTGAGGGACGAAGGCTAGGGGAGCGAACCGG
## 3242                                                         ACGTAGGGTGCAAGCGTTAATCGGAATTACTGGGCGTAAAGCGTGCGCAGGCGGACTTTTAAGCCAGATGTGAAAGCCCCGAGCTTAACTTGGGAATTGCGTTTGGAACTGAGCATCTAGAGTCTGTCAGAGGGGGATGGAATTCCACGTGTAGCAGTGAAATGCGTAGAGATGTGGAGGAACACCGATGGCGAAGGCAGCCCCCTGGGCCAATACTGACGCTCATGCACGAAAGCGTGGGGAGCAAACAGG
## 3253                                                         ACGGAGGATCCAAGCGTTATCCGGATTTATTGGGTTTAAAGGGTCCGTAGGCGGGTCTTTAAGTCAGTGGTGAAAGCCAAGGGCTCAACCCTTGAACTGCATCTGATACTGGTATTCTTGAGTTCGGAAGAGGAAGGCGGAATTCCCGGTGTAGCGGTGAAATGCGTAGATATCGGGAAGAACACCGGTGGCGAAGGCGGCCTTCTGGTCTAGGACTGACGCTGACATGCGAAAGTGTGGGGATCAAACGGG
## 3266                                                         ACGTAGGGTGCAAGCGTTAATCGGAATTACTGGGCGTAAAGCGTGCGCAGGCGGTTATGCAAGACAGATGTGAAATCCCCGGGCTCAACCTGGGAACTGCATTTGTGACTGCATAGCTAGAGTACGGTAGAGGGGGATGGAATTCCGCGTGTAGCAGTGAAATGCGTAGATATGCGGAGGAACACCGATGGCGAAGGCAGTCCCCTGGGATAAGACTGACGCTCATGCACGAAAGCGTGGGGAGCAAACAGG
## 3278                                                         ACGTAGGGTGCAAGCGTTGTTCGGAATCATTGGGCGTAAAGGGTCCGTAGGCGGACAGACAAGTCAGGTGTGAAATCCCGCAGCTTAACTGCGGAACTGCGCTTGAAACTATCTGTCTAGAGGATGGTAGAGGTAACTGGAATTCCTAGTGTAGAGGTGAAATTCGTAGATATTAGGAGGAACACTCGTGGCGAAGGCGGGTTACTGGGCCATTCCTGACGCTGATGGACGAAAGCATGGGTAGCAAACAGG
## 3280                                                         ACGTATGGTGCAAGCGTTGTTCGGAATTATTGGGTGTAAAGGGTATGTAGGCGGAATTGTAAGTCAGATGTGAAACCTAAGGGCTCAACCCTTAGCCTGCATTTGATACTGTGATTCTAGAGTTTGGGAGAGGCAAGCGGAATTCCTGGTGTAGCGGTGAAATGCGTAGATATCAGGAAGAACACCGGTGGCGAAAGCGGCTTGCTGGCCTAAAACTGACGCTGAGATACGAAAGTGTGGGGAGCGAACGGG
## 3318                                                         ACAGAGGATGCAAGCGTTATCCGGAATGATTGGGCGTAAAGCGTCTGTAGGTGGCCTGACAAGTCTTCTGTCAAAGATCAGGGCTTAACCCTGGACAGGCAGGAGAAACTATCAGGCTAGAGTTCGGTAGAGGCAGAGGGAATTCCCAGTGAAACGGTGAAATGTGTAGATATTGGGAGGAACACCAACGGCGAAAGCACTCTGCTGGGCCGAGACTGACGCTCATGCACGAAAGCGTGGGGAGCAAACAGG
## 3330                                                         ACGTAGGGTGCGAGCGTTAATCGGAATTACTGGGCGTAAAGCGTGCGCAGGCGGTTATATAAGACAGTTGTGAAATCCCCGGGCTCAACCTGGGAATTGCATCTGTGACTGTATAGCTAGAGTACGGTAGAGGGGGATGGAATTCCGCGTGTAGCAGTGAAATGCGTAGATATGCGGAGGAACACCGATGGCGAAGGCAATCCCCTGGGCCTGTACTGACGCTCATGCACGAAAGCGTGGGGAGCAAACAGG
## 3338                                                         ACGTAGGGTGCAAGCGTTAATCGGAATTACTGGGCGTAAAGCGTGCGCAGGCGGTTCGGAAAGAGAGGTGTGAAATCCCAGGGCTCAACCTTGGAACTGCACTTTTAACTACCGAGCTAGAGTATGTCAGAGGGGGGTGGAATTCCGCGTGTAGCAGTGAAATGCGTAGATATGCGGAGGAACACCGATGGCGAAGGCAATCCCCTGGACCTGTACTGACGCTCATGCACGAAAGCGTGGGGAGCAAACAGG
## 3361                                                         ACGTAGGGTGCAAGCGTTAATCGGAATTACTGGGCGTAAAGCGTGCGCAGGCGGTTATGCAAGACAGATGTGAAATCCCCGGGCTCAACCTGGGAACTGCATTTGTGACTGCATAGCTAGAGTACGGTAGAGGGGGATGGAATTCCGCGTGTAGCAGTGAAATGCGTAGATATGCGGAGGAACACCGATGGCGAAGGCAATCCCCTGGGCCTGCACTGACGCTCATGCACGAAAGCGTGGGGAGCAAACAGG
## 3384                                                        ACGTAGGAGGCGAGCGTTATCCGGATTTACTGGGCGTAAAGCGCGTGTAGGCGGTTTCATAAGTCGTGCGTGAAAGCTCCCTGGCTCAACCGGGAGAGGTCGTGCGATACTGCGAAACTTGAGTCAGACAGAGGCAAGTAGAATTCCGGGTGTAGTGGTGAAATGCGTAGATATCCGGAGGAATACCAGTGGCGAAGGCGACTTGCTGGGTCTGTACTGACGCTCAGACGCGAAAGCATGGGGAGCGAACGGG
## 3391                                                         ACGTAGGTGTCAAGCGTTGTTCGGATTTATTGGGCGTAAAGGGTCCGCAGGTGGCCTGACAAGTCTGATGTGAAATCCCGGAGCTCAACTCCGGAACTGCATTGGAAACTGTCGGGCTGGAGTATCGGAGAGGAGATTGGAATTCCAGGTGTAGCGATAAAATGCGTAGATATCTGGAAGAACACCGAAAGCGTAGGCAGATCTCTGGACGATCACTGACACTCAGGGACGAAAGCATGGGGAGCAAACAGG
## 3430                                                          ACAGAGGTCCCGAGCGTTATCCGGATTTACTGGGCGTAAAGAGTCCGTAGCTGGCATTGTAAGTCTTGATTCAAATACTCTGACTCACTCAGAGAAAGGGTTGAGATACTGCAAAGCTAGAGATCATAGGGGGTTAGTGGAATTTCCAGTGGAGGGGTGAAATCCGTTGATATTGGAAGGAATGCCGAAAGCGAAAGCAGCTAACTACTACATTCTGACAGTCAAGGACGAAAGCTTGGGGATCGAACGGG
## 3438                                                         ACAGAGGGTGCAAGCGTTGTTCGGAATCACTGGGCGTAAAGAGTTCGTAGGTGGACTTATTAGTACAGGGTTAAATACCGGGGCCTAACCCCGGACCTGCTTTGTAAACGGTAAGTCTAGAGTACTAGAGAGGATAGCGGAATTCCTGGTGTAGAGGTGAAATTCGTAGATATCAGGAGGAACATCTGAGGCGAAGGCGGCTATCTGGCTAGGCACTGACACTAAAGAACGAAAGCATGGGGATCAAACAGG
## 3449                                                         ACGTAGGGTGCGAGCGTTAATCGGAATTACTGGGCGTAAAGCGTGCGCAGGCGGTTATGTAAGACAGTTGTGAAATCCCCGGGCTCAACCTGGGAATTGCATCTGTGACTGCATAGCTAGAGTACGGTAGAGGGGGATGGAATTCCGCGTGTAGCAGTGAAATGCGTAGATATGCGGAGGAACACCGATGGCGAAGGCAATCCCCTGGGCCTGTACTGACGCTCATGCACGAAAGCGTGGGGAGCAAACAGG
## 3453                                                       CCCGGCAGTCCAAGTCGCATCCACTATTATTGGGTCTAAAACATCCGTAGCTTGCAATTTAAGTTCCTTGTGAAATCCTACGTCTCAAGCGTAGGGCGAGCGGGGAATACTGAATTGCTAGAGATCGGTAGACGTAAGAAGTACGCTGGAGGTAGTGGTTAAATACGTTAATCTTCGGTGGACTAACAATGAGCGAAGGCATCTTACGAGGACGAATCTGACAGTGAGGGATGAAGGCCAGGGGCGCAAAACGG
## 3466                                                         ACGTAGGGTGCAAGCGTTAATCGGAATTACTGGGCGTAAAGCGTGCGCAGGCGGTTGTGTAAGTCAGATGTGAAATCCCCGGGCTCAACCTGGGAACTGCTTTCAAAACTGCTAGTCTTGAGTGAAGTAGGGGATGATGGAATTCCTAGTGTAGAGGTGAAATTCTTAGATATTAGGAGGAACACCGGTGGCGAAGGCGGTCATCTGGACTTCAACTGACGCTGAGGCGCGAAAGCGTGGGGAGCAAACAGG
## 3484                                                         ACGTAGGGTGCGAGCGTTGTTCGGAATTACTGGGTGTAAAGGGTCCGCAGGCGGGCGAGCAAGTCGGTGGTGAAAGTTCATGGCTTAACTGTGAAATTGCCTTCGATACTGCTCGTCTTGAGTTCGAGAGAGGATGATGGAATTCATGGTGTAGCAGTGAAATGCGTAGAGATCATGAAGAACACCAGTTGCGAAGGCGGTCATCTGGCTCGATACTGACGCTCAGGGACGAAAGCGTGGGTAGCAAACAGG
## 3485                                                        ACCAGCGCCACGAGTGGGGGCCGTATTTATTGGGCCTAAAGCGTCCGTAGCCGGCTTTGTAAATTTCTTGTGAAATTGTCAGGCTTAACCTGACAGATGGCAAGGAACACTGCAGAGCTAGGGACCGGGAGGGGTCAGAGGTATTACAGGGGTAGCGGTAAAATGTTATAATCCTTGTAAGACCACCTGTGGCGAAGGCGTCTGACTAGAACGGTTCCGACGGTGAGGGACGAAAGCTAGGGGAGCGATCCGG
## 3489                                                         ACGTAGGGTGCAAGCGTTAATCGGAATTACTGGGCGTAAAGCGTGCGCAGGCGGTTATGTAAGACAGAGGTGAAATCCCCGGGCTCAACCTGGGAACTGCCTTTGTGACTGCGAGGCTAGAGTATGGCAGAGGGGGGTGGAATTCCACGTGTAGCAGTGAAATGCGTAGAGATGTGGAGGAACACCGATGGCGAAGGCAGCCCCCTGGGCCAATACTGACACTGAGGAGCGAAAGCGTGGGGAGCGAACAGG
## 3491                                                         ACAGAGGATGCAAGCGTTATCCGGAATCACTGGGCATAAAGCGTCTGTAGGTGGTTTGGTAAGTCTGCTGTTAAAGACTGGGGCTCAACCTGGGAACTGCATTTGTGACTGCAAGGCTAGAGTACGGTAGAGGGGGATGGAATTCCGCGTGTAGCAGTGAAATGCGTAGATATGCGGAGGAACACCAATGGCGAAGGCAATCCCCTGGACCTGTACTGACGCTCATGCACGAAAGCGTGGGGAGCAAACAGG
## 3506                                                         ACAGAGACCTCAAGCGTTATCCGGATTTATTGGGCGTAAAGCGCAGGTAGGCGGGTTTATTAGTCGGATGTCAAAGACCGGGGCTCAACCCTGGAACTGCCTTCGATACTGAAAGTCTTGAGTCCGGAAGAGGTAAGTGGAACTCCGAGTGTAGAGGTGAAATTCGTAGATATTCGGAAGAACACCAGTGGCGAAGGCGGCTTACTGGTCCGGTACTGACGCTGAGGTGCGAAAGCGTGGGGAGCAAACAGG
## 3522                                                         ACGTAGGATCCAAGCGTTATCCGAATTCACTGGGCGTAAAGCGCTTGCAGGCGGTCTGGTAAGTTGGATATGAAAGCTCCTGGCTCAACTGGGAGAGGCTGTTCAAGACTGTCAGACTAGAGTACGACAGAGGGAGGTGGAATTCCAGGTGTAGTGGTGAAATGCGTAGATATCTGGAGGAACACCAGAGGCGAAAGCGGCCTCCTGGGTCGCAACTGACGCTCAGAAGCGAAAGCTAGGGGAGCGAACGGG
## 3545                                                                                                              ACAGGAGGGGCAAGCTTAGAGCGAATTTACTAGGTATACAGCGCATCGAAGCGTACTAGTATCAGATAATAGTAGGGCAACACTAGGAGTATCGGTCAAATGATTATATACCTGGTGGAAAGCCGATGGCGAAGGCACCTACAAGTCTATACTGAAGCTTAAATGCGGAAGCTTGGGGAGCAAAGCGGATTAGATACCC
## 3565                                                         ACGTAGGGTGCAAGCGTTAATCGGAATTACTGGGCGTAAAGCGTGCGCAGGCGGTTATGTAAGACAGTTGTGAAATCCCCGGGCTCAACCTGGGAACTGCATCTGTGACTGCATAGCTAGAGTACGGCAGAGGGGGATGGAATTCCGCGTGTAGCAGTGAAATGCGTAGATATGCGGAGGAACACCGATGGCGAAGGCAATCCCCTGGGCCTGCACTGACGCTCATGCACGAAAGCGTGGGGAGCAAACAGG
## 3577                                                         ACGAAGGGGGCTAGCGTTGTTCGGATTTACTGGGCGTAAAGCGCACGTAGGCGGATCTTTAAGTCAGAGGTGAAATCCCAAGGCTCAACCTTGGAACTGCCTTTGATACTGGGGATCTCGAGTCCGGGAGAGGTGAGTGGAACTGCGAGTGTAGAGGTGAAATTCGTAGATATTCGCAAGAACACCAGTGGCGAAGGCGGCTCACTGGCCCGGTACTGACGCTGAGGTGCGAAAGCGTGGGGAGCAAACAGG
## 3579                                                         ACGTAGGTGGCAAGCGTTGTCCGGATTTATTGGGTTTAAAGGGTGCGTAGGTGGCTTGGTAAGTCAGCGGTAAAAGACGGCAGCTTAACTGTTAGTGCGCCGTTGATACTGCCAAGCTTGAGTAAAGTTGAGGTGGGCGGAATGAATAGTGTAGCGGTGAAATGCATAGATATTATTCAGAACGCCAATTGCGAAGGCAGCTCACTTAGTTTTCACTGACACTGAGGCACGAAAGCGTGGGGATCAAACAGG
## 3595                                                         ACGAAGGTGGCGAATGTTGTTCGGATTTACTGGGCGTAAAGCGTCCGCAGGTGGTTTATTACGTCAGGTGTGAAAGCCCCGTGCTCAACATGGGAACTGCACTTGAAACGGATTGACTTGAAGGTAAGAGAGGTTGGCAGAATATCTGGTGTAGAGGTGAAATTCGTAGATATCAGATGGAATACCCATGGCGAAGGCAGCTAACTGGCTTATTCTTGACACTCATGGACGACAGCGTGGGGAGCAAACAGG
## 3612                                                         ACAGAGGTGGCGAGCATTGTCCGGAATTACTGGGTGTAAAGGGTGCGTAGGCTGTTTTGTAAGTCTGAGGTTAAAGACTTTCGCTTAACGAAAGGACTGCTTTGGATACTGCAAGACTAGAGAACCAAAGGGGAAAATAGAATTCCCGGTGTAGCGGTGGAATGTGTAGAGATCGGGAGGAATACCAGTGGCGAAGGCAGTTTTCTGGTTGGTTTCTGACGCTGAGGCACGAAAGCATGGGTAGCAAACAGG
## 3625                                                        ACGTAGGTCCCAAGCATTATCCGAAATTACTGGGCGTAAAGCGATCAAAGGCGGTGTAATAAGTCTGATGTTAAAGCCCTCGGCTCAACTGAGGAATCGTATCGGAAACTGTTATACTAGAGATATGTAGGGGTAGCTGGAATTTCTATTGTAGGGGTTAAATCCGTAGATAATAGAAGGAACACCAAAAGCGAAGGCAGGCTACTGGACATTTTCTGACGCTGCTTGATCGAAAGCGTGGGGAGCAAAACGG
## 3651                                                         ACGGAGGGTGCGAGCGTTACCCGGATTTACTGGGCGTAAAGGGCGTGTAGGCGGTTTGTTAAGTCCGATGTGAAAGACCGGGGCTCAACCCCGGGAACGCGTTGGATACTGGCAGACTAGACGGCTGGAGGGGGTAGCGGAATTTCCGGAGTAGCGGTGAAATGCGCAGATACCGGAAGGAACGCCAATAGCGAAGGCAGCTACCTAGACAGCACGTGACGCTGAGGCGCGAAAGCATGGGGAGCAAACCGG
## 3658                                                        ACAGAGGTTACAAGCATTGTCCGGAATCACTGGGCGTAAAACGTTTCGCAGGTGTTTTCTCAAGTTATGTTTGAAAGACCGAGGCTCAACCTCGGAAACGGGCGTAATACTGGGAGAATTGAGGGATATTGGGGCTACTGGAACTGTCGGTGTAGGGGTGAAATCCGTTAATATCGACAGGAACTCCGAGGGCGTAGGCAGGTAGCTAAGTATCTCCTGACACTCATGAACGAAAGCTAGGGTAGCGAAAGGG
## 3702                                                        ACGAAGGGGGCAAGCGTTATTCGGATTTATTGGGCGTAAAGGGCGCGTAGGTGGCTGGGAAAGTCTGGTGTGAAAACGCTACCGCTTAACGGAGCGAGGTCGCTGGAAACTTCCCAACTTGAGAGACGTAGAGGTGAGCAGAATGCCTGGTGTAGCGGTGGAATGCTTAGAGATCAGGCAGAATACCCGAGGCGAAGGCGGCTCACTGGACGTTTTCTGACACTAATGCGCGAAAGCGTGGGGATCAAACGGG
## 3704                                                          CAGAGACCTCAAGCGTTATCCGGAGTTATTGGGCGTAAAGGGTCCGCAGGCGGCCTTGCGCGTCATACGTTAAATGTTTGGGCTCAACCCAGACACCGCGTGTGATACGGCAAAGCTAGAGACTGGGAGAGGCACATGGAATTGTCGGTGTAGCGGTAAAATGCGTTAATATCGACAAGAACACCAAAGGCGAAGGCAGTGTGCTAGAACAGTTCTGACGCTCAGGGACGAAAGCGTGGGGAGCGAATGAG
## 3728                                                         ACAGAGGGTGCAAACGTTGCTCGGAATTACTGGGCGTAAAGCGCGTGTAGGTGGACCAGCAAGTCAGACGTGAAAGCCCTGGGCTCAACCCAGGAACTGCGTTTCGAACTGATTAGCTAGAGTGCAGTAGAGGGTGGTGGAATTTCCGGTGTAGCGGTGAAATGCGTAGAGATCGGAAGGAACATCAGTGGCGAAGGCGACTGCCTGGACTGACACTGACGCTGAGGTGCGAAAGCGTGGGGAGCAAACAGG
## 3738                                                         ACGTAGGGTGCGAGCGTTAATCGGAATTACTGGGCGTAAAGCGTGCGCAGGCGGTTATATAAGACAGTTGTGAAATCCCCGGGCTCAACCTGGGAATTGCATCTGTGACTGTATAGCTAGAGTACGGTAGAGGGGGATGGAATTCCGCGTGTAGCAGTGAAATGCGTAGATATGCGGAGGAACACCGATGGCGAAGGCAATCCCCTGGGCCTGCACTGACGCTCATGCACGAAAGCGTGGGGAGCAAACAGG
## 3739                                                         ACGTAGGGTGCGAGCGTTAATCGGAATTACTGGGCGTAAAGCGTGCGCAGGCGGTCTTGCAAGACAGGTGTGAAATCCCCGGGCTTAACCTGGGAACTGCACTTGTGACTGCAAGGCTGGAGTACGGCAGAGGGGGATGGAATTCCGCGTGTAGCAGTGAAATGCGTAGATATGCGGAGGAACACCGATGGCGAAGGCAGGTCTCTGGGCCAATACTGACACTGAGGAGCGAAAGCGTGGGGAGCGAACAGG
## 3762                                                         ACGTAGGGTGCAAGCGTTAATCGGAATTACTGGGCGTAAAGCGTGCGCAGGCGGCTTTGCAAGACAGATGTGAAATCCCCGGGCTCAACCTGGGAACTGCATTTGAGACTGTATAGCTAGAGTACGGTAGAGGGGGATGGAATTCCGCGTGTAGCAGTGAAATGCGTAGATATGCGGAGGAACACCGATGGCGAAGGCAATCCCCTGGGCCTGTACTGACGCTCATGCACGAAAGCGTGGGGAGCAAACAGG
## 3780                                                         ACGGAGGGGGCAAGTGTTACTCATCATGACTAGGCGTAAAGGGTGTTAAGATAGCTAGAGGTTGTTCTGATTAAAAAGTCAACTGAATTGTTGAAACGTATTAAGTAAACAACCTAGCTTGAGTAGTTTGGAGAATAGAGTTACTAGTTTTCAAGGGGTAGAATCCAACGAGATGATTAGGAGAAACAAAGGCGAAGGCAACTATTTATGGACAACTGACATTAATACACAAAAGCGTGGGGAGCGAACAGG
## 3785                                                         ACGGAGGGTGCAAGCGTTATCCGGATTTACTGGGTTTAAAGGGTGCGTAGGCGGGCAGGTAAGTCAGTGGTGAAATCCCCGAGCTTAACTCGGGAACTGCCGTTGATACTATCTGTCTTGAATATCGTGGAGGTAAGCGGAATATGTCATGTAGCGGTGAAATGCTTAGATATGACATAGAACACCAATTGCGAAGGCAGCTTGCTACACGGTCATTGACGCTGAGGCACGAAAGCGTGGGGATCAAACAGG
## 3811                                                         ACAGAGACCTCGAGCGTTATCCGGAATTACTGGGCGTAAAGCGCGAGCAGCCGGTTGAGTTAGTCCGATGTTAAATCCCAAGGCTTAACTTTGGAACCGCATTGGAAACGGCTCAACTCGAGTGGGTAAGAGGCACGCGGAACGTGCAGTGTAGGGGTGAAATCCATTGATATTGCACGGAACACCAAAAGCGAAGGCAGCACACTGGTGCACTCCTGACACTGAAACAAGAAAGCGTAGGTAGCGAATGGG
## 3812                                                         ACGGAGGGTGCAAGCGTTATCCGGATTCACTGGGTTTAAAGGGTGCGTAGGTGGATCTGTAAGTCAGAGGTGAAATCCCCGAGCTTAACTTGGGAACTGCCTTTGATACTATAGATCTTGAATATCGTGGAGGTAAGCGGAATATGTCATGTAGCGGTGAAATGCATAGAGATGACATAGAACACCAATTGCGAAGGCAGGTTACTACCAGTGGATTGACGCTGATGGACGAAAGCGTGGGGAGCGAACAGG
## 3823                                                        ACGTAGGGTGCAAGCGTTGTTCGGAATTACTGGGCGTAAAGGACGCGTAGGCGGGCCTATAAGTTAGTTGTGAAATCCCTGGGCTTAACCCAGGACGTGCAACTAAAACTGTATGGTCTTGAGTATTGGAGAGGATGGCGGAATTCCAGGTGTAGAAGTGAAATTCGTAGATATCTGGAAGAACACCGGAGGCGAAGGCGGCTGTCTGGCCAAATACTGACGCTAAGGCGTGAAAGCATGGGGATCAAACAGG
## 3830                                                         ACGTAGGGTGCAAGCGTTAATCGGAATTACTGGGCGTAAAGGGCGTGTAGGCGGGATGTTAAGTCTGACTTTAAAGACTGTGGCTCAACCACAGGAGTGGGCTGGATACTGGCATTCTAGACAACTGGAGAGGCGATTGGAATTACTGGTGTAGCGGTGGAATGCGTAGATACCAGTAGGAACGTCAATGGCGAAGGCAGGTCGCTGGACAGTTTGTGACGCTGAGGCGCGAAAGTGTGGGGAGCAAACCGG
## 3845                                                         ACAGAGACCACGAGCGTTATCCGGATTTATTGGGCGTAAAGCGTCCGCAGGCGGTTTGAAAAGTCTTGTGTTAAATTTCTGGGCTCAACCCAGAACCTGCACGAGATACTCTCAAACTAGAGGATGGGAGAGGTACACGGAATTGTTGGTGTAGCGGTTAAATGCGTTAATATCAACAAGAACACCAAATGCGAAAGCAGTGTACTGGAACATTTCTGACGCTCAGGGACGAAAGCGTGGGGAGCGAACGGG
## 3851                                                         ACAGAGGGCTCGAGCGTTAATCGGAATCACTGGGCTTAAAGCGTGCGTAGGCGGGCGTGTAAGCATCTTGTGAAAGCCCCCGGCTCAACCGGGGAATTGCTTGGTGAACTGCACGTCTTGAGGCAAGTAGAGGTGGTCGGAACGATAGGTGGAGCGGTGAAATGCGTAGATATCTATCGGAACGCCAACGGAGAAGTCAGGCCACTGGGCTTGTTCTGACGCTGAGGCACGAAAGCGTGGGTAGCGAACGGG
## 3871                                                          ACGTAGGTGGCGAGCGTTATCCGGATTTATTGGGCGTAAAGCGTGCGCAGACGGTTTATTAAGTCTAAGATGAAAGCCCGAAGCTTAACTTCGGATTGTTTTAGAAACTGGTAGGCTTGAGTATAGTAGAGGCAAATGGAATTTCTAGTGTAGCGGTAAAATGTGTAGATATTAGAAGGAACACCAGTGGCGAAAGCGATTTGCTAGGCTATTACTGACGTTCATGCACGAAAGCGTGGGGAGCAAATAGG
## 3924                                                         ACATAGGGTGCAAGCGTTGTCCGGAATTATTGGGCGTAAAGAGCTCGTAGGTGGTTCGTTACGTCGGATGTGAAAATCTGGGGCTCAACCCCAGACCTGCATCCGATACGGTCGAGCTAGAGTTTGGTAGGGGAGACTGGAATTCCTGGTGTAGCGGTGGAATGCGCAGATATCAGGAGGAACACCAATGGCGAAGGCAGGACTCTGGGCCATTACTGACACTGAGGAGCGAAAGCGTGGGGAGCGAACAGG
## 3929                                                         ACAGAGGGTGCAAGCGTTAATCGGAATTACTGGGCGTAAAGCGCGCGTAGGCGGCTGATTAAGTCGGATGTGAAATCCCCGGGCTTAACCTGGGAACTGCGTTCGATACTGATCGGCTAGAGTATGAGAGAGGGAGGTAGAATTCCACGTGTAGCGGTGAAATGCGTAGATATGTGGAGGAATACCGGTGGCGAAGGCGGCCTCCTGGCTTAATACTGACGCTCATGCACGAAAGCGTGGGGAGCAAACAGG
## 3932                                                          ACGGGGAGGACAAGTGTTGTTCGGATTTATTGGGCGTAAAGGGAGCGTAGGTGGATATATAAGTGAAATTTGAAATATTCATTCATGTTGAATCAAGGATTTCATACTATAATATCTTGAGTGATTTCGGGGAAAATAGAATTATTGGTGGAGTAGTTACATGCGTTGATCTCAATAGGAATTCCGGTACTGGAGAGTATTTTCTAGAAATTTACTGACACTGAGGCTCGAAAGCATGGATAGTCAAAGGG
## 3955                                                         ACGGAGGGTGCAAGCGTTGTTCGGAATTACTGGGCGTAAAGCGTGTCTAGGTGGATATGTAAGTCAGGTGTGAAAGACCTTGGCTCAACCAAGGAATTGCGCCTGAAACTGCGTATCTTGAACAAGGAAGAGGTGGGTAGAATACCTAGTGCAGCAGTGAAATGCGTAGATATTAGGTAGAACACCGGAGGCGAAGGCGGCCCACTGGTCCTTTGTTGACACTGAAGCACGAAAGCGTGGGGAGCAAACAGG
## 3984                                                        ACGGAGGATCCAAGCGTTATCCGGAATTACTGGGCGTAAAGAGTTGCGTAGGCGGCCAAGTAAGCAGAATGTGAAATGATACGGCTCAACCGTATCCCCATGTTTTGAACTACTTGGCTAGAGAATGGAAGAGGTCGCTAGAATTCCTGGTGTAGGGGTGAAATCCGTAGATATCAGGAGGAATACCGATGGCGTAGGCAGGCGACTGGTCCATTTCTGACGCTCAGGCACGAAAGCGTGGGGAGCGATCGGG
## 3998                                                         ACGAGGGGAGCGAGCGTTATACAGTCAAACTGGGCGTAAAGGGTCCTTCGACGAAAAAGTGTAAGAAAGTGGCGAAAACTTAGGGGGTAACTTTAAGTGTTTACTTTTACTACTTTTTTAGAATTTTATAAGGGTTAACGTTATTCCGTCAAGAGAGTTAAAATGCATAGACTGACGGACGAGGGCCTTGGGCGAAAGCAGTTAACTATATAAGATTGACGTTTAGGGACGAAAGCATGGGGAGCGATCGGG
## 4002                                                         ACGTAGGGTGCGAGCGTTATCCGGAATTACTGGGCGTAAAGCGTCCGCAGCCGGACTTGTAAGTCTGATGTTAAATCCCGCAGCTCAACTGCGGAATCGTATCGGAAACTGCAAGTCTAGAGTGATGTAGAGGTACATGGAATTCATAGTGTAGGGGTAAAATCCGTTGATACTATGAGGAACACCAAAAGCGAAGGCAGTGTACTGGACATTTACTGACGGTCAGGGACGAAAGCGTGGGGATCAAAAAGG
## 4004                                                         ACGTAGGTGGCAAGCGTTATCCGGAATTACTGGGCGTAAAGCGCACGCAGGTGGCTATCTAAGATTGACGTGAAAGCTTCTGGCTTAACTGGAAGAAGTCGTTGATGACTGGATGGCTTGAGACGAGGAGAGGGGTGCAGAATTCCCGGTGTAGTGGTGGAATGCGTAGATATCGGGAGGAATACCAGTGGCGAAAGCGGCACCCTGGCCTTAGTCTGACGCTCAGGTGCGAAAGCGTGGGGAGCGAACGGG
## 4008                                                         ACGGGAGTGGCAAGCGTTATCCGGAATTATTGGGCGTAAAGCGTCCGCAGGCGGTCTTGTAAGTCTGTCGTTAAAGCGTGGAGCTTAACTCCATTTGAGCGATGGAAACTGCAAGACTAGAGTGTGGTAGGGGCAGAGGGAATTCCCGGTGTAGCGGTGAAATGCGTAGATATCGGGAAGAACACCAGTGGCGAAGGCGCTCTGCTGGGCCATAACTGACGCTCATGGACGAAAGCGTGGGGAGCAAACAGG
## 4010                                                         ACGGAAGGTCCTAGCGTTGTTCGGAATTACTGGGCGTAAAGCGCATGTAGGCGGAACAAAAAGTTAGAAGTGAAATCCCTGGGCTCAACCTAGGAATTGCTTTTAAAACTTTTGTTCTGGAATTCAGGAGAGGAAAGTGGAATTTCCAGTGTAGAGGTGAAATTCGTAGATATTGGAAGGAACACCAGTGGCGAAGGCGACTTTCTGGACTGATATTGACGCTGAGATGCGAAGGCATGGGTAGCAAACGGG
## 4013                                                         ACGTAGGGTGCAAGCGTTGTCCGGAATTACTGGGTTTAAAGGGTGCGTAGGCGGTTAAATAAGTCCTTGGTGAAATTTTGTAGCTTAACTATAAACATGCCAATGATACTGTATGACTAGAGTGGAATTGAGGTACTTGGAACGAGTGGTGTAGCGGTGAAATGCATAGAGATCACTCAGAACACCAATTGCGAAGGCAGAGTACTAAATTTTAACTGACGCTGAGGCACGAAAGTGTGGGGATCAAACAGG
## 4017                                                         ACGAGTGCCCCGAGCGTTATCCGGATTAACTGGGCGTAAAGGGTCCGTAGGGGGTTTTGTTAGTCCCGCGTTAAATTTATGGGCTCAACCCATAAGCCGCGCGGGAAACGGCAAAACTAGAGAATGCGAGAGGTTCGCGGAACCTATGGTGTAGGGGTGAAATCCGTTGATATCATAGGGAACACCGAAGGCGAAGGCAGCGAACTGGCGCATTTCTGACCCTGAGGGACGAAAGCGTGGGGAGCAAAAAGG
## 4018                                                         ACGTAGGGTGCGAGCGTTAATCGGAATTACTGGGCGTAAAGCGTGCGCAGGCGGTTATATAAGACAGTTGTGAAATCCCCGGGCTCAACCTGGGAATTGCATCTGTGACTGTATAGCTAGAGTACGGTAGAGGGGGATGGAATTCCGCGTGTAGCAGTGAAATGCGTAGATATGCGGAGGAACACCGATGGCGAAGGCAGTCCCCTGGGATAAGACTGACGCTCATGCACGAAAGCGTGGGGAGCAAACAGG
## 4019                                                         ACGGAGGATGCAAGCGTTATCCGGAATTATTGGGTTTAAAGGGTCCGCAGGCGGCCCGTTAAGTCAGTAGTGAAAACCCGGGGCTCAACCCCGGACGTGCTACAGATACTGGCGGGCTAGAGAAAGTCTGGAGGCAGCGGAATGTGACAAGTAGCGGTGAAATGCATAGATATGTCACAGAACACCGATAGCGAAAGCAGCTGCCTAAGACTTATCTGACGCTCAGGGACGAAAGCGTGGGGAGCAAACAGG
## 4049                                                         ACGGAGGATCCGAGCGTTATCCGGATTTATTGGGTTTAAAGGGAGCGTAGGTTGACGTATAAGTCAGCTGTGAAAGTTTACGGCTCAACCGTGAAATTGCAGTTGATACTGTATGTCTTGAGTGTACAAGAGGTGGGCGGAATTCGTGGTGTAGCGGTGAAATGCTTAGATATCACGAAGAACTCCAATTGCGAAGGCAGGACTCTGGGCCATTACTGACACTGAGGAGCGAAAGCGTGGGGAGCGAACAGG
## 4067                                                         ACGAACCGAGCGAACGTTGTTCGGAATCACTGGGCTTAAAGGGCGCGTAGGCGGGCTGCCGCGTCCGGGGTGAAGTCCTCCGGCTCAACCGGAGAACGGCCCCGGGTACGGGCGGCCTCGAGGGGGGTAGGGGCGTGCGGAACTGTGGGTGGAGCGGTGAAATGCGTTGATATCCACAGGAACTCCGGTGGCGAAGGCGGCACGCTGGACCCCATCTGACGCTGAGGCGCGAAAGCCAGGGGAGCGAACGGG
## 4079                                                    ACGAAGGGGGCAAGCGTTATTCGCCTTGACTGGGCGTAAAGGGTACGTAGGCTGTGTGTTTTATTTATGTAAATTAAATCCGGGCATAACCCTTGAAAATTATATAAGTGTAAAGCATTCTTTGAGTATATGCGAGGTATGTCTTATCTACGGTGTAAAGGTAAAATTTGACAATATCGTAGGGCAGTCCAAAAGGTGAAAACTACATACTAGCTATATACTGACGCTGAATTGCGAAAGCATGGGGTTCAAACAGG
## 4084                                                         ACGGGGGGGGCAAGCGTTGCTCGGAATTACTGGGCGTAAAGGGTCCGCAGGTGGCCTCGTAAGTTAGATGTGAAATCTTAGGGCTTAACCCTGAAACTGCATCTAATACTGTGAGGCTCGAGTCCGGGAGAGGTTGGCGGAATTCCAGGTGTAGCGGTGAAATGCGTAGATATCTGGAGGAACACCAGTGGCGAAGGCGGCCAACTGGCCCGGTACTGACACTCAGGGACGAAAGCGTGGGTAGCGAACCGG
## 4086                                                         ACGTAGGTAGCGAGCGTTATTCGGATTTACTGGGCGTAAAGGGTCCGCAGGAGGCCTGCCAAGTCTGATGTGAAAGCCCATGGCTCAACTATGGAACTGCATCGGAAACTGGCGGGCTCGAGTGCACGAGGGGAAAGCGGAATTCCAGGTGTAGCGGTGGAATGCGTAGATATCTGGAGGAACACCGATGGCGAAGGCGGCTTTCTGGCGTGCAACTGACTCTCATGGACGAAAGTGTGGGTAGCAAACAGG
## 4097                                                        CCTAGCGGTCCAAGTCGCATCCATCTTTATTGGGTCTAAAACATCCGTAGCTTGCTTAGTAAGTTCCTTGTGAAATCCTATGTCTCAAGCATAGGGCGGGCAAGGAGTACTGCTGAGCTAGAGACTGGAAGGCGTAGGAAGTACGTTTAAAGTAACGGTAAAATGTGTTAATCTTGGACGGACTCACAATAGCGAAGGCATCCTACGAGGACAGTTCTGACAGTGAGGGATGAAAGCCAGGGTCGCGAAACGG
## 4121                                                   ACGTAAGGTCTAAGTTTTTATCAGAAAAGGCGTAAAGGGTGTGTAGGCAGGATCCCATCGGCAAGGCGTCACCTACGGCGGCATCTATGATGCCGCTGGGAGGCTATGCCTCGGACCCATGGGATGTGAGGTTTAAAGGGGTAAATGGAACTTTTATGTAGCGGTAGAATGCTTTAATATAAAATGGAACACCGGAGGCGAAGGCAATTTACTAATTAAATCTGACGCTGAGACACGAAAGTAAGGGGAGCAAACAGG
## 4133                                                         ACGAGGGGGACAAGCGTTATTCACCATGACTGGGCGTAAAGAGTTCATAGACAGTAAAGTCAGTTTTTTGTTAAAATTTGAAGCAAAACTTTAAAATAGCATTTAATACAGCTTTACTTGAGTTATATACAAGAGATATACAATTTTGTTTGTAAGGGTAAAATCTAACTATAAACAAAGGAATGCCGTTAGCGAAGGCGAGTTTCTAGTAAAAACTGACGTTGTAGAACGAAAGTATGGGTAGCAAACAGG
## 4137                                                                                                    ACAGATTGGACAAGCGTTTTTCATCTTGAATAGGCAGAAAGTGTGATAAGATATGAACATATCAGATATGTTTCTAGAGTTTTATAAAAGTTTTTGGAATAGTTAAAGGAAGGATGAAATTTTCAGATCTTAATTGGAATACCAAAGGTGTAGACACTTTTGCTTTTTTTAACTGACATTGACTCACTAAAGTTTGGGTAGCAAACGGG
## 4151                                                         ACGTAAGGGGCGAGCGTTGTTCGGAATTATTGGGCGTAAAGGGCGTGCAGGCGGTCATGTAAGCCTGTTGTGAAATCCCGGGGCTTAACTCCGGAGGCGCGATGGAAACTGTGTGACTAGAATCTTGGAGGGGGAGTTGGAATTCCTAGTGTAGGGGTGAAATCTGTTGATATTAGGAAGAACACCGGTGGCGAAGGCGAACTCCTGGCCGAAGATTGACGCTGAGGCGCGAAAGCGTAGGTAGCAAACAGG
## 4195                                                         ACAGAGACCTCAAACGTTATCCGGATTTACTGGGCGTAAAGCGTCCGTAGGTGGATTGATAAGTTAGTGGTTAAATCCTGGGGCTTAACTTCAGGACTGCCGCTAATACTGTTAATCTAGGGGTTGGTAGAAGCAAGCGGAACTACCGGTGTAGTCGTAATAAGCGCTGATATCGGTAGGAACACCAAAGGCAAAGGCAGCTTGCTATAACAAACCTGACACTCAAGGACTAAAGCGTGGGGAGCGAATAGG
## 4204                       ACGGGGGGGACCAGTGTTATTCGGAGTAATTAGGCGTAAAGAGTTTGAAGATTGTTTAAATAAAATTAATAGATTTTCTCTTAATTTTTCACATAATAAATGAAAACTTAAAATTATCCAAATATTAAGCAATTTTAAAATGTATTTAAACTTGAGATTATAAGACAATAGAAGAATTTTAAATGTAGAGATCAAATTTTTTAATATTTAAAAGAAAACCAAAAGCGAAGGCCTCTATTGATTATAAATCTGACATTGTAAAACGAAAGTATAGGTATCGAATAGG
## 4207                                                         ACGTAGGTGGCAAGCGTTGTCCGGATTTACTGGGCGTAAAGCGAACGCAGGCGGATCTTTAAGTAGGAAGTGAAAGGTTGGAGCTCAACTCTAACACTGCTCCCTATACTGGAGATCTTGAGTGCCGGAGAGGAAAACGGAACAAGTTGTGTAGCGGTGAAATGCGTTGATATAACTTGGAACACCAATGGCGAAAGCAGTTTTCTGGACGGATACTGACGCTCATGTTCGAAAGCCAAGGTAGCAAACAGG
## 4213                                                               CGGAGGGGGCTAGCGTTATTCATCCTTATTGGGCGTAAAGGGTATGTAGGTAATGAAAAGTGTTTTTTTTTAAATCTCAAATACCAGTTTGAACGGGAAAAAACATTGATTAATTCGGGATGTAGGGGGTTGGCGGTATTACTAATGTAAAGGTAAAATCTAACGAGATTAGTACGAATACCAAAGGCGCAGGCGGCTTTCTTTACCATCTGACACTGAGATACTAAAGCGTGGGGATCAAACAGG
## 4226                                                         ACGTAGGGTGCAAGCGTTAATCGGAATTACTGGGCGTAAAGCGTGCGCAGGCGGTTATGTAAGACAGAGGTGAAATCCCCGGGCTCAACCTGGGAACTGCCTTTGTGACTGCGAGGCTAGAGTATGGCAGAGGGGGGTGGAATTCCACGTGTAGCAGTGAAATGCGTAGAGATGTGGAGGAACACCGATGGCGAAGGCAATCCCCTGGACCTGTACTGACGCTCATGCACGAAAGCGTGGGGAGCAAACAGG
## 4249                                                         ACGTAGGGTGCGAGCGTTAATCGGAATTACTGGGCGTAAAGCGTGCGCAGGCGGTTTCGTAAGACAGAGGTGAAATCCCCGGGCTCAACCTGGGAACTGCCTTTGTGACTGCATAGCTAGAGTACGGCAGAGGGGGATGGAATTCCGCGTGTAGCAGTGAAATGCGTAGATATGCGGAGGAACACCGATGGCGAAGGCAATCCCCTGGACCTGTACTGACGCTCATGCACGAAAGCGTGGGGAGCAAACAGG
## 4659                                                         ACGTAGGGTGCAAGCGTTAATCGGAATTACTGGGCGTAAAGCGTGCGCAGGCGGTTATGTAAGACAGAGGTGAAATCCCCGGGCTCAACCTGGGAACTGCCTTTGTGACTGCGAGGCTAGAGTATGGCAGAGGGGGGTGGAATTCCACGTGTAGCAGTGAAATGCGTAGAGATGTGGAGGAACACCGATGGCGAAGGCAGCCCCCTGGGTCGACACTGACGCTCATGCACGAAAGCGTGGGGAGCAAACAGG
## 4668                                                         ACGTAGGGTGCGAGCGTTAATCGGAATTACTGGGCGTAAAGCGTGCGCAGGCGGTTATGTAAGACAGATGTGAAATCCCCGGGCTCAACCTGGGACCTGCATTTGTGACTGCATAGCTAGAGTACGGTAGAGGGGGATGGAATTCCGCGTGTAGCAGTGAAATGCGTAGATATGCGGAGGAACACCGATGGCGAAGGCAGCTTACTACGCATATATTGACGCTGAGGCACGAAAGCGTGGGGATCAAACAGG
## 4710                                                         ACGGAGGATCCAAGCGTTATCCGGAATCATTGGGTTTAAAGGGTCCGTAGGCGGTTTTATAAGTCAGTGGTGAAATCCGGCAGCTCAACTGTCGAACTGCCATTGATACTGTAGAACTTGAATTACTTGGAAGTAACTAGAATATGTAGTGTAGCGGTGAAATGCTTAGAGATTACATGGAATACCAATTGCGAAGGCAGGTTACTACAAGTGGATTGACGCTGATGGACGAAAGCGTGGGGAGCGAACAGG
## 4873                                                         ACGTAGGGTGCGAGCGTTAATCGGAATTACTGGGCGTAAAGCGTGCGCAGGCGGTTATGTAAGACAGATGTGAAATCCCCGGGCTCAACCTGGGACCTGCATTTGTGACTGCATAGCTAGAGTACGGTAGAGGGGGATGGAATTCCGCGTGTAGCAGTGAAATGCGTAGATATGCGGAGGAACACCGATGGCGAAGGCAGTCCCCTGGGATAAGACTGACGCTCATGCACGAAAGCGTGGGGAGCAAACAGG
## 5025                                                         ACGTAGGGTGCAAGCGTTAATCGGAATTACTGGGCGTAAAGCGTGCGCAGGCGGTTATGTAAGACAGATGTGAAATCCCCGGGCTCAACCTGGGAACTGCATTTGTGACTGCATAGCTAGAGTACGGTAGAGGGGGATGGAATTCCGCGTGTAGCAGTGAAATGCGTAGATATGCGGAGGAACACCGATGGCGAAGGCAGCCTCCTGGGCCAGTACTGACGCTCATGCACGAAAGCGTGGGGAGCAAACAGG
## 5197                                                         ACGTAGGGTGCGAGCGTTAATCGGAATTACTGGGCGTAAAGCGTGCGCAGGCGGTTATATAAGACAGATGTGAAATCCCCGGGCTTAACCTGGGACCTGCATTTGTGACTGTATAGCTAGAGTACGGTAGAGGGGGATGGAATTCCGCGTGTAGCAGTGAAATGCGTAGATATGCGGAGGAACACCGATGGCGAAGGCAGTCCCCTGGGATAAGACTGACGCTCATGCACGAAAGCGTGGGGAGCAAACAGG
## 5258                                                         ACGTAGGGTGCGAGCGTTAATCGGAATTACTGGGCGTAAAGCGTGCGCAGGCGGTTATGCAAGACAGAGGTGAAATCCCCGGGCTCAACCTGGGAACTGCCTTTGTGACTGCATAGCTAGAGTACGGCAGAGGGGGATGGAATTCCGCGTGTAGCAGTGAAATGCGTAGATATGCGGAGGAACACCGATGGCGAAGGCAATCCCCTGGACCTGTACTGACGCTCATGCACGAAAGCGTGGGGAGCAAACAGG
## 5308                                                         ACGGGAGTGGCAAGCGTTATCCGGAATTATTGGGCGTAAAGCGTCCGCAGGCGGTCTTTTAAGTCTGCTGTTAAAGCGTGGAGCTTAACTCCATTTCGGCAGTGGAAACTGGAAGACTAGAGTGTGGTAGGGGCAGAGGGAATTCCCGGTGTAGCGGTGAAATGCGTAGATATCGGGAAGAACACCAGTGGCGAAGGCGCTCTGCTGGGCCATAACTGACGCTCATGGACGAAAGCGTGGGGAGCAAACAGG
## 5656                                                         ACGTAGGGTGCAAGCGTTAATCGGAATTACTGGGCGTAAAGCGTGCGCAGGCGGTTTTGTAAGACAGAGGTGAAATCCCCGGGCTCAACCTGGGAACTGCCTTTGTGACTGCAAGGCTGGAGTACGGCAGAGGGGGATGGAATTCCGCGTGTAGCAGTGAAATGCGTAGATATGCGGAGGAACACCGATGGCGAAGGCAGGTCTCTGGGCCAATACTGACACTGAGGAGCGAAAGCGTGGGGAGCGAACAGG
## 5691                                                         ACGTAGGGTGCGAGCGTTAATCGGAATTACTGGGCGTAAAGCGTGCGCAGGCGGACTTTTAAGCCAGATGTGAAAGCCCCGAGCTTAACTTGGGAATTGCGTTTGGAACTGGGAGTCTAGAGTCTGTCAGAGGGGGATGGAATTCCACGTGTAGCAGTGAAATGCGTAGAGATGTGGAGGAACACCGATGGCGAAGGCAGTCCCCTGGGCCTGCACTGACGCTCATGCACGAAAGCGTGGGGAGCAAACAGG
## 5711                                                         ACAGAGGGTGCAAGCGTTAATCGGAATTACTGGGCGTAAAGCGTGCGTAGACGGTTATCTAAGTCGGATGTGAAATCCCCGGGCTCAACCTGGGAACTGCCTTTGTGACTGCGAGGCTAGAGTATGGCAGAGGGGGGTGGAATTCCACGTGTAGCAGTGAAATGCGTAGAGATGTGGAGGAACACCGATGGCGAAGGCAGCCCCCTGGGCCAATACTGACACTGAGGAGCGAAAGCGTGGGGAGCGAACAGG
## 5802                                                         ACGTAGGGTGCAAGCGTTAATCGGAATTACTGGGCGTAAAGCGTGCGCAGGCGGTTATGTAAGACAGAGGTGAAATCCCCGGGCTCAACCTGGGAACGGCCTTTGTGACTGCATAGCTAGAGTACGGTAGAGGGGGATGGAATTCCGCGTGTAGCAGTGAAATGCGTAGATATGCGGAGGAACACCGATGGCGAAGGCAGCCTCCTGGGCCAGTACTGACGCTCATGCACGAAAGCGTGGGGAGCAAACAGG
## 5854                                                         ACGTAGGGTGCAAGCGTTAATCGGAATTACTGGGCGTAAAGCGTGCGCAGGCGGTTTCGTAAGACAGACGTGAAATCCCCGGGCTCAACCTGGGAACTGCGTTTGTGACTGCGAGGCTAGAGTATGGCAGAGGGGGGTGGAATTCCACGTGTAGCAGTGAAATGCGTAGAGATGTGGAGGAACACCGATGGCGAAGGCAGCCCCCTGGGTCGACACTGACGCTCATGCACGAAAGCGTGGGGAGCAAACAGG
## 6106                                                         ACGAAGGATGCTAGCGTTACTCGGAATTACTGGGCGTAAAGAATGCGTAGGCGGATTATTAAGTTGATAGTGAAATCCCTGAGCCTAACTTGGGAACTGCTGTCAAAACTGATAGTCTAGAATATGGTAGAGAGTAAGGGAATTCTTGGTGTAGGGGTAAAATCCGTAGATATCAGGAGGAAGACCGGAAGCGAAAGCGCTTACTTGGGCCATCATTGACGCTGAGATATGAAAGCGTGGGGATCGAACAGG
## 6115                                                         ACGTAGGGTGCAAGCGTTAATCGGAATTACTGGGCGTAAAGCGTGCGCAGGCGGTTTCGTAAGACAGACGTGAAATCCCCGGGCTCAACCTGGGAACTGCGTTTGTGACTGCGAGGCTAGAGTATGGCAGAGGGGGGTGGAATTCCACGTGTAGCAGTGAAATGCGTAGAGATGTGGAGGAACACCGATGGCGAAGGCAATCCCCTGGACCTGTACTGACGCTCATGCACGAAAGCGTGGGGAGCAAACAGG
## 6336                                                         ACGGAGGGTGCGAGCGTTATCCGGATTCACTGGGTTTAAAGGGTGCGTAGGCGGGCTAGTAAGTCAGTGGTGAAATCCCCGAGCTTAACTTGGGAACTGCCGTTGATACTATTAGTCTTGAATATCGTGGAGGTAAGCGGAATATGTCATGTAGCGGTGAAATGCTTAGATATGACATAGAACACCAATTGCGAAGGCAGCTTGCTACACGGTCATTGACACTGAGGCACGAAAGCGTGGGGAGCAAACAGG
## 6598                                                         ACGTAGGGTGCAAGCGTTAATCGGAATTACTGGGCGTAAAGCGTGCGCAGGCGGTTATATAAGTCAGATGTGAAATCCCCGGGCTCAACCTGGGAACTGCATTTGAGACTGTATAGCTAGAGTACGGTAGAGGGGGATGGAATTCCGCGTGTAGCAGTGAAATGCGTAGATATGCGGAGGAACACCGATGGCGAAGGCAGCTTACTACGCATATATTGACGCTGAGGCACGAAAGCGTGGGGATCAAACAGG
## 6644                                                         ACGGAGGGTGCAAGCGTTATCCGGAATCATTGGGTTTAAAGGGTCCGCAGGCGGATTTATAAGTCAGTGGTGAAAGCCTACAGCTCAACTGTAGAACTGCCATTGATACTGTAAGTCTTGAATTCGGTCGAAGTGGGCGGAATGTGTAGTGTAGCGGTGAAATGCTTAGATATTACACAGAACACCGATAGCGAAGGCAGCTCACTAGGCCTGGATTGACGCTCAGGGACGAAAGCGTGGGGAGCAAACAGG
## 6667                                                         ACGTAGGGTGCAAGCGTTAATCGGAATTACTGGGCGTAAAGCGTGCGCAGGCGGTTTCGTAAGACAGACGTGAAATCCCCGGGCTCAACCTGGGAACTGCGTTTGTGACTGCGAGGCTAGAGTATGGCAGAGGGGGGTGGAATTCCACGTGTAGCAGTGAAATGCGTAGAGATGTGGAGGAACACCGATGGCGAAGGCAGTCCCCTGGGATAAGACTGACGCTCATGCACGAAAGCGTGGGGAGCAAACAGG
## 7077                                                         ACGTAGGGTGCGAGCGTTAATCGGAATTACTGGGCGTAAAGCGTGCGCAGGCGGACTTTTAAGCCAGATGTGAAAGCCCCGAGCTTAACTTGGGAATTGCGTTTGGAACTGGGAGTCTAGAGTCTGTCAGAGGGGGATGGAATTCCACGTGTAGCAGTGAAATGCGTAGAGATGTGGAGGAACACCGATGGCGAAGGCAGCCCCCTGGGCCAATACTGACGCTCATGCACGAAAGCGTGGGGAGCAAACAGG
## 7178                                                         ACAGAGGGTGCAAGCGTTAATCGGAATTACTGGGCGTAAAGCGTGCGTAGACGGTTATCTAAGTCGGATGTGAAATCCCCGGGCTCAACCTGGGAACTGCCTTTGTGACTGCGAGGCTAGAGTATGGCAGAGGGGGGTGGAATTCCACGTGTAGCAGTGAAATGCGTAGAGATGTGGAGGAACACCGATGGCGAAGGCAGTCCCCTGGGATAAGACTGACGCTCATGCACGAAAGCGTGGGGAGCAAACAGG
## 7201                                                         ACGTAGGGTGCGAGCGTTAATCGGAATTACTGGGCGTAAAGCGTGCGCAGGCGGTTATGTAAGACAGTTGTGAAATCCCCGGGCTCAACCTGGGAATTGCATCTGTGACTGCATAGCTAGAGTACGGTAGAGGGGGATGGAATTCCGCGTGTAGCAGTGAAATGCGTAGATATGCGGAGGAACACCGATGGCGAAGGCAATCCCCTGGGCCTGCACTGACGCTCATGCACGAAAGCGTGGGGAGCAAACAGG
## 7572                                                         ACGTAGGGTGCAAGCGTTAATCGGAATTACTGGGCGTAAAGCGTGCGCAGGCGGTTATGTAAGACAGAGGTGAAATCCCCGGGCTCAACCTGGGAACGGCCTTTGTGACTGCATAGCTAGAGTACGGTAGAGGGGGATGGAATTCCGCGTGTAGCAGTGAAATGCGTAGATATGCGGAGGAACACCGATGGCGAAGGCAGTCCCCTGGGATAAGACTGACGCTCATGCACGAAAGCGTGGGGAGCAAACAGG
## 7798                                                         ACGTAGGGTGCGAGCGTTAATCGGAATTACTGGGCGTAAAGCGTGCGCAGGCGGTTTGATAAGACAGGCGTGAAATCCCCGGGCTTAACCTGGGAACTGCGCTTGTGACTGTCTCACTAGAGTACGGCAGAGGGGGGTGGAATTCCACGTGTAGCAGTGAAATGCGTAGAGATGTGGAGGAACACCGATGGCGAAGGCAGCCCCCTGGGCCGATACTGACGCTCATGCACGAAAGCGTGGGTAGCAAACAGG
## 7893                                                         ACGTAGGGTGCAAGCGTTAATCGGAATTACTGGGCGTAAAGCGTGCGCAGGCGGTTATGCAAGACAGATGTGAAATCCCCGGGCTCAACCTGGGAACTGCATTTGTGACTGCATAGCTAGAGTACGGTAGAGGGGGATGGAATTCCGCGTGTAGCAGTGAAATGCGTAGATATGCGGAGGAACACCGATGGCGAAGGCAGCCTCCTGGGCCAGTACTGACGCTCATGCACGAAAGCGTGGGGAGCAAACAGG
## 7974                                                         ACATAGGGTGCAAGCGTTGTCCGGAATTATTGGGCGTAAAGAGCTCGTAGGTCGTTTGACACGTCGGATGTGAAAATCTGAGGCTTAACCTCAGACCTGCATTCGATACGGTCAAACTAGAGTGTGGTAGGGGAGACTGGAATTCCTGGTGTAGCGGTGGAATGCGCAGATATCAGGAGGAACACCAATGGCGAAGGCAGGTCTCTGGGCCATAACTGACACTGAGGAGCGAAAGTGCGGGGAGCGAACAGG
##      abundance forward reverse nmatch nmismatch nindel prefer accept
## 1         3694       1       3     87         0      0      2   TRUE
## 2         3356       2       4     87         0      0      2   TRUE
## 3         3026       3       6     87         0      0      2   TRUE
## 4         2873       4       7     87         0      0      2   TRUE
## 5         2721       6       1     87         0      0      2   TRUE
## 6         2448       5       8     87         0      0      2   TRUE
## 7         1558       7       5     87         0      0      2   TRUE
## 8         1530      10       9     87         0      0      2   TRUE
## 9         1232       8      12     87         0      0      2   TRUE
## 10        1221      11       2     87         0      0      2   TRUE
## 11        1210       9      10     87         0      0      2   TRUE
## 12         977      27       2     87         0      0      2   TRUE
## 13         936      18       1     87         0      0      2   TRUE
## 14         871      12      14     87         0      0      2   TRUE
## 15         825      38      18     87         0      0      2   TRUE
## 16         800      13      16     87         0      0      2   TRUE
## 17         795      24      25     87         0      0      2   TRUE
## 18         782      40       1     87         0      0      2   TRUE
## 19         777      22      20     87         0      0      2   TRUE
## 20         771      15       5     87         0      0      2   TRUE
## 21         753      17      26     87         0      0      2   TRUE
## 22         751      14      19     87         0      0      2   TRUE
## 23         740      23      17     87         0      0      2   TRUE
## 24         734      33       2     87         0      0      2   TRUE
## 25         727      16      24     87         0      0      2   TRUE
## 26         721      21      23     87         0      0      2   TRUE
## 27         686      19      22     87         0      0      2   TRUE
## 28         664      20      15     87         0      0      2   TRUE
## 29         662      25      13     87         0      0      2   TRUE
## 30         659      43      29     87         0      0      2   TRUE
## 31         657      36      31     87         0      0      2   TRUE
## 32         635      29      21     87         0      0      2   TRUE
## 33         626      28      32     87         0      0      2   TRUE
## 34         620      47      33     87         0      0      2   TRUE
## 35         616      26      30     87         0      0      2   TRUE
## 36         593     924      11     87         0      0      2   TRUE
## 37         578      44       2     87         0      0      2   TRUE
## 38         571      49      11     87         0      0      2   TRUE
## 39         548      30      15     87         0      0      2   TRUE
## 40         538      39      34     87         0      0      2   TRUE
## 41         532      32      36     87         0      0      2   TRUE
## 42         522      34      39     87         0      0      2   TRUE
## 43         520      31      35     87         0      0      2   TRUE
## 44         519      35      42     87         0      0      2   TRUE
## 45         502      37      48     87         0      0      1   TRUE
## 46         501      57      41     87         0      0      2   TRUE
## 47         466      41      47     87         0      0      2   TRUE
## 48         422      60      40     87         0      0      2   TRUE
## 49         422      42      51     87         0      0      2   TRUE
## 50         422      56      44     87         0      0      2   TRUE
## 51         417      63      49     87         0      0      2   TRUE
## 52         394      53      52     87         0      0      2   TRUE
## 53         393      59      45     87         0      0      2   TRUE
## 54         362      51      53     87         0      0      2   TRUE
## 55         354      48      13     87         0      0      2   TRUE
## 56         353      64      54     87         0      0      2   TRUE
## 57         340      55      43     87         0      0      2   TRUE
## 58         338      68      28     87         0      0      2   TRUE
## 59         336      45      14     87         0      0      2   TRUE
## 60         328      54      64     87         0      0      2   TRUE
## 61         318      75      56     87         0      0      2   TRUE
## 62         315      52      65     87         0      0      2   TRUE
## 63         312      50      59     87         0      0      2   TRUE
## 64         307      77      63     87         0      0      2   TRUE
## 65         306      46      55     86         0      0      2   TRUE
## 66         286      62      58     87         0      0      2   TRUE
## 67         279      76      67     86         0      0      2   TRUE
## 68         263      58     655     87         0      0      2   TRUE
## 69         261     101       1     87         0      0      2   TRUE
## 70         257     928      38     87         0      0      2   TRUE
## 71         255     188      72     87         0      0      2   TRUE
## 72         249    1501       5     87         0      0      2   TRUE
## 73         247      85      70     87         0      0      2   TRUE
## 74         247      66      38     87         0      0      2   TRUE
## 75         244      87      50     87         0      0      2   TRUE
## 76         241      80      71     87         0      0      2   TRUE
## 77         240    1168       1     87         0      0      2   TRUE
## 78         239      61      27     87         0      0      2   TRUE
## 79         236      65      46     87         0      0      2   TRUE
## 80         235      67      76     86         0      0      2   TRUE
## 81         234      69      80     87         0      0      2   TRUE
## 82         233      70      77     87         0      0      2   TRUE
## 83         216      73      75     86         0      0      2   TRUE
## 84         215      72      68     87         0      0      2   TRUE
## 85         215      74      88     87         0      0      1   TRUE
## 86         210    1014      61     87         0      0      2   TRUE
## 87         206      71      81     87         0      0      2   TRUE
## 88         205    1051     667     87         0      0      2   TRUE
## 89         200      86      82     87         0      0      2   TRUE
## 90         199     121      27     87         0      0      2   TRUE
## 91         199      81      87     87         0      0      2   TRUE
## 92         198    1414      37     87         0      0      2   TRUE
## 93         198      92      93     87         0      0      2   TRUE
## 94         197    1385      28     87         0      0      2   TRUE
## 95         195      79      57     87         0      0      2   TRUE
## 96         195      93      86     87         0      0      2   TRUE
## 97         195    1709       3     87         0      0      2   TRUE
## 98         188    1095      19     87         0      0      2   TRUE
## 99         187      83     102     87         0      0      2   TRUE
## 100        187      78      90     87         0      0      2   TRUE
## 101        184     103      84     87         0      0      2   TRUE
## 102        182     952       9     87         0      0      2   TRUE
## 103        180      82      21     87         0      0      2   TRUE
## 104        177      99      78     87         0      0      2   TRUE
## 105        177      84      91     85         0      0      2   TRUE
## 106        175      94      99     87         0      0      2   TRUE
## 107        174      95      97     87         0      0      2   TRUE
## 108        172      91     103     87         0      0      2   TRUE
## 109        172     123     125     87         0      0      2   TRUE
## 110        170     132     105     86         0      0      2   TRUE
## 111        169    1216      13     87         0      0      2   TRUE
## 112        168     156       1     87         0      0      2   TRUE
## 113        166     140      98     87         0      0      2   TRUE
## 114        166     111     115     87         0      0      2   TRUE
## 115        165      96      96     87         0      0      2   TRUE
## 116        164     102     104     87         0      0      2   TRUE
## 117        163    1482       5     87         0      0      2   TRUE
## 118        163      89     116     87         0      0      1   TRUE
## 119        161      97     101     87         0      0      2   TRUE
## 120        159    1627       1     87         0      0      2   TRUE
## 121        158     112      66     87         0      0      2   TRUE
## 122        157     171     128     87         0      0      2   TRUE
## 123        157     152      73     87         0      0      2   TRUE
## 124        156     124     107     87         0      0      2   TRUE
## 125        156    1419      50     87         0      0      2   TRUE
## 126        155     104     109     87         0      0      2   TRUE
## 127        155    1969       5     87         0      0      2   TRUE
## 128        154      88      83     87         0      0      2   TRUE
## 129        153     138     130     87         0      0      2   TRUE
## 130        153      98     108     87         0      0      2   TRUE
## 131        151     109      46     87         0      0      2   TRUE
## 132        148     117      23     87         0      0      2   TRUE
## 133        147     115     119     87         0      0      2   TRUE
## 134        146     120     120     87         0      0      2   TRUE
## 135        146     108     113     87         0      0      2   TRUE
## 136        145     295      62     87         0      0      2   TRUE
## 137        145     118     820     87         0      0      2   TRUE
## 138        145     170     100     87         0      0      2   TRUE
## 139        142     113     132     87         0      0      2   TRUE
## 140        142     128     111     87         0      0      2   TRUE
## 141        142      90      85     87         0      0      2   TRUE
## 142        141     146     129     87         0      0      2   TRUE
## 143        138     105      46     87         0      0      2   TRUE
## 144        138     127      95     87         0      0      2   TRUE
## 145        137     110     122     87         0      0      2   TRUE
## 146        137     201     124     87         0      0      2   TRUE
## 147        135     107      74     87         0      0      2   TRUE
## 148        135     166      89     87         0      0      2   TRUE
## 149        134     147     127     87         0      0      2   TRUE
## 150        134     969      37     87         0      0      2   TRUE
## 151        134     114     121     87         0      0      2   TRUE
## 152        133    1390      27     87         0      0      2   TRUE
## 153        133     106     126     87         0      0      1   TRUE
## 154        133    1401      61     87         0      0      2   TRUE
## 155        133     100     114     87         0      0      2   TRUE
## 156        132    1145      57     87         0      0      2   TRUE
## 157        132     157     139     87         0      0      2   TRUE
## 158        131     151     136     87         0      0      2   TRUE
## 159        131    1743      27     87         0      0      2   TRUE
## 160        130     116     117     86         0      0      2   TRUE
## 161        130     122     155     87         0      0      1   TRUE
## 162        128     144     152     87         0      0      2   TRUE
## 163        124     137      43     87         0      0      2   TRUE
## 164        122     977     299     87         0      0      1   TRUE
## 165        122     148     133     87         0      0      2   TRUE
## 166        122     149     142     87         0      0      2   TRUE
## 167        122    2165      17     87         0      0      2   TRUE
## 168        122    1319       1     87         0      0      2   TRUE
## 169        121    1443      16     87         0      0      2   TRUE
## 170        120     183     904     87         0      0      2   TRUE
## 171        119    2081      10     87         0      0      2   TRUE
## 172        117     164     977     87         0      0      2   TRUE
## 173        117     135     131     86         0      0      2   TRUE
## 174        117    1277     895     87         0      0      2   TRUE
## 175        117     126     145     87         0      0      1   TRUE
## 176        116     142     123     87         0      0      2   TRUE
## 177        115     205     161     87         0      0      2   TRUE
## 178        114     150     146     87         0      0      2   TRUE
## 179        114     141     118     87         0      0      2   TRUE
## 180        114     134     106     87         0      0      2   TRUE
## 181        113    1719       2     87         0      0      2   TRUE
## 182        113     200     137     87         0      0      2   TRUE
## 183        113    1147       9     87         0      0      2   TRUE
## 184        113     173     164     87         0      0      2   TRUE
## 185        112    1191     144     87         0      0      2   TRUE
## 186        112     161     135     87         0      0      2   TRUE
## 187        111    1355      16     87         0      0      2   TRUE
## 188        111     180     197     87         0      0      1   TRUE
## 189        111    1514      21     87         0      0      2   TRUE
## 190        111     131    1134     87         0      0      2   TRUE
## 191        110     129      79     87         0      0      2   TRUE
## 192        109     179     163     87         0      0      2   TRUE
## 193        109     225     165     87         0      0      2   TRUE
## 194        109     130      60     87         0      0      2   TRUE
## 195        108    2231      22     87         0      0      2   TRUE
## 196        108     258      92     87         0      0      2   TRUE
## 197        107     172     204     87         0      0      1   TRUE
## 198        107    1073      79     87         0      0      2   TRUE
## 199        107     143     173     87         0      0      1   TRUE
## 200        107     125     147     87         0      0      1   TRUE
## 201        107     210     935     87         0      0      2   TRUE
## 202        106     175     149     87         0      0      2   TRUE
## 203        106     331     159     87         0      0      2   TRUE
## 204        105     158     148     87         0      0      2   TRUE
## 205        105     209     112     87         0      0      2   TRUE
## 206        105    2062       2     87         0      0      2   TRUE
## 207        105     154     151     87         0      0      2   TRUE
## 208        104     216     632     87         0      0      2   TRUE
## 209        104    1784      11     87         0      0      2   TRUE
## 210        103     190     168     87         0      0      2   TRUE
## 211        103    1773    1713     87         0      0      2   TRUE
## 212        102     159     866     87         0      0      2   TRUE
## 213        102     176     141     87         0      0      2   TRUE
## 214        102    1655      66     87         0      0      2   TRUE
## 215        101     169     158     87         0      0      2   TRUE
## 216        101     162     178     87         0      0      2   TRUE
## 217        100     153     160     87         0      0      2   TRUE
## 218        100     133     169     87         0      0      1   TRUE
## 219        100     139     157     87         0      0      2   TRUE
## 220        100     178     166     87         0      0      2   TRUE
## 221        100     182     830     87         0      0      2   TRUE
## 222         99     165     179     87         0      0      2   TRUE
## 223         98    2082       9     87         0      0      2   TRUE
## 224         97     174     189     87         0      0      1   TRUE
## 225         97     136     638     87         0      0      2   TRUE
## 226         97     160     646     87         0      0      2   TRUE
## 227         97    1416      22     87         0      0      2   TRUE
## 228         96    1516       1     87         0      0      2   TRUE
## 229         96     204    1050     87         0      0      2   TRUE
## 230         95     192     184     87         0      0      2   TRUE
## 231         95     187     150     87         0      0      2   TRUE
## 232         95     237      69     87         0      0      2   TRUE
## 233         95     186     154     87         0      0      2   TRUE
## 234         95     155     153     87         0      0      2   TRUE
## 235         95     184     175     87         0      0      2   TRUE
## 236         94     119     172     87         0      0      1   TRUE
## 237         94     935     192     87         0      0      2   TRUE
## 238         94     199     190     87         0      0      2   TRUE
## 239         93    2229    1046     87         0      0      2   TRUE
## 240         93     228     200     87         0      0      2   TRUE
## 241         93     189     187     87         0      0      2   TRUE
## 242         91     271     208     87         0      0      2   TRUE
## 243         91     163     162     83         0      0      2   TRUE
## 244         91     203     194     85         0      0      2   TRUE
## 245         91    1163     185     87         0      0      2   TRUE
## 246         90     215     188     87         0      0      2   TRUE
## 247         89     919     186     87         0      0      2   TRUE
## 248         88     288     213     87         0      0      2   TRUE
## 249         88     198     237     87         0      0      2   TRUE
## 250         88    1243     649     87         0      0      1   TRUE
## 251         88     247     721     87         0      0      2   TRUE
## 252         88    1263      28     87         0      0      2   TRUE
## 253         88     167     183     83         0      0      2   TRUE
## 254         87     251     171     87         0      0      2   TRUE
## 255         86     304     249     87         0      0      2   TRUE
## 256         86    1044     947     87         0      0      2   TRUE
## 257         85     202     201     87         0      0      2   TRUE
## 258         85     177     170     87         0      0      2   TRUE
## 259         84     253     205     87         0      0      2   TRUE
## 260         84       1     853     87         0      0      1   TRUE
## 261         84     181     215     88         0      0      1   TRUE
## 262         84    1601      58     87         0      0      2   TRUE
## 263         84     197    1184     87         0      0      2   TRUE
## 264         84     218     193     87         0      0      2   TRUE
## 265         83     221     167     87         0      0      2   TRUE
## 266         83     196     225     87         0      0      1   TRUE
## 267         82     223     176     87         0      0      2   TRUE
## 268         81     226     207     87         0      0      2   TRUE
## 269         81     194     217     87         0      0      1   TRUE
## 270         81     310     182     87         0      0      2   TRUE
## 271         81     955    1079     87         0      0      2   TRUE
## 272         80     168     174     86         0      0      2   TRUE
## 273         79     978    1505     87         0      0      2   TRUE
## 274         79     222     211     87         0      0      2   TRUE
## 275         79     195     219     87         0      0      1   TRUE
## 276         79     238     673     87         0      0      2   TRUE
## 277         79     929      37     87         0      0      2   TRUE
## 278         78    1432     921     87         0      0      2   TRUE
## 279         78     942     110     86         0      0      2   TRUE
## 280         78     294     220     87         0      0      2   TRUE
## 281         78    1190      94     87         0      0      2   TRUE
## 282         78     207     236     87         0      0      1   TRUE
## 283         78     252     250     87         0      0      2   TRUE
## 284         78     249     221     87         0      0      2   TRUE
## 285         77     264     199     87         0      0      2   TRUE
## 286         77     208     214     87         0      0      2   TRUE
## 287         77     185     110     86         0      0      2   TRUE
## 288         77     193     206     83         0      0      2   TRUE
## 289         77     347    1206     87         0      0      2   TRUE
## 290         76    1560       9     87         0      0      2   TRUE
## 291         76     211     196     87         0      0      2   TRUE
## 293         75     316     140     87         0      0      2   TRUE
## 294         75     236     216     87         0      0      2   TRUE
## 295         75    1015      60     87         0      0      2   TRUE
## 296         74    1086     802     87         0      0      2   TRUE
## 297         74     273    1233     87         0      0      2   TRUE
## 298         74     280      27     87         0      0      2   TRUE
## 299         73     227     229     87         0      0      2   TRUE
## 300         73    1649     841     87         0      0      2   TRUE
## 301         73    1863    1032     87         0      0      2   TRUE
## 302         73     262     198     87         0      0      2   TRUE
## 303         72     921     921     87         0      0      2   TRUE
## 304         72     212     134     87         0      0      2   TRUE
## 305         72    1182      94     87         0      0      2   TRUE
## 306         71     268     212     87         0      0      2   TRUE
## 307         71       6    1486     87         0      0      1   TRUE
## 308         71     278     263     87         0      0      2   TRUE
## 309         71     231     231     87         0      0      2   TRUE
## 310         71     191     209     87         0      0      2   TRUE
## 311         71     293     350     87         0      0      1   TRUE
## 312         71     284     242     87         0      0      2   TRUE
## 313         71     217     218     87         0      0      2   TRUE
## 314         71     235     235     87         0      0      2   TRUE
## 315         71      98    1083     87         0      0      1   TRUE
## 316         70     246     851     87         0      0      1   TRUE
## 317         70     286     234     87         0      0      2   TRUE
## 318         70     214     233     87         0      0      1   TRUE
## 319         70     241     156     87         0      0      2   TRUE
## 320         70     239     246     87         0      0      2   TRUE
## 321         70     234     995     87         0      0      2   TRUE
## 322         69     352     659     87         0      0      2   TRUE
## 323         69       6     633     87         0      0      1   TRUE
## 324         69    1779    1432     87         0      0      2   TRUE
## 325         68    1510      62     87         0      0      2   TRUE
## 326         68    1964    1059     87         0      0      2   TRUE
## 327         68    1705      78     87         0      0      2   TRUE
## 328         68     145     223     87         0      0      1   TRUE
## 329         67     301     191     87         0      0      2   TRUE
## 330         67     320     283     87         0      0      2   TRUE
## 331         67     302     760     87         0      0      2   TRUE
## 332         67     213     143     87         0      0      2   TRUE
## 333         66     330     270     87         0      0      2   TRUE
## 334         66     248     238     87         0      0      2   TRUE
## 335         66     206     241     87         0      0      1   TRUE
## 336         66     232     631     87         0      0      1   TRUE
## 337         66     244     251     87         0      0      2   TRUE
## 338         65     220     230     86         0      0      2   TRUE
## 339         65    1499       1     87         0      0      2   TRUE
## 340         65     321     210     87         0      0      2   TRUE
## 341         65    1018      69     87         0      0      2   TRUE
## 342         65     412     239     87         0      0      2   TRUE
## 343         65     233     222     87         0      0      2   TRUE
## 344         65     240     195     87         0      0      2   TRUE
## 345         64     224     232     87         0      0      2   TRUE
## 346         64    1036     277     87         0      0      2   TRUE
## 347         64     270    1131     87         0      0      1   TRUE
## 348         64     979      74     87         0      0      2   TRUE
## 349         64     283     273     87         0      0      1   TRUE
## 350         63     985      68     87         0      0      2   TRUE
## 351         63    1053      20     87         0      0      2   TRUE
## 352         63    1064     692     87         0      0      2   TRUE
## 353         63     272     224     87         0      0      2   TRUE
## 354         63     219     243     87         0      0      2   TRUE
## 355         63     314     252     87         0      0      2   TRUE
## 356         63     990    1043     87         0      0      2   TRUE
## 357         63     378     343     87         0      0      2   TRUE
## 358         62     447     383     87         0      0      2   TRUE
## 359         62     242    1351     87         0      0      2   TRUE
## 360         61     306     272     87         0      0      2   TRUE
## 361         61     367     266     87         0      0      2   TRUE
## 362         61    1255     711     87         0      0      2   TRUE
## 363         61    1558     203     87         0      0      2   TRUE
## 364         61     351     309     87         0      0      2   TRUE
## 365         60    1916     735     87         0      0      2   TRUE
## 366         60     275     658     87         0      0      1   TRUE
## 367         60     947     720     87         0      0      2   TRUE
## 368         60    1288     177     87         0      0      2   TRUE
## 369         60     281     253     87         0      0      2   TRUE
## 370         60     265     245     87         0      0      2   TRUE
## 371         60     245     240     88         0      0      2   TRUE
## 373         59     296     259     87         0      0      2   TRUE
## 374         59     440     281     87         0      0      2   TRUE
## 375         58     300     286     87         0      0      2   TRUE
## 376         58    1096      60     87         0      0      2   TRUE
## 377         58     255     247     87         0      0      2   TRUE
## 378         58     260     256     88         0      0      2   TRUE
## 379         58     340     181     87         0      0      2   TRUE
## 380         58    1986    1418     87         0      0      2   TRUE
## 381         57     259     268     87         0      0      1   TRUE
## 382         57     307     255     87         0      0      2   TRUE
## 383         57     277     290     87         0      0      1   TRUE
## 384         57    1977      69     87         0      0      2   TRUE
## 385         57     958    1720     87         0      0      2   TRUE
## 386         56     266     140     87         0      0      2   TRUE
## 387         56    1528      73     87         0      0      2   TRUE
## 388         56       2    1815     87         0      0      1   TRUE
## 389         56     303     983     87         0      0      1   TRUE
## 390         56     256     282     87         0      0      1   TRUE
## 392         56     230     258     87         0      0      1   TRUE
## 393         55     349     388     87         0      0      1   TRUE
## 394         55     229     262     87         0      0      1   TRUE
## 395         55    1879     134     87         0      0      2   TRUE
## 396         55     335     301     87         0      0      2   TRUE
## 397         55     932     865     87         0      0      2   TRUE
## 398         55     119    1474     87         0      0      1   TRUE
## 399         55    1119     877     87         0      0      1   TRUE
## 400         54     292     685     87         0      0      2   TRUE
## 401         54     357     288     88         0      0      2   TRUE
## 402         54    1504     138     87         0      0      2   TRUE
## 403         54      10     948     87         0      0      1   TRUE
## 404         54     473     358     87         0      0      2   TRUE
## 405         54     404     318     87         0      0      2   TRUE
## 407         54     282     274     87         0      0      1   TRUE
## 408         54     290     269     87         0      0      2   TRUE
## 409         53     299     285     87         0      0      2   TRUE
## 410         53    1219     728     87         0      0      2   TRUE
## 411         53    2239     112     87         0      0      2   TRUE
## 412         53     279     227     87         0      0      2   TRUE
## 413         53    1698      11     87         0      0      2   TRUE
## 414         53     243     244     87         0      0      2   TRUE
## 415         53    2034     177     87         0      0      2   TRUE
## 416         53     358     265     87         0      0      2   TRUE
## 417         52     324     303     88         0      0      2   TRUE
## 418         52      11    1465     87         0      0      1   TRUE
## 419         52     250    1266     87         0      0      2   TRUE
## 420         52    2068      60     87         0      0      2   TRUE
## 421         52     313     291     87         0      0      2   TRUE
## 422         52    1900       2     87         0      0      2   TRUE
## 423         52     309     261     87         0      0      2   TRUE
## 424         51       1    1388     87         0      0      1   TRUE
## 425         51     982      60     87         0      0      2   TRUE
## 426         51     285     275     87         0      0      2   TRUE
## 427         51    1487      34     87         0      0      2   TRUE
## 428         51    1076     888     87         0      0      2   TRUE
## 429         51      21    1019     87         0      0      1   TRUE
## 430         51     274     287     87         0      0      1   TRUE
## 431         51     287     300     87         0      0      1   TRUE
## 432         51     338     342     87         0      0      1   TRUE
## 433         51    1241     882     87         0      0      2   TRUE
## 434         51     145    1152     87         0      0      1   TRUE
## 435         50    1948     802     87         0      0      2   TRUE
## 436         50    2076    1058     87         0      0      2   TRUE
## 437         50     377     289     87         0      0      2   TRUE
## 439         49    2178     106     87         0      0      2   TRUE
## 440         49     334     780     87         0      0      2   TRUE
## 441         49    2051      28     87         0      0      2   TRUE
## 442         49     448     180     87         0      0      2   TRUE
## 443         49     477     310     87         0      0      2   TRUE
## 444         49     315     311     87         0      0      2   TRUE
## 445         49     267     674     86         0      0      1   TRUE
## 447         48     361     825     87         0      0      1   TRUE
## 448         48     319     292     87         0      0      2   TRUE
## 449         48    1059      92     87         0      0      2   TRUE
## 450         48     430     409     87         0      0      2   TRUE
## 451         48     269     276     87         0      0      2   TRUE
## 452         48     998     202     87         0      0      2   TRUE
## 453         48     254     260     88         0      0      2   TRUE
## 454         48     429     320     87         0      0      2   TRUE
## 455         48     333     308     87         0      0      2   TRUE
## 456         48     276     317     87         0      0      1   TRUE
## 457         48     263     271     87         0      0      2   TRUE
## 458         48     466     347     87         0      0      2   TRUE
## 459         48     966     636     87         0      0      2   TRUE
## 460         48     567     399     87         0      0      2   TRUE
## 461         47      18    1486     87         0      0      1   TRUE
## 462         47     346     280     87         0      0      2   TRUE
## 463         47    1100      83     87         0      0      2   TRUE
## 464         47     308     316     87         0      0      1   TRUE
## 466         47     322     278     87         0      0      2   TRUE
## 467         47     257     707     87         0      0      1   TRUE
## 468         47     972    1646     87         0      0      2   TRUE
## 469         47     405     357     87         0      0      2   TRUE
## 470         47     380     408     87         0      0      1   TRUE
## 471         47    1146    1793     87         0      0      2   TRUE
## 472         46     305     313     87         0      0      1   TRUE
## 473         46    1965     635     87         0      0      2   TRUE
## 474         46     298     264     88         0      0      2   TRUE
## 475         46     375     295     87         0      0      2   TRUE
## 476         46     936     968     87         0      0      2   TRUE
## 478         46    2136    1752     87         0      0      2   TRUE
## 480         45     344     279     87         0      0      2   TRUE
## 481         45    1005    1572     87         0      0      2   TRUE
## 482         45     363     389     87         0      0      1   TRUE
## 483         45     383     360     87         0      0      2   TRUE
## 484         45    1198      69     87         0      0      2   TRUE
## 485         45     399     341     87         0      0      2   TRUE
## 486         45    1509    1719     87         0      0      2   TRUE
## 487         44    1956       2     87         0      0      2   TRUE
## 488         44     427      85     87         0      0      2   TRUE
## 490         44     317     138     87         0      0      2   TRUE
## 491         44     437     325     87         0      0      2   TRUE
## 492         44    1530     100     87         0      0      2   TRUE
## 493         44      11    1423     87         0      0      1   TRUE
## 494         44     968     293     87         0      0      2   TRUE
## 495         44     326     334     87         0      0      1   TRUE
## 496         43    1137    1215     87         0      0      1   TRUE
## 497         43     354     943     87         0      0      2   TRUE
## 498         43     371     321     87         0      0      2   TRUE
## 499         43     311     298     87         0      0      2   TRUE
## 500         43       1    1194     87         0      0      1   TRUE
## 501         43    1308    1033     87         0      0      1   TRUE
## 502         43    1643    1743     87         0      0      1   TRUE
## 503         43     369     335     87         0      0      2   TRUE
## 504         43     486     361     87         0      0      2   TRUE
## 505         43     328     284     87         0      0      2   TRUE
## 506         43     432     346     87         0      0      2   TRUE
## 508         43     426     747     87         0      0      1   TRUE
## 510         42     291     340     87         0      0      1   TRUE
## 511         42     327     999     87         0      0      2   TRUE
## 512         42    1298    1103     87         0      0      2   TRUE
## 513         42     933     294     87         0      0      2   TRUE
## 514         42     421     381     87         0      0      2   TRUE
## 515         42     409     634     87         0      0      2   TRUE
## 516         42     336     336     87         0      0      1   TRUE
## 517         42     385     344     87         0      0      2   TRUE
## 518         42    1829     143     87         0      0      2   TRUE
## 519         42     425     324     87         0      0      2   TRUE
## 520         42     967     932     87         0      0      2   TRUE
## 521         41     400     352     87         0      0      2   TRUE
## 522         41     387     371     87         0      0      2   TRUE
## 524         41    1714    1409     87         0      0      2   TRUE
## 525         41    2002      62     87         0      0      2   TRUE
## 526         41     353     328     87         0      0      2   TRUE
## 527         41      18     633     87         0      0      1   TRUE
## 528         41     332     332     87         0      0      2   TRUE
## 529         41    1738      89     87         0      0      2   TRUE
## 530         41     428     228     87         0      0      2   TRUE
## 531         41     339     307     87         0      0      2   TRUE
## 532         40     439     367     87         0      0      2   TRUE
## 533         40     494     415     87         0      0      2   TRUE
## 534         40     370     305     87         0      0      2   TRUE
## 535         40     925    1028     87         0      0      2   TRUE
## 536         40    1000     382     87         0      0      2   TRUE
## 537         40     329     333     87         0      0      2   TRUE
## 539         40     342     329     87         0      0      2   TRUE
## 540         40    1224     180     87         0      0      2   TRUE
## 541         40    1011     755     87         0      0      2   TRUE
## 542         40     312     362     87         0      0      1   TRUE
## 543         40    1328    1023     87         0      0      2   TRUE
## 544         40     586     436     87         0      0      2   TRUE
## 545         40     372     345     87         0      0      2   TRUE
## 546         40    1953      92     87         0      0      2   TRUE
## 547         40     345     365     87         0      0      1   TRUE
## 548         40     355     296     88         0      0      2   TRUE
## 549         39     365     370     86         0      0      1   TRUE
## 550         39     423     421     87         0      0      1   TRUE
## 551         39     297     306     85         0      0      1   TRUE
## 552         39     382     304     87         0      0      2   TRUE
## 553         39     376     322     87         0      0      2   TRUE
## 554         39    1013    1767     87         0      0      2   TRUE
## 555         38     325     337     86         0      0      1   TRUE
## 556         38     318     302     88         0      0      2   TRUE
## 557         38     362     356     87         0      0      1   TRUE
## 559         38     359     326     87         0      0      2   TRUE
## 560         38    1069    1457     87         0      0      2   TRUE
## 561         38     379     385     87         0      0      1   TRUE
## 562         38    1665    1412     87         0      0      2   TRUE
## 563         38     386     351     87         0      0      2   TRUE
## 564         38     401     394     87         0      0      1   TRUE
## 566         37    1343     824     87         0      0      2   TRUE
## 567         37    1792     267     87         0      0      2   TRUE
## 569         37     364     364     87         0      0      1   TRUE
## 571         37    2228    1795     87         0      0      2   TRUE
## 572         37    2237     257    140         0      0      2   TRUE
## 574         37     954    1027     87         0      0      2   TRUE
## 575         37     667     338     87         0      0      2   TRUE
## 576         37      12     941     87         0      0      1   TRUE
## 577         37     388     366     87         0      0      2   TRUE
## 578         37    1871     726     87         0      0      2   TRUE
## 579         37     490     801     87         0      0      2   TRUE
## 580         36     443     429     87         0      0      2   TRUE
## 581         36     389     339     86         0      0      2   TRUE
## 582         36    2105     850     87         0      0      2   TRUE
## 583         36     341     330     86         0      0      2   TRUE
## 584         36    1294    1683     87         0      0      2   TRUE
## 585         36     926     703     87         0      0      1   TRUE
## 586         36     390     312     87         0      0      2   TRUE
## 587         36    1974      17     87         0      0      2   TRUE
## 588         36    1061     782     87         0      0      1   TRUE
## 589         36    1052     859     87         0      0      2   TRUE
## 590         36    1734    1650     87         0      0      2   TRUE
## 592         36    1115     117     86         0      0      2   TRUE
## 594         36     462     390     87         0      0      2   TRUE
## 595         36     368    1438     87         0      0      2   TRUE
## 596         36     360     327     87         0      0      2   TRUE
## 597         36     350     348     87         0      0      2   TRUE
## 598         36     514     414     87         0      0      2   TRUE
## 599         36     373     314     87         0      0      2   TRUE
## 600         35    2175     228     87         0      0      2   TRUE
## 601         35      27    1465     87         0      0      1   TRUE
## 602         35    1046    1746     87         0      0      2   TRUE
## 603         35     384     432     87         0      0      1   TRUE
## 604         35     415     319     87         0      0      2   TRUE
## 605         35     394     254     87         0      0      2   TRUE
## 606         35    1090    1239     87         0      0      1   TRUE
## 607         35     923    1093     87         0      0      2   TRUE
## 608         34     406     257    140         0      0      2   TRUE
## 609         34     397    1105     87         0      0      2   TRUE
## 610         34     391     354     87         0      0      2   TRUE
## 613         34     323     323     87         0      0      2   TRUE
## 614         34     395     378     87         0      0      1   TRUE
## 615         34    1131     698     87         0      0      2   TRUE
## 616         34    1628    1483     87         0      0      2   TRUE
## 617         34    1688      11     87         0      0      2   TRUE
## 618         34     625     495     87         0      0      2   TRUE
## 620         34     356     359     87         0      0      2   TRUE
## 621         34       5     800     87         0      0      1   TRUE
## 622         34     418     355     87         0      0      2   TRUE
## 623         34    1398     226     87         0      0      2   TRUE
## 624         34    1569    1299     87         0      0      2   TRUE
## 625         34    1218       2     87         0      0      2   TRUE
## 626         34     424     315     87         0      0      2   TRUE
## 627         33     402     375     87         0      0      2   TRUE
## 628         33     441     954     87         0      0      1   TRUE
## 629         33     460     395     87         0      0      2   TRUE
## 630         33    1642     439     87         0      0      2   TRUE
## 631         33     465     387     86         0      0      2   TRUE
## 632         33     289    1470     87         0      0      1   TRUE
## 633         33     348     349     87         0      0      1   TRUE
## 634         33     337     353     87         0      0      1   TRUE
## 635         33    1760     549     87         0      0      2   TRUE
## 636         33     463     405     87         0      0      2   TRUE
## 637         33    1873      24     87         0      0      2   TRUE
## 639         33      63    1262     87         0      0      1   TRUE
## 640         33    1898     141     87         0      0      2   TRUE
## 641         33     407     374     87         0      0      2   TRUE
## 642         33     451     648     87         0      0      2   TRUE
## 643         33     677     418     87         0      0      2   TRUE
## 644         33     261    1266     87         0      0      2   TRUE
## 646         32     416     380     87         0      0      2   TRUE
## 647         32     589     248     87         0      0      2   TRUE
## 648         32     482     331     87         0      0      2   TRUE
## 649         32    1448    1159     87         0      0      2   TRUE
## 650         32     922    1178     87         0      0      2   TRUE
## 651         32    1333     267     87         0      0      2   TRUE
## 652         32    2202     649     87         0      0      2   TRUE
## 653         32    1380    1743     87         0      0      2   TRUE
## 655         32     413     297     87         0      0      2   TRUE
## 656         32     289     657     87         0      0      1   TRUE
## 658         31     525     427     87         0      0      2   TRUE
## 659         31    1542    1122     87         0      0      2   TRUE
## 662         31    1175    1444     87         0      0      2   TRUE
## 663         31    1228     938     87         0      0      2   TRUE
## 664         31     410     392     87         0      0      2   TRUE
## 665         31    1049     818     87         0      0      2   TRUE
## 666         31     450     416     87         0      0      2   TRUE
## 667         31    1187     138     87         0      0      2   TRUE
## 668         31     261    1074     87         0      0      1   TRUE
## 669         31     436     372     87         0      0      2   TRUE
## 670         31    1794     869     87         0      0      2   TRUE
## 671         30     419     420     87         0      0      1   TRUE
## 672         30     951    1292     87         0      0      2   TRUE
## 674         30     510     398     87         0      0      2   TRUE
## 675         30      16    1687     87         0      0      1   TRUE
## 676         30     445     681     87         0      0      1   TRUE
## 677         30     458     417     87         0      0      2   TRUE
## 678         30    1023    1108     87         0      0      2   TRUE
## 679         30     381     644     87         0      0      1   TRUE
## 680         29     446     404     87         0      0      2   TRUE
## 681         29     392     437     87         0      0      1   TRUE
## 682         29    1106    1344     87         0      0      1   TRUE
## 684         29     449     431     87         0      0      2   TRUE
## 685         29    1612     701     87         0      0      2   TRUE
## 686         29     480     428     87         0      0      2   TRUE
## 687         29    1151    1231     87         0      0      1   TRUE
## 688         29     433     369     86         0      0      2   TRUE
## 689         29     920    1318     87         0      0      1   TRUE
## 690         29     478     254     87         0      0      2   TRUE
## 691         29    1962      62     87         0      0      2   TRUE
## 692         29     403     386     88         0      0      2   TRUE
## 693         29     431    1078     87         0      0      2   TRUE
## 696         28     474     434     87         0      0      2   TRUE
## 697         28     417     880     87         0      0      2   TRUE
## 698         28     579     410    140         0      0      2   TRUE
## 699         28     444     413     87         0      0      2   TRUE
## 700         28     461     407     87         0      0      2   TRUE
## 701         28     464     384     87         0      0      2   TRUE
## 702         28     472     379     87         0      0      2   TRUE
## 704         28     934     650     87         0      0      2   TRUE
## 705         28     640     487     87         0      0      2   TRUE
## 706         28     621     719     87         0      0      2   TRUE
## 707         28    1722      56     87         0      0      2   TRUE
## 708         28     457     401     87         0      0      2   TRUE
## 709         28      33    1423     87         0      0      1   TRUE
## 710         28    1648     100     87         0      0      2   TRUE
## 711         28     456     433     87         0      0      2   TRUE
## 712         27     476     443     87         0      0      2   TRUE
## 713         27     946    1548     87         0      0      1   TRUE
## 714         27     964     722     87         0      0      1   TRUE
## 715         27     531     426     87         0      0      2   TRUE
## 716         27     393     376     87         0      0      2   TRUE
## 717         27     467     368     87         0      0      2   TRUE
## 718         27    1438     363     87         0      0      2   TRUE
## 719         27    1496    1461     87         0      0      2   TRUE
## 720         27     492     419     87         0      0      2   TRUE
## 721         27    1478     397     87         0      0      2   TRUE
## 722         27     484     406     87         0      0      2   TRUE
## 723         27     601     482     87         0      0      2   TRUE
## 724         27    2050       1     87         0      0      2   TRUE
## 725         27     374     377     86         0      0      1   TRUE
## 726         27      16    1640     87         0      0      1   TRUE
## 727         27     524     457     87         0      0      2   TRUE
## 728         27     414     373     86         0      0      2   TRUE
## 729         27     438     425     87         0      0      2   TRUE
## 731         27    1203    1590     87         0      0      2   TRUE
## 732         27    1314    1009     87         0      0      2   TRUE
## 733         27     422     411     87         0      0      1   TRUE
## 734         27       6     913     87         0      0      1   TRUE
## 735         26    2128       2     87         0      0      2   TRUE
## 736         26     989     883     87         0      0      2   TRUE
## 737         26    1550    1673     87         0      0      2   TRUE
## 740         26       7     668     87         0      0      1   TRUE
## 741         26       5    1826     87         0      0      1   TRUE
## 742         26    1158     671     87         0      0      2   TRUE
## 744         26     971     842     87         0      0      2   TRUE
## 745         26     560     471     87         0      0      2   TRUE
## 747         26     442     412     87         0      0      2   TRUE
## 748         26      27    1423     87         0      0      1   TRUE
## 749         26    1684    1365     87         0      0      2   TRUE
## 750         26     965     665     87         0      0      2   TRUE
## 751         26    1417     195     87         0      0      2   TRUE
## 752         26     659     492     87         0      0      2   TRUE
## 753         26     542    1024     87         0      0      2   TRUE
## 755         25     930     695     87         0      0      1   TRUE
## 756         25       7    1760     87         0      0      1   TRUE
## 757         25    1954     182     87         0      0      2   TRUE
## 759         25    1927    1557     87         0      0      1   TRUE
## 761         25     513     838     87         0      0      2   TRUE
## 762         25    2115       5     87         0      0      2   TRUE
## 763         25     475     645     87         0      0      1   TRUE
## 764         25    1143    1011     87         0      0      2   TRUE
## 766         25    1754     831     87         0      0      2   TRUE
## 769         25    1992    1140     87         0      0      2   TRUE
## 770         25     454     400     87         0      0      2   TRUE
## 771         25     583     539     87         0      0      1   TRUE
## 772         25    1065     973     87         0      0      2   TRUE
## 773         25     978    1302     87         0      0      1   TRUE
## 775         25     434     493     87         0      0      1   TRUE
## 778         25      49    1362     87         0      0      1   TRUE
## 780         24     420     396     88         0      0      1   TRUE
## 781         24     535     504     87         0      0      1   TRUE
## 783         24    1693     150     87         0      0      2   TRUE
## 784         24     994     452     87         0      0      2   TRUE
## 785         24     396     403     86         0      0      1   TRUE
## 786         24     546     393     87         0      0      2   TRUE
## 787         24     500     444     87         0      0      2   TRUE
## 788         24    1071    1041     87         0      0      2   TRUE
## 789         24    1481     771     87         0      0      2   TRUE
## 791         24    1035     732     87         0      0      2   TRUE
## 793         24    1160    1565     87         0      0      1   TRUE
## 794         24    2021     156     87         0      0      2   TRUE
## 795         24    1066    1049     87         0      0      1   TRUE
## 796         24    2025     248     87         0      0      2   TRUE
## 797         24     507     469     86         0      0      2   TRUE
## 798         24     581     451     87         0      0      2   TRUE
## 799         24     435     391     87         0      0      2   TRUE
## 800         24    1643    1578     87         0      0      1   TRUE
## 801         24    1125    1033     87         0      0      2   TRUE
## 802         24     590     447     87         0      0      2   TRUE
## 803         24     469     470     88         0      0      1   TRUE
## 805         23     485    1146     87         0      0      2   TRUE
## 806         23      12     797     87         0      0      1   TRUE
## 807         23    1184     991     87         0      0      1   TRUE
## 808         23    1394    1755     87         0      0      2   TRUE
## 809         23    1322      45     87         0      0      2   TRUE
## 810         23    1248    1611     87         0      0      2   TRUE
## 811         23    1387      92     87         0      0      2   TRUE
## 812         23    1043    1561     87         0      0      2   TRUE
## 813         23    1058     454     87         0      0      2   TRUE
## 814         23     519     455     87         0      0      2   TRUE
## 815         23     619     568     87         0      0      1   TRUE
## 816         23    2186     831     87         0      0      2   TRUE
## 817         23     508     402     87         0      0      2   TRUE
## 818         23     506     423     87         0      0      2   TRUE
## 821         23    1862      69     87         0      0      2   TRUE
## 822         23    1700    1758     87         0      0      1   TRUE
## 823         23    1128    1054     87         0      0      2   TRUE
## 824         23    1104     762     87         0      0      2   TRUE
## 825         23     600     656     87         0      0      2   TRUE
## 827         23      47    1666     87         0      0      1   TRUE
## 828         23      40    1486     87         0      0      1   TRUE
## 830         23    1087     683     83         0      0      2   TRUE
## 831         23     522     898     87         0      0      1   TRUE
## 832         23     411     738     87         0      0      1   TRUE
## 833         23      69    1588     87         0      0      1   TRUE
## 834         23     523     467     87         0      0      2   TRUE
## 835         23     973     922     87         0      0      2   TRUE
## 836         22     136       7     87         0      0      2   TRUE
## 837         22    1235    1139     87         0      0      1   TRUE
## 839         22    2164     227     87         0      0      2   TRUE
## 840         22    1098     757     87         0      0      2   TRUE
## 843         22     521     424     88         0      0      2   TRUE
## 844         22    1350    1117     87         0      0      2   TRUE
## 845         22     528     652     87         0      0      1   TRUE
## 846         22    1140     986     87         0      0      2   TRUE
## 847         22     481     445     87         0      0      2   TRUE
## 848         22     538     462     87         0      0      2   TRUE
## 849         22    1029     731     87         0      0      2   TRUE
## 850         22       4    1229     87         0      0      1   TRUE
## 851         22    1359     917     87         0      0      2   TRUE
## 852         22    1056     806     87         0      0      1   TRUE
## 853         22     961    1545     87         0      0      1   TRUE
## 854         22     491    1166     87         0      0      1   TRUE
## 857         22     692     524     87         0      0      2   TRUE
## 858         22    1678    1322     87         0      0      2   TRUE
## 859         22     540     693     87         0      0      2   TRUE
## 860         21    1472    1172     87         0      0      2   TRUE
## 864         21     501     458     87         0      0      2   TRUE
## 865         21     483     453     87         0      0      2   TRUE
## 866         21    1048     440     87         0      0      2   TRUE
## 868         21     516     937     88         0      0      2   TRUE
## 869         21    1302    1370     87         0      0      1   TRUE
## 871         21     499     637     88         0      0      2   TRUE
## 872         21    1138     852     87         0      0      2   TRUE
## 875         21     701     500     87         0      0      2   TRUE
## 876         21    2219      13     87         0      0      2   TRUE
## 877         21     970    1042     87         0      0      2   TRUE
## 881         21     562    1359     87         0      0      1   TRUE
## 882         21      40     633     87         0      0      1   TRUE
## 883         21    1085    1407     87         0      0      2   TRUE
## 884         21     940     714     87         0      0      1   TRUE
## 885         21    1578     118     87         0      0      2   TRUE
## 886         21    1027    1238     87         0      0      1   TRUE
## 887         21     518     449     87         0      0      2   TRUE
## 888         21    1866    1332     87         0      0      2   TRUE
## 890         21     615     473     87         0      0      2   TRUE
## 891         20     452     461     87         0      0      1   TRUE
## 892         20     585     867     87         0      0      1   TRUE
## 895         20    1809     854     86         0      0      2   TRUE
## 896         20     453     430     88         0      0      2   TRUE
## 897         20    1113     456     87         0      0      2   TRUE
## 898         20     512     664     87         0      0      2   TRUE
## 899         20     517     486     87         0      0      2   TRUE
## 900         20    1422     442     87         0      0      2   TRUE
## 902         20    1226    1460     87         0      0      1   TRUE
## 903         20     587     525     86         0      0      2   TRUE
## 904         20     675     843     88         0      0      2   TRUE
## 905         20    1924    1585     87         0      0      2   TRUE
## 906         20    1032    1647     87         0      0      1   TRUE
## 908         20     498    1144     87         0      0      2   TRUE
## 909         20     408     925     87         0      0      1   TRUE
## 910         20     530     468     87         0      0      2   TRUE
## 913         20     489     446     87         0      0      2   TRUE
## 914         20     657     490     87         0      0      2   TRUE
## 915         20     496     459     85         0      0      2   TRUE
## 916         20    1465     849     87         0      0      2   TRUE
## 919         20     504     441     87         0      0      2   TRUE
## 920         20    1045    1374     87         0      0      2   TRUE
## 921         20     999     902     87         0      0      2   TRUE
## 922         20    1042    1476     87         0      0      1   TRUE
## 923         20     602     479     87         0      0      2   TRUE
## 924         20     534     759     87         0      0      1   TRUE
## 925         20     553     660     87         0      0      1   TRUE
## 927         20     920     852     87         0      0      1   TRUE
## 928         20     459     472     87         0      0      1   TRUE
## 929         20     497     680     87         0      0      1   TRUE
## 930         20    1177     992     87         0      0      2   TRUE
## 931         20      33    1465     87         0      0      1   TRUE
## 932         19     691     485     87         0      0      2   TRUE
## 933         19    1877      69     87         0      0      2   TRUE
## 934         19    1031    1378     87         0      0      2   TRUE
## 936         19     257    1739     87         0      0      1   TRUE
## 937         19     554     829     87         0      0      1   TRUE
## 938         19     717     828     87         0      0      2   TRUE
## 941         19     556     498     86         0      0      2   TRUE
## 942         19    1141     706     87         0      0      2   TRUE
## 943         19     526     460     87         0      0      2   TRUE
## 944         19    1020     640     87         0      0      2   TRUE
## 945         19     983     499     87         0      0      2   TRUE
## 946         19     552     474     86         0      0      2   TRUE
## 947         19     470     435     86         0      0      2   TRUE
## 949         19     653     548     87         0      0      2   TRUE
## 952         19     529     464     87         0      0      2   TRUE
## 953         19     471    1038     86         0      0      1   TRUE
## 954         19    1658    1692     87         0      0      2   TRUE
## 955         19     611     477     87         0      0      2   TRUE
## 957         19    1136    1063     87         0      0      2   TRUE
## 958         19       6    1259     87         0      0      1   TRUE
## 959         19     928    1448     87         0      0      1   TRUE
## 960         19    1605    1175     87         0      0      2   TRUE
## 961         19    1012     803     87         0      0      1   TRUE
## 962         19     455     422     87         0      0      2   TRUE
## 964         19    1233     718     87         0      0      1   TRUE
## 966         19    1050    1583     87         0      0      2   TRUE
## 969         18    1362     463     87         0      0      2   TRUE
## 972         18     987     643     87         0      0      2   TRUE
## 973         18    1657    1533     87         0      0      2   TRUE
## 975         18    2159    1755     87         0      0      2   TRUE
## 976         18     541     478     87         0      0      2   TRUE
## 977         18       5    1502     87         0      0      1   TRUE
## 978         18     511     521     87         0      0      1   TRUE
## 980         18     537     691     87         0      0      1   TRUE
## 981         18     334    1803     87         0      0      1   TRUE
## 982         18     559     501     86         0      0      2   TRUE
## 983         18     527     647     87         0      0      1   TRUE
## 984         18    1057     678     87         0      0      2   TRUE
## 985         18     706     494     87         0      0      2   TRUE
## 986         18    1497    1022     87         0      0      2   TRUE
## 988         18    1975    1287     87         0      0      2   TRUE
## 989         18     468     491     87         0      0      1   TRUE
## 992         18    1170     799     87         0      0      2   TRUE
## 993         18     479     669     87         0      0      1   TRUE
## 994         18     641     574     87         0      0      1   TRUE
## 995         18    1152     994     87         0      0      2   TRUE
## 996         18     558     465     87         0      0      2   TRUE
## 998         18     679     710     87         0      0      2   TRUE
## 999         18    1936       6     87         0      0      2   TRUE
## 1000        18     984     979     87         0      0      2   TRUE
## 1001        18     509     489     87         0      0      1   TRUE
## 1003        18     487    1021     87         0      0      1   TRUE
## 1005        18     569     507     87         0      0      2   TRUE
## 1006        18     696     514     86         0      0      2   TRUE
## 1007        17    1038     846     87         0      0      1   TRUE
## 1009        17     557     448     87         0      0      2   TRUE
## 1010        17    1229     804     87         0      0      2   TRUE
## 1012        17    1839    1829     87         0      0      2   TRUE
## 1013        17    2203    1624     87         0      0      1   TRUE
## 1014        17     520     520     86         0      0      1   TRUE
## 1015        17     536     505     87         0      0      1   TRUE
## 1016        17     503     466     87         0      0      2   TRUE
## 1017        17    1323     920     87         0      0      2   TRUE
## 1018        17    1105    1297     87         0      0      2   TRUE
## 1019        17     623     557     87         0      0      2   TRUE
## 1020        17      44    1465     87         0      0      1   TRUE
## 1021        17    1556    1199     87         0      0      2   TRUE
## 1022        17     963     890     87         0      0      2   TRUE
## 1024        17     670     786     87         0      0      2   TRUE
## 1026        17     545     496     87         0      0      1   TRUE
## 1027        17     626     553     88         0      0      1   TRUE
## 1028        17    1783     947     87         0      0      2   TRUE
## 1029        17     549     639     87         0      0      2   TRUE
## 1030        17     613     534     87         0      0      2   TRUE
## 1031        17     614     503     86         0      0      2   TRUE
## 1033        17    2124    1757     87         0      0      1   TRUE
## 1034        17    1762    1354     87         0      0      2   TRUE
## 1035        17     622    1586     87         0      0      1   TRUE
## 1038        17    1348     694     87         0      0      2   TRUE
## 1041        17     488     713     87         0      0      1   TRUE
## 1042        17    2154      17     87         0      0      2   TRUE
## 1043        17    1732      95     87         0      0      2   TRUE
## 1044        17    1245     787     87         0      0      2   TRUE
## 1045        17    1276     654     87         0      0      1   TRUE
## 1047        17     628     653     87         0      0      1   TRUE
## 1048        17    1864    1070     87         0      0      2   TRUE
## 1049        16     740     483     88         0      0      2   TRUE
## 1050        16     709    1276     87         0      0      1   TRUE
## 1051        16     594     763     87         0      0      1   TRUE
## 1052        16    1725    1695     87         0      0      1   TRUE
## 1053        16     937     481     87         0      0      2   TRUE
## 1055        16    1737     438     86         0      0      2   TRUE
## 1056        16    1894    1288     87         0      0      2   TRUE
## 1057        16    1211     642     87         0      0      2   TRUE
## 1058        16     939     484     87         0      0      2   TRUE
## 1059        16    1135     945     86         0      0      2   TRUE
## 1060        16     708     965     87         0      0      2   TRUE
## 1062        16     678     506     87         0      0      2   TRUE
## 1063        16    1003     894     87         0      0      2   TRUE
## 1064        16    1434    1185     87         0      0      2   TRUE
## 1065        16    1553     689     87         0      0      2   TRUE
## 1066        16     673     513     87         0      0      2   TRUE
## 1068        16    1112     909     87         0      0      2   TRUE
## 1069        16    1010    1025     88         0      0      2   TRUE
## 1070        16    1123     912     87         0      0      2   TRUE
## 1071        16     495     450     87         0      0      2   TRUE
## 1072        16    1173     753     87         0      0      2   TRUE
## 1073        16     502     893     87         0      0      1   TRUE
## 1075        16    2114     976     87         0      0      2   TRUE
## 1076        16    1895    1832     87         0      0      1   TRUE
## 1077        16     945     725     87         0      0      2   TRUE
## 1080        16    1254     809     87         0      0      2   TRUE
## 1082        16     645     522     86         0      0      2   TRUE
## 1083        16     544     480     87         0      0      2   TRUE
## 1085        16     555     488     87         0      0      2   TRUE
## 1086        16     680     676     88         0      0      2   TRUE
## 1088        16    1089    1685     87         0      0      1   TRUE
## 1089        16     666     672     87         0      0      1   TRUE
## 1091        16    1488     875     87         0      0      2   TRUE
## 1092        16       5    1316     87         0      0      1   TRUE
## 1093        16    1363    1485     87         0      0      2   TRUE
## 1094        16     997    1065     87         0      0      2   TRUE
## 1096        16    1963    1059     87         0      0      2   TRUE
## 1097        16     607     580     87         0      0      1   TRUE
## 1099        16    1028    1368     87         0      0      1   TRUE
## 1100        16     591     754     87         0      0      2   TRUE
## 1101        16     612     518     86         0      0      2   TRUE
## 1102        16     656     705     87         0      0      1   TRUE
## 1103        15     572     727     87         0      0      2   TRUE
## 1104        15    1025     857     87         0      0      2   TRUE
## 1105        15     563    1303     87         0      0      1   TRUE
## 1107        15     988     936     87         0      0      1   TRUE
## 1109        15     943     933     87         0      0      2   TRUE
## 1110        15       5    1433     87         0      0      1   TRUE
## 1111        15    1129     964     87         0      0      2   TRUE
## 1113        15     573    1158     87         0      0      1   TRUE
## 1114        15     493     700     87         0      0      1   TRUE
## 1116        15    1236    1077     86         0      0      2   TRUE
## 1117        15     660     543     87         0      0      2   TRUE
## 1118        15    2192       1     87         0      0      2   TRUE
## 1121        15     669     558     87         0      0      2   TRUE
## 1122        15    1441    1240     87         0      0      2   TRUE
## 1123        15    1623    1275     87         0      0      2   TRUE
## 1124        15     505     709     87         0      0      2   TRUE
## 1125        15     576     529     87         0      0      1   TRUE
## 1126        15    1037    1799     87         0      0      1   TRUE
## 1127        15    1139    1208     88         0      0      1   TRUE
## 1128        15     608     540     87         0      0      2   TRUE
## 1130        15     539     476     87         0      0      1   TRUE
## 1133        15    1604    1602     87         0      0      2   TRUE
## 1134        15    1300    1790     86         0      0      1   TRUE
## 1136        15    1718     657     87         0      0      2   TRUE
## 1137        15    1399    1393     88         0      0      2   TRUE
## 1138        15     533     515     87         0      0      1   TRUE
## 1139        15     627     884     87         0      0      1   TRUE
## 1141        15     710     919     87         0      0      2   TRUE
## 1142        15     532     512     87         0      0      1   TRUE
## 1143        15    1325    1668     87         0      0      2   TRUE
## 1144        15    2174     858     87         0      0      2   TRUE
## 1145        15     624     651     87         0      0      2   TRUE
## 1146        15    1915    1146     87         0      0      2   TRUE
## 1147        15    1946    1510     87         0      0      2   TRUE
## 1148        15    1972    1581     87         0      0      2   TRUE
## 1150        15    1960    1676     87         0      0      1   TRUE
## 1151        15       4     638     87         0      0      1   TRUE
## 1152        15     944    1168     87         0      0      2   TRUE
## 1153        15     991     666     87         0      0      2   TRUE
## 1154        15    1315    1118     87         0      0      2   TRUE
## 1155        14     547    1012     87         0      0      1   TRUE
## 1157        14      81    1397     87         0      0      1   TRUE
## 1158        14    1230    1367     87         0      0      2   TRUE
## 1161        14    1019     687     87         0      0      2   TRUE
## 1163        14    1468     934     87         0      0      2   TRUE
## 1164        14    1001     526     87         0      0      2   TRUE
## 1168        14    1268    1090     87         0      0      2   TRUE
## 1169        14    1892    1052     87         0      0      2   TRUE
## 1171        14    1469    1223     87         0      0      2   TRUE
## 1173        14     993     550     87         0      0      2   TRUE
## 1174        14    1309     992     87         0      0      2   TRUE
## 1176        14     959     677     87         0      0      2   TRUE
## 1177        14       7    1774     87         0      0      1   TRUE
## 1178        14    1110     970     86         0      0      2   TRUE
## 1180        14     599     905     86         0      0      2   TRUE
## 1181        14    1088     740     87         0      0      2   TRUE
## 1182        14       7    1669     87         0      0      1   TRUE
## 1185        14    2088      74     87         0      0      2   TRUE
## 1186        14    1220    1171     87         0      0      2   TRUE
## 1187        14     588     724     87         0      0      1   TRUE
## 1188        14     661     523     87         0      0      2   TRUE
## 1189        14      15    1760     87         0      0      1   TRUE
## 1190        14     561     781     87         0      0      1   TRUE
## 1192        14    1573     516     87         0      0      2   TRUE
## 1195        14    1256    1473     87         0      0      1   TRUE
## 1196        14    1074     874     87         0      0      2   TRUE
## 1198        14     564     641     87         0      0      1   TRUE
## 1199        14     592     748     87         0      0      1   TRUE
## 1200        14     668     712     87         0      0      2   TRUE
## 1202        14     618     497     86         0      0      2   TRUE
## 1206        14    1274     907     87         0      0      2   TRUE
## 1208        14    1021     827     87         0      0      2   TRUE
## 1209        14     980     908     87         0      0      1   TRUE
## 1210        14    1080    1732     87         0      0      2   TRUE
## 1211        14     577     560     87         0      0      1   TRUE
## 1212        14    1097     837     87         0      0      2   TRUE
## 1213        14       5    1736     87         0      0      1   TRUE
## 1214        14     566     775     86         0      0      2   TRUE
## 1217        14     568     508     87         0      0      2   TRUE
## 1218        14       2    1214     87         0      0      1   TRUE
## 1219        14    1156     930     87         0      0      2   TRUE
## 1220        14    2121    1659     87         0      0      2   TRUE
## 1221        14    1009     662     87         0      0      2   TRUE
## 1222        13     596     761     87         0      0      1   TRUE
## 1224        13     976     765     87         0      0      2   TRUE
## 1225        13     725     562     88         0      0      2   TRUE
## 1226        13    1103     702     87         0      0      2   TRUE
## 1228        13      45     797     87         0      0      1   TRUE
## 1230        13    1843    1663     87         0      0      2   TRUE
## 1231        13    1991      95     87         0      0      2   TRUE
## 1232        13     654     511     87         0      0      2   TRUE
## 1234        13     543     736     87         0      0      1   TRUE
## 1235        13     582     545     87         0      0      1   TRUE
## 1236        13     550     819     87         0      0      1   TRUE
## 1239        13     652     871     87         0      0      1   TRUE
## 1240        13     778     599     87         0      0      1   TRUE
## 1241        13     683     823     86         0      0      2   TRUE
## 1242        13     684     475     86         0      0      2   TRUE
## 1243        13    2117       8     87         0      0      2   TRUE
## 1245        13    1575     704     87         0      0      2   TRUE
## 1246        13     630     730     87         0      0      2   TRUE
## 1247        13    1161    1475     87         0      0      1   TRUE
## 1248        13     574     530     87         0      0      1   TRUE
## 1249        13    1062     798     87         0      0      1   TRUE
## 1250        13    1130    1430     87         0      0      2   TRUE
## 1251        13     580     510     87         0      0      2   TRUE
## 1254        13    1763    1570     87         0      0      2   TRUE
## 1255        13    1329    1216     87         0      0      1   TRUE
## 1256        13    1529     891     87         0      0      2   TRUE
## 1258        13     743     963     87         0      0      1   TRUE
## 1259        13    1196    1006     87         0      0      2   TRUE
## 1261        13     565     899     85         0      0      1   TRUE
## 1265        13    1311    1382     87         0      0      1   TRUE
## 1266        13     632    1071     87         0      0      1   TRUE
## 1268        13    1907     820     87         0      0      2   TRUE
## 1270        13     571     915     86         0      0      1   TRUE
## 1272        13    1107     650     87         0      0      2   TRUE
## 1276        13    1681    1723     87         0      0      2   TRUE
## 1277        13    1379    1290     87         0      0      1   TRUE
## 1278        13      45     941     87         0      0      1   TRUE
## 1279        13      18     913     87         0      0      1   TRUE
## 1280        13    2227     191     87         0      0      2   TRUE
## 1281        13     924    1362     87         0      0      1   TRUE
## 1282        13     734     716     87         0      0      2   TRUE
## 1283        13    1695     863     87         0      0      2   TRUE
## 1284        13    1807    1109     87         0      0      2   TRUE
## 1285        13    1181    1315     87         0      0      2   TRUE
## 1286        13     578     535     85         0      0      1   TRUE
## 1289        12    1150     923     87         0      0      2   TRUE
## 1291        12    1188    1031     87         0      0      1   TRUE
## 1292        12    1897    1755     87         0      0      2   TRUE
## 1293        12     996     769     87         0      0      2   TRUE
## 1294        12    1108     729     87         0      0      2   TRUE
## 1295        12     570    1702     87         0      0      2   TRUE
## 1297        12    1400     873     87         0      0      1   TRUE
## 1298        12    1077     737     87         0      0      2   TRUE
## 1299        12     598     697     87         0      0      1   TRUE
## 1300        12      30    1558     87         0      0      1   TRUE
## 1301        12     597     807     87         0      0      1   TRUE
## 1302        12    1075     752     87         0      0      2   TRUE
## 1304        12    1006     751     87         0      0      2   TRUE
## 1306        12     616     519     85         0      0      2   TRUE
## 1308        12     787     715     87         0      0      1   TRUE
## 1310        12    1366     822     87         0      0      2   TRUE
## 1311        12     593    1057     87         0      0      1   TRUE
## 1313        12     633    1176     86         0      0      2   TRUE
## 1314        12     744     551     85         0      0      2   TRUE
## 1315        12     643     739     87         0      0      1   TRUE
## 1316        12    1460    1429     87         0      0      1   TRUE
## 1317        12    1890     952     87         0      0      2   TRUE
## 1320        12     584     542     87         0      0      1   TRUE
## 1321        12    1257    1443     87         0      0      1   TRUE
## 1323        12    1166    1406     87         0      0      2   TRUE
## 1326        12     575     663     87         0      0      1   TRUE
## 1328        12     737     541     86         0      0      2   TRUE
## 1330        12    1375     896     87         0      0      2   TRUE
## 1331        12     747     773     87         0      0      1   TRUE
## 1332        12    1927     150     87         0      0      2   TRUE
## 1335        12    1591     960     87         0      0      2   TRUE
## 1336        12     762     563     89         0      0      2   TRUE
## 1337        12    2155    1330     87         0      0      2   TRUE
## 1338        12    1157     910     87         0      0      2   TRUE
## 1339        12     646    1342     87         0      0      1   TRUE
## 1340        12     663     536     86         0      0      2   TRUE
## 1341        12      18    1259     87         0      0      1   TRUE
## 1343        12    1590    1100     87         0      0      2   TRUE
## 1344        12    1127     974     87         0      0      1   TRUE
## 1345        12     962    1143     87         0      0      1   TRUE
## 1346        12     603    1789     87         0      0      1   TRUE
## 1347        12    1209    1095     87         0      0      2   TRUE
## 1350        12     605     861     87         0      0      1   TRUE
## 1352        12    1391     795     87         0      0      2   TRUE
## 1353        12    1164    1326     87         0      0      1   TRUE
## 1354        12    2196    1517     87         0      0      1   TRUE
## 1355        12    2033    1814     87         0      0      2   TRUE
## 1356        12    1593    1610     87         0      0      1   TRUE
## 1357        12     794     856     87         0      0      2   TRUE
## 1364        12    1789    1200     87         0      0      2   TRUE
## 1365        12     649     756     87         0      0      2   TRUE
## 1368        12    1159    1030     87         0      0      2   TRUE
## 1369        12     715    1226     87         0      0      2   TRUE
## 1370        12    1720     532     87         0      0      2   TRUE
## 1371        12     639     566     87         0      0      1   TRUE
## 1372        12    1445    1066     87         0      0      2   TRUE
## 1374        12    1169     661     86         0      0      2   TRUE
## 1376        12     700     609     86         0      0      1   TRUE
## 1377        11     941    1114     87         0      0      1   TRUE
## 1378        11    1073    1801     87         0      0      1   TRUE
## 1380        11    1079    1174     87         0      0      1   TRUE
## 1381        11    1024    1405     87         0      0      1   TRUE
## 1382        11     610     509     86         0      0      2   TRUE
## 1384        11    1082     805     87         0      0      2   TRUE
## 1385        11     992    1020     87         0      0      1   TRUE
## 1387        11    1179     939     87         0      0      1   TRUE
## 1388        11    1713    1127     87         0      0      2   TRUE
## 1389        11    1316    1440     87         0      0      1   TRUE
## 1391        11     250    1074     87         0      0      1   TRUE
## 1394        11    1638    1511     87         0      0      1   TRUE
## 1396        11    1562    1582     87         0      0      1   TRUE
## 1397        11      40     913     87         0      0      1   TRUE
## 1400        11    1332    1467     87         0      0      1   TRUE
## 1401        11    1622    1267     88         0      0      2   TRUE
## 1403        11     664     517     86         0      0      2   TRUE
## 1404        11     729     531     86         0      0      2   TRUE
## 1406        11    1627    1486     87         0      0      2   TRUE
## 1408        11     604     791     87         0      0      1   TRUE
## 1409        11     651     554     87         0      0      2   TRUE
## 1410        11     938    1008     87         0      0      1   TRUE
## 1412        11    2017    1298     87         0      0      1   TRUE
## 1417        11    1008     758     87         0      0      2   TRUE
## 1418        11    1869     726     87         0      0      2   TRUE
## 1419        11     634     741     86         0      0      2   TRUE
## 1421        11    1330    1119     87         0      0      2   TRUE
## 1423        11    1002     792     87         0      0      1   TRUE
## 1425        11     636     533     87         0      0      2   TRUE
## 1427        11    1790    1633     87         0      0      2   TRUE
## 1428        11     638     810     87         0      0      1   TRUE
## 1429        11    1367    1566     87         0      0      1   TRUE
## 1430        11     620     696     87         0      0      2   TRUE
## 1432        11      15     668     87         0      0      1   TRUE
## 1434        11     773     586     87         0      0      2   TRUE
## 1436        11    1117     868     87         0      0      1   TRUE
## 1443        11     648     544     87         0      0      2   TRUE
## 1444        11    2103    1753     87         0      0      2   TRUE
## 1449        10     686     537    140         0      0      2   TRUE
## 1452        10    1770    1701     87         0      0      1   TRUE
## 1453        10     617     552     86         0      0      1   TRUE
## 1454        10    1566    1781     87         0      0      2   TRUE
## 1456        10    1162    1092     86         0      0      1   TRUE
## 1457        10    2110     137     87         0      0      2   TRUE
## 1460        10    1697    1802     87         0      0      2   TRUE
## 1462        10     742     690     87         0      0      1   TRUE
## 1464        10    1180    1421     87         0      0      2   TRUE
## 1466        10    1769    1765     87         0      0      1   TRUE
## 1467        10    1132     768     86         0      0      1   TRUE
## 1470        10    1234     708     87         0      0      2   TRUE
## 1471        10    1153    1419     87         0      0      1   TRUE
## 1472        10    1134     833     87         0      0      2   TRUE
## 1473        10    1632    1088     87         0      0      2   TRUE
## 1476        10     714     953     87         0      0      1   TRUE
## 1479        10     749    1138     88         0      0      1   TRUE
## 1484        10    1304    1634     88         0      0      1   TRUE
## 1487        10     609     570     85         0      0      1   TRUE
## 1488        10    1485    1707     87         0      0      2   TRUE
## 1490        10    1461    1559     87         0      0      2   TRUE
## 1491        10     736     788     87         0      0      2   TRUE
## 1493        10     650     528     86         0      0      2   TRUE
## 1494        10    1595     593     87         0      0      1   TRUE
## 1495        10     631     675     87         0      0      1   TRUE
## 1496        10    1547    1149     87         0      0      2   TRUE
## 1497        10    1944    1770     87         0      0      2   TRUE
## 1498        10     953     862     87         0      0      1   TRUE
## 1501        10    1232    1047     87         0      0      2   TRUE
## 1503        10     950     926     87         0      0      1   TRUE
## 1504        10     697     575     87         0      0      1   TRUE
## 1506        10    1857     684     87         0      0      2   TRUE
## 1508        10    1099    1400     86         0      0      1   TRUE
## 1510        10    1984    1716     87         0      0      1   TRUE
## 1511        10    1711     540     87         0      0      2   TRUE
## 1512        10     644     546     87         0      0      2   TRUE
## 1515        10    1114    1016     87         0      0      1   TRUE
## 1522         9     629    1840     88         0      0      1   TRUE
## 1523         9     674     745     86         0      0      2   TRUE
## 1527         9    1247     876     88         0      0      1   TRUE
## 1528         9    1092     903     87         0      0      2   TRUE
## 1530         9    1451    1096     87         0      0      2   TRUE
## 1531         9    1116     996     87         0      0      1   TRUE
## 1534         9    1833     946     88         0      0      2   TRUE
## 1537         9    1091     438     86         0      0      2   TRUE
## 1545         9    2090     686     87         0      0      1   TRUE
## 1547         9    1716    1698     87         0      0      2   TRUE
## 1550         9    1978      69     87         0      0      2   TRUE
## 1551         9    1033    1338     86         0      0      1   TRUE
## 1553         9    1221    1173     86         0      0      2   TRUE
## 1554         9    1272    1693     87         0      0      1   TRUE
## 1557         9     702     779     87         0      0      1   TRUE
## 1558         9     685     931     87         0      0      1   TRUE
## 1561         9     718     790     87         0      0      1   TRUE
## 1565         9    1144     897     87         0      0      1   TRUE
## 1569         9    1680    1132     87         0      0      2   TRUE
## 1570         9    1799     929     87         0      0      2   TRUE
## 1571         9    1926    1637     87         0      0      2   TRUE
## 1574         9     724     582     87         0      0      2   TRUE
## 1576         9     595     813     86         0      0      1   TRUE
## 1577         9    1600    1306     87         0      0      2   TRUE
## 1579         9     949     812     87         0      0      2   TRUE
## 1581         9    1022     892     87         0      0      2   TRUE
## 1582         9      15    1774     87         0      0      1   TRUE
## 1584         9    1614     969     87         0      0      2   TRUE
## 1585         9     647     916     86         0      0      1   TRUE
## 1586         9     635     770     85         0      0      1   TRUE
## 1587         9    1094    1124     87         0      0      1   TRUE
## 1588         9     757     573     87         0      0      2   TRUE
## 1589         9    1004     789     87         0      0      2   TRUE
## 1592         9    1937    1327     87         0      0      2   TRUE
## 1593         9     655     811     87         0      0      1   TRUE
## 1595         9     798     610     87         0      0      2   TRUE
## 1598         9    1249    1142     87         0      0      2   TRUE
## 1599         9    1576    1373     87         0      0      2   TRUE
## 1600         9    1239    1102     87         0      0      2   TRUE
## 1601         9     986    1527     86         0      0      1   TRUE
## 1603         9    1101    1390     87         0      0      1   TRUE
## 1604         9    1016    1277     87         0      0      1   TRUE
## 1606         9     956     555     87         0      0      2   TRUE
## 1607         9    1489    1818     87         0      0      1   TRUE
## 1612         9     792     591     87         0      0      2   TRUE
## 1613         9    1493    1788     87         0      0      1   TRUE
## 1614         9     658     749     85         0      0      2   TRUE
## 1615         9     791     988     87         0      0      1   TRUE
## 1618         9    1067    1323     87         0      0      1   TRUE
## 1619         8     690     772     87         0      0      1   TRUE
## 1620         8    1412    1313     86         0      0      2   TRUE
## 1621         8     720     900     87         0      0      1   TRUE
## 1622         8    2187       1     87         0      0      2   TRUE
## 1626         8     676    1234     86         0      0      1   TRUE
## 1628         8    1526    1585     87         0      0      2   TRUE
## 1629         8    1495    1157     87         0      0      2   TRUE
## 1632         8     606     559     86         0      0      1   TRUE
## 1633         8    1941    1714     87         0      0      1   TRUE
## 1634         8    1111    1086     87         0      0      2   TRUE
## 1635         8     793    1308     87         0      0      1   TRUE
## 1636         8     975    1104     87         0      0      1   TRUE
## 1638         8      14     851     87         0      0      1   TRUE
## 1639         8    1118     688     87         0      0      2   TRUE
## 1641         8    1026     889     87         0      0      1   TRUE
## 1642         8    1476    1060     87         0      0      2   TRUE
## 1643         8    1598    1097     87         0      0      2   TRUE
## 1644         8    1630    1488     87         0      0      2   TRUE
## 1647         8    1376    1675     87         0      0      1   TRUE
## 1650         8     783    1480     87         0      0      1   TRUE
## 1651         8     864    1147     87         0      0      2   TRUE
## 1654         8    1083     967     87         0      0      1   TRUE
## 1656         8    2095    1293     87         0      0      2   TRUE
## 1658         8    1494     808     87         0      0      2   TRUE
## 1660         8    1637    1587     87         0      0      1   TRUE
## 1661         8     777     860     86         0      0      2   TRUE
## 1664         8     682     887     87         0      0      1   TRUE
## 1666         8    1545     733     86         0      0      2   TRUE
## 1669         8     681    1619     86         0      0      1   TRUE
## 1672         8    1212     848     87         0      0      1   TRUE
## 1674         8     695     565     87         0      0      2   TRUE
## 1675         8    1030     864     87         0      0      1   TRUE
## 1679         8     707    1151     87         0      0      1   TRUE
## 1680         8    1017    1269     87         0      0      1   TRUE
## 1681         8    1749     993     87         0      0      2   TRUE
## 1682         8    1318    1183     87         0      0      1   TRUE
## 1683         8    1149    1571     87         0      0      1   TRUE
## 1685         8    1192    1592     87         0      0      2   TRUE
## 1686         8    1646    1056     87         0      0      1   TRUE
## 1689         8    1208    1494     87         0      0      1   TRUE
## 1690         8     750     734     87         0      0      2   TRUE
## 1692         8    1654    1417     87         0      0      2   TRUE
## 1694         8    1060     826     87         0      0      1   TRUE
## 1697         8    1840     140     87         0      0      2   TRUE
## 1698         8     665     870     87         0      0      1   TRUE
## 1704         8     671    1135     87         0      0      1   TRUE
## 1705         8    1990    1691     87         0      0      1   TRUE
## 1706         8    1252    1101     87         0      0      1   TRUE
## 1707         8     705     985     85         0      0      2   TRUE
## 1709         8     704     585     87         0      0      1   TRUE
## 1711         8    1167     984     87         0      0      1   TRUE
## 1713         8      15    1669     87         0      0      1   TRUE
## 1714         8     719     901     87         0      0      2   TRUE
## 1716         8     141    1730     87         0      0      1   TRUE
## 1718         8      44    1423     87         0      0      1   TRUE
## 1720         8     688    1196     87         0      0      1   TRUE
## 1722         8    1739     987     87         0      0      2   TRUE
## 1724         8    1656    1580     87         0      0      1   TRUE
## 1725         8    1371    1817     87         0      0      1   TRUE
## 1728         7     947    1650     87         0      0      1   TRUE
## 1730         7    1767    1648     87         0      0      2   TRUE
## 1731         7    2116    1555     87         0      0      2   TRUE
## 1732         7    1572    1794     87         0      0      1   TRUE
## 1734         7    1289    1204     87         0      0      1   TRUE
## 1737         7    1246    1257     87         0      0      2   TRUE
## 1739         7    1039     878     87         0      0      1   TRUE
## 1740         7    2080    1775     87         0      0      2   TRUE
## 1743         7     687     886     87         0      0      1   TRUE
## 1752         7     698    1478     87         0      0      1   TRUE
## 1753         7    1653    1522     87         0      0      1   TRUE
## 1755         7     731     564     86         0      0      2   TRUE
## 1757         7    1473    1002     88         0      0      1   TRUE
## 1759         7     730    1534     87         0      0      1   TRUE
## 1760         7    1901    1212     87         0      0      1   TRUE
## 1761         7    1306    1285     87         0      0      2   TRUE
## 1762         7     711     817     87         0      0      1   TRUE
## 1763         7    1729     949     87         0      0      2   TRUE
## 1764         7    1155    1235     87         0      0      2   TRUE
## 1770         7     789     577     83         0      0      2   TRUE
## 1771         7    1741    1198     87         0      0      2   TRUE
## 1773         7    1776    1427     87         0      0      2   TRUE
## 1774         7    1334    1482     87         0      0      1   TRUE
## 1775         7     739     855     87         0      0      1   TRUE
## 1776         7    1679    1356     87         0      0      2   TRUE
## 1778         7    1726     583     87         0      0      2   TRUE
## 1779         7    1172    1218     87         0      0      2   TRUE
## 1782         7    1818    1531     87         0      0      2   TRUE
## 1787         7     769     966     87         0      0      2   TRUE
## 1788         7    1791    1469     87         0      0      1   TRUE
## 1790         7     852    1509     87         0      0      1   TRUE
## 1791         7     721     578     86         0      0      2   TRUE
## 1798         7    1582    1350     87         0      0      2   TRUE
## 1800         7    1645    1797     87         0      0      2   TRUE
## 1802         7    1253    1773     88         0      0      1   TRUE
## 1803         7     960    1014     87         0      0      1   TRUE
## 1804         7     751     885     86         0      0      2   TRUE
## 1806         7     637     567     87         0      0      1   TRUE
## 1812         7    1397    1307     87         0      0      1   TRUE
## 1813         7    1259     604     84         0      0      2   TRUE
## 1815         7     728     679     88         0      0      1   TRUE
## 1817         7    1408    1371     86         0      0      2   TRUE
## 1818         7     804    1455     86         0      0      2   TRUE
## 1819         7    1757     958     87         0      0      2   TRUE
## 1820         7    1040     616     87         0      0      2   TRUE
## 1821         7    1852    1664     87         0      0      1   TRUE
## 1823         7    1508    1026     87         0      0      2   TRUE
## 1824         7    2131    1777     87         0      0      2   TRUE
## 1825         7    1518     951     87         0      0      2   TRUE
## 1827         7     723    1162     87         0      0      1   TRUE
## 1829         7     755     581     87         0      0      2   TRUE
## 1831         7     780     561     87         0      0      2   TRUE
## 1832         7    1747      28     87         0      0      2   TRUE
## 1837         7    2167       9     87         0      0      2   TRUE
## 1840         7     713     906     87         0      0      2   TRUE
## 1843         7    1759     972     87         0      0      2   TRUE
## 1844         7    1815      11     87         0      0      2   TRUE
## 1845         7    1200    1563     88         0      0      1   TRUE
## 1848         7    1766     785     86         0      0      2   TRUE
## 1851         7     853    1348     86         0      0      1   TRUE
## 1856         7     776     613     89         0      0      1   TRUE
## 1857         7     957    1148     87         0      0      1   TRUE
## 1858         7    1402    1319     87         0      0      2   TRUE
## 1859         7     854    1375     87         0      0      1   TRUE
## 1861         7     662    1180     87         0      0      1   TRUE
## 1862         7    1287    1831     87         0      0      2   TRUE
## 1864         7    1636    1516     87         0      0      2   TRUE
## 1866         7     732    1317     87         0      0      1   TRUE
## 1867         7    2241    1532     87         0      0      1   TRUE
## 1868         7     689     742     87         0      0      1   TRUE
## 1869         7    1242    1211     87         0      0      2   TRUE
## 1872         7     758     588     87         0      0      2   TRUE
## 1874         7    1520    1179     87         0      0      2   TRUE
## 1878         6    1943    1560     87         0      0      1   TRUE
## 1880         6    1435    1202     87         0      0      2   TRUE
## 1882         6    1262    1484     87         0      0      2   TRUE
## 1883         6     835    1626     87         0      0      1   TRUE
## 1885         6     995    1165     87         0      0      1   TRUE
## 1886         6    1356     956     87         0      0      1   TRUE
## 1888         6    1312    1656     87         0      0      1   TRUE
## 1890         6    1617    1696     87         0      0      1   TRUE
## 1893         6    1168     633     87         0      0      1   TRUE
## 1895         6    1421    1539     87         0      0      2   TRUE
## 1897         6     779    1833     87         0      0      1   TRUE
## 1900         6    1225     942     87         0      0      2   TRUE
## 1902         6    1505     823     86         0      0      2   TRUE
## 1905         6    1606    1064     87         0      0      2   TRUE
## 1911         6    1120    1255     87         0      0      2   TRUE
## 1912         6    1530    1793     87         0      0      2   TRUE
## 1914         6     738     836     87         0      0      1   TRUE
## 1917         6     722    1221     87         0      0      1   TRUE
## 1919         6    1340     598     87         0      0      2   TRUE
## 1920         6     774    1210     87         0      0      1   TRUE
## 1925         6     767     595     89         0      0      2   TRUE
## 1926         6     821     717     88         0      0      2   TRUE
## 1930         6     735     594     87         0      0      1   TRUE
## 1933         6     703     670     88         0      0      1   TRUE
## 1934         6    1174    1681     87         0      0      1   TRUE
## 1936         6    1546    1372     87         0      0      1   TRUE
## 1938         6    2109    1230     87         0      0      2   TRUE
## 1939         6    1297    1512     87         0      0      1   TRUE
## 1942         6     693     579     81         0      0      1   TRUE
## 1944         6    1436    1340     87         0      0      2   TRUE
## 1946         6     865    1242     87         0      0      2   TRUE
## 1947         6    1484    1347     87         0      0      1   TRUE
## 1948         6     886     624     84         0      0      2   TRUE
## 1951         6    1041    1621     88         0      0      1   TRUE
## 1954         6    1183    1441     87         0      0      1   TRUE
## 1956         6    1360    1003     87         0      0      2   TRUE
## 1963         6    1537    1068     87         0      0      1   TRUE
## 1965         6     914    1048     87         0      0      1   TRUE
## 1967         6     815     980     87         0      0      1   TRUE
## 1968         6    1206    1479     86         0      0      1   TRUE
## 1969         6    1784    1362     87         0      0      1   TRUE
## 1975         6     752     957     87         0      0      1   TRUE
## 1976         6    1047    1254     87         0      0      1   TRUE
## 1978         6     712     834     87         0      0      1   TRUE
## 1979         6     772     783     87         0      0      1   TRUE
## 1984         6    1603     776     87         0      0      2   TRUE
## 1985         6     756    1203     87         0      0      1   TRUE
## 1986         6     818     576     86         0      0      2   TRUE
## 1988         6     746    1346     87         0      0      1   TRUE
## 1989         6    1479    1312     87         0      0      2   TRUE
## 1991         6    2151    1201     87         0      0      2   TRUE
## 1996         6    1883    1341     87         0      0      2   TRUE
## 1998         6     805     832     86         0      0      1   TRUE
## 1999         6    1403    1034     87         0      0      2   TRUE
## 2001         6    2215    1249     86         0      0      1   TRUE
## 2003         6     760     607     86         0      0      1   TRUE
## 2009         6    2129      28     87         0      0      2   TRUE
## 2010         6    1699    1567     87         0      0      1   TRUE
## 2011         6     763    1398     87         0      0      1   TRUE
## 2016         6     790     587     87         0      0      2   TRUE
## 2017         6    1821    1540     86         0      0      2   TRUE
## 2018         6    1452    1749     86         0      0      1   TRUE
## 2020         6     129    1801     87         0      0      1   TRUE
## 2022         6     901     589     86         0      0      2   TRUE
## 2024         6     741     571     88         0      0      1   TRUE
## 2026         6    1034    1190     87         0      0      1   TRUE
## 2030         6    1186     602     88         0      0      2   TRUE
## 2034         6    1568    1055     87         0      0      1   TRUE
## 2036         5    1449     872     87         0      0      2   TRUE
## 2037         5    1899    1017     86         0      0      2   TRUE
## 2038         5     733    1544     87         0      0      1   TRUE
## 2039         5     810     600     88         0      0      2   TRUE
## 2045         5    2024      97     87         0      0      2   TRUE
## 2052         5     672    1349     87         0      0      1   TRUE
## 2053         5    1579    1672     87         0      0      1   TRUE
## 2056         5    1532    1782     86         0      0      1   TRUE
## 2058         5    1307     682     86         0      0      2   TRUE
## 2060         5     869     630    140         0      0      1   TRUE
## 2061         5    1812    1153     88         0      0      2   TRUE
## 2063         5    1299    1641     87         0      0      1   TRUE
## 2065         5     867     590     31         0      0      2   TRUE
## 2066         5    1447    1631     87         0      0      1   TRUE
## 2067         5    1633    1364     87         0      0      2   TRUE
## 2069         5    1506    1529     87         0      0      2   TRUE
## 2076         5     753    1324     87         0      0      1   TRUE
## 2078         5    1765    1431     87         0      0      2   TRUE
## 2083         5    1456    1500     87         0      0      2   TRUE
## 2086         5     952     948     87         0      0      1   TRUE
## 2089         5     856    1688     87         0      0      1   TRUE
## 2092         5    2012      46     87         0      0      2   TRUE
## 2093         5    1319     633     87         0      0      2   TRUE
## 2096         5     828     750     87         0      0      1   TRUE
## 2107         5    1321    1766     87         0      0      1   TRUE
## 2108         5      29    1170     87         0      0      1   TRUE
## 2112         5     839    1329     87         0      0      1   TRUE
## 2113         5      97     735     87         0      0      1   TRUE
## 2114         5    1054     839     87         0      0      1   TRUE
## 2117         5    1290    1738     87         0      0      1   TRUE
## 2122         5    1368    1449     87         0      0      1   TRUE
## 2126         5    1644    1577     87         0      0      1   TRUE
## 2131         5    1440     723     87         0      0      2   TRUE
## 2133         5    1338    1036     87         0      0      2   TRUE
## 2135         5     765    1270     87         0      0      2   TRUE
## 2139         5    1310    1550     87         0      0      1   TRUE
## 2140         5    2122    1355     87         0      0      2   TRUE
## 2143         5    1372    1805     87         0      0      1   TRUE
## 2144         5      18    1562     87         0      0      1   TRUE
## 2147         5     782     844     87         0      0      1   TRUE
## 2150         5    1178    1035     87         0      0      1   TRUE
## 2152         5     876     794     87         0      0      1   TRUE
## 2154         5     817    1764     87         0      0      1   TRUE
## 2155         5    1501     668     87         0      0      1   TRUE
## 2156         5    1597    1188     87         0      0      2   TRUE
## 2159         5    1093    1010     87         0      0      2   TRUE
## 2161         5    1240    1335     86         0      0      1   TRUE
## 2164         5     811    1401     87         0      0      1   TRUE
## 2173         5    1629    1245     87         0      0      2   TRUE
## 2174         5      40    1259     87         0      0      1   TRUE
## 2175         5    1764    1301     87         0      0      2   TRUE
## 2177         5    1237    1094     87         0      0      1   TRUE
## 2179         5    2055    1110     87         0      0      2   TRUE
## 2180         5    1171    1209     87         0      0      1   TRUE
## 2186         5    1516    1486     87         0      0      2   TRUE
## 2188         5    1920    1694     87         0      0      2   TRUE
## 2190         5    2161    1632     87         0      0      2   TRUE
## 2191         5     104    1489     87         0      0      1   TRUE
## 2193         5    1198    1526     87         0      0      1   TRUE
## 2199         5    1717    1649     87         0      0      1   TRUE
## 2203         5    1548    1593     87         0      0      1   TRUE
## 2205         5    1810    1538     87         0      0      2   TRUE
## 2206         5     748    1353     88         0      0      1   TRUE
## 2207         5    1574    1128     87         0      0      1   TRUE
## 2208         5    2232    1706     87         0      0      2   TRUE
## 2210         5    1957     816     87         0      0      2   TRUE
## 2216         5     727     592     87         0      0      1   TRUE
## 2217         5    1386     940     87         0      0      2   TRUE
## 2219         5    1124    1261     87         0      0      1   TRUE
## 2220         5    2163    1107     87         0      0      2   TRUE
## 2221         5    1418     955     87         0      0      2   TRUE
## 2224         5     842     627     88         0      0      1   TRUE
## 2229         5      26    1552     87         0      0      1   TRUE
## 2230         5    1561    1018     86         0      0      2   TRUE
## 2234         5      20    1558     87         0      0      1   TRUE
## 2236         5     726    1045     87         0      0      1   TRUE
## 2238         5    1409    1189     87         0      0      2   TRUE
## 2239         5    2029    1635     87         0      0      2   TRUE
## 2241         5    1826    1654     87         0      0      1   TRUE
## 2242         5    2050     913     87         0      0      2   TRUE
## 2244         5    2180    1652     87         0      0      2   TRUE
## 2246         5    1475    1391     87         0      0      1   TRUE
## 2248         5    1889    1082     87         0      0      2   TRUE
## 2250         5    1864     214     87         0      0      2   TRUE
## 2257         5     770     606     87         0      0      1   TRUE
## 2258         5     849     608     86         0      0      2   TRUE
## 2263         4    1142    1627     87         0      0      1   TRUE
## 2265         4     759     596     87         0      0      1   TRUE
## 2271         4    1250    1250     88         0      0      1   TRUE
## 2273         4     768     611     87         0      0      1   TRUE
## 2279         4    1931    1105     87         0      0      2   TRUE
## 2280         4    1147     948     87         0      0      2   TRUE
## 2281         4    1874    1684     87         0      0      1   TRUE
## 2283         4    1324    1325     87         0      0      1   TRUE
## 2285         4    2082     948     87         0      0      1   TRUE
## 2288         4    1195    1434     87         0      0      1   TRUE
## 2292         4    2071    1653     87         0      0      2   TRUE
## 2301         4     906     612     89         0      0      2   TRUE
## 2306         4    1501    1774     87         0      0      1   TRUE
## 2324         4    1213    1154     86         0      0      1   TRUE
## 2327         4    1275    1000     88         0      0      2   TRUE
## 2329         4    1676    1839     86         0      0      1   TRUE
## 2331         4    1450    1733     87         0      0      1   TRUE
## 2341         4     786     614     87         0      0      1   TRUE
## 2343         4     785     840     87         0      0      1   TRUE
## 2345         4     101    1486     87         0      0      2   TRUE
## 2351         4    1068    1377     87         0      0      1   TRUE
## 2359         4     850     628     86         0      0      1   TRUE
## 2365         4    1293    1113     87         0      0      1   TRUE
## 2370         4    1559     814     87         0      0      1   TRUE
## 2371         4    1838    1763     87         0      0      1   TRUE
## 2375         4     841    1735     87         0      0      1   TRUE
## 2377         4    1146     100     87         0      0      2   TRUE
## 2382         4     101     633     87         0      0      2   TRUE
## 2385         4    1380    1578     87         0      0      1   TRUE
## 2388         4     812    1005     84         0      0      1   TRUE
## 2392         4     904    1098     86         0      0      1   TRUE
## 2394         4    1168    1486     87         0      0      2   TRUE
## 2398         4    2125    1584     86         0      0      1   TRUE
## 2402         4    1133    1013     88         0      0      1   TRUE
## 2403         4    2182    1785     87         0      0      2   TRUE
## 2406         4    1395    1831     87         0      0      2   TRUE
## 2409         4    1084    1518     87         0      0      1   TRUE
## 2415         4    1199    1616     87         0      0      1   TRUE
## 2421         4     764     597     85         0      0      1   TRUE
## 2423         4     847     626     87         0      0      1   TRUE
## 2424         4     187    1557     87         0      0      1   TRUE
## 2425         4    1635    1630     87         0      0      1   TRUE
## 2426         4    1341    1530     90         0      0      2   TRUE
## 2431         4    1095     851     87         0      0      1   TRUE
## 2435         4    2156    1490     87         0      0      2   TRUE
## 2439         4    1491    1192     87         0      0      2   TRUE
## 2440         4     868    1004     85         0      0      1   TRUE
## 2441         4    1320     924     87         0      0      2   TRUE
## 2452         4    1078    1515     87         0      0      1   TRUE
## 2454         4    1768    1085     87         0      0      1   TRUE
## 2457         4    1823    1311     87         0      0      1   TRUE
## 2464         4    1554    1721     87         0      0      1   TRUE
## 2465         4    1446    1251     87         0      0      2   TRUE
## 2467         4    2171    1321     88         0      0      2   TRUE
## 2468         4     840     615     87         0      0      2   TRUE
## 2470         4    1978    1526     87         0      0      2   TRUE
## 2472         4     820    1268     87         0      0      1   TRUE
## 2473         4    1647    1690     86         0      0      2   TRUE
## 2476         4    1796    1264     87         0      0      1   TRUE
## 2479         4     766     815     87         0      0      1   TRUE
## 2481         4    1347    1331     87         0      0      1   TRUE
## 2482         4    1382    1645     87         0      0      1   TRUE
## 2484         4    1552    1576     87         0      0      1   TRUE
## 2486         4    1344    1464     87         0      0      2   TRUE
## 2488         4    1918    1598     87         0      0      2   TRUE
## 2489         4    1507    1705     87         0      0      2   TRUE
## 2490         4    1471    1603     86         0      0      2   TRUE
## 2491         4     761     774     87         0      0      1   TRUE
## 2497         4    1923     981     87         0      0      2   TRUE
## 2498         4      46    1252     86         0      0      1   TRUE
## 2501         4    2230    1108     87         0      0      2   TRUE
## 2502         4     893    1053     86         0      0      1   TRUE
## 2503         4    1541    1291     87         0      0      1   TRUE
## 2509         4    1273    1280     85         0      0      1   TRUE
## 2511         4    2212    1358     87         0      0      2   TRUE
## 2515         4     165    1686     87         0      0      1   TRUE
## 2516         4    1389    1300     87         0      0      1   TRUE
## 2522         4    2049     152     87         0      0      2   TRUE
## 2523         4    1346    1394     87         0      0      1   TRUE
## 2525         4     754    1724     87         0      0      1   TRUE
## 2527         4     915     620     85         0      0      2   TRUE
## 2530         4     775     601     84         0      0      1   TRUE
## 2531         4     781     746     87         0      0      1   TRUE
## 2532         4    2036    1089     87         0      0      2   TRUE
## 2534         4    1270    1181     86         0      0      1   TRUE
## 2535         4    1070     845     87         0      0      1   TRUE
## 2538         4     796    1771     87         0      0      1   TRUE
## 2539         4    1712       2     87         0      0      2   TRUE
## 2541         4    1641    1725     87         0      0      1   TRUE
## 2549         4    1470       1     87         0      0      2   TRUE
## 2555         3    1292    1660     86         0      0      1   TRUE
## 2558         3    2094    1601     87         0      0      2   TRUE
## 2560         3    1189    1220     86         0      0      1   TRUE
## 2568         3    2063    1360     87         0      0      1   TRUE
## 2574         3     909     918     87         0      0      1   TRUE
## 2575         3    1197    1129     88         0      0      2   TRUE
## 2591         3    2062    1423     87         0      0      2   TRUE
## 2593         3    1464    1437     87         0      0      2   TRUE
## 2596         3    1499     633     87         0      0      2   TRUE
## 2600         3    1349     928     88         0      0      1   TRUE
## 2601         3    1342    1471     87         0      0      2   TRUE
## 2602         3    1406    1537     87         0      0      1   TRUE
## 2604         3     851     623     95         0      0      1   TRUE
## 2605         3    1947    1425     87         0      0      1   TRUE
## 2630         3     866    1205     87         0      0      1   TRUE
## 2634         3     917     617     86         0      0      2   TRUE
## 2636         3    1295    1636     87         0      0      1   TRUE
## 2637         3    1351     784     87         0      0      2   TRUE
## 2647         3    1282    1424     87         0      0      1   TRUE
## 2660         3    1336    1389     87         0      0      1   TRUE
## 2665         3    1624    1352     87         0      0      1   TRUE
## 2669         3     809    1222     85         0      0      1   TRUE
## 2675         3    1210    1806     87         0      0      1   TRUE
## 2677         3    1319    1486     87         0      0      2   TRUE
## 2696         3     858     881     87         0      0      1   TRUE
## 2704         3    1877    1526     87         0      0      1   TRUE
## 2706         3     825     777     87         0      0      1   TRUE
## 2709         3     803     618     87         0      0      1   TRUE
## 2712         3      31    1680     87         0      0      1   TRUE
## 2716         3     802    1123     87         0      0      1   TRUE
## 2719         3     830     847     87         0      0      1   TRUE
## 2723         3    1662    1761     87         0      0      2   TRUE
## 2727         3     834    1451     87         0      0      1   TRUE
## 2736         3     913    1039     87         0      0      1   TRUE
## 2740         3     816    1217     87         0      0      1   TRUE
## 2744         3     874     621     86         0      0      2   TRUE
## 2753         3     807    1076     86         0      0      1   TRUE
## 2754         3    1620    1606     87         0      0      1   TRUE
## 2767         3    1983     787     87         0      0      2   TRUE
## 2770         3     808    1029     87         0      0      1   TRUE
## 2775         3    1393    1609     87         0      0      1   TRUE
## 2778         3     878     629     86         0      0      2   TRUE
## 2790         3    2057    1289     87         0      0      1   TRUE
## 2796         3    1055     796     86         0      0      1   TRUE
## 2801         3    1820       2     87         0      0      2   TRUE
## 2802         3    2026    1286     88         0      0      1   TRUE
## 2812         3    1803    1483     87         0      0      2   TRUE
## 2816         3    1223    1328     87         0      0      2   TRUE
## 2822         3    1585    1836     87         0      0      1   TRUE
## 2827         3    1938    1258     87         0      0      2   TRUE
## 2839         3    1989    1547     87         0      0      2   TRUE
## 2842         3    1327     744     86         0      0      2   TRUE
## 2846         3    1740    1126     87         0      0      2   TRUE
## 2861         3    1390    1272     87         0      0      1   TRUE
## 2864         3     814    1044     84         0      0      1   TRUE
## 2869         3    2195     525     86         0      0      2   TRUE
## 2870         3    1744    1697     87         0      0      2   TRUE
## 2873         3     892     622     84         0      0      2   TRUE
## 2875         3    1358    1256     87         0      0      2   TRUE
## 2878         3    1660    1376     87         0      0      1   TRUE
## 2890         3    1581    1402     87         0      0      1   TRUE
## 2899         3    1154    1282     87         0      0      1   TRUE
## 2903         3    1501    1760     87         0      0      1   TRUE
## 2921         3    2073    1710     88         0      0      2   TRUE
## 2922         3    1831    1497     87         0      0      1   TRUE
## 2926         3     426    1523     87         0      0      1   TRUE
## 2928         3    1516     633     87         0      0      2   TRUE
## 2937         3     912    1379     87         0      0      2   TRUE
## 2941         3     907     625     80         0      0      2   TRUE
## 2950         3    1588    1115     87         0      0      2   TRUE
## 2961         3    1704    1772     87         0      0      1   TRUE
## 2963         3     237    1526     87         0      0      1   TRUE
## 2970         3    1018    1526     87         0      0      1   TRUE
## 2981         3     832    1061     87         0      0      1   TRUE
## 2982         3     862     766     87         0      0      1   TRUE
## 2983         3    1721    1617     87         0      0      1   TRUE
## 2992         3    2035    1677     87         0      0      1   TRUE
## 2999         3     831    1594     87         0      0      1   TRUE
## 3000         3     848     989     87         0      0      1   TRUE
## 3005         3    1813    1535     87         0      0      2   TRUE
## 3007         3     861    1358     87         0      0      2   TRUE
## 3010         3    1755    1726     87         0      0      1   TRUE
## 3014         3    1527    1477     86         0      0      1   TRUE
## 3018         3     193    1620     84         0      0      1   TRUE
## 3021         3    2000    1644     87         0      0      1   TRUE
## 3035         3    2130    1295     87         0      0      1   TRUE
## 3037         3    1517    1573     88         0      0      2   TRUE
## 3038         3    1222    1361     88         0      0      1   TRUE
## 3039         3    1463    1496     87         0      0      1   TRUE
## 3042         3     844    1310     87         0      0      1   TRUE
## 3045         3    1467    1073     88         0      0      2   TRUE
## 3048         3    1659    1728     87         0      0      1   TRUE
## 3049         3    1982    1792     87         0      0      2   TRUE
## 3050         3     799    1456     87         0      0      1   TRUE
## 3064         3    1613    1167     87         0      0      1   TRUE
## 3079         3    2077    1589     87         0      0      2   TRUE
## 3085         3     859    1265     87         0      0      1   TRUE
## 3090         3     829    1410     88         0      0      1   TRUE
## 3091         3     101     913     87         0      0      1   TRUE
## 3094         3      30    1834     87         0      0      1   TRUE
## 3097         3    1217    1253     87         0      0      1   TRUE
## 3104         3    1748    1699     87         0      0      1   TRUE
## 3107         3    1609    1392     87         0      0      1   TRUE
## 3108         3    1715    1121     87         0      0      1   TRUE
## 3110         3    1512    1426     86         0      0      1   TRUE
## 3112         3    1081    1248     87         0      0      1   TRUE
## 3114         3     813    1001     87         0      0      1   TRUE
## 3119         3    2054      48     87         0      0      2   TRUE
## 3124         3     885    1468     86         0      0      2   TRUE
## 3163         2     208    1070     87         0      0      1   TRUE
## 3179         2     882    1007     85         0      0      1   TRUE
## 3182         2    2128    1465     87         0      0      2   TRUE
## 3229         2     903    1116     86         0      0      1   TRUE
## 3242         2    1873    1687     87         0      0      1   TRUE
## 3253         2    1728    1337     87         0      0      2   TRUE
## 3266         2    1627     913     87         0      0      1   TRUE
## 3278         2    1677    1709     87         0      0      1   TRUE
## 3280         2    1413    1343     87         0      0      1   TRUE
## 3318         2    1299    1595     87         0      0      1   TRUE
## 3330         2     156    1486     87         0      0      2   TRUE
## 3338         2     117    1019     87         0      0      1   TRUE
## 3361         2    1627     633     87         0      0      2   TRUE
## 3384         2    1905    1628     86         0      0      1   TRUE
## 3391         2    1429    1737     87         0      0      1   TRUE
## 3430         2    1331    1120     88         0      0      1   TRUE
## 3438         2    1424    1414     87         0      0      1   TRUE
## 3449         2    2050    1486     87         0      0      2   TRUE
## 3453         2     899    1080     85         0      0      1   TRUE
## 3466         2    2092     197     87         0      0      2   TRUE
## 3484         2    1411    1442     87         0      0      1   TRUE
## 3485         2     896    1111     86         0      0      1   TRUE
## 3489         2    1482    1669     87         0      0      1   TRUE
## 3491         2    2007      10     87         0      0      2   TRUE
## 3506         2    2003     221     87         0      0      2   TRUE
## 3522         2    1524    1495     87         0      0      1   TRUE
## 3545         2     905    1091    140         0      0      1   TRUE
## 3565         2    1218    1423     87         0      0      2   TRUE
## 3577         2     398     267     87         0      0      2   TRUE
## 3579         2    1251    1783     87         0      0      1   TRUE
## 3595         2    1265    1458     87         0      0      1   TRUE
## 3612         2    1345    1278     87         0      0      1   TRUE
## 3625         2    1361    1284     86         0      0      1   TRUE
## 3651         2    1515    1689     87         0      0      1   TRUE
## 3658         2     895    1237     86         0      0      1   TRUE
## 3702         2    1313    1339     86         0      0      1   TRUE
## 3704         2     889    1811     88         0      0      1   TRUE
## 3728         2    2238      34     87         0      0      2   TRUE
## 3738         2     156     633     87         0      0      2   TRUE
## 3739         2     280    1272     87         0      0      1   TRUE
## 3762         2    1499    1486     87         0      0      2   TRUE
## 3780         2     877    1125     87         0      0      1   TRUE
## 3785         2    1916     101     87         0      0      2   TRUE
## 3811         2    1079    1776     87         0      0      1   TRUE
## 3812         2     251    1661     87         0      0      1   TRUE
## 3823         2    1903    1274     86         0      0      1   TRUE
## 3830         2    1377    1241     87         0      0      1   TRUE
## 3845         2     887    1718     87         0      0      1   TRUE
## 3851         2    1888    1246     87         0      0      1   TRUE
## 3871         2    1998    1415     88         0      0      1   TRUE
## 3924         2     131    1745     87         0      0      1   TRUE
## 3929         2    1578    1730     87         0      0      1   TRUE
## 3932         2     897     971     88         0      0      1   TRUE
## 3955         2    1431    1403     87         0      0      1   TRUE
## 3984         2    1388    1345     86         0      0      1   TRUE
## 3998         2     888    1062     87         0      0      1   TRUE
## 4002         2    1577    1296     87         0      0      1   TRUE
## 4004         2    1378    1435     87         0      0      1   TRUE
## 4008         2    1977    1526     87         0      0      1   TRUE
## 4010         2    1827    1244     87         0      0      1   TRUE
## 4013         2     881    1137     87         0      0      1   TRUE
## 4017         2     873    1236     87         0      0      1   TRUE
## 4018         2     156     913     87         0      0      1   TRUE
## 4019         2    1303    1841     87         0      0      1   TRUE
## 4049         2      82    1170     87         0      0      1   TRUE
## 4067         2     879    1163     87         0      0      1   TRUE
## 4079         2     891     990     82         0      0      1   TRUE
## 4084         2    1415    1504     87         0      0      1   TRUE
## 4086         2    1551    1396     87         0      0      1   TRUE
## 4097         2     908    1219     86         0      0      1   TRUE
## 4121         2    1280    1156     81         0      0      1   TRUE
## 4133         2     880    1067     87         0      0      1   TRUE
## 4137         2     900     978    130         0      0      1   TRUE
## 4151         2    1458    1366     87         0      0      1   TRUE
## 4195         2     898    1722     87         0      0      1   TRUE
## 4204         2     911     950     53         0      0      1   TRUE
## 4207         2    1337    1197     87         0      0      1   TRUE
## 4213         2     918    1072     93         0      0      1   TRUE
## 4226         2    1482    1760     87         0      0      1   TRUE
## 4249         2    1956    1465     87         0      0      2   TRUE
## 4659         1    1482    1774     87         0      0      1   TRUE
## 4668         1    1516    1562     87         0      0      1   TRUE
## 4710         1    1138    1318     87         0      0      2   TRUE
## 4873         1    1516     913     87         0      0      1   TRUE
## 5025         1     101    1259     87         0      0      1   TRUE
## 5197         1    1319     913     87         0      0      1   TRUE
## 5258         1    2062    1465     87         0      0      2   TRUE
## 5308         1    1862    1526     87         0      0      1   TRUE
## 5656         1     121    1272     87         0      0      1   TRUE
## 5691         1    1709    1194     87         0      0      1   TRUE
## 5711         1    2115    1669     87         0      0      2   TRUE
## 5802         1    1168    1259     87         0      0      1   TRUE
## 5854         1    1969    1774     87         0      0      1   TRUE
## 6106         1    1694     739     87         0      0      2   TRUE
## 6115         1    1969    1760     87         0      0      1   TRUE
## 6336         1      66    1448     87         0      0      1   TRUE
## 6598         1       6    1562     87         0      0      1   TRUE
## 6644         1    1693    1557     87         0      0      2   TRUE
## 6667         1    1969     668     87         0      0      1   TRUE
## 7077         1    1709     853     87         0      0      1   TRUE
## 7178         1    2115     668     87         0      0      2   TRUE
## 7201         1    2050     633     87         0      0      2   TRUE
## 7572         1    1168     913     87         0      0      1   TRUE
## 7798         1    1734     720     87         0      0      2   TRUE
## 7893         1    1627    1259     87         0      0      1   TRUE
## 7974         1     246      19     87         0      0      2   TRUE
```

#### Including Plots

```
seqtab <- makeSequenceTable(mergers)
dim(seqtab)
```

```
## [1]    47 24787
```

```
## inspect distribution of sequence lengths
table(nchar(getSequences(seqtab)))
```

```
## 
##   199   201   202   203   204   205   206   207   208   209   210   212   217 
##    78     3     4     1     1     3     3     3     3     1     2     1     2 
##   219   220   221   222   223   224   226   228   229   230   231   233   234 
##     2     4     1     1     2     2     1     3     1     2     7     1     2 
##   235   236   237   238   240   241   243   244   245   246   247   248   249 
##     1     1     2     1     2     2     1     4     4    10    11    20    54 
##   250   251   252   253   254   255   256   257   258   259   260   261   262 
##   136   668 21788  1456   206    86    61    18    12     7     6     2     2 
##   263   264   267   269   270   271   272   273   275   276   277   278   279 
##     2     1     3     1     7     3    10     1     3     6     1     6     1 
##   280   281   282   283   285   286   287   288   291   293   300   302   303 
##     5     1     1     1     3     7     1     1     1     1     2     1     4 
##   304   307   308   310   311   312   313   314 
##     1     1     1     1     5     6     1     2
```

#### Remove non-target lengths

```
##  
seqtab2 <- seqtab[,nchar(colnames(seqtab)) %in% seq(251,253)]
table(nchar(getSequences(seqtab2)))
```

```
## 
##   251   252   253 
##   668 21788  1456
```

#### Remove chimers

```
seqtab.nochim <- removeBimeraDenovo(seqtab2, method="consensus", multithread=TRUE, verbose=TRUE)
```

```
## Identified 10776 bimeras out of 23912 input sequences.
```

```
dim(seqtab.nochim)
```

```
## [1]    47 13136
```

```
table(nchar(getSequences(seqtab.nochim)))
```

```
## 
##   251   252   253 
##   632 11305  1199
```

#### Track pipeline

```
## define function getN
#getN <- function(x) sum(getUniques(x))
## track reads through pipeline
#track <- cbind(outHH, sapply(dadaFs, getN), sapply(dadaRs, getN), sapply(mergers, getN), rowSums(seqtab.nochim))
# If processing a single sample, remove the sapply calls: e.g. replace sapply(dadaFs, getN) with getN(dadaFs)
#colnames(track) <- c("input", "filtered", "denoisedF", "denoisedR", "merged", "nonchim")
#rownames(track) <- sample.names
## see how much data you have left
#write.csv(as.matrix(track), "track.csv")
track <- read.csv("track.csv")
head(track)
```

```
##          X  input filtered denoisedF denoisedR merged nonchim
## 1      A6A 151699   139654    134476    136263 115712  108304
## 2 C34SEP25 142049   132852    129868    130807 115246  108448
## 3  C35OCT2 164534   152672    149318    150114 133830  126840
## 4      C6C 188069   176819    173357    174375 158066  146915
## 5     CA25 191428   178167    174541    175580 154969  138841
## 6     CA30 153976   140764    134816    137099 115047  107337
```

#### Classify with Decipher

```
dna <- DNAStringSet(getSequences(seqtab.nochim)) # Create a DNAStringSet from the ASVs
#load("C:/Users/mglam/Dropbox/BIOINFO/Rscripts/SILVA_SSU_r138_2019.RData") # CHANGE TO THE PATH OF YOUR TRAINING SET
#ids <- IdTaxa(dna, trainingSet, strand="top", processors=NULL, verbose=FALSE) # use all processors
#ranks <- c("domain", "phylum", "class", "order", "family", "genus", "species") # ranks of interest
# Convert the output object of class "Taxa" to a matrix analogous to the output from assignTaxonomy
#taxid <- t(sapply(ids, function(x) {
#        m <- match(ranks, x$rank)
#        taxa <- x$taxon[m]
#       taxa[startsWith(taxa, "unclassified_")] <- NA
 #       taxa
#}))
# saveRDS(taxid, "taxIDHH.rds")
taxid <- readRDS("taxIDHH.rds")
#colnames(taxid) <- ranks; rownames(taxid) <- getSequences(seqtab.nochim)
```

#### Create and save phyloseq object for demo

```
ps <- phyloseq(otu_table(seqtab.nochim, taxa_are_rows=FALSE), 
               tax_table(taxid))
ps
```

```
## phyloseq-class experiment-level object
## otu_table()   OTU Table:         [ 13136 taxa and 47 samples ]
## tax_table()   Taxonomy Table:    [ 13136 taxa by 7 taxonomic ranks ]
```

```
# saveRDS(ps, "psHH.rds")
```

#### Import phyloseq object and start here

```
ps <- readRDS("psHH.rds")
#taxa.print <- taxid # Removing sequence rownames for display only
#rownames(taxa.print) <- NULL
#head(taxa.print)
ps
```

```
## phyloseq-class experiment-level object
## otu_table()   OTU Table:         [ 13136 taxa and 47 samples ]
## tax_table()   Taxonomy Table:    [ 13136 taxa by 7 taxonomic ranks ]
```

#### Remove ASV with uncertain phyla

```
ps1 <- subset_taxa(ps, !is.na(phylum) & !phylum %in% c("", "uncharacterized"))
# prune OTUs that are not present in at least one sample
ps1 <- prune_taxa(taxa_sums(ps1) > 0, ps1)
ps1
```

```
## phyloseq-class experiment-level object
## otu_table()   OTU Table:         [ 7491 taxa and 47 samples ]
## tax_table()   Taxonomy Table:    [ 7491 taxa by 7 taxonomic ranks ]
```

#### Get sequences from phyloseq object and align them

```
seqtabPS <- otu_table(ps1)
## get sequences, this is a dada2 function
seqs <- getSequences(seqtabPS)
names(seqs) <- seqs # This propagates to the tip labels of the tree
alignment <- AlignSeqs(DNAStringSet(seqs), anchor=NA)
```

```
## Determining distance matrix based on shared 8-mers:
## ================================================================================
## 
## Time difference of 265.24 secs
## 
## Clustering into groups by similarity:
## ================================================================================
## 
## Time difference of 8.82 secs
## 
## Aligning Sequences:
## ================================================================================
## 
## Time difference of 50.29 secs
## 
## Iteration 1 of 2:
## 
## Determining distance matrix based on alignment:
## ================================================================================
## 
## Time difference of 27.69 secs
## 
## Reclustering into groups by similarity:
## ================================================================================
## 
## Time difference of 4.8 secs
## 
## Realigning Sequences:
## ================================================================================
## 
## Time difference of 29.21 secs
## 
## Iteration 2 of 2:
## 
## Determining distance matrix based on alignment:
## ================================================================================
## 
## Time difference of 27.81 secs
## 
## Reclustering into groups by similarity:
## ================================================================================
## 
## Time difference of 4.86 secs
## 
## Realigning Sequences:
## ================================================================================
## 
## Time difference of 4.23 secs
```

```
#saveRDS (alignment, "alignmentHH.rds")
head(alignment)
```

```
## DNAStringSet object of length 6:
##     width seq                                               names               
## [1]   281 -ACGGAGGATGCAAGTGTTATCC...GAAAGCTAGGGTAGCAAATGGG- ACGGAGGATGCAAGTGT...
## [2]   281 -ACGGGAGTGGCAAGCGTTATCC...GAAAGCCAGGGGAGCGAAAGGG- ACGGGAGTGGCAAGCGT...
## [3]   281 -ACGGGAGTGGCAAGCGTTATCC...GAAAGCCAGGGGAGCGAAAGGG- ACGGGAGTGGCAAGCGT...
## [4]   281 -ACATAGGGTGCAAGCGTTGTCC...GAAAGCGCGGGGAGCGAACAGG- ACATAGGGTGCAAGCGT...
## [5]   281 -ACGGGAGTGGCAAGCGTTATCC...GAAAGCCAGGGGAGCGAAAGGG- ACGGGAGTGGCAAGCGT...
## [6]   281 -ACGGAGGGTGCAAGCGTTGTCC...GAAAGTGTGGGTATCAAACAGG- ACGGAGGGTGCAAGCGT...
```

#### Make tree (skip in markdown because of processing time)

```
#phang.align <- phyDat(as(alignment, "matrix"), type="DNA")
#dm <- dist.ml(phang.align)
#treeNJ <- NJ(dm) # Note, tip order != sequence order
#fit = pml(treeNJ, data=phang.align)
## negative edges length changed to 0!
#fitGTR <- update(fit, k=4, inv=0.2)
#fitGTR <- optim.pml(fitGTR, model="GTR", optInv=TRUE, optGamma=TRUE,
#                  rearrangement = "stochastic", control = #pml.control(trace = 0))
#saveRDS(fitGTR, "fitGTRhh.rds")
fitGTR <- readRDS("fitGTRhh.rds")
fitGTR
```

```
## 
##  loglikelihood: -289808.7 
## 
## unconstrained loglikelihood: -1581.605 
## Proportion of invariant sites: 0.2158931 
## Discrete gamma model
## Number of rate categories: 4 
## Shape parameter: 0.5493491 
## 
## Rate matrix:
##           a         c         g        t
## a 0.0000000 0.7979219 3.3132184 1.404095
## c 0.7979219 0.0000000 0.5859925 3.999548
## g 3.3132184 0.5859925 0.0000000 1.000000
## t 1.4040950 3.9995478 1.0000000 0.000000
## 
## Base frequencies:  
## 0.2377295 0.1945061 0.2837799 0.2839846
```

#### Add sample data and tree and save phyloseq object

```
hhID <- read.csv2("hhID.csv", header = TRUE, row.names = 1, sep =",", strip.white = T)
sampleMeta = sample_data(hhID)
ps2 = merge_phyloseq(ps, sampleMeta, phy_tree(fitGTR$tree))
ps2
```

```
## phyloseq-class experiment-level object
## otu_table()   OTU Table:         [ 7491 taxa and 47 samples ]
## sample_data() Sample Data:       [ 47 samples by 4 sample variables ]
## tax_table()   Taxonomy Table:    [ 7491 taxa by 7 taxonomic ranks ]
## phy_tree()    Phylogenetic Tree: [ 7491 tips and 7489 internal nodes ]
```

```
##saveRDS(ps2, "psHH2.rds")
```

#### Subset bacteria

```
# ps2 <- readRDS("psHH2.rds")
psB <- subset_taxa(ps2, domain=="Bacteria")
psB <- prune_taxa(taxa_sums(psB) > 0, psB)
psB
```

```
## phyloseq-class experiment-level object
## otu_table()   OTU Table:         [ 7410 taxa and 47 samples ]
## sample_data() Sample Data:       [ 47 samples by 4 sample variables ]
## tax_table()   Taxonomy Table:    [ 7410 taxa by 7 taxonomic ranks ]
## phy_tree()    Phylogenetic Tree: [ 7410 tips and 7408 internal nodes ]
```

#### Subset Archaea

```
psA <- subset_taxa(ps2, domain=="Archaea")
psA <- prune_taxa(taxa_sums(psA) > 0, psA)
psA
```

```
## phyloseq-class experiment-level object
## otu_table()   OTU Table:         [ 81 taxa and 47 samples ]
## sample_data() Sample Data:       [ 47 samples by 4 sample variables ]
## tax_table()   Taxonomy Table:    [ 81 taxa by 7 taxonomic ranks ]
## phy_tree()    Phylogenetic Tree: [ 81 tips and 80 internal nodes ]
```

#### Remove sewage samples

```
ps3 <- subset_samples(psB, type!="sewage")
ps3
```

```
## phyloseq-class experiment-level object
## otu_table()   OTU Table:         [ 7410 taxa and 45 samples ]
## sample_data() Sample Data:       [ 45 samples by 4 sample variables ]
## tax_table()   Taxonomy Table:    [ 7410 taxa by 7 taxonomic ranks ]
## phy_tree()    Phylogenetic Tree: [ 7410 tips and 7408 internal nodes ]
```

#### Plot alpha diversity of bacteria by type

```
pAlpha <- plot_richness(ps3, x= "type", measures=c("Observed", "Shannon"))
#write.csv(as.matrix((pAlpha$data)), "pAlphaB.csv")
pAlpha + theme(panel.background = element_blank(), axis.line = element_line(colour = "black"))
```

#### ANOVA suggests difference in diversity between sample type but not by season.

```
pAlphaData <- pAlpha$data
pShannon <- subset(pAlphaData, variable=="Shannon")
pShannon <- pShannon[order(pShannon[,"sampleID"], decreasing = FALSE),]
fit <- aov(value ~  type, data=pShannon)
summary(fit)
```

```
##             Df Sum Sq Mean Sq F value Pr(>F)  
## type         2  1.130  0.5652   3.899  0.028 *
## Residuals   42  6.088  0.1450                 
## ---
## Signif. codes:  0 '***' 0.001 '**' 0.01 '*' 0.05 '.' 0.1 ' ' 1
```

#### Honest significant difference test from the agricolae package

```
# 
tukey.test2 <- HSD.test(fit, trt = 'type')
tukey.test2
```

```
## $statistics
##     MSerror Df    Mean       CV
##   0.1449615 42 4.83806 7.869643
## 
## $parameters
##    test name.t ntr StudentizedRange alpha
##   Tukey   type   3         3.435823  0.05
## 
## $means
##         value       std  r      Min      Max      Q25      Q50      Q75
## HH   4.951180 0.3348537  6 4.475244 5.295158 4.718026 5.017075 5.217410
## post 4.890346 0.3865404 33 4.091065 5.653430 4.656304 4.900314 5.129454
## pre  4.437367 0.3863976  6 3.773884 4.850057 4.299098 4.524435 4.676664
## 
## $comparison
## NULL
## 
## $groups
##         value groups
## HH   4.951180      a
## post 4.890346      a
## pre  4.437367     ab
## 
## attr(,"class")
## [1] "group"
```

#### Plot Shannon diversity by type

```
# compare means
Hplot <- ggplot(pShannon, aes(x = type, y = value, color = type)) +
  geom_violin(fill = NA) +
  geom_jitter(position = position_jitter(0.2)) +
  xlab(element_blank()) + ylab("Shannon Diversity") + theme_bw() +
 theme(panel.grid.major = element_blank(), panel.grid.minor = element_blank()) +
  theme(legend.position = "none")
Hplot +scale_x_discrete(limits = c("pre", "HH", "post"))
```

#### Save plot for Shannon diversity

```
ggsave("Fig_S8_Shannon.png")
```

```
## Saving 7 x 5 in image
```

#### ANOVA suggests difference in diversity between sample type but not by season.

```
pAlphaData <- pAlpha$data
pRich <- subset(pAlphaData, variable=="Observed")
pRich <- pRich[order(pRich[,"sampleID"], decreasing = FALSE),]
fit <- aov(value ~  type, data=pRich)
summary(fit)
```

```
##             Df Sum Sq Mean Sq F value  Pr(>F)   
## type         2 227003  113501   6.673 0.00304 **
## Residuals   42 714416   17010                   
## ---
## Signif. codes:  0 '***' 0.001 '**' 0.01 '*' 0.05 '.' 0.1 ' ' 1
```

#### Honest significant difference test from the agricolae package

```
# 
tukey.test2 <- HSD.test(fit, trt = 'type')
tukey.test2
```

```
## $statistics
##   MSerror Df     Mean       CV
##   17009.9 42 632.3778 20.62406
## 
## $parameters
##    test name.t ntr StudentizedRange alpha
##   Tukey   type   3         3.435823  0.05
## 
## $means
##         value      std  r Min  Max    Q25 Q50    Q75
## HH   764.8333 167.8492  6 529 1034 706.25 739 824.25
## post 634.1818 132.0978 33 431 1010 566.00 613 678.00
## pre  490.0000  55.0527  6 434  556 441.25 483 538.25
## 
## $comparison
## NULL
## 
## $groups
##         value groups
## HH   764.8333      a
## post 634.1818      a
## pre  490.0000      b
## 
## attr(,"class")
## [1] "group"
```

#### Plot richness by type

```
# compare means
Hplot <- ggplot(pRich, aes(x = type, y = value, color = type)) +
  geom_violin(fill = NA) +
  geom_jitter(position = position_jitter(0.2)) +
  xlab(element_blank()) + ylab("Richness") + theme_bw() +
 theme(panel.grid.major = element_blank(), panel.grid.minor = element_blank()) +
  theme(legend.position = "none")
Hplot +scale_x_discrete(limits = c("pre", "HH", "post"))
```

#### save ASV richness

```
ggsave("Fig_S9_richness.png")
```

```
## Saving 7 x 5 in image
```

#### merge by sampleID

```
mergePS <- merge_samples(ps2, "sampleID")
mergePS
```

```
## phyloseq-class experiment-level object
## otu_table()   OTU Table:         [ 7491 taxa and 44 samples ]
## sample_data() Sample Data:       [ 44 samples by 4 sample variables ]
## tax_table()   Taxonomy Table:    [ 7491 taxa by 7 taxonomic ranks ]
## phy_tree()    Phylogenetic Tree: [ 7491 tips and 7489 internal nodes ]
```

#### Import meta data

```
metaHH <- read.csv2("metaHHfactor.csv", header = TRUE, row.names = 1, sep =",", strip.white = T)
# define function to convert to numeric data
metaHH$month <- as.numeric(metaHH$month)
metaHH$Date <- as.Date.numeric(metaHH$Date, origin = "1900-01-01")
metaHH$NOxuM <- as.numeric(as.character(metaHH$NOxuM))
metaHH$NH4uM <- as.numeric(as.character(metaHH$NH4uM))
metaHH$PuM <- as.numeric(as.character(metaHH$PuM))
metaHH$pH <- as.numeric(as.character(metaHH$pH))
metaHH$Conductivity <- as.numeric(as.character(metaHH$Conductivity))
metaHH$TDS <- as.numeric(as.character(metaHH$TDS))
metaHH$Salinity <- as.numeric(as.character(metaHH$Salinity))
metaHH$Oxygen <- as.numeric(as.character(metaHH$Oxygen))
metaHH$Temp <- as.numeric(as.character(metaHH$Temp))
metaHH$Total <- as.numeric(as.character(metaHH$Total))
metaHH$Ecoli <- as.numeric(as.character(metaHH$Ecoli))
metaHH$Enterococii <- as.numeric(as.character(metaHH$Enterococii))
head(metaHH)
```

```
##    Station SegmentID type month season       Date NOxuM NH4uM  PuM   pH
## aC     C34      2425  pre     8 summer 2017-08-27   1.1   0.5  4.0 7.77
## aH     H08     1113B  pre     8 summer 2017-08-27    NA    NA   NA 7.87
## aJ     J76     2425B  pre     8 summer 2017-08-27   1.0   0.6  2.7 7.47
## aK     K45      2425  pre     8 summer 2017-08-27   1.1   1.4  4.1 7.39
## aR     R75      1101  pre     8 summer 2017-08-27  59.3   5.9 14.6 7.74
## bC     C34      2425   HH     8 summer 2017-09-01   1.6   1.4  2.0 7.58
##    Conductivity   TDS Salinity SterivexML Oxygen Temp Total Ecoli Enterococii
## aC       14.475 13059      8.8         90   5.70 32.6    NA    NA          NA
## aH        1.181  1063      0.6        150   6.99 31.4    NA    NA          NA
## aJ       18.260 16796     11.5        140   7.40 32.7    NA    NA          NA
## aK       19.530 17595     12.1        120   6.00 31.7    NA    NA          NA
## aR        5.395  4854      3.0        165   8.10 32.0    NA    NA          NA
## bC        0.105    90      0.0         95   6.00 24.5 43520  1203          NA
##    factorDate   Day
## aC     25-Aug 42972
## aH     25-Aug 42972
## aJ     25-Aug 42972
## aK     25-Aug 42972
## aR     25-Aug 42972
## bC     30-Aug 42977
```

```
#saveRDS(metaHH, "MetaHHfactor.rds")
```

#### Create refseq and rename

```
ps4 = merge_phyloseq(otu_table(mergePS), tax_table(mergePS), sample_data(metaHH), phy_tree(mergePS))
ps4
```

```
## phyloseq-class experiment-level object
## otu_table()   OTU Table:         [ 7491 taxa and 44 samples ]
## sample_data() Sample Data:       [ 44 samples by 21 sample variables ]
## tax_table()   Taxonomy Table:    [ 7491 taxa by 7 taxonomic ranks ]
## phy_tree()    Phylogenetic Tree: [ 7491 tips and 7489 internal nodes ]
```

```
saveRDS(ps4, "ps4HH.rds")
```

#### Show sample data

```
head(sample_data(ps4))
```

```
##    Station SegmentID type month season       Date NOxuM NH4uM  PuM   pH
## aC     C34      2425  pre     8 summer 2017-08-27   1.1   0.5  4.0 7.77
## aH     H08     1113B  pre     8 summer 2017-08-27    NA    NA   NA 7.87
## aJ     J76     2425B  pre     8 summer 2017-08-27   1.0   0.6  2.7 7.47
## aK     K45      2425  pre     8 summer 2017-08-27   1.1   1.4  4.1 7.39
## aR     R75      1101  pre     8 summer 2017-08-27  59.3   5.9 14.6 7.74
## bC     C34      2425   HH     8 summer 2017-09-01   1.6   1.4  2.0 7.58
##    Conductivity   TDS Salinity SterivexML Oxygen Temp Total Ecoli Enterococii
## aC       14.475 13059      8.8         90   5.70 32.6    NA    NA          NA
## aH        1.181  1063      0.6        150   6.99 31.4    NA    NA          NA
## aJ       18.260 16796     11.5        140   7.40 32.7    NA    NA          NA
## aK       19.530 17595     12.1        120   6.00 31.7    NA    NA          NA
## aR        5.395  4854      3.0        165   8.10 32.0    NA    NA          NA
## bC        0.105    90      0.0         95   6.00 24.5 43520  1203          NA
##    factorDate   Day
## aC     25-Aug 42972
## aH     25-Aug 42972
## aJ     25-Aug 42972
## aK     25-Aug 42972
## aR     25-Aug 42972
## bC     30-Aug 42977
```

#### Remove sewage samples

```
ps4B <- subset_samples(ps4, type!="sewage")
ps4B
```

```
## phyloseq-class experiment-level object
## otu_table()   OTU Table:         [ 7491 taxa and 43 samples ]
## sample_data() Sample Data:       [ 43 samples by 21 sample variables ]
## tax_table()   Taxonomy Table:    [ 7491 taxa by 7 taxonomic ranks ]
## phy_tree()    Phylogenetic Tree: [ 7491 tips and 7489 internal nodes ]
```

#### Plot alpha diversity of microbes by type

```
pAlphaB <- plot_richness(ps4B, x= "type", measures=c("Observed", "Shannon"))
# write.csv(as.matrix((pAlpha$data)), "pAlphaB.csv")
pAlphaB + theme(panel.background = element_blank(), axis.line = element_line(colour = "black"))
```

#### ANOVA suggests difference in diversity between sample type but not by season.

```
pAlphaData <- pAlphaB$data
pShannon <- subset(pAlphaData, variable=="Shannon")
pShannon <- pShannon[order(pShannon[,"SegmentID"], decreasing = FALSE),]
fit <- aov(value ~  SegmentID + type, data=pShannon)
summary(fit)
```

```
##             Df Sum Sq Mean Sq F value Pr(>F)  
## SegmentID    4  0.875  0.2188    1.56 0.2060  
## type         2  1.111  0.5556    3.96 0.0279 *
## Residuals   36  5.051  0.1403                 
## ---
## Signif. codes:  0 '***' 0.001 '**' 0.01 '*' 0.05 '.' 0.1 ' ' 1
```

#### Tukey test shannon

```
tukey.test2 <- HSD.test(fit, trt = 'type')
tukey.test2
```

```
## $statistics
##     MSerror Df     Mean       CV
##   0.1403156 36 4.852362 7.719688
## 
## $parameters
##    test name.t ntr StudentizedRange alpha
##   Tukey   type   3         3.456758  0.05
## 
## $means
##         value       std  r      Min      Max      Q25      Q50      Q75
## HH   4.954070 0.3359039  6 4.476893 5.296815 4.720021 5.019879 5.223136
## post 4.899553 0.3901293 32 4.091466 5.654662 4.642762 4.916744 5.133923
## pre  4.428292 0.4251928  5 3.773884 4.855365 4.263028 4.560822 4.688364
## 
## $comparison
## NULL
## 
## $groups
##         value groups
## HH   4.954070      a
## post 4.899553      a
## pre  4.428292     ab
## 
## attr(,"class")
## [1] "group"
```

#### Set factors

```
pShannon$SegmentID <- as.factor(pShannon$SegmentID)
pShannon$type <- as.factor(pShannon$type)
pShannon$Day <- as.factor(pShannon$Day)
pShannon$month <- as.factor(pShannon$month)
pShannon$factorDate <- as.factor(pShannon$factorDate)
head(pShannon)
```

```
##    Station SegmentID type month season       Date NOxuM NH4uM  PuM   pH
## 48     R75      1101  pre     8 summer 2017-08-27  59.3   5.9 14.6 7.74
## 54     R75      1101   HH     8 summer 2017-09-01   1.3   1.5  1.9 7.42
## 60     R75      1101 post     9 summer 2017-09-10  20.2   0.6  3.5 7.61
## 66     R75      1101 post     9 summer 2017-09-20  37.1   0.0  4.2 7.72
## 72     R75      1101 post     9 summer 2017-09-27  27.9   3.2  7.4 7.58
## 81     R75      1101 post    10   fall 2017-10-19  15.7   0.0 10.3 7.85
##    Conductivity  TDS Salinity SterivexML Oxygen   Temp Total Ecoli Enterococii
## 48        5.395 4854      3.0        165  8.100 32.000    NA    NA          NA
## 54        0.081   73      0.0        105  5.350 26.100 41060  1046          NA
## 60        0.271  253      0.1        192  5.725 27.195  6212    69          NA
## 66        0.605  545      0.3        116  5.630 29.060  4020    19        1120
## 72        0.276  248      0.1         80  3.830 28.415  7490    35        1120
## 81        1.750 1578      0.9        128  6.450 26.140  6310    18          63
##    factorDate   Day samples variable    value se
## 48     25-Aug 42972      aR  Shannon 3.773884 NA
## 54     30-Aug 42977      bR  Shannon 4.978579 NA
## 60      8-Sep 42986      cR  Shannon 5.170954 NA
## 66     18-Sep 42996      dR  Shannon 4.548674 NA
## 72     25-Sep 43003      eR  Shannon 5.290946 NA
## 81     17-Oct 43025      gR  Shannon 4.417358 NA
```

#### Plot Shannon

```
pShannon %>%
  dplyr::mutate(type = fct_reorder(type, -dplyr::desc(Day))) %>%
ggplot(aes(x = SegmentID, y = value, color = type, shape = SegmentID)) + geom_boxplot(fill = NA) + geom_point(size = 2) +
  geom_jitter(position = position_jitter(0.01)) +
  xlab("Water Body Segment") + ylab("Shannon Diversity Index") + theme_bw() +
  theme(panel.grid.major = element_blank(), panel.grid.minor = element_blank()) +
  theme(legend.position = "right")
```

```
## Error: Problem with `mutate()` column `type`.
## i `type = fct_reorder(type, -dplyr::desc(Day))`.
## x could not find function "fct_reorder"
```

#### Plot Shannon

```
# ggsave("Fig_6_Shannon.png")
```

#### Plot abundant taxa

```
top20 <- names(sort(taxa_sums(ps4), decreasing=TRUE))[1:20]
ps3top20 <- prune_taxa(top20, ps4)
ps3top20 <- transform_sample_counts(ps3top20, function(OTU) OTU/sum(OTU))
plot_bar(ps3top20, fill = "phylum")
```

#### Inspect tax table

```
head(tax_table(ps4))
```

```
## Taxonomy Table:     [6 taxa by 7 taxonomic ranks]:
##                                                                                                                                                                                                                                                              domain    
## ACGGAGGATGCAAGTGTTATCCGGAATCACTGGGCGTAAAGCGTCTGTAGGTGGTTTAATAAGTCAACTGTTAAATCTTGAGGCTCAACTTCAAAATCGCAGCTGAAACTATTAGACTAGAGTATAGTAGAGGTAAAGGGAATTTCCAGTGGAGCGGTGAAATGCGTAGATATTGGAAAGAACACCGATGGCGAAAGCACTTTACTGGGCTATTACTAACACTCAGAGACGAAAGCTAGGGTAGCAAATGGG "Bacteria"
## ACGGGAGTGGCAAGCGTTATCCGGAATTATTGGGCGTAAAGCGTCCGCAGGCGGTCTTGTAAGTCTGTTGTTAAAGCGTGGAGCTTAACTCCATTTCAGCAATGGAAACTGTAAGACTAGAGTGTGGTAGGGGCAGAGGGAATTCCCGGTGTAGCGGTGAAATGCGTAGATATCGGGAAGAACACCAGTGGCGAAGGCGCTCTGCTGGGCCATAACTGACGCTCATGGACGAAAGCCAGGGGAGCGAAAGGG "Bacteria"
## ACGGGAGTGGCAAGCGTTATCCGGAATTATTGGGCGTAAAGCGTCCGCAGGCGGTCTTTTAAGTCTGCTGTTAAAGCGTGGAGCTTAACTCCATTTCGGCAGTGGAAACTGGAAGACTAGAGTGTGGTAGGGGCAGAGGGAATTCCCGGTGTAGCGGTGAAATGCGTAGATATCGGGAAGAACACCAGTGGCGAAGGCGCTCTGCTGGGCCATAACTGACGCTCATGGACGAAAGCCAGGGGAGCGAAAGGG "Bacteria"
## ACATAGGGTGCAAGCGTTGTCCGGAATTATTGGGCGTAAAGAGCTCGTAGGTCGTTTGCCGCGTCGATTGTGAAAATCTGAGGCTCAACCTCAGACCTGCAGTCGATACGGGCAAACTAGAGTGTGGTAGGGGAGACTGGAATTCCTGGTGTAGCGGTGGAATGCGCAGATATCAGGAGGAACACCAATGGCGAAGGCAGGTCTCTGGGCCATAACTGACACTGAGGAGCGAAAGCGCGGGGAGCGAACAGG "Bacteria"
## ACGGGAGTGGCAAGCGTTATCCGGAATTATTGGGCGTAAAGCGTCCGCAGGCGGTCTTGTAAGTCTGTCGTTAAAGCGTGGAGCTTAACTCCATTTCAGCGATGGAAACTGCAAGACTAGAGTGTGGTAGGGGCAGAGGGAATTCCCGGTGTAGCGGTGAAATGCGTAGATATCGGGAAGAACACCAGTGGCGAAGGCGCTCTGCTGGGCCATAACTGACGCTCATGGACGAAAGCCAGGGGAGCGAAAGGG "Bacteria"
## ACGGAGGGTGCAAGCGTTGTCCGGATTTATTGGGTTTAAAGGGTGCGCAGGTGGTTTATTAAGTCAGTGGTGAAAGACGGTCGCTCAACGATTGCAGTGCCATTGAAACTGGTAGACTTGAGTAAAGTAGAGGTGGGCGGAATTGATAGTGTAGCGGTGAAATGCATAGATATTATCAAGAACTCCAATTGCGTAGGCAGCTCACTTGGCTTTTACTGACACTCATGCACGAAAGTGTGGGTATCAAACAGG "Bacteria"
##                                                                                                                                                                                                                                                              phylum            
## ACGGAGGATGCAAGTGTTATCCGGAATCACTGGGCGTAAAGCGTCTGTAGGTGGTTTAATAAGTCAACTGTTAAATCTTGAGGCTCAACTTCAAAATCGCAGCTGAAACTATTAGACTAGAGTATAGTAGAGGTAAAGGGAATTTCCAGTGGAGCGGTGAAATGCGTAGATATTGGAAAGAACACCGATGGCGAAAGCACTTTACTGGGCTATTACTAACACTCAGAGACGAAAGCTAGGGTAGCAAATGGG "Cyanobacteria"   
## ACGGGAGTGGCAAGCGTTATCCGGAATTATTGGGCGTAAAGCGTCCGCAGGCGGTCTTGTAAGTCTGTTGTTAAAGCGTGGAGCTTAACTCCATTTCAGCAATGGAAACTGTAAGACTAGAGTGTGGTAGGGGCAGAGGGAATTCCCGGTGTAGCGGTGAAATGCGTAGATATCGGGAAGAACACCAGTGGCGAAGGCGCTCTGCTGGGCCATAACTGACGCTCATGGACGAAAGCCAGGGGAGCGAAAGGG "Cyanobacteria"   
## ACGGGAGTGGCAAGCGTTATCCGGAATTATTGGGCGTAAAGCGTCCGCAGGCGGTCTTTTAAGTCTGCTGTTAAAGCGTGGAGCTTAACTCCATTTCGGCAGTGGAAACTGGAAGACTAGAGTGTGGTAGGGGCAGAGGGAATTCCCGGTGTAGCGGTGAAATGCGTAGATATCGGGAAGAACACCAGTGGCGAAGGCGCTCTGCTGGGCCATAACTGACGCTCATGGACGAAAGCCAGGGGAGCGAAAGGG "Cyanobacteria"   
## ACATAGGGTGCAAGCGTTGTCCGGAATTATTGGGCGTAAAGAGCTCGTAGGTCGTTTGCCGCGTCGATTGTGAAAATCTGAGGCTCAACCTCAGACCTGCAGTCGATACGGGCAAACTAGAGTGTGGTAGGGGAGACTGGAATTCCTGGTGTAGCGGTGGAATGCGCAGATATCAGGAGGAACACCAATGGCGAAGGCAGGTCTCTGGGCCATAACTGACACTGAGGAGCGAAAGCGCGGGGAGCGAACAGG "Actinobacteriota"
## ACGGGAGTGGCAAGCGTTATCCGGAATTATTGGGCGTAAAGCGTCCGCAGGCGGTCTTGTAAGTCTGTCGTTAAAGCGTGGAGCTTAACTCCATTTCAGCGATGGAAACTGCAAGACTAGAGTGTGGTAGGGGCAGAGGGAATTCCCGGTGTAGCGGTGAAATGCGTAGATATCGGGAAGAACACCAGTGGCGAAGGCGCTCTGCTGGGCCATAACTGACGCTCATGGACGAAAGCCAGGGGAGCGAAAGGG "Cyanobacteria"   
## ACGGAGGGTGCAAGCGTTGTCCGGATTTATTGGGTTTAAAGGGTGCGCAGGTGGTTTATTAAGTCAGTGGTGAAAGACGGTCGCTCAACGATTGCAGTGCCATTGAAACTGGTAGACTTGAGTAAAGTAGAGGTGGGCGGAATTGATAGTGTAGCGGTGAAATGCATAGATATTATCAAGAACTCCAATTGCGTAGGCAGCTCACTTGGCTTTTACTGACACTCATGCACGAAAGTGTGGGTATCAAACAGG "Bacteroidota"    
##                                                                                                                                                                                                                                                              class           
## ACGGAGGATGCAAGTGTTATCCGGAATCACTGGGCGTAAAGCGTCTGTAGGTGGTTTAATAAGTCAACTGTTAAATCTTGAGGCTCAACTTCAAAATCGCAGCTGAAACTATTAGACTAGAGTATAGTAGAGGTAAAGGGAATTTCCAGTGGAGCGGTGAAATGCGTAGATATTGGAAAGAACACCGATGGCGAAAGCACTTTACTGGGCTATTACTAACACTCAGAGACGAAAGCTAGGGTAGCAAATGGG "Cyanobacteriia"
## ACGGGAGTGGCAAGCGTTATCCGGAATTATTGGGCGTAAAGCGTCCGCAGGCGGTCTTGTAAGTCTGTTGTTAAAGCGTGGAGCTTAACTCCATTTCAGCAATGGAAACTGTAAGACTAGAGTGTGGTAGGGGCAGAGGGAATTCCCGGTGTAGCGGTGAAATGCGTAGATATCGGGAAGAACACCAGTGGCGAAGGCGCTCTGCTGGGCCATAACTGACGCTCATGGACGAAAGCCAGGGGAGCGAAAGGG "Cyanobacteriia"
## ACGGGAGTGGCAAGCGTTATCCGGAATTATTGGGCGTAAAGCGTCCGCAGGCGGTCTTTTAAGTCTGCTGTTAAAGCGTGGAGCTTAACTCCATTTCGGCAGTGGAAACTGGAAGACTAGAGTGTGGTAGGGGCAGAGGGAATTCCCGGTGTAGCGGTGAAATGCGTAGATATCGGGAAGAACACCAGTGGCGAAGGCGCTCTGCTGGGCCATAACTGACGCTCATGGACGAAAGCCAGGGGAGCGAAAGGG "Cyanobacteriia"
## ACATAGGGTGCAAGCGTTGTCCGGAATTATTGGGCGTAAAGAGCTCGTAGGTCGTTTGCCGCGTCGATTGTGAAAATCTGAGGCTCAACCTCAGACCTGCAGTCGATACGGGCAAACTAGAGTGTGGTAGGGGAGACTGGAATTCCTGGTGTAGCGGTGGAATGCGCAGATATCAGGAGGAACACCAATGGCGAAGGCAGGTCTCTGGGCCATAACTGACACTGAGGAGCGAAAGCGCGGGGAGCGAACAGG "Actinobacteria"
## ACGGGAGTGGCAAGCGTTATCCGGAATTATTGGGCGTAAAGCGTCCGCAGGCGGTCTTGTAAGTCTGTCGTTAAAGCGTGGAGCTTAACTCCATTTCAGCGATGGAAACTGCAAGACTAGAGTGTGGTAGGGGCAGAGGGAATTCCCGGTGTAGCGGTGAAATGCGTAGATATCGGGAAGAACACCAGTGGCGAAGGCGCTCTGCTGGGCCATAACTGACGCTCATGGACGAAAGCCAGGGGAGCGAAAGGG "Cyanobacteriia"
## ACGGAGGGTGCAAGCGTTGTCCGGATTTATTGGGTTTAAAGGGTGCGCAGGTGGTTTATTAAGTCAGTGGTGAAAGACGGTCGCTCAACGATTGCAGTGCCATTGAAACTGGTAGACTTGAGTAAAGTAGAGGTGGGCGGAATTGATAGTGTAGCGGTGAAATGCATAGATATTATCAAGAACTCCAATTGCGTAGGCAGCTCACTTGGCTTTTACTGACACTCATGCACGAAAGTGTGGGTATCAAACAGG "Bacteroidia"   
##                                                                                                                                                                                                                                                              order            
## ACGGAGGATGCAAGTGTTATCCGGAATCACTGGGCGTAAAGCGTCTGTAGGTGGTTTAATAAGTCAACTGTTAAATCTTGAGGCTCAACTTCAAAATCGCAGCTGAAACTATTAGACTAGAGTATAGTAGAGGTAAAGGGAATTTCCAGTGGAGCGGTGAAATGCGTAGATATTGGAAAGAACACCGATGGCGAAAGCACTTTACTGGGCTATTACTAACACTCAGAGACGAAAGCTAGGGTAGCAAATGGG "Chloroplast"    
## ACGGGAGTGGCAAGCGTTATCCGGAATTATTGGGCGTAAAGCGTCCGCAGGCGGTCTTGTAAGTCTGTTGTTAAAGCGTGGAGCTTAACTCCATTTCAGCAATGGAAACTGTAAGACTAGAGTGTGGTAGGGGCAGAGGGAATTCCCGGTGTAGCGGTGAAATGCGTAGATATCGGGAAGAACACCAGTGGCGAAGGCGCTCTGCTGGGCCATAACTGACGCTCATGGACGAAAGCCAGGGGAGCGAAAGGG "Synechococcales"
## ACGGGAGTGGCAAGCGTTATCCGGAATTATTGGGCGTAAAGCGTCCGCAGGCGGTCTTTTAAGTCTGCTGTTAAAGCGTGGAGCTTAACTCCATTTCGGCAGTGGAAACTGGAAGACTAGAGTGTGGTAGGGGCAGAGGGAATTCCCGGTGTAGCGGTGAAATGCGTAGATATCGGGAAGAACACCAGTGGCGAAGGCGCTCTGCTGGGCCATAACTGACGCTCATGGACGAAAGCCAGGGGAGCGAAAGGG "Synechococcales"
## ACATAGGGTGCAAGCGTTGTCCGGAATTATTGGGCGTAAAGAGCTCGTAGGTCGTTTGCCGCGTCGATTGTGAAAATCTGAGGCTCAACCTCAGACCTGCAGTCGATACGGGCAAACTAGAGTGTGGTAGGGGAGACTGGAATTCCTGGTGTAGCGGTGGAATGCGCAGATATCAGGAGGAACACCAATGGCGAAGGCAGGTCTCTGGGCCATAACTGACACTGAGGAGCGAAAGCGCGGGGAGCGAACAGG "Frankiales"     
## ACGGGAGTGGCAAGCGTTATCCGGAATTATTGGGCGTAAAGCGTCCGCAGGCGGTCTTGTAAGTCTGTCGTTAAAGCGTGGAGCTTAACTCCATTTCAGCGATGGAAACTGCAAGACTAGAGTGTGGTAGGGGCAGAGGGAATTCCCGGTGTAGCGGTGAAATGCGTAGATATCGGGAAGAACACCAGTGGCGAAGGCGCTCTGCTGGGCCATAACTGACGCTCATGGACGAAAGCCAGGGGAGCGAAAGGG "Synechococcales"
## ACGGAGGGTGCAAGCGTTGTCCGGATTTATTGGGTTTAAAGGGTGCGCAGGTGGTTTATTAAGTCAGTGGTGAAAGACGGTCGCTCAACGATTGCAGTGCCATTGAAACTGGTAGACTTGAGTAAAGTAGAGGTGGGCGGAATTGATAGTGTAGCGGTGAAATGCATAGATATTATCAAGAACTCCAATTGCGTAGGCAGCTCACTTGGCTTTTACTGACACTCATGCACGAAAGTGTGGGTATCAAACAGG "Cytophagales"   
##                                                                                                                                                                                                                                                              family           
## ACGGAGGATGCAAGTGTTATCCGGAATCACTGGGCGTAAAGCGTCTGTAGGTGGTTTAATAAGTCAACTGTTAAATCTTGAGGCTCAACTTCAAAATCGCAGCTGAAACTATTAGACTAGAGTATAGTAGAGGTAAAGGGAATTTCCAGTGGAGCGGTGAAATGCGTAGATATTGGAAAGAACACCGATGGCGAAAGCACTTTACTGGGCTATTACTAACACTCAGAGACGAAAGCTAGGGTAGCAAATGGG NA               
## ACGGGAGTGGCAAGCGTTATCCGGAATTATTGGGCGTAAAGCGTCCGCAGGCGGTCTTGTAAGTCTGTTGTTAAAGCGTGGAGCTTAACTCCATTTCAGCAATGGAAACTGTAAGACTAGAGTGTGGTAGGGGCAGAGGGAATTCCCGGTGTAGCGGTGAAATGCGTAGATATCGGGAAGAACACCAGTGGCGAAGGCGCTCTGCTGGGCCATAACTGACGCTCATGGACGAAAGCCAGGGGAGCGAAAGGG "Cyanobiaceae"   
## ACGGGAGTGGCAAGCGTTATCCGGAATTATTGGGCGTAAAGCGTCCGCAGGCGGTCTTTTAAGTCTGCTGTTAAAGCGTGGAGCTTAACTCCATTTCGGCAGTGGAAACTGGAAGACTAGAGTGTGGTAGGGGCAGAGGGAATTCCCGGTGTAGCGGTGAAATGCGTAGATATCGGGAAGAACACCAGTGGCGAAGGCGCTCTGCTGGGCCATAACTGACGCTCATGGACGAAAGCCAGGGGAGCGAAAGGG "Cyanobiaceae"   
## ACATAGGGTGCAAGCGTTGTCCGGAATTATTGGGCGTAAAGAGCTCGTAGGTCGTTTGCCGCGTCGATTGTGAAAATCTGAGGCTCAACCTCAGACCTGCAGTCGATACGGGCAAACTAGAGTGTGGTAGGGGAGACTGGAATTCCTGGTGTAGCGGTGGAATGCGCAGATATCAGGAGGAACACCAATGGCGAAGGCAGGTCTCTGGGCCATAACTGACACTGAGGAGCGAAAGCGCGGGGAGCGAACAGG "Sporichthyaceae"
## ACGGGAGTGGCAAGCGTTATCCGGAATTATTGGGCGTAAAGCGTCCGCAGGCGGTCTTGTAAGTCTGTCGTTAAAGCGTGGAGCTTAACTCCATTTCAGCGATGGAAACTGCAAGACTAGAGTGTGGTAGGGGCAGAGGGAATTCCCGGTGTAGCGGTGAAATGCGTAGATATCGGGAAGAACACCAGTGGCGAAGGCGCTCTGCTGGGCCATAACTGACGCTCATGGACGAAAGCCAGGGGAGCGAAAGGG "Cyanobiaceae"   
## ACGGAGGGTGCAAGCGTTGTCCGGATTTATTGGGTTTAAAGGGTGCGCAGGTGGTTTATTAAGTCAGTGGTGAAAGACGGTCGCTCAACGATTGCAGTGCCATTGAAACTGGTAGACTTGAGTAAAGTAGAGGTGGGCGGAATTGATAGTGTAGCGGTGAAATGCATAGATATTATCAAGAACTCCAATTGCGTAGGCAGCTCACTTGGCTTTTACTGACACTCATGCACGAAAGTGTGGGTATCAAACAGG "Spirosomaceae"  
##                                                                                                                                                                                                                                                              genus               
## ACGGAGGATGCAAGTGTTATCCGGAATCACTGGGCGTAAAGCGTCTGTAGGTGGTTTAATAAGTCAACTGTTAAATCTTGAGGCTCAACTTCAAAATCGCAGCTGAAACTATTAGACTAGAGTATAGTAGAGGTAAAGGGAATTTCCAGTGGAGCGGTGAAATGCGTAGATATTGGAAAGAACACCGATGGCGAAAGCACTTTACTGGGCTATTACTAACACTCAGAGACGAAAGCTAGGGTAGCAAATGGG NA                  
## ACGGGAGTGGCAAGCGTTATCCGGAATTATTGGGCGTAAAGCGTCCGCAGGCGGTCTTGTAAGTCTGTTGTTAAAGCGTGGAGCTTAACTCCATTTCAGCAATGGAAACTGTAAGACTAGAGTGTGGTAGGGGCAGAGGGAATTCCCGGTGTAGCGGTGAAATGCGTAGATATCGGGAAGAACACCAGTGGCGAAGGCGCTCTGCTGGGCCATAACTGACGCTCATGGACGAAAGCCAGGGGAGCGAAAGGG "Cyanobium PCC-6307"
## ACGGGAGTGGCAAGCGTTATCCGGAATTATTGGGCGTAAAGCGTCCGCAGGCGGTCTTTTAAGTCTGCTGTTAAAGCGTGGAGCTTAACTCCATTTCGGCAGTGGAAACTGGAAGACTAGAGTGTGGTAGGGGCAGAGGGAATTCCCGGTGTAGCGGTGAAATGCGTAGATATCGGGAAGAACACCAGTGGCGAAGGCGCTCTGCTGGGCCATAACTGACGCTCATGGACGAAAGCCAGGGGAGCGAAAGGG "Cyanobium PCC-6307"
## ACATAGGGTGCAAGCGTTGTCCGGAATTATTGGGCGTAAAGAGCTCGTAGGTCGTTTGCCGCGTCGATTGTGAAAATCTGAGGCTCAACCTCAGACCTGCAGTCGATACGGGCAAACTAGAGTGTGGTAGGGGAGACTGGAATTCCTGGTGTAGCGGTGGAATGCGCAGATATCAGGAGGAACACCAATGGCGAAGGCAGGTCTCTGGGCCATAACTGACACTGAGGAGCGAAAGCGCGGGGAGCGAACAGG "hgcI clade"        
## ACGGGAGTGGCAAGCGTTATCCGGAATTATTGGGCGTAAAGCGTCCGCAGGCGGTCTTGTAAGTCTGTCGTTAAAGCGTGGAGCTTAACTCCATTTCAGCGATGGAAACTGCAAGACTAGAGTGTGGTAGGGGCAGAGGGAATTCCCGGTGTAGCGGTGAAATGCGTAGATATCGGGAAGAACACCAGTGGCGAAGGCGCTCTGCTGGGCCATAACTGACGCTCATGGACGAAAGCCAGGGGAGCGAAAGGG "Cyanobium PCC-6307"
## ACGGAGGGTGCAAGCGTTGTCCGGATTTATTGGGTTTAAAGGGTGCGCAGGTGGTTTATTAAGTCAGTGGTGAAAGACGGTCGCTCAACGATTGCAGTGCCATTGAAACTGGTAGACTTGAGTAAAGTAGAGGTGGGCGGAATTGATAGTGTAGCGGTGAAATGCATAGATATTATCAAGAACTCCAATTGCGTAGGCAGCTCACTTGGCTTTTACTGACACTCATGCACGAAAGTGTGGGTATCAAACAGG "Pseudarcicella"    
##                                                                                                                                                                                                                                                              species
## ACGGAGGATGCAAGTGTTATCCGGAATCACTGGGCGTAAAGCGTCTGTAGGTGGTTTAATAAGTCAACTGTTAAATCTTGAGGCTCAACTTCAAAATCGCAGCTGAAACTATTAGACTAGAGTATAGTAGAGGTAAAGGGAATTTCCAGTGGAGCGGTGAAATGCGTAGATATTGGAAAGAACACCGATGGCGAAAGCACTTTACTGGGCTATTACTAACACTCAGAGACGAAAGCTAGGGTAGCAAATGGG NA     
## ACGGGAGTGGCAAGCGTTATCCGGAATTATTGGGCGTAAAGCGTCCGCAGGCGGTCTTGTAAGTCTGTTGTTAAAGCGTGGAGCTTAACTCCATTTCAGCAATGGAAACTGTAAGACTAGAGTGTGGTAGGGGCAGAGGGAATTCCCGGTGTAGCGGTGAAATGCGTAGATATCGGGAAGAACACCAGTGGCGAAGGCGCTCTGCTGGGCCATAACTGACGCTCATGGACGAAAGCCAGGGGAGCGAAAGGG NA     
## ACGGGAGTGGCAAGCGTTATCCGGAATTATTGGGCGTAAAGCGTCCGCAGGCGGTCTTTTAAGTCTGCTGTTAAAGCGTGGAGCTTAACTCCATTTCGGCAGTGGAAACTGGAAGACTAGAGTGTGGTAGGGGCAGAGGGAATTCCCGGTGTAGCGGTGAAATGCGTAGATATCGGGAAGAACACCAGTGGCGAAGGCGCTCTGCTGGGCCATAACTGACGCTCATGGACGAAAGCCAGGGGAGCGAAAGGG NA     
## ACATAGGGTGCAAGCGTTGTCCGGAATTATTGGGCGTAAAGAGCTCGTAGGTCGTTTGCCGCGTCGATTGTGAAAATCTGAGGCTCAACCTCAGACCTGCAGTCGATACGGGCAAACTAGAGTGTGGTAGGGGAGACTGGAATTCCTGGTGTAGCGGTGGAATGCGCAGATATCAGGAGGAACACCAATGGCGAAGGCAGGTCTCTGGGCCATAACTGACACTGAGGAGCGAAAGCGCGGGGAGCGAACAGG NA     
## ACGGGAGTGGCAAGCGTTATCCGGAATTATTGGGCGTAAAGCGTCCGCAGGCGGTCTTGTAAGTCTGTCGTTAAAGCGTGGAGCTTAACTCCATTTCAGCGATGGAAACTGCAAGACTAGAGTGTGGTAGGGGCAGAGGGAATTCCCGGTGTAGCGGTGAAATGCGTAGATATCGGGAAGAACACCAGTGGCGAAGGCGCTCTGCTGGGCCATAACTGACGCTCATGGACGAAAGCCAGGGGAGCGAAAGGG NA     
## ACGGAGGGTGCAAGCGTTGTCCGGATTTATTGGGTTTAAAGGGTGCGCAGGTGGTTTATTAAGTCAGTGGTGAAAGACGGTCGCTCAACGATTGCAGTGCCATTGAAACTGGTAGACTTGAGTAAAGTAGAGGTGGGCGGAATTGATAGTGTAGCGGTGAAATGCATAGATATTATCAAGAACTCCAATTGCGTAGGCAGCTCACTTGGCTTTTACTGACACTCATGCACGAAAGTGTGGGTATCAAACAGG NA
```

#### Create refseq and rename ASV

```
dna <- Biostrings::DNAStringSet(taxa_names(ps4))
names(dna) <- taxa_names(ps4)
ps5 <- merge_phyloseq(ps4, dna)
taxa_names(ps5) <- paste0("ASV", seq(ntaxa(ps5)))
head(t(otu_table(ps5)))
```

```
## OTU Table:          [6 taxa and 44 samples]
##                      taxa are rows
##         aC   aH    aJ    aK    aR    bC   bH   bJ   bK   bN   bR   cC   cH   cJ
## ASV1    59 1200     0    21   272   115   25    0   26   27    0 2366  750 3384
## ASV2 18705  518 39293 10875  6280    73   43    0   40    0   24  565    0  512
## ASV3  6928 4989 13328  4633 18019    48   56   39   31    0    9  311   19  603
## ASV4  2934 1936  2113  1112  6283     0  106   52  204    0   80 2963 1229 2527
## ASV5  3189 8033  1507   562 20708    63   88    0   49    0    0  680   93  445
## ASV6     0   76     0     0     0 17623 8848 4448 4309 9771 8615   86  184   61
##        cK    cN   cR   dC   dH    dJ   dK   dN    dR    eC   eH    eJ    eK
## ASV1 1787 22392 4367 4645  387  3989 4760  893 11355 29889  338   833   372
## ASV2  370   245  349 1187   63   984 2317 3101   427   384    0  8058  4722
## ASV3  380    88  200 2226  234  4224 3440 3928  2387  1288  158 13164 13910
## ASV4 2580   876 2088 7145 2361  3433 5847 3476  6389  2663  408  8092  8106
## ASV5  269   236  706 2346  877 16272 1206 3225  7988  2598  401  3442  3026
## ASV6   58   265  625    0  220     0    0    0     0    53 1887     0     0
##        eN   eR    fC   fH   fK    gC    gH   gJ   gK    gN    gR    hC   hJ
## ASV1 6873 7802 21047 8470 2633 13638 10750  349  313 44980 25548  3323 2807
## ASV2  860  383  1174  228 5773  4976   124 8258 7101   788   813 10392 3915
## ASV3 1955 1646  3211 1033 9669  6183   345 4322 2000  1734  3156  9610 4100
## ASV4 2913 3988  6230 2321 2627  5854  1856 5826 5593  8461  5547  3559 2304
## ASV5 6884 5772  2446 2522 1548  3008  1038  981  545  2454  3473  2304 1037
## ASV6  139  807     0  124    0     0   123    0    0   119    47    42   17
##        hK  iH    jH   kS
## ASV1  811 662   164    0
## ASV2 3752  95    66  141
## ASV3 4217  23    16    0
## ASV4 2483   0     0    0
## ASV5  861  65     0    0
## ASV6   15 800 24693 3703
```

```
# saveRDS(ps5, "psHHasv.rds")
```

#### View phyloseq object and export refseq with function from Nicholas Hathaway

```
#ps5 <- readRDS("psHHasv.rds")
ps5
```

```
## phyloseq-class experiment-level object
## otu_table()   OTU Table:         [ 7491 taxa and 44 samples ]
## sample_data() Sample Data:       [ 44 samples by 21 sample variables ]
## tax_table()   Taxonomy Table:    [ 7491 taxa by 7 taxonomic ranks ]
## phy_tree()    Phylogenetic Tree: [ 7491 tips and 7489 internal nodes ]
## refseq()      DNAStringSet:      [ 7491 reference sequences ]
```

```
ps4ref <- as.data.frame(refseq(ps5))
ps4ref$name <- row.names(ps4ref)
ps4ref$seq <- ps4ref$x
ps5ref <- ps4ref[,2:3]
writeFasta<-function(data, filename){
  fastaLines = c()
  for (rowNum in 1:nrow(data)){
    fastaLines = c(fastaLines, as.character(paste(">", data[rowNum,"name"], sep = "")))
    fastaLines = c(fastaLines,as.character(data[rowNum,"seq"]))
  }
  fileConn<-file(filename)
  writeLines(fastaLines, fileConn)
  close(fileConn)
}
writeFasta(ps5ref, "HHps5.fna")
write.csv(otu_table(ps5), "ps5OTU.csv")
```

```
## Error in file(file, ifelse(append, "a", "w")): cannot open the connection
```

```
write.csv(tax_table(ps5), "ps5tax.csv")
```

#### Rarefaction

```
rarecurve((otu_table(ps5)), col = "blue", step=50, cex=0.5)
```

```
# p5rare
```
