## Supplemental File S08 for "Hurricane Harvey Impacts on Water Quality and Microbial Communities in Houston, TX Waterbodies": S08_NMDS.html

HHphyloseq


### HHphyloseq

###### Michael G. LaMontagne

#### 12/31/2021

#### This is R Markdown document presents multivariate analysis, starting with a phyloseq object. The phyloseq will be used to identify environmental variables associated with bacterial community structure.

#### install ggordiplots

```
library(devtools)
```

```
## Loading required package: usethis
```

```
#install_github("jfq3/ggordiplots")
```

#### R Load packages

```
library(philentropy)
packageVersion("philentropy")
```

```
## [1] '0.5.0'
```

```
library(dada2)
```

```
## Loading required package: Rcpp
```

```
packageVersion("dada2")
```

```
## [1] '1.20.0'
```

```
library(biomformat)
packageVersion("biomformat")
```

```
## [1] '1.20.0'
```

```
library(gridExtra)
packageVersion("gridExtra")
```

```
## [1] '2.3'
```

```
library(ggplot2)
packageVersion("ggplot2")
```

```
## [1] '3.3.5'
```

```
library(phyloseq)
```

```
## 
## Attaching package: 'phyloseq'
```

```
## The following object is masked from 'package:philentropy':
## 
##     distance
```

```
packageVersion("phyloseq")
```

```
## [1] '1.36.0'
```

```
library(DECIPHER); packageVersion("DECIPHER")
```

```
## Loading required package: Biostrings
```

```
## Loading required package: BiocGenerics
```

```
## Loading required package: parallel
```

```
## 
## Attaching package: 'BiocGenerics'
```

```
## The following objects are masked from 'package:parallel':
## 
##     clusterApply, clusterApplyLB, clusterCall, clusterEvalQ,
##     clusterExport, clusterMap, parApply, parCapply, parLapply,
##     parLapplyLB, parRapply, parSapply, parSapplyLB
```

```
## The following object is masked from 'package:gridExtra':
## 
##     combine
```

```
## The following objects are masked from 'package:biomformat':
## 
##     colnames, rownames
```

```
## The following objects are masked from 'package:stats':
## 
##     IQR, mad, sd, var, xtabs
```

```
## The following objects are masked from 'package:base':
## 
##     anyDuplicated, append, as.data.frame, basename, cbind, colnames,
##     dirname, do.call, duplicated, eval, evalq, Filter, Find, get, grep,
##     grepl, intersect, is.unsorted, lapply, Map, mapply, match, mget,
##     order, paste, pmax, pmax.int, pmin, pmin.int, Position, rank,
##     rbind, Reduce, rownames, sapply, setdiff, sort, table, tapply,
##     union, unique, unsplit, which.max, which.min
```

```
## Loading required package: S4Vectors
```

```
## Loading required package: stats4
```

```
## 
## Attaching package: 'S4Vectors'
```

```
## The following objects are masked from 'package:base':
## 
##     expand.grid, I, unname
```

```
## Loading required package: IRanges
```

```
## 
## Attaching package: 'IRanges'
```

```
## The following object is masked from 'package:phyloseq':
## 
##     distance
```

```
## The following object is masked from 'package:philentropy':
## 
##     distance
```

```
## The following object is masked from 'package:grDevices':
## 
##     windows
```

```
## Loading required package: XVector
```

```
## Loading required package: GenomeInfoDb
```

```
## 
## Attaching package: 'Biostrings'
```

```
## The following object is masked from 'package:base':
## 
##     strsplit
```

```
## Loading required package: RSQLite
```

```
## [1] '2.20.0'
```

```
library(phangorn)
```

```
## Loading required package: ape
```

```
## 
## Attaching package: 'ape'
```

```
## The following object is masked from 'package:Biostrings':
## 
##     complement
```

```
packageVersion("phangorn")
```

```
## [1] '2.7.1'
```

```
library(vegan)
```

```
## Loading required package: permute
```

```
## 
## Attaching package: 'permute'
```

```
## The following object is masked from 'package:devtools':
## 
##     check
```

```
## Loading required package: lattice
```

```
## This is vegan 2.5-7
```

```
## 
## Attaching package: 'vegan'
```

```
## The following objects are masked from 'package:phangorn':
## 
##     diversity, treedist
```

```
packageVersion("vegan")
```

```
## [1] '2.5.7'
```

```
library(biomformat)
packageVersion("biomformat")
```

```
## [1] '1.20.0'
```

```
library(ggordiplots)
```

```
## Loading required package: glue
```

```
## 
## Attaching package: 'glue'
```

```
## The following object is masked from 'package:Biostrings':
## 
##     collapse
```

```
## The following objects are masked from 'package:IRanges':
## 
##     collapse, trim
```

```
packageVersion("ggordiplots")
```

```
## [1] '0.4.0'
```

```
sessionInfo()
```

```
## R version 4.1.1 (2021-08-10)
## Platform: x86_64-w64-mingw32/x64 (64-bit)
## Running under: Windows 10 x64 (build 22000)
## 
## Matrix products: default
## 
## locale:
## [1] LC_COLLATE=English_United States.1252 
## [2] LC_CTYPE=English_United States.1252   
## [3] LC_MONETARY=English_United States.1252
## [4] LC_NUMERIC=C                          
## [5] LC_TIME=English_United States.1252    
## 
## attached base packages:
## [1] stats4    parallel  stats     graphics  grDevices utils     datasets 
## [8] methods   base     
## 
## other attached packages:
##  [1] ggordiplots_0.4.0   glue_1.4.2          vegan_2.5-7        
##  [4] lattice_0.20-44     permute_0.9-5       phangorn_2.7.1     
##  [7] ape_5.5             DECIPHER_2.20.0     RSQLite_2.2.8      
## [10] Biostrings_2.60.2   GenomeInfoDb_1.28.2 XVector_0.32.0     
## [13] IRanges_2.26.0      S4Vectors_0.30.0    BiocGenerics_0.38.0
## [16] phyloseq_1.36.0     ggplot2_3.3.5       gridExtra_2.3      
## [19] biomformat_1.20.0   dada2_1.20.0        Rcpp_1.0.7         
## [22] philentropy_0.5.0   devtools_2.4.2      usethis_2.1.3      
## 
## loaded via a namespace (and not attached):
##   [1] colorspace_2.0-2            hwriter_1.3.2              
##   [3] ellipsis_0.3.2              rprojroot_2.0.2            
##   [5] GenomicRanges_1.44.0        fs_1.5.0                   
##   [7] rstudioapi_0.13             remotes_2.4.1              
##   [9] bit64_4.0.5                 fansi_0.5.0                
##  [11] codetools_0.2-18            splines_4.1.1              
##  [13] cachem_1.0.6                knitr_1.36                 
##  [15] pkgload_1.2.3               ade4_1.7-18                
##  [17] jsonlite_1.7.2              Rsamtools_2.8.0            
##  [19] cluster_2.1.2               png_0.1-7                  
##  [21] compiler_4.1.1              assertthat_0.2.1           
##  [23] Matrix_1.3-4                fastmap_1.1.0              
##  [25] cli_3.0.1                   htmltools_0.5.2            
##  [27] prettyunits_1.1.1           tools_4.1.1                
##  [29] igraph_1.2.7                gtable_0.3.0               
##  [31] GenomeInfoDbData_1.2.6      reshape2_1.4.4             
##  [33] dplyr_1.0.7                 fastmatch_1.1-3            
##  [35] ShortRead_1.50.0            Biobase_2.52.0             
##  [37] jquerylib_0.1.4             vctrs_0.3.8                
##  [39] rhdf5filters_1.4.0          multtest_2.48.0            
##  [41] nlme_3.1-152                iterators_1.0.13           
##  [43] xfun_0.25                   stringr_1.4.0              
##  [45] ps_1.6.0                    testthat_3.1.0             
##  [47] lifecycle_1.0.1             zlibbioc_1.38.0            
##  [49] MASS_7.3-54                 scales_1.1.1               
##  [51] MatrixGenerics_1.4.3        SummarizedExperiment_1.22.0
##  [53] rhdf5_2.36.0                RColorBrewer_1.1-2         
##  [55] yaml_2.2.1                  memoise_2.0.0              
##  [57] sass_0.4.0                  latticeExtra_0.6-29        
##  [59] stringi_1.7.5               desc_1.4.0                 
##  [61] foreach_1.5.1               pkgbuild_1.2.0             
##  [63] BiocParallel_1.26.2         rlang_0.4.11               
##  [65] pkgconfig_2.0.3             matrixStats_0.61.0         
##  [67] bitops_1.0-7                evaluate_0.14              
##  [69] purrr_0.3.4                 Rhdf5lib_1.14.2            
##  [71] GenomicAlignments_1.28.0    bit_4.0.4                  
##  [73] processx_3.5.2              tidyselect_1.1.1           
##  [75] plyr_1.8.6                  magrittr_2.0.1             
##  [77] R6_2.5.1                    generics_0.1.1             
##  [79] DelayedArray_0.18.0         DBI_1.1.1                  
##  [81] pillar_1.6.4                withr_2.4.2                
##  [83] mgcv_1.8-36                 survival_3.2-11            
##  [85] RCurl_1.98-1.5              tibble_3.1.4               
##  [87] crayon_1.4.1                utf8_1.2.2                 
##  [89] rmarkdown_2.11              jpeg_0.1-9                 
##  [91] grid_4.1.1                  data.table_1.14.2          
##  [93] blob_1.2.2                  callr_3.7.0                
##  [95] digest_0.6.27               RcppParallel_5.1.4         
##  [97] munsell_0.5.0               bslib_0.3.1                
##  [99] quadprog_1.5-8              sessioninfo_1.1.1
```

#### Read phyloseq object

```
ps5 <- readRDS("psHHasv.rds")
ps5
```

```
## phyloseq-class experiment-level object
## otu_table()   OTU Table:         [ 7491 taxa and 44 samples ]
## sample_data() Sample Data:       [ 44 samples by 21 sample variables ]
## tax_table()   Taxonomy Table:    [ 7491 taxa by 7 taxonomic ranks ]
## phy_tree()    Phylogenetic Tree: [ 7491 tips and 7489 internal nodes ]
## refseq()      DNAStringSet:      [ 7491 reference sequences ]
```

#### Compute prevalence of each feature, store as data.frame

```
prevdf = apply(X = otu_table(ps5),
               MARGIN = ifelse(taxa_are_rows(ps5), yes = 1, no = 2),
               FUN = function(x){sum(x > 0)})
head(prevdf)
```

```
## ASV1 ASV2 ASV3 ASV4 ASV5 ASV6 
##   40   40   42   39   39   28
```

#### Add taxonomy and total read counts to this data.frame

```
prevdfB = data.frame(Prevalence = prevdf,
                    TotalAbundance = taxa_sums(ps5),
                    tax_table(ps5), refseq(ps5))
head(prevdfB)
```

```
##      Prevalence TotalAbundance   domain           phylum          class
## ASV1         40         244422 Bacteria    Cyanobacteria Cyanobacteriia
## ASV2         40         147974 Bacteria    Cyanobacteria Cyanobacteriia
## ASV3         42         147860 Bacteria    Cyanobacteria Cyanobacteriia
## ASV4         39         134565 Bacteria Actinobacteriota Actinobacteria
## ASV5         39         112947 Bacteria    Cyanobacteria Cyanobacteriia
## ASV6         28          87758 Bacteria     Bacteroidota    Bacteroidia
##                order          family              genus species
## ASV1     Chloroplast            <NA>               <NA>    <NA>
## ASV2 Synechococcales    Cyanobiaceae Cyanobium PCC-6307    <NA>
## ASV3 Synechococcales    Cyanobiaceae Cyanobium PCC-6307    <NA>
## ASV4      Frankiales Sporichthyaceae         hgcI clade    <NA>
## ASV5 Synechococcales    Cyanobiaceae Cyanobium PCC-6307    <NA>
## ASV6    Cytophagales   Spirosomaceae     Pseudarcicella    <NA>
##                                                                                                                                                                                                                                                       refseq.ps5.
## ASV1 ACGGAGGATGCAAGTGTTATCCGGAATCACTGGGCGTAAAGCGTCTGTAGGTGGTTTAATAAGTCAACTGTTAAATCTTGAGGCTCAACTTCAAAATCGCAGCTGAAACTATTAGACTAGAGTATAGTAGAGGTAAAGGGAATTTCCAGTGGAGCGGTGAAATGCGTAGATATTGGAAAGAACACCGATGGCGAAAGCACTTTACTGGGCTATTACTAACACTCAGAGACGAAAGCTAGGGTAGCAAATGGG
## ASV2 ACGGGAGTGGCAAGCGTTATCCGGAATTATTGGGCGTAAAGCGTCCGCAGGCGGTCTTGTAAGTCTGTTGTTAAAGCGTGGAGCTTAACTCCATTTCAGCAATGGAAACTGTAAGACTAGAGTGTGGTAGGGGCAGAGGGAATTCCCGGTGTAGCGGTGAAATGCGTAGATATCGGGAAGAACACCAGTGGCGAAGGCGCTCTGCTGGGCCATAACTGACGCTCATGGACGAAAGCCAGGGGAGCGAAAGGG
## ASV3 ACGGGAGTGGCAAGCGTTATCCGGAATTATTGGGCGTAAAGCGTCCGCAGGCGGTCTTTTAAGTCTGCTGTTAAAGCGTGGAGCTTAACTCCATTTCGGCAGTGGAAACTGGAAGACTAGAGTGTGGTAGGGGCAGAGGGAATTCCCGGTGTAGCGGTGAAATGCGTAGATATCGGGAAGAACACCAGTGGCGAAGGCGCTCTGCTGGGCCATAACTGACGCTCATGGACGAAAGCCAGGGGAGCGAAAGGG
## ASV4 ACATAGGGTGCAAGCGTTGTCCGGAATTATTGGGCGTAAAGAGCTCGTAGGTCGTTTGCCGCGTCGATTGTGAAAATCTGAGGCTCAACCTCAGACCTGCAGTCGATACGGGCAAACTAGAGTGTGGTAGGGGAGACTGGAATTCCTGGTGTAGCGGTGGAATGCGCAGATATCAGGAGGAACACCAATGGCGAAGGCAGGTCTCTGGGCCATAACTGACACTGAGGAGCGAAAGCGCGGGGAGCGAACAGG
## ASV5 ACGGGAGTGGCAAGCGTTATCCGGAATTATTGGGCGTAAAGCGTCCGCAGGCGGTCTTGTAAGTCTGTCGTTAAAGCGTGGAGCTTAACTCCATTTCAGCGATGGAAACTGCAAGACTAGAGTGTGGTAGGGGCAGAGGGAATTCCCGGTGTAGCGGTGAAATGCGTAGATATCGGGAAGAACACCAGTGGCGAAGGCGCTCTGCTGGGCCATAACTGACGCTCATGGACGAAAGCCAGGGGAGCGAAAGGG
## ASV6 ACGGAGGGTGCAAGCGTTGTCCGGATTTATTGGGTTTAAAGGGTGCGCAGGTGGTTTATTAAGTCAGTGGTGAAAGACGGTCGCTCAACGATTGCAGTGCCATTGAAACTGGTAGACTTGAGTAAAGTAGAGGTGGGCGGAATTGATAGTGTAGCGGTGAAATGCATAGATATTATCAAGAACTCCAATTGCGTAGGCAGCTCACTTGGCTTTTACTGACACTCATGCACGAAAGTGTGGGTATCAAACAGG
```

```
write.csv(as.matrix((prevdfB)), "prevdfB.csv")
```

#### Plot Prevalence

```
plot(prevdfB$Prevalence, prevdfB$TotalAbundance, log="y")
```

#### Create table, number of features for each phyla

```
pHH <- table(tax_table(ps5)[, "phylum"], exclude = NULL)
dim(pHH)
```

```
## [1] 59
```

```
head(pHH)
```

```
## 
## Abditibacteriota     Acetothermia  Acidobacteriota Actinobacteriota 
##                4                1              126              259 
##  Aenigmarchaeota   Armatimonadota 
##                1               12
```

#### Plot dominant phyla

```
top20 <- names(sort(taxa_sums(ps5), decreasing=TRUE))[1:20]
ps.top20 <- transform_sample_counts(ps5, function(OTU) OTU/sum(OTU))
ps.top20 <- prune_taxa(top20, ps.top20)
HHphylBar <- plot_bar(ps.top20, fill = "phylum")
HHphylBar
```

#### Plot top 20

```
ps20 = transform_sample_counts(ps.top20, function(x) x / sum(x) )
p <- plot_bar(ps20, fill = "phylum")
p + theme_minimal() + geom_text(x=3.0, y=0.5, label="pre", size = 12, color="black") + geom_text(x=8.5, y=0.9, label="- HH -", size = 12, color="black")+ geom_text(x=14.5, y=0.5, label="- post", size = 12, color="black")
```

#### Define prevalence threshold as 10% of total samples

```
prevalenceThreshold = 0.1 * nsamples(ps5)
prevalenceThreshold
```

```
## [1] 4.4
```

```
keepTaxa = rownames(prevdfB)[(prevdfB$Prevalence >= prevalenceThreshold)]
```

#### Prune by prevelance 10%

```
prevalenceThreshold = 0.1 * nsamples(ps5)
prevalenceThreshold
```

```
## [1] 4.4
```

```
keepTaxa = rownames(prevdfB)[(prevdfB$Prevalence >= prevalenceThreshold)]
ps6 = prune_taxa(keepTaxa, ps5)
saveRDS(ps6, "psHHasv6.rds")
ps6
```

```
## phyloseq-class experiment-level object
## otu_table()   OTU Table:         [ 1616 taxa and 44 samples ]
## sample_data() Sample Data:       [ 44 samples by 21 sample variables ]
## tax_table()   Taxonomy Table:    [ 1616 taxa by 7 taxonomic ranks ]
## phy_tree()    Phylogenetic Tree: [ 1616 tips and 1615 internal nodes ]
## refseq()      DNAStringSet:      [ 1616 reference sequences ]
```

#### Write Prev

```
ps62 = transform_sample_counts(ps6, function(x) x / sum(x) )
plot_bar(ps62, fill = "domain")
```

#### Extract abundance matrix from the phyloseq object

```
ps6OTU = as(otu_table(ps6), "matrix")
ps6chem = as(sample_data(ps6), "matrix")
write.csv(ps6chem, "ps6chem.csv")
ps6chem = ps6chem[,5:14]
# Coerce to data.frame
ps6OTUdf = as.data.frame(ps6OTU)
ps6chemDF <- as.data.frame(apply(ps6chem, 2, as.numeric))  # Convert all variable types to numeric
```

```
## Warning in apply(ps6chem, 2, as.numeric): NAs introduced by coercion

## Warning in apply(ps6chem, 2, as.numeric): NAs introduced by coercion
```

```
head(ps6chemDF)
```

```
##   season Date NOxuM NH4uM  PuM   pH Conductivity   TDS Salinity SterivexML
## 1     NA   NA   1.1   0.5  4.0 7.77       14.475 13059      8.8         90
## 2     NA   NA    NA    NA   NA 7.87        1.181  1063      0.6        150
## 3     NA   NA   1.0   0.6  2.7 7.47       18.260 16796     11.5        140
## 4     NA   NA   1.1   1.4  4.1 7.39       19.530 17595     12.1        120
## 5     NA   NA  59.3   5.9 14.6 7.74        5.395  4854      3.0        165
## 6     NA   NA   1.6   1.4  2.0 7.58        0.105    90      0.0         95
```

#### Archive preprocessed phyloseq object

```
tail(ps6chemDF)
```

```
##    season Date NOxuM NH4uM PuM   pH Conductivity   TDS Salinity SterivexML
## 39     NA   NA   1.5   1.8 2.1 7.83        12.16 10935      7.0        112
## 40     NA   NA   0.9   3.7 1.4 7.48        15.30 13772      9.0        125
## 41     NA   NA   1.3   2.9 1.0 7.31        18.07 16263     10.8        146
## 42     NA   NA    NA    NA  NA   NA           NA    NA       NA         NA
## 43     NA   NA    NA    NA  NA   NA           NA    NA       NA         NA
## 44     NA   NA    NA    NA  NA   NA           NA    NA       NA         NA
```

#### Run NMDS on Hellinger transformed data

```
ps6OTUrel <-         
  decostand(ps6OTU, method = "total")
# Calculate distance matrix
ps6OTUdistmat <- vegdist(decostand(ps6OTUrel, "hellinger"), "bray")
# Creating easy to view matrix and writing .csv
ps6OTUdistmat <- 
  as.matrix(ps6OTUdistmat, labels = T)
write.csv(ps6OTUdistmat, "ps6OTUdistmat.csv")
ps6NMS <-
  vegan::metaMDS(ps6OTUdistmat,
          distance = "bray",
          k = 3,
          maxit = 999, 
          trymax = 500,
          wascores = TRUE)
```

```
## Run 0 stress 0.03622557 
## Run 1 stress 0.03622558 
## ... Procrustes: rmse 0.0003373262  max resid 0.0009140127 
## ... Similar to previous best
## Run 2 stress 0.03622558 
## ... Procrustes: rmse 0.0003338497  max resid 0.0009088591 
## ... Similar to previous best
## Run 3 stress 0.03622562 
## ... Procrustes: rmse 0.0003537178  max resid 0.0009548466 
## ... Similar to previous best
## Run 4 stress 0.03622539 
## ... New best solution
## ... Procrustes: rmse 0.0002470455  max resid 0.0006677073 
## ... Similar to previous best
## Run 5 stress 0.0404628 
## Run 6 stress 0.04046292 
## Run 7 stress 0.03622536 
## ... New best solution
## ... Procrustes: rmse 2.534491e-05  max resid 6.780369e-05 
## ... Similar to previous best
## Run 8 stress 0.04046296 
## Run 9 stress 0.04568821 
## Run 10 stress 0.04204516 
## Run 11 stress 0.04204502 
## Run 12 stress 0.03622562 
## ... Procrustes: rmse 0.0001295251  max resid 0.0003545126 
## ... Similar to previous best
## Run 13 stress 0.04204505 
## Run 14 stress 0.03622552 
## ... Procrustes: rmse 9.613068e-05  max resid 0.0002593197 
## ... Similar to previous best
## Run 15 stress 0.04569526 
## Run 16 stress 0.04046281 
## Run 17 stress 0.03622537 
## ... Procrustes: rmse 9.918511e-06  max resid 2.831006e-05 
## ... Similar to previous best
## Run 18 stress 0.04046287 
## Run 19 stress 0.04204513 
## Run 20 stress 0.04046284 
## *** Solution reached
```

#### Shepards test/goodness of fit. Produces a results of test statistics for goodness of fit for each point

```
##  [1] 0.004651985 0.006174455 0.004243652 0.004740741 0.005482613 0.004695361
##  [7] 0.006675539 0.005536475 0.004594166 0.005408768 0.004243294 0.003576584
## [13] 0.006098621 0.004377600 0.005239927 0.005520910 0.003924155 0.005789899
## [19] 0.004999366 0.005482978 0.004735721 0.005117853 0.005176527 0.005521471
## [25] 0.004227606 0.004400464 0.003579852 0.006328594 0.007033387 0.007785439
## [31] 0.005501049 0.004193429 0.007619319 0.005566072 0.004629565 0.005063451
## [37] 0.005351386 0.005917205 0.004163441 0.004030614 0.004086450 0.009123976
## [43] 0.007712492 0.006211058
```

#### Plot Model

```
p1 <- plot(ps6NMS)
```

```
## species scores not available
```

```
p1Sites <- as.data.frame(p1$sites)
psGroup <- as.data.frame(sample_data(ps6))
p1Sites <- cbind(p1Sites, psGroup$type)
p1Sites <- cbind(p1Sites, psGroup$season)
names(p1Sites)[3] <- "type"
names(p1Sites)[4] <- "season"
head(p1Sites)
```

```
##         NMDS1      NMDS2 type season
## aC -0.4395558 -0.1339471  pre summer
## aH -0.0154260  0.1265597  pre summer
## aJ -0.4654323 -0.2166353  pre summer
## aK -0.4812832 -0.2268656  pre summer
## aR -0.2773364 -0.0096675  pre summer
## bC  0.3951293 -0.2493230   HH summer
```

#### Ordination Plot

```
gg_ordiplot(ps6NMS, groups = p1Sites$type, kind = "sd", conf = 0.95, pt.size = 3) + theme_bw()
```

```
## Warning in chol.default(cov, pivot = TRUE): the matrix is either rank-deficient
## or indefinite
```

```
## NULL
```

#### Remove samples without nutrients

```
ps7 = subset_samples(ps6, NOxuM != "NA")
ps7
```

```
## phyloseq-class experiment-level object
## otu_table()   OTU Table:         [ 1616 taxa and 40 samples ]
## sample_data() Sample Data:       [ 40 samples by 21 sample variables ]
## tax_table()   Taxonomy Table:    [ 1616 taxa by 7 taxonomic ranks ]
## phy_tree()    Phylogenetic Tree: [ 1616 tips and 1615 internal nodes ]
## refseq()      DNAStringSet:      [ 1616 reference sequences ]
```

```
# saveRDS(ps7, "ps7HH.rds")
```

#### Extract abundance matrix from the phyloseq object

```
ps6OTU = as(otu_table(ps7), "matrix")
ps6chem = as(sample_data(ps7), "matrix")
write.csv(ps6chem, "ps6chem.csv")
ps6chem2 = ps6chem[,7:14]
p6type <-  as.data.frame(ps6chem[,3])
names(p6type)[1] <- "types"
# Coerce to data.frame
ps6OTUdf = as.data.frame(ps6OTU)
ps6chemDF <- as.data.frame(apply(ps6chem2, 2, as.numeric))  # Convert all variable types to numeric
head(ps6chemDF)
```

```
##   NOxuM NH4uM  PuM   pH Conductivity   TDS Salinity SterivexML
## 1   1.1   0.5  4.0 7.77       14.475 13059      8.8         90
## 2   1.0   0.6  2.7 7.47       18.260 16796     11.5        140
## 3   1.1   1.4  4.1 7.39       19.530 17595     12.1        120
## 4  59.3   5.9 14.6 7.74        5.395  4854      3.0        165
## 5   1.6   1.4  2.0 7.58        0.105    90      0.0         95
## 6   9.1   1.6  4.4 7.81        0.158   142      0.0         75
```

#### Run NMDS on Hellinger transformed data

```
ps6OTUrel <-         
  decostand(ps6OTU, method = "total")
# Calculate distance matrix
ps6OTUdistmat <- vegdist(decostand(ps6OTUrel, "hellinger"), "bray")
# Creating easy to view matrix and writing .csv
ps6OTUdistmat <- 
  as.matrix(ps6OTUdistmat, labels = T)
write.csv(ps6OTUdistmat, "ps6OTUdistmat.csv")
ps6NMS <-
  metaMDS(ps6OTUdistmat,
          distance = "bray",
          k = 3,
          maxit = 999, 
          trymax = 500,
          wascores = TRUE)
```

```
## Run 0 stress 0.03159978 
## Run 1 stress 0.03159966 
## ... New best solution
## ... Procrustes: rmse 0.0001881148  max resid 0.0006146769 
## ... Similar to previous best
## Run 2 stress 0.03544438 
## Run 3 stress 0.03159972 
## ... Procrustes: rmse 5.252977e-05  max resid 0.0001721951 
## ... Similar to previous best
## Run 4 stress 0.03159982 
## ... Procrustes: rmse 0.0001023533  max resid 0.0003371035 
## ... Similar to previous best
## Run 5 stress 0.03159972 
## ... Procrustes: rmse 5.20556e-05  max resid 0.0001713764 
## ... Similar to previous best
## Run 6 stress 0.03544449 
## Run 7 stress 0.03159976 
## ... Procrustes: rmse 7.411649e-05  max resid 0.0002428389 
## ... Similar to previous best
## Run 8 stress 0.03544425 
## Run 9 stress 0.03159972 
## ... Procrustes: rmse 4.935694e-05  max resid 0.0001618637 
## ... Similar to previous best
## Run 10 stress 0.03159984 
## ... Procrustes: rmse 0.0001106656  max resid 0.000368278 
## ... Similar to previous best
## Run 11 stress 0.03631769 
## Run 12 stress 0.03159987 
## ... Procrustes: rmse 0.000121236  max resid 0.0003988983 
## ... Similar to previous best
## Run 13 stress 0.0315999 
## ... Procrustes: rmse 0.00013202  max resid 0.0004297002 
## ... Similar to previous best
## Run 14 stress 0.0324302 
## Run 15 stress 0.03631796 
## Run 16 stress 0.03159977 
## ... Procrustes: rmse 7.653114e-05  max resid 0.0002516646 
## ... Similar to previous best
## Run 17 stress 0.03243032 
## Run 18 stress 0.03544455 
## Run 19 stress 0.03243012 
## Run 20 stress 0.03243021 
## *** Solution reached
```

#### Ordination Plot

```
goodness(ps6NMS)
```

```
##  [1] 0.004262361 0.003820235 0.004469449 0.004283770 0.003647276 0.005064128
##  [7] 0.004355580 0.003609018 0.005270571 0.003941746 0.003798690 0.007021765
## [13] 0.004111714 0.004753323 0.005148209 0.003171176 0.005252961 0.005374848
## [19] 0.005690611 0.003972307 0.005078202 0.005274292 0.006032157 0.004800211
## [25] 0.004625421 0.003652393 0.006295545 0.006958527 0.007431847 0.005385061
## [31] 0.004073178 0.007363405 0.006053172 0.004546170 0.004959209 0.005046586
## [37] 0.005743995 0.003781406 0.003270658 0.003538688
```

```
stressplot(ps6NMS)
```

#### Ordination Plot

```
ef <- envfit(ps6NMS, ps6chemDF, permu = 999)
scores(ef, "vectors")
```

```
##                   NMDS1      NMDS2
## NOxuM         0.2350880 -0.3761545
## NH4uM         0.2271009 -0.1027414
## PuM           0.2341593 -0.2382207
## pH           -0.1249150 -0.2882091
## Conductivity -0.7601465  0.5270600
## TDS          -0.7592745  0.5276726
## Salinity     -0.7432550  0.5386200
## SterivexML    0.1562087  0.2209990
```

```
p1 <- plot(ps6NMS)
```

```
## species scores not available
```

```
# orditorp(ps6NMS, "sites")
# plot(ef)
plotEF <- plot(ef, p.max = 0.1, col = "red")
```

```
plotEF
```

```
## NULL
```

#### Test fit of environmental variables

```
gg_ordiplot(ps6NMS, groups = p6type$types, kind = "sd", conf = 0.95, pt.size = 3)
```

#### Test fit of environmental variables

```
gg_envfit(ord = ps6NMS, env = ps6chemDF, groups = p6type$types, perm = 9999, pt.size = 2, alpha = 0.1) + theme_bw()
```

```
## NULL
```

#### Test fit of environmental variables

```
ef <- envfit(ps6NMS ~ Salinity + NOxuM + PuM, ps6chemDF, permu = 999)
ef
```

```
## 
## ***VECTORS
## 
##             NMDS1    NMDS2     r2 Pr(>r)    
## Salinity -0.80973  0.58680 0.8425  0.001 ***
## NOxuM     0.52999 -0.84801 0.1968  0.014 *  
## PuM       0.70100 -0.71316 0.1116  0.099 .  
## ---
## Signif. codes:  0 '***' 0.001 '**' 0.01 '*' 0.05 '.' 0.1 ' ' 1
## Permutation: free
## Number of permutations: 999
```
