## Supplemental File S12 for "Hurricane Harvey Impacts on Water Quality and Microbial Communities in Houston, TX Waterbodies": S12_HH_picrust.html

biomFormat


### biomFormat

###### Michael G. LaMontagne

#### 11/15/2021

#### R Markdown

This is R Markdown document presents exporting Hurricane Harvey microbiome data in phyloseq format to picrust2 and then importing pathway data for creating differential abundance heatmaps.

```
library(biomformat)
packageVersion("biomformat")
```

```
## [1] '1.20.0'
```

```
library(phyloseq)
packageVersion("biomformat")
```

```
## [1] '1.20.0'
```

#### Import phyloseq object

```
## phyloseq-class experiment-level object
## otu_table()   OTU Table:         [ 1616 taxa and 44 samples ]
## sample_data() Sample Data:       [ 44 samples by 21 sample variables ]
## tax_table()   Taxonomy Table:    [ 1616 taxa by 7 taxonomic ranks ]
## phy_tree()    Phylogenetic Tree: [ 1616 tips and 1615 internal nodes ]
## refseq()      DNAStringSet:      [ 1616 reference sequences ]
```

#### Convert phyloseq object to biom format

```
otu <- t(as(otu_table(ps6), "matrix"))
head(otu)
```

```
##         aC   aH    aJ    aK    aR    bC   bH   bJ   bK   bN   bR   cC   cH   cJ
## ASV1    59 1200     0    21   272   115   25    0   26   27    0 2366  750 3384
## ASV2 18705  518 39293 10875  6280    73   43    0   40    0   24  565    0  512
## ASV3  6928 4989 13328  4633 18019    48   56   39   31    0    9  311   19  603
## ASV4  2934 1936  2113  1112  6283     0  106   52  204    0   80 2963 1229 2527
## ASV5  3189 8033  1507   562 20708    63   88    0   49    0    0  680   93  445
## ASV6     0   76     0     0     0 17623 8848 4448 4309 9771 8615   86  184   61
##        cK    cN   cR   dC   dH    dJ   dK   dN    dR    eC   eH    eJ    eK
## ASV1 1787 22392 4367 4645  387  3989 4760  893 11355 29889  338   833   372
## ASV2  370   245  349 1187   63   984 2317 3101   427   384    0  8058  4722
## ASV3  380    88  200 2226  234  4224 3440 3928  2387  1288  158 13164 13910
## ASV4 2580   876 2088 7145 2361  3433 5847 3476  6389  2663  408  8092  8106
## ASV5  269   236  706 2346  877 16272 1206 3225  7988  2598  401  3442  3026
## ASV6   58   265  625    0  220     0    0    0     0    53 1887     0     0
##        eN   eR    fC   fH   fK    gC    gH   gJ   gK    gN    gR    hC   hJ
## ASV1 6873 7802 21047 8470 2633 13638 10750  349  313 44980 25548  3323 2807
## ASV2  860  383  1174  228 5773  4976   124 8258 7101   788   813 10392 3915
## ASV3 1955 1646  3211 1033 9669  6183   345 4322 2000  1734  3156  9610 4100
## ASV4 2913 3988  6230 2321 2627  5854  1856 5826 5593  8461  5547  3559 2304
## ASV5 6884 5772  2446 2522 1548  3008  1038  981  545  2454  3473  2304 1037
## ASV6  139  807     0  124    0     0   123    0    0   119    47    42   17
##        hK  iH    jH   kS
## ASV1  811 662   164    0
## ASV2 3752  95    66  141
## ASV3 4217  23    16    0
## ASV4 2483   0     0    0
## ASV5  861  65     0    0
## ASV6   15 800 24693 3703
```

#### Convert phyloseq object to biom format

```
otu_biomTemp <- make_biom(data=otu)
as(biom_data(otu_biomTemp), "matrix")
write_biom(otu_biomTemp,"HHasv.biom")
head(otu_biomTemp)
plot(biom_data(otu_biomTemp))
```

#### Export sequences from phyloseq object

```
ps4ref <- as.data.frame(refseq(ps6))
ps4ref$name <- row.names(ps4ref)
ps4ref$seq <- ps4ref$x
ps5ref <- ps4ref[,2:3]
writeFasta<-function(data, filename){
  fastaLines = c()
  for (rowNum in 1:nrow(data)){
    fastaLines = c(fastaLines, as.character(paste(">", data[rowNum,"name"], sep = "")))
    fastaLines = c(fastaLines,as.character(data[rowNum,"seq"]))
  }
  fileConn<-file(filename)
  writeLines(fastaLines, fileConn)
  close(fileConn)
}
writeFasta(ps5ref, "ps6.fna")
head(ps5ref)
```

```
##      name
## ASV1 ASV1
## ASV2 ASV2
## ASV3 ASV3
## ASV4 ASV4
## ASV5 ASV5
## ASV6 ASV6
##                                                                                                                                                                                                                                                               seq
## ASV1 ACGGAGGATGCAAGTGTTATCCGGAATCACTGGGCGTAAAGCGTCTGTAGGTGGTTTAATAAGTCAACTGTTAAATCTTGAGGCTCAACTTCAAAATCGCAGCTGAAACTATTAGACTAGAGTATAGTAGAGGTAAAGGGAATTTCCAGTGGAGCGGTGAAATGCGTAGATATTGGAAAGAACACCGATGGCGAAAGCACTTTACTGGGCTATTACTAACACTCAGAGACGAAAGCTAGGGTAGCAAATGGG
## ASV2 ACGGGAGTGGCAAGCGTTATCCGGAATTATTGGGCGTAAAGCGTCCGCAGGCGGTCTTGTAAGTCTGTTGTTAAAGCGTGGAGCTTAACTCCATTTCAGCAATGGAAACTGTAAGACTAGAGTGTGGTAGGGGCAGAGGGAATTCCCGGTGTAGCGGTGAAATGCGTAGATATCGGGAAGAACACCAGTGGCGAAGGCGCTCTGCTGGGCCATAACTGACGCTCATGGACGAAAGCCAGGGGAGCGAAAGGG
## ASV3 ACGGGAGTGGCAAGCGTTATCCGGAATTATTGGGCGTAAAGCGTCCGCAGGCGGTCTTTTAAGTCTGCTGTTAAAGCGTGGAGCTTAACTCCATTTCGGCAGTGGAAACTGGAAGACTAGAGTGTGGTAGGGGCAGAGGGAATTCCCGGTGTAGCGGTGAAATGCGTAGATATCGGGAAGAACACCAGTGGCGAAGGCGCTCTGCTGGGCCATAACTGACGCTCATGGACGAAAGCCAGGGGAGCGAAAGGG
## ASV4 ACATAGGGTGCAAGCGTTGTCCGGAATTATTGGGCGTAAAGAGCTCGTAGGTCGTTTGCCGCGTCGATTGTGAAAATCTGAGGCTCAACCTCAGACCTGCAGTCGATACGGGCAAACTAGAGTGTGGTAGGGGAGACTGGAATTCCTGGTGTAGCGGTGGAATGCGCAGATATCAGGAGGAACACCAATGGCGAAGGCAGGTCTCTGGGCCATAACTGACACTGAGGAGCGAAAGCGCGGGGAGCGAACAGG
## ASV5 ACGGGAGTGGCAAGCGTTATCCGGAATTATTGGGCGTAAAGCGTCCGCAGGCGGTCTTGTAAGTCTGTCGTTAAAGCGTGGAGCTTAACTCCATTTCAGCGATGGAAACTGCAAGACTAGAGTGTGGTAGGGGCAGAGGGAATTCCCGGTGTAGCGGTGAAATGCGTAGATATCGGGAAGAACACCAGTGGCGAAGGCGCTCTGCTGGGCCATAACTGACGCTCATGGACGAAAGCCAGGGGAGCGAAAGGG
## ASV6 ACGGAGGGTGCAAGCGTTGTCCGGATTTATTGGGTTTAAAGGGTGCGCAGGTGGTTTATTAAGTCAGTGGTGAAAGACGGTCGCTCAACGATTGCAGTGCCATTGAAACTGGTAGACTTGAGTAAAGTAGAGGTGGGCGGAATTGATAGTGTAGCGGTGAAATGCATAGATATTATCAAGAACTCCAATTGCGTAGGCAGCTCACTTGGCTTTTACTGACACTCATGCACGAAAGTGTGGGTATCAAACAGG
```

#### Plot biom format

```
heatmap(as(biom_data(otu_biomTemp), "matrix"))
```

#### run picrust2 (bash commands not run in R markdown)

```
#cp /home/mgl111/winhome/Documents/preHH.biom /home/mgl111/HHpi
#cp /home/mgl111/winhome/Documents/preHH.fna /home/mgl111/HHpi
#biom head -i preHH.biom
#get more information about the rest of the table by using this command:
#biom summarize-table -i preHH.biom
#place_seqs.py -s preHH.fna -o preHH.tre -p 2 --verbose
# determine sequenced neighbor
#hsp.py -i 16S -t preHH.tre -o marker_predicted_and_nsti_preHH.tsv.gz -p 2 -n --verbose
#hsp.py -i EC -t preHH.tre -o EC_predicted_preHH.tsv.gz -p 2 --verbose
#hsp.py -i KO -t preHH.tre -o KO_predicted_preHH.tsv.gz -p 2 --verbose
#
#metagenome_pipeline.py -i preHH.biom -f EC_predicted_preHH.tsv.gz -m marker_predicted_and_nsti_preHH.tsv.gz --max_nsti 2.0 -o metagenome_out_preHH
#metagenome_pipeline.py -i preHH.biom 
#  -m marker_predicted_and_nsti_preHH.tsv.gz 
#  -f KO_predicted_preHH.tsv.gz --strat_out -o KO_metagenome_out

#
#pathway_pipeline.py -i metagenome_out_preHH/pred_metagenome_unstrat.tsv.gz -o pathways_out -p 2
#add_descriptions.py -i metagenome_out_preHH/pred_metagenome_unstrat.tsv.gz -m EC -o metagenome_out/pred_metagenome_unstrat_descrip_preHH.tsv.gz
#add_descriptions.py -i pathways_out/path_abun_unstrat.tsv.gz -m METACYC -o pathways_out/path_abun_unstrat_descrip_preHH.tsv.gz
```
