## Supplemental File S15 for "Hurricane Harvey Impacts on Water Quality and Microbial Communities in Houston, TX Waterbodies": S15_ANCOMBC_HH_pathways.html

ANCOMBC of Hurricane Harvey Pathways


### ANCOMBC of Hurricane Harvey Pathways

###### Michael G. LaMontagne

#### 11/1/2021

#### R Markdown

This is R Markdown identifies differentially abundant archaea in HH data set following ANCOMBC (https://www.nature.com/articles/s41467-020-17041-7).

```
library(ANCOMBC)
library(phyloseq)
library(microbiome)
```

```
## Loading required package: ggplot2
```

```
## 
## microbiome R package (microbiome.github.com)
##     
## 
## 
##  Copyright (C) 2011-2020 Leo Lahti, 
##     Sudarshan Shetty et al. <microbiome.github.io>
```

```
## 
## Attaching package: 'microbiome'
```

```
## The following object is masked from 'package:ggplot2':
## 
##     alpha
```

```
## The following object is masked from 'package:base':
## 
##     transform
```

```
library(dplyr)
```

```
## 
## Attaching package: 'dplyr'
```

```
## The following objects are masked from 'package:stats':
## 
##     filter, lag
```

```
## The following objects are masked from 'package:base':
## 
##     intersect, setdiff, setequal, union
```

```
library(qwraps2)
library(DT)
library("tibble")
sessionInfo()
```

```
## R version 4.1.1 (2021-08-10)
## Platform: x86_64-w64-mingw32/x64 (64-bit)
## Running under: Windows 10 x64 (build 22000)
## 
## Matrix products: default
## 
## locale:
## [1] LC_COLLATE=English_United States.1252 
## [2] LC_CTYPE=English_United States.1252   
## [3] LC_MONETARY=English_United States.1252
## [4] LC_NUMERIC=C                          
## [5] LC_TIME=English_United States.1252    
## 
## attached base packages:
## [1] stats     graphics  grDevices utils     datasets  methods   base     
## 
## other attached packages:
## [1] tibble_3.1.4      DT_0.19           qwraps2_0.5.2     dplyr_1.0.7      
## [5] microbiome_1.14.0 ggplot2_3.3.5     phyloseq_1.36.0   ANCOMBC_1.2.2    
## 
## loaded via a namespace (and not attached):
##  [1] Biobase_2.52.0         sass_0.4.0             tidyr_1.1.4           
##  [4] jsonlite_1.7.2         splines_4.1.1          foreach_1.5.1         
##  [7] bslib_0.3.1            Rdpack_2.1.2           assertthat_0.2.1      
## [10] stats4_4.1.1           GenomeInfoDbData_1.2.6 yaml_2.2.1            
## [13] pillar_1.6.4           lattice_0.20-44        glue_1.4.2            
## [16] digest_0.6.27          XVector_0.32.0         rbibutils_2.2.4       
## [19] colorspace_2.0-2       htmltools_0.5.2        Matrix_1.3-4          
## [22] plyr_1.8.6             pkgconfig_2.0.3        zlibbioc_1.38.0       
## [25] purrr_0.3.4            scales_1.1.1           Rtsne_0.15            
## [28] mgcv_1.8-36            generics_0.1.1         IRanges_2.26.0        
## [31] ellipsis_0.3.2         withr_2.4.2            BiocGenerics_0.38.0   
## [34] survival_3.2-11        magrittr_2.0.1         crayon_1.4.1          
## [37] evaluate_0.14          fansi_0.5.0            nlme_3.1-152          
## [40] MASS_7.3-54            vegan_2.5-7            tools_4.1.1           
## [43] data.table_1.14.2      lifecycle_1.0.1        stringr_1.4.0         
## [46] Rhdf5lib_1.14.2        S4Vectors_0.30.0       munsell_0.5.0         
## [49] cluster_2.1.2          Biostrings_2.60.2      ade4_1.7-18           
## [52] compiler_4.1.1         jquerylib_0.1.4        GenomeInfoDb_1.28.2   
## [55] rlang_0.4.11           rhdf5_2.36.0           grid_4.1.1            
## [58] RCurl_1.98-1.5         nloptr_1.2.2.2         iterators_1.0.13      
## [61] rhdf5filters_1.4.0     biomformat_1.20.0      htmlwidgets_1.5.4     
## [64] igraph_1.2.7           bitops_1.0-7           rmarkdown_2.11        
## [67] gtable_0.3.0           codetools_0.2-18       multtest_2.48.0       
## [70] DBI_1.1.1              reshape2_1.4.4         R6_2.5.1              
## [73] knitr_1.36             fastmap_1.1.0          utf8_1.2.2            
## [76] permute_0.9-5          ape_5.5                stringi_1.7.5         
## [79] parallel_4.1.1         Rcpp_1.0.7             vctrs_0.3.8           
## [82] tidyselect_1.1.1       xfun_0.25
```

#### Import phyloseq object

```
ps6 <- readRDS("psHHasv6.rds")
ps6
```

```
## phyloseq-class experiment-level object
## otu_table()   OTU Table:         [ 1616 taxa and 44 samples ]
## sample_data() Sample Data:       [ 44 samples by 21 sample variables ]
## tax_table()   Taxonomy Table:    [ 1616 taxa by 7 taxonomic ranks ]
## phy_tree()    Phylogenetic Tree: [ 1616 tips and 1615 internal nodes ]
## refseq()      DNAStringSet:      [ 1616 reference sequences ]
```

#### View sample data

```
head(tax_table(ps6))
```

```
## Taxonomy Table:     [6 taxa by 7 taxonomic ranks]:
##      domain     phylum             class            order            
## ASV1 "Bacteria" "Cyanobacteria"    "Cyanobacteriia" "Chloroplast"    
## ASV2 "Bacteria" "Cyanobacteria"    "Cyanobacteriia" "Synechococcales"
## ASV3 "Bacteria" "Cyanobacteria"    "Cyanobacteriia" "Synechococcales"
## ASV4 "Bacteria" "Actinobacteriota" "Actinobacteria" "Frankiales"     
## ASV5 "Bacteria" "Cyanobacteria"    "Cyanobacteriia" "Synechococcales"
## ASV6 "Bacteria" "Bacteroidota"     "Bacteroidia"    "Cytophagales"   
##      family            genus                species
## ASV1 NA                NA                   NA     
## ASV2 "Cyanobiaceae"    "Cyanobium PCC-6307" NA     
## ASV3 "Cyanobiaceae"    "Cyanobium PCC-6307" NA     
## ASV4 "Sporichthyaceae" "hgcI clade"         NA     
## ASV5 "Cyanobiaceae"    "Cyanobium PCC-6307" NA     
## ASV6 "Spirosomaceae"   "Pseudarcicella"     NA
```

#### View sample data

```
head(sample_data(ps6))
```

```
##    Station SegmentID type month season       Date NOxuM NH4uM  PuM   pH
## aC     C34      2425 Apre     8 summer 2017-08-27   1.1   0.5  4.0 7.77
## aH     H08     1113B Apre     8 summer 2017-08-27    NA    NA   NA 7.87
## aJ     J76     2425B Apre     8 summer 2017-08-27   1.0   0.6  2.7 7.47
## aK     K45      2425 Apre     8 summer 2017-08-27   1.1   1.4  4.1 7.39
## aR     R75      1101 Apre     8 summer 2017-08-27  59.3   5.9 14.6 7.74
## bC     C34      2425   HH     8 summer 2017-09-01   1.6   1.4  2.0 7.58
##    Conductivity   TDS Salinity SterivexML Oxygen Temp Total Ecoli Enterococii
## aC       14.475 13059      8.8         90   5.70 32.6    NA    NA          NA
## aH        1.181  1063      0.6        150   6.99 31.4    NA    NA          NA
## aJ       18.260 16796     11.5        140   7.40 32.7    NA    NA          NA
## aK       19.530 17595     12.1        120   6.00 31.7    NA    NA          NA
## aR        5.395  4854      3.0        165   8.10 32.0    NA    NA          NA
## bC        0.105    90      0.0         95   6.00 24.5 43520  1203          NA
##    factorDate   Day
## aC     25-Aug 42972
## aH     25-Aug 42972
## aJ     25-Aug 42972
## aK     25-Aug 42972
## aR     25-Aug 42972
## bC     30-Aug 42977
```

```
ps6meta <- as.data.frame(sample_data(ps6))
```

#### Import pathway data and merge to phyloseq object

```
hhPath <- read.csv2("HHpath.csv", header = TRUE, row.names = 1, sep =",", strip.white = T)
hhPathM <- data.matrix(hhPath)
otu6path = otu_table(hhPathM, taxa_are_rows = TRUE)
ps6path = merge_phyloseq(otu6path, sample_data(ps6))
ps6path
```

```
## phyloseq-class experiment-level object
## otu_table()   OTU Table:         [ 418 taxa and 44 samples ]
## sample_data() Sample Data:       [ 44 samples by 21 sample variables ]
```

#### Show pathway abundances

```
head(otu_table(ps6path))
```

```
## OTU Table:          [6 taxa and 44 samples]
##                      taxa are rows
##                                         aC  aH  aJ  aK  aR  bC  bH  bJ  bK  bN
## 1CMET2-PWY                             222 172  37 242 169   2  20 318 335 295
## 3-HYDROXYPHENYLACETATE-DEGRADATION-PWY 166 339 165   6 105 192 306 218 216 239
## AEROBACTINSYN-PWY                      180   1 346 150  53   1   1   1   1   1
## ALL-CHORISMATE-PWY                     211 220 202  47 329 246 106 231 294 317
## ANAEROFRUCAT-PWY                       206 196  27 236 199 278 314 244 263 213
## ANAGLYCOLYSIS-PWY                      294 254 104 304 255 355 358 280 304 251
##                                         bR  cC  cH  cJ  cK  cN  cR  dC  dH  dJ
## 1CMET2-PWY                             347 257 285 275 278 248  14 247 248 192
## 3-HYDROXYPHENYLACETATE-DEGRADATION-PWY 227 237  92 305 291 227 113  45   9  27
## AEROBACTINSYN-PWY                        1   1   1   1   1   1   1   1 121   1
## ALL-CHORISMATE-PWY                     268 243 251 348  31 242 151  40 314 272
## ANAEROFRUCAT-PWY                       270 268 292 281 272 280  16 273 277 228
## ANAGLYCOLYSIS-PWY                      310 319 337 338 336 337  65 321 320 289
##                                         dK  dN  dR  eC  eH  eJ  eK  eN  eR  fC
## 1CMET2-PWY                             265 207 274 206 303 226 260 239 338 275
## 3-HYDROXYPHENYLACETATE-DEGRADATION-PWY 146 101 344 171 107  21  75 284   9 180
## AEROBACTINSYN-PWY                       63   1 125   1 288  73 308   1 124 270
## ALL-CHORISMATE-PWY                      34 329  38 324  64 277 217  52  82 117
## ANAEROFRUCAT-PWY                       269 214 312 230 257 244 277 250 349 298
## ANAGLYCOLYSIS-PWY                      337 284  13 319 316 309 335 308  40  24
##                                         fH  fK  gC  gH  gJ  gK  gN  gR  hC  hJ
## 1CMET2-PWY                             238 222 275 265 298 313 357 272 313 316
## 3-HYDROXYPHENYLACETATE-DEGRADATION-PWY 311 297 178  26  49  51  26 240 307  45
## AEROBACTINSYN-PWY                        1 144 263 246 285 240   1   1   1 241
## ALL-CHORISMATE-PWY                     339  86  93  37 349 220 157  75  33 302
## ANAEROFRUCAT-PWY                       272 228 302 293 317 338  37 301 317 311
## ANAGLYCOLYSIS-PWY                      328 309  17 347   5  24 104  12  48  48
##                                         hK  iH  jH  kS
## 1CMET2-PWY                             329 238 133   7
## 3-HYDROXYPHENYLACETATE-DEGRADATION-PWY  32 119 288 143
## AEROBACTINSYN-PWY                       92   1   1   1
## ALL-CHORISMATE-PWY                     302  51  79 223
## ANAEROFRUCAT-PWY                       338 214  51 317
## ANAGLYCOLYSIS-PWY                       49 278 110  14
```

#### Add pathway descriptions

```
HHpathD <- read.csv2("HHpathDscrp.csv", header = TRUE, row.names = 1, sep =",", strip.white = T)
#HHpathD <- data.matrix(HHpathD)
HH6pathD = tax_table(HHpathD)
```

```
## Warning in .local(object): Coercing from data.frame class to character matrix 
## prior to building taxonomyTable. 
## This could introduce artifacts. 
## Check your taxonomyTable, or coerce to matrix manually.
```

```
taxa_names(HH6pathD) <- row.names(HHpathD)
ps6pathD = merge_phyloseq(otu6path, sample_data(ps6), HH6pathD)
ps6pathD
```

```
## phyloseq-class experiment-level object
## otu_table()   OTU Table:         [ 418 taxa and 44 samples ]
## sample_data() Sample Data:       [ 44 samples by 21 sample variables ]
## tax_table()   Taxonomy Table:    [ 418 taxa by 1 taxonomic ranks ]
```

#### View pathway descriptions

```
head(tax_table(ps6pathD))
```

```
## Taxonomy Table:     [6 taxa by 1 taxonomic ranks]:
##                                        ta1                                       
## 1CMET2-PWY                             "N10-formyl-tetrahydrofolate biosynthesis"
## 3-HYDROXYPHENYLACETATE-DEGRADATION-PWY "4-hydroxyphenylacetate degradation"      
## AEROBACTINSYN-PWY                      "aerobactin biosynthesis"                 
## ALL-CHORISMATE-PWY                     "superpathway of chorismate metabolism"   
## ANAEROFRUCAT-PWY                       "homolactic fermentation"                 
## ANAGLYCOLYSIS-PWY                      "glycolysis III (from glucose)"
```

#### Create list to summarize data

```
options(qwraps2_markup = "markdown")
summary_template =
  list("salinity" =
         list("min" = ~ min(Salinity, na.rm = TRUE),
              "max" = ~ max(Salinity, na.rm = TRUE),
              "mean (sd)" = ~ mean_sd(Salinity, na_rm = TRUE, 
                                      show_n = "never")),
       "Type" =
         list("Pre" = ~ n_perc0(type == "Apre", na_rm = TRUE),
              "Hurricane" = ~ n_perc0(type == "HH", na_rm = TRUE),
              "Post" = ~ n_perc0(type == "post", na_rm = TRUE),
              "Sewage" = ~ n_perc0(type == "sewage", na_rm = TRUE)),
       "Season" =
         list("Summer" = ~ n_perc0(season == "summer", na_rm = TRUE),
              "Fall" = ~ n_perc0(season == "fall", na_rm = TRUE),
              "Spring" = ~ n_perc0(season == "spring", na_rm = TRUE))
       )
data_summary = summary_table(meta(ps6pathD), summary_template)
data_summary
```

```
## 
## 
## |                       |meta(ps6pathD) (N = 44) |
## |:----------------------|:-----------------------|
## |**salinity**           |&nbsp;&nbsp;            |
## |&nbsp;&nbsp; min       |0                       |
## |&nbsp;&nbsp; max       |12.1                    |
## |&nbsp;&nbsp; mean (sd) |2.65 &plusmn; 3.63      |
## |**Type**               |&nbsp;&nbsp;            |
## |&nbsp;&nbsp; Pre       |5 (11)                  |
## |&nbsp;&nbsp; Hurricane |6 (14)                  |
## |&nbsp;&nbsp; Post      |32 (73)                 |
## |&nbsp;&nbsp; Sewage    |1 (2)                   |
## |**Season**             |&nbsp;&nbsp;            |
## |&nbsp;&nbsp; Summer    |29 (66)                 |
## |&nbsp;&nbsp; Fall      |12 (27)                 |
## |&nbsp;&nbsp; Spring    |3 (7)                   |
```

#### Remove samples with no nutrient data

```
ps7pathD = subset_samples(ps6pathD, NOxuM != "NA")
ps7pathD
```

```
## phyloseq-class experiment-level object
## otu_table()   OTU Table:         [ 418 taxa and 40 samples ]
## sample_data() Sample Data:       [ 40 samples by 21 sample variables ]
## tax_table()   Taxonomy Table:    [ 418 taxa by 1 taxonomic ranks ]
```

#### Run ancombc function

```
head(sample_data(ps7pathD))
```

```
##    Station SegmentID type month season       Date NOxuM NH4uM  PuM   pH
## aC     C34      2425 Apre     8 summer 2017-08-27   1.1   0.5  4.0 7.77
## aJ     J76     2425B Apre     8 summer 2017-08-27   1.0   0.6  2.7 7.47
## aK     K45      2425 Apre     8 summer 2017-08-27   1.1   1.4  4.1 7.39
## aR     R75      1101 Apre     8 summer 2017-08-27  59.3   5.9 14.6 7.74
## bC     C34      2425   HH     8 summer 2017-09-01   1.6   1.4  2.0 7.58
## bH     H08     1113B   HH     8 summer 2017-09-01   9.1   1.6  4.4 7.81
##    Conductivity   TDS Salinity SterivexML Oxygen Temp Total Ecoli Enterococii
## aC       14.475 13059      8.8         90    5.7 32.6    NA    NA          NA
## aJ       18.260 16796     11.5        140    7.4 32.7    NA    NA          NA
## aK       19.530 17595     12.1        120    6.0 31.7    NA    NA          NA
## aR        5.395  4854      3.0        165    8.1 32.0    NA    NA          NA
## bC        0.105    90      0.0         95    6.0 24.5 43520  1203          NA
## bH        0.158   142      0.0         75    4.9 28.1 57940  1070          NA
##    factorDate   Day
## aC     25-Aug 42972
## aJ     25-Aug 42972
## aK     25-Aug 42972
## aR     25-Aug 42972
## bC     30-Aug 42977
## bH     30-Aug 42977
```

#### Run ancombc function

```
outPath = ancombc(phyloseq = ps7pathD, formula = "Salinity + NOxuM + type", 
              p_adj_method = "holm", zero_cut = 1, lib_cut = 0, 
              group = "type", struc_zero = TRUE, neg_lb = TRUE, tol = 1e-5, 
              max_iter = 100, conserve = TRUE, alpha = 0.05, global = TRUE)
```

```
## Warning: Small sample size detected for the following group(s): 
## Apre
## ANCOM-BC results would be unstable when the sample size is < 5 per group
```

```
saveRDS(outPath, "ancmbcHHpath.Rds")
outPath <- readRDS("ancmbcHHpath.Rds")
resPath = outPath$res
res_globalPath = outPath$res_global
write.csv(outPath$feature_table, "outFeatureTablePath.csv")
write.csv(outPath$zero_ind, "outZeroIndPath.csv")
write.csv(res_globalPath, "resGlobalPath.csv")
head(res_globalPath)
```

```
##                                                W       p_val q_val diff_abn
## 1CMET2-PWY                              4.373146 0.224601880     1    FALSE
## 3-HYDROXYPHENYLACETATE-DEGRADATION-PWY  1.621420 0.889084844     1    FALSE
## AEROBACTINSYN-PWY                       9.744140 0.015314998     1    FALSE
## ALL-CHORISMATE-PWY                      0.891916 0.719578710     1    FALSE
## ANAEROFRUCAT-PWY                        2.938808 0.460125114     1    FALSE
## ANAGLYCOLYSIS-PWY                      11.868931 0.005293274     1    FALSE
```

#### Coefficients from the Primary Result

```
tab_coef = resPath$beta
col_name = c("Salinity", "NOx", "pre-HH", "pre-post")
colnames(tab_coef) = col_name
tab_coef %>% datatable(caption = "Coefficients from the Primary Result") %>%
      formatRound(col_name, digits = 2)
```

#### SEs from the Primary Result

```
tab_se = resPath$se
colnames(tab_se) = col_name
tab_se %>% datatable(caption = "SEs from the Primary Result") %>%
      formatRound(col_name, digits = 2)
```

#### Test statistics

```
tab_w = resPath$W
colnames(tab_w) = col_name
tab_w %>% datatable(caption = "Test Statistics from the Primary Result") %>%
      formatRound(col_name, digits = 2)
```

#### P-values

```
tab_p = resPath$p_val
colnames(tab_p) = col_name
tab_p %>% datatable(caption = "P-values from the Primary Result") %>%
      formatRound(col_name, digits = 2)
```

#### Adjusted p-values

```
tab_q = resPath$q
colnames(tab_q) = col_name
tab_q %>% datatable(caption = "Adjusted p-values from the Primary Result") %>%
      formatRound(col_name, digits = 2)
```

#### Differentially abundant pathways

```
tab_diff = resPath$diff_abn
colnames(tab_diff) = col_name
tab_diff %>% 
  datatable(caption = "Differentially Abundant Taxa 
            from the Primary Result")
```

```
write.csv(tab_diff, "tabDiffHHpath.csv")
```

#### Join differential abundances to taxa table

```
taxTable7 <- tax_table(ps7pathD)
tax7 <- cbind(taxTable7,tab_diff)
names(tax7)[match(names(tax7), 'ta1')] <- 'Pathway'
names(tax7)[4] <- 'HH'
names(tax7)[5] <- 'post'
tax8 <- cbind(tax7, res_globalPath)
head(tax8)
```

```
##                                                                         Pathway
## 1CMET2-PWY                             N10-formyl-tetrahydrofolate biosynthesis
## 3-HYDROXYPHENYLACETATE-DEGRADATION-PWY       4-hydroxyphenylacetate degradation
## AEROBACTINSYN-PWY                                       aerobactin biosynthesis
## ALL-CHORISMATE-PWY                        superpathway of chorismate metabolism
## ANAEROFRUCAT-PWY                                        homolactic fermentation
## ANAGLYCOLYSIS-PWY                                 glycolysis III (from glucose)
##                                        Salinity   NOx    HH  post         W
## 1CMET2-PWY                                FALSE FALSE FALSE FALSE  4.373146
## 3-HYDROXYPHENYLACETATE-DEGRADATION-PWY    FALSE FALSE FALSE FALSE  1.621420
## AEROBACTINSYN-PWY                          TRUE FALSE FALSE FALSE  9.744140
## ALL-CHORISMATE-PWY                        FALSE FALSE FALSE FALSE  0.891916
## ANAEROFRUCAT-PWY                          FALSE FALSE FALSE FALSE  2.938808
## ANAGLYCOLYSIS-PWY                         FALSE FALSE FALSE FALSE 11.868931
##                                              p_val q_val diff_abn
## 1CMET2-PWY                             0.224601880     1    FALSE
## 3-HYDROXYPHENYLACETATE-DEGRADATION-PWY 0.889084844     1    FALSE
## AEROBACTINSYN-PWY                      0.015314998     1    FALSE
## ALL-CHORISMATE-PWY                     0.719578710     1    FALSE
## ANAEROFRUCAT-PWY                       0.460125114     1    FALSE
## ANAGLYCOLYSIS-PWY                      0.005293274     1    FALSE
```

#### Adjust for Bias

```
samp_frac = outPath$samp_frac
# Replace NA with 0
samp_frac[is.na(samp_frac)] = 0 

# Add pesudo-count (1) to avoid taking the log of 0
log_obs_abn = log(abundances(ps7pathD) + 1) 
# Adjust the log observed abundances
log_obs_abn_adj = t(t(log_obs_abn) - samp_frac)
# Show the first 6 samples
round(log_obs_abn_adj[, 1:6], 2) %>% 
  datatable(caption = "Bias-adjusted log observed abundances")
```

```
# write.csv(log_obs_abn_adj, "logObsAbnAdjHHpath.csv")
```

#### Create phyloseq object with bias adjusted pathway abundances

```
ps8pathD <- phyloseq(otu_table(log_obs_abn_adj, taxa_are_rows = TRUE))
tax8m <- as.matrix(tax8)
ps8tax <- phyloseq(tax_table(tax8m)) 
ps8pathM = merge_phyloseq(ps8pathD, sample_data(ps7pathD), ps8tax)
ps8pathM
```

```
## phyloseq-class experiment-level object
## otu_table()   OTU Table:         [ 418 taxa and 40 samples ]
## sample_data() Sample Data:       [ 40 samples by 21 sample variables ]
## tax_table()   Taxonomy Table:    [ 418 taxa by 9 taxonomic ranks ]
```

#### Visualizations for “Salinity”

```
df_fig1 = data.frame(resPath$beta * resPath$diff_abn, check.names = FALSE) %>% 
  rownames_to_column("taxon_id")
df_fig2 = data.frame(resPath$se * resPath$diff_abn, check.names = FALSE) %>% 
  rownames_to_column("taxon_id")
colnames(df_fig2)[-1] = paste0(colnames(df_fig2)[-1], "SD")
df_fig = df_fig1 %>% left_join(df_fig2, by = "taxon_id") %>%
  transmute(taxon_id, Salinity, SalinitySD) %>%
  filter(Salinity != 0) %>% arrange(desc(Salinity)) %>%
  mutate(group = ifelse(Salinity > 0, "g1", "g2"))
df_fig$taxon_id = factor(df_fig$taxon_id, levels = df_fig$taxon_id)
  
p = ggplot(data = df_fig, 
           aes(x = taxon_id, y = Salinity, fill = group, color = group)) + 
  geom_bar(stat = "identity", width = 0.7, 
           position = position_dodge(width = 0.4)) +
  geom_errorbar(aes(ymin = Salinity - SalinitySD, ymax = Salinity + SalinitySD), width = 0.2,
                position = position_dodge(0.05), color = "black") + 
  labs(x = NULL, y = "Log fold change", 
       title = "Waterfall Plot for the Salinity Effect") + 
  theme_bw() + 
  theme(legend.position = "none",
        plot.title = element_text(hjust = 0.5),
        panel.grid.minor.y = element_blank(),
        axis.text.x = element_text(angle = 60, hjust = 1))
p
```

#### Save plot

```
ggsave("Salinity_path.png")
```

```
## Saving 7 x 5 in image
```

#### Visualizations for Nitrate/Nitrite

```
df_fig1 = data.frame(resPath$beta * resPath$diff_abn, check.names = FALSE) %>% 
  rownames_to_column("taxon_id")
df_fig2 = data.frame(resPath$se * resPath$diff_abn, check.names = FALSE) %>% 
  rownames_to_column("taxon_id")
colnames(df_fig2)[-1] = paste0(colnames(df_fig2)[-1], "SD")
df_fig = df_fig1 %>% left_join(df_fig2, by = "taxon_id") %>%
  transmute(taxon_id, NOxuM, NOxuMSD) %>%
  filter(NOxuM != 0) %>% arrange(desc(NOxuM)) %>%
  mutate(group = ifelse(NOxuM > 0, "g1", "g2"))
df_fig$taxon_id = factor(df_fig$taxon_id, levels = df_fig$taxon_id)
  
p = ggplot(data = df_fig, 
           aes(x = taxon_id, y = NOxuM, fill = group, color = group)) + 
  geom_bar(stat = "identity", width = 0.7, 
           position = position_dodge(width = 0.4)) +
  geom_errorbar(aes(ymin = NOxuM - NOxuMSD, ymax = NOxuM + NOxuMSD), width = 0.2,
                position = position_dodge(0.05), color = "black") + 
  labs(x = NULL, y = "Log fold change", 
       title = "Waterfall Plot for the Nitrogen Effect") + 
  theme_bw() + 
  theme(legend.position = "none",
        plot.title = element_text(hjust = 0.5),
        panel.grid.minor.y = element_blank(),
        axis.text.x = element_text(angle = 60, hjust = 1))
p
```

#### Visualizations for nitrat/nitrite

```
ggsave("NOx_path.png")
```

```
## Saving 7 x 5 in image
```

#### Visualizations for Hurricane Harvey

```
df_fig1 = data.frame(resPath$beta * resPath$diff_abn, check.names = FALSE) %>% 
  rownames_to_column("taxon_id")
df_fig2 = data.frame(resPath$se * resPath$diff_abn, check.names = FALSE) %>% 
  rownames_to_column("taxon_id")
colnames(df_fig2)[-1] = paste0(colnames(df_fig2)[-1], "SD")
df_fig = df_fig1 %>% left_join(df_fig2, by = "taxon_id") %>%
  transmute(taxon_id, typeHH, typeHHSD) %>%
  filter(typeHH != 0) %>% arrange(desc(typeHH)) %>%
  mutate(group = ifelse(typeHH > 0, "g1", "g2"))
df_fig$taxon_id = factor(df_fig$taxon_id, levels = df_fig$taxon_id)
df_fig <- subset(df_fig, df_fig$typeHH > 1 | df_fig$typeHH < -1)
# write.csv(df_fig, "dfFigHHKO.csv")
p = ggplot(data = df_fig, 
           aes(x = taxon_id, y = typeHH, fill = group, color = group)) + 
  geom_bar(stat = "identity", width = 0.7, 
           position = position_dodge(width = 0.4)) +
  geom_errorbar(aes(ymin = typeHH - typeHHSD, ymax = typeHH + typeHHSD), width = 0.2,
                position = position_dodge(0.05), color = "black") + 
  labs(x = NULL, y = "Log fold change", 
       title = "Waterfall Plot for the Hurricane Effect") + 
  theme_bw() + 
  theme(legend.position = "none",
        plot.title = element_text(hjust = 0.5),
        panel.grid.minor.y = element_blank(),
        axis.text.x = element_text(angle = 60, hjust = 1))
p
```

#### ASV

```
ggsave("HH_path.png")
```

```
## Saving 7 x 5 in image
```

#### ASV

```
ps8pathDpreHH <- subset_taxa(ps8pathM, HH=="TRUE")
ps8pathDpreHH
```

```
## phyloseq-class experiment-level object
## otu_table()   OTU Table:         [ 29 taxa and 40 samples ]
## sample_data() Sample Data:       [ 40 samples by 21 sample variables ]
## tax_table()   Taxonomy Table:    [ 29 taxa by 9 taxonomic ranks ]
```

#### Plot differnetially abundant pathways

```
head(tax_table(ps8pathDpreHH))
```

```
## Taxonomy Table:     [6 taxa by 9 taxonomic ranks]:
##                     Pathway                                                                   
## ARGDEG-PWY          "superpathway of L-arginine, putrescine, and 4-aminobutanoate degradation"
## FASYN-ELONG-PWY     "fatty acid elongation -- saturated"                                      
## FUC-RHAMCAT-PWY     "superpathway of fucose and rhamnose degradation"                         
## GLYCOL-GLYOXDEG-PWY "superpathway of glycol metabolism and degradation"                       
## METHGLYUT-PWY       "superpathway of methylglyoxal degradation"                               
## ORNARGDEG-PWY       "superpathway of L-arginine and L-ornithine degradation"                  
##                     Salinity NOx     HH     post   W             
## ARGDEG-PWY          "FALSE"  "FALSE" "TRUE" "TRUE" " 42.01076129"
## FASYN-ELONG-PWY     "FALSE"  "FALSE" "TRUE" "TRUE" " 22.62283899"
## FUC-RHAMCAT-PWY     "FALSE"  "FALSE" "TRUE" "TRUE" " 35.08045351"
## GLYCOL-GLYOXDEG-PWY "FALSE"  "FALSE" "TRUE" "TRUE" " 32.29926507"
## METHGLYUT-PWY       "FALSE"  "FALSE" "TRUE" "TRUE" " 57.87332725"
## ORNARGDEG-PWY       "FALSE"  "FALSE" "TRUE" "TRUE" " 42.01076129"
##                     p_val           q_val           diff_abn
## ARGDEG-PWY          " 1.508374e-09" " 5.988246e-07" "TRUE"  
## FASYN-ELONG-PWY     " 2.446487e-05" " 8.611635e-03" "TRUE"  
## FUC-RHAMCAT-PWY     " 4.823989e-08" " 1.866884e-05" "TRUE"  
## GLYCOL-GLYOXDEG-PWY " 1.937910e-07" " 7.402818e-05" "TRUE"  
## METHGLYUT-PWY       " 5.419967e-13" " 2.189667e-10" "TRUE"  
## ORNARGDEG-PWY       " 1.508374e-09" " 5.988246e-07" "TRUE"
```

#### Plot differnetially abundant phyla

```
top15 <- names(sort(taxa_sums(ps8pathDpreHH), decreasing=TRUE))[1:15]
ps.top15 <- transform_sample_counts(ps8pathDpreHH, function(OTU) OTU/sum(OTU))
ps.top15 <- prune_taxa(top15, ps.top15)
HHphylBar <- plot_bar(ps.top15, fill = "Pathway")
```

```
## Warning in psmelt(physeq): The sample variables: 
## Salinity
##  have been renamed to: 
## sample_Salinity
## to avoid conflicts with taxonomic rank names.
```

```
HHphylBar
```

#### Plot pathways

```
ps15 = transform_sample_counts(ps.top15, function(x) x / sum(x) )
p <- plot_bar(ps15, fill = "Pathway")
```

```
## Warning in psmelt(physeq): The sample variables: 
## Salinity
##  have been renamed to: 
## sample_Salinity
## to avoid conflicts with taxonomic rank names.
```

```
p + theme_minimal() + geom_text(x=8.5, y=0.9, label="HH", size = 8, color="black")
```

` ## Plot global test result

```
ResHH <- subset(res_globalPath, res_globalPath$q_val < 0.05 & res_globalPath$W > 20)
pairs(ResHH[,1:3], pch = 19)
```
