## Supplemental File S16 for "Hurricane Harvey Impacts on Water Quality and Microbial Communities in Houston, TX Waterbodies": S16_heatMapHHpcrst.html

Making heatmaps with R for microbiome analysis


### Making heatmaps with R for microbiome analysis

###### Michael G. LaMontagne

#### 10/28/2021

#### R Markdown

This is R Markdown document follows https://www.molecularecologist.com/2013/08/20/making-heatmaps-with-r-for-microbiome-analysis/.

```
library(gplots)
```

```
## 
## Attaching package: 'gplots'
```

```
## The following object is masked from 'package:stats':
## 
##     lowess
```

```
library(Heatplus)
library(vegan)
```

```
## Loading required package: permute
```

```
## Loading required package: lattice
```

```
## This is vegan 2.5-7
```

```
# load the RColorBrewer package for better colour options
library(RColorBrewer)
sessionInfo()
```

```
## R version 4.1.1 (2021-08-10)
## Platform: x86_64-w64-mingw32/x64 (64-bit)
## Running under: Windows 10 x64 (build 22000)
## 
## Matrix products: default
## 
## locale:
## [1] LC_COLLATE=English_United States.1252 
## [2] LC_CTYPE=English_United States.1252   
## [3] LC_MONETARY=English_United States.1252
## [4] LC_NUMERIC=C                          
## [5] LC_TIME=English_United States.1252    
## 
## attached base packages:
## [1] stats     graphics  grDevices utils     datasets  methods   base     
## 
## other attached packages:
## [1] RColorBrewer_1.1-2 vegan_2.5-7        lattice_0.20-44    permute_0.9-5     
## [5] Heatplus_3.0.0     gplots_3.1.1      
## 
## loaded via a namespace (and not attached):
##  [1] cluster_2.1.2      knitr_1.36         magrittr_2.0.1     splines_4.1.1     
##  [5] MASS_7.3-54        R6_2.5.1           rlang_0.4.11       fastmap_1.1.0     
##  [9] stringr_1.4.0      caTools_1.18.2     tools_4.1.1        parallel_4.1.1    
## [13] grid_4.1.1         nlme_3.1-152       mgcv_1.8-36        xfun_0.25         
## [17] KernSmooth_2.23-20 jquerylib_0.1.4    htmltools_0.5.2    gtools_3.9.2      
## [21] yaml_2.2.1         digest_0.6.27      Matrix_1.3-4       sass_0.4.0        
## [25] bitops_1.0-7       evaluate_0.14      rmarkdown_2.11     stringi_1.7.5     
## [29] compiler_4.1.1     bslib_0.3.1        jsonlite_1.7.2
```

#### Read phyloseq object from ANCOMBC\_HH\_paths

```
ps8pathM <- readRDS("ps8pathM.rds")
ps8pathM
```

```
## Loading required package: phyloseq
```

```
## phyloseq-class experiment-level object
## otu_table()   OTU Table:         [ 418 taxa and 40 samples ]
## sample_data() Sample Data:       [ 40 samples by 21 sample variables ]
## tax_table()   Taxonomy Table:    [ 418 taxa by 9 taxonomic ranks ]
```

#### Convert phyloseq object into dataframes

```
HHadjPathOTU <- as.data.frame(otu_table(ps8pathM))
HHpathSample <- as.data.frame(sample_data(ps8pathM))
HHpathTax <- as.data.frame(tax_table(ps8pathM))
dim(HHadjPathOTU)
```

```
## [1] 418  40
```

#### join tax\_table and otu\_table

```
HHpathMerge <- cbind(HHpathTax,HHadjPathOTU)
saveRDS(HHpathMerge, "HHpathMerge.rds")
dim(HHpathMerge)
```

```
## [1] 418  49
```

#### Select differentially abundant pathways

```
HHpathQ <- subset(HHpathMerge, HHpathMerge$diff_abn == TRUE)
dataHH <- t(HHpathQ[10:49])
dataHH[1:3, 1:4]
```

```
##    ARGDEG-PWY ARGSYN-PWY DENOVOPURINE2-PWY  FAO-PWY
## aC  0.5737205   5.348633          5.246549 5.168840
## aJ  0.2503616   3.651559          5.423682 3.363877
## aK  0.3571667   5.220848          5.043917 5.197409
```

#### normalize data to proportion

```
dataHHprop <- dataHH/rowSums(dataHH)
dataHHprop[1:3, 1:3]
```

```
##     ARGDEG-PWY ARGSYN-PWY DENOVOPURINE2-PWY
## aC 0.002135995 0.01991327        0.01953321
## aJ 0.001101979 0.01607251        0.02387260
## aK 0.001369096 0.02001263        0.01933441
```

#### Do average linkage hierarchical clustering on Bray-Curtis dissimilarity matrix on the full dataset

```
dataHHdist <- vegdist(dataHHprop, method = "bray")
```

```
## Warning in vegdist(dataHHprop, method = "bray"): results may be meaningless
## because data have negative entries in method "bray"
```

```
row.clus <- hclust(dataHHdist, "aver")
heatmap(as.matrix(dataHHprop), Rowv = as.dendrogram(row.clus), Colv = NA, margins = c(10, 3))
```

#### Add a column dendrogram to cluster by pathway

```
# you have to transpose the dataset to get the genera as rows
dataHHdist.g <- vegdist(t(dataHHprop), method = "bray")
```

```
## Warning in vegdist(t(dataHHprop), method = "bray"): results may be meaningless
## because data have negative entries in method "bray"
```

```
col.clus <- hclust(dataHHdist.g, "aver")
heatmap(as.matrix(dataHHprop), Rowv = as.dendrogram(row.clus), Colv = as.dendrogram(col.clus), margins = c(10, 3))
```

#### Assign colors to samples

```
varHH <- HHpathSample$type
varHH <- replace(varHH, which(varHH == "Apre"), "deepskyblue")
varHH <- replace(varHH, which(varHH == "HH"), "red")
varHH <- replace(varHH, which(varHH == "post"), "green")
cbind(row.names(dataHHprop), varHH)
```

```
##            varHH        
##  [1,] "aC" "deepskyblue"
##  [2,] "aJ" "deepskyblue"
##  [3,] "aK" "deepskyblue"
##  [4,] "aR" "deepskyblue"
##  [5,] "bC" "red"        
##  [6,] "bH" "red"        
##  [7,] "bJ" "red"        
##  [8,] "bK" "red"        
##  [9,] "bN" "red"        
## [10,] "bR" "red"        
## [11,] "cC" "green"      
## [12,] "cH" "green"      
## [13,] "cJ" "green"      
## [14,] "cK" "green"      
## [15,] "cN" "green"      
## [16,] "cR" "green"      
## [17,] "dC" "green"      
## [18,] "dH" "green"      
## [19,] "dJ" "green"      
## [20,] "dK" "green"      
## [21,] "dN" "green"      
## [22,] "dR" "green"      
## [23,] "eC" "green"      
## [24,] "eH" "green"      
## [25,] "eJ" "green"      
## [26,] "eK" "green"      
## [27,] "eN" "green"      
## [28,] "eR" "green"      
## [29,] "fC" "green"      
## [30,] "fH" "green"      
## [31,] "fK" "green"      
## [32,] "gC" "green"      
## [33,] "gH" "green"      
## [34,] "gJ" "green"      
## [35,] "gK" "green"      
## [36,] "gN" "green"      
## [37,] "gR" "green"      
## [38,] "hC" "green"      
## [39,] "hJ" "green"      
## [40,] "hK" "green"
```

#### Plot heatmap

```
heatmap.2(as.matrix(dataHHprop),Rowv = as.dendrogram(row.clus), Colv = as.dendrogram(col.clus), col = 'heat.colors', RowSideColors = varHH, margins = c(7, 3))
```
